# Identifying Fibrogenic Cells Following Salivary Gland Obstructive Injury

**DOI:** 10.1101/2023.03.09.531751

**Authors:** Amber L. Altrieth, Kevin J. O’Keefe, Victoria A. Gellatly, Joey R. Tavarez, Sage M. Feminella, Nicholas L. Moskwa, Carmalena V. Cordi, Judy C. Turrieta, Deirdre A. Nelson, Melinda Larsen

## Abstract

Fibrosis results from excess extracellular matrix accumulation, which alters normal tissue architecture and impedes function. In the salivary gland, fibrosis can be induced by irradiation treatment for cancer therapy, Sjögren’s Disease, and other causes; however, it is unclear which stromal cells and signals participate in injury responses and disease progression. As hedgehog signaling has been implicated in fibrosis of the salivary gland and other organs, we examined contributions of the hedgehog effector, Gli1, to fibrotic responses in salivary glands. To experimentally induce a fibrotic response in female murine submandibular salivary glands, we performed ductal ligation surgery. We detected a progressive fibrotic response where both extracellular matrix accumulation and actively remodeled collagen trended upwards at 7 days and significantly increased at 14 days post- ligation. Macrophages, which participate in extracellular matrix remodeling, Gli1^+^ and PDGFRα^+^ stromal cells, which may deposit extracellular matrix, both increased with injury. Using single-cell RNA-sequencing, we found that a majority of *Gli1*^+^ cells at embryonic day 16 also express *Pdgfra* and/or *Pdgfrb.* However, in adult mice, only a small subset of Gli1^+^ cells express PDGFRα and/or PDGFRβ at the protein level. Using lineage-tracing mice, we found that Gli1-derived cells expand with ductal ligation injury. Although some of the Gli1 lineage-traced tdTomato^+^ cells expressed vimentin and PDGFRβ following injury, there was no increase in the classic myofibroblast marker, smooth muscle alpha-actin. Additionally, there was little change in extracellular matrix area, remodeled collagen area, PDGFRα, PDGFRβ, endothelial cells, neurons, or macrophages in Gli1 null salivary glands following injury when compared with controls, suggesting that Gli1 signaling and Gli1^+^ cells have only a minor contribution to mechanical injury-induced fibrotic changes in the salivary gland. We used scRNA-seq to examine cell populations that expand with ligation and/or showed increased expression of matrisome genes. *Pdgfra^+^/Pdgfrb^+^* stromal cell subpopulations both expanded in response to ligation, showed increased expression and a greater diversity of matrisome genes expressed, consistent with these cells being fibrogenic. Defining the signaling pathways driving fibrotic responses in stromal cell sub-types could reveal future therapeutic targets.

## 1 Introduction

Although extracellular matrix (ECM) remodeling is required for organ development and wound repair (Gonzalez et al., 2016), if not resolved it leads to prolonged excess deposition of ECM, known as fibrosis (Gurtner et al., 2008). Following normal wound healing responses, ECM production transiently increases but is ultimately degraded as the wound heals. However, following a persistent injury, the presence of excess ECM causes an alteration of tissue architecture and impedes organ function, ultimately leading to a decline in systemic health. Fibrosis contributes to up to 45% of all deaths in the developed world due to its prevalence in many diseases (Wynn, 2008). Understanding the molecular mechanisms that drive fibrosis can provide insight into therapeutics that slow progression or reverse fibrotic responses, which interfere with normal organ function.

One prevalent organ that is subject to fibrosis is the salivary gland. Salivary gland fibrosis has many causes including ductal obstruction (Lau et al., 2017), radiation therapy as a treatment for head and neck cancer (Straub et al., 2015), and a subset of patients with the autoimmune disease, Sjögren’s Syndrome (Bookman et al., 2011a; Llamas-Gutierrez et al., 2014). Salivary gland obstruction typically occurs following formation of intraluminal calcifications in the striated and excretory ducts, leading to inflammation, fibrosis, and acinar cell atrophy (Lau et al., 2017). Radiation-induced fibrosis is one of the significant long-term side effects following radiation therapy and is thought to be caused by chronic inflammation (Straub et al., 2015). In patients with primary Sjögren’s Syndrome, there is a correlation between stimulated saliva flow and the degree of fibrosis, where 34% of patients with higher fibrotic indices tended to have a greater decrease in saliva flow (Bookman et al., 2011). In all cases, salivary gland fibrosis is associated with a decrease in gland function and an overall decrease in quality of life for patients. The underlying mechanisms of salivary gland fibrosis are complex and poorly understood.

The cells contributing and the mechanisms propagating fibrosis need to be defined to facilitate therapeutic development. Many cells that reside in the connective tissue are known to produce ECM, including stromal fibroblast cells, pericytes, endothelial cells, and myofibroblasts (Weiskirchen et al., 2019). One stromal population expressing the transcription factor, glioma-associated oncogene 1 (Gli1), has been associated with fibrosis. Gli1 is an effector of canonical hedgehog signaling, and Shh is implicated in not only fibrosis but also in development and regeneration in many contexts (Jaskoll et al., 2004; Zhou et al., 2016). Genetic ablation of Gli1^+^ cells attenuated fibrosis in the heart, kidney, and bone marrow (Kramann et al., 2015a; Schneider et al., 2017a) implicating these cells in fibrosis. In addition, signaling though Gli family transcription factors have been shown contributing to fibrosis in multiple organs. Specifically, the Gli1^+^ cell population expands following fibrotic injury and has been directly implicated in fibrosis by differentiating into myofibroblast cells in the heart, kidney, lung, and bone marrow (Cassandras et al., 2020; Kramann et al., 2015b; Schneider et al., 2017b). Gli1^+^ cells constitute a small subset of a larger platelet derived growth factor receptor beta (PDGFRβ)-expressing stromal cell population which, in some organs, also expresses platelet derived growth factor receptor alpha (PDGFRα) (Kramann et al., 2015b). Whether signaling through the Gli family transcription factors contributes broadly to fibrotic responses and specifically to salivary gland ductal ligation induced fibrosis is unknown.

In this study, we used the salivary gland ductal ligation model to induce a fibrotic response in adult female wild type (WT), Gli1 null mice, and Gli-1 lineage traced mice. This model mimics the reversible form of fibrosis that occurs in obstructive disease. Ligation of the primary ducts leading into the submandibular and sublingual salivary glands was previously reported to lead to downstream salivary gland atrophy, acinar cell loss, and fibrosis (Cotroneo et al., 2008; Woods et al., 2015a). We examined the time-course of the fibrotic response and the stromal cell populations that expanded with injury using IHC. We specifically examined the contribution of Gli1 signaling to fibrosis and the changes in the Gli1^+^ cells following injury. Using a genetic null mouse for Gli1 (Gli1^lz/lz^), we examined levels of ECM deposition, ECM remodeling, PDGFRα^+^ cells and macrophages with IHC. Using a single-cell RNA-sequencing (scRNA-seq) approach, we examined the cell populations that expand with ligation injury, and more specifically examined stromal cell subtypes which increased expression of ECM genes.

## 2 Materials and Methods

### 2.1 Animal husbandry

All animal husbandry, surgical procedures, and tissue collection were performed in accordance with protocols approved by the University at Albany, SUNY IACUC committee. Mice were housed in 12-hour light/dark cycle with access to water and dry food. C57BL/6J (JAX #000664) mice were purchased from The Jackson Laboratory. The colony starters for the Gli1^tm2Alj^/J (Gli1^lz^) (JAX #008211) mice and Gli1^tm3(cre/ERT2)Alj^/J (Gli1-CreER^T2^) (JAX #007913) mice were provided to us by Alexandera Joyner. The Gli1^lz/wt^ littermates were crossed to generate Gli1^lz/lz^ mice, null for Gli1 function (Bai et al., 2002a). Gli1^wt/wt^ littermates were used as controls for the Gli1^lz/lz^ mice. To create double transgenic Gli1 reporter mice, Gli1-CreER^T2^ males were crossed with B6.Cg-Gt(ROSA)26Sor^tm9 (CAG-tdTomato)Hze^ /J (R26^tdT^) female mice to generate Gli1-CreER^T2^;R26^tdT^ double transgenic mice. Although both strains are on a mixed background, male Gli1^lz/wt^ and Gli-CreER^T2^ males were crossed with C57BL/6J (JAX #000664) females for line maintenance. Mice were assigned a unique identifier between postnatal day 7 and 10 and were genotyped with PCR to detect LacZ for Gli1^lz^ or Cre and tdTomato for Gli1-CreER^T2^; R26^tdT^.

### 2.2 Tamoxifen induction

For lineage tracing experiments, Gli1-CreER^T2^; R26^tdT^ female mice were induced at 11 weeks old with 10 mg/kg tamoxifen (Sigma cat #T5648) dissolved in corn oil for three consecutive days. As Gli1-CreER^T2^;R26^tdT^ mice do not show leaky expression (Kramann et al., 2015a), non-induced mice were not evaluated. Surgeries were performed on mice one week after the first induction.

### 2.3 Salivary gland ductal ligation

Only female mice were used for surgical manipulations due to their similarities in tissue architecture with human salivary glands (Amano et al., 2012; Maruyama et al., 2019). Adult female mice between 10- and 18-weeks old were used for surgeries. A unique identifier was assigned to each mouse either 7 to 10 days after birth or at the time of surgery. The mice were anesthetized using 100 mg/kg ketamine and 10 mg/kg xylazine by intraperitoneal injection from a stock concentration of 10 mg/mL ketamine and 1 mg/mL xylazine solution in sterile water for injection and were given 100 µL of buprenorphine at a concentration of 0.015mg/mL by subcutaneous injection as an analgesic for post-operative pain management. An incision was made to visualize the main ducts of the submandibular and sublingual glands, Wharton’s and Bartholin’s ducts, respectively. A vascular clamp (Vitalitec/Peter’s Surgical) was applied to these ducts, and the incision was closed using two to four interrupted sutures. Mice were continuously monitored under anesthesia and post-operatively for pain, distress, and changes in weight for a minimum of 48 hours following surgery. The ducts were ligated for either 7 or 14 days. Successful 14-day ligation was determined if the gland weight was between 30 and 70% of the average historical control gland weight, and samples outside of the range were excluded from downstream analyses; no cutoff criteria was used for 7-day ligation surgeries. As controls for the C57BL/6J experiments, time point zero (T_0_) mice were euthanized at 12-weeks old and had no surgical manipulations. Mock surgery control mice received an incision, and the ducts were located, but not ligated, and mice were euthanized at 14-days post-surgery. As controls for the Gli1^lz^ mouse strain experiments non-surgically manipulated, age-, gender-, and strain-matched mice were used. As controls for the Gli1-CreER^T2^; R26^tdT^ experiments 3-week post-induction mice were used. All mice were euthanized at the desired time point under CO_2_ with secondary cervical dislocation. The submandibular and sublingual glands were immediately weighed upon removal. All subsequent sample processing was performed blind with reference only to the unique identifiers.

### 2.4 Preparation of cryosections

Submandibular and sublingual salivary glands were removed together, weighed, and fixed in 4% paraformaldehyde (PFA) in 1X phosphate buffered saline (1X PBS) for 2 hours at 4°C. Glands were washed in three consecutive 1X PBS washes, and incubated in an increasing sucrose gradient progressing from 5%, to 10%, to 15% sucrose for 1 hour each and then transferred to 30% sucrose overnight at 4°C. After overnight incubation, glands were transferred to 15% sucrose, 50% tissue freezing media (Tissue Freezing Medium, Electron Microscopy Sciences) overnight at 4°C. Glands were transferred to tissue freezing medium and then frozen over liquid nitrogen. Ten-micron (10 µm) sections were collected on Superfrost Plus glass slides (Electron Microscopy Sciences) using a Leica CM 1860 cryostat. Slides were collected by taking three consecutive 10 µm sections per slide until the entire tissue was sectioned. Sections were dried for 30 minutes at room temperature and stored at -80°C. Slides were returned to room temperature before staining.

### 2.5 Immunohistochemistry

Slides were post-fixed in 4% PFA in 1X PBS for 18 minutes at room temperature or in pre-chilled 100% methanol at -20°C for 20 minutes followed by two three-minute rinses in 1X PBS. Slides were permeabilized in 0.5% Triton-X-100 in 1X PBS for 18 minutes, followed by two three-minute rinses in 1X PBS. Sections on slides were encircled with a hydrophobic pen and blocked using 0.5% Triton-X-100 in 3% bovine serum albumin (BSA) in 1X PBS (3%BSA-0.5%TX-PBS) for 60 minutes at room temperature in a humidified chamber. Primary antibodies were diluted in 3% BSA in 1X PBS and incubated on sections for 60 minutes at room temperature or overnight at 4°C. Primary antibodies were used as documented (**Table S1**). Four five-minute rinses were performed after each round of primary antibody. Secondary antibodies (Jackson ImmunoResearch) were applied at a concentration of 1:500 in 3% BSA-0.5%TX-PBS for 60 minutes at room temperature, followed by two two-minute 1X PBS rinses. DAPI staining (Life Technologies, 1 ug/ml) was performed for ten minutes followed by two additional three-minute 1X PBS rinses. Coverslips were applied with a glycerol-based mounting media containing 4% n-propyl gallate and triethylenediamine (DABCO) as an antifade (Nelson et al. 2013). Serial multiplexed immunohistochemistry (MX-IHC) was performed for a subset of Gli1-CreER^T2^; R26^tdT^ slides, as described previously (Gerdes et al., 2013; Gervais et al., 2015; Nelson et al., 2013), with red fluorescence protein (RFP) and beta-tubulin III (βIII) in round one, vimentin, RFP, and CD31 in round two, and smooth muscle actin alpha (SMA) in round three.

### 2.6 Masson’s trichrome staining

Room temperature cryosections were fixed in Bouin’s Fixative (Polysciences) for 20 minutes. Slides were rinsed in cold tap water for five minutes and then stained in Weigert’s Iron Hematoxylin working solution (Polysciences) for five seconds. Slides were rinsed in cold tap water for two minutes and then stained in Beibrich Scarlett-Acid Fuchsin Solution (Polysciences) for 15 seconds. Slides were rinsed in deionized water (DI) until the water running off the slide was clear. Phosphotungstic/Phosphomolybdic acid (Polysciences) was added directly to the tissue sections, incubated for ten minutes, and then drained onto a Kimwipe. Slides were stained in Aniline Blue (Polysciences) for 30 seconds and rinsed in DI three times for 30 seconds. 1% Acetic Acid (Polysciences) was pipetted directly onto the sections for one minute, and then rinsed in DI. Slides were dehydrated in 95% and 100% ethanol for two minutes each, cleared in xylene for two minutes, and then mounted with Permount mounting medium (Electron Microscopy Sciences).

### 2.7 Hematoxylin and eosin (H&E) staining

Room temperature cryosections were rehydrated in 1X PBS for five minutes. Slides were rinsed under running tap water for one minute prior to staining. Slides were then stained with Hematoxylin 7211 (Richard-Allan Scientific) for ten minutes and rinsed under tap water for one minute and 30 seconds. Slides were then stained with Eosin-Y alcoholic (Richard-Allan Scientific) for three minutes and rinsed under tap water for one minute. The slides were then incubated in 100% ethanol for three minutes, allowed to air dry briefly, and mounted with Permount mounting medium (Electron Microscopy Sciences).

### 2.8 Collagen hybridizing peptide (CHP) staining

Collagen hybridizing peptide (CHP) conjugated to 5-carboxyfluorescein (5-FAM) (Advanced BioMatrix) was solubilized per manufacturer’s recommendations. For 5-FAM- CHP multiplexing with fluorescent antibodies, slides were fixed in 100% pre-chilled methanol and immunostained. While slides were in the last 1X PBS wash or after slides were stained, imaged, and had coverslips removed, 5-FAM-CHP was diluted to 20µM in 1X PBS and heated to 80°C on a heat block. After five minutes, 5-FAM-CHP was cooled in an ice water bath for one minute. One tissue section per slide was encircled with a hydrophobic pen. Each tissue section received 40µL of the 5-FAM-CHP working solution for two hours at room temperature or overnight at 4°C. Slides were rinsed with two two-minute 1X PBS washes. When needed, TrueBlack (Biotium) was applied to quench autofluorescence. When TrueBlack was applied, it was diluted to 1X in 70% ethanol and was incubated on sections for one minute at room temperature. Slides were then rinsed with two two-minute 1X PBS washes.

### 2.9 Image processing and quantification

For quantification of each marker by IHC or histological stain, sections obtained from similar tissue depths and similar regions of the submandibular glands were compared. For H&E and Masson’s-stained whole gland images, both submandibular and sublingual glands were imaged. For both Masson’s trichrome and IHC both submandibular and sublingual glands were imaged but only the submandibular gland was included in quantifications. For IHC staining, sections at the approximate midpoint of the gland were stained, imaged, and quantified. On average, three images were collected, one from the proximal, medial, and distal regions of the submandibular gland from each of three tissue sections on a slide, for a total of 9 images per mouse. IHC sections were imaged on an Olympus IX-81 microscope with a Peltier-cooled CCD camera (Q imaging, RET 4000DC-F-M-12) using a PLAN S-APO 20X, 0.75 NA objective or Zeiss Z1 Cell Observer widefield with an Axio712 mono camera (Carl Zeiss, LLC) using a Plan-Neofluar 20X/0.50 Ph2 M27 objective. All images for one imaging session were captured using one microscope with identical microscope settings and regions containing folds, tears, or unequal staining were avoided. Sections stained with Masson’s trichrome were imaged for quantification on a Hamamatsu Nanozoomer 2.0 with a LX2000 light source and TDI camera. Sections stained with H&E and Masson’s trichrome were imaged on a Leica DM 4000 B LED scope with a DFC 310 FX RGB color camera (Leica) for whole gland stitched images and 10X magnification images.

All image processing and quantification was performed using the freeware, FIJI, an imaging processing package of ImageJ (Schindelin et al., 2012). To estimate the area positive for the expression of a marker, the image was background subtracted using rolling ball background subtraction. Images from each day imaged on the same microscope were equally thresholded, and FIJI was used to quantify the tissue area that was positive for a given marker relative to total tissue area. For fluorescent images, the green channel was used to detect the total issue area by autofluorescence with no background subtraction. Colocalization analysis was performed with background subtracted images using the image calculator with the AND function to determine area present in both images. The resulting AND images were equally thresholded, and FIJI was used to quantify the area co-positive for both markers.

For trichrome quantification, the submandibular gland regions of control and ligated glands were traced using the freeform shape tool in FIJI to measure to total submandibular gland area and create a region of interest (ROI), which was defined as the total tissue area. Within the ROI a color threshold was selected to detect the blue-stained area within the ROI and was performed equally across staining groups. Positive trichrome area was normalized to the total submandibular gland area and graphed as a percent of total area. On average between 2 and 3 sections were used per mouse with a minimum of 1 section per mouse. Sections were excluded from analysis if the tissue had uneven staining that would affect the accurate measurement of blue-stained area or had folds or tears affecting greater than 50% of the section.

### 2.10 Statistical analysis

All tabulation of quantifications was performed in Microsoft Excel with graphs generated and statistics run using GraphPad Prism. For comparison of two conditions, an unpaired two-tailed t-test was performed using GraphPad Prism version 9.4.1 for Mac or Windows, GraphPad Software, San Diego, California USA, www.graphpad.com. For comparisons of more than two conditions, an Ordinary one-way analysis of variance (ANOVA) with Tukey’s multiple comparisons test was performed using GraphPad Prism. For Gli1^lz^ studies comparing both genotype and condition, a Two-Way ANOVA with Tukey’s multiple comparisons test was performed using GraphPad Prism. Statistical significance is indicated with stars over the corresponding bars with p values indicated in the figure legends. Analyses with no statistical significance (p > 0.05) are unmarked.

### 2.11 Single-cell RNA-sequencing analysis for embryonic day 16 mouse salivary glands

Single-cell RNA-sequencing of E16 stromal enriched submandibular salivary gland cells and Seurat-based clustering was previously performed (Moskwa et al., 2022). Raw data can be found on the Gene Expression Omnibus (GEO) repository under GSE181425. Percent of *Gli1*^+^ cells expressing *Gli1* only, *Gli1* and *Pdgfra*, *Gli1 and Pdgfrb*, and *Gli1* and both *Pdgfra* and *Pdgfrb* were calculated using an in-house generated function within R that calculates the number of cells expressing a given gene or combination of genes. Scripts can be accessed through GitHub: https://github.com/MLarsenLab.

### 2.12 Single-cell isolation for mock and ligated mouse salivary glands

Two ligated or mock surgery submandibular and sublingual glands were harvested, and excess fat and interstitial tissue was removed. The glands were then transferred to a dish containing 1X PBS, liberase TL Research Grade low Thermolysin (Roche), DNase I, and dispase and microdissected for 7 minutes. The sample was then incubated in a 37°C incubator for 15 minutes and triturated. After trituration, the sample was incubated for an additional 5 minutes in a 37°C incubator and triturated once more. A 15 mL conical tube was placed on ice and the entire sample was transferred to the conical tube and incubated for 10 minutes. The supernatant was isolated and transferred to a fresh conical tube and was centrifuged for 5 minutes at 450xg. The supernatant was removed and discarded, and the cell pellet was resuspended in isolation buffer composed of sterile-filtered Ca^2+^- and Mg^2+^-free 1X PBS, 0.1% bovine serum albumin, and 2mM ethylenediaminetetraacetic acid. 5 μg of EpCAM-A647 (catalog: 118212, BioLegend) and Ter119 (catalog: 50-133-27, eBioscience/ThermoFisher) was added to the cell suspension and incubated at 4°C for 10 minutes for subsequent depletion of epithelial and red blood cells, respectively. The cell suspension was washed using isolation buffer and centrifuged at 450xg for 5 minutes. The supernatant was discarded, and the cell pellet was resuspended in 1 mL of isolation buffer and 25 μL of sheep anti-rat Dynabeads, which bind to the EpCAM and Ter119 labeled cells, (catalog: 11035, Invitrogen) before incubating on ice for 20 minutes on a rocker. The sample was placed on a microcentrifuge magnet for 2 minutes, and the supernatant was transferred to a fresh microcentrifuge tube to remove labeled epithelial and red blood cells. One additional epithelial and red blood cell depletion was performed. The sample was then depleted of dead cells using a dead cell removal kit (catalog: 130-090-101, Miltenyi Biotec) following manufacturer’s instructions. Cells were counted and resuspended to a concentration of 1000 cells/μL. Following the manufacturer’s protocol for the Chromium Next GEM Single Cell 3’ Reagent kits v3.1, scRNA-seq libraries were generated.

### 2.13 Single-cell RNA-sequencing analysis for mock and ligated mouse salivary glands

Mock and ligated samples were sequenced (the ligated sample was sequenced on the Nextseq500 and mock on Nextseq2000), and initial processing of the datasets was performed at the Center for Functional Genomics at the University at Albany. Initial processing steps included, generating FASTQ files, aligning to the genome and generating counts files, which were completed using CellRanger version 6.0.1. Data files were imported using Seurat v3.1.5 in R v3.6.3 (“R: The R Project for Statistical Computing,” n.d.; Stuart et al., 2019). Data clusters were calculated following the default pipeline (“Seurat - Guided Clustering Tutorial,” n.d.). Dead or apoptotic cells were removed if >5% of unique molecular identifiers (UMIs) mapped to mitochondrial genes. Any cells with <200 or >9000 genes were excluded to select for single cells. The processed datasets were saved as RDS files and imported into another workspace using Seurat v4.1.1 in R v4.1.2 (Hao et al., 2021). After importing, the mock and ligated datasets were merged. We were unable to perform dataset integration as there were dropout reads for important transcripts such as *Gli1* and *PDGFRβ*. The functions ‘ElbowPlot’, principal component ‘Heatmaps’, and ‘JackStrawPlot’ were used to determine the principal components for unsupervised cluster modeling. Cell clusters were determined using linear dimensional reduction and visualization was performed using uniform manifold projection (UMAP). 14 dimensions were used for the merged dataset. Further analysis was performed on the stromal subset of this dataset containing cells expressing *Pdgfra* >0.5, or *Pdgfrb* >0.5, or *Gli1* >0. For the stromal subset, 14 dimensions were used, generating 5 clusters. After clustering, the Seurat V4.1.1 package was used to create plots based on single-cell gene expression.

## 3 Results

### 3.1 Stromal cells and ECM area are expanded with fibrotic injury

We performed ductal ligation surgeries on submandibular (SMG) and sublingual (SLG) glands of 12-week-old female C57BL/6J mice. We harvested glands from mice at 7- days post-ligation (7-Day) or 14-days post-ligation (14-Day) (**Figure 1A**). We compared the ligated glands to negative controls, including both unmanipulated glands harvested at the time of surgery, or time-point zero (T_0_), and glands subjected to a mock surgery with no ligation and harvested at the 14-day timepoint (Mock). Upon harvest, we weighed each SMG and SLG together and normalized their weight to total mouse weight. We found a 36% decrease in gland weight for 14-Day glands when compared to T_0_ controls and a 44.8% decrease in normalized gland weight when compared to Mock controls (**Figure 1C**). The average normalized and unnormalized weight was similar in 7-Day and 14-Day ligated glands (**Figures 1C** **and S1B**). Following ligation injury, we detected acinar cell atrophy, as indicated by hematoxylin and eosin (H&E)-staining in comparable tissue regions with mock controls (**Figures 1B** **and S1A**), similar to previous reports (Cotroneo et al., 2008).

**Figure 1.**
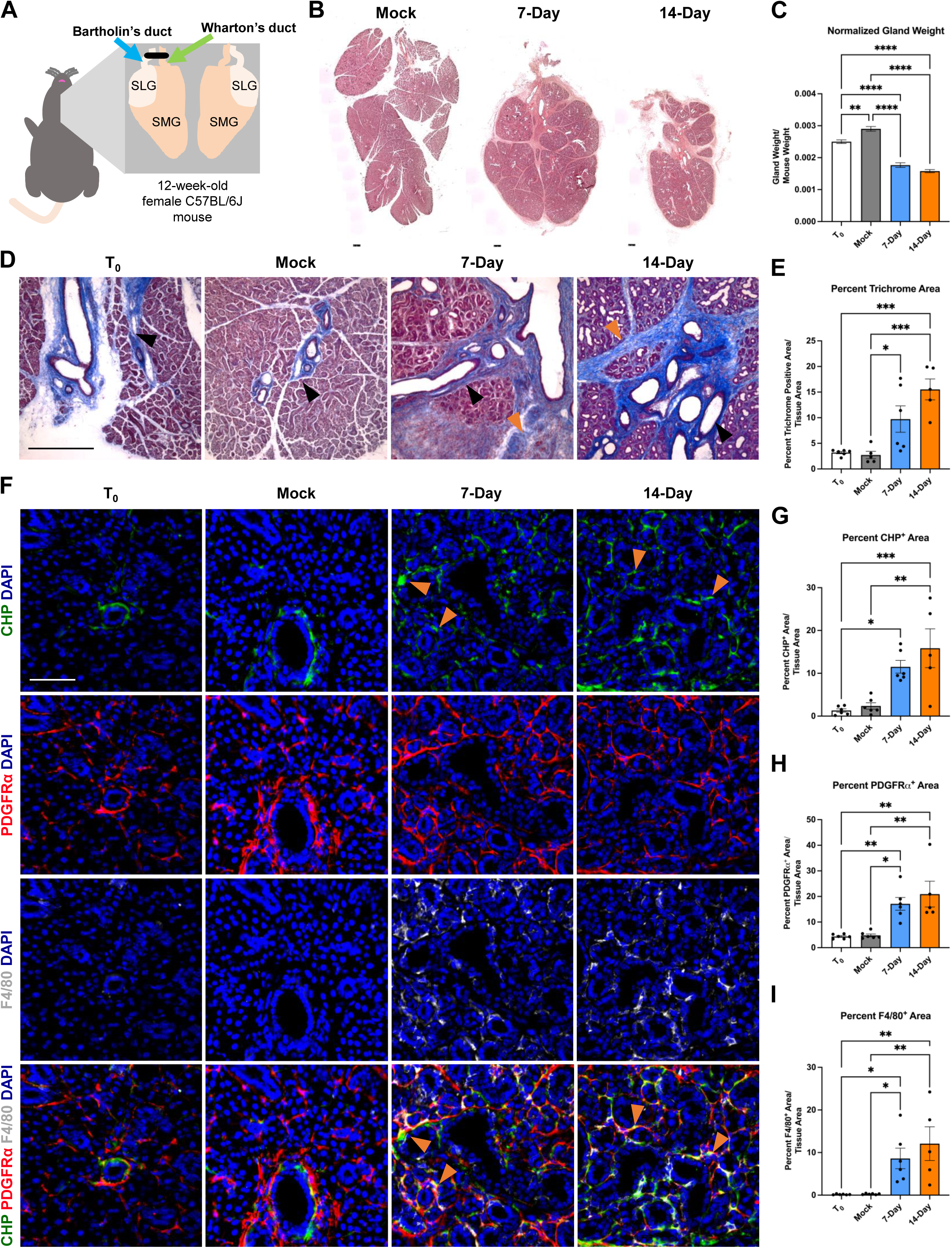
A progressive and dynamic fibrotic stromal response occurs in response to ductal ligation surgery. (A) Schematic of ductal ligation surgery showing clip application on the main ducts of the left submandibular and sublingual glands of a 12-week-old female C57BL/6J mouse. (B) H&E-stained whole gland images of 14-day mock (Mock), 7-Day ligated (7-Day), and 14-Day ligated (14-Day) mice. Scale bar 250 µm. (**C**) Gland weights normalized to mouse weight in time-point zero (T_0_), Mock, 7-Day, and 14-Day glands. N = 10, 25, 6, and 34, respectively. (**D**) Trichrome staining shows deposited ECM in blue with a trend of increasing ECM from 7- to 14-Day ligation. Black arrowheads denote regions of ECM present in both ligated and control conditions adjacent to large ducts and vasculature, while orange arrowheads denote expanded regions of ECM present only in the ligated glands, specifically in interlobular regions. (**E**) Quantification of percent trichrome area relative to tissue area in T_0_, Mock, 7-Day, and 14-Day glands. N = 6, 5, 6, 5 respectively. (**F**) Collagen hybridizing peptide (CHP), in green, labeling actively remodeled collagen, together with immunohistochemistry to detect platelet-derived growth factor receptor alpha (PDGFRα), in red, labeling a subset of the stromal cells, and F4/80, in grey, labeling macrophages with nuclear staining (DAPI) in blue. A subset of actively remodeled collagen is located near PDGFRα^+^ cells and/or F4/80^+^ macrophages (orange arrowheads). Quantification of percent stain area normalized to total tissue area for (**G**) CHP (**H**) PDGFRα and (**I**) F4/80 N = 6, 6, 6, 5 respectively. Error bars: S.E.M. One-way ANOVA followed by Tukey’s multiple comparisons test was performed using GraphPad Prism version 9.4.1. *p≤ 0.05, **p≤ 0.01, ***p≤ 0.001, ****p≤ 0.0001. Scale bars 100 µm.

Previous studies reported an increase in extracellular matrix (ECM) accumulation seven days after ductal ligation (Woods et al., 2015a); however, the extent of ECM deposition over time was not examined. To quantify the fibrotic progression following ligation we stained cryosections from T_0_, Mock, 7-Day, and 14-Day salivary glands using Masson’s trichrome and quantified the percent trichrome-positive area relative to total tissue area. We found a significant increase in ECM present in both 7-Day and 14-Day glands compared to Mock (**Figures 1E** **and S1C**). 7-Day glands had a three-fold increase in the percent trichrome-positive area, as compared with T_0_ and Mock glands, while 14-Day glands had an approximate five-fold increase in trichrome-positive area compared with T_0_ and Mock, and a 1.6-fold increase compared to 7-Day glands (**Figure 1E**). This suggests that the ECM accumulation trends upwards from 7-day ligation to 14-day ligation. Ductal ligation injury not only increased ECM accumulation around large ducts and vasculature, where it is normally present (**Figure 1D, black arrowheads**), but also expanded the ECM-positive area in regions between the epithelial lobes of the gland (**Figure 1D, orange arrowheads**).

Fibrosis can also be characterized by aberrant ECM remodeling, including active collagen remodeling (L. Li et al., 2018; Wu et al., 2021). To evaluate active collagen remodeling, we exposed tissue sections to a collagen hybridizing peptide (CHP) chemically conjugated to the fluorophore 5-carboxyfluorescein (5-FAM). The single chain collagen-like peptide hybridizes with denatured or partially denatured fibrillar collagen (**Figure 1F** **top panel**). In the SMG, we measured approximately a 10-fold increase in active collagen remodeling in 14-Day glands compared with T_0_ glands (**Figure 1G**). The increase in active collagen remodeling with 14-Day glands was present across distal, medial, and proximal regions of the gland and across medial and proximal regions in 7-Day glands (**Figures S2A-C**). Together, this data indicates that there is an upward trend of increased ECM deposition and active collagen remodeling from 7-days to 14-days post-ligation.

The cells that contribute to the fibrotic response following salivary gland ductal ligation injury have not been defined. However, stromal cell populations are known to contribute to fibrosis directly or indirectly via changes in myofibroblast populations (Bonner, 2004; Buhl et al., 2020; Deng et al., 2021; Kishi et al., 2018; R. Li et al., 2018). We investigated the stromal cell population expressing platelet-derived growth factor receptor alpha (PDGFRα) because PDGFRα^+^ stromal cells produce ECM in the embryonic salivary gland (Moskwa et al., 2022) and drive fibrotic responses in other organs (Santini et al., 2020). To determine if PDGFRα^+^ cells expand following ductal ligation injury, we performed immunohistochemistry (IHC) and quantified the regions of the gland positive for PDGFRa signal. The percent area of the SMG positive for PDGFRα increased three-fold in 7-Day glands and four-fold in 14-Day glands compared to T_0_ and Mock controls (**Figure 1F** **2^nd^ panel and 1H**). This increase was seen across the distal, medial, and proximal regions of the gland (**Figures S2D-F**). These data suggest that the number of cells expressing PDGFRα increases in response to injury.

Macrophages are active contributors to injury and fibrosis by promoting inflammatory responses and stimulating ECM remodeling (Vasse et al., 2021). We quantified the percent gland area positive for the pan-macrophage membrane marker, F4/80, and found that the F4/80^+^ area increased 49-fold in 7-Day glands and 69-fold in 14-Day glands compared to T_0_ controls. However, the response was variable, with some glands retaining high levels of macrophages and some having lower levels (**Figure 1F** **3^rd^ panel and 1I**). There was also regional variability in the F4/80^+^ area with higher levels of macrophages detected in the proximal region at 7-days and higher levels in the medial regions by 14-days with similar levels in the distal region at both time points (**Figures S2 G-I**). Although there was little co- localization of PDGFRα or F4/80 with CHP (**Figure 1F** **4^th^ panel**), PDGFRα^+^ cells and macrophages were in the proximity of the actively remodeled collagen (**Figure 1F** **4^th^ panel, orange arrowheads**), consistent with a contribution of both cell types to the fibrotic response and active ECM remodeling we detect in this model.

### 3.2 Gli1^+^ stromal cells are a subset of the PDGFRα/β^+^ cell populations at E16 and expand with ligation injury in adult SMG

In many injury and disease conditions, it is the activation of specific stromal cell subpopulations that induces organ fibrosis (Deng et al., 2021; El Agha et al., 2017). A subset of the PDGFRβ^+^ stromal population that expresses the sonic hedgehog (Shh) effector protein, Gli1, have been implicated in organ fibrosis in the lung (Cassandras et al., 2020), bone marrow (Manshouri et al., 2022; Schneider et al., 2017b), kidney, liver, and heart (Kramann et al., 2015b). Gli1^+^ cells have previously been shown to comprise a small subset of the PDGFRβ^+^ cell population in the adult lung, heart, kidney, and bone chips and have been associated with PDGFRα^+^ expression in the heart and kidney (Kramann et al., 2015b). To identify the Gli1^+^ cells in the salivary gland, we utilized single-cell RNA-sequencing (scRNA-seq) data from embryonic day 16 salivary glands (Moskwa et al. 2022). We identified a small population of 103 *Gli1*^+^ cells in the embryonic day 16 (E16) SMG stromal populations, of which 48.54% co-express *Pdgfra*, 8.74% co-express *Pdgfrb,* and 33.01% co-express both *Pdgfra* and *Pdgfrb* (**Figures 2A-D**). *Gli1*^+^ cells thus represent a small subpopulation of stromal cells in the developing salivary gland.

**Figure 2.**
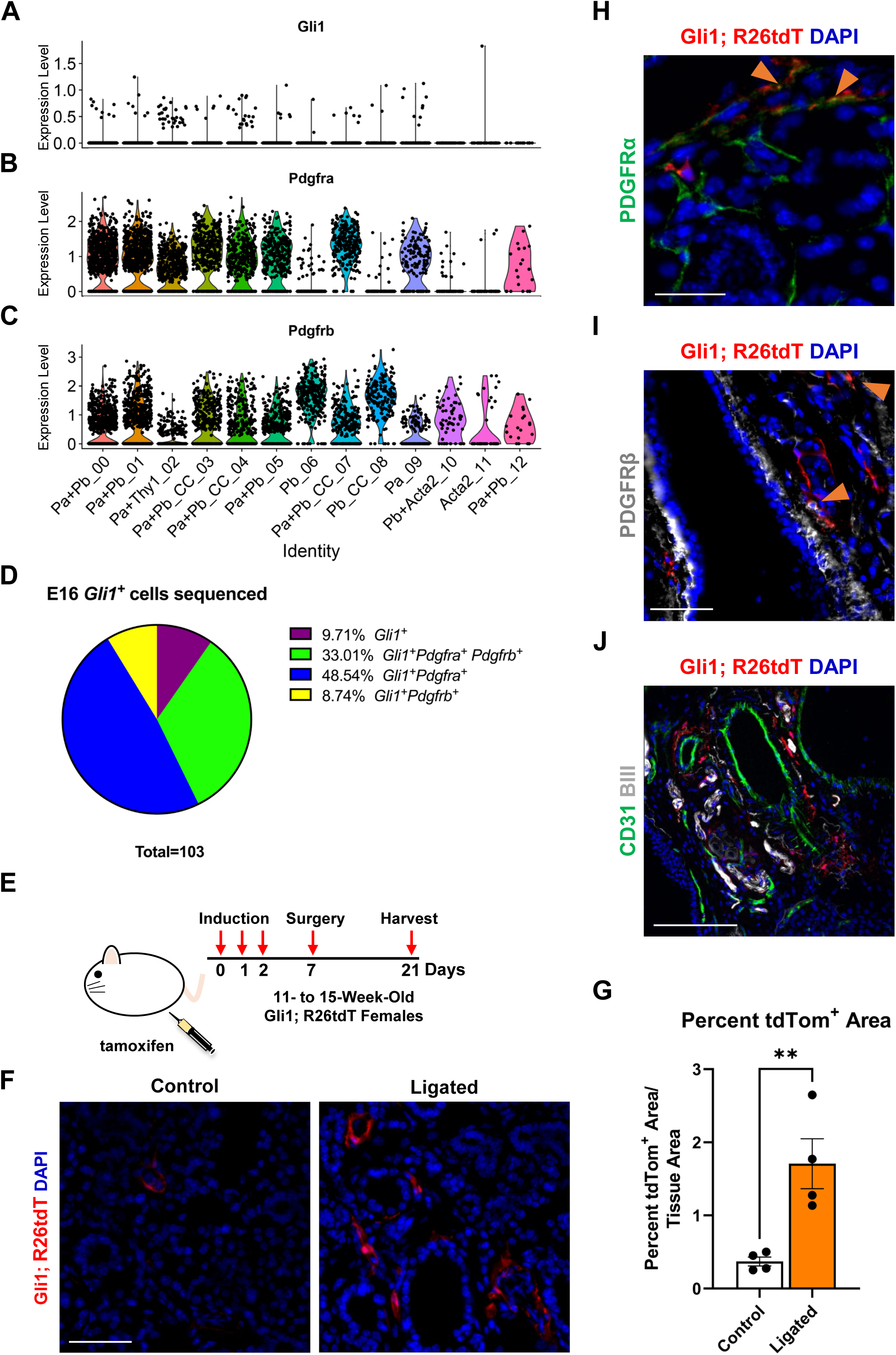
Gli1^+^ cells, which are a subset of PDGFRα^+^ and PDGFRβ^+^ cells, reside in the neurovascular region and expand with injury. Violin plots of stromal enriched scRNASeq data subsetted for stromal cells from an embryonic day 16 (E16) SMG and SLG, highlighting cells expressing transcripts for (**A**) *Gli1* (**B**) *Pdgfra* and (**C**) *Pdgfrb*. (**D**) Pie chart depicting percentages of *Gli1*^+^ cells that express *Gli1* only, *Gli1* and *Pdgfra*, *Gli1* and *Pdgfrb*, or *Gli1*, *Pdgfra*, and *Pdgfrb* from E16 stromal enriched scRNASeq subsetted for stromal cells. (**E**) Schematic showing tamoxifen-induction scheme for 11- to 15-week-old adult female Gli1^tm3(cre/ERT2)Alj^/J;(CAG)ROSA26^tdTomato^ (Gli1;R26tdT) lineage reporter mice harvested 3 weeks post-initial induction or with 14 day ductal ligation surgery performed 1 week after induction. (**F**) Immunohistochemistry (IHC) to detect Gli1;R26tdT lineage-traced cells (red) with nuclei (DAPI) in blue in control 3 week induced and 14 day ligated mouse SMG. (**G**) Quantification of percent Gli1;R26tdT^+^ area normalized to tissue area in control 3 week induced and 14 day ligated mouse SMG. (**H**) IHC to detect Gli1;R26tdT lineage-traced cells (red) in adult SMG together with PDGFRα (green) with nuclei (DAPI) in blue. Orange arrowheads represent colocalization of Gli1;R26tdT with PDGFRα. Scale bar 25 µm. (**I**) IHC to detect Gli1;R26tdT lineage-traced cells (red) in adult SMG together with PDGFRβ (grey) with nuclei (DAPI) in blue. Orange arrowheads represent colocalization of Gli1;R26tdT with PDGFRβ. (**J**) IHC to detect Gli1;R26tdT (red), CD31 (green), and βIII (grey), shows that Gli1;R26tdT-derived cells are enriched in areas near large ducts, large vessels, and nerve bundles in the proximal region of the adult SMG. Scale bar 100 µm. N=4. Error bars: S.E.M. Statistical tests were performed using GraphPad Prism version 9.4.1. ** p≤0.01.

To determine if Gli1^+^ cells expand with injury in adult SMGs we used a tamoxifen-inducible Gli1-CreER^T2^;R26^tdT^ lineage tracing mouse (**Figure 2E**) to trace cells after ductal ligation injury. We used Gli1^tm3(cre/ERT2)Alj^/J (Gli1-CreER^T2^) mice, where tamoxifen-sensitive Cre recombinase is driven by the Gli1 promoter (Ahn and Joyner, 2004) crossed to a reporter strain B6.Cg-Gt(ROSA)26Sor^tm9 (CAG-tdTomato)Hze^ /J to label all Gli1-expressing cells prior to surgery. As three doses of tamoxifen were administered one week prior to surgery, cells that express Gli1 at the time of induction and all their subsequent progeny were labeled with tdTomato (tdTom). Glands were harvested 14-days after ductal ligation and compared to 3-week induced controls with no surgical manipulation. After ligation injury, there was a 4.6-fold increase in percent tdTom^+^ area compared to 3-week induced controls, suggesting that Gli1 cells respond to ductal ligation injury (**Figure 2F and G**). Next, to determine if the Gli1^+^ cells express PDGFRα and/or PDGFRβ in the adult, similar to the embryonic mice, we again used the tamoxifen-inducible Gli1-CreER^T2^;R26^tdT^ lineage tracing mouse. In SMGs of mice three weeks after induction, we found that approximately 5% of the tdTomato-expressing cells also expressed PDGFRα (**Figure 2H**) and 8% of tdTomato expressing cells also expressed PDGFRβ (**Figure 2I**) by IHC (Quantification not shown). Gli1^+^ cells have been shown to localize near nerves and vasculature in other organs (Kramann et al., 2015b; Zhao et al., 2014a). To determine where Gli1^+^ cells are located in the salivary glands of adult mice, we performed IHC in Gli1-CreER^T2^;R26^tdT^ glands three weeks after tamoxifen induction. We identified Gli1 lineage-traced cells within regions close to βIII^+^ neurons and CD31^+^ endothelial cells in the vasculature (**Figure 2J**). These data indicate that Gli1^+^ cells are retained in adult salivary glands and can be detected near the neurovascular regions of homeostatic glands, similar to other organs (Kramann et al., 2015a; Schneider et al., 2017b).

### 3.3 Gli1 null mice have decreasing trends of ECM deposition and active collagen remodeling following ductal ligation

As the Gli1 lineage-traced cells expanded with ductal ligation injury, they may have a function in the injury response. Previous studies of the mouse incisor demonstrated that Gli1^+^ cells contribute to injury repair (Zhao et al., 2014a), suggesting that Gli1 cells may contribute to ligation injury. In addition, inhibition of Gli1 signaling can inhibit fibrosis (Schneider et al., 2017b), and several components of hedgehog signaling including *Smo*, *Ptch*, *Gli1*, *2*, and *3,* were increased at the mRNA level, in the whole gland of male mice 6 days after ligation (Hai et al., 2010). We utilized a genetic null Gli1 line and examined the adult female SMG in a homeostatic state and after ligation injury. In the Gli1^tm2Alj^/J (Gli1^lz^) mouse strain, the zinc-finger domains and Gli1 N-terminal fragment in exons 2-7 are replaced with the LacZ gene (Bai et al., 2002b), mice homozygous for *LacZ (lz*) are *Gli1* null as they do not express a functional Gli1 protein (**Figure 3A**). We examined the weight of 10- to 19-week-old adult salivary glands from Gli1^wt/wt^, Gli1^wt/lz^, and Gli1^lz/lz^ animals with no surgical manipulation (T_0_) and found no difference (**Figure 3B**). We next performed ductal ligation on 10- to 18-week-old Gli1^wt/wt^, Gli1^wt/lz^, and Gli1^lz/lz^ salivary glands (Ligated). Although all ligated glands weighed significantly less than control glands (**Figure 3B**), there was no significant difference in gland weight as a function of genotype.

**Figure 3.**
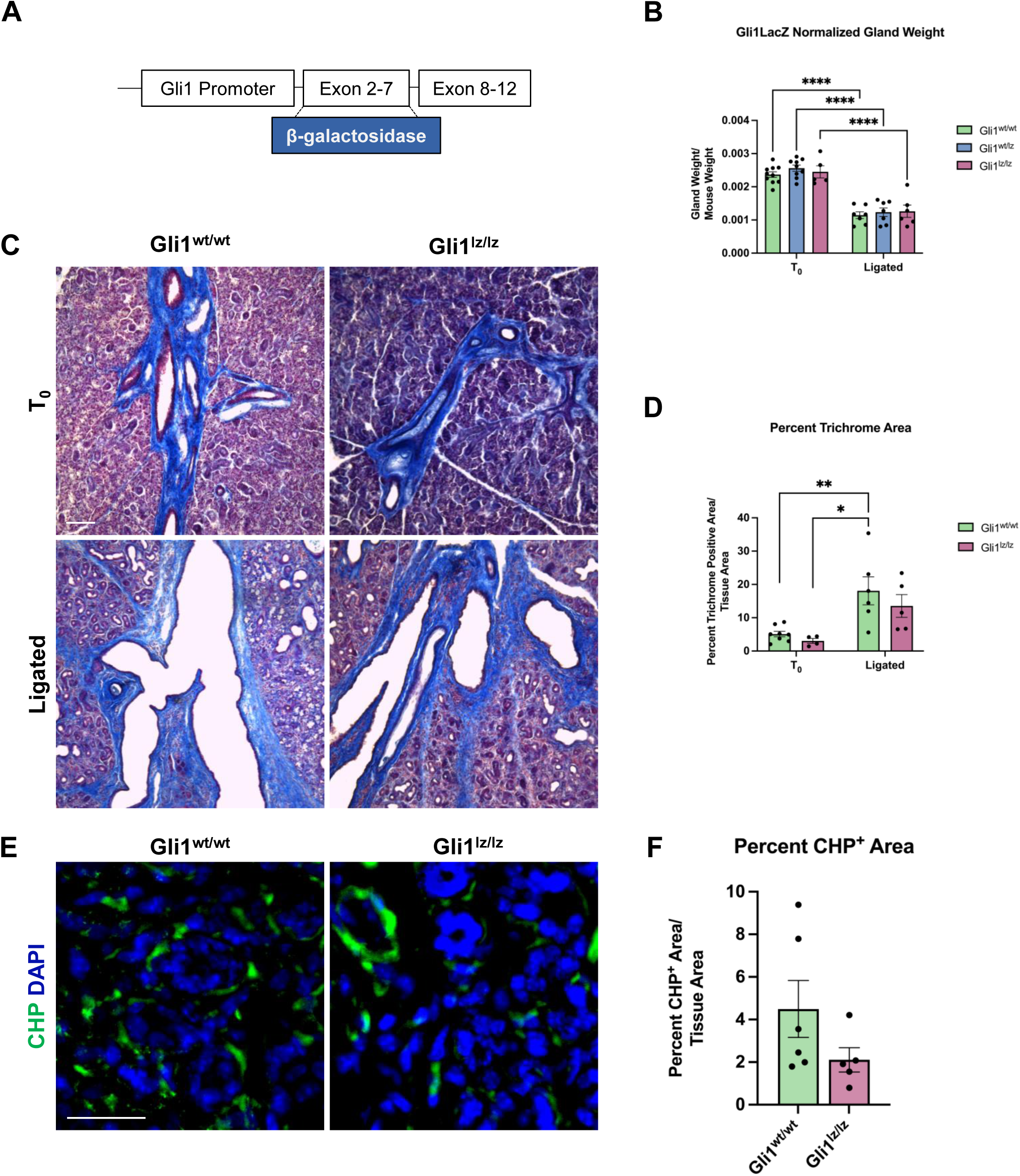
Genetic knockout of Gli1 reduces trend in ECM deposition and remodeling. (**A**) Schematic of the Gli1 gene, showing insertion of the beta-galactosidase gene (β- galactosidase) in exons 2-7 of the gene, resulting in a non-functional protein in Gli1^lz/lz^ mice. (**B**) Left SMG and SLG weight normalized to mouse weight for control (T_0_) and 14- day ligated (Ligated) wildtype (Gli1^wt/wt^), heterozygous (Gli1^lz/wt^) and homozygous knockout (Gli1^lz/lz^) mice. N = 10, 9, and 5 for T_0_ and 7, 7, and 6 for Ligated, respectively. All T_0_ glands are statistically larger than ligated glands and for simplicity only comparisons from mice of the same genotype are displayed on the graph. Statistical Test: Two-way ANOVA followed by Tukey’s multiple comparisons test ****p≤ 0.0001. (**C**) 10X magnification of Masson’s trichrome-stained T_0_ and Ligated Gli1^wt/wt^and Gli1^lz/lz^ mice. Scale bar 100 µm. (**D**) Quantification of percent trichrome area normalized to total tissue area. Statistical Test: Two-way ANOVA followed by Tukey’s multiple comparisons test *p≤ 0.05 ** p≤ 0.01. N = 8 and 4 for T_0_ and 6 and 5 for Ligated, respectively. (**E**) Immunohistochemistry on Ligated Gli1^wt/wt^ and Gli1^lz/lz^ knockout mice for collagen hybridizing peptide (CHP) in green with nuclear staining (DAPI) in blue. Scale bar 25 µm. (**F**) Quantification of percent stain area normalized to total tissue area for CHP. Statistical Test: Unpaired two-tailed t-test was performed. Ligated Gli1^wt/wt^ N = 6 and Gli1^lz/lz^ N = 5. Error bars: S.E.M. Statistical tests were performed using GraphPad Prism version 9.4.1.

To determine if ECM deposition after injury was affected by the lack of Gli1, we performed Masson’s trichrome and quantified the percent trichrome positive area to total tissue area for Gli1^wt/wt^ and Gli1^lz/lz^ T_0_ and Ligated mice. We found that Gli1^wt/wt^ mice showed a 3.7-fold increase in trichrome positive area following ligation and Gli1^lz/lz^ mice showed a 4.43-fold increase in trichrome positive area compared to the respective T_0_ controls (**Figures 3C and 3D**). There was no significant difference in ECM deposition across genotypes in T_0_ glands or ligated glands; however, Gli1^lz/lz^ glands displayed a downward trend in ECM deposition compared to Gli1^wt/wt^ (**Figure 3C and D**). Since active collagen remodeling occurs following ductal ligation, we examined active collagen remodeling in the absence of Gli1. We exposed ligated Gli1^wt/wt^ and Gli1^lz/lz^ gland sections to CHP, but we found no statistical difference in active collagen remodeling between the two genotypes within the whole gland or in regional analysis (**Figures 3E-F** **and S3A-C).** However, there was a trend downwards of active collagen remodeling in the proximal, medial, and distal regions of the gland, following a similar patten to total ECM. This data suggests that Gli1 signaling has a minor role in inducing ECM production and collagen remodeling following ductal ligation injury.

### 3.4 Genetic knockout of Gli1 does not significantly alter cell populations in ligated SMGs

Since the PDGFRα^+^ area of the gland expands with ligation, and some Gli1 cells express PDGFRα and β, we next determined if the absence of Gli1 affected the amount of PDGFRα^+^ or PDGFRβ^+^ cells in the SMG following ductal ligation. Relative to Gli^wt/wt^ mice, Gli1^lz/lz^ null mice showed no significant change in PDGFRα^+^ or PDGFRβ^+^ area within the total SMG, however in the medial region, PDGFRα^+^ area was significantly lower in Gli1^lz/lz^ null mice compared to Gli^wt/wt^ mice (**Figures 4A** **panel 1, 4B, 4C, 4E, and S4A-F**). We also examined cells that may be affected by lack of Gli1 signaling in Gli1 cells, including F4/80^+^ macrophages (**Figure 4A** **panel 2 and 4D**), βIII^+^ neurons (**Figure 4F** **panel 1 and 4G**), and CD31^+^ endothelial cells (**Figure 4F** **panel 2 and 4H**). We found no difference in percent area of the gland expressing these cell type markers in Gli1 null mice relative to WT mice following ligation. These data suggest that the lack of Gli1 signaling does not significantly alter specific subsets of stromal populations after ligation but does reduce the PDGFRa^+^ cells in the medial region of the gland.

**Figure 4.**
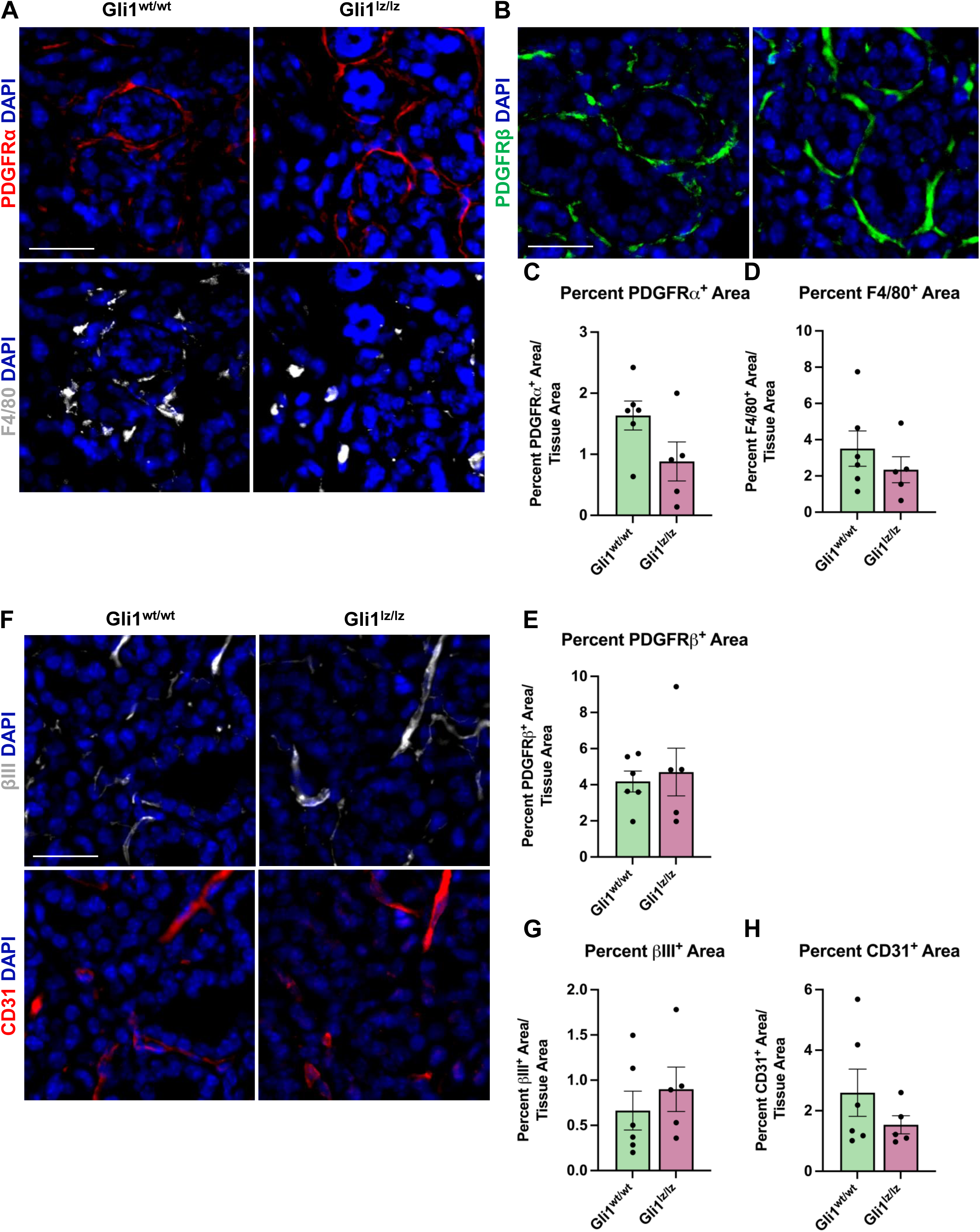
Genetic knockout of Gli1 does not significantly alter stromal cell populations. (**A**) Immunohistochemistry on 14-day ligated wildtype (Gli1^wt/wt^) and homozygous knockout (Gli1^lz/lz^) mice to detect platelet-derived growth factor receptor alpha (PDGFRα) in red and F4/80 in grey or (**B**) platelet-derived growth factor beta (PDGFRβ) in green or (**F**) beta-III tubulin (βIII) in grey and platelet endothelial cell adhesion molecule (CD31 or PECAM1) in red with nuclear staining (DAPI) in blue. Scale bar 25 µm. Quantification of percent stain area normalized to total tissue area for (**C**) PDGFRα (**D**) F4/80 (**E**) PDGFRβ (**G**) βIII and (**H**) CD31. Statistical Test: Unpaired two-tailed t-test was performed. Gli1^wt/wt^ N = 6 and Gli1^lz/lz^ N = 5. Error bars: S.E.M. Statistical tests were performed using GraphPad Prism version 9.4.1.

### 3.5 Gli1 lineage-traced cells show increased colocalization with vimentin and PDGFRβ following ligation injury

As previous studies in the kidney and heart found that Gli1^+^ cells expand following injury and can differentiate into myofibroblasts, thereby directly contributing to organ fibrosis (Kramann et al., 2015b), we looked for evidence of expansion of Gli1-derived cells with expression of myofibroblast markers. To examine changes in cell fate, we used Gli1^tm3(cre/ERT2)Alj^/J (Gli1-CreER^T2^) mice. We induced 11- to 15-week-old female Gli1- CreER^T2^;R26^tdT^ mice with tamoxifen one week prior to ductal ligation surgery (**Figure 2E**). Since Gli1 or Gli1-derived cells can undergo a myofibroblast conversion in response to injury, we looked for myofibroblast differentiation using the classic myofibroblast marker, smooth muscle alpha actin (SMA). We examined cells that expressed Gli1 at the time of ligation or were derived from such cells based on their tdTomato expression and performed IHC to detect SMA but found no significant change in colocalization between Gli1-derived cells and SMA in the whole gland or regionally (**Figures 5A and 5B**). Additionally, in the whole gland, we detected no significant change in SMA following ligation (**Figure S1D**), suggesting that stromal cells in general and Gli1-expressing cells specifically, may not undergo a conventional myofibroblast conversion during ductal ligation injury. However, as a subset of myofibroblasts have been reported to express the mesenchymal marker vimentin (Bagalad et al., 2017), we performed IHC to detect vimentin and measured the colocalization with tdTomato. Overall, vimentin area did not significantly change with ligation injury (**Figure S1E**). However, we measured a 1.4-fold significant increase in colocalization between vimentin and Gli1-lineage traced cells following ductal ligation with 16% of Gli1- derived cells expressing vimentin in the whole gland (**Figure 5C and 5D**) and regional differences in the distal and medial regions of the gland (**Figures S5A-C**). These data reveal that a small subset of Gli1-derived cells increase vimentin in response to injury, which may be indicative of these Gli1^+^ cells undergoing a partial myofibroblast transition.

**Figure 5.**
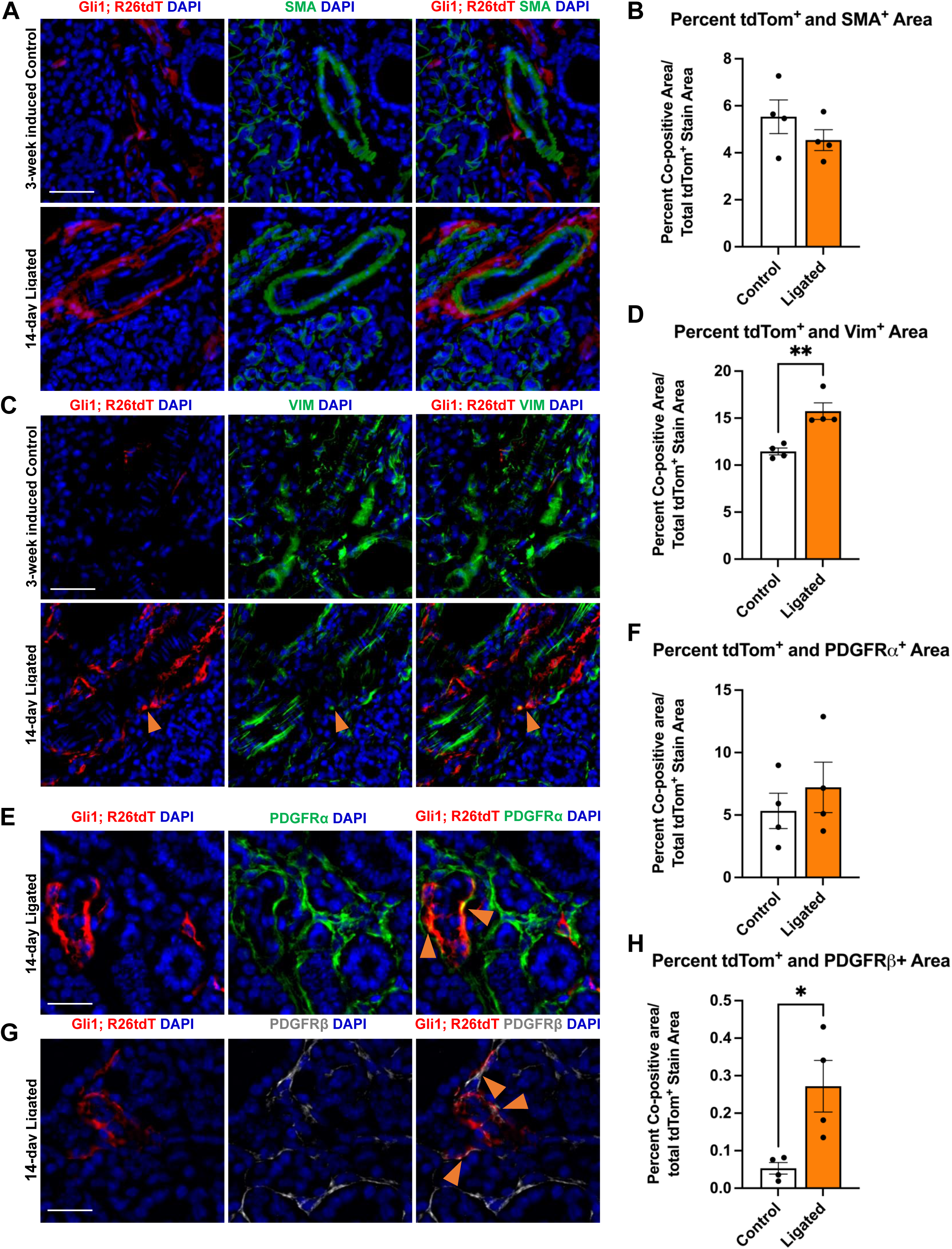
Gli1-linage traced cells show increased colocalization with vimentin and PDGFRβ after ligation injury. Immunohistochemistry (IHC) to detect Gli1; R26tdT-lineage traced cells (red) in SMG together with (**A**) SMA (green) or (**C**) vimentin (green) with nuclei (DAPI) in blue in 3- week induced control (Control) or 14-day ligated (Ligated) mice. Quantification of Gli1; R26tdT lineage traced cells with (**B**) SMA and (**D**) vimentin. IHC to detect Gli1; R26tdT- lineage traced cells (red) in SMG together with (**E**) PDGFRα (green) or (**G**) PDGFRβ (grey) with nuclei (DAPI) in blue in 14-day Ligated mice. Orange arrows represent colocalization between Gli1; R26tdT^+^ cells and PDGFRα or β. Quantification of Gli1; R26tdT lineage-traced cells with (**F**) PDGFRα and (**H**) PDGFRβ, showing a significant increase in colocalization between Gli1; R26tdT and PDGFRβ. N = 4. Scale bars 50 µm. Error bars: S.E.M. Statistical Test: Unpaired two-tailed t-test was performed using GraphPad Prism version 9.4.1. * p≤0.05 and **p≤ 0.01.

Since *Gli1*^+^ cells are a subset of the *Pdgfra* and *Pdgfrb* population in the embryonic gland and in other organs, we used IHC to determine if the Gli1-derived cells themselves show altered colocalization of PDGFRα or PDGFRβ following ductal ligation injury. We performed IHC to detect colocalization of PDGFRα and PDGFRβ in Gli1 lineage-traced cells. While the Gli1 and PDGFRα co-positive cells were maintained at similar levels in control and ligated mice in the whole gland (**Figures 5E and 5F**), the Gli1-derived cell population that co-expresses PDGFRβ expanded five-fold post-ligation surgery in the whole gland (**Figures 5G and 5H**), with a significant increase being present in the distal region of the gland and an increasing trend in the medial and proximal regions of the gland (**Figures S5D-F**). These data indicate that a subset of Gli1-derived cells increase PDGFRβ expression in response to ductal ligation injury.

### 3.6 *Pdgfra* expressing stromal cell subsets highly express matrisome-associated genes

To identify transcriptome changes that may correlate with the partial change in phenotype following injury, we performed scRNA-seq on 14 day-ligated and mock glands. The two datasets were merged and analyzed to look at differentially expressed genes between similar cell populations in ligated or mock samples, and specifically the *Gli1*- expressing population. Using Seurat and limiting to 14 dimensions, we generated uniform manifold approximation and projections (UMAPs) to produce 12 clusters of cells and primarily used these known marker genes *Pdgfra* (stromal cells)*, Pdgfrb* (stromal cells)*, Pecam1* (endothelial cells)*, Adgre1* (macrophage cells)*, Col1a1* (fibroblast cells)*, Epcam* (epithelial cells)*, Ptprcap* (immune cells)*, Cd74* (macrophages), and *Gli1* to identify the populations (**Figures S6A-C**). *Gli1* cells were mainly localized to cluster 3, which was composed of stromal cells (**Figure S6C**). Overall, there were very few cells expressing *Gli1* in both conditions, but we did detect an overall increase in the number of *Gli1*-expressing cells following ligation (mock n=17 cells, ligated n=57 cells) (**Figure 6B**), consistent with IHC analysis. We performed differential gene expression analysis using the *Gli1* cells and examined a subset of genes expressed in the matrisome. The matrisome is defined as known and bioinformatically predicted ECM components and associated proteins which includes collagens, proteoglycans, and glycoproteins (Hynes and Naba, 2012; Naba et al., 2012, 2016, 2017; Shao et al., 2020). Since specific components of the matrisome are increased in fibrotic responses, we decided to see if changes were present in *Gli1* cells (Ghatak et al., 2015; Ricard-Blum et al., 2018). There was not a change in expression of ECM genes encoding fibrillar proteins or glycoproteins by *Gli1*-expressing cells following ductal ligation (**Table S2**).

**Figure 6.**
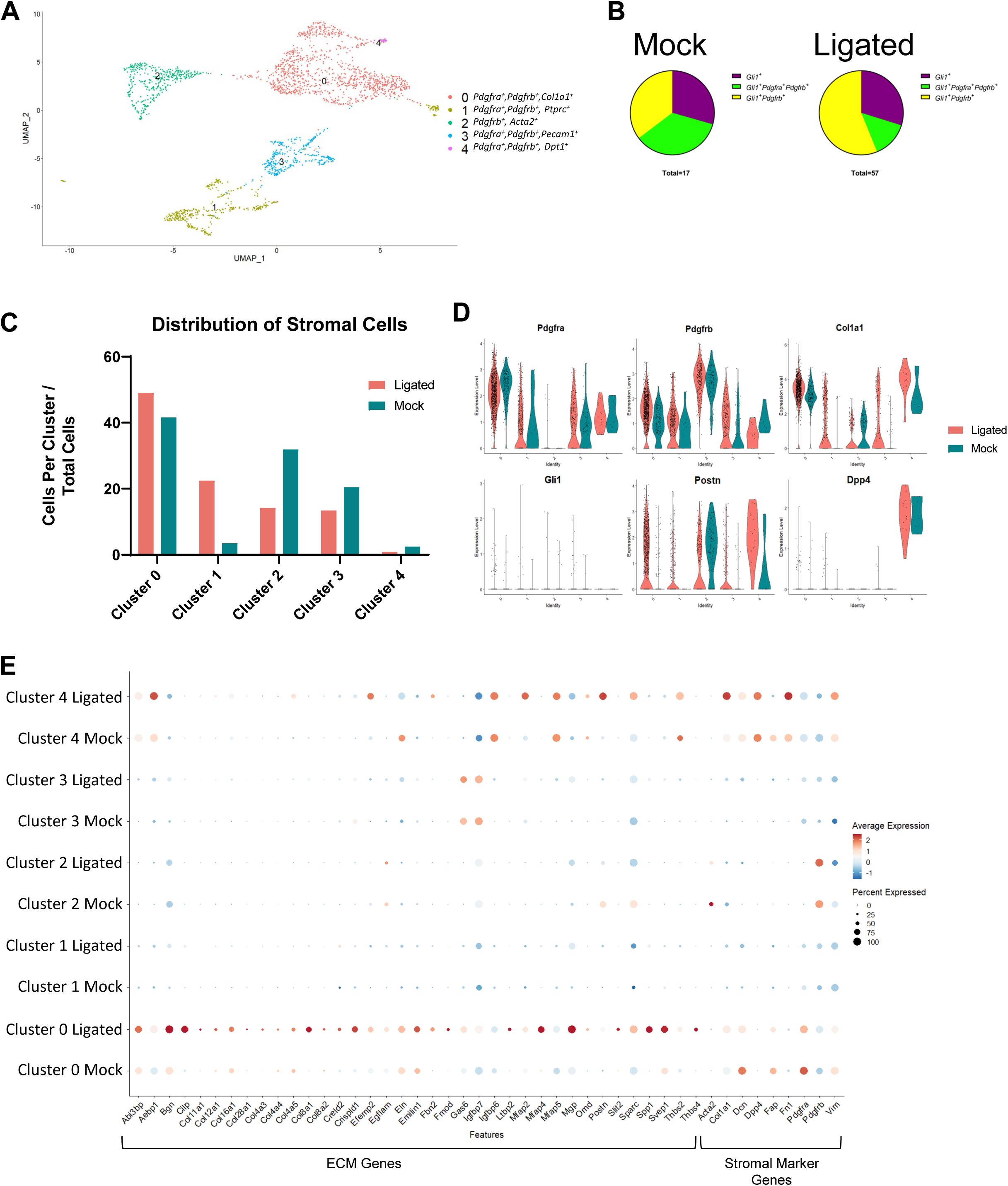
A *Pdgfra* expressing subset of stromal cells shows enrichment of ECM and ECM-associated genes after ductal ligation injury. (**A**) UMAP from stromal cells subsetted based on expression of the stromal cell identity markers *Pdgfra, Pdgfrb, and/or Gli1*. (**B**) Pie charts showing the proportion of *Gli1^+^* cells that express *Gli1* only, *Gli1* and *Pdgfra* only, or *Gli1*, *Pdgfra* and *Pdgfrb* in 14-day mock (Mock) and 14-day ligated (Ligated) glands. (**C**) Percent distribution of stromal cells was calculated by dividing the number of cells in each cluster by the total number of cells in their sample of origin and multiplying by 100. (**D**) Violin plots showing expression levels of *Pdgfra*, *Pdgfrb*, *Col1a1*, *Gli1, Postn, and Dpp4* in Mock and Ligated glands. (**E**) Dot plot showing expression levels and percent of cells expressing ECM-associated genes that were differentially expressed in cluster 0 post-ligation.

Since Gli1-expressing cells appear to have only a minor contribution to ductal ligation-induced salivary gland fibrotic responses and do not display increased expression of fibrillar collagens or proteoglycans after ligation, we wanted to determine which cell population may be responsible for the extracellular matrix deposition. Using a matrisome genelist (Naba et al., 2012), a heat map was generated to display significantly increased genes in clusters based on sample origin and we found that cells in cluster 3 highly express many genes in the matrisome categories of collagens, proteoglycans, and glycoproteins, suggesting they directly contributing to the fibrotic injury (**Figure S6D**). Interestingly, cluster 3 is a *Pdgfra^+^* and *Pdgfrb^+^* stromal cell population (**Figure S6C**), suggesting that the stromal fibroblast may be responsible for ECM production.

As there is known heterogeneity among stromal fibroblast cells (Deng et al., 2021; Zou et al., 2021) we subsetted the stromal populations based on expression of *Pdgfra*, *Pdgfrb,* or *Gli1* to create a new stromal only dataset, which consists of 5 subpopulations (**Figure 6A**). To examine any changes in abundance of cell subpopulations following ligation injury, we calculated the percent of cells per sample. Clusters 0 and 1 displayed increased percentages of cells in ligated glands compared to mock glands, while clusters 2, 3, and 4 had a higher percentage of cells in mock glands compared to ligated glands (**Figure 6C**). Known stromal genes and significantly differentially expressed marker genes were used to identify stromal subpopulations (**Table S3)**. While many of the clusters expressed *Pdgfra* and *Pdgfrb,* Cluster 0 expressed the highest levels of *Pdgfra* and cluster 2, the highest levels of *Pdgfrb.* Cluster 0 showed high levels of expression of *Col1a1 and Postn*, while cluster 4 showed increased expression of *Postn* and *Dpp4*, which are known to contribute to fibrosis (**Figure 6D**) (Chen et al., 2020; Deng et al., 2021). To identify which ECM genes were significantly altered (q score <0.05) following ligation injury we used the “FindAllMarkers” command in Seurat to compare differential gene expression tables with the matrisome gene list. We found the highest expression of ECM genes and the largest changes in gene expression in cluster 0 (**Table S3 and S4).** We then used the “dotplot” command in Seurat to examine the differentially expressed matrisome genes from cluster 0 that were increased in 14 day-ligated cells relative to mock control along with stromal marker genes (**Figure 6E**). The difference in matrisome gene expression levels suggests that cluster 0 is the stromal cell subset that contributes most significantly to the ductal ligation-induced fibrotic response.

## 4 Discussion

Here we report that ligation of the salivary gland ducts produces a progressive salivary gland fibrotic response accompanied by dynamic ECM remodeling up to 14 days after injury. We investigated whether Gli1 signaling contributes to this ductal ligation-induced fibrotic response. Contrary to prior reports that implicated Gli1 signaling in other fibrosis models (Cassandras et al., 2020; Kramann et al., 2015b; Schneider et al., 2017b), knockout of Gli1 had little effect on the fibrotic response following ductal ligation, although there was a downward trend in ECM deposition and active collagen remodeling following Gli1 knockout, suggesting a minor contribution for Gli1 signaling. Lineage tracing of cells that expressed Gli1 at the time of injury showed a small increase in the number of Gli1-derived cells, and a fraction of these cells showed increased co-expression of vimentin and PDGFRβ. Given the extensive fibrotic response, the very small numbers of Gli1^+^ cells, and the inconsistent transcriptional responses of these cells, it’s unlikely that the Gli1^+^ cells are significant contributors to this fibrotic response. We also examined cells expressing PDGFRα and/or β, as these cells have previously been identified as fibrogenic (Ajay et al., 2022; Buhl et al., 2020; Kikuchi et al., 2020; Kocabayoglu et al., 2015; R. Li et al., 2018; Santini et al., 2020; Yao et al., 2022). In wild type glands, there was an increase in PDGFRα^+^ stromal cells after injury. Single-cell RNA-sequencing of stromal cells obtained from glands following 14-day mock surgery or 14 days ligation surgery revealed a subset of *Pdgfra/Pdgfrb* cells, representing the largest stromal cell cohort, which show increased expression and an increased diversity of matrisomal-associated gene expression. This dynamic ECM remodeling was accompanied by a more than a 10-fold increase in macrophages by tissue area. Together our results are consistent with the hypothesis that a PDGFRα/PDGFRβ stomal cell subset is the major fibrogenic cell type and suggest that macrophages contribute to the dynamic ECM remodeling that occurs in this reversible fibrotic injury model.

Our results imply that Gli1 signaling and Gli1 cells have only a minor contribution to salivary gland ductal ligation-induced fibrosis. In our model, while their expansion with injury is 4.6-fold, the number of Gli1^+^ cells is very small, and we found little evidence of a myofibroblast conversion. While *Acta2*/SMA has historically been used as a myofibroblast marker, and SMG stromal cells can undergo a myofibroblast conversion in organoid culture and express SMA (Moskwa et al., 2022), recent studies employing scRNA-seq have defined myofibroblasts based primarily on ECM mRNA expression (Kuppe et al., 2021). scRNA-seq showed no consistent increase in expression of matrisome genes by Gli1 lineage-traced cells. Few Gli1^+^ cells expressed *Acta2*, the gene encoding SMA, and little to no Gli1-derived cells co-expressed SMA at the protein level, nor did SMA protein itself increase after ligation. The increased expression of vimentin, which can be expressed by myofibroblasts, and increased PDGFRβ in a subset of Gli1 lineage-traced cells, suggests that some Gli1- expressing cells may undergo a partial myofibroblast conversion. Although our data suggest that Gli1 signaling is not required for ductal ligation-induced fibrosis, there may be other contributions of Gli1-expressing cells to the fibrotic response in this model, which remains to be investigated. Alternatively, Gli1-expressing cells or Gli1 signaling may contribute to regenerative responses, which can be investigated in this model following removal of the clip. Targeted expression of Shh in keratin 5^+^ (*Krt5*^+^) ductal epithelium protected male mice from hyposalivation (Hai et al., 2014). Additionally, gene delivery of Shh into mice after irradiation prevented damage to microvasculature and parasympathetic innervation (Hai et al., 2016), and its delivery to irradiated minipigs was reported to restore salivary function (Hu et al., 2018; Zhao et al., 2020). As *Gli1^+^* cells in the mouse incisor are progenitor cells that populate the dental mesenchyme in homeostasis and after injury (Zhao et al., 2014b), and *Gli1^+^* cells regenerate the pancreas following ligation (Mathew et al., 2014), Gli1^+^ cells may contribute to salivary gland regeneration after SMG ductal ligation.

PDGFRs were previously shown to have an important function in salivary gland development in regulation of branching morphogenesis upstream of fibroblast growth factor (Yamamoto et al., 2008), and we previously showed that PDGFRα-expressing cells support salivary gland proacinar organoid differentiation (Moskwa et al., 2022). However, there is little understood regarding the contribution of PDGFRα-expressing cells in adult salivary gland. Our finding that *Pdgfra/Pdgfrb*-expressing cells express ECM genes in ligated salivary glands is consistent with scRNA-seq studies of kidney fibrosis that identified similar populations of cells producing ECM (Kuppe et al., 2021). The fact that these cells do not express *Acta2* but do express *Col1a1* and other matrisomal genes is also consistent with recent studies indicating that fibroblasts can transition through a pre-activated proto-myofibroblast state before reaching a functional myofibroblast state (Younesi et al., 2021). As the ductal ligation model is reversible, the fibroblasts may not fully differentiate into myofibroblasts. The *Pdgfra/Pdgfrb* subsets might also be referred to as fibrogenic cells on the basis of their expression of matrisomal genes at the mRNA level (Sun et al., 2016). Our data shows that subsets of stromal cells expressing *Pdgfra* and *Pdgfrb* expand and express an increased diversity of matrisome genes after ligation injury. Our data suggest that the *Pdgfra/Pdgfrb*-expressing cells are important fibrogenic cells in salivary gland ductal ligation-induced fibrosis. Their contribution to other injury and disease states involving fibrosis will be important to examine as well as their contribution to regenerative responses.

Examining the differentially expressed genes in *Pdgfra* and *Pdgfrb*-expressing cells by scRNA-seq in 14 day-ligated glands compared to mock revealed many genes with known associations with fibrosis or disease, including periostin (*Postn*), secreted phosphoprotein 1 (*Spp1*), and latent TGF-β binding protein-2 (*Ltbp2*). *Postn* is an ECM protein that has been shown to be upregulated in pulmonary fibrosis, fibroblasts in keloids, and patients with liver cirrhosis (Chen et al., 2020; Deng et al., 2021; Du et al., 2022). Upregulation of *Spp1* has been shown to increase atrial fibrosis by activating the TGFβ signaling pathway (Du et al., 2022). *Ltbp2* is involved in the TGFβ signaling pathway as it is required for the formation of the large latent complex (LLC) which binds latent TGFβ to the ECM (Munger et al., 1997) and is secreted by myofibroblasts in pulmonary fibrosis (Enomoto et al., 2018). TGFβ is a master regulator of fibrosis, and inhibition of TGFβ1 signaling in ductal ligation-induced fibrosis of the mouse SMG 7 days after injury significantly reduced the fibrotic response (Woods et al., 2015b). Macrophages are known to produce TGFβ in response to injury (Meng et al., 2016), and the macrophages may be a source of TGFβ in this model. Consistent with this, we previously reported on a persistent fibrotic injury following partial gland resection, in which TGFβ1 levels and numbers of M2-like macrophages were significantly increased (O’Keefe et al., 2020). Elevated levels of TGFβ signaling are also detected in patients with Sjögren’s syndrome (Sisto et al., 2018), which is also associated with fibrosis in some patients (Bookman et al., 2011; Leehan et al., 2018; Llamas-Gutierrez et al., 2014). PDGFR^+^ cells are known to respond to TGFβ signaling and have been implicated in fibrosis in the kidney, bone marrow, liver, heart and the lung (Klinkhammer et al., 2018). Receptor tyrosine kinase inhibitors affecting PDGFRs have shown promise in reducing fibrosis in the bone marrow, liver and lung (Klinkhammer et al., 2018). Future studies focusing on PDGFR-expressing cells and PDGFR signaling could define contributions of these stromal subsets and PDGFR signaling in driving fibrosis and in regenerative responses.

## 6 Author Contributions

A.L.A., K.J.O., V.A.G., J.R.T., S.M.F., D.A.N. and M.L. designed experiments, analyzed data, assembled figures, and wrote and revised the manuscript. A.L.A, K.J.O., V.A.G., J.R.T., S.M.F., and N.L.M. performed experiments, analyzed the data, and revised the manuscript. C.V.C. and J.C.T. performed experiments and revised the manuscript.

## 8 Acknowledgments

The authors thank Steve Higgins and Paul Higgins from Albany Medical College for imaging and allowing use of the Nanozoomer. The authors would also like to thank Dr. Paolo Forni for use of his Leica DM 4000 B LED microscope system. The authors are grateful to Dr. Catherine Ovitt, Antigone McKenna, DVM and Timothy Quinn for helpful suggestions, insight, and assistance in implementing this surgical model and Dr. Alexandra Joyner for providing the Gli1^CreERT2^ and Gli1^lz^ founder mice.

## 9 Data Availability Statement

The raw datasets generated for this study can be found in GEO under SuperSeries GSE226640. Scripts used to analyze the data can be found on GitHub (https://github.com/MLarsenLab).

