## Supplementary figures and tables for "Identifying Fibrogenic Cells Following Salivary Gland Obstructive Injury"

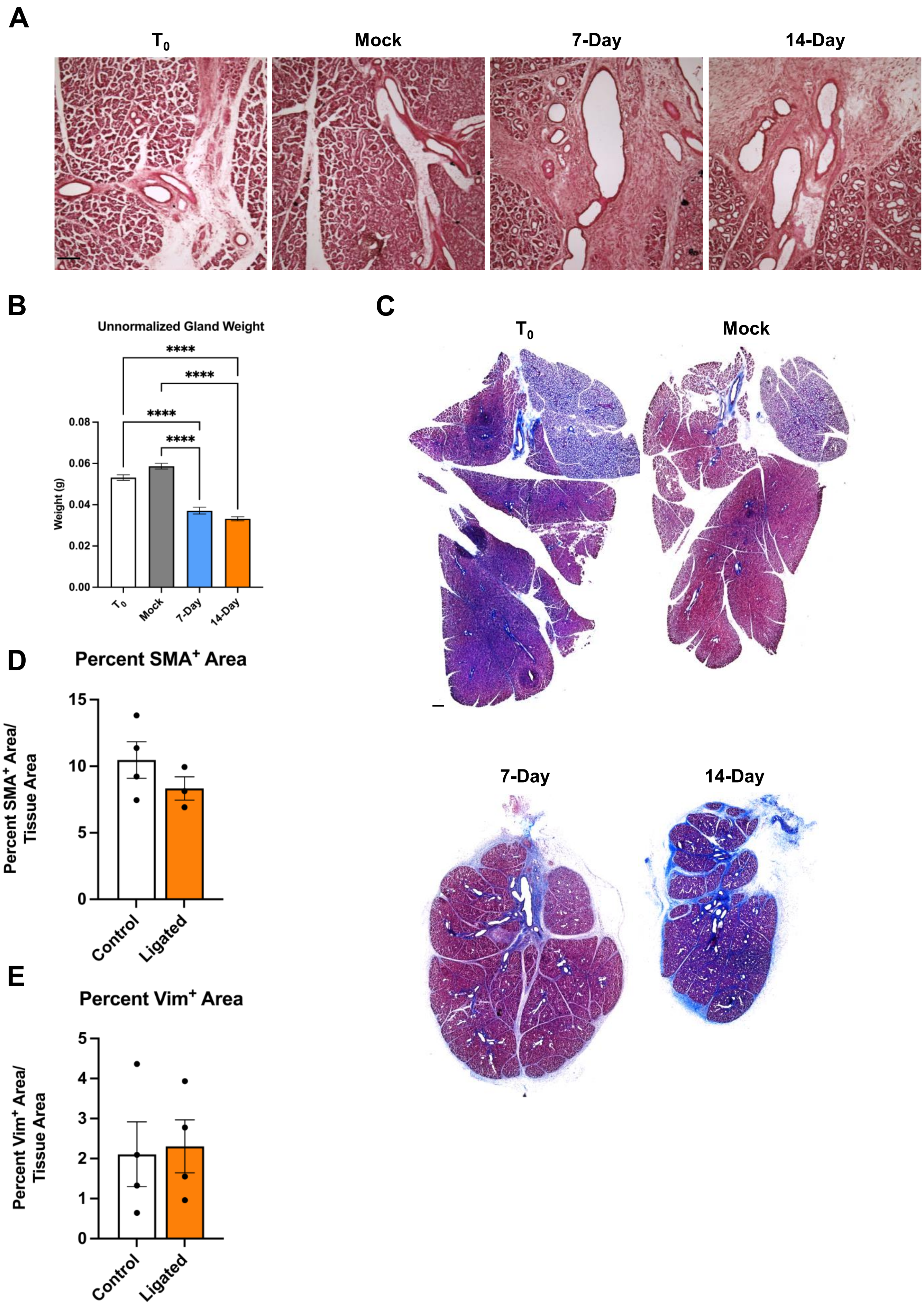
A
T0
Mock
7-Day
14-Day
B
Unnormalized Gland Weight
****
****
****
****
0.08
0.06
0.04
0.02
0.00
Weight (g)
T0
Mock
7-Day
14-Day
C
T0
Mock
7-Day
14-Day
D
Percent SMA+ Area
15
10
5
0
Percent SMA+ Area/
Tissue Area
Control
Ligated
E
Percent Vim+ Area
5
4
3
2
1
0
Percent Vim+ Area/
Tissue Area
Control
Ligated

**Supplementary Figure 1. The stromal and ECM response to ductal ligation surgery**

(**A**) 10X magnification images of H&E-stained glands from time zero ($T_0$),14-day mock (Mock), 7-day ligated (7-Day), and 14-day ligated (14-Day) mice. Scale bar 100 µm. (**B**) Raw gland weights in $T_0$, Mock, 7-Day, and 14-Day glands. N = 10, 25, 6, 34 respectively. (**C**) Masson's trichrome-stained whole gland images of $T_0$, Mock, 7-Day, and 14-Day mice. Scale bar 250 µm. Quantification of percent stain area normalized to total tissue area in Gli1$^{tm3(cre/ERT2)Alj}$/J;(CAG)ROSA26$^{tdTomato}$ (Gli1; R26tdT) 3-week induced control (Control) and 14-day ligated (Ligated) mice for (**D**) SMA (N = 4 and 3 respectively) and (**E**) vimentin (N = 4). Error bars: S.E.M. Statistical Test: Unpaired two-tailed t-test was performed using GraphPad Prism version 9.4.1. ****p≤ 0.0001.

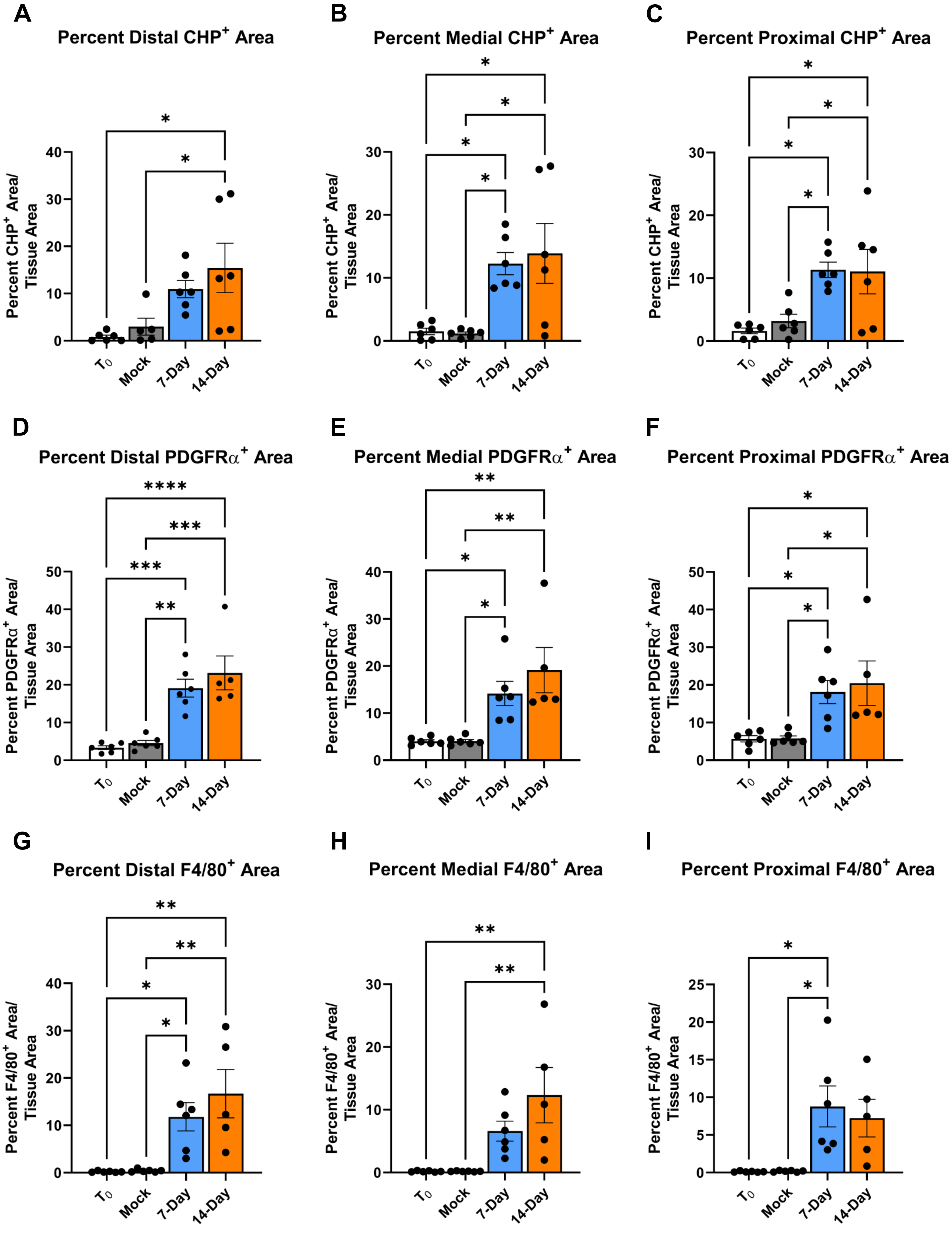

A
Percent Distal CHP+ Area
B
Percent Medial CHP+ Area
C
Percent Proximal CHP+ Area
D
Percent Distal PDGFRα+ Area
E
Percent Medial PDGFRα+ Area
F
Percent Proximal PDGFRα+ Area
G
Percent Distal F4/80+ Area
H
Percent Medial F4/80+ Area
I
Percent Proximal F4/80+ Area
Percent CHP+ Area/ Tissue Area
Percent PDGFRα+ Area/ Tissue Area
Percent F4/80+ Area/ Tissue Area
T0
Mock
7-Day
14-Day

**Supplementary Figure 2. Regional stromal and ECM response to ductal ligation surgery**

Quantification of percent stain area normalized to total tissue area for distal, medial, and proximal regions in the submandibular salivary gland for (**A, B, C**) CHP, (**D, E, F**) PDGFRα, and (**G, H, I**) F4/80. N = 6, 6, 6, and 5 respectively from 2 to 3 images per region. Error bars: S.E.M. Statistics: One-way ANOVA followed by Tukey's multiple comparisons test was performed using GraphPad Prism version 9.4.1. *$p \leq 0.05$, **$p \leq 0.01$, ***$p \leq 0.001$, ****$p \leq 0.0001$.

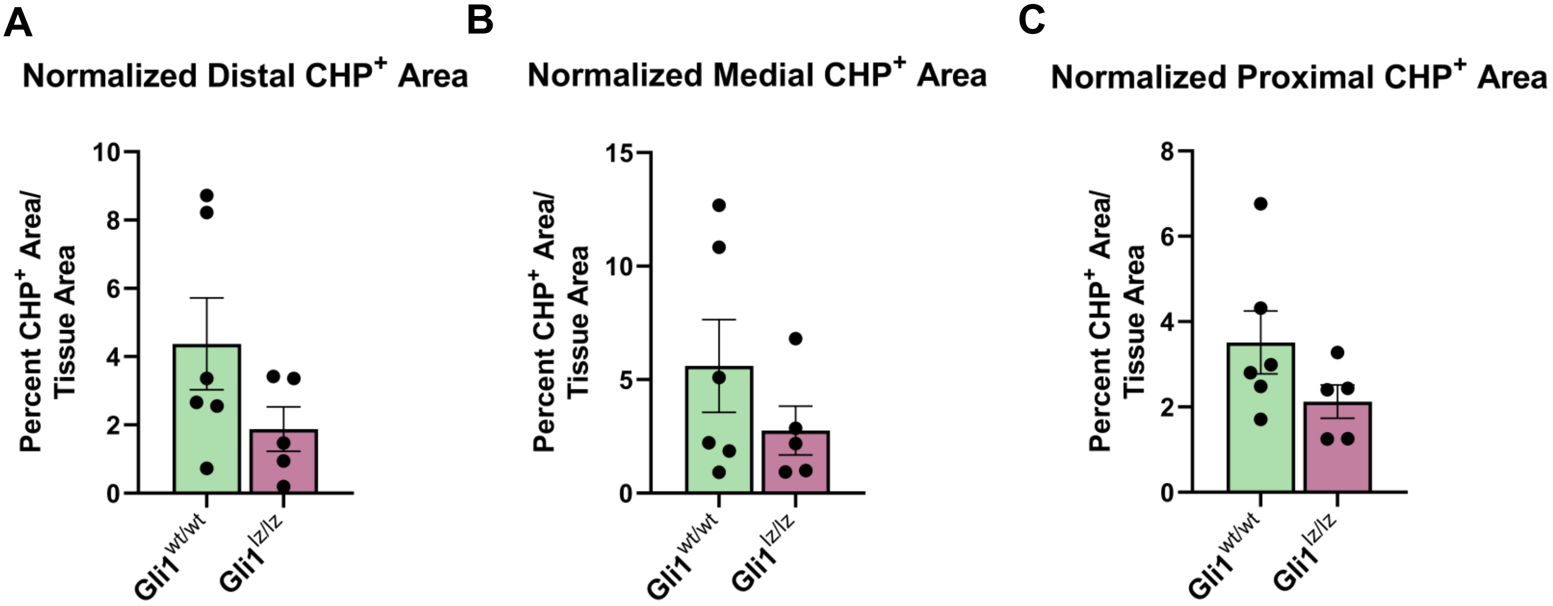
A
Normalized Distal CHP+ Area
Percent CHP+ Area/ Tissue Area
0
2
4
6
8
10
Gli1wt/wt
Gli1lz/lz
B
Normalized Medial CHP+ Area
Percent CHP+ Area/ Tissue Area
0
5
10
15
Gli1wt/wt
Gli1lz/lz
C
Normalized Proximal CHP+ Area
Percent CHP+ Area/ Tissue Area
0
2
4
6
8
Gli1wt/wt
Gli1lz/lz

**Supplementary Figure 3. Regional CHP in 14-day ligated Gli1 null and wildtype mice**

Quantification of percent stain area normalized to total tissue area for (**A**) distal, (**B**) medial, and (**C**) proximal regions in the submandibular salivary gland for CHP in 14-day ligated Gli1 wildtype ($Gli1^{wt/wt}$) and Gli1 null ($Gli1^{lz/lz}$) mice from 1 image per region. Statistical Test: Unpaired two-tailed t-test was performed using GraphPad Prism version 9.4.1. $Gli1^{wt/wt}$ N=6 and $Gli1^{lz/lz}$ N=5.

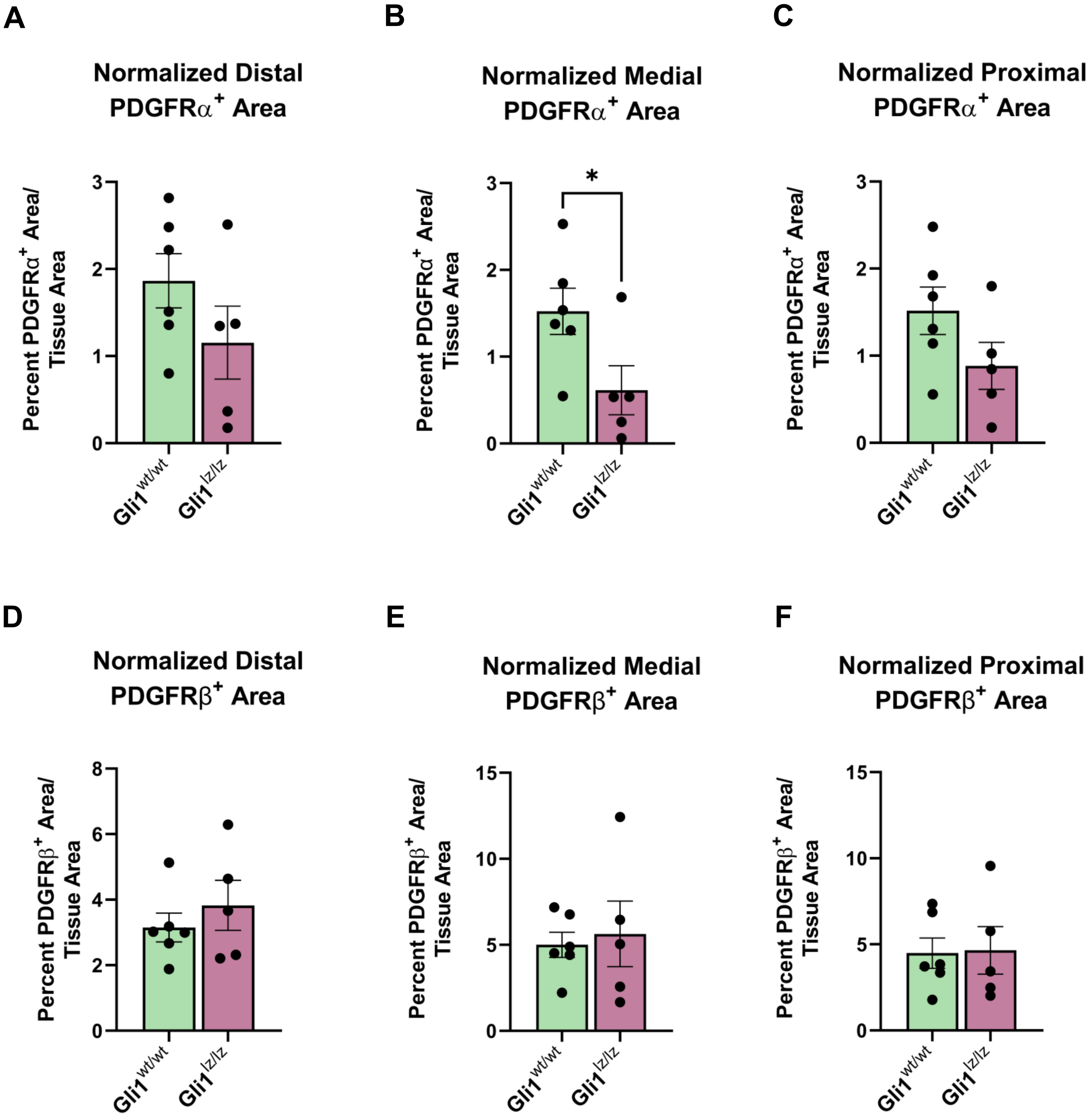

A
Normalized Distal PDGFRα+ Area
Percent PDGFRα+ Area/ Tissue Area
0
1
2
3
Gli1wt/wt
Gli1lz/lz
B
Normalized Medial PDGFRα+ Area
*
C
Normalized Proximal PDGFRα+ Area
D
Normalized Distal PDGFRβ+ Area
Percent PDGFRβ+ Area/ Tissue Area
0
2
4
6
8
E
Normalized Medial PDGFRβ+ Area
0
5
10
15
F
Normalized Proximal PDGFRβ+ Area

**Supplementary Figure 4. Regional changes in PDGFRα and PDGFRβ in 14-day ligated Gli1 null and wildtype mice**

Quantification of percent stain area normalized to total tissue area for distal, medial, and proximal regions in the submandibular salivary gland for (**A, B, C**) PDGFRα and (**D, E, F**) PDGFRβ in 14-day ligated Gli1 wildtype ($Gli1^{wt/wt}$) and Gli1 null ($Gli1^{lz/lz}$) mice from 2-3 images per region. Statistical Test: Unpaired two-tailed t-test was performed using GraphPad Prism version 9.4.1. *$p \leq 0.05$. $Gli1^{wt/wt}$ N=6 and $Gli1^{lz/lz}$ N=5.

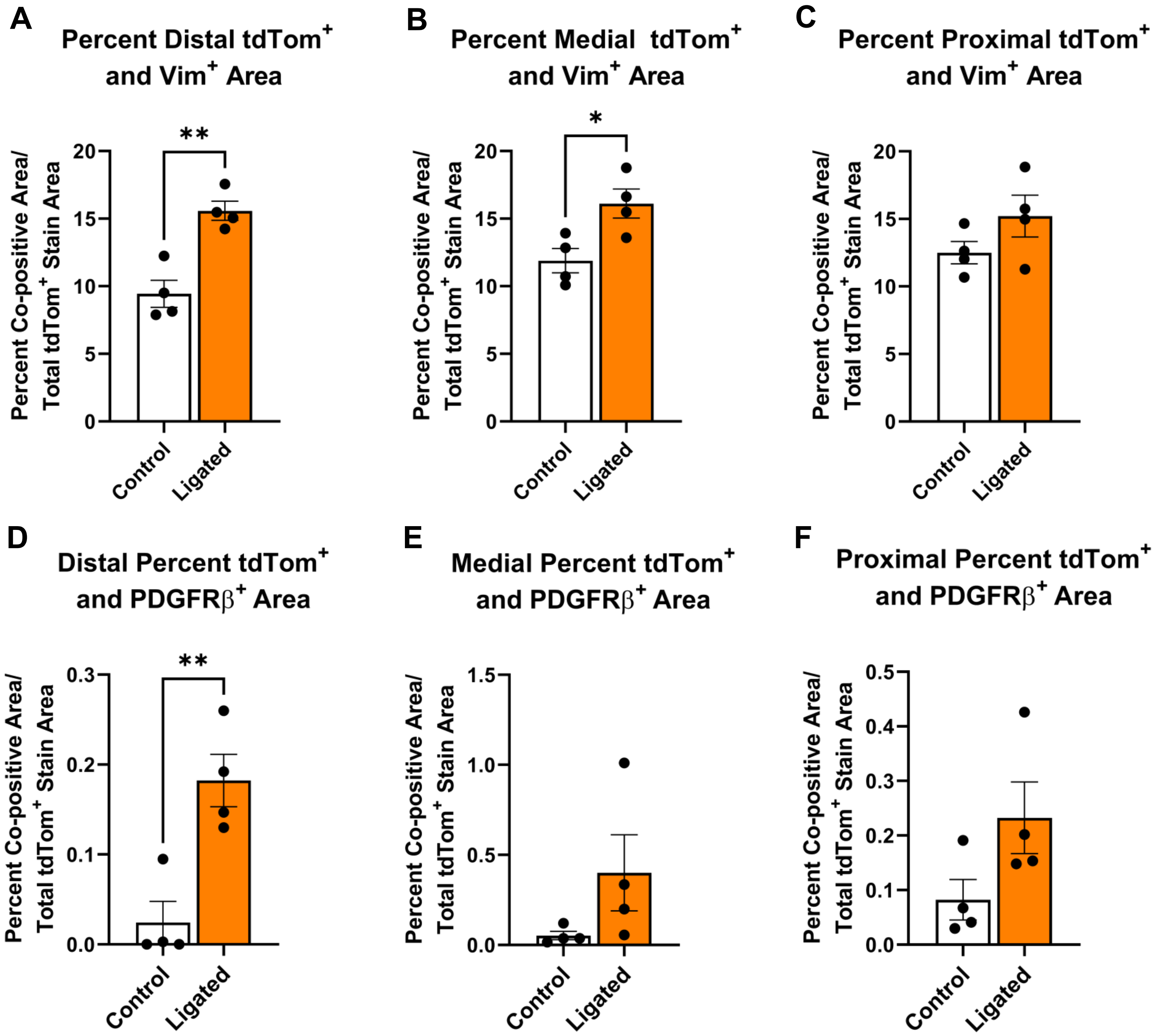
A
Percent Distal tdTom+ and Vim+ Area
B
Percent Medial tdTom+ and Vim+ Area
C
Percent Proximal tdTom+ and Vim+ Area
D
Distal Percent tdTom+ and PDGFRβ+ Area
E
Medial Percent tdTom+ and PDGFRβ+ Area
F
Proximal Percent tdTom+ and PDGFRβ+ Area
Percent Co-positive Area/ Total tdTom+ Stain Area
Control
Ligated
**
*
0
5
10
15
20
0.0
0.1
0.2
0.3
0.5
1.0
1.5
0.4

**Supplementary Figure 5. Gli1-lineage derived cells show regional differences in vimentin and PDGFRβ colocalization**

Quantification of percent colocalized stain area normalized to total Gli1;R26tdT$^+$ stain area for distal, medial, and proximal regions in the submandibular salivary gland for (**A, B, C**) vimentin (Vim), and (**D, E, F**) PDGFRβ in 3-week induced control (Control) or 14-day ligated (Ligated) Gli1;R26tdT mice from 2-3 images per region. Statistical Test: Unpaired two-tailed t-test was performed using GraphPad Prism version 9.4.1. *$p \leq 0.05$ and ** $p \leq 0.01$. N = 4

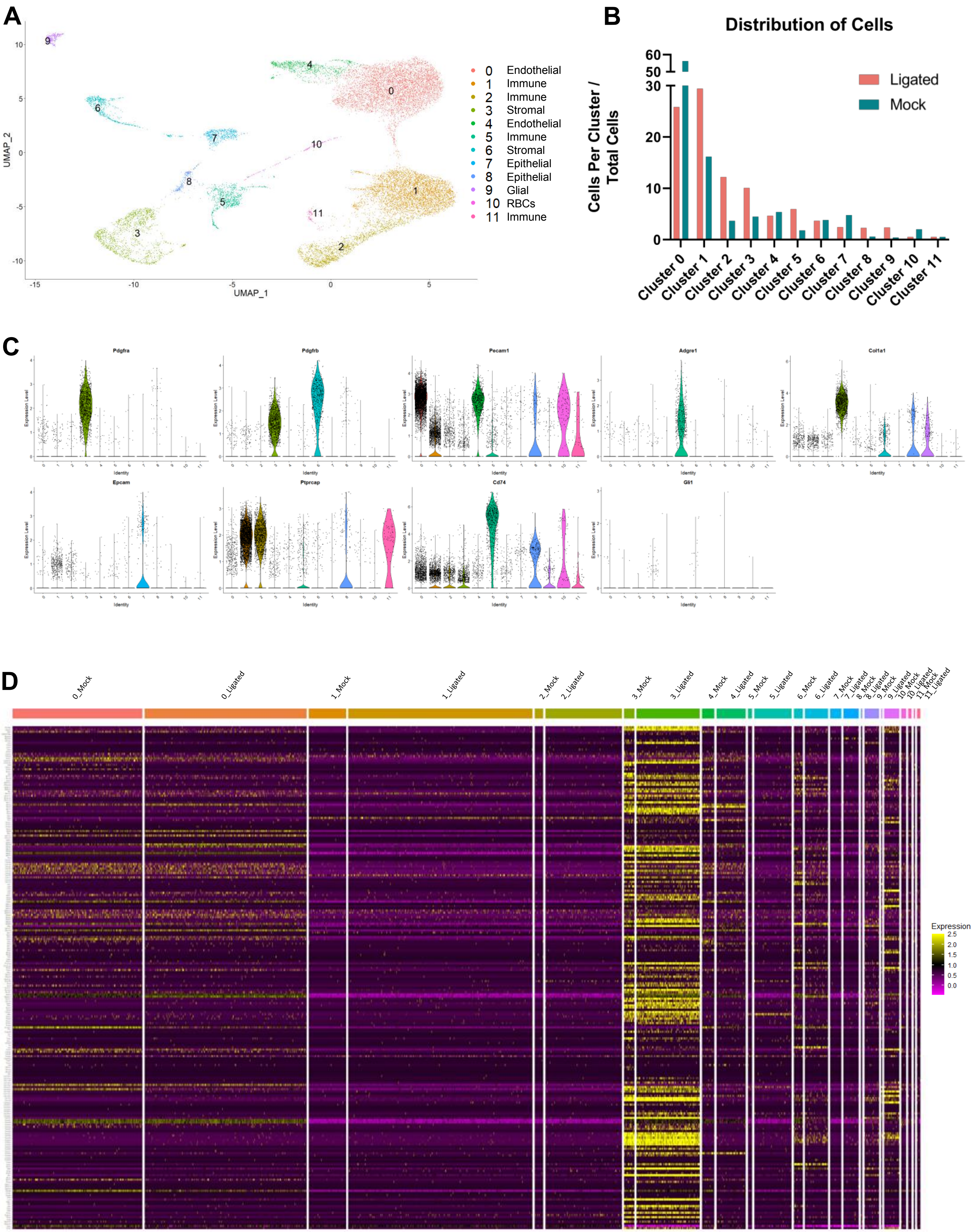
A
UMAP_1
UMAP_2
0 Endothelial
1 Immune
2 Immune
3 Stromal
4 Endothelial
5 Immune
6 Stromal
7 Epithelial
8 Epithelial
9 Glial
10 RBCs
11 Immune
B
Distribution of Cells
Cells Per Cluster / Total Cells
Ligated
Mock
Cluster 0
Cluster 1
Cluster 2
Cluster 3
Cluster 4
Cluster 5
Cluster 6
Cluster 7
Cluster 8
Cluster 9
Cluster 10
Cluster 11
C
Pdgfra
Pdgfrb
Pecam1
Adgre1
Col1a1
Epcam
Ptprcap
Cd74
Gli1
Expression Level
Identity
D
0_Mock
0_Ligated
1_Mock
1_Ligated
2_Mock
2_Ligated
3_Mock
3_Ligated
4_Mock
4_Ligated
5_Mock
5_Ligated
6_Mock
6_Ligated
7_Mock
7_Ligated
8_Mock
8_Ligated
9_Mock
9_Ligated
10_Mock
10_Ligated
11_Mock
11_Ligated
Expression

**Supplemental Figure 6. scRNAseq reveals cluster 3 expresses high levels of stromal identity markers and ECM-associated genes**

(**A**) UMAP of merged 14-day ligated and 14-day mock datasets showing 12 different clusters of cells sequenced. (**B**) Percent distribution of cells was calculated by dividing the number of cells in each cluster by the total number of cells in their sample of origin multiplied by 100. (**C**) Violin plots showing the expression of *Pdgfra, Pdgfrb, Pecam1, Adgre1, Col1a1, Epcam, Ptprcap, Cd74,* and *Gli1* which were used for identification of clusters. (**D**) A heatmap was generated using "matrisome-associated" genes that encode for collagens, proteoglycans and glycoproteins, with cluster 3, the stromal population, expressing the highest levels in both mock and ligated conditions.

**Supplementary Table 1. Antibodies and dilutions used**

Table showing the primary antibodies, direct conjugates, secondary antibodies, and other fluorescent peptides used for staining including the antibody name, the company name, the catalog number, the RRID number and the dilution used for staining, where applicable.

| | Antibody | Company | Catalog Number | RRID Number | Dilution Used |
|---|---|---|---|---|---|
| Primary Antibodies | PDGFRα goat | R & D Systems | AF1062 | AB_2236897 | 1:100 |
| | Living Colors DSRed polyclonal antibody | Takara Biosciences | 632496 | AB_10013483 | 1:1000 |
| | Anti-mouse CD140b (PDGFRb) | Invitrogen | 14-1402-82 | AB_467493 | 1:200 |
| | CD31 (clone MEC13.3) | BD Pharmingen | 553370 | AB_394816 | 1:200 |
| | F4/80 clone BM8 rat | Biolegend | 123101 | AB_893504 | 1:200 |
| | RFP | Rockland | 200-101-379 | AB_2744552 | 1:1000 |
| Direct Conjugates | A488 vimentin | Cell Signaling | 9854 | AB_10829352 | 1:100 |
| | A488 smooth muscle actin (clone 1A4) | Invitrogen | 53-9760-80 | AB_2574460 | 1:200 |
| | Cy3 smooth muscle actin (clone 1A4) | Sigma | C6198 | AB_476856 | 1:600 |
| | Alexa Fluor 647 anti-Tubulin β 3 (TUBB3) | Biolegend | 801210 | AB_2686931 | 1:200 |
| Secondary Antibodies | Anti-rabbit cy 2 | Jackson ImmunoResearch | 711-546-152 | | 1:500 |
| | anti-goat 488 | Jackson ImmunoResearch | 705-546-147 | | 1:500 |
| | Anti-rat no mouse cy 3 | Jackson ImmunoResearch | 712-166-153 | | 1:500 |

| | | | | | |
|---|---|---|---|---|---|
| | Anti-rabbit cy 3 | Jackson ImmunoResearch | 711-166-152 | | 1:500 |
| | Anti-goat cy 5 | Jackson ImmunoResearch | 705-606-147 | | 1:500 |
| | anti-rat no mouse Cy5 | Jackson ImmunoResearch | 712-606-153 | | 1:500 |
| Other | 5-FAM-CHP | Advanced Biomatrix | 5264-60ug | | 40μL of 20 μM stock |

**Supplementary Table 2. Positively upregulated genes in *Gli1*$^+$ cells**

The FindAllMarkers command from the Seurat package in Rstudio was used to examine all differentially expressed genes between 14-day ligated and 14-day mock *Gli1*$^+$ cells.

| | | | | | | | |
|---|---|---|---|---|---|---|---|
| Dirc2.1 | 4.75E-07 | -0.66283 | 0 | 0.412 | 0.009567 | Ligated_Gli+ | Dirc2 |
| Skiv2l2.1 | 4.75E-07 | -0.7214 | 0 | 0.412 | 0.009567 | Ligated_Gli+ | Skiv2l2 |
| Zcchc11.1 | 4.75E-07 | -0.75053 | 0 | 0.412 | 0.009567 | Ligated_Gli+ | Zcchc11 |
| Taf6l.1 | 3.64E-07 | -0.81203 | 0.035 | 0.529 | 0.007321 | Ligated_Gli+ | Taf6l |
| Coro1b.1 | 6.10E-09 | -0.84145 | 0.018 | 0.588 | 0.000123 | Ligated_Gli+ | Coro1b |
| Tmem5.1 | 6.87E-09 | -0.84841 | 0 | 0.529 | 0.000138 | Ligated_Gli+ | Tmem5 |
| 2410015M20Rik.1 | 7.65E-10 | -0.89785 | 0 | 0.588 | 1.54E-05 | Ligated_Gli+ | 2410015M20Rik |
| BC003331.1 | 6.87E-09 | -0.9096 | 0 | 0.529 | 0.000138 | Ligated_Gli+ | BC003331 |
| Fam208a.1 | 5.85E-08 | -0.93118 | 0 | 0.471 | 0.001178 | Ligated_Gli+ | Fam208a |
| Fam96b.1 | 7.65E-10 | -1.11944 | 0 | 0.588 | 1.54E-05 | Ligated_Gli+ | Fam96b |
| 2010111I01Rik.1 | 7.65E-10 | -1.13755 | 0 | 0.588 | 1.54E-05 | Ligated_Gli+ | 2010111I01Rik |
| Trove2.1 | 5.85E-08 | -1.1583 | 0 | 0.471 | 0.001178 | Ligated_Gli+ | Trove2 |
| Zcchc6.1 | 8.08E-11 | -1.20512 | 0 | 0.647 | 1.63E-06 | Ligated_Gli+ | Zcchc6 |
| 1810022K09Rik.1 | 7.59E-13 | -1.21291 | 0 | 0.765 | 1.53E-08 | Ligated_Gli+ | 1810022K09Rik |
| 1700020I14Rik.1 | 6.87E-09 | -1.22248 | 0 | 0.529 | 0.000138 | Ligated_Gli+ | 1700020I14Rik |
| 2010107E04Rik.1 | 7.65E-10 | -1.33406 | 0 | 0.588 | 1.54E-05 | Ligated_Gli+ | 2010107E04Rik |
| 1110008F13Rik.1 | 8.07E-12 | -1.34601 | 0 | 0.706 | 1.62E-07 | Ligated_Gli+ | 1110008F13Rik |
| Usmg5.1 | 8.07E-12 | -1.59894 | 0 | 0.706 | 1.62E-07 | Ligated_Gli+ | Usmg5 |
| Ndufb1-ps | 1.77E-06 | -1.60746 | 0 | 0.737 | 0.035645 | Mock_Gli+ | Ndufb1-ps |
| Atp5mpl | 1.77E-06 | -1.64217 | 0 | 0.737 | 0.035645 | Mock_Gli+ | Atp5mpl |
| Fam198b.1 | 6.87E-09 | -1.64632 | 0 | 0.529 | 0.000138 | Ligated_Gli+ | Fam198b |
| Atp5o.1.1 | 6.73E-14 | -1.65189 | 0 | 0.824 | 1.35E-09 | Ligated_Gli+ | Atp5o.1 |
| Atp5o | 8.62E-08 | -1.66918 | 0 | 0.842 | 0.001737 | Mock_Gli+ | Atp5o |
| Minos1.1 | 8.07E-12 | -1.67088 | 0 | 0.706 | 1.62E-07 | Ligated_Gli+ | Minos1 |
| Aes.1 | 6.87E-09 | -1.67792 | 0 | 0.529 | 0.000138 | Ligated_Gli+ | Aes |
| Grcc10 | 1.92E-06 | -1.7219 | 0.059 | 0.754 | 0.038709 | Mock_Gli+ | Grcc10 |
| Atp5md | 4.12E-07 | -1.78051 | 0 | 0.789 | 0.008302 | Mock_Gli+ | Atp5md |
| Rnasek | 1.47E-07 | -1.78407 | 0 | 0.825 | 0.002962 | Mock_Gli+ | Rnasek |
| mt-Atp8.1 | 3.37E-08 | -1.80531 | 0.614 | 1 | 0.000678 | Ligated_Gli+ | mt-Atp8 |
| Micos10 | 2.48E-07 | -1.90341 | 0 | 0.807 | 0.004989 | Mock_Gli+ | Micos10 |
| Tle5 | 8.62E-08 | -2.21558 | 0 | 0.842 | 0.001737 | Mock_Gli+ | Tle5 |
| Atp6v0c | 2.32E-08 | -2.2503 | 0.059 | 0.895 | 0.000467 | Mock_Gli+ | Atp6v0c |
| Gas5 | 8.62E-08 | -2.43705 | 0 | 0.842 | 0.001737 | Mock_Gli+ | Gas5 |
| Rpl13a | 2.44E-08 | -2.55719 | 0.588 | 0.947 | 0.000492 | Mock_Gli+ | Rpl13a |
| AY036118.1 | 8.09E-07 | -2.95973 | 0.561 | 0.941 | 0.016296 | Ligated_Gli+ | AY036118 |
| Nme2 | 1.15E-08 | -2.96905 | 0.059 | 0.912 | 0.000232 | Mock_Gli+ | Nme2 |

| | p_val | avg_log2FC | pct.1 | pct.2 | p_val_adj | cluster | gene |
|---|---|---|---|---|---|---|---|
| Nme2.1 | 1.15E-08 | 2.969054 | 0.912 | 0.059 | 0.000232 | Ligated_Gli+ | Nme2 |
| AY036118 | 8.09E-07 | 2.959726 | 0.941 | 0.561 | 0.016296 | Mock_Gli+ | AY036118 |
| Rpl13a.1 | 2.44E-08 | 2.557193 | 0.947 | 0.588 | 0.000492 | Ligated_Gli+ | Rpl13a |
| Gas5.1 | 8.62E-08 | 2.437051 | 0.842 | 0 | 0.001737 | Ligated_Gli+ | Gas5 |
| Atp6v0c.1 | 2.32E-08 | 2.250302 | 0.895 | 0.059 | 0.000467 | Ligated_Gli+ | Atp6v0c |
| Tle5.1 | 8.62E-08 | 2.215576 | 0.842 | 0 | 0.001737 | Ligated_Gli+ | Tle5 |
| Micos10.1 | 2.48E-07 | 1.90341 | 0.807 | 0 | 0.004989 | Ligated_Gli+ | Micos10 |
| mt-Atp8 | 3.37E-08 | 1.805314 | 1 | 0.614 | 0.000678 | Mock_Gli+ | mt-Atp8 |
| Rnasek.1 | 1.47E-07 | 1.784067 | 0.825 | 0 | 0.002962 | Ligated_Gli+ | Rnasek |
| Atp5md.1 | 4.12E-07 | 1.78051 | 0.789 | 0 | 0.008302 | Ligated_Gli+ | Atp5md |
| Grcc10.1 | 1.92E-06 | 1.721904 | 0.754 | 0.059 | 0.038709 | Ligated_Gli+ | Grcc10 |
| Aes | 6.87E-09 | 1.677924 | 0.529 | 0 | 0.000138 | Mock_Gli+ | Aes |
| Minos1 | 8.07E-12 | 1.670877 | 0.706 | 0 | 1.62E-07 | Mock_Gli+ | Minos1 |
| Atp5o.2 | 8.62E-08 | 1.669184 | 0.842 | 0 | 0.001737 | Ligated_Gli+ | Atp5o |
| Atp5o.1 | 6.73E-14 | 1.651892 | 0.824 | 0 | 1.35E-09 | Mock_Gli+ | Atp5o.1 |
| Fam198b | 6.87E-09 | 1.646321 | 0.529 | 0 | 0.000138 | Mock_Gli+ | Fam198b |
| Atp5mpl.1 | 1.77E-06 | 1.642173 | 0.737 | 0 | 0.035645 | Ligated_Gli+ | Atp5mpl |
| Ndufb1-ps.1 | 1.77E-06 | 1.607464 | 0.737 | 0 | 0.035645 | Ligated_Gli+ | Ndufb1-ps |
| Usmg5 | 8.07E-12 | 1.598941 | 0.706 | 0 | 1.62E-07 | Mock_Gli+ | Usmg5 |
| 1110008F13Rik | 8.07E-12 | 1.346015 | 0.706 | 0 | 1.62E-07 | Mock_Gli+ | 1110008F13Rik |
| 2010107E04Rik | 7.65E-10 | 1.334062 | 0.588 | 0 | 1.54E-05 | Mock_Gli+ | 2010107E04Rik |
| 1700020I14Rik | 6.87E-09 | 1.222484 | 0.529 | 0 | 0.000138 | Mock_Gli+ | 1700020I14Rik |
| 1810022K09Rik | 7.59E-13 | 1.212908 | 0.765 | 0 | 1.53E-08 | Mock_Gli+ | 1810022K09Rik |
| Zcchc6 | 8.08E-11 | 1.205116 | 0.647 | 0 | 1.63E-06 | Mock_Gli+ | Zcchc6 |
| Trove2 | 5.85E-08 | 1.158301 | 0.471 | 0 | 0.001178 | Mock_Gli+ | Trove2 |
| 2010111I01Rik | 7.65E-10 | 1.137548 | 0.588 | 0 | 1.54E-05 | Mock_Gli+ | 2010111I01Rik |
| Fam96b | 7.65E-10 | 1.119438 | 0.588 | 0 | 1.54E-05 | Mock_Gli+ | Fam96b |
| Fam208a | 5.85E-08 | 0.931175 | 0.471 | 0 | 0.001178 | Mock_Gli+ | Fam208a |
| BC003331 | 6.87E-09 | 0.909603 | 0.529 | 0 | 0.000138 | Mock_Gli+ | BC003331 |
| 2410015M20Rik | 7.65E-10 | 0.89785 | 0.588 | 0 | 1.54E-05 | Mock_Gli+ | 2410015M20Rik |
| Tmem5 | 6.87E-09 | 0.848406 | 0.529 | 0 | 0.000138 | Mock_Gli+ | Tmem5 |
| Coro1b | 6.10E-09 | 0.841451 | 0.588 | 0.018 | 0.000123 | Mock_Gli+ | Coro1b |
| Taf6l | 3.64E-07 | 0.812025 | 0.529 | 0.035 | 0.007321 | Mock_Gli+ | Taf6l |
| Zcchc11 | 4.75E-07 | 0.750525 | 0.412 | 0 | 0.009567 | Mock_Gli+ | Zcchc11 |
| Skiv2l2 | 4.75E-07 | 0.721403 | 0.412 | 0 | 0.009567 | Mock_Gli+ | Skiv2l2 |
| Dirc2 | 4.75E-07 | 0.66283 | 0.412 | 0 | 0.009567 | Mock_Gli+ | Dirc2 |
| 0610037L13Rik | 4.75E-07 | 0.634829 | 0.412 | 0 | 0.009567 | Mock_Gli+ | 0610037L13Rik |
| Fbxo18 | 4.75E-07 | 0.54309 | 0.412 | 0 | 0.009567 | Mock_Gli+ | Fbxo18 |
| Fbxo18.1 | 4.75E-07 | -0.54309 | 0 | 0.412 | 0.009567 | Ligated_Gli+ | Fbxo18 |
| 0610037L13Rik.1 | 4.75E-07 | -0.63483 | 0 | 0.412 | 0.009567 | Ligated_Gli+ | 0610037L13Rik |

**Supplementary Table 3. Positively upregulated genes in stromal subsetted cells**

The FindAllMarkers command from the Seurat package in Rstudio was used to determine the positively upregulated genes in the stromal subsetted cells expressing *Pdgfra, Pdgfrb,* or *Gli1*. Clusters are divided into 14-day ligated cells or mock ligated cells denoted by the “L” or “M” respectively, followed by their cluster number.

| | p_val | avg_log2FC | pct.1 | pct.2 | p_val_adj | cluster | gene |
|---|---|---|---|---|---|---|---|
| Mucl2 | 7.38E-08 | 0.622332 | 0.74 | 0.738 | 0.001487 | 0 L1 _ | Mucl2 |
| Iigp11 | 1.27E-08 | 0.452436 | 0.509 | 0.368 | 0.000257 | 0 L1 _ | Iigp1 |
| Sh3glb11 | 1.12E-09 | 0.252942 | 0.878 | 0.77 | 2.26E-05 | 0 L1 _ | Sh3glb1 |
| Hlf | 1.56E-12 | 0.255726 | 0.25 | 0.136 | 3.15E-08 | 0 L1 _ | Hlf |
| Smdt1 | 6.22E-13 | 0.255473 | 0.883 | 0.748 | 1.25E-08 | 0 L1 _ | Smdt1 |
| F830016B08Rik | 7.21E-15 | 0.261458 | 0.164 | 0.063 | 1.45E-10 | 0 L1 _ | F830016B08Rik |
| Pid1 | 6.66E-15 | 0.253918 | 0.423 | 0.257 | 1.34E-10 | 0 L1 _ | Pid1 |
| Abca11 | 4.25E-15 | 0.273139 | 0.467 | 0.304 | 8.55E-11 | 0 L1 _ | Abca1 |
| Hcfc1r1 | 1.86E-15 | 0.254016 | 0.738 | 0.559 | 3.74E-11 | 0 L1 _ | Hcfc1r1 |
| Ier3ip11 | 1.60E-15 | 0.265152 | 0.761 | 0.592 | 3.23E-11 | 0 L1 _ | Ier3ip1 |
| Imp3 | 2.46E-16 | 0.255102 | 0.796 | 0.606 | 4.95E-12 | 0 L1 _ | Imp3 |
| Tm9sf31 | 2.26E-16 | 0.25314 | 0.847 | 0.685 | 4.55E-12 | 0 L1 _ | Tm9sf3 |
| Col4a4 | 2.32E-17 | 0.254517 | 0.249 | 0.118 | 4.67E-13 | 0 L1 _ | Col4a4 |
| Arcn11 | 1.23E-17 | 0.258555 | 0.647 | 0.448 | 2.47E-13 | 0 L1 _ | Arcn1 |
| Ddb1 | 1.18E-17 | 0.265922 | 0.623 | 0.427 | 2.38E-13 | 0 L1 _ | Ddb1 |
| Tmem64 | 7.30E-18 | 0.271098 | 0.332 | 0.182 | 1.47E-13 | 0 L1 _ | Tmem64 |
| Adh11 | 6.83E-18 | 0.329914 | 0.252 | 0.116 | 1.38E-13 | 0 L1 _ | Adh1 |
| Serpine2 | 6.28E-18 | 0.345982 | 0.669 | 0.455 | 1.26E-13 | 0 L1 _ | Serpine2 |
| Resf1 | 2.20E-18 | 0.267046 | 0.585 | 0.366 | 4.43E-14 | 0 L1 _ | Resf1 |
| Dnajc1 | 1.84E-18 | 0.275029 | 0.737 | 0.528 | 3.71E-14 | 0 L1 _ | Dnajc1 |
| Nrip1 | 1.54E-18 | 0.292644 | 0.733 | 0.532 | 3.11E-14 | 0 L1 _ | Nrip1 |
| Grina1 | 1.50E-18 | 0.256986 | 0.71 | 0.472 | 3.01E-14 | 0 L1 _ | Grina |
| Sec61b1 | 1.03E-18 | 0.29495 | 0.902 | 0.723 | 2.08E-14 | 0 L1 _ | Sec61b |
| Adamts151 | 7.24E-19 | 0.312005 | 0.342 | 0.182 | 1.46E-14 | 0 L1 _ | Adamts15 |
| C1d | 6.90E-19 | 0.259023 | 0.582 | 0.389 | 1.39E-14 | 0 L1 _ | C1d |
| Uap1l1 | 6.18E-19 | 0.251401 | 0.347 | 0.189 | 1.25E-14 | 0 L1 _ | Uap1l1 |
| Itpr11 | 5.44E-19 | 0.253808 | 0.602 | 0.395 | 1.09E-14 | 0 L1 _ | Itpr1 |
| Dnajb11 | 5.01E-19 | 0.270581 | 0.631 | 0.441 | 1.01E-14 | 0 L1 _ | Dnajb11 |
| Aprt | 3.41E-19 | 0.253091 | 0.683 | 0.477 | 6.86E-15 | 0 L1 _ | Aprt |
| Acbd3 | 2.97E-19 | 0.251901 | 0.592 | 0.395 | 5.98E-15 | 0 L1 _ | Acbd3 |
| Sfr1 | 1.52E-19 | 0.312919 | 0.882 | 0.684 | 3.07E-15 | 0 L1 _ | Sfr1 |
| Ccl81 | 4.30E-20 | 0.398268 | 0.191 | 0.068 | 8.67E-16 | 0 L1 _ | Ccl8 |
| Glg11 | 3.68E-20 | 0.286973 | 0.798 | 0.561 | 7.41E-16 | 0 L1 _ | Glg1 |
| Prrg31 | 3.37E-20 | 0.266923 | 0.247 | 0.109 | 6.79E-16 | 0 L1 _ | Prrg3 |
| Vegfb | 3.00E-20 | 0.272464 | 0.377 | 0.206 | 6.05E-16 | 0 L1 _ | Vegfb |
| Ssh21 | 1.92E-20 | 0.258583 | 0.61 | 0.391 | 3.87E-16 | 0 L1 _ | Ssh2 |
| Irs21 | 1.87E-20 | 0.274087 | 0.297 | 0.142 | 3.78E-16 | 0 L1 _ | Irs2 |
| Svbp | 1.46E-20 | 0.268379 | 0.625 | 0.422 | 2.94E-16 | 0 L1 _ | Svbp |
| 1700025G04Rik | 1.38E-20 | 0.301204 | 0.603 | 0.393 | 2.77E-16 | 0 L1 _ | 1700025G04Rik |
| Trf1 | 1.32E-20 | 0.295855 | 0.372 | 0.195 | 2.66E-16 | 0 L1 _ | Trf |

| | | | | | | | |
|---|---|---|---|---|---|---|---|
| Cebpb1 | 1.06E-20 | 0.311476 | 0.555 | 0.346 | 2.13E-16 | 0 L1 _ | Cebpb |
| Selenok | 8.02E-21 | 0.261249 | 0.929 | 0.782 | 1.62E-16 | 0 L1 _ | Selenok |
| Cdc16 | 7.63E-21 | 0.255957 | 0.399 | 0.221 | 1.54E-16 | 0 L1 _ | Cdc16 |
| Cxxc5 | 5.52E-21 | 0.259985 | 0.469 | 0.281 | 1.11E-16 | 0 L1 _ | Cxxc5 |
| Kdelr11 | 3.77E-21 | 0.289052 | 0.77 | 0.554 | 7.60E-17 | 0 L1 _ | Kdelr1 |
| Runx1t11 | 3.54E-21 | 0.268239 | 0.426 | 0.239 | 7.14E-17 | 0 L1 _ | Runx1t1 |
| Tril1 | 2.50E-21 | 0.262195 | 0.585 | 0.363 | 5.03E-17 | 0 L1 _ | Tril |
| Hmgcs1 | 2.40E-21 | 0.355503 | 0.375 | 0.207 | 4.84E-17 | 0 L1 _ | Hmgcs1 |
| Smad31 | 1.54E-21 | 0.275853 | 0.319 | 0.158 | 3.10E-17 | 0 L1 _ | Smad3 |
| Olfm1 | 1.22E-21 | 0.252538 | 0.444 | 0.251 | 2.47E-17 | 0 L1 _ | Olfm1 |
| Pnp1 | 1.01E-21 | 0.261646 | 0.73 | 0.484 | 2.03E-17 | 0 L1 _ | Pnp |
| Tcf42 | 7.17E-22 | 0.339371 | 0.975 | 0.83 | 1.44E-17 | 0 L1 _ | Tcf4 |
| Fnbp11 | 4.15E-22 | 0.288331 | 0.665 | 0.44 | 8.36E-18 | 0 L1 _ | Fnbp1 |
| Tex264 | 3.72E-22 | 0.251855 | 0.577 | 0.36 | 7.48E-18 | 0 L1 _ | Tex264 |
| Sult5a11 | 3.44E-22 | 0.259445 | 0.243 | 0.099 | 6.92E-18 | 0 L1 _ | Sult5a1 |
| Slc38a101 | 3.04E-22 | 0.282089 | 0.538 | 0.334 | 6.12E-18 | 0 L1 _ | Slc38a10 |
| Yipf3 | 2.91E-22 | 0.257142 | 0.663 | 0.431 | 5.86E-18 | 0 L1 _ | Yipf3 |
| Ufc1 | 2.55E-22 | 0.265587 | 0.712 | 0.482 | 5.14E-18 | 0 L1 _ | Ufc1 |
| Ptn | 2.39E-22 | 0.392363 | 0.174 | 0.051 | 4.80E-18 | 0 L1 _ | Ptn |
| Rnf10 | 2.24E-22 | 0.251477 | 0.671 | 0.443 | 4.51E-18 | 0 L1 _ | Rnf10 |
| Iah1 | 2.03E-22 | 0.278792 | 0.511 | 0.309 | 4.09E-18 | 0 L1 _ | Iah1 |
| Dpy19l11 | 1.94E-22 | 0.28224 | 0.426 | 0.237 | 3.92E-18 | 0 L1 _ | Dpy19l1 |
| Clptm1l | 1.69E-22 | 0.252353 | 0.553 | 0.353 | 3.40E-18 | 0 L1 _ | Clptm1l |
| Sgcb1 | 1.61E-22 | 0.250534 | 0.452 | 0.26 | 3.25E-18 | 0 L1 _ | Sgcb |
| Plod3 | 1.27E-22 | 0.256651 | 0.493 | 0.293 | 2.56E-18 | 0 L1 _ | Plod3 |
| Fign | 1.23E-22 | 0.275755 | 0.262 | 0.11 | 2.48E-18 | 0 L1 _ | Fign |
| Ccn1 | 1.10E-22 | 0.286291 | 0.227 | 0.084 | 2.22E-18 | 0 L1 _ | Ccn1 |
| Gpc4 | 1.02E-22 | 0.291207 | 0.263 | 0.111 | 2.05E-18 | 0 L1 _ | Gpc4 |
| Asah1 | 1.01E-22 | 0.26483 | 0.63 | 0.407 | 2.03E-18 | 0 L1 _ | Asah1 |
| Trim471 | 9.17E-23 | 0.272084 | 0.533 | 0.321 | 1.85E-18 | 0 L1 _ | Trim47 |
| Acvr1 | 8.84E-23 | 0.265286 | 0.303 | 0.141 | 1.78E-18 | 0 L1 _ | Acvr1 |
| Xbp11 | 8.16E-23 | 0.268084 | 0.748 | 0.506 | 1.64E-18 | 0 L1 _ | Xbp1 |
| Ccl19 | 7.04E-23 | 0.382558 | 0.173 | 0.049 | 1.42E-18 | 0 L1 _ | Ccl19 |
| Casp12 | 5.64E-23 | 0.281813 | 0.299 | 0.136 | 1.14E-18 | 0 L1 _ | Casp12 |
| Smarcb1 | 5.45E-23 | 0.259156 | 0.65 | 0.43 | 1.10E-18 | 0 L1 _ | Smarcb1 |
| Dio2 | 5.37E-23 | 0.321242 | 0.204 | 0.069 | 1.08E-18 | 0 L1 _ | Dio2 |
| Ang | 5.11E-23 | 0.266325 | 0.28 | 0.12 | 1.03E-18 | 0 L1 _ | Ang |
| C1qtnf6 | 4.14E-23 | 0.259474 | 0.442 | 0.248 | 8.34E-19 | 0 L1 _ | C1qtnf6 |
| Copg11 | 3.41E-23 | 0.304437 | 0.513 | 0.308 | 6.87E-19 | 0 L1 _ | Copg1 |
| Creld2 | 3.35E-23 | 0.274587 | 0.429 | 0.242 | 6.75E-19 | 0 L1 _ | Creld2 |
| Pcsk5 | 3.34E-23 | 0.325153 | 0.301 | 0.134 | 6.72E-19 | 0 L1 _ | Pcsk5 |

| | | | | | | | |
|---|---|---|---|---|---|---|---|
| Col4a3 | 3.15E-23 | 0.264496 | 0.218 | 0.079 | 6.34E-19 | 0 L1 _ | Col4a3 |
| Fkbp21 | 1.97E-23 | 0.316328 | 0.709 | 0.475 | 3.97E-19 | 0 L1 _ | Fkbp2 |
| Mcrip1 | 1.73E-23 | 0.314454 | 0.721 | 0.51 | 3.48E-19 | 0 L1 _ | Mcrip1 |
| Man2a11 | 1.42E-23 | 0.270258 | 0.715 | 0.458 | 2.86E-19 | 0 L1 _ | Man2a1 |
| Spred1 | 1.38E-23 | 0.281439 | 0.733 | 0.474 | 2.78E-19 | 0 L1 _ | Spred1 |
| Lrp61 | 1.34E-23 | 0.308409 | 0.589 | 0.365 | 2.70E-19 | 0 L1 _ | Lrp6 |
| Loxl21 | 1.33E-23 | 0.265551 | 0.457 | 0.244 | 2.69E-19 | 0 L1 _ | Loxl2 |
| St131 | 1.00E-23 | 0.321324 | 0.868 | 0.66 | 2.02E-19 | 0 L1 _ | St13 |
| Zeb1 | 7.45E-24 | 0.34569 | 0.742 | 0.516 | 1.50E-19 | 0 L1 _ | Zeb1 |
| Copa1 | 6.83E-24 | 0.299162 | 0.756 | 0.526 | 1.38E-19 | 0 L1 _ | Copa |
| Asap11 | 6.19E-24 | 0.316423 | 0.76 | 0.522 | 1.25E-19 | 0 L1 _ | Asap1 |
| Mitf | 4.55E-24 | 0.297219 | 0.302 | 0.136 | 9.16E-20 | 0 L1 _ | Mitf |
| Tspan311 | 3.72E-24 | 0.280628 | 0.594 | 0.368 | 7.50E-20 | 0 L1 _ | Tspan31 |
| Ssc5d | 2.95E-24 | 0.25459 | 0.236 | 0.088 | 5.94E-20 | 0 L1 _ | Ssc5d |
| Mettl9 | 2.49E-24 | 0.259824 | 0.533 | 0.314 | 5.01E-20 | 0 L1 _ | Mettl9 |
| Fdps | 2.28E-24 | 0.439952 | 0.378 | 0.197 | 4.59E-20 | 0 L1 _ | Fdps |
| Serf2 | 2.09E-24 | 0.310459 | 0.972 | 0.932 | 4.20E-20 | 0 L1 _ | Serf2 |
| Atxn10 | 2.07E-24 | 0.254685 | 0.677 | 0.428 | 4.17E-20 | 0 L1 _ | Atxn10 |
| Ctsa | 2.05E-24 | 0.285459 | 0.65 | 0.419 | 4.14E-20 | 0 L1 _ | Ctsa |
| Cdkn2b | 1.22E-24 | 0.251604 | 0.164 | 0.043 | 2.45E-20 | 0 L1 _ | Cdkn2b |
| Lsp11 | 1.02E-24 | 0.334709 | 0.669 | 0.401 | 2.05E-20 | 0 L1 _ | Lsp1 |
| Rhoq1 | 1.00E-24 | 0.277989 | 0.543 | 0.32 | 2.02E-20 | 0 L1 _ | Rhoq |
| Lamc31 | 9.54E-25 | 0.301445 | 0.444 | 0.228 | 1.92E-20 | 0 L1 _ | Lamc3 |
| Maf1 | 8.10E-25 | 0.265071 | 0.693 | 0.426 | 1.63E-20 | 0 L1 _ | Maf |
| Myo101 | 7.37E-25 | 0.269899 | 0.601 | 0.353 | 1.49E-20 | 0 L1 _ | Myo10 |
| Synpo | 7.32E-25 | 0.282973 | 0.472 | 0.26 | 1.47E-20 | 0 L1 _ | Synpo |
| Pink1 | 7.14E-25 | 0.298326 | 0.638 | 0.415 | 1.44E-20 | 0 L1 _ | Pink1 |
| Naglu | 7.01E-25 | 0.276239 | 0.301 | 0.134 | 1.41E-20 | 0 L1 _ | Naglu |
| Rtn41 | 6.64E-25 | 0.325925 | 0.855 | 0.671 | 1.34E-20 | 0 L1 _ | Rtn4 |
| Sox4 | 6.16E-25 | 0.289551 | 0.584 | 0.343 | 1.24E-20 | 0 L1 _ | Sox4 |
| Bet1l | 6.07E-25 | 0.323767 | 0.63 | 0.405 | 1.22E-20 | 0 L1 _ | Bet1l |
| Mme1 | 5.77E-25 | 0.253092 | 0.253 | 0.093 | 1.16E-20 | 0 L1 _ | Mme |
| Bmpr1a1 | 5.52E-25 | 0.292488 | 0.514 | 0.288 | 1.11E-20 | 0 L1 _ | Bmpr1a |
| Pard6g | 5.48E-25 | 0.281867 | 0.346 | 0.161 | 1.10E-20 | 0 L1 _ | Pard6g |
| Gcsh | 4.48E-25 | 0.263694 | 0.345 | 0.161 | 9.03E-21 | 0 L1 _ | Gcsh |
| Nrxn11 | 4.13E-25 | 0.402682 | 0.208 | 0.066 | 8.31E-21 | 0 L1 _ | Nrxn1 |
| Pam1 | 4.02E-25 | 0.259884 | 0.674 | 0.423 | 8.10E-21 | 0 L1 _ | Pam |
| 1110065P20Rik | 3.81E-25 | 0.250709 | 0.466 | 0.255 | 7.67E-21 | 0 L1 _ | 1110065P20Rik |
| Dnm11 | 2.67E-25 | 0.251967 | 0.329 | 0.148 | 5.37E-21 | 0 L1 _ | Dnm1 |
| Smad5 | 2.46E-25 | 0.357792 | 0.488 | 0.278 | 4.96E-21 | 0 L1 _ | Smad5 |
| Cdk8 | 2.15E-25 | 0.357108 | 0.497 | 0.281 | 4.34E-21 | 0 L1 _ | Cdk8 |

| | | | | | | | |
|---|---|---|---|---|---|---|---|
| Siva1 | 1.99E-25 | 0.294877 | 0.555 | 0.332 | 4.02E-21 | 0 L1 _ | Siva1 |
| Rab7b | 1.81E-25 | 0.27483 | 0.298 | 0.129 | 3.64E-21 | 0 L1 _ | Rab7b |
| Ssr4 | 1.74E-25 | 0.36512 | 0.883 | 0.714 | 3.51E-21 | 0 L1 _ | Ssr4 |
| Vmp1 | 1.08E-25 | 0.299458 | 0.565 | 0.34 | 2.18E-21 | 0 L1 _ | Vmp1 |
| App2 | 1.05E-25 | 0.282016 | 0.972 | 0.75 | 2.11E-21 | 0 L1 _ | App |
| Rhod | 9.76E-26 | 0.261997 | 0.269 | 0.103 | 1.97E-21 | 0 L1 _ | Rhod |
| Il15 | 8.54E-26 | 0.277283 | 0.285 | 0.116 | 1.72E-21 | 0 L1 _ | Il15 |
| Gm300751 | 7.93E-26 | 0.327801 | 0.259 | 0.099 | 1.60E-21 | 0 L1 _ | Gm30075 |
| Map1lc3b | 6.89E-26 | 0.309047 | 0.923 | 0.804 | 1.39E-21 | 0 L1 _ | Map1lc3b |
| Sqle1 | 6.48E-26 | 0.315338 | 0.263 | 0.102 | 1.30E-21 | 0 L1 _ | Sqle |
| Lefty1 | 6.25E-26 | 0.250901 | 0.247 | 0.092 | 1.26E-21 | 0 L1 _ | Lefty1 |
| Tslp1 | 5.86E-26 | 0.276434 | 0.238 | 0.085 | 1.18E-21 | 0 L1 _ | Tslp |
| Serpinb6a2 | 4.96E-26 | 0.27088 | 0.861 | 0.613 | 9.98E-22 | 0 L1 _ | Serpinb6a |
| Atp1a21 | 4.19E-26 | 0.301899 | 0.361 | 0.166 | 8.43E-22 | 0 L1 _ | Atp1a2 |
| Smo | 4.18E-26 | 0.272541 | 0.438 | 0.227 | 8.42E-22 | 0 L1 _ | Smo |
| Tnfsf12 | 4.00E-26 | 0.310075 | 0.788 | 0.559 | 8.06E-22 | 0 L1 _ | Tnfsf12 |
| Atp5o1 | 3.55E-26 | 0.279946 | 0.806 | 0.492 | 7.16E-22 | 0 L1 _ | Atp5o |
| Tmem97 | 3.10E-26 | 0.255978 | 0.317 | 0.138 | 6.24E-22 | 0 L1 _ | Tmem97 |
| Tceal9 | 2.74E-26 | 0.288085 | 0.849 | 0.625 | 5.52E-22 | 0 L1 _ | Tceal9 |
| Srpr | 2.64E-26 | 0.347801 | 0.764 | 0.541 | 5.33E-22 | 0 L1 _ | Srpr |
| Susd2 | 2.53E-26 | 0.276367 | 0.249 | 0.089 | 5.09E-22 | 0 L1 _ | Susd2 |
| Tmem9 | 2.41E-26 | 0.252284 | 0.383 | 0.186 | 4.86E-22 | 0 L1 _ | Tmem9 |
| Tmed9 | 2.38E-26 | 0.321306 | 0.886 | 0.707 | 4.80E-22 | 0 L1 _ | Tmed9 |
| Usp9x1 | 2.19E-26 | 0.326513 | 0.673 | 0.434 | 4.40E-22 | 0 L1 _ | Usp9x |
| Creb3l2 | 2.12E-26 | 0.358585 | 0.564 | 0.341 | 4.27E-22 | 0 L1 _ | Creb3l2 |
| Herpud11 | 2.09E-26 | 0.299825 | 0.708 | 0.442 | 4.21E-22 | 0 L1 _ | Herpud1 |
| Tomm6 | 1.98E-26 | 0.260664 | 0.77 | 0.492 | 3.98E-22 | 0 L1 _ | Tomm6 |
| Tmem59 | 1.80E-26 | 0.295274 | 0.935 | 0.834 | 3.62E-22 | 0 L1 _ | Tmem59 |
| Magi2 | 1.70E-26 | 0.258134 | 0.347 | 0.154 | 3.43E-22 | 0 L1 _ | Magi2 |
| Sigmar1 | 1.61E-26 | 0.345736 | 0.464 | 0.258 | 3.25E-22 | 0 L1 _ | Sigmar1 |
| Oasl2 | 1.61E-26 | 0.368031 | 0.476 | 0.257 | 3.24E-22 | 0 L1 _ | Oasl2 |
| Layn1 | 1.04E-26 | 0.295179 | 0.374 | 0.181 | 2.10E-22 | 0 L1 _ | Layn |
| 1500011B03Rik | 9.93E-27 | 0.251454 | 0.321 | 0.138 | 2.00E-22 | 0 L1 _ | 1500011B03Rik |
| Sra1 | 5.76E-27 | 0.335271 | 0.688 | 0.46 | 1.16E-22 | 0 L1 _ | Sra1 |
| Parp3 | 4.87E-27 | 0.302002 | 0.581 | 0.349 | 9.81E-23 | 0 L1 _ | Parp3 |
| Gulp11 | 4.19E-27 | 0.299604 | 0.501 | 0.275 | 8.43E-23 | 0 L1 _ | Gulp1 |
| Spcs1 | 2.65E-27 | 0.318497 | 0.844 | 0.658 | 5.34E-23 | 0 L1 _ | Spcs1 |
| Hist1h2bc1 | 2.62E-27 | 0.330413 | 0.635 | 0.397 | 5.27E-23 | 0 L1 _ | Hist1h2bc |
| Bace2 | 2.37E-27 | 0.281911 | 0.339 | 0.149 | 4.78E-23 | 0 L1 _ | Bace2 |
| Ubtd1 | 2.06E-27 | 0.2694 | 0.325 | 0.138 | 4.16E-23 | 0 L1 _ | Ubtd1 |
| Gadd45b | 1.56E-27 | 0.450121 | 0.462 | 0.245 | 3.14E-23 | 0 L1 _ | Gadd45b |

| | | | | | | | |
|---|---|---|---|---|---|---|---|
| Glmp | 1.39E-27 | 0.291119 | 0.672 | 0.421 | 2.79E-23 | 0 L1 _ | Glmp |
| Rnd31 | 1.31E-27 | 0.380772 | 0.515 | 0.29 | 2.63E-23 | 0 L1 _ | Rnd3 |
| Atraid | 1.30E-27 | 0.311202 | 0.802 | 0.59 | 2.61E-23 | 0 L1 _ | Atraid |
| Gpnmb | 9.30E-28 | 0.28622 | 0.175 | 0.042 | 1.87E-23 | 0 L1 _ | Gpnmb |
| Ptov1 | 8.55E-28 | 0.257406 | 0.645 | 0.389 | 1.72E-23 | 0 L1 _ | Ptov1 |
| Tubb2b | 7.97E-28 | 0.299681 | 0.216 | 0.065 | 1.60E-23 | 0 L1 _ | Tubb2b |
| Dock71 | 7.13E-28 | 0.294577 | 0.453 | 0.238 | 1.44E-23 | 0 L1 _ | Dock7 |
| Rhoj2 | 3.74E-28 | 0.352309 | 0.89 | 0.642 | 7.53E-24 | 0 L1 _ | Rhoj |
| Rab2a | 2.84E-28 | 0.348772 | 0.906 | 0.733 | 5.73E-24 | 0 L1 _ | Rab2a |
| Itga81 | 2.54E-28 | 0.352501 | 0.31 | 0.13 | 5.11E-24 | 0 L1 _ | Itga8 |
| Adprh | 2.19E-28 | 0.337465 | 0.695 | 0.466 | 4.41E-24 | 0 L1 _ | Adprh |
| Sar1a1 | 1.59E-28 | 0.384565 | 0.729 | 0.494 | 3.20E-24 | 0 L1 _ | Sar1a |
| Ror1 | 1.56E-28 | 0.259321 | 0.313 | 0.125 | 3.15E-24 | 0 L1 _ | Ror1 |
| Pbx11 | 1.42E-28 | 0.368083 | 0.685 | 0.416 | 2.86E-24 | 0 L1 _ | Pbx1 |
| Kctd11 | 1.40E-28 | 0.25456 | 0.299 | 0.115 | 2.81E-24 | 0 L1 _ | Kctd11 |
| Tram11 | 6.90E-29 | 0.350835 | 0.748 | 0.504 | 1.39E-24 | 0 L1 _ | Tram1 |
| Naxd | 6.08E-29 | 0.277608 | 0.62 | 0.381 | 1.23E-24 | 0 L1 _ | Naxd |
| Ttc3 | 5.33E-29 | 0.369448 | 0.721 | 0.475 | 1.07E-24 | 0 L1 _ | Ttc3 |
| Nid21 | 5.02E-29 | 0.263262 | 0.517 | 0.264 | 1.01E-24 | 0 L1 _ | Nid2 |
| Fgf2 | 4.69E-29 | 0.280301 | 0.322 | 0.134 | 9.44E-25 | 0 L1 _ | Fgf2 |
| Tmed1 | 4.24E-29 | 0.293251 | 0.487 | 0.257 | 8.54E-25 | 0 L1 _ | Tmed1 |
| Sec24d1 | 3.42E-29 | 0.327657 | 0.45 | 0.236 | 6.89E-25 | 0 L1 _ | Sec24d |
| Ggcx | 2.89E-29 | 0.269503 | 0.362 | 0.162 | 5.82E-25 | 0 L1 _ | Ggcx |
| Stt3a1 | 2.51E-29 | 0.326671 | 0.628 | 0.366 | 5.05E-25 | 0 L1 _ | Stt3a |
| Hmox1 | 2.21E-29 | 0.268725 | 0.255 | 0.089 | 4.46E-25 | 0 L1 _ | Hmox1 |
| Rgma1 | 2.15E-29 | 0.274422 | 0.528 | 0.277 | 4.32E-25 | 0 L1 _ | Rgma |
| Piezo21 | 1.76E-29 | 0.328931 | 0.51 | 0.258 | 3.54E-25 | 0 L1 _ | Piezo2 |
| Ro60 | 1.39E-29 | 0.312108 | 0.425 | 0.202 | 2.80E-25 | 0 L1 _ | Ro60 |
| Rab3il1 | 8.57E-30 | 0.323038 | 0.38 | 0.183 | 1.73E-25 | 0 L1 _ | Rab3il1 |
| Ahnak1 | 7.76E-30 | 0.360577 | 0.975 | 0.841 | 1.56E-25 | 0 L1 _ | Ahnak |
| Icam1 | 6.48E-30 | 0.349442 | 0.41 | 0.198 | 1.31E-25 | 0 L1 _ | Icam1 |
| Ift20 | 5.77E-30 | 0.326757 | 0.808 | 0.574 | 1.16E-25 | 0 L1 _ | Ift20 |
| Ddost | 5.09E-30 | 0.37825 | 0.79 | 0.539 | 1.03E-25 | 0 L1 _ | Ddost |
| Sertad4 | 4.94E-30 | 0.292628 | 0.258 | 0.087 | 9.94E-26 | 0 L1 _ | Sertad4 |
| Tmem167 | 3.52E-30 | 0.361852 | 0.786 | 0.565 | 7.08E-26 | 0 L1 _ | Tmem167 |
| Pofut21 | 3.11E-30 | 0.317925 | 0.611 | 0.356 | 6.27E-26 | 0 L1 _ | Pofut2 |
| Col28a1 | 3.03E-30 | 0.379532 | 0.176 | 0.038 | 6.10E-26 | 0 L1 _ | Col28a1 |
| Anxa52 | 2.99E-30 | 0.326196 | 0.961 | 0.837 | 6.03E-26 | 0 L1 _ | Anxa5 |
| Ifi2071 | 2.64E-30 | 0.319604 | 0.481 | 0.238 | 5.31E-26 | 0 L1 _ | Ifi207 |
| Cbr31 | 2.37E-30 | 0.30862 | 0.504 | 0.262 | 4.78E-26 | 0 L1 _ | Cbr3 |
| S1pr21 | 2.30E-30 | 0.306245 | 0.373 | 0.167 | 4.63E-26 | 0 L1 _ | S1pr2 |

| | | | | | | | |
|---|---|---|---|---|---|---|---|
| Cyb5a1 | 2.02E-30 | 0.384345 | 0.9 | 0.684 | 4.07E-26 | 0 L1 _ | Cyb5a |
| Lrrc581 | 1.68E-30 | 0.386142 | 0.827 | 0.567 | 3.38E-26 | 0 L1 _ | Lrrc58 |
| Igsf31 | 1.60E-30 | 0.327991 | 0.357 | 0.149 | 3.23E-26 | 0 L1 _ | Igsf3 |
| Tm7sf3 | 1.28E-30 | 0.347559 | 0.481 | 0.254 | 2.58E-26 | 0 L1 _ | Tm7sf3 |
| Dse1 | 9.20E-31 | 0.312441 | 0.412 | 0.198 | 1.85E-26 | 0 L1 _ | Dse |
| Art4 | 6.58E-31 | 0.345157 | 0.22 | 0.061 | 1.32E-26 | 0 L1 _ | Art4 |
| Atp6ap1 | 6.12E-31 | 0.373285 | 0.726 | 0.462 | 1.23E-26 | 0 L1 _ | Atp6ap1 |
| Sec61g1 | 5.17E-31 | 0.405536 | 0.927 | 0.814 | 1.04E-26 | 0 L1 _ | Sec61g |
| Tnfrsf19 | 5.06E-31 | 0.263608 | 0.164 | 0.03 | 1.02E-26 | 0 L1 _ | Tnfrsf19 |
| Ltbp11 | 5.00E-31 | 0.431057 | 0.566 | 0.314 | 1.01E-26 | 0 L1 _ | Ltbp1 |
| Rnf150 | 4.52E-31 | 0.308283 | 0.314 | 0.121 | 9.10E-27 | 0 L1 _ | Rnf150 |
| Tubb2a | 4.36E-31 | 0.467919 | 0.581 | 0.338 | 8.78E-27 | 0 L1 _ | Tubb2a |
| Tspan4 | 2.94E-31 | 0.35287 | 0.605 | 0.352 | 5.92E-27 | 0 L1 _ | Tspan4 |
| Tnfrsf12a | 2.75E-31 | 0.472882 | 0.252 | 0.081 | 5.54E-27 | 0 L1 _ | Tnfrsf12a |
| Tm2d21 | 2.72E-31 | 0.318905 | 0.642 | 0.391 | 5.47E-27 | 0 L1 _ | Tm2d2 |
| Ldlr | 2.16E-31 | 0.336384 | 0.356 | 0.15 | 4.36E-27 | 0 L1 _ | Ldlr |
| Dsel1 | 2.13E-31 | 0.354192 | 0.455 | 0.226 | 4.29E-27 | 0 L1 _ | Dsel |
| Fscn1 | 1.76E-31 | 0.277231 | 0.481 | 0.239 | 3.55E-27 | 0 L1 _ | Fscn1 |
| Sh3pxd2b1 | 1.62E-31 | 0.318067 | 0.474 | 0.229 | 3.26E-27 | 0 L1 _ | Sh3pxd2b |
| Itih2 | 1.62E-31 | 0.256243 | 0.182 | 0.039 | 3.25E-27 | 0 L1 _ | Itih2 |
| Slc1a5 | 1.33E-31 | 0.365237 | 0.658 | 0.399 | 2.67E-27 | 0 L1 _ | Slc1a5 |
| Fgl21 | 1.30E-31 | 0.493045 | 0.557 | 0.31 | 2.62E-27 | 0 L1 _ | Fgl2 |
| Cul1 | 8.69E-32 | 0.400327 | 0.739 | 0.495 | 1.75E-27 | 0 L1 _ | Cul1 |
| Mocs2 | 7.60E-32 | 0.312141 | 0.619 | 0.366 | 1.53E-27 | 0 L1 _ | Mocs2 |
| Surf41 | 6.47E-32 | 0.371374 | 0.709 | 0.441 | 1.30E-27 | 0 L1 _ | Surf4 |
| Rcn2 | 6.11E-32 | 0.296443 | 0.658 | 0.39 | 1.23E-27 | 0 L1 _ | Rcn2 |
| Shox21 | 3.42E-32 | 0.326858 | 0.386 | 0.166 | 6.90E-28 | 0 L1 _ | Shox2 |
| Sgce1 | 3.29E-32 | 0.293746 | 0.586 | 0.314 | 6.62E-28 | 0 L1 _ | Sgce |
| Nr2f12 | 3.16E-32 | 0.274628 | 0.8 | 0.465 | 6.37E-28 | 0 L1 _ | Nr2f1 |
| 2900097C17Rik1 | 3.08E-32 | 0.360997 | 0.717 | 0.446 | 6.20E-28 | 0 L1 _ | 2900097C17Rik |
| Rgmb1 | 3.04E-32 | 0.386333 | 0.438 | 0.215 | 6.13E-28 | 0 L1 _ | Rgmb |
| Sec22b | 2.89E-32 | 0.300287 | 0.566 | 0.322 | 5.83E-28 | 0 L1 _ | Sec22b |
| Pthlh | 2.65E-32 | 0.404841 | 0.249 | 0.078 | 5.34E-28 | 0 L1 _ | Pthlh |
| Scml4 | 2.49E-32 | 0.321786 | 0.294 | 0.107 | 5.02E-28 | 0 L1 _ | Scml4 |
| Fndc3b1 | 2.26E-32 | 0.312099 | 0.71 | 0.413 | 4.55E-28 | 0 L1 _ | Fndc3b |
| Mettl11 | 2.11E-32 | 0.369095 | 0.421 | 0.202 | 4.24E-28 | 0 L1 _ | Mettl1 |
| Ndfip11 | 2.09E-32 | 0.326261 | 0.88 | 0.621 | 4.21E-28 | 0 L1 _ | Ndfip1 |
| Zfyve21 | 1.99E-32 | 0.275414 | 0.462 | 0.219 | 4.02E-28 | 0 L1 _ | Zfyve21 |
| Bnc2 | 1.69E-32 | 0.314359 | 0.264 | 0.087 | 3.40E-28 | 0 L1 _ | Bnc2 |
| Ext11 | 1.33E-32 | 0.358087 | 0.515 | 0.272 | 2.68E-28 | 0 L1 _ | Ext1 |
| P2rx4 | 1.11E-32 | 0.317812 | 0.473 | 0.233 | 2.23E-28 | 0 L1 _ | P2rx4 |

| | | | | | | | |
|---|---|---|---|---|---|---|---|
| Plbd2 | 1.07E-32 | 0.353692 | 0.49 | 0.252 | 2.15E-28 | 0 L1 _ | Plbd2 |
| Fkbp101 | 1.02E-32 | 0.297084 | 0.443 | 0.214 | 2.06E-28 | 0 L1 _ | Fkbp10 |
| Itga2b | 9.89E-33 | 0.298974 | 0.269 | 0.089 | 1.99E-28 | 0 L1 _ | Itga2b |
| Colgalt1 | 7.77E-33 | 0.388645 | 0.702 | 0.445 | 1.57E-28 | 0 L1 _ | Colgalt1 |
| Tcea3 | 6.05E-33 | 0.296314 | 0.258 | 0.08 | 1.22E-28 | 0 L1 _ | Tcea3 |
| Myh101 | 5.70E-33 | 0.36138 | 0.361 | 0.151 | 1.15E-28 | 0 L1 _ | Myh10 |
| Swi5 | 5.51E-33 | 0.375544 | 0.887 | 0.689 | 1.11E-28 | 0 L1 _ | Swi5 |
| Apbb1ip1 | 4.28E-33 | 0.365905 | 0.61 | 0.335 | 8.63E-29 | 0 L1 _ | Apbb1ip |
| Rbis | 3.86E-33 | 0.272803 | 0.622 | 0.329 | 7.77E-29 | 0 L1 _ | Rbis |
| Csf11 | 2.95E-33 | 0.384718 | 0.549 | 0.298 | 5.94E-29 | 0 L1 _ | Csf1 |
| Boc1 | 2.93E-33 | 0.337575 | 0.332 | 0.132 | 5.91E-29 | 0 L1 _ | Boc |
| Calu1 | 2.18E-33 | 0.37926 | 0.809 | 0.513 | 4.39E-29 | 0 L1 _ | Calu |
| Gpx31 | 2.16E-33 | 0.707527 | 0.551 | 0.284 | 4.36E-29 | 0 L1 _ | Gpx3 |
| Ap3s1 | 1.85E-33 | 0.34379 | 0.54 | 0.297 | 3.73E-29 | 0 L1 _ | Ap3s1 |
| Pkig1 | 1.78E-33 | 0.384395 | 0.777 | 0.514 | 3.58E-29 | 0 L1 _ | Pkig |
| Serpine1 | 1.59E-33 | 0.490481 | 0.223 | 0.058 | 3.20E-29 | 0 L1 _ | Serpine1 |
| Czib | 1.25E-33 | 0.272375 | 0.442 | 0.199 | 2.53E-29 | 0 L1 _ | Czib |
| Bsg | 1.13E-33 | 0.358594 | 0.933 | 0.775 | 2.28E-29 | 0 L1 _ | Bsg |
| Gab2 | 1.12E-33 | 0.330537 | 0.37 | 0.155 | 2.25E-29 | 0 L1 _ | Gab2 |
| Dclk11 | 1.00E-33 | 0.388478 | 0.344 | 0.136 | 2.01E-29 | 0 L1 _ | Dclk1 |
| Eif1a | 8.73E-34 | 0.353354 | 0.617 | 0.366 | 1.76E-29 | 0 L1 _ | Eif1a |
| Ghr | 7.38E-34 | 0.340721 | 0.429 | 0.2 | 1.49E-29 | 0 L1 _ | Ghr |
| Lman11 | 5.91E-34 | 0.369935 | 0.781 | 0.525 | 1.19E-29 | 0 L1 _ | Lman1 |
| Antxr11 | 5.81E-34 | 0.31791 | 0.45 | 0.21 | 1.17E-29 | 0 L1 _ | Antxr1 |
| Tgfbr1 | 4.93E-34 | 0.326 | 0.551 | 0.302 | 9.93E-30 | 0 L1 _ | Tgfbr1 |
| Cst3 | 4.82E-34 | 0.738882 | 0.983 | 0.933 | 9.71E-30 | 0 L1 _ | Cst3 |
| Dnajc31 | 2.40E-34 | 0.407233 | 0.86 | 0.615 | 4.83E-30 | 0 L1 _ | Dnajc3 |
| Sema3a1 | 2.27E-34 | 0.377966 | 0.307 | 0.108 | 4.56E-30 | 0 L1 _ | Sema3a |
| Chic2 | 2.25E-34 | 0.375521 | 0.718 | 0.452 | 4.54E-30 | 0 L1 _ | Chic2 |
| Bace11 | 2.20E-34 | 0.334104 | 0.525 | 0.266 | 4.42E-30 | 0 L1 _ | Bace1 |
| Emp31 | 1.61E-34 | 0.407318 | 0.926 | 0.725 | 3.25E-30 | 0 L1 _ | Emp3 |
| Hsp90b11 | 1.35E-34 | 0.368895 | 0.965 | 0.845 | 2.71E-30 | 0 L1 _ | Hsp90b1 |
| Gm2a | 4.97E-35 | 0.351974 | 0.529 | 0.273 | 1.00E-30 | 0 L1 _ | Gm2a |
| Yipf51 | 4.41E-35 | 0.368495 | 0.65 | 0.386 | 8.88E-31 | 0 L1 _ | Yipf5 |
| Slc7a2 | 3.72E-35 | 0.329619 | 0.368 | 0.141 | 7.49E-31 | 0 L1 _ | Slc7a2 |
| Abat | 3.08E-35 | 0.328223 | 0.314 | 0.114 | 6.21E-31 | 0 L1 _ | Abat |
| Igfbp71 | 2.92E-35 | 0.339385 | 0.992 | 0.912 | 5.87E-31 | 0 L1 _ | Igfbp7 |
| Twsg11 | 2.70E-35 | 0.371901 | 0.535 | 0.274 | 5.44E-31 | 0 L1 _ | Twsg1 |
| Odr4 | 1.97E-35 | 0.320081 | 0.435 | 0.188 | 3.97E-31 | 0 L1 _ | Odr4 |
| Slc39a13 | 1.94E-35 | 0.327492 | 0.501 | 0.25 | 3.90E-31 | 0 L1 _ | Slc39a13 |
| Wipi11 | 1.54E-35 | 0.343704 | 0.561 | 0.291 | 3.09E-31 | 0 L1 _ | Wipi1 |

| | | | | | | | |
|---|---|---|---|---|---|---|---|
| Ap2a2 | 1.28E-35 | 0.415372 | 0.695 | 0.433 | 2.58E-31 | 0 L1 _ | Ap2a2 |
| Gm26682 | 1.14E-35 | 0.31179 | 0.224 | 0.053 | 2.30E-31 | 0 L1 _ | Gm26682 |
| Nnmt | 1.11E-35 | 0.307821 | 0.449 | 0.2 | 2.24E-31 | 0 L1 _ | Nnmt |
| Plpp5 | 1.02E-35 | 0.342683 | 0.39 | 0.17 | 2.06E-31 | 0 L1 _ | Plpp5 |
| Enho | 9.72E-36 | 0.362781 | 0.345 | 0.13 | 1.96E-31 | 0 L1 _ | Enho |
| Csad | 8.31E-36 | 0.330284 | 0.438 | 0.205 | 1.67E-31 | 0 L1 _ | Csad |
| Arf5 | 6.88E-36 | 0.428006 | 0.893 | 0.72 | 1.38E-31 | 0 L1 _ | Arf5 |
| Pld3 | 6.57E-36 | 0.356913 | 0.593 | 0.329 | 1.32E-31 | 0 L1 _ | Pld3 |
| Erp29 | 5.63E-36 | 0.380619 | 0.815 | 0.572 | 1.13E-31 | 0 L1 _ | Erp29 |
| Myo7a | 4.10E-36 | 0.29052 | 0.224 | 0.054 | 8.26E-32 | 0 L1 _ | Myo7a |
| Mir22hg | 2.78E-36 | 0.304102 | 0.274 | 0.08 | 5.60E-32 | 0 L1 _ | Mir22hg |
| Tmed10 | 2.67E-36 | 0.38526 | 0.934 | 0.776 | 5.37E-32 | 0 L1 _ | Tmed10 |
| Tmem2631 | 2.52E-36 | 0.345058 | 0.472 | 0.227 | 5.07E-32 | 0 L1 _ | Tmem263 |
| Robo11 | 1.77E-36 | 0.337397 | 0.361 | 0.141 | 3.56E-32 | 0 L1 _ | Robo1 |
| Cnn3 | 1.60E-36 | 0.403368 | 0.777 | 0.512 | 3.23E-32 | 0 L1 _ | Cnn3 |
| Ctsd1 | 1.18E-36 | 0.363363 | 0.947 | 0.765 | 2.37E-32 | 0 L1 _ | Ctsd |
| Fam102b1 | 1.17E-36 | 0.379626 | 0.47 | 0.212 | 2.35E-32 | 0 L1 _ | Fam102b |
| Atp6v0c | 1.05E-36 | 0.370607 | 0.885 | 0.541 | 2.12E-32 | 0 L1 _ | Atp6v0c |
| Lmna1 | 8.64E-37 | 0.443007 | 0.847 | 0.593 | 1.74E-32 | 0 L1 _ | Lmna |
| Sned11 | 8.58E-37 | 0.494788 | 0.511 | 0.249 | 1.73E-32 | 0 L1 _ | Sned1 |
| Yif1a | 7.74E-37 | 0.359983 | 0.543 | 0.285 | 1.56E-32 | 0 L1 _ | Yif1a |
| Dap | 7.60E-37 | 0.433961 | 0.663 | 0.401 | 1.53E-32 | 0 L1 _ | Dap |
| Tmem223 | 6.80E-37 | 0.313339 | 0.542 | 0.267 | 1.37E-32 | 0 L1 _ | Tmem223 |
| Ifitm3 | 6.26E-37 | 0.423562 | 0.991 | 0.905 | 1.26E-32 | 0 L1 _ | Ifitm3 |
| Clcn51 | 6.13E-37 | 0.352832 | 0.384 | 0.153 | 1.24E-32 | 0 L1 _ | Clcn5 |
| Anxa21 | 5.81E-37 | 0.451283 | 0.929 | 0.794 | 1.17E-32 | 0 L1 _ | Anxa2 |
| Ifngr11 | 5.71E-37 | 0.424423 | 0.888 | 0.629 | 1.15E-32 | 0 L1 _ | Ifngr1 |
| Tiam1 | 5.51E-37 | 0.306455 | 0.243 | 0.063 | 1.11E-32 | 0 L1 _ | Tiam1 |
| Ryk | 4.32E-37 | 0.408056 | 0.678 | 0.41 | 8.71E-33 | 0 L1 _ | Ryk |
| Aldh7a11 | 3.57E-37 | 0.343624 | 0.508 | 0.242 | 7.18E-33 | 0 L1 _ | Aldh7a1 |
| Npc2 | 2.43E-37 | 0.432293 | 0.882 | 0.668 | 4.89E-33 | 0 L1 _ | Npc2 |
| Dipk1b | 2.42E-37 | 0.273078 | 0.257 | 0.069 | 4.88E-33 | 0 L1 _ | Dipk1b |
| Echdc2 | 2.16E-37 | 0.286134 | 0.236 | 0.059 | 4.36E-33 | 0 L1 _ | Echdc2 |
| Slc8a1 | 1.54E-37 | 0.331766 | 0.264 | 0.073 | 3.11E-33 | 0 L1 _ | Slc8a1 |
| Axl2 | 1.51E-37 | 0.322925 | 0.781 | 0.443 | 3.03E-33 | 0 L1 _ | Axl |
| Desi2 | 1.43E-37 | 0.367016 | 0.614 | 0.342 | 2.88E-33 | 0 L1 _ | Desi2 |
| Tnfaip2 | 1.43E-37 | 0.503769 | 0.59 | 0.321 | 2.88E-33 | 0 L1 _ | Tnfaip2 |
| Nrxn21 | 1.31E-37 | 0.445505 | 0.43 | 0.18 | 2.64E-33 | 0 L1 _ | Nrxn2 |
| Dapk11 | 1.11E-37 | 0.363171 | 0.534 | 0.249 | 2.24E-33 | 0 L1 _ | Dapk1 |
| Lgmn | 1.01E-37 | 0.321089 | 0.563 | 0.283 | 2.04E-33 | 0 L1 _ | Lgmn |
| Slc39a7 | 9.29E-38 | 0.363748 | 0.611 | 0.345 | 1.87E-33 | 0 L1 _ | Slc39a7 |

| | | | | | | | |
|---|---|---|---|---|---|---|---|
| Cdk141 | 8.79E-38 | 0.444507 | 0.581 | 0.312 | 1.77E-33 | 0 L1 _ | Cdk14 |
| Cks1b | 8.19E-38 | 0.305026 | 0.337 | 0.118 | 1.65E-33 | 0 L1 _ | Cks1b |
| Golim41 | 4.80E-38 | 0.406181 | 0.745 | 0.466 | 9.67E-34 | 0 L1 _ | Golim4 |
| Celf21 | 3.37E-38 | 0.381368 | 0.784 | 0.438 | 6.79E-34 | 0 L1 _ | Celf2 |
| Dact3 | 3.20E-38 | 0.338404 | 0.346 | 0.125 | 6.44E-34 | 0 L1 _ | Dact3 |
| Krtcap2 | 2.17E-38 | 0.378432 | 0.919 | 0.756 | 4.36E-34 | 0 L1 _ | Krtcap2 |
| Nr1d1 | 2.12E-38 | 0.476857 | 0.468 | 0.22 | 4.28E-34 | 0 L1 _ | Nr1d1 |
| Egflam | 9.44E-39 | 0.314591 | 0.607 | 0.309 | 1.90E-34 | 0 L1 _ | Egflam |
| Tmem176a | 8.18E-39 | 0.470658 | 0.94 | 0.73 | 1.65E-34 | 0 L1 _ | Tmem176a |
| Lix1l | 7.30E-39 | 0.376286 | 0.577 | 0.312 | 1.47E-34 | 0 L1 _ | Lix1l |
| Nrep | 6.88E-39 | 0.562221 | 0.438 | 0.201 | 1.39E-34 | 0 L1 _ | Nrep |
| Arhgap6 | 6.75E-39 | 0.402554 | 0.589 | 0.305 | 1.36E-34 | 0 L1 _ | Arhgap6 |
| Lamc11 | 6.65E-39 | 0.374548 | 0.807 | 0.482 | 1.34E-34 | 0 L1 _ | Lamc1 |
| Cyp51 | 6.48E-39 | 0.457675 | 0.351 | 0.13 | 1.31E-34 | 0 L1 _ | Cyp51 |
| Prrc1 | 4.66E-39 | 0.416478 | 0.549 | 0.286 | 9.38E-35 | 0 L1 _ | Prrc1 |
| Zbtb201 | 4.61E-39 | 0.452824 | 0.973 | 0.851 | 9.29E-35 | 0 L1 _ | Zbtb20 |
| Cacna1g | 3.98E-39 | 0.349742 | 0.285 | 0.085 | 8.02E-35 | 0 L1 _ | Cacna1g |
| Postn1 | 3.41E-39 | 0.799799 | 0.632 | 0.36 | 6.86E-35 | 0 L1 _ | Postn |
| Pcdh181 | 1.73E-39 | 0.393057 | 0.623 | 0.328 | 3.48E-35 | 0 L1 _ | Pcdh18 |
| Olfm2 | 1.69E-39 | 0.3483 | 0.201 | 0.035 | 3.41E-35 | 0 L1 _ | Olfm2 |
| Actn1 | 1.61E-39 | 0.347164 | 0.675 | 0.375 | 3.24E-35 | 0 L1 _ | Actn1 |
| Adgra1 | 1.37E-39 | 0.425652 | 0.197 | 0.032 | 2.76E-35 | 0 L1 _ | Adgra1 |
| Idh11 | 1.32E-39 | 0.380009 | 0.511 | 0.244 | 2.66E-35 | 0 L1 _ | Idh1 |
| Slc39a11 | 5.13E-40 | 0.440676 | 0.613 | 0.337 | 1.03E-35 | 0 L1 _ | Slc39a1 |
| Phyhd1 | 4.48E-40 | 0.293759 | 0.257 | 0.063 | 9.02E-36 | 0 L1 _ | Phyhd1 |
| Ptges1 | 3.29E-40 | 0.386806 | 0.41 | 0.165 | 6.63E-36 | 0 L1 _ | Ptges |
| Eri3 | 3.14E-40 | 0.33489 | 0.479 | 0.218 | 6.33E-36 | 0 L1 _ | Eri3 |
| Itgav1 | 3.07E-40 | 0.464908 | 0.652 | 0.368 | 6.19E-36 | 0 L1 _ | Itgav |
| H6pd1 | 2.34E-40 | 0.41147 | 0.455 | 0.201 | 4.72E-36 | 0 L1 _ | H6pd |
| Scd21 | 2.14E-40 | 0.3689 | 0.711 | 0.392 | 4.30E-36 | 0 L1 _ | Scd2 |
| Runx2 | 1.83E-40 | 0.376752 | 0.369 | 0.141 | 3.69E-36 | 0 L1 _ | Runx2 |
| Morf4l21 | 1.78E-40 | 0.424583 | 0.758 | 0.471 | 3.58E-36 | 0 L1 _ | Morf4l2 |
| Naalad21 | 1.39E-40 | 0.372571 | 0.666 | 0.35 | 2.79E-36 | 0 L1 _ | Naalad2 |
| Rftn1 | 1.07E-40 | 0.453884 | 0.643 | 0.37 | 2.15E-36 | 0 L1 _ | Rftn1 |
| Wnt9a | 9.95E-41 | 0.391846 | 0.226 | 0.048 | 2.00E-36 | 0 L1 _ | Wnt9a |
| Mmp191 | 6.53E-41 | 0.301026 | 0.472 | 0.197 | 1.32E-36 | 0 L1 _ | Mmp19 |
| Calhm5 | 5.47E-41 | 0.330681 | 0.251 | 0.059 | 1.10E-36 | 0 L1 _ | Calhm5 |
| Plod11 | 5.31E-41 | 0.412097 | 0.736 | 0.42 | 1.07E-36 | 0 L1 _ | Plod1 |
| Pkd21 | 4.64E-41 | 0.442724 | 0.712 | 0.393 | 9.35E-37 | 0 L1 _ | Pkd2 |
| Micos10 | 4.13E-41 | 0.382826 | 0.834 | 0.482 | 8.32E-37 | 0 L1 _ | Micos10 |
| Robo2 | 2.47E-41 | 0.431018 | 0.302 | 0.093 | 4.96E-37 | 0 L1 _ | Robo2 |

| | | | | | | | |
|---|---|---|---|---|---|---|---|
| Laptm4a1 | 2.39E-41 | 0.33044 | 0.983 | 0.887 | 4.82E-37 | 0 L1 _ | Laptm4a |
| Fxyd11 | 1.49E-41 | 0.342776 | 0.884 | 0.539 | 3.00E-37 | 0 L1 _ | Fxyd1 |
| Rspo31 | 7.31E-42 | 0.486007 | 0.433 | 0.167 | 1.47E-37 | 0 L1 _ | Rspo3 |
| Akr1b8 | 6.17E-42 | 0.357812 | 0.371 | 0.133 | 1.24E-37 | 0 L1 _ | Akr1b8 |
| P3h1 | 4.64E-42 | 0.378762 | 0.395 | 0.151 | 9.35E-38 | 0 L1 _ | P3h1 |
| Mpg | 3.93E-42 | 0.390685 | 0.499 | 0.234 | 7.91E-38 | 0 L1 _ | Mpg |
| Plod21 | 3.19E-42 | 0.448834 | 0.471 | 0.21 | 6.42E-38 | 0 L1 _ | Plod2 |
| Manf | 2.55E-42 | 0.612428 | 0.81 | 0.595 | 5.14E-38 | 0 L1 _ | Manf |
| Gng10 | 1.81E-42 | 0.407547 | 0.535 | 0.256 | 3.65E-38 | 0 L1 _ | Gng10 |
| Sesn3 | 1.80E-42 | 0.451399 | 0.658 | 0.377 | 3.63E-38 | 0 L1 _ | Sesn3 |
| Stc2 | 1.63E-42 | 0.374943 | 0.242 | 0.051 | 3.28E-38 | 0 L1 _ | Stc2 |
| Adamts22 | 1.41E-42 | 0.363511 | 0.474 | 0.197 | 2.84E-38 | 0 L1 _ | Adamts2 |
| Lgals3bp | 1.34E-42 | 0.488085 | 0.8 | 0.499 | 2.70E-38 | 0 L1 _ | Lgals3bp |
| Rxylt1 | 1.33E-42 | 0.317821 | 0.536 | 0.231 | 2.68E-38 | 0 L1 _ | Rxylt1 |
| Cpz | 1.04E-42 | 0.389758 | 0.198 | 0.029 | 2.09E-38 | 0 L1 _ | Cpz |
| Dpp7 | 7.87E-43 | 0.332773 | 0.348 | 0.115 | 1.59E-38 | 0 L1 _ | Dpp7 |
| Fam20c | 6.85E-43 | 0.348723 | 0.315 | 0.097 | 1.38E-38 | 0 L1 _ | Fam20c |
| Tmem176b | 5.77E-43 | 0.526158 | 0.96 | 0.768 | 1.16E-38 | 0 L1 _ | Tmem176b |
| Arl11 | 4.05E-43 | 0.46719 | 0.818 | 0.559 | 8.15E-39 | 0 L1 _ | Arl1 |
| Tspan33 | 4.03E-43 | 0.384009 | 0.845 | 0.547 | 8.11E-39 | 0 L1 _ | Tspan3 |
| Sspn1 | 3.80E-43 | 0.406127 | 0.523 | 0.241 | 7.65E-39 | 0 L1 _ | Sspn |
| Antxr22 | 2.27E-43 | 0.292113 | 0.589 | 0.264 | 4.57E-39 | 0 L1 _ | Antxr2 |
| Dnajc251 | 2.26E-43 | 0.429266 | 0.492 | 0.224 | 4.55E-39 | 0 L1 _ | Dnajc25 |
| Serpina3n | 2.16E-43 | 0.565649 | 0.306 | 0.088 | 4.34E-39 | 0 L1 _ | Serpina3n |
| Adamts10 | 1.98E-43 | 0.486734 | 0.6 | 0.311 | 3.98E-39 | 0 L1 _ | Adamts10 |
| P4ha11 | 1.95E-43 | 0.437917 | 0.669 | 0.368 | 3.93E-39 | 0 L1 _ | P4ha1 |
| Fads11 | 1.90E-43 | 0.420001 | 0.473 | 0.205 | 3.82E-39 | 0 L1 _ | Fads1 |
| Plekhf11 | 1.38E-43 | 0.380971 | 0.404 | 0.154 | 2.78E-39 | 0 L1 _ | Plekhf1 |
| Col11a1 | 1.13E-43 | 0.411844 | 0.223 | 0.04 | 2.28E-39 | 0 L1 _ | Col11a1 |
| Ostc | 1.09E-43 | 0.500587 | 0.816 | 0.585 | 2.19E-39 | 0 L1 _ | Ostc |
| Tafa5 | 9.81E-44 | 0.277909 | 0.207 | 0.032 | 1.98E-39 | 0 L1 _ | Tafa5 |
| Timm23 | 8.59E-44 | 0.397747 | 0.587 | 0.282 | 1.73E-39 | 0 L1 _ | Timm23 |
| Sdc41 | 5.72E-44 | 0.51043 | 0.532 | 0.251 | 1.15E-39 | 0 L1 _ | Sdc4 |
| Ccpg1 | 3.37E-44 | 0.421135 | 0.612 | 0.311 | 6.79E-40 | 0 L1 _ | Ccpg1 |
| Tmem150a | 3.31E-44 | 0.361482 | 0.387 | 0.141 | 6.66E-40 | 0 L1 _ | Tmem150a |
| Prxl2c | 2.38E-44 | 0.373297 | 0.49 | 0.205 | 4.80E-40 | 0 L1 _ | Prxl2c |
| Thbs11 | 1.71E-44 | 0.570495 | 0.405 | 0.151 | 3.45E-40 | 0 L1 _ | Thbs1 |
| Fkbp11 | 1.32E-44 | 0.375738 | 0.325 | 0.099 | 2.65E-40 | 0 L1 _ | Fkbp11 |
| Slc29a11 | 1.27E-44 | 0.488532 | 0.803 | 0.486 | 2.55E-40 | 0 L1 _ | Slc29a1 |
| Slc36a2 | 1.01E-44 | 0.335272 | 0.241 | 0.047 | 2.03E-40 | 0 L1 _ | Slc36a2 |
| Msx1 | 7.27E-45 | 0.478827 | 0.419 | 0.171 | 1.46E-40 | 0 L1 _ | Msx1 |

| | | | | | | | |
|---|---|---|---|---|---|---|---|
| Slit31 | 5.07E-45 | 0.445498 | 0.555 | 0.262 | 1.02E-40 | 0 L1 _ | Slit3 |
| Fbln51 | 2.20E-45 | 0.297222 | 0.577 | 0.262 | 4.43E-41 | 0 L1 _ | Fbln5 |
| P2ry11 | 2.13E-45 | 0.440015 | 0.426 | 0.159 | 4.29E-41 | 0 L1 _ | P2ry1 |
| Hmgn3 | 1.43E-45 | 0.475058 | 0.689 | 0.39 | 2.87E-41 | 0 L1 _ | Hmgn3 |
| Scarb21 | 1.41E-45 | 0.438167 | 0.734 | 0.415 | 2.85E-41 | 0 L1 _ | Scarb2 |
| Mmp231 | 1.07E-45 | 0.388527 | 0.351 | 0.112 | 2.15E-41 | 0 L1 _ | Mmp23 |
| Myo1d | 9.67E-46 | 0.456817 | 0.443 | 0.178 | 1.95E-41 | 0 L1 _ | Myo1d |
| Ifi27l2a | 7.11E-46 | 0.62753 | 0.793 | 0.479 | 1.43E-41 | 0 L1 _ | Ifi27l2a |
| Penk1 | 5.13E-46 | 0.848753 | 0.401 | 0.148 | 1.03E-41 | 0 L1 _ | Penk |
| Rcn1 | 4.13E-46 | 0.47059 | 0.732 | 0.432 | 8.32E-42 | 0 L1 _ | Rcn1 |
| Tmem98 | 4.13E-46 | 0.297289 | 0.272 | 0.062 | 8.32E-42 | 0 L1 _ | Tmem98 |
| Arhgef401 | 3.77E-46 | 0.496255 | 0.63 | 0.333 | 7.60E-42 | 0 L1 _ | Arhgef40 |
| Gata6 | 2.90E-46 | 0.375132 | 0.346 | 0.107 | 5.84E-42 | 0 L1 _ | Gata6 |
| Tmem200a | 2.15E-46 | 0.328866 | 0.22 | 0.034 | 4.33E-42 | 0 L1 _ | Tmem200a |
| Casc4 | 1.47E-46 | 0.405641 | 0.563 | 0.262 | 2.95E-42 | 0 L1 _ | Casc4 |
| Poglut3 | 1.24E-46 | 0.380757 | 0.407 | 0.142 | 2.50E-42 | 0 L1 _ | Poglut3 |
| Arf4 | 1.05E-46 | 0.482856 | 0.899 | 0.679 | 2.11E-42 | 0 L1 _ | Arf4 |
| Lamp1 | 8.50E-47 | 0.444617 | 0.959 | 0.804 | 1.71E-42 | 0 L1 _ | Lamp1 |
| Ifi2041 | 7.83E-47 | 0.475572 | 0.535 | 0.239 | 1.58E-42 | 0 L1 _ | Ifi204 |
| Wwtr1 | 7.22E-47 | 0.482602 | 0.772 | 0.472 | 1.45E-42 | 0 L1 _ | Wwtr1 |
| Rin2 | 4.50E-47 | 0.548286 | 0.597 | 0.306 | 9.05E-43 | 0 L1 _ | Rin2 |
| Ikbip1 | 4.32E-47 | 0.413051 | 0.672 | 0.345 | 8.70E-43 | 0 L1 _ | Ikbip |
| Cgnl1 | 3.80E-47 | 0.49747 | 0.421 | 0.164 | 7.65E-43 | 0 L1 _ | Cgnl1 |
| Tmem158 | 3.64E-47 | 0.487017 | 0.423 | 0.161 | 7.33E-43 | 0 L1 _ | Tmem158 |
| Fuca1 | 1.48E-47 | 0.458091 | 0.767 | 0.462 | 2.97E-43 | 0 L1 _ | Fuca1 |
| Tcf7l21 | 1.36E-47 | 0.532046 | 0.65 | 0.342 | 2.73E-43 | 0 L1 _ | Tcf7l2 |
| Sh3d191 | 1.27E-47 | 0.457119 | 0.6 | 0.284 | 2.56E-43 | 0 L1 _ | Sh3d19 |
| Ak1 | 1.08E-47 | 0.426743 | 0.315 | 0.087 | 2.18E-43 | 0 L1 _ | Ak1 |
| Fam20a | 1.00E-47 | 0.422065 | 0.444 | 0.174 | 2.02E-43 | 0 L1 _ | Fam20a |
| Ecm11 | 7.94E-48 | 0.400653 | 0.677 | 0.352 | 1.60E-43 | 0 L1 _ | Ecm1 |
| Gabarap | 5.32E-48 | 0.464473 | 0.942 | 0.847 | 1.07E-43 | 0 L1 _ | Gabarap |
| Grn | 3.92E-48 | 0.505742 | 0.861 | 0.562 | 7.89E-44 | 0 L1 _ | Grn |
| Tmem2041 | 3.64E-48 | 0.545822 | 0.773 | 0.448 | 7.34E-44 | 0 L1 _ | Tmem204 |
| Nfic1 | 1.22E-48 | 0.491767 | 0.895 | 0.596 | 2.47E-44 | 0 L1 _ | Nfic |
| Gm10115 | 9.85E-49 | 0.355361 | 0.236 | 0.038 | 1.98E-44 | 0 L1 _ | Gm10115 |
| Ptprd | 9.07E-49 | 0.469522 | 0.349 | 0.104 | 1.83E-44 | 0 L1 _ | Ptprd |
| Frem1 | 4.67E-49 | 0.395761 | 0.287 | 0.067 | 9.41E-45 | 0 L1 _ | Frem1 |
| Igfbp51 | 3.49E-49 | 0.815748 | 0.709 | 0.388 | 7.02E-45 | 0 L1 _ | Igfbp5 |
| Chp2 | 1.04E-49 | 0.564202 | 0.325 | 0.092 | 2.09E-45 | 0 L1 _ | Chp2 |
| Nucb21 | 1.02E-49 | 0.487981 | 0.594 | 0.285 | 2.05E-45 | 0 L1 _ | Nucb2 |
| Gng12 | 9.45E-50 | 0.499341 | 0.727 | 0.413 | 1.90E-45 | 0 L1 _ | Gng12 |

| | | | | | | | |
|---|---|---|---|---|---|---|---|
| Twist11 | 8.05E-50 | 0.437152 | 0.682 | 0.343 | 1.62E-45 | 0 L1 _ | Twist1 |
| Gm17167 | 4.89E-50 | 0.483071 | 0.272 | 0.055 | 9.84E-46 | 0 L1 _ | Gm17167 |
| Fat1 | 4.89E-50 | 0.452986 | 0.427 | 0.152 | 9.84E-46 | 0 L1 _ | Fat1 |
| Kdelr21 | 6.92E-51 | 0.520452 | 0.748 | 0.455 | 1.39E-46 | 0 L1 _ | Kdelr2 |
| Medag2 | 5.41E-51 | 0.454756 | 0.518 | 0.207 | 1.09E-46 | 0 L1 _ | Medag |
| Vcan1 | 5.08E-51 | 0.661805 | 0.481 | 0.19 | 1.02E-46 | 0 L1 _ | Vcan |
| Selenom1 | 4.60E-51 | 0.521262 | 0.952 | 0.735 | 9.27E-47 | 0 L1 _ | Selenom |
| Runx1 | 4.28E-51 | 0.56586 | 0.496 | 0.205 | 8.62E-47 | 0 L1 _ | Runx1 |
| Tpbg | 2.04E-51 | 0.418354 | 0.322 | 0.087 | 4.11E-47 | 0 L1 _ | Tpbg |
| P4hb1 | 1.43E-51 | 0.517861 | 0.869 | 0.577 | 2.89E-47 | 0 L1 _ | P4hb |
| Hmgn1 | 3.55E-52 | 0.543578 | 0.885 | 0.64 | 7.15E-48 | 0 L1 _ | Hmgn1 |
| Fkbp91 | 3.18E-52 | 0.48793 | 0.732 | 0.396 | 6.41E-48 | 0 L1 _ | Fkbp9 |
| Cdc42ep5 | 3.04E-52 | 0.482304 | 0.554 | 0.249 | 6.12E-48 | 0 L1 _ | Cdc42ep5 |
| Lima11 | 2.86E-52 | 0.523835 | 0.82 | 0.466 | 5.75E-48 | 0 L1 _ | Lima1 |
| Il341 | 2.43E-52 | 0.589682 | 0.697 | 0.362 | 4.89E-48 | 0 L1 _ | Il34 |
| Bpgm | 1.80E-52 | 0.828904 | 0.403 | 0.143 | 3.62E-48 | 0 L1 _ | Bpgm |
| Ar1 | 1.09E-52 | 0.579885 | 0.773 | 0.438 | 2.20E-48 | 0 L1 _ | Ar |
| Tmsb101 | 6.49E-53 | 0.433645 | 0.98 | 0.885 | 1.31E-48 | 0 L1 _ | Tmsb10 |
| mt-Cytb | 5.95E-53 | 0.332434 | 0.987 | 0.991 | 1.20E-48 | 0 L1 _ | mt-Cytb |
| Man1c1 | 3.03E-53 | 0.443535 | 0.437 | 0.161 | 6.10E-49 | 0 L1 _ | Man1c1 |
| Hspg22 | 3.01E-53 | 0.485016 | 0.93 | 0.541 | 6.06E-49 | 0 L1 _ | Hspg2 |
| Cstb | 1.41E-53 | 0.597694 | 0.761 | 0.463 | 2.85E-49 | 0 L1 _ | Cstb |
| Rhbdf1 | 1.24E-53 | 0.473296 | 0.536 | 0.231 | 2.51E-49 | 0 L1 _ | Rhbdf1 |
| Mmp11 | 1.21E-53 | 0.622683 | 0.519 | 0.216 | 2.44E-49 | 0 L1 _ | Mmp11 |
| Col12a1 | 1.10E-53 | 0.427737 | 0.333 | 0.083 | 2.21E-49 | 0 L1 _ | Col12a1 |
| Col8a2 | 8.68E-54 | 0.403236 | 0.317 | 0.074 | 1.75E-49 | 0 L1 _ | Col8a2 |
| Rasl11b | 5.11E-54 | 0.523711 | 0.367 | 0.105 | 1.03E-49 | 0 L1 _ | Rasl11b |
| Clmp1 | 2.52E-54 | 0.473427 | 0.486 | 0.18 | 5.07E-50 | 0 L1 _ | Clmp |
| Fap1 | 2.20E-54 | 0.462526 | 0.682 | 0.332 | 4.43E-50 | 0 L1 _ | Fap |
| Oat1 | 1.33E-54 | 0.518437 | 0.84 | 0.511 | 2.69E-50 | 0 L1 _ | Oat |
| Fmo11 | 8.37E-55 | 0.49606 | 0.704 | 0.348 | 1.69E-50 | 0 L1 _ | Fmo1 |
| Hspa5 | 6.99E-55 | 0.65556 | 0.956 | 0.831 | 1.41E-50 | 0 L1 _ | Hspa5 |
| Cxcl16 | 5.43E-55 | 0.38333 | 0.509 | 0.197 | 1.09E-50 | 0 L1 _ | Cxcl16 |
| Clec11a1 | 5.14E-55 | 0.601881 | 0.408 | 0.133 | 1.04E-50 | 0 L1 _ | Clec11a |
| Scd11 | 4.44E-55 | 0.403307 | 0.505 | 0.192 | 8.94E-51 | 0 L1 _ | Scd1 |
| Abca91 | 3.16E-55 | 0.508778 | 0.51 | 0.201 | 6.35E-51 | 0 L1 _ | Abca9 |
| Egfr2 | 2.44E-55 | 0.475827 | 0.498 | 0.187 | 4.92E-51 | 0 L1 _ | Egfr |
| Erlec1 | 1.60E-55 | 0.517357 | 0.689 | 0.36 | 3.21E-51 | 0 L1 _ | Erlec1 |
| Setbp1 | 1.29E-55 | 0.541783 | 0.516 | 0.211 | 2.59E-51 | 0 L1 _ | Setbp1 |
| Gria3 | 1.01E-55 | 0.44296 | 0.372 | 0.107 | 2.04E-51 | 0 L1 _ | Gria3 |
| Pdia61 | 9.43E-56 | 0.580587 | 0.88 | 0.641 | 1.90E-51 | 0 L1 _ | Pdia6 |

| | | | | | | | |
|---|---|---|---|---|---|---|---|
| Ssr2 | 9.41E-56 | 0.537197 | 0.811 | 0.53 | 1.90E-51 | 0 L1 _ | Ssr2 |
| Fgf71 | 5.86E-56 | 0.522212 | 0.408 | 0.133 | 1.18E-51 | 0 L1 _ | Fgf7 |
| Tgfbr22 | 4.70E-56 | 0.563055 | 0.837 | 0.528 | 9.47E-52 | 0 L1 _ | Tgfbr2 |
| Ptgis1 | 3.63E-56 | 0.53406 | 0.426 | 0.141 | 7.31E-52 | 0 L1 _ | Ptgis |
| Msc | 2.82E-56 | 0.448791 | 0.331 | 0.08 | 5.69E-52 | 0 L1 _ | Msc |
| Gm15867 | 2.00E-56 | 0.371049 | 0.313 | 0.068 | 4.03E-52 | 0 L1 _ | Gm15867 |
| Prss12 | 1.36E-56 | 0.448344 | 0.329 | 0.078 | 2.73E-52 | 0 L1 _ | Prss12 |
| Lrrc32 | 1.22E-56 | 0.610782 | 0.576 | 0.258 | 2.45E-52 | 0 L1 _ | Lrrc32 |
| Cyth31 | 9.69E-57 | 0.592454 | 0.864 | 0.54 | 1.95E-52 | 0 L1 _ | Cyth3 |
| Rora1 | 4.44E-57 | 0.592926 | 0.731 | 0.389 | 8.95E-53 | 0 L1 _ | Rora |
| Pdzrn3 | 4.34E-57 | 0.468839 | 0.429 | 0.142 | 8.74E-53 | 0 L1 _ | Pdzrn3 |
| Tceal8 | 4.08E-57 | 0.494587 | 0.642 | 0.314 | 8.22E-53 | 0 L1 _ | Tceal8 |
| Wdr66 | 2.85E-57 | 0.571711 | 0.404 | 0.128 | 5.75E-53 | 0 L1 _ | Wdr66 |
| Nenf | 2.20E-57 | 0.539449 | 0.926 | 0.671 | 4.44E-53 | 0 L1 _ | Nenf |
| Sec31a | 1.62E-57 | 0.550871 | 0.793 | 0.476 | 3.26E-53 | 0 L1 _ | Sec31a |
| Ckap41 | 1.33E-57 | 0.585683 | 0.727 | 0.386 | 2.68E-53 | 0 L1 _ | Ckap4 |
| Cx3cl1 | 1.29E-57 | 0.552269 | 0.582 | 0.258 | 2.60E-53 | 0 L1 _ | Cx3cl1 |
| Socs21 | 8.93E-58 | 0.529257 | 0.698 | 0.36 | 1.80E-53 | 0 L1 _ | Socs2 |
| Fam114a11 | 5.53E-58 | 0.518787 | 0.668 | 0.327 | 1.11E-53 | 0 L1 _ | Fam114a1 |
| Selenos1 | 5.50E-58 | 0.609309 | 0.83 | 0.53 | 1.11E-53 | 0 L1 _ | Selenos |
| Rbms31 | 4.56E-58 | 0.439394 | 0.815 | 0.408 | 9.18E-54 | 0 L1 _ | Rbms3 |
| Bhlhe41 | 4.15E-58 | 0.560627 | 0.357 | 0.093 | 8.35E-54 | 0 L1 _ | Bhlhe41 |
| F2r | 3.08E-58 | 0.742139 | 0.551 | 0.249 | 6.20E-54 | 0 L1 _ | F2r |
| Basp12 | 2.84E-58 | 0.411069 | 0.586 | 0.235 | 5.72E-54 | 0 L1 _ | Basp1 |
| Col27a11 | 2.35E-58 | 0.553833 | 0.53 | 0.206 | 4.72E-54 | 0 L1 _ | Col27a1 |
| Pdlim4 | 8.46E-59 | 0.47175 | 0.44 | 0.147 | 1.70E-54 | 0 L1 _ | Pdlim4 |
| Ctsz | 5.34E-59 | 0.489619 | 0.909 | 0.658 | 1.08E-54 | 0 L1 _ | Ctsz |
| Ccn5 | 4.21E-59 | 1.34691 | 0.257 | 0.033 | 8.48E-55 | 0 L1 _ | Ccn5 |
| Crtap | 3.98E-59 | 0.504133 | 0.699 | 0.351 | 8.01E-55 | 0 L1 _ | Crtap |
| Htra11 | 9.14E-60 | 0.604275 | 0.875 | 0.503 | 1.84E-55 | 0 L1 _ | Htra1 |
| Pi161 | 4.19E-60 | 0.49353 | 0.45 | 0.144 | 8.43E-56 | 0 L1 _ | Pi16 |
| Fam180a | 4.17E-60 | 0.630666 | 0.291 | 0.05 | 8.40E-56 | 0 L1 _ | Fam180a |
| Ctsb1 | 2.74E-60 | 0.457834 | 0.933 | 0.667 | 5.52E-56 | 0 L1 _ | Ctsb |
| Lrrk1 | 2.27E-60 | 0.599504 | 0.624 | 0.297 | 4.58E-56 | 0 L1 _ | Lrrk1 |
| Plac81 | 3.69E-61 | 0.861323 | 0.567 | 0.229 | 7.43E-57 | 0 L1 _ | Plac8 |
| Rap2b | 2.64E-61 | 0.531105 | 0.563 | 0.24 | 5.31E-57 | 0 L1 _ | Rap2b |
| Stimate | 1.61E-61 | 0.378006 | 0.377 | 0.094 | 3.24E-57 | 0 L1 _ | Stimate |
| Ank21 | 1.35E-61 | 0.58673 | 0.53 | 0.199 | 2.72E-57 | 0 L1 _ | Ank2 |
| Svil1 | 8.90E-62 | 0.635492 | 0.713 | 0.359 | 1.79E-57 | 0 L1 _ | Svil |
| Plau1 | 7.23E-62 | 0.569043 | 0.521 | 0.194 | 1.46E-57 | 0 L1 _ | Plau |
| Tgfb3 | 7.00E-62 | 0.532183 | 0.487 | 0.168 | 1.41E-57 | 0 L1 _ | Tgfb3 |

| | | | | | | | |
|---|---|---|---|---|---|---|---|
| Ergic3 | 3.60E-62 | 0.595992 | 0.809 | 0.488 | 7.25E-58 | 0 L1 _ | Ergic3 |
| Snhg18 | 1.50E-62 | 0.553893 | 0.831 | 0.484 | 3.02E-58 | 0 L1 _ | Snhg18 |
| Hdlbp1 | 1.48E-62 | 0.63946 | 0.865 | 0.588 | 2.98E-58 | 0 L1 _ | Hdlbp |
| Col4a5 | 1.47E-62 | 0.642708 | 0.454 | 0.151 | 2.97E-58 | 0 L1 _ | Col4a5 |
| Cp1 | 1.44E-62 | 0.667959 | 0.845 | 0.436 | 2.89E-58 | 0 L1 _ | Cp |
| Pros1 | 1.39E-62 | 0.569805 | 0.769 | 0.423 | 2.80E-58 | 0 L1 _ | Pros1 |
| Shc4 | 1.23E-62 | 0.508597 | 0.336 | 0.072 | 2.48E-58 | 0 L1 _ | Shc4 |
| Copz2 | 1.07E-62 | 0.546996 | 0.668 | 0.319 | 2.15E-58 | 0 L1 _ | Copz2 |
| Cnpy2 | 8.39E-63 | 0.559167 | 0.854 | 0.542 | 1.69E-58 | 0 L1 _ | Cnpy2 |
| Lrpap1 | 6.34E-63 | 0.577199 | 0.818 | 0.484 | 1.28E-58 | 0 L1 _ | Lrpap1 |
| Gas61 | 6.34E-63 | 0.637658 | 0.795 | 0.417 | 1.28E-58 | 0 L1 _ | Gas6 |
| Cmklr11 | 5.93E-63 | 0.524459 | 0.496 | 0.174 | 1.20E-58 | 0 L1 _ | Cmklr1 |
| Efemp12 | 3.72E-63 | 0.4159 | 0.577 | 0.207 | 7.50E-59 | 0 L1 _ | Efemp1 |
| Dnm3os | 2.16E-63 | 0.688621 | 0.491 | 0.172 | 4.35E-59 | 0 L1 _ | Dnm3os |
| S100a161 | 1.96E-63 | 0.660339 | 0.806 | 0.455 | 3.94E-59 | 0 L1 _ | S100a16 |
| Eva1b | 1.62E-63 | 0.601085 | 0.893 | 0.602 | 3.26E-59 | 0 L1 _ | Eva1b |
| Rhobtb31 | 1.27E-63 | 0.539326 | 0.584 | 0.239 | 2.56E-59 | 0 L1 _ | Rhobtb3 |
| Prrx11 | 9.39E-64 | 0.634169 | 0.934 | 0.572 | 1.89E-59 | 0 L1 _ | Prrx1 |
| Gsto1 | 4.83E-64 | 0.547508 | 0.463 | 0.155 | 9.73E-60 | 0 L1 _ | Gsto1 |
| Creb3l11 | 2.21E-64 | 0.551878 | 0.56 | 0.215 | 4.44E-60 | 0 L1 _ | Creb3l1 |
| Gadd45a1 | 1.68E-64 | 0.655646 | 0.494 | 0.177 | 3.38E-60 | 0 L1 _ | Gadd45a |
| Npdc1 | 1.29E-64 | 0.604716 | 0.803 | 0.45 | 2.60E-60 | 0 L1 _ | Npdc1 |
| Snap47 | 2.21E-65 | 0.566403 | 0.528 | 0.2 | 4.44E-61 | 0 L1 _ | Snap47 |
| Gpx81 | 1.43E-65 | 0.460618 | 0.869 | 0.505 | 2.89E-61 | 0 L1 _ | Gpx8 |
| Gfpt22 | 8.60E-66 | 0.525974 | 0.523 | 0.185 | 1.73E-61 | 0 L1 _ | Gfpt2 |
| Mgst12 | 7.75E-66 | 0.515019 | 0.742 | 0.338 | 1.56E-61 | 0 L1 _ | Mgst1 |
| Gpx7 | 6.27E-66 | 0.547088 | 0.543 | 0.207 | 1.26E-61 | 0 L1 _ | Gpx7 |
| Mfap2 | 5.34E-66 | 0.654532 | 0.693 | 0.332 | 1.07E-61 | 0 L1 _ | Mfap2 |
| Adamtsl11 | 3.75E-66 | 0.639448 | 0.547 | 0.209 | 7.55E-62 | 0 L1 _ | Adamtsl1 |
| Fmo21 | 3.56E-66 | 0.863983 | 0.684 | 0.318 | 7.17E-62 | 0 L1 _ | Fmo2 |
| Rab34 | 3.32E-66 | 0.532986 | 0.597 | 0.246 | 6.68E-62 | 0 L1 _ | Rab34 |
| Capg | 2.58E-66 | 0.729235 | 0.768 | 0.446 | 5.20E-62 | 0 L1 _ | Capg |
| Atf51 | 5.37E-67 | 0.651218 | 0.718 | 0.345 | 1.08E-62 | 0 L1 _ | Atf5 |
| Ifi2111 | 3.73E-67 | 0.577348 | 0.571 | 0.216 | 7.51E-63 | 0 L1 _ | Ifi211 |
| Hacd4 | 3.46E-67 | 0.683432 | 0.546 | 0.211 | 6.97E-63 | 0 L1 _ | Hacd4 |
| Gstt11 | 2.58E-67 | 0.669783 | 0.615 | 0.259 | 5.21E-63 | 0 L1 _ | Gstt1 |
| Ppib | 1.64E-68 | 0.508968 | 0.964 | 0.882 | 3.30E-64 | 0 L1 _ | Ppib |
| C32 | 1.18E-68 | 0.578528 | 0.629 | 0.242 | 2.38E-64 | 0 L1 _ | C3 |
| Gas11 | 7.88E-69 | 0.599676 | 0.879 | 0.441 | 1.59E-64 | 0 L1 _ | Gas1 |
| Slc40a1 | 3.74E-69 | 0.585512 | 0.378 | 0.083 | 7.53E-65 | 0 L1 _ | Slc40a1 |
| Adra2c | 1.97E-69 | 0.919473 | 0.332 | 0.061 | 3.96E-65 | 0 L1 _ | Adra2c |

| | | | | | | | |
|---|---|---|---|---|---|---|---|
| Ubtd2 | 1.30E-69 | 0.541248 | 0.446 | 0.131 | 2.63E-65 | 0 L1 _ | Ubtd2 |
| Spry11 | 4.21E-70 | 0.691256 | 0.765 | 0.383 | 8.48E-66 | 0 L1 _ | Spry1 |
| Ndn1 | 1.91E-70 | 0.556773 | 0.709 | 0.308 | 3.85E-66 | 0 L1 _ | Ndn |
| Gas51 | 1.65E-70 | 0.678147 | 0.926 | 0.571 | 3.32E-66 | 0 L1 _ | Gas5 |
| Ctsf | 1.48E-70 | 0.554836 | 0.548 | 0.202 | 2.98E-66 | 0 L1 _ | Ctsf |
| C1s11 | 1.29E-70 | 0.616343 | 0.955 | 0.562 | 2.60E-66 | 0 L1 _ | C1s1 |
| Sulf1 | 4.44E-71 | 0.622966 | 0.558 | 0.208 | 8.93E-67 | 0 L1 _ | Sulf1 |
| Dchs2 | 2.91E-71 | 0.627288 | 0.412 | 0.104 | 5.86E-67 | 0 L1 _ | Dchs2 |
| Pdlim2 | 2.31E-71 | 0.702984 | 0.783 | 0.435 | 4.66E-67 | 0 L1 _ | Pdlim2 |
| Lbp1 | 1.36E-71 | 0.58729 | 0.473 | 0.146 | 2.74E-67 | 0 L1 _ | Lbp |
| Inmt1 | 1.19E-71 | 1.050032 | 0.508 | 0.165 | 2.40E-67 | 0 L1 _ | Inmt |
| Txndc51 | 9.06E-72 | 0.625512 | 0.662 | 0.309 | 1.82E-67 | 0 L1 _ | Txndc5 |
| Olfml1 | 5.37E-72 | 0.556406 | 0.436 | 0.117 | 1.08E-67 | 0 L1 _ | Olfml1 |
| Bmp12 | 1.91E-72 | 0.581753 | 0.675 | 0.284 | 3.86E-68 | 0 L1 _ | Bmp1 |
| Maged11 | 1.87E-72 | 0.644648 | 0.777 | 0.405 | 3.76E-68 | 0 L1 _ | Maged1 |
| Plekha61 | 3.38E-73 | 0.625277 | 0.582 | 0.216 | 6.80E-69 | 0 L1 _ | Plekha6 |
| Mt22 | 2.02E-73 | 0.93679 | 0.557 | 0.2 | 4.07E-69 | 0 L1 _ | Mt2 |
| Vim1 | 1.80E-73 | 0.629265 | 0.967 | 0.828 | 3.62E-69 | 0 L1 _ | Vim |
| F31 | 1.09E-73 | 0.645312 | 0.488 | 0.146 | 2.19E-69 | 0 L1 _ | F3 |
| Omd | 4.37E-74 | 0.610391 | 0.38 | 0.081 | 8.80E-70 | 0 L1 _ | Omd |
| Steap31 | 4.11E-74 | 0.552103 | 0.593 | 0.207 | 8.28E-70 | 0 L1 _ | Steap3 |
| Fmod | 3.83E-74 | 0.640824 | 0.339 | 0.055 | 7.72E-70 | 0 L1 _ | Fmod |
| B3gnt9 | 2.17E-74 | 0.558786 | 0.471 | 0.143 | 4.36E-70 | 0 L1 _ | B3gnt9 |
| Adamtsl21 | 5.82E-75 | 0.735505 | 0.511 | 0.16 | 1.17E-70 | 0 L1 _ | Adamtsl2 |
| Nav31 | 4.84E-75 | 0.664015 | 0.558 | 0.2 | 9.76E-71 | 0 L1 _ | Nav3 |
| 1810058I24Rik | 4.26E-75 | 0.741603 | 0.856 | 0.563 | 8.57E-71 | 0 L1 _ | 1810058I24Rik |
| Pdia3 | 3.82E-75 | 0.645103 | 0.961 | 0.799 | 7.69E-71 | 0 L1 _ | Pdia3 |
| Auts21 | 1.24E-75 | 0.650864 | 0.725 | 0.327 | 2.50E-71 | 0 L1 _ | Auts2 |
| Zfhx41 | 1.19E-75 | 0.645501 | 0.647 | 0.253 | 2.40E-71 | 0 L1 _ | Zfhx4 |
| Oaf1 | 2.75E-76 | 0.685813 | 0.75 | 0.336 | 5.55E-72 | 0 L1 _ | Oaf |
| Adcy7 | 6.12E-77 | 0.67849 | 0.662 | 0.28 | 1.23E-72 | 0 L1 _ | Adcy7 |
| Ctsh1 | 4.54E-77 | 0.706061 | 0.832 | 0.394 | 9.14E-73 | 0 L1 _ | Ctsh |
| Cyp1b11 | 3.10E-77 | 0.68319 | 0.567 | 0.191 | 6.23E-73 | 0 L1 _ | Cyp1b1 |
| Creg1 | 2.36E-77 | 0.696508 | 0.782 | 0.428 | 4.75E-73 | 0 L1 _ | Creg1 |
| Mrc21 | 1.19E-77 | 0.598944 | 0.668 | 0.26 | 2.40E-73 | 0 L1 _ | Mrc2 |
| Calr1 | 6.45E-78 | 0.643308 | 0.966 | 0.812 | 1.30E-73 | 0 L1 _ | Calr |
| Arap11 | 1.49E-78 | 0.674895 | 0.636 | 0.267 | 3.00E-74 | 0 L1 _ | Arap1 |
| Scara52 | 1.27E-78 | 0.608516 | 0.665 | 0.235 | 2.55E-74 | 0 L1 _ | Scara5 |
| Lbhd2 | 6.52E-79 | 0.771685 | 0.336 | 0.047 | 1.31E-74 | 0 L1 _ | Lbhd2 |
| Scarf21 | 6.03E-79 | 0.624274 | 0.689 | 0.288 | 1.21E-74 | 0 L1 _ | Scarf2 |
| Lrrn2 | 4.80E-79 | 0.602445 | 0.473 | 0.131 | 9.66E-75 | 0 L1 _ | Lrrn2 |

| | | | | | | | |
|---|---|---|---|---|---|---|---|
| Barx1 | 3.16E-79 | 0.672303 | 0.829 | 0.414 | 6.37E-75 | 0 L1 _ | Barx1 |
| Fcgrt1 | 2.20E-79 | 0.711178 | 0.914 | 0.584 | 4.44E-75 | 0 L1 _ | Fcgrt |
| Gaa1 | 4.80E-80 | 0.624923 | 0.743 | 0.349 | 9.67E-76 | 0 L1 _ | Gaa |
| Chpf1 | 3.73E-80 | 0.67428 | 0.714 | 0.315 | 7.50E-76 | 0 L1 _ | Chpf |
| Ftl11 | 1.08E-80 | 0.538984 | 0.982 | 0.95 | 2.17E-76 | 0 L1 _ | Ftl1 |
| P3h31 | 5.00E-81 | 0.664944 | 0.739 | 0.34 | 1.01E-76 | 0 L1 _ | P3h3 |
| Slit2 | 2.29E-81 | 0.593326 | 0.438 | 0.097 | 4.61E-77 | 0 L1 _ | Slit2 |
| Lamp21 | 1.05E-81 | 0.702922 | 0.934 | 0.632 | 2.11E-77 | 0 L1 _ | Lamp2 |
| Ppic | 1.43E-82 | 0.704719 | 0.808 | 0.438 | 2.89E-78 | 0 L1 _ | Ppic |
| Kdelr31 | 8.72E-83 | 0.644593 | 0.632 | 0.238 | 1.76E-78 | 0 L1 _ | Kdelr3 |
| Col6a61 | 3.17E-83 | 0.759847 | 0.547 | 0.169 | 6.38E-79 | 0 L1 _ | Col6a6 |
| Tenm3 | 1.46E-83 | 0.647462 | 0.492 | 0.134 | 2.93E-79 | 0 L1 _ | Tenm3 |
| Gstm21 | 1.41E-83 | 0.661399 | 0.738 | 0.317 | 2.85E-79 | 0 L1 _ | Gstm2 |
| Fzd2 | 9.00E-84 | 0.70641 | 0.516 | 0.154 | 1.81E-79 | 0 L1 _ | Fzd2 |
| Efemp2 | 3.14E-84 | 0.677274 | 0.717 | 0.314 | 6.33E-80 | 0 L1 _ | Efemp2 |
| Eid1 | 1.15E-84 | 0.77603 | 0.924 | 0.691 | 2.32E-80 | 0 L1 _ | Eid1 |
| Gpr1531 | 9.56E-85 | 0.673637 | 0.623 | 0.223 | 1.93E-80 | 0 L1 _ | Gpr153 |
| Nme2 | 6.05E-85 | 0.775946 | 0.952 | 0.661 | 1.22E-80 | 0 L1 _ | Nme2 |
| Vgll3 | 1.31E-85 | 0.690389 | 0.54 | 0.159 | 2.65E-81 | 0 L1 _ | Vgll3 |
| Tmem1191 | 8.88E-86 | 0.643282 | 0.63 | 0.218 | 1.79E-81 | 0 L1 _ | Tmem119 |
| Serf1 | 6.69E-86 | 0.717612 | 0.585 | 0.208 | 1.35E-81 | 0 L1 _ | Serf1 |
| Phlda31 | 1.44E-86 | 0.82148 | 0.789 | 0.397 | 2.90E-82 | 0 L1 _ | Phlda3 |
| Marcks1 | 1.23E-86 | 0.786908 | 0.975 | 0.721 | 2.48E-82 | 0 L1 _ | Marcks |
| Lamb11 | 1.07E-86 | 0.74018 | 0.918 | 0.48 | 2.16E-82 | 0 L1 _ | Lamb1 |
| Snx9 | 4.06E-87 | 0.727385 | 0.741 | 0.348 | 8.18E-83 | 0 L1 _ | Snx9 |
| Ccbe1 | 2.78E-87 | 0.655706 | 0.434 | 0.094 | 5.59E-83 | 0 L1 _ | Ccbe1 |
| Ltbp2 | 4.12E-88 | 1.036489 | 0.419 | 0.076 | 8.29E-84 | 0 L1 _ | Ltbp2 |
| Fhl2 | 9.57E-89 | 0.775438 | 0.629 | 0.232 | 1.93E-84 | 0 L1 _ | Fhl2 |
| Adk | 1.07E-89 | 0.689955 | 0.612 | 0.218 | 2.16E-85 | 0 L1 _ | Adk |
| Vkorc1 | 4.33E-90 | 0.75413 | 0.835 | 0.451 | 8.72E-86 | 0 L1 _ | Vkorc1 |
| Prrx21 | 2.02E-91 | 0.676642 | 0.554 | 0.164 | 4.07E-87 | 0 L1 _ | Prrx2 |
| Thbs4 | 1.95E-91 | 2.246591 | 0.464 | 0.102 | 3.93E-87 | 0 L1 _ | Thbs4 |
| Aldh1a11 | 2.62E-92 | 0.901601 | 0.582 | 0.188 | 5.27E-88 | 0 L1 _ | Aldh1a1 |
| Ltbp31 | 2.53E-93 | 0.736957 | 0.687 | 0.275 | 5.10E-89 | 0 L1 _ | Ltbp3 |
| Nbl1 | 1.28E-93 | 1.148849 | 0.878 | 0.498 | 2.57E-89 | 0 L1 _ | Nbl1 |
| Islr1 | 6.45E-94 | 0.717973 | 0.747 | 0.285 | 1.30E-89 | 0 L1 _ | Islr |
| Col5a12 | 2.58E-94 | 0.81985 | 0.797 | 0.323 | 5.21E-90 | 0 L1 _ | Col5a1 |
| Lpar11 | 1.19E-94 | 0.683374 | 0.684 | 0.234 | 2.40E-90 | 0 L1 _ | Lpar1 |
| Col16a1 | 9.16E-95 | 0.744286 | 0.636 | 0.225 | 1.84E-90 | 0 L1 _ | Col16a1 |
| Tspan111 | 1.08E-95 | 0.718336 | 0.647 | 0.213 | 2.18E-91 | 0 L1 _ | Tspan11 |
| Ptgfrn1 | 3.72E-96 | 0.69828 | 0.658 | 0.234 | 7.49E-92 | 0 L1 _ | Ptgfrn |

| | | | | | | | |
|---|---|---|---|---|---|---|---|
| Igsf101 | 3.34E-96 | 0.793971 | 0.761 | 0.296 | 6.73E-92 | 0 L1 _ | Igsf10 |
| Ank | 2.57E-96 | 0.801368 | 0.574 | 0.184 | 5.18E-92 | 0 L1 _ | Ank |
| Mt11 | 1.77E-96 | 1.125912 | 0.868 | 0.459 | 3.57E-92 | 0 L1 _ | Mt1 |
| Nav11 | 1.21E-96 | 0.864728 | 0.891 | 0.435 | 2.44E-92 | 0 L1 _ | Nav1 |
| Hexa | 4.26E-97 | 0.790928 | 0.84 | 0.451 | 8.59E-93 | 0 L1 _ | Hexa |
| Clec3b2 | 2.11E-97 | 0.663031 | 0.727 | 0.239 | 4.25E-93 | 0 L1 _ | Clec3b |
| Sdc21 | 2.07E-97 | 0.809182 | 0.827 | 0.382 | 4.17E-93 | 0 L1 _ | Sdc2 |
| Col14a12 | 7.38E-99 | 0.709951 | 0.752 | 0.26 | 1.49E-94 | 0 L1 _ | Col14a1 |
| Fgfr11 | 5.35E-99 | 0.727153 | 0.764 | 0.304 | 1.08E-94 | 0 L1 _ | Fgfr1 |
| Cpxm2 | 2.68E-99 | 1.35466 | 0.541 | 0.147 | 5.40E-95 | 0 L1 _ | Cpxm2 |
| Flrt21 | 1.71E-99 | 0.914739 | 0.626 | 0.213 | 3.45E-95 | 0 L1 _ | Flrt2 |
| Lsamp1 | 1.05E-99 | 0.761365 | 0.793 | 0.306 | 2.12E-95 | 0 L1 _ | Lsamp |
| Pla1a | 8.93E-100 | 0.871566 | 0.684 | 0.236 | 1.80E-95 | 0 L1 _ | Pla1a |
| Thbs2 | 8.14E-101 | 0.795729 | 0.63 | 0.206 | 1.64E-96 | 0 L1 _ | Thbs2 |
| Selenop1 | 5.24E-101 | 0.918764 | 0.961 | 0.632 | 1.06E-96 | 0 L1 _ | Selenop |
| Tent5a | 4.18E-101 | 0.88937 | 0.672 | 0.229 | 8.42E-97 | 0 L1 _ | Tent5a |
| Ckb | 1.63E-101 | 1.114994 | 0.852 | 0.49 | 3.28E-97 | 0 L1 _ | Ckb |
| Tuba1a1 | 4.33E-102 | 0.870055 | 0.964 | 0.738 | 8.71E-98 | 0 L1 _ | Tuba1a |
| Mmp141 | 1.43E-102 | 0.829303 | 0.824 | 0.348 | 2.89E-98 | 0 L1 _ | Mmp14 |
| Vcam1 | 1.42E-102 | 0.834692 | 0.573 | 0.159 | 2.86E-98 | 0 L1 _ | Vcam1 |
| Ifitm2 | 2.78E-103 | 0.756229 | 0.976 | 0.833 | 5.60E-99 | 0 L1 _ | Ifitm2 |
| Pcdh71 | 1.87E-103 | 0.808696 | 0.633 | 0.2 | 3.76E-99 | 0 L1 _ | Pcdh7 |
| Nid12 | 1.10E-103 | 0.768597 | 0.957 | 0.583 | 2.21E-99 | 0 L1 _ | Nid1 |
| Fbn12 | 3.40E-104 | 0.786002 | 0.888 | 0.441 | 6.84E-100 | 0 L1 _ | Fbn1 |
| Ahnak2 | 1.42E-104 | 0.824469 | 0.611 | 0.177 | 2.85E-100 | 0 L1 _ | Ahnak2 |
| Tmed31 | 1.30E-104 | 0.867406 | 0.88 | 0.531 | 2.61E-100 | 0 L1 _ | Tmed3 |
| Ms4a4d1 | 9.59E-105 | 0.809893 | 0.736 | 0.253 | 1.93E-100 | 0 L1 _ | Ms4a4d |
| Fbn2 | 8.02E-105 | 0.885648 | 0.51 | 0.11 | 1.62E-100 | 0 L1 _ | Fbn2 |
| Emilin1 | 4.28E-105 | 0.831476 | 0.76 | 0.311 | 8.62E-101 | 0 L1 _ | Emilin1 |
| Prss23 | 2.38E-105 | 1.054602 | 0.933 | 0.578 | 4.80E-101 | 0 L1 _ | Prss23 |
| Mmp31 | 7.62E-106 | 1.755535 | 0.609 | 0.176 | 1.53E-101 | 0 L1 _ | Mmp3 |
| Fkbp71 | 6.04E-106 | 0.765686 | 0.842 | 0.381 | 1.22E-101 | 0 L1 _ | Fkbp7 |
| Col5a31 | 5.28E-106 | 0.796803 | 0.859 | 0.365 | 1.06E-101 | 0 L1 _ | Col5a3 |
| Wls | 6.96E-107 | 0.889509 | 0.884 | 0.5 | 1.40E-102 | 0 L1 _ | Wls |
| Sfrp1 | 4.80E-108 | 1.041802 | 0.617 | 0.175 | 9.66E-104 | 0 L1 _ | Sfrp1 |
| Nfix1 | 1.57E-108 | 0.84602 | 0.947 | 0.513 | 3.16E-104 | 0 L1 _ | Nfix |
| Lhfp | 5.97E-109 | 0.85442 | 0.926 | 0.508 | 1.20E-104 | 0 L1 _ | Lhfp |
| Cd91 | 2.75E-109 | 0.879345 | 0.937 | 0.58 | 5.54E-105 | 0 L1 _ | Cd9 |
| Rrbp11 | 1.01E-111 | 0.828402 | 0.974 | 0.823 | 2.04E-107 | 0 L1 _ | Rrbp1 |
| Pdpn1 | 4.58E-114 | 0.834582 | 0.652 | 0.195 | 9.22E-110 | 0 L1 _ | Pdpn |
| Nkd2 | 3.98E-114 | 0.930401 | 0.589 | 0.143 | 8.01E-110 | 0 L1 _ | Nkd2 |

| | | | | | | | |
|---|---|---|---|---|---|---|---|
| Scpep11 | 2.71E-114 | 0.839711 | 0.738 | 0.278 | 5.45E-110 | 0 L1 _ | Scpep1 |
| Cped11 | 1.59E-115 | 1.004832 | 0.946 | 0.53 | 3.21E-111 | 0 L1 _ | Cped1 |
| Srpx21 | 5.65E-117 | 0.889295 | 0.64 | 0.189 | 1.14E-112 | 0 L1 _ | Srpx2 |
| Enpp1 | 1.33E-117 | 0.989238 | 0.646 | 0.193 | 2.68E-113 | 0 L1 _ | Enpp1 |
| Srpx1 | 1.37E-118 | 0.671195 | 0.792 | 0.256 | 2.76E-114 | 0 L1 _ | Srpx |
| Igf11 | 2.44E-119 | 0.937027 | 0.82 | 0.297 | 4.91E-115 | 0 L1 _ | Igf1 |
| Abca8a1 | 1.64E-119 | 0.898219 | 0.85 | 0.313 | 3.30E-115 | 0 L1 _ | Abca8a |
| Fth11 | 3.93E-120 | 0.832326 | 0.994 | 0.992 | 7.92E-116 | 0 L1 _ | Fth1 |
| Tgfbi1 | 6.01E-121 | 1.100849 | 0.729 | 0.237 | 1.21E-116 | 0 L1 _ | Tgfbi |
| Pla2g4a | 1.26E-121 | 0.862246 | 0.655 | 0.19 | 2.53E-117 | 0 L1 _ | Pla2g4a |
| Ccn4 | 3.39E-122 | 0.930507 | 0.518 | 0.093 | 6.83E-118 | 0 L1 _ | Ccn4 |
| C1ra1 | 3.79E-125 | 0.96479 | 0.86 | 0.342 | 7.64E-121 | 0 L1 _ | C1ra |
| Il331 | 9.78E-126 | 0.9606 | 0.757 | 0.229 | 1.97E-121 | 0 L1 _ | Il33 |
| Plat | 3.39E-126 | 0.917989 | 0.823 | 0.324 | 6.83E-122 | 0 L1 _ | Plat |
| Cpq1 | 3.38E-126 | 0.878146 | 0.75 | 0.264 | 6.80E-122 | 0 L1 _ | Cpq |
| Col15a11 | 2.17E-126 | 1.085166 | 0.938 | 0.428 | 4.38E-122 | 0 L1 _ | Col15a1 |
| Dpep11 | 1.43E-127 | 0.948814 | 0.849 | 0.271 | 2.87E-123 | 0 L1 _ | Dpep1 |
| Mir100hg | 1.32E-127 | 0.890357 | 0.72 | 0.2 | 2.66E-123 | 0 L1 _ | Mir100hg |
| Fzd1 | 8.16E-129 | 1.03609 | 0.691 | 0.216 | 1.64E-124 | 0 L1 _ | Fzd1 |
| Sparc1 | 4.19E-129 | 1.081525 | 0.991 | 0.877 | 8.44E-125 | 0 L1 _ | Sparc |
| Crispld1 | 1.87E-130 | 1.161827 | 0.768 | 0.272 | 3.77E-126 | 0 L1 _ | Crispld1 |
| Mafb1 | 1.51E-130 | 1.164671 | 0.883 | 0.389 | 3.03E-126 | 0 L1 _ | Mafb |
| Timp22 | 4.37E-131 | 0.79585 | 0.962 | 0.531 | 8.80E-127 | 0 L1 _ | Timp2 |
| Ifi30 | 1.24E-131 | 0.966826 | 0.699 | 0.196 | 2.50E-127 | 0 L1 _ | Ifi30 |
| Kcnk2 | 1.08E-131 | 0.970619 | 0.653 | 0.167 | 2.18E-127 | 0 L1 _ | Kcnk2 |
| Spats2l1 | 5.82E-132 | 1.026578 | 0.785 | 0.294 | 1.17E-127 | 0 L1 _ | Spats2l |
| Ccn2 | 5.33E-132 | 1.515107 | 0.655 | 0.159 | 1.07E-127 | 0 L1 _ | Ccn2 |
| Cygb | 3.85E-132 | 1.048926 | 0.979 | 0.614 | 7.75E-128 | 0 L1 _ | Cygb |
| Mtch1 | 1.38E-132 | 1.086124 | 0.923 | 0.63 | 2.78E-128 | 0 L1 _ | Mtch1 |
| Rnase42 | 2.80E-134 | 0.904407 | 0.938 | 0.373 | 5.63E-130 | 0 L1 _ | Rnase4 |
| Cd3021 | 1.89E-135 | 1.046942 | 0.925 | 0.417 | 3.80E-131 | 0 L1 _ | Cd302 |
| Mmp21 | 6.59E-136 | 0.898987 | 0.926 | 0.314 | 1.33E-131 | 0 L1 _ | Mmp2 |
| Lgals1 | 9.69E-138 | 1.217545 | 0.974 | 0.708 | 1.95E-133 | 0 L1 _ | Lgals1 |
| Itih51 | 7.06E-138 | 1.12874 | 0.95 | 0.492 | 1.42E-133 | 0 L1 _ | Itih5 |
| S100a61 | 4.73E-138 | 1.062297 | 0.989 | 0.887 | 9.53E-134 | 0 L1 _ | S100a6 |
| Crlf11 | 1.42E-138 | 1.236727 | 0.725 | 0.213 | 2.87E-134 | 0 L1 _ | Crlf1 |
| C1qtnf2 | 3.61E-140 | 1.02657 | 0.869 | 0.371 | 7.27E-136 | 0 L1 _ | C1qtnf2 |
| Serpinh11 | 3.35E-140 | 1.026806 | 0.966 | 0.706 | 6.75E-136 | 0 L1 _ | Serpinh1 |
| C2 | 2.04E-140 | 1.086392 | 0.693 | 0.191 | 4.12E-136 | 0 L1 _ | C2 |
| Scn7a1 | 2.16E-143 | 0.989791 | 0.844 | 0.296 | 4.36E-139 | 0 L1 _ | Scn7a |
| Eln | 3.18E-145 | 1.42667 | 0.891 | 0.428 | 6.41E-141 | 0 L1 _ | Eln |

| | | | | | | | |
|---|---|---|---|---|---|---|---|
| Itgb5 | 2.14E-146 | 1.180656 | 0.904 | 0.489 | 4.31E-142 | 0 L1 _ | Itgb5 |
| Hsd11b11 | 1.33E-149 | 1.647699 | 0.821 | 0.314 | 2.67E-145 | 0 L1 _ | Hsd11b1 |
| Gpm6b1 | 1.83E-150 | 1.124157 | 0.875 | 0.285 | 3.68E-146 | 0 L1 _ | Gpm6b |
| Cfb | 1.76E-151 | 1.355532 | 0.655 | 0.146 | 3.54E-147 | 0 L1 _ | Cfb |
| Timp1 | 5.07E-153 | 2.053152 | 0.71 | 0.183 | 1.02E-148 | 0 L1 _ | Timp1 |
| Lama21 | 7.58E-154 | 1.174344 | 0.948 | 0.406 | 1.53E-149 | 0 L1 _ | Lama2 |
| Mir99ahg | 2.10E-154 | 1.033698 | 0.686 | 0.149 | 4.24E-150 | 0 L1 _ | Mir99ahg |
| Col8a1 | 4.13E-155 | 1.783269 | 0.689 | 0.169 | 8.31E-151 | 0 L1 _ | Col8a1 |
| Pdgfrl | 2.19E-156 | 1.220712 | 0.66 | 0.13 | 4.41E-152 | 0 L1 _ | Pdgfrl |
| Mxra7 | 1.75E-156 | 1.128999 | 0.926 | 0.503 | 3.53E-152 | 0 L1 _ | Mxra7 |
| Epha3 | 1.57E-156 | 1.156966 | 0.712 | 0.182 | 3.16E-152 | 0 L1 _ | Epha3 |
| Adamts52 | 1.39E-157 | 1.389822 | 0.875 | 0.288 | 2.80E-153 | 0 L1 _ | Adamts5 |
| Fstl11 | 7.46E-158 | 1.175573 | 0.96 | 0.522 | 1.50E-153 | 0 L1 _ | Fstl1 |
| Fibin | 3.08E-158 | 1.295021 | 0.751 | 0.212 | 6.21E-154 | 0 L1 _ | Fibin |
| Crispld21 | 7.74E-159 | 1.24837 | 0.932 | 0.352 | 1.56E-154 | 0 L1 _ | Crispld2 |
| Lrp11 | 9.90E-162 | 1.20577 | 0.973 | 0.512 | 1.99E-157 | 0 L1 _ | Lrp1 |
| Itm2a1 | 1.27E-162 | 1.305694 | 0.935 | 0.426 | 2.56E-158 | 0 L1 _ | Itm2a |
| Cd63 | 1.01E-163 | 1.106349 | 0.981 | 0.691 | 2.02E-159 | 0 L1 _ | Cd63 |
| Gsn1 | 3.11E-165 | 1.084509 | 0.989 | 0.675 | 6.26E-161 | 0 L1 _ | Gsn |
| Pmepa1 | 5.99E-169 | 1.477315 | 0.95 | 0.566 | 1.21E-164 | 0 L1 _ | Pmepa1 |
| Dpysl31 | 5.88E-169 | 1.201258 | 0.865 | 0.283 | 1.18E-164 | 0 L1 _ | Dpysl3 |
| Mxra81 | 4.41E-169 | 1.233364 | 0.939 | 0.41 | 8.89E-165 | 0 L1 _ | Mxra8 |
| Htra31 | 3.76E-169 | 1.34169 | 0.924 | 0.324 | 7.58E-165 | 0 L1 _ | Htra3 |
| Ogn1 | 9.40E-170 | 1.551174 | 0.891 | 0.312 | 1.89E-165 | 0 L1 _ | Ogn |
| Col6a21 | 1.92E-170 | 1.318897 | 0.958 | 0.468 | 3.87E-166 | 0 L1 _ | Col6a2 |
| Pdgfra1 | 1.95E-171 | 1.370209 | 0.978 | 0.469 | 3.93E-167 | 0 L1 _ | Pdgfra |
| Igfbp41 | 3.77E-172 | 1.348912 | 0.975 | 0.57 | 7.60E-168 | 0 L1 _ | Igfbp4 |
| Il11ra11 | 1.27E-172 | 1.30427 | 0.932 | 0.396 | 2.56E-168 | 0 L1 _ | Il11ra1 |
| Igfbp61 | 3.61E-173 | 1.35674 | 0.848 | 0.244 | 7.28E-169 | 0 L1 _ | Igfbp6 |
| Rcn31 | 5.34E-177 | 1.225402 | 0.932 | 0.43 | 1.08E-172 | 0 L1 _ | Rcn3 |
| Col6a11 | 9.28E-179 | 1.331116 | 0.973 | 0.537 | 1.87E-174 | 0 L1 _ | Col6a1 |
| Serping11 | 2.76E-179 | 1.366648 | 0.982 | 0.604 | 5.56E-175 | 0 L1 _ | Serping1 |
| Spon11 | 2.21E-179 | 1.378563 | 0.849 | 0.254 | 4.45E-175 | 0 L1 _ | Spon1 |
| Podn1 | 1.88E-179 | 1.505721 | 0.818 | 0.23 | 3.78E-175 | 0 L1 _ | Podn |
| Pcolce1 | 1.61E-180 | 1.325548 | 0.973 | 0.604 | 3.24E-176 | 0 L1 _ | Pcolce |
| Col6a31 | 1.11E-181 | 1.538754 | 0.95 | 0.355 | 2.23E-177 | 0 L1 _ | Col6a3 |
| Col5a21 | 8.02E-183 | 1.466613 | 0.96 | 0.442 | 1.62E-178 | 0 L1 _ | Col5a2 |
| Plxdc21 | 8.85E-185 | 1.373988 | 0.953 | 0.337 | 1.78E-180 | 0 L1 _ | Plxdc2 |
| Entpd21 | 4.70E-185 | 1.308035 | 0.91 | 0.298 | 9.47E-181 | 0 L1 _ | Entpd2 |
| Ccdc801 | 6.06E-186 | 1.440915 | 0.941 | 0.401 | 1.22E-181 | 0 L1 _ | Ccdc80 |
| Prelp1 | 1.10E-186 | 1.505862 | 0.882 | 0.292 | 2.22E-182 | 0 L1 _ | Prelp |

| | | | | | | | |
|---|---|---|---|---|---|---|---|
| Mfap51 | 2.60E-189 | 1.746988 | 0.888 | 0.298 | 5.23E-185 | 0 L1 _ | Mfap5 |
| Dcn1 | 9.02E-191 | 1.619709 | 0.992 | 0.461 | 1.82E-186 | 0 L1 _ | Dcn |
| Ctsl1 | 1.94E-191 | 1.32455 | 0.975 | 0.701 | 3.91E-187 | 0 L1 _ | Ctsl |
| Abi3bp | 1.12E-191 | 1.591996 | 0.876 | 0.276 | 2.25E-187 | 0 L1 _ | Abi3bp |
| Gask1b | 2.08E-192 | 1.371254 | 0.785 | 0.177 | 4.18E-188 | 0 L1 _ | Gask1b |
| Lox | 1.53E-194 | 1.587288 | 0.797 | 0.195 | 3.09E-190 | 0 L1 _ | Lox |
| Col3a11 | 4.69E-198 | 1.710564 | 0.987 | 0.601 | 9.45E-194 | 0 L1 _ | Col3a1 |
| Mdk | 1.15E-205 | 1.524995 | 0.907 | 0.309 | 2.32E-201 | 0 L1 _ | Mdk |
| Lum1 | 9.84E-208 | 1.641907 | 0.98 | 0.355 | 1.98E-203 | 0 L1 _ | Lum |
| Col1a21 | 1.20E-208 | 1.736493 | 0.984 | 0.592 | 2.42E-204 | 0 L1 _ | Col1a2 |
| Colec121 | 4.09E-210 | 1.501374 | 0.94 | 0.322 | 8.24E-206 | 0 L1 _ | Colec12 |
| Emp11 | 1.92E-211 | 1.604539 | 0.961 | 0.453 | 3.86E-207 | 0 L1 _ | Emp1 |
| Smoc21 | 8.02E-212 | 1.725647 | 0.908 | 0.264 | 1.61E-207 | 0 L1 _ | Smoc2 |
| Mfap4 | 1.14E-212 | 1.923695 | 0.802 | 0.163 | 2.29E-208 | 0 L1 _ | Mfap4 |
| Fbln71 | 4.30E-213 | 1.448524 | 0.88 | 0.243 | 8.66E-209 | 0 L1 _ | Fbln7 |
| Spp1 | 7.09E-214 | 2.938211 | 0.842 | 0.224 | 1.43E-209 | 0 L1 _ | Spp1 |
| Ctsk | 4.18E-216 | 1.548095 | 0.92 | 0.323 | 8.41E-212 | 0 L1 _ | Ctsk |
| Fxyd6 | 2.01E-217 | 1.81976 | 0.854 | 0.224 | 4.05E-213 | 0 L1 _ | Fxyd6 |
| Bicc11 | 9.34E-218 | 1.576652 | 0.956 | 0.354 | 1.88E-213 | 0 L1 _ | Bicc1 |
| Col1a11 | 2.10E-218 | 1.853802 | 0.98 | 0.465 | 4.23E-214 | 0 L1 _ | Col1a1 |
| Lpl1 | 2.05E-218 | 1.848937 | 0.979 | 0.484 | 4.12E-214 | 0 L1 _ | Lpl |
| Cxcl141 | 6.98E-220 | 2.089792 | 0.965 | 0.379 | 1.41E-215 | 0 L1 _ | Cxcl14 |
| Fbln11 | 6.20E-220 | 1.589965 | 0.923 | 0.27 | 1.25E-215 | 0 L1 _ | Fbln1 |
| Olfml3 | 4.80E-220 | 1.609825 | 0.898 | 0.286 | 9.66E-216 | 0 L1 _ | Olfml3 |
| Svep1 | 2.02E-224 | 1.573931 | 0.891 | 0.246 | 4.08E-220 | 0 L1 _ | Svep1 |
| Cpxm11 | 3.57E-225 | 1.671395 | 0.924 | 0.271 | 7.18E-221 | 0 L1 _ | Cpxm1 |
| Cilp | 3.66E-230 | 2.479287 | 0.878 | 0.218 | 7.38E-226 | 0 L1 _ | Cilp |
| Nupr11 | 7.05E-232 | 1.928896 | 0.955 | 0.373 | 1.42E-227 | 0 L1 _ | Nupr1 |
| Apod1 | 3.32E-238 | 2.350431 | 0.986 | 0.421 | 6.68E-234 | 0 L1 _ | Apod |
| Cfh | 2.01E-238 | 1.853023 | 0.969 | 0.399 | 4.06E-234 | 0 L1 _ | Cfh |
| Cdh111 | 5.00E-242 | 1.801704 | 0.955 | 0.27 | 1.01E-237 | 0 L1 _ | Cdh11 |
| Aebp1 | 1.19E-243 | 1.782552 | 0.972 | 0.399 | 2.40E-239 | 0 L1 _ | Aebp1 |
| Itgbl1 | 5.21E-245 | 1.808423 | 0.893 | 0.215 | 1.05E-240 | 0 L1 _ | Itgbl1 |
| Loxl1 | 3.63E-247 | 1.787859 | 0.962 | 0.306 | 7.32E-243 | 0 L1 _ | Loxl1 |
| Rbp1 | 2.32E-265 | 2.130634 | 0.96 | 0.396 | 4.67E-261 | 0 L1 _ | Rbp1 |
| Serpinf11 | 2.84E-266 | 2.052953 | 0.98 | 0.336 | 5.72E-262 | 0 L1 _ | Serpinf1 |
| Bgn | 1.40E-279 | 2.173885 | 0.988 | 0.645 | 2.82E-275 | 0 L1 _ | Bgn |
| Mgp | 8.11E-292 | 2.765209 | 0.993 | 0.736 | 1.63E-287 | 0 L1 _ | Mgp |
| Ccdc88a | 1.86E-06 | 0.282521 | 0.599 | 0.412 | 0.037451 | 0 M1 _ | Ccdc88a |
| Xbp1 | 1.83E-06 | 0.267449 | 0.784 | 0.591 | 0.036758 | 0 M1 _ | Xbp1 |
| Snx6 | 1.75E-06 | 0.319531 | 0.737 | 0.539 | 0.035201 | 0 M1 _ | Snx6 |

| | | | | | | | |
|---|---|---|---|---|---|---|---|
| Sfrp2 | 1.53E-06 | 0.557054 | 0.204 | 0.088 | 0.030768 | 0 M1 _ | Sfrp2 |
| C4b | 1.38E-06 | 0.473834 | 0.192 | 0.081 | 0.027788 | 0 M1 _ | C4b |
| Lamc3 | 1.26E-06 | 0.311435 | 0.485 | 0.302 | 0.025298 | 0 M1 _ | Lamc3 |
| Ncbp2 | 1.17E-06 | 0.33131 | 0.587 | 0.382 | 0.023591 | 0 M1 _ | Ncbp2 |
| Irgm2 | 1.16E-06 | 0.260315 | 0.365 | 0.203 | 0.023379 | 0 M1 _ | Irgm2 |
| Ptgis | 1.03E-06 | 0.321014 | 0.413 | 0.244 | 0.02081 | 0 M1 _ | Ptgis |
| Sec63 | 9.58E-07 | 0.276401 | 0.772 | 0.55 | 0.019287 | 0 M1 _ | Sec63 |
| Slc25a39 | 9.01E-07 | 0.266498 | 0.563 | 0.369 | 0.018143 | 0 M1 _ | Slc25a39 |
| Pdap1 | 8.65E-07 | 0.264643 | 0.904 | 0.711 | 0.017428 | 0 M1 _ | Pdap1 |
| Nucb2 | 8.01E-07 | 0.254867 | 0.593 | 0.396 | 0.016127 | 0 M1 _ | Nucb2 |
| Matr3 | 7.96E-07 | 0.250398 | 0.749 | 0.517 | 0.016031 | 0 M1 _ | Matr3 |
| Spon1 | 7.23E-07 | 0.465567 | 0.683 | 0.481 | 0.014553 | 0 M1 _ | Spon1 |
| Errfi1 | 6.59E-07 | 0.37925 | 0.814 | 0.615 | 0.013269 | 0 M1 _ | Errfi1 |
| Lmna | 3.86E-07 | 0.365582 | 0.91 | 0.68 | 0.00777 | 0 M1 _ | Lmna |
| Hist1h2bc | 3.80E-07 | 0.353975 | 0.701 | 0.478 | 0.007657 | 0 M1 _ | Hist1h2bc |
| Rex1bd | 3.32E-07 | 0.289285 | 0.808 | 0.58 | 0.006692 | 0 M1 _ | Rex1bd |
| Ace | 3.13E-07 | 0.312419 | 0.593 | 0.384 | 0.006314 | 0 M1 _ | Ace |
| Cpxm1 | 2.80E-07 | 0.291627 | 0.802 | 0.516 | 0.005647 | 0 M1 _ | Cpxm1 |
| Lhfpl2 | 2.77E-07 | 0.251662 | 0.431 | 0.247 | 0.005589 | 0 M1 _ | Lhfpl2 |
| Ikbip | 1.91E-07 | 0.345953 | 0.671 | 0.463 | 0.003854 | 0 M1 _ | Ikbip |
| Tuba1b | 1.85E-07 | 0.312288 | 0.952 | 0.813 | 0.003736 | 0 M1 _ | Tuba1b |
| Fam120a | 1.75E-07 | 0.256668 | 0.641 | 0.429 | 0.00353 | 0 M1 _ | Fam120a |
| Rpl4 | 1.73E-07 | 0.314828 | 0.976 | 0.83 | 0.003492 | 0 M1 _ | Rpl4 |
| Nupr1 | 1.67E-07 | 0.314685 | 0.964 | 0.582 | 0.003369 | 0 M1 _ | Nupr1 |
| Lbp | 1.59E-07 | 0.26101 | 0.461 | 0.265 | 0.003195 | 0 M1 _ | Lbp |
| Golim4 | 1.54E-07 | 0.349714 | 0.79 | 0.563 | 0.003102 | 0 M1 _ | Golim4 |
| Capn1 | 1.42E-07 | 0.261089 | 0.443 | 0.261 | 0.002853 | 0 M1 _ | Capn1 |
| Sec62 | 1.38E-07 | 0.295436 | 0.964 | 0.819 | 0.002784 | 0 M1 _ | Sec62 |
| Pofut2 | 1.33E-07 | 0.30597 | 0.671 | 0.444 | 0.00267 | 0 M1 _ | Pofut2 |
| Pds5b | 1.32E-07 | 0.279246 | 0.587 | 0.367 | 0.002653 | 0 M1 _ | Pds5b |
| Kdelr3 | 1.31E-07 | 0.253068 | 0.629 | 0.38 | 0.002633 | 0 M1 _ | Kdelr3 |
| Hdlbp | 1.27E-07 | 0.294505 | 0.934 | 0.682 | 0.002564 | 0 M1 _ | Hdlbp |
| Sspn | 1.25E-07 | 0.262326 | 0.551 | 0.34 | 0.002522 | 0 M1 _ | Sspn |
| Tm2d2 | 1.24E-07 | 0.276199 | 0.713 | 0.476 | 0.002494 | 0 M1 _ | Tm2d2 |
| Tspan31 | 1.22E-07 | 0.264161 | 0.659 | 0.445 | 0.00246 | 0 M1 _ | Tspan31 |
| Ogn | 1.16E-07 | 0.433338 | 0.772 | 0.53 | 0.002331 | 0 M1 _ | Ogn |
| Anxa2 | 1.04E-07 | 0.288292 | 0.976 | 0.839 | 0.002085 | 0 M1 _ | Anxa2 |
| Dhrs3 | 9.84E-08 | 0.266609 | 0.635 | 0.405 | 0.001983 | 0 M1 _ | Dhrs3 |
| Tm9sf3 | 9.84E-08 | 0.281227 | 0.922 | 0.738 | 0.001982 | 0 M1 _ | Tm9sf3 |
| Rcn3 | 9.37E-08 | 0.273383 | 0.928 | 0.611 | 0.001886 | 0 M1 _ | Rcn3 |
| Yipf5 | 9.36E-08 | 0.263762 | 0.707 | 0.477 | 0.001884 | 0 M1 _ | Yipf5 |

| | | | | | | | |
|---|---|---|---|---|---|---|---|
| Eif4g1 | 9.10E-08 | 0.282231 | 0.784 | 0.574 | 0.001834 | 0 M1 _ | Eif4g1 |
| Ier3ip1 | 8.80E-08 | 0.263036 | 0.88 | 0.644 | 0.001772 | 0 M1 _ | Ier3ip1 |
| Uba1 | 7.15E-08 | 0.264951 | 0.653 | 0.438 | 0.001439 | 0 M1 _ | Uba1 |
| Mgat2 | 6.55E-08 | 0.298038 | 0.671 | 0.43 | 0.001319 | 0 M1 _ | Mgat2 |
| Sec61g | 6.49E-08 | 0.330024 | 0.97 | 0.852 | 0.001308 | 0 M1 _ | Sec61g |
| Shox2 | 5.86E-08 | 0.272303 | 0.437 | 0.241 | 0.00118 | 0 M1 _ | Shox2 |
| Socs4 | 4.79E-08 | 0.26334 | 0.407 | 0.209 | 0.000964 | 0 M1 _ | Socs4 |
| Ccdc80 | 4.41E-08 | 0.305476 | 0.904 | 0.598 | 0.000889 | 0 M1 _ | Ccdc80 |
| Wipf1 | 4.39E-08 | 0.283841 | 0.647 | 0.438 | 0.000884 | 0 M1 _ | Wipf1 |
| Dync1h1 | 3.98E-08 | 0.29105 | 0.832 | 0.618 | 0.000801 | 0 M1 _ | Dync1h1 |
| Rab6a | 3.66E-08 | 0.260849 | 0.737 | 0.504 | 0.000738 | 0 M1 _ | Rab6a |
| Fxyd1 | 3.52E-08 | 0.258569 | 0.946 | 0.658 | 0.000709 | 0 M1 _ | Fxyd1 |
| Nrxn1 | 3.43E-08 | 0.46357 | 0.257 | 0.113 | 0.000692 | 0 M1 _ | Nrxn1 |
| Vcp | 3.22E-08 | 0.320936 | 0.94 | 0.752 | 0.000649 | 0 M1 _ | Vcp |
| Anxa5 | 3.18E-08 | 0.289226 | 0.994 | 0.879 | 0.00064 | 0 M1 _ | Anxa5 |
| Dusp1 | 3.04E-08 | 0.281266 | 0.557 | 0.322 | 0.000611 | 0 M1 _ | Dusp1 |
| Twf1 | 2.93E-08 | 0.291049 | 0.617 | 0.401 | 0.00059 | 0 M1 _ | Twf1 |
| Gabarapl1 | 2.62E-08 | 0.269372 | 0.707 | 0.469 | 0.000527 | 0 M1 _ | Gabarapl1 |
| Klf6 | 2.55E-08 | 0.339043 | 0.778 | 0.543 | 0.000514 | 0 M1 _ | Klf6 |
| Chpt1 | 2.40E-08 | 0.298438 | 0.587 | 0.36 | 0.000484 | 0 M1 _ | Chpt1 |
| Cebpb | 1.92E-08 | 0.311682 | 0.647 | 0.414 | 0.000387 | 0 M1 _ | Cebpb |
| Svil | 1.84E-08 | 0.413352 | 0.701 | 0.488 | 0.000371 | 0 M1 _ | Svil |
| Cdh11 | 1.81E-08 | 0.317812 | 0.85 | 0.525 | 0.000366 | 0 M1 _ | Cdh11 |
| Tor1aip1 | 1.72E-08 | 0.387919 | 0.79 | 0.576 | 0.000347 | 0 M1 _ | Tor1aip1 |
| Lman1 | 1.58E-08 | 0.321131 | 0.844 | 0.612 | 0.000319 | 0 M1 _ | Lman1 |
| Copa | 1.55E-08 | 0.27564 | 0.838 | 0.602 | 0.000313 | 0 M1 _ | Copa |
| Dram1 | 1.43E-08 | 0.250177 | 0.335 | 0.165 | 0.000289 | 0 M1 _ | Dram1 |
| Sp3 | 1.43E-08 | 0.275016 | 0.671 | 0.436 | 0.000287 | 0 M1 _ | Sp3 |
| Aldh1a2 | 1.42E-08 | 0.274226 | 0.162 | 0.053 | 0.000286 | 0 M1 _ | Aldh1a2 |
| Mettl1 | 1.27E-08 | 0.252904 | 0.491 | 0.276 | 0.000256 | 0 M1 _ | Mettl1 |
| Phactr2 | 1.20E-08 | 0.257365 | 0.665 | 0.42 | 0.000242 | 0 M1 _ | Phactr2 |
| Fn1 | 1.16E-08 | 0.443684 | 0.317 | 0.154 | 0.000234 | 0 M1 _ | Fn1 |
| Amotl1 | 1.11E-08 | 0.30247 | 0.856 | 0.622 | 0.000224 | 0 M1 _ | Amotl1 |
| Adamts4 | 1.09E-08 | 0.25811 | 0.329 | 0.158 | 0.000219 | 0 M1 _ | Adamts4 |
| Phtf1 | 9.40E-09 | 0.303957 | 0.341 | 0.161 | 0.000189 | 0 M1 _ | Phtf1 |
| Rab1a | 9.28E-09 | 0.342343 | 0.802 | 0.558 | 0.000187 | 0 M1 _ | Rab1a |
| Cdkn1c | 7.74E-09 | 0.582819 | 0.605 | 0.397 | 0.000156 | 0 M1 _ | Cdkn1c |
| Derl1 | 6.87E-09 | 0.252808 | 0.671 | 0.428 | 0.000138 | 0 M1 _ | Derl1 |
| Layn | 6.60E-09 | 0.305305 | 0.449 | 0.245 | 0.000133 | 0 M1 _ | Layn |
| Lrrc8a | 6.32E-09 | 0.289384 | 0.587 | 0.353 | 0.000127 | 0 M1 _ | Lrrc8a |
| Sntb2 | 6.21E-09 | 0.325288 | 0.802 | 0.533 | 0.000125 | 0 M1 _ | Sntb2 |

| | | | | | | | |
|---|---|---|---|---|---|---|---|
| Nucb1 | 5.45E-09 | 0.367833 | 0.826 | 0.596 | 0.00011 | 0 M1 _ | Nucb1 |
| Tmed3 | 5.36E-09 | 0.328585 | 0.922 | 0.654 | 0.000108 | 0 M1 _ | Tmed3 |
| Sqle | 4.81E-09 | 0.274008 | 0.329 | 0.155 | 9.69E-05 | 0 M1 _ | Sqle |
| 5730455P16Rik | 4.72E-09 | 0.259898 | 0.359 | 0.175 | 9.51E-05 | 0 M1 _ | 5730455P16Rik |
| Mia3 | 4.52E-09 | 0.311111 | 0.743 | 0.525 | 9.10E-05 | 0 M1 _ | Mia3 |
| Scarf2 | 4.39E-09 | 0.324912 | 0.689 | 0.433 | 8.85E-05 | 0 M1 _ | Scarf2 |
| Scn1b | 4.38E-09 | 0.265022 | 0.79 | 0.518 | 8.82E-05 | 0 M1 _ | Scn1b |
| Egr1 | 4.37E-09 | 0.549175 | 0.563 | 0.349 | 8.81E-05 | 0 M1 _ | Egr1 |
| Arhgef40 | 4.15E-09 | 0.259064 | 0.713 | 0.434 | 8.35E-05 | 0 M1 _ | Arhgef40 |
| Phldb1 | 3.95E-09 | 0.270957 | 0.557 | 0.328 | 7.96E-05 | 0 M1 _ | Phldb1 |
| Trip12 | 3.80E-09 | 0.286211 | 0.719 | 0.473 | 7.66E-05 | 0 M1 _ | Trip12 |
| Atp6v0d1 | 3.78E-09 | 0.254554 | 0.665 | 0.408 | 7.60E-05 | 0 M1 _ | Atp6v0d1 |
| Crlf1 | 3.45E-09 | 0.397817 | 0.671 | 0.402 | 6.95E-05 | 0 M1 _ | Crlf1 |
| Mcfd2 | 3.41E-09 | 0.291884 | 0.485 | 0.276 | 6.86E-05 | 0 M1 _ | Mcfd2 |
| Tmem159 | 3.26E-09 | 0.280782 | 0.545 | 0.316 | 6.57E-05 | 0 M1 _ | Tmem159 |
| Grsf1 | 3.19E-09 | 0.257205 | 0.371 | 0.18 | 6.42E-05 | 0 M1 _ | Grsf1 |
| Gaa | 3.18E-09 | 0.326776 | 0.766 | 0.489 | 6.41E-05 | 0 M1 _ | Gaa |
| Furin | 3.17E-09 | 0.267647 | 0.539 | 0.314 | 6.38E-05 | 0 M1 _ | Furin |
| Nudcd2 | 3.11E-09 | 0.28652 | 0.461 | 0.256 | 6.25E-05 | 0 M1 _ | Nudcd2 |
| Cand1 | 2.89E-09 | 0.276805 | 0.419 | 0.223 | 5.82E-05 | 0 M1 _ | Cand1 |
| Emp1 | 2.88E-09 | 0.287438 | 0.97 | 0.636 | 5.80E-05 | 0 M1 _ | Emp1 |
| Cblb | 2.84E-09 | 0.314972 | 0.497 | 0.283 | 5.72E-05 | 0 M1 _ | Cblb |
| Herpud1 | 2.72E-09 | 0.310871 | 0.778 | 0.533 | 5.48E-05 | 0 M1 _ | Herpud1 |
| Ssbp3 | 2.54E-09 | 0.299615 | 0.575 | 0.343 | 5.12E-05 | 0 M1 _ | Ssbp3 |
| Golga2 | 2.39E-09 | 0.280112 | 0.539 | 0.304 | 4.81E-05 | 0 M1 _ | Golga2 |
| Lnpep | 2.15E-09 | 0.300458 | 0.737 | 0.476 | 4.33E-05 | 0 M1 _ | Lnpep |
| mt-Nd3 | 2.04E-09 | 0.348547 | 0.988 | 0.915 | 4.10E-05 | 0 M1 _ | mt-Nd3 |
| Caprin1 | 2.04E-09 | 0.293698 | 0.784 | 0.53 | 4.10E-05 | 0 M1 _ | Caprin1 |
| Sdcbp | 2.01E-09 | 0.361468 | 0.838 | 0.603 | 4.04E-05 | 0 M1 _ | Sdcbp |
| Tmed2 | 1.99E-09 | 0.342122 | 0.94 | 0.727 | 4.01E-05 | 0 M1 _ | Tmed2 |
| App | 1.97E-09 | 0.360029 | 0.994 | 0.828 | 3.96E-05 | 0 M1 _ | App |
| Rarres2 | 1.95E-09 | 0.334096 | 0.916 | 0.674 | 3.92E-05 | 0 M1 _ | Rarres2 |
| Tcf7l2 | 1.82E-09 | 0.300391 | 0.719 | 0.448 | 3.66E-05 | 0 M1 _ | Tcf7l2 |
| Copg1 | 1.81E-09 | 0.251813 | 0.623 | 0.373 | 3.65E-05 | 0 M1 _ | Copg1 |
| Wdr26 | 1.71E-09 | 0.270393 | 0.719 | 0.45 | 3.45E-05 | 0 M1 _ | Wdr26 |
| Esyt2 | 1.62E-09 | 0.285146 | 0.772 | 0.523 | 3.27E-05 | 0 M1 _ | Esyt2 |
| Aida | 1.55E-09 | 0.347042 | 0.641 | 0.389 | 3.12E-05 | 0 M1 _ | Aida |
| Clec11a | 1.33E-09 | 0.266834 | 0.455 | 0.229 | 2.68E-05 | 0 M1 _ | Clec11a |
| Ttc28 | 1.28E-09 | 0.32566 | 0.82 | 0.553 | 2.57E-05 | 0 M1 _ | Ttc28 |
| S100a16 | 1.09E-09 | 0.315308 | 0.904 | 0.574 | 2.20E-05 | 0 M1 _ | S100a16 |
| Tmed7 | 1.08E-09 | 0.312843 | 0.569 | 0.343 | 2.17E-05 | 0 M1 _ | Tmed7 |

| | | | | | | | |
|---|---|---|---|---|---|---|---|
| Bace1 | 9.24E-10 | 0.318497 | 0.593 | 0.354 | 1.86E-05 | 0 M1 _ | Bace1 |
| Mtmr6 | 9.23E-10 | 0.255548 | 0.461 | 0.245 | 1.86E-05 | 0 M1 _ | Mtmr6 |
| Rnf144a | 8.84E-10 | 0.343289 | 0.551 | 0.323 | 1.78E-05 | 0 M1 _ | Rnf144a |
| Selenos | 8.70E-10 | 0.326486 | 0.91 | 0.632 | 1.75E-05 | 0 M1 _ | Selenos |
| Csde1 | 8.26E-10 | 0.333544 | 0.91 | 0.686 | 1.66E-05 | 0 M1 _ | Csde1 |
| Osbpl1a | 8.01E-10 | 0.270233 | 0.437 | 0.225 | 1.61E-05 | 0 M1 _ | Osbpl1a |
| Srpx2 | 7.88E-10 | 0.314041 | 0.605 | 0.354 | 1.59E-05 | 0 M1 _ | Srpx2 |
| Scpep1 | 7.33E-10 | 0.304406 | 0.731 | 0.444 | 1.48E-05 | 0 M1 _ | Scpep1 |
| Pbx1 | 6.25E-10 | 0.351571 | 0.743 | 0.509 | 1.26E-05 | 0 M1 _ | Pbx1 |
| P4ha1 | 6.16E-10 | 0.318887 | 0.76 | 0.469 | 1.24E-05 | 0 M1 _ | P4ha1 |
| Nckap1 | 6.09E-10 | 0.28308 | 0.76 | 0.5 | 1.23E-05 | 0 M1 _ | Nckap1 |
| Spry2 | 6.00E-10 | 0.258477 | 0.431 | 0.218 | 1.21E-05 | 0 M1 _ | Spry2 |
| Kdelr1 | 5.79E-10 | 0.348862 | 0.868 | 0.624 | 1.17E-05 | 0 M1 _ | Kdelr1 |
| Rtn4 | 5.45E-10 | 0.391999 | 0.964 | 0.728 | 1.10E-05 | 0 M1 _ | Rtn4 |
| Vps35 | 5.40E-10 | 0.3031 | 0.707 | 0.441 | 1.09E-05 | 0 M1 _ | Vps35 |
| Prrc2a | 5.32E-10 | 0.279182 | 0.641 | 0.391 | 1.07E-05 | 0 M1 _ | Prrc2a |
| Trf | 5.22E-10 | 0.84732 | 0.443 | 0.253 | 1.05E-05 | 0 M1 _ | Trf |
| Pea15a | 5.17E-10 | 0.321935 | 0.826 | 0.549 | 1.04E-05 | 0 M1 _ | Pea15a |
| Grina | 4.95E-10 | 0.37259 | 0.808 | 0.55 | 9.97E-06 | 0 M1 _ | Grina |
| Thbs3 | 4.74E-10 | 0.280516 | 0.323 | 0.145 | 9.55E-06 | 0 M1 _ | Thbs3 |
| Smarca2 | 4.58E-10 | 0.38136 | 0.838 | 0.618 | 9.22E-06 | 0 M1 _ | Smarca2 |
| Ppp3r1 | 4.57E-10 | 0.258975 | 0.509 | 0.274 | 9.20E-06 | 0 M1 _ | Ppp3r1 |
| Ncor2 | 4.55E-10 | 0.291778 | 0.665 | 0.388 | 9.15E-06 | 0 M1 _ | Ncor2 |
| Cpd | 4.51E-10 | 0.313278 | 0.725 | 0.466 | 9.08E-06 | 0 M1 _ | Cpd |
| Naalad2 | 4.32E-10 | 0.29957 | 0.737 | 0.459 | 8.70E-06 | 0 M1 _ | Naalad2 |
| Ppp3cb | 3.74E-10 | 0.292812 | 0.677 | 0.403 | 7.53E-06 | 0 M1 _ | Ppp3cb |
| St13 | 3.65E-10 | 0.345527 | 0.928 | 0.73 | 7.35E-06 | 0 M1 _ | St13 |
| Slc25a24 | 3.33E-10 | 0.29444 | 0.551 | 0.307 | 6.71E-06 | 0 M1 _ | Slc25a24 |
| Scoc | 3.27E-10 | 0.257236 | 0.605 | 0.331 | 6.58E-06 | 0 M1 _ | Scoc |
| Camkk2 | 3.22E-10 | 0.265997 | 0.491 | 0.255 | 6.49E-06 | 0 M1 _ | Camkk2 |
| Pik3r1 | 3.11E-10 | 0.341714 | 0.85 | 0.652 | 6.26E-06 | 0 M1 _ | Pik3r1 |
| Nmb | 3.08E-10 | 0.298247 | 0.24 | 0.089 | 6.19E-06 | 0 M1 _ | Nmb |
| Fcgrt | 2.93E-10 | 0.35477 | 0.952 | 0.7 | 5.90E-06 | 0 M1 _ | Fcgrt |
| Tmem100 | 2.76E-10 | 0.276869 | 0.515 | 0.269 | 5.56E-06 | 0 M1 _ | Tmem100 |
| Amfr | 2.75E-10 | 0.272772 | 0.509 | 0.283 | 5.55E-06 | 0 M1 _ | Amfr |
| P3h3 | 2.71E-10 | 0.320907 | 0.796 | 0.479 | 5.45E-06 | 0 M1 _ | P3h3 |
| Xdh | 2.69E-10 | 0.33637 | 0.683 | 0.414 | 5.42E-06 | 0 M1 _ | Xdh |
| Srebf1 | 2.63E-10 | 0.277699 | 0.461 | 0.243 | 5.29E-06 | 0 M1 _ | Srebf1 |
| Rnd3 | 2.54E-10 | 0.300583 | 0.641 | 0.361 | 5.12E-06 | 0 M1 _ | Rnd3 |
| Arl1 | 2.50E-10 | 0.378406 | 0.88 | 0.648 | 5.04E-06 | 0 M1 _ | Arl1 |
| Ppp1r2 | 2.44E-10 | 0.376769 | 0.898 | 0.672 | 4.92E-06 | 0 M1 _ | Ppp1r2 |

| | | | | | | | |
|---|---|---|---|---|---|---|---|
| Gadd45a | 2.43E-10 | 0.266596 | 0.539 | 0.288 | 4.90E-06 | 0 M1 _ | Gadd45a |
| Mtpn | 2.36E-10 | 0.29518 | 0.76 | 0.498 | 4.76E-06 | 0 M1 _ | Mtpn |
| Gfpt1 | 2.34E-10 | 0.325382 | 0.521 | 0.288 | 4.71E-06 | 0 M1 _ | Gfpt1 |
| Ext1 | 2.28E-10 | 0.371614 | 0.599 | 0.353 | 4.58E-06 | 0 M1 _ | Ext1 |
| Dnajc25 | 2.26E-10 | 0.295884 | 0.563 | 0.315 | 4.56E-06 | 0 M1 _ | Dnajc25 |
| Eif4g2 | 2.21E-10 | 0.331426 | 0.922 | 0.769 | 4.44E-06 | 0 M1 _ | Eif4g2 |
| Zfp503 | 2.16E-10 | 0.408121 | 0.653 | 0.41 | 4.36E-06 | 0 M1 _ | Zfp503 |
| Srxn1 | 2.12E-10 | 0.292786 | 0.287 | 0.118 | 4.27E-06 | 0 M1 _ | Srxn1 |
| Itsn1 | 1.99E-10 | 0.271604 | 0.413 | 0.204 | 4.00E-06 | 0 M1 _ | Itsn1 |
| Carm1 | 1.82E-10 | 0.258576 | 0.377 | 0.174 | 3.67E-06 | 0 M1 _ | Carm1 |
| Mafb | 1.63E-10 | 0.403294 | 0.868 | 0.568 | 3.28E-06 | 0 M1 _ | Mafb |
| Kdm7a | 1.61E-10 | 0.267735 | 0.473 | 0.236 | 3.24E-06 | 0 M1 _ | Kdm7a |
| Smoc2 | 1.46E-10 | 0.697342 | 0.826 | 0.503 | 2.94E-06 | 0 M1 _ | Smoc2 |
| Spats2l | 1.45E-10 | 0.31157 | 0.778 | 0.472 | 2.92E-06 | 0 M1 _ | Spats2l |
| Tgfbr2 | 1.35E-10 | 0.407704 | 0.892 | 0.635 | 2.72E-06 | 0 M1 _ | Tgfbr2 |
| Wipi1 | 1.17E-10 | 0.366046 | 0.629 | 0.383 | 2.35E-06 | 0 M1 _ | Wipi1 |
| Usp9x | 1.09E-10 | 0.406087 | 0.766 | 0.513 | 2.19E-06 | 0 M1 _ | Usp9x |
| Cyp26b1 | 1.06E-10 | 0.462133 | 0.263 | 0.099 | 2.13E-06 | 0 M1 _ | Cyp26b1 |
| Slc38a10 | 1.04E-10 | 0.355253 | 0.653 | 0.399 | 2.09E-06 | 0 M1 _ | Slc38a10 |
| Col4a1 | 1.01E-10 | 0.399724 | 0.982 | 0.865 | 2.04E-06 | 0 M1 _ | Col4a1 |
| Reep3 | 1.00E-10 | 0.393701 | 0.91 | 0.681 | 2.02E-06 | 0 M1 _ | Reep3 |
| Rnf6 | 9.97E-11 | 0.33119 | 0.623 | 0.373 | 2.01E-06 | 0 M1 _ | Rnf6 |
| Map3k7 | 9.70E-11 | 0.275826 | 0.401 | 0.192 | 1.95E-06 | 0 M1 _ | Map3k7 |
| Fam91a1 | 8.83E-11 | 0.271628 | 0.425 | 0.21 | 1.78E-06 | 0 M1 _ | Fam91a1 |
| Taok1 | 8.41E-11 | 0.32921 | 0.772 | 0.503 | 1.69E-06 | 0 M1 _ | Taok1 |
| Csf1 | 8.26E-11 | 0.313487 | 0.659 | 0.38 | 1.66E-06 | 0 M1 _ | Csf1 |
| Zbtb4 | 7.93E-11 | 0.320854 | 0.617 | 0.353 | 1.60E-06 | 0 M1 _ | Zbtb4 |
| Slc48a1 | 7.88E-11 | 0.340898 | 0.587 | 0.335 | 1.59E-06 | 0 M1 _ | Slc48a1 |
| Arap1 | 7.68E-11 | 0.344929 | 0.683 | 0.396 | 1.55E-06 | 0 M1 _ | Arap1 |
| Plscr4 | 7.22E-11 | 0.325099 | 0.677 | 0.402 | 1.45E-06 | 0 M1 _ | Plscr4 |
| Dnajc3 | 7.18E-11 | 0.404332 | 0.922 | 0.698 | 1.45E-06 | 0 M1 _ | Dnajc3 |
| Negr1 | 7.16E-11 | 0.256539 | 0.246 | 0.091 | 1.44E-06 | 0 M1 _ | Negr1 |
| Rhoq | 6.88E-11 | 0.302509 | 0.647 | 0.392 | 1.39E-06 | 0 M1 _ | Rhoq |
| Myadm | 6.78E-11 | 0.295994 | 0.575 | 0.32 | 1.36E-06 | 0 M1 _ | Myadm |
| BC005537 | 6.62E-11 | 0.323535 | 0.707 | 0.432 | 1.33E-06 | 0 M1 _ | BC005537 |
| Acadsb | 6.59E-11 | 0.35656 | 0.617 | 0.368 | 1.33E-06 | 0 M1 _ | Acadsb |
| Smim1 | 6.26E-11 | 0.289525 | 0.425 | 0.211 | 1.26E-06 | 0 M1 _ | Smim1 |
| Wdfy2 | 6.20E-11 | 0.255429 | 0.341 | 0.15 | 1.25E-06 | 0 M1 _ | Wdfy2 |
| Snx18 | 5.98E-11 | 0.299659 | 0.665 | 0.398 | 1.20E-06 | 0 M1 _ | Snx18 |
| Ugdh | 5.45E-11 | 0.377646 | 0.449 | 0.23 | 1.10E-06 | 0 M1 _ | Ugdh |
| Tmem53 | 5.42E-11 | 0.270954 | 0.24 | 0.087 | 1.09E-06 | 0 M1 _ | Tmem53 |

| | | | | | | | |
|---|---|---|---|---|---|---|---|
| Ddah2 | 4.93E-11 | 0.391618 | 0.772 | 0.505 | 9.94E-07 | 0 M1 _ | Ddah2 |
| Col1a1 | 4.75E-11 | 0.389034 | 0.988 | 0.65 | 9.56E-07 | 0 M1 _ | Col1a1 |
| Mob1a | 4.74E-11 | 0.275927 | 0.629 | 0.356 | 9.54E-07 | 0 M1 _ | Mob1a |
| Adamtsl2 | 4.24E-11 | 0.720112 | 0.509 | 0.287 | 8.54E-07 | 0 M1 _ | Adamtsl2 |
| Vma21 | 4.03E-11 | 0.308109 | 0.587 | 0.334 | 8.13E-07 | 0 M1 _ | Vma21 |
| Plxnc1 | 3.96E-11 | 0.308207 | 0.293 | 0.122 | 7.98E-07 | 0 M1 _ | Plxnc1 |
| Fasn | 3.82E-11 | 0.328854 | 0.395 | 0.182 | 7.69E-07 | 0 M1 _ | Fasn |
| Il4ra | 3.52E-11 | 0.287566 | 0.461 | 0.231 | 7.09E-07 | 0 M1 _ | Il4ra |
| Mmp3 | 3.40E-11 | 0.729089 | 0.599 | 0.333 | 6.85E-07 | 0 M1 _ | Mmp3 |
| Rora | 3.19E-11 | 0.394031 | 0.814 | 0.506 | 6.42E-07 | 0 M1 _ | Rora |
| Tnrc18 | 2.97E-11 | 0.359869 | 0.581 | 0.325 | 5.98E-07 | 0 M1 _ | Tnrc18 |
| Epb41l2 | 2.81E-11 | 0.375077 | 0.904 | 0.663 | 5.66E-07 | 0 M1 _ | Epb41l2 |
| Dag1 | 2.45E-11 | 0.331228 | 0.838 | 0.573 | 4.92E-07 | 0 M1 _ | Dag1 |
| Lpp | 2.38E-11 | 0.324101 | 0.874 | 0.612 | 4.79E-07 | 0 M1 _ | Lpp |
| Igf1r | 2.36E-11 | 0.353543 | 0.593 | 0.338 | 4.76E-07 | 0 M1 _ | Igf1r |
| Lmo4 | 2.34E-11 | 0.35695 | 0.796 | 0.557 | 4.71E-07 | 0 M1 _ | Lmo4 |
| Peg3 | 2.23E-11 | 0.755577 | 0.371 | 0.178 | 4.48E-07 | 0 M1 _ | Peg3 |
| Podn | 2.22E-11 | 0.304435 | 0.778 | 0.445 | 4.48E-07 | 0 M1 _ | Podn |
| Gm30075 | 1.63E-11 | 0.289401 | 0.353 | 0.15 | 3.29E-07 | 0 M1 _ | Gm30075 |
| Fbln7 | 1.50E-11 | 0.331748 | 0.862 | 0.474 | 3.02E-07 | 0 M1 _ | Fbln7 |
| Arcn1 | 1.47E-11 | 0.371364 | 0.772 | 0.51 | 2.97E-07 | 0 M1 _ | Arcn1 |
| Emp2 | 1.43E-11 | 0.35754 | 0.82 | 0.543 | 2.87E-07 | 0 M1 _ | Emp2 |
| Acvr2a | 1.41E-11 | 0.272214 | 0.365 | 0.16 | 2.83E-07 | 0 M1 _ | Acvr2a |
| Plod2 | 1.29E-11 | 0.395691 | 0.551 | 0.298 | 2.60E-07 | 0 M1 _ | Plod2 |
| Acp2 | 1.27E-11 | 0.334514 | 0.467 | 0.238 | 2.55E-07 | 0 M1 _ | Acp2 |
| Clcn4 | 1.18E-11 | 0.317068 | 0.725 | 0.463 | 2.38E-07 | 0 M1 _ | Clcn4 |
| Agtrap | 1.00E-11 | 0.358189 | 0.617 | 0.343 | 2.02E-07 | 0 M1 _ | Agtrap |
| Magt1 | 1.00E-11 | 0.396568 | 0.713 | 0.436 | 2.02E-07 | 0 M1 _ | Magt1 |
| Hsd11b1 | 9.99E-12 | 0.520129 | 0.796 | 0.498 | 2.01E-07 | 0 M1 _ | Hsd11b1 |
| C1qtnf7 | 9.86E-12 | 0.27988 | 0.407 | 0.181 | 1.99E-07 | 0 M1 _ | C1qtnf7 |
| Pdia6 | 9.82E-12 | 0.482237 | 0.934 | 0.723 | 1.98E-07 | 0 M1 _ | Pdia6 |
| Prrg3 | 9.49E-12 | 0.262677 | 0.359 | 0.15 | 1.91E-07 | 0 M1 _ | Prrg3 |
| Mbtps1 | 9.34E-12 | 0.324986 | 0.701 | 0.419 | 1.88E-07 | 0 M1 _ | Mbtps1 |
| Nr3c1 | 8.56E-12 | 0.388893 | 0.952 | 0.721 | 1.72E-07 | 0 M1 _ | Nr3c1 |
| Tgfb2 | 8.36E-12 | 0.367176 | 0.605 | 0.331 | 1.68E-07 | 0 M1 _ | Tgfb2 |
| Rpn2 | 8.20E-12 | 0.365354 | 0.886 | 0.659 | 1.65E-07 | 0 M1 _ | Rpn2 |
| Tcf4 | 7.80E-12 | 0.443271 | 0.988 | 0.881 | 1.57E-07 | 0 M1 _ | Tcf4 |
| Rrbp1 | 7.79E-12 | 0.38054 | 0.988 | 0.876 | 1.57E-07 | 0 M1 _ | Rrbp1 |
| Trps1 | 7.71E-12 | 0.309637 | 0.551 | 0.296 | 1.55E-07 | 0 M1 _ | Trps1 |
| Pdpn | 7.52E-12 | 0.385465 | 0.641 | 0.361 | 1.51E-07 | 0 M1 _ | Pdpn |
| Gnaq | 7.30E-12 | 0.27587 | 0.731 | 0.425 | 1.47E-07 | 0 M1 _ | Gnaq |

| | | | | | | | |
|---|---|---|---|---|---|---|---|
| Atp11a | 6.58E-12 | 0.253412 | 0.443 | 0.207 | 1.33E-07 | 0 M1 _ | Atp11a |
| Pxdn | 6.25E-12 | 0.308987 | 0.515 | 0.266 | 1.26E-07 | 0 M1 _ | Pxdn |
| Sobp | 6.16E-12 | 0.304913 | 0.491 | 0.242 | 1.24E-07 | 0 M1 _ | Sobp |
| Ube4b | 5.97E-12 | 0.288518 | 0.497 | 0.253 | 1.20E-07 | 0 M1 _ | Ube4b |
| Sod1 | 5.76E-12 | 0.320209 | 0.796 | 0.514 | 1.16E-07 | 0 M1 _ | Sod1 |
| Sqor | 5.75E-12 | 0.282899 | 0.431 | 0.203 | 1.16E-07 | 0 M1 _ | Sqor |
| Rgma | 5.63E-12 | 0.386839 | 0.617 | 0.361 | 1.13E-07 | 0 M1 _ | Rgma |
| Acap2 | 5.46E-12 | 0.380091 | 0.808 | 0.542 | 1.10E-07 | 0 M1 _ | Acap2 |
| Cdk14 | 5.44E-12 | 0.365532 | 0.683 | 0.401 | 1.10E-07 | 0 M1 _ | Cdk14 |
| Tmem263 | 5.31E-12 | 0.367657 | 0.581 | 0.307 | 1.07E-07 | 0 M1 _ | Tmem263 |
| Smarcd2 | 4.81E-12 | 0.272776 | 0.467 | 0.227 | 9.68E-08 | 0 M1 _ | Smarcd2 |
| Rpn1 | 4.50E-12 | 0.386729 | 0.844 | 0.609 | 9.06E-08 | 0 M1 _ | Rpn1 |
| Chpf | 4.08E-12 | 0.392625 | 0.754 | 0.456 | 8.23E-08 | 0 M1 _ | Chpf |
| Tiparp | 3.93E-12 | 0.276135 | 0.401 | 0.18 | 7.92E-08 | 0 M1 _ | Tiparp |
| Tmem106b | 3.88E-12 | 0.337263 | 0.701 | 0.418 | 7.82E-08 | 0 M1 _ | Tmem106b |
| Cd81 | 3.86E-12 | 0.417246 | 0.994 | 0.889 | 7.78E-08 | 0 M1 _ | Cd81 |
| Gulp1 | 3.72E-12 | 0.294679 | 0.629 | 0.347 | 7.49E-08 | 0 M1 _ | Gulp1 |
| Sgpl1 | 2.85E-12 | 0.283117 | 0.461 | 0.216 | 5.74E-08 | 0 M1 _ | Sgpl1 |
| Cyth3 | 2.74E-12 | 0.385909 | 0.922 | 0.652 | 5.51E-08 | 0 M1 _ | Cyth3 |
| Ank3 | 2.60E-12 | 0.374337 | 0.527 | 0.271 | 5.23E-08 | 0 M1 _ | Ank3 |
| Fkbp10 | 2.53E-12 | 0.258411 | 0.569 | 0.287 | 5.09E-08 | 0 M1 _ | Fkbp10 |
| Tmem184c | 2.40E-12 | 0.263904 | 0.377 | 0.16 | 4.83E-08 | 0 M1 _ | Tmem184c |
| Irs2 | 2.27E-12 | 0.282177 | 0.419 | 0.189 | 4.58E-08 | 0 M1 _ | Irs2 |
| Selenot | 2.12E-12 | 0.305731 | 0.671 | 0.371 | 4.28E-08 | 0 M1 _ | Selenot |
| Sgce | 2.01E-12 | 0.336368 | 0.689 | 0.404 | 4.05E-08 | 0 M1 _ | Sgce |
| Emp3 | 2.01E-12 | 0.437544 | 0.964 | 0.794 | 4.04E-08 | 0 M1 _ | Emp3 |
| Ccdc8 | 1.97E-12 | 0.264549 | 0.335 | 0.136 | 3.98E-08 | 0 M1 _ | Ccdc8 |
| Itgav | 1.84E-12 | 0.331048 | 0.766 | 0.462 | 3.71E-08 | 0 M1 _ | Itgav |
| Map1a | 1.51E-12 | 0.303866 | 0.503 | 0.245 | 3.04E-08 | 0 M1 _ | Map1a |
| Arl8b | 1.49E-12 | 0.32389 | 0.593 | 0.311 | 3.01E-08 | 0 M1 _ | Arl8b |
| Aff1 | 1.49E-12 | 0.316616 | 0.449 | 0.206 | 2.99E-08 | 0 M1 _ | Aff1 |
| Notch2 | 1.48E-12 | 0.26938 | 0.437 | 0.196 | 2.99E-08 | 0 M1 _ | Notch2 |
| Ifi211 | 1.37E-12 | 0.428869 | 0.605 | 0.342 | 2.76E-08 | 0 M1 _ | Ifi211 |
| Plekhf1 | 1.20E-12 | 0.309941 | 0.485 | 0.237 | 2.42E-08 | 0 M1 _ | Plekhf1 |
| Rgs7bp | 1.15E-12 | 0.282562 | 0.365 | 0.148 | 2.33E-08 | 0 M1 _ | Rgs7bp |
| Arhgap24 | 1.13E-12 | 0.350727 | 0.425 | 0.195 | 2.27E-08 | 0 M1 _ | Arhgap24 |
| Ccl8 | 1.03E-12 | 0.383702 | 0.287 | 0.105 | 2.07E-08 | 0 M1 _ | Ccl8 |
| Fkbp7 | 9.63E-13 | 0.447613 | 0.85 | 0.546 | 1.94E-08 | 0 M1 _ | Fkbp7 |
| Mn1 | 9.20E-13 | 0.279354 | 0.287 | 0.103 | 1.85E-08 | 0 M1 _ | Mn1 |
| Cbx3 | 9.07E-13 | 0.409879 | 0.934 | 0.716 | 1.83E-08 | 0 M1 _ | Cbx3 |
| Slc43a3 | 8.32E-13 | 0.307814 | 0.784 | 0.488 | 1.68E-08 | 0 M1 _ | Slc43a3 |

| | | | | | | | |
|---|---|---|---|---|---|---|---|
| Ptp4a2 | 8.20E-13 | 0.439022 | 0.88 | 0.703 | 1.65E-08 | 0 M1 _ | Ptp4a2 |
| Copb2 | 8.10E-13 | 0.468214 | 0.79 | 0.545 | 1.63E-08 | 0 M1 _ | Copb2 |
| Kdelr2 | 6.70E-13 | 0.427796 | 0.874 | 0.551 | 1.35E-08 | 0 M1 _ | Kdelr2 |
| Zadh2 | 6.51E-13 | 0.259905 | 0.467 | 0.216 | 1.31E-08 | 0 M1 _ | Zadh2 |
| Spry1 | 5.80E-13 | 0.430363 | 0.808 | 0.517 | 1.17E-08 | 0 M1 _ | Spry1 |
| Man1a | 5.77E-13 | 0.485865 | 0.581 | 0.32 | 1.16E-08 | 0 M1 _ | Man1a |
| Fgd4 | 5.50E-13 | 0.286645 | 0.365 | 0.148 | 1.11E-08 | 0 M1 _ | Fgd4 |
| Sgcb | 5.42E-13 | 0.325985 | 0.605 | 0.317 | 1.09E-08 | 0 M1 _ | Sgcb |
| Rev3l | 5.28E-13 | 0.261538 | 0.479 | 0.219 | 1.06E-08 | 0 M1 _ | Rev3l |
| Tlr4 | 4.44E-13 | 0.252348 | 0.293 | 0.105 | 8.95E-09 | 0 M1 _ | Tlr4 |
| S1pr2 | 4.32E-13 | 0.271943 | 0.491 | 0.232 | 8.70E-09 | 0 M1 _ | S1pr2 |
| Sult5a1 | 4.24E-13 | 0.35806 | 0.353 | 0.142 | 8.54E-09 | 0 M1 _ | Sult5a1 |
| Thbd | 4.10E-13 | 0.703324 | 0.641 | 0.399 | 8.25E-09 | 0 M1 _ | Thbd |
| Lama4 | 4.08E-13 | 0.49543 | 0.79 | 0.5 | 8.22E-09 | 0 M1 _ | Lama4 |
| Fam114a1 | 4.08E-13 | 0.41626 | 0.737 | 0.444 | 8.21E-09 | 0 M1 _ | Fam114a1 |
| Klhl13 | 3.95E-13 | 0.262897 | 0.347 | 0.137 | 7.96E-09 | 0 M1 _ | Klhl13 |
| Lrrc58 | 3.93E-13 | 0.449355 | 0.904 | 0.655 | 7.91E-09 | 0 M1 _ | Lrrc58 |
| Glis2 | 2.70E-13 | 0.279854 | 0.431 | 0.189 | 5.43E-09 | 0 M1 _ | Glis2 |
| Tmem63a | 2.64E-13 | 0.293424 | 0.515 | 0.24 | 5.32E-09 | 0 M1 _ | Tmem63a |
| Slfn5 | 2.32E-13 | 0.397785 | 0.916 | 0.701 | 4.68E-09 | 0 M1 _ | Slfn5 |
| Ltbp3 | 2.03E-13 | 0.365602 | 0.754 | 0.419 | 4.09E-09 | 0 M1 _ | Ltbp3 |
| Selenon | 1.96E-13 | 0.329482 | 0.539 | 0.265 | 3.95E-09 | 0 M1 _ | Selenon |
| Tgoln1 | 1.89E-13 | 0.395542 | 0.82 | 0.548 | 3.81E-09 | 0 M1 _ | Tgoln1 |
| Setd7 | 1.89E-13 | 0.348049 | 0.569 | 0.294 | 3.81E-09 | 0 M1 _ | Setd7 |
| Chp1 | 1.85E-13 | 0.393323 | 0.491 | 0.249 | 3.72E-09 | 0 M1 _ | Chp1 |
| Zfp592 | 1.74E-13 | 0.286678 | 0.419 | 0.184 | 3.51E-09 | 0 M1 _ | Zfp592 |
| Serping1 | 1.66E-13 | 0.484398 | 0.988 | 0.74 | 3.35E-09 | 0 M1 _ | Serping1 |
| Vim | 1.62E-13 | 0.562593 | 0.988 | 0.877 | 3.27E-09 | 0 M1 _ | Vim |
| Pank3 | 1.59E-13 | 0.289248 | 0.491 | 0.224 | 3.20E-09 | 0 M1 _ | Pank3 |
| Lamp2 | 1.41E-13 | 0.412471 | 0.97 | 0.738 | 2.85E-09 | 0 M1 _ | Lamp2 |
| Mab21l2 | 1.41E-13 | 0.288772 | 0.275 | 0.095 | 2.85E-09 | 0 M1 _ | Mab21l2 |
| Pttg1ip | 1.32E-13 | 0.396416 | 0.898 | 0.634 | 2.66E-09 | 0 M1 _ | Pttg1ip |
| Runx1t1 | 1.29E-13 | 0.39732 | 0.569 | 0.295 | 2.60E-09 | 0 M1 _ | Runx1t1 |
| Col23a1 | 1.28E-13 | 0.313127 | 0.317 | 0.118 | 2.57E-09 | 0 M1 _ | Col23a1 |
| Ywhag | 1.21E-13 | 0.325917 | 0.635 | 0.346 | 2.43E-09 | 0 M1 _ | Ywhag |
| Cped1 | 1.03E-13 | 0.435527 | 0.952 | 0.68 | 2.07E-09 | 0 M1 _ | Cped1 |
| Itpripl2 | 8.80E-14 | 0.417359 | 0.802 | 0.545 | 1.77E-09 | 0 M1 _ | Itpripl2 |
| Fmnl2 | 8.78E-14 | 0.359235 | 0.539 | 0.271 | 1.77E-09 | 0 M1 _ | Fmnl2 |
| Itga8 | 8.76E-14 | 0.276825 | 0.443 | 0.185 | 1.76E-09 | 0 M1 _ | Itga8 |
| Stt3a | 8.72E-14 | 0.382967 | 0.743 | 0.451 | 1.76E-09 | 0 M1 _ | Stt3a |
| Mlec | 8.59E-14 | 0.408515 | 0.808 | 0.518 | 1.73E-09 | 0 M1 _ | Mlec |

| | | | | | | | |
|---|---|---|---|---|---|---|---|
| Hunk | 8.27E-14 | 0.289456 | 0.222 | 0.064 | 1.67E-09 | 0 M1 _ | Hunk |
| Serpinf1 | 7.60E-14 | 0.584958 | 0.988 | 0.568 | 1.53E-09 | 0 M1 _ | Serpinf1 |
| Dsel | 7.38E-14 | 0.360749 | 0.581 | 0.299 | 1.49E-09 | 0 M1 _ | Dsel |
| Dpy19l1 | 7.02E-14 | 0.356736 | 0.569 | 0.294 | 1.41E-09 | 0 M1 _ | Dpy19l1 |
| Prelp | 6.28E-14 | 0.423839 | 0.862 | 0.506 | 1.26E-09 | 0 M1 _ | Prelp |
| Olfml2b | 5.97E-14 | 0.269004 | 0.329 | 0.116 | 1.20E-09 | 0 M1 _ | Olfml2b |
| Cdh19 | 5.82E-14 | 0.250078 | 0.353 | 0.133 | 1.17E-09 | 0 M1 _ | Cdh19 |
| S100a6 | 4.80E-14 | 0.486608 | 0.994 | 0.923 | 9.67E-10 | 0 M1 _ | S100a6 |
| Morf4l2 | 4.62E-14 | 0.427786 | 0.844 | 0.568 | 9.30E-10 | 0 M1 _ | Morf4l2 |
| S100a10 | 4.39E-14 | 0.555504 | 0.94 | 0.674 | 8.84E-10 | 0 M1 _ | S100a10 |
| Adcy5 | 3.55E-14 | 0.277437 | 0.329 | 0.121 | 7.16E-10 | 0 M1 _ | Adcy5 |
| Tpcn1 | 3.40E-14 | 0.279887 | 0.461 | 0.202 | 6.84E-10 | 0 M1 _ | Tpcn1 |
| Sh3pxd2a | 3.38E-14 | 0.411843 | 0.683 | 0.387 | 6.81E-10 | 0 M1 _ | Sh3pxd2a |
| Abl1 | 3.37E-14 | 0.269887 | 0.521 | 0.242 | 6.79E-10 | 0 M1 _ | Abl1 |
| C1qtnf1 | 3.11E-14 | 0.252259 | 0.389 | 0.15 | 6.26E-10 | 0 M1 _ | C1qtnf1 |
| Tslp | 2.98E-14 | 0.307799 | 0.347 | 0.132 | 6.01E-10 | 0 M1 _ | Tslp |
| Txndc5 | 2.86E-14 | 0.370994 | 0.76 | 0.428 | 5.76E-10 | 0 M1 _ | Txndc5 |
| Zfhx4 | 2.58E-14 | 0.415778 | 0.701 | 0.391 | 5.20E-10 | 0 M1 _ | Zfhx4 |
| Extl3 | 2.51E-14 | 0.350406 | 0.587 | 0.301 | 5.05E-10 | 0 M1 _ | Extl3 |
| Tmem123 | 2.48E-14 | 0.3303 | 0.635 | 0.338 | 4.99E-10 | 0 M1 _ | Tmem123 |
| Aldh7a1 | 2.20E-14 | 0.459823 | 0.593 | 0.331 | 4.42E-10 | 0 M1 _ | Aldh7a1 |
| Adamtsl1 | 2.06E-14 | 0.515243 | 0.617 | 0.325 | 4.15E-10 | 0 M1 _ | Adamtsl1 |
| Lrp3 | 2.04E-14 | 0.287521 | 0.443 | 0.183 | 4.11E-10 | 0 M1 _ | Lrp3 |
| Asph | 1.92E-14 | 0.350842 | 0.701 | 0.379 | 3.88E-10 | 0 M1 _ | Asph |
| Samd4b | 1.89E-14 | 0.2846 | 0.599 | 0.286 | 3.81E-10 | 0 M1 _ | Samd4b |
| Cbr3 | 1.78E-14 | 0.452273 | 0.629 | 0.34 | 3.59E-10 | 0 M1 _ | Cbr3 |
| Obsl1 | 1.77E-14 | 0.264889 | 0.353 | 0.135 | 3.56E-10 | 0 M1 _ | Obsl1 |
| Basp1 | 1.57E-14 | 0.461535 | 0.665 | 0.355 | 3.17E-10 | 0 M1 _ | Basp1 |
| Fstl1 | 1.35E-14 | 0.524186 | 0.982 | 0.678 | 2.72E-10 | 0 M1 _ | Fstl1 |
| Tshz1 | 1.30E-14 | 0.338861 | 0.497 | 0.23 | 2.61E-10 | 0 M1 _ | Tshz1 |
| Vangl1 | 1.23E-14 | 0.262382 | 0.515 | 0.234 | 2.48E-10 | 0 M1 _ | Vangl1 |
| Flrt2 | 1.22E-14 | 0.264131 | 0.725 | 0.354 | 2.45E-10 | 0 M1 _ | Flrt2 |
| Idh1 | 1.21E-14 | 0.432294 | 0.617 | 0.332 | 2.44E-10 | 0 M1 _ | Idh1 |
| Ckap4 | 8.96E-15 | 0.445442 | 0.832 | 0.501 | 1.80E-10 | 0 M1 _ | Ckap4 |
| Bicc1 | 8.88E-15 | 0.454865 | 0.952 | 0.571 | 1.79E-10 | 0 M1 _ | Bicc1 |
| 8-Sep | 7.69E-15 | 0.388572 | 0.689 | 0.37 | 1.55E-10 | 0 M1 _ | 8-Sep |
| Ifi27 | 7.32E-15 | 0.498041 | 0.946 | 0.785 | 1.47E-10 | 0 M1 _ | Ifi27 |
| Rgmb | 6.98E-15 | 0.368858 | 0.575 | 0.285 | 1.41E-10 | 0 M1 _ | Rgmb |
| Dock7 | 6.98E-15 | 0.327254 | 0.617 | 0.303 | 1.40E-10 | 0 M1 _ | Dock7 |
| Myh10 | 6.73E-15 | 0.424193 | 0.479 | 0.218 | 1.36E-10 | 0 M1 _ | Myh10 |
| Efs | 5.99E-15 | 0.270888 | 0.293 | 0.098 | 1.21E-10 | 0 M1 _ | Efs |

| | | | | | | | |
|---|---|---|---|---|---|---|---|
| Tmem106a | 5.58E-15 | 0.311465 | 0.365 | 0.14 | 1.12E-10 | 0 M1 _ | Tmem106a |
| Jade1 | 4.62E-15 | 0.425283 | 0.617 | 0.331 | 9.30E-11 | 0 M1 _ | Jade1 |
| Lima1 | 4.40E-15 | 0.552748 | 0.868 | 0.59 | 8.86E-11 | 0 M1 _ | Lima1 |
| Fads1 | 3.09E-15 | 0.441855 | 0.569 | 0.294 | 6.23E-11 | 0 M1 _ | Fads1 |
| Lpgat1 | 2.45E-15 | 0.37922 | 0.611 | 0.303 | 4.94E-11 | 0 M1 _ | Lpgat1 |
| Ergic1 | 2.13E-15 | 0.375452 | 0.581 | 0.291 | 4.28E-11 | 0 M1 _ | Ergic1 |
| Impad1 | 2.02E-15 | 0.396051 | 0.665 | 0.35 | 4.06E-11 | 0 M1 _ | Impad1 |
| Fkbp9 | 1.97E-15 | 0.416163 | 0.856 | 0.508 | 3.96E-11 | 0 M1 _ | Fkbp9 |
| Ski | 1.96E-15 | 0.442536 | 0.838 | 0.539 | 3.94E-11 | 0 M1 _ | Ski |
| Rbfox2 | 1.96E-15 | 0.390652 | 0.689 | 0.377 | 3.94E-11 | 0 M1 _ | Rbfox2 |
| Rnf145 | 1.82E-15 | 0.295973 | 0.449 | 0.186 | 3.67E-11 | 0 M1 _ | Rnf145 |
| Pcolce | 1.75E-15 | 0.543779 | 0.988 | 0.736 | 3.52E-11 | 0 M1 _ | Pcolce |
| Serinc1 | 1.57E-15 | 0.502372 | 0.922 | 0.709 | 3.16E-11 | 0 M1 _ | Serinc1 |
| Apod | 1.55E-15 | 0.39124 | 0.982 | 0.625 | 3.12E-11 | 0 M1 _ | Apod |
| Fkbp14 | 1.55E-15 | 0.312177 | 0.455 | 0.193 | 3.12E-11 | 0 M1 _ | Fkbp14 |
| Dnajc10 | 1.28E-15 | 0.40979 | 0.808 | 0.471 | 2.58E-11 | 0 M1 _ | Dnajc10 |
| Rhobtb1 | 1.17E-15 | 0.259422 | 0.275 | 0.085 | 2.35E-11 | 0 M1 _ | Rhobtb1 |
| Prrx2 | 1.11E-15 | 0.397816 | 0.617 | 0.3 | 2.23E-11 | 0 M1 _ | Prrx2 |
| Nectin1 | 1.04E-15 | 0.317282 | 0.365 | 0.136 | 2.09E-11 | 0 M1 _ | Nectin1 |
| Irf2bp2 | 9.41E-16 | 0.54036 | 0.856 | 0.568 | 1.90E-11 | 0 M1 _ | Irf2bp2 |
| Reep5 | 8.33E-16 | 0.525141 | 0.97 | 0.727 | 1.68E-11 | 0 M1 _ | Reep5 |
| Abca1 | 7.32E-16 | 0.45805 | 0.665 | 0.348 | 1.47E-11 | 0 M1 _ | Abca1 |
| Irak1 | 6.35E-16 | 0.446384 | 0.683 | 0.385 | 1.28E-11 | 0 M1 _ | Irak1 |
| Nav3 | 5.36E-16 | 0.411328 | 0.647 | 0.323 | 1.08E-11 | 0 M1 _ | Nav3 |
| Col27a1 | 5.02E-16 | 0.477552 | 0.617 | 0.316 | 1.01E-11 | 0 M1 _ | Col27a1 |
| Prrx1 | 4.64E-16 | 0.522448 | 0.958 | 0.7 | 9.36E-12 | 0 M1 _ | Prrx1 |
| Pik3r2 | 3.91E-16 | 0.334824 | 0.521 | 0.232 | 7.88E-12 | 0 M1 _ | Pik3r2 |
| Gfpt2 | 3.85E-16 | 0.48297 | 0.599 | 0.301 | 7.75E-12 | 0 M1 _ | Gfpt2 |
| Twist1 | 3.40E-16 | 0.474043 | 0.784 | 0.457 | 6.86E-12 | 0 M1 _ | Twist1 |
| Aoc3 | 2.81E-16 | 0.254876 | 0.473 | 0.183 | 5.67E-12 | 0 M1 _ | Aoc3 |
| Arhgap20 | 2.73E-16 | 0.44008 | 0.509 | 0.225 | 5.50E-12 | 0 M1 _ | Arhgap20 |
| Boc | 2.70E-16 | 0.385015 | 0.461 | 0.194 | 5.44E-12 | 0 M1 _ | Boc |
| Oat | 2.33E-16 | 0.453761 | 0.952 | 0.621 | 4.70E-12 | 0 M1 _ | Oat |
| Ppp3ca | 2.06E-16 | 0.520259 | 0.784 | 0.497 | 4.14E-12 | 0 M1 _ | Ppp3ca |
| Naaa | 1.78E-16 | 0.347354 | 0.335 | 0.116 | 3.59E-12 | 0 M1 _ | Naaa |
| Olfml2a | 1.77E-16 | 0.397567 | 0.425 | 0.172 | 3.57E-12 | 0 M1 _ | Olfml2a |
| Tuba1a | 1.74E-16 | 0.508306 | 0.976 | 0.819 | 3.50E-12 | 0 M1 _ | Tuba1a |
| Slc29a1 | 1.58E-16 | 0.582896 | 0.862 | 0.596 | 3.19E-12 | 0 M1 _ | Slc29a1 |
| Vldlr | 1.52E-16 | 0.31019 | 0.228 | 0.058 | 3.05E-12 | 0 M1 _ | Vldlr |
| Mt1 | 1.45E-16 | 0.820666 | 0.952 | 0.6 | 2.91E-12 | 0 M1 _ | Mt1 |
| Slc16a2 | 1.26E-16 | 0.458466 | 0.599 | 0.297 | 2.53E-12 | 0 M1 _ | Slc16a2 |

| | | | | | | | |
|---|---|---|---|---|---|---|---|
| Sec61a1 | 1.17E-16 | 0.481242 | 0.778 | 0.476 | 2.36E-12 | 0 M1 _ | Sec61a1 |
| Eps15 | 1.13E-16 | 0.444044 | 0.653 | 0.343 | 2.28E-12 | 0 M1 _ | Eps15 |
| Por | 1.13E-16 | 0.431895 | 0.653 | 0.329 | 2.27E-12 | 0 M1 _ | Por |
| Sec24d | 9.99E-17 | 0.375398 | 0.611 | 0.301 | 2.01E-12 | 0 M1 _ | Sec24d |
| Fbln1 | 9.89E-17 | 0.593274 | 0.916 | 0.506 | 1.99E-12 | 0 M1 _ | Fbln1 |
| Igf2r | 8.37E-17 | 0.614263 | 0.695 | 0.405 | 1.68E-12 | 0 M1 _ | Igf2r |
| Trabd2b | 7.33E-17 | 0.263189 | 0.323 | 0.101 | 1.48E-12 | 0 M1 _ | Trabd2b |
| Cd44 | 6.60E-17 | 0.382242 | 0.455 | 0.187 | 1.33E-12 | 0 M1 _ | Cd44 |
| Stat5b | 6.49E-17 | 0.3112 | 0.479 | 0.195 | 1.31E-12 | 0 M1 _ | Stat5b |
| Ptgir | 6.40E-17 | 0.399219 | 0.467 | 0.194 | 1.29E-12 | 0 M1 _ | Ptgir |
| Cdr2l | 5.05E-17 | 0.264384 | 0.431 | 0.162 | 1.02E-12 | 0 M1 _ | Cdr2l |
| Arhgap21 | 3.65E-17 | 0.420579 | 0.491 | 0.211 | 7.35E-13 | 0 M1 _ | Arhgap21 |
| Dmpk | 3.65E-17 | 0.275804 | 0.323 | 0.1 | 7.35E-13 | 0 M1 _ | Dmpk |
| Ssr3 | 2.82E-17 | 0.486196 | 0.874 | 0.559 | 5.69E-13 | 0 M1 _ | Ssr3 |
| Tppp3 | 2.77E-17 | 1.211634 | 0.413 | 0.176 | 5.58E-13 | 0 M1 _ | Tppp3 |
| Ahnak | 2.48E-17 | 0.580946 | 0.988 | 0.888 | 5.00E-13 | 0 M1 _ | Ahnak |
| Sar1a | 2.43E-17 | 0.509941 | 0.88 | 0.567 | 4.90E-13 | 0 M1 _ | Sar1a |
| Col6a6 | 2.40E-17 | 0.731964 | 0.605 | 0.301 | 4.82E-13 | 0 M1 _ | Col6a6 |
| Tram1 | 2.16E-17 | 0.514602 | 0.88 | 0.582 | 4.34E-13 | 0 M1 _ | Tram1 |
| Fkbp2 | 2.08E-17 | 0.531936 | 0.856 | 0.548 | 4.20E-13 | 0 M1 _ | Fkbp2 |
| Tubb6 | 2.00E-17 | 0.451903 | 0.617 | 0.298 | 4.03E-13 | 0 M1 _ | Tubb6 |
| Fam111a | 1.71E-17 | 0.361044 | 0.539 | 0.234 | 3.44E-13 | 0 M1 _ | Fam111a |
| Vwa5a | 1.62E-17 | 0.372517 | 0.653 | 0.33 | 3.26E-13 | 0 M1 _ | Vwa5a |
| Ftl1 | 1.48E-17 | 0.448882 | 1 | 0.96 | 2.99E-13 | 0 M1 _ | Ftl1 |
| Arrdc3 | 1.39E-17 | 0.376407 | 0.509 | 0.21 | 2.80E-13 | 0 M1 _ | Arrdc3 |
| Hook3 | 1.31E-17 | 0.499406 | 0.916 | 0.633 | 2.64E-13 | 0 M1 _ | Hook3 |
| Pam | 1.27E-17 | 0.650834 | 0.808 | 0.503 | 2.56E-13 | 0 M1 _ | Pam |
| Golph3 | 1.22E-17 | 0.407698 | 0.659 | 0.322 | 2.45E-13 | 0 M1 _ | Golph3 |
| Pdia4 | 9.29E-18 | 0.513024 | 0.713 | 0.399 | 1.87E-13 | 0 M1 _ | Pdia4 |
| Il1r1 | 9.22E-18 | 0.507709 | 0.79 | 0.434 | 1.86E-13 | 0 M1 _ | Il1r1 |
| Krt10 | 9.14E-18 | 0.287841 | 0.401 | 0.144 | 1.84E-13 | 0 M1 _ | Krt10 |
| Slc39a1 | 8.83E-18 | 0.509465 | 0.743 | 0.426 | 1.78E-13 | 0 M1 _ | Slc39a1 |
| Aldh1a1 | 8.54E-18 | 0.35464 | 0.683 | 0.322 | 1.72E-13 | 0 M1 _ | Aldh1a1 |
| Psap | 7.66E-18 | 0.460347 | 0.982 | 0.848 | 1.54E-13 | 0 M1 _ | Psap |
| Eps8 | 6.71E-18 | 0.43364 | 0.743 | 0.403 | 1.35E-13 | 0 M1 _ | Eps8 |
| Pmp22 | 5.42E-18 | 0.523955 | 0.94 | 0.69 | 1.09E-13 | 0 M1 _ | Pmp22 |
| Tgfbi | 4.07E-18 | 0.818401 | 0.754 | 0.413 | 8.21E-14 | 0 M1 _ | Tgfbi |
| Canx | 3.40E-18 | 0.538714 | 0.934 | 0.721 | 6.85E-14 | 0 M1 _ | Canx |
| mt-Nd4l | 3.19E-18 | 0.588582 | 0.97 | 0.817 | 6.43E-14 | 0 M1 _ | mt-Nd4l |
| Abca6 | 2.91E-18 | 0.298158 | 0.341 | 0.108 | 5.86E-14 | 0 M1 _ | Abca6 |
| Cd9 | 2.50E-18 | 0.60522 | 0.946 | 0.708 | 5.03E-14 | 0 M1 _ | Cd9 |

| | | | | | | | |
|---|---|---|---|---|---|---|---|
| Foxo3 | 2.05E-18 | 0.309733 | 0.509 | 0.207 | 4.12E-14 | 0 M1 _ | Foxo3 |
| Txn1 | 1.91E-18 | 0.560895 | 0.958 | 0.824 | 3.85E-14 | 0 M1 _ | Txn1 |
| Surf4 | 1.75E-18 | 0.553466 | 0.832 | 0.528 | 3.52E-14 | 0 M1 _ | Surf4 |
| Dpysl2 | 1.61E-18 | 0.476748 | 0.778 | 0.456 | 3.25E-14 | 0 M1 _ | Dpysl2 |
| Plod1 | 1.60E-18 | 0.546245 | 0.844 | 0.525 | 3.22E-14 | 0 M1 _ | Plod1 |
| Atp2a2 | 1.54E-18 | 0.499217 | 0.934 | 0.665 | 3.11E-14 | 0 M1 _ | Atp2a2 |
| Mt2 | 1.34E-18 | 0.943851 | 0.677 | 0.32 | 2.69E-14 | 0 M1 _ | Mt2 |
| Dse | 9.38E-19 | 0.432197 | 0.581 | 0.262 | 1.89E-14 | 0 M1 _ | Dse |
| Adamts1 | 7.57E-19 | 0.645582 | 0.659 | 0.319 | 1.52E-14 | 0 M1 _ | Adamts1 |
| Armcx2 | 6.74E-19 | 0.347099 | 0.491 | 0.195 | 1.36E-14 | 0 M1 _ | Armcx2 |
| Bmpr1a | 6.64E-19 | 0.518404 | 0.695 | 0.355 | 1.34E-14 | 0 M1 _ | Bmpr1a |
| Cdon | 6.63E-19 | 0.347749 | 0.317 | 0.094 | 1.34E-14 | 0 M1 _ | Cdon |
| Pnp | 6.32E-19 | 0.612799 | 0.862 | 0.562 | 1.27E-14 | 0 M1 _ | Pnp |
| Marcks | 6.15E-19 | 0.613721 | 0.988 | 0.811 | 1.24E-14 | 0 M1 _ | Marcks |
| Tmem119 | 6.10E-19 | 0.595329 | 0.707 | 0.361 | 1.23E-14 | 0 M1 _ | Tmem119 |
| Zhx3 | 5.54E-19 | 0.468094 | 0.635 | 0.308 | 1.12E-14 | 0 M1 _ | Zhx3 |
| Colec12 | 5.33E-19 | 0.581703 | 0.952 | 0.544 | 1.07E-14 | 0 M1 _ | Colec12 |
| Col5a2 | 5.06E-19 | 0.617845 | 0.97 | 0.628 | 1.02E-14 | 0 M1 _ | Col5a2 |
| Hikeshi | 4.77E-19 | 0.443394 | 0.689 | 0.362 | 9.60E-15 | 0 M1 _ | Hikeshi |
| Nxf1 | 4.76E-19 | 0.402334 | 0.575 | 0.253 | 9.58E-15 | 0 M1 _ | Nxf1 |
| Il6st | 3.53E-19 | 0.528713 | 0.856 | 0.558 | 7.12E-15 | 0 M1 _ | Il6st |
| Nr2f1 | 3.41E-19 | 0.669129 | 0.886 | 0.579 | 6.87E-15 | 0 M1 _ | Nr2f1 |
| Tbc1d2b | 2.96E-19 | 0.417185 | 0.689 | 0.337 | 5.96E-15 | 0 M1 _ | Tbc1d2b |
| Lrp6 | 2.62E-19 | 0.536027 | 0.754 | 0.433 | 5.29E-15 | 0 M1 _ | Lrp6 |
| Glg1 | 2.57E-19 | 0.534134 | 0.916 | 0.637 | 5.18E-15 | 0 M1 _ | Glg1 |
| Dclk1 | 2.51E-19 | 0.60264 | 0.491 | 0.2 | 5.06E-15 | 0 M1 _ | Dclk1 |
| Col3a1 | 2.44E-19 | 0.673313 | 0.988 | 0.74 | 4.91E-15 | 0 M1 _ | Col3a1 |
| Npas2 | 2.31E-19 | 0.255196 | 0.341 | 0.104 | 4.65E-15 | 0 M1 _ | Npas2 |
| Sash1 | 2.29E-19 | 0.559735 | 0.844 | 0.559 | 4.60E-15 | 0 M1 _ | Sash1 |
| Dcun1d3 | 2.24E-19 | 0.394106 | 0.347 | 0.112 | 4.52E-15 | 0 M1 _ | Dcun1d3 |
| Cd47 | 1.95E-19 | 0.504275 | 0.982 | 0.854 | 3.92E-15 | 0 M1 _ | Cd47 |
| Fgf7 | 1.52E-19 | 0.346523 | 0.551 | 0.221 | 3.05E-15 | 0 M1 _ | Fgf7 |
| Fap | 1.41E-19 | 0.501526 | 0.826 | 0.447 | 2.84E-15 | 0 M1 _ | Fap |
| Ssr1 | 1.11E-19 | 0.574286 | 0.814 | 0.505 | 2.23E-15 | 0 M1 _ | Ssr1 |
| Uap1 | 1.08E-19 | 0.456371 | 0.563 | 0.245 | 2.18E-15 | 0 M1 _ | Uap1 |
| Maged2 | 8.84E-20 | 0.546627 | 0.76 | 0.412 | 1.78E-15 | 0 M1 _ | Maged2 |
| Akap12 | 8.71E-20 | 0.744988 | 0.838 | 0.523 | 1.75E-15 | 0 M1 _ | Akap12 |
| Ssh2 | 7.17E-20 | 0.653575 | 0.766 | 0.458 | 1.44E-15 | 0 M1 _ | Ssh2 |
| Ncoa1 | 5.99E-20 | 0.422618 | 0.545 | 0.228 | 1.21E-15 | 0 M1 _ | Ncoa1 |
| Smad3 | 5.99E-20 | 0.389671 | 0.503 | 0.202 | 1.21E-15 | 0 M1 _ | Smad3 |
| Srgap1 | 5.64E-20 | 0.26431 | 0.251 | 0.06 | 1.14E-15 | 0 M1 _ | Srgap1 |

| | | | | | | | |
|---|---|---|---|---|---|---|---|
| Ecm1 | 4.24E-20 | 0.739364 | 0.784 | 0.461 | 8.53E-16 | 0 M1 _ | Ecm1 |
| Gpc3 | 3.12E-20 | 0.376386 | 0.281 | 0.07 | 6.29E-16 | 0 M1 _ | Gpc3 |
| Scarb2 | 2.41E-20 | 0.590698 | 0.856 | 0.521 | 4.86E-16 | 0 M1 _ | Scarb2 |
| Gnas | 2.25E-20 | 0.415036 | 1 | 0.959 | 4.53E-16 | 0 M1 _ | Gnas |
| Fam129a | 1.91E-20 | 0.529736 | 0.707 | 0.373 | 3.85E-16 | 0 M1 _ | Fam129a |
| Angptl4 | 1.57E-20 | 0.322589 | 0.275 | 0.068 | 3.16E-16 | 0 M1 _ | Angptl4 |
| Gdf10 | 1.32E-20 | 0.643176 | 0.341 | 0.103 | 2.67E-16 | 0 M1 _ | Gdf10 |
| Adgra2 | 1.00E-20 | 0.456788 | 0.563 | 0.237 | 2.02E-16 | 0 M1 _ | Adgra2 |
| Steap4 | 8.55E-21 | 0.27817 | 0.832 | 0.422 | 1.72E-16 | 0 M1 _ | Steap4 |
| Cat | 7.04E-21 | 0.522869 | 0.605 | 0.274 | 1.42E-16 | 0 M1 _ | Cat |
| Gja1 | 5.68E-21 | 0.640662 | 0.671 | 0.33 | 1.14E-16 | 0 M1 _ | Gja1 |
| Flnc | 5.36E-21 | 0.339696 | 0.347 | 0.103 | 1.08E-16 | 0 M1 _ | Flnc |
| Adamts5 | 3.65E-21 | 0.688397 | 0.904 | 0.497 | 7.35E-17 | 0 M1 _ | Adamts5 |
| Tmem198b | 3.05E-21 | 0.268886 | 0.335 | 0.096 | 6.15E-17 | 0 M1 _ | Tmem198b |
| Thbs1 | 3.01E-21 | 0.651933 | 0.557 | 0.231 | 6.07E-17 | 0 M1 _ | Thbs1 |
| Col1a2 | 2.35E-21 | 0.753453 | 0.994 | 0.733 | 4.73E-17 | 0 M1 _ | Col1a2 |
| Lifr | 2.27E-21 | 0.741901 | 0.707 | 0.389 | 4.58E-17 | 0 M1 _ | Lifr |
| G6pdx | 1.73E-21 | 0.335294 | 0.401 | 0.128 | 3.48E-17 | 0 M1 _ | G6pdx |
| Fth1 | 1.55E-21 | 0.550057 | 0.994 | 0.993 | 3.13E-17 | 0 M1 _ | Fth1 |
| Abca9 | 1.49E-21 | 0.561376 | 0.653 | 0.302 | 3.00E-17 | 0 M1 _ | Abca9 |
| Bmp1 | 1.44E-21 | 0.564674 | 0.766 | 0.418 | 2.91E-17 | 0 M1 _ | Bmp1 |
| Fgf18 | 1.25E-21 | 0.334461 | 0.305 | 0.081 | 2.52E-17 | 0 M1 _ | Fgf18 |
| Cpq | 1.23E-21 | 0.650541 | 0.82 | 0.434 | 2.47E-17 | 0 M1 _ | Cpq |
| Grb10 | 1.23E-21 | 0.631652 | 0.671 | 0.333 | 2.47E-17 | 0 M1 _ | Grb10 |
| Fam171b | 1.20E-21 | 0.466961 | 0.371 | 0.116 | 2.41E-17 | 0 M1 _ | Fam171b |
| Reck | 1.06E-21 | 0.426034 | 0.521 | 0.202 | 2.14E-17 | 0 M1 _ | Reck |
| Gstm2 | 7.38E-22 | 0.674125 | 0.796 | 0.464 | 1.49E-17 | 0 M1 _ | Gstm2 |
| Nfic | 7.14E-22 | 0.675465 | 0.958 | 0.699 | 1.44E-17 | 0 M1 _ | Nfic |
| Atrnl1 | 5.78E-22 | 0.395625 | 0.497 | 0.185 | 1.16E-17 | 0 M1 _ | Atrnl1 |
| Slc16a9 | 5.71E-22 | 0.340124 | 0.491 | 0.169 | 1.15E-17 | 0 M1 _ | Slc16a9 |
| Eif4g3 | 4.74E-22 | 0.679851 | 0.886 | 0.602 | 9.55E-18 | 0 M1 _ | Eif4g3 |
| Calr | 4.73E-22 | 0.638024 | 0.982 | 0.866 | 9.52E-18 | 0 M1 _ | Calr |
| Dennd2a | 4.64E-22 | 0.39944 | 0.581 | 0.235 | 9.34E-18 | 0 M1 _ | Dennd2a |
| H6pd | 3.83E-22 | 0.467997 | 0.641 | 0.278 | 7.72E-18 | 0 M1 _ | H6pd |
| Cacna1a | 3.58E-22 | 0.478431 | 0.497 | 0.189 | 7.21E-18 | 0 M1 _ | Cacna1a |
| Arhgap32 | 3.41E-22 | 0.463225 | 0.479 | 0.178 | 6.87E-18 | 0 M1 _ | Arhgap32 |
| Hgsnat | 3.32E-22 | 0.48698 | 0.683 | 0.327 | 6.69E-18 | 0 M1 _ | Hgsnat |
| Sdc2 | 3.03E-22 | 0.650443 | 0.91 | 0.536 | 6.11E-18 | 0 M1 _ | Sdc2 |
| Gstm1 | 1.63E-22 | 0.734664 | 0.91 | 0.647 | 3.29E-18 | 0 M1 _ | Gstm1 |
| P4hb | 1.54E-22 | 0.635424 | 0.952 | 0.676 | 3.10E-18 | 0 M1 _ | P4hb |
| Maged1 | 1.44E-22 | 0.604048 | 0.904 | 0.529 | 2.90E-18 | 0 M1 _ | Maged1 |

| | | | | | | | |
|---|---|---|---|---|---|---|---|
| Cyb5a | 1.33E-22 | 0.57813 | 0.982 | 0.756 | 2.68E-18 | 0 M1 _ | Cyb5a |
| Sh3d19 | 1.23E-22 | 0.621614 | 0.743 | 0.387 | 2.48E-18 | 0 M1 _ | Sh3d19 |
| Col6a1 | 9.27E-23 | 0.6889 | 0.982 | 0.693 | 1.87E-18 | 0 M1 _ | Col6a1 |
| Abcc4 | 9.22E-23 | 0.308773 | 0.323 | 0.084 | 1.86E-18 | 0 M1 _ | Abcc4 |
| Bmper | 9.20E-23 | 0.321729 | 0.269 | 0.061 | 1.85E-18 | 0 M1 _ | Bmper |
| Slit3 | 8.28E-23 | 0.538639 | 0.725 | 0.355 | 1.67E-18 | 0 M1 _ | Slit3 |
| Rhobtb3 | 3.80E-23 | 0.599995 | 0.719 | 0.353 | 7.66E-19 | 0 M1 _ | Rhobtb3 |
| Cxcl14 | 3.29E-23 | 1.108858 | 0.886 | 0.596 | 6.63E-19 | 0 M1 _ | Cxcl14 |
| 9330159F19Rik | 2.65E-23 | 0.29297 | 0.192 | 0.031 | 5.33E-19 | 0 M1 _ | 9330159F19Rik |
| Lgi2 | 1.77E-23 | 0.445149 | 0.371 | 0.106 | 3.57E-19 | 0 M1 _ | Lgi2 |
| Spag7 | 1.45E-23 | 0.35293 | 0.479 | 0.161 | 2.92E-19 | 0 M1 _ | Spag7 |
| Sema3a | 9.19E-24 | 0.495632 | 0.479 | 0.166 | 1.85E-19 | 0 M1 _ | Sema3a |
| Mxra8 | 7.85E-24 | 0.721046 | 0.946 | 0.6 | 1.58E-19 | 0 M1 _ | Mxra8 |
| Dpysl3 | 5.95E-24 | 0.682463 | 0.91 | 0.49 | 1.20E-19 | 0 M1 _ | Dpysl3 |
| Lgi4 | 5.08E-24 | 0.416336 | 0.413 | 0.133 | 1.02E-19 | 0 M1 _ | Lgi4 |
| Bcl6 | 4.54E-24 | 0.397417 | 0.437 | 0.14 | 9.14E-20 | 0 M1 _ | Bcl6 |
| Adamtsl3 | 3.91E-24 | 0.507769 | 0.371 | 0.108 | 7.88E-20 | 0 M1 _ | Adamtsl3 |
| Nfib | 3.85E-24 | 0.606699 | 0.958 | 0.746 | 7.76E-20 | 0 M1 _ | Nfib |
| Ifngr1 | 3.27E-24 | 0.772974 | 0.94 | 0.718 | 6.59E-20 | 0 M1 _ | Ifngr1 |
| Reps2 | 2.94E-24 | 0.355325 | 0.371 | 0.104 | 5.93E-20 | 0 M1 _ | Reps2 |
| Mrap | 2.46E-24 | 0.414314 | 0.311 | 0.076 | 4.95E-20 | 0 M1 _ | Mrap |
| Tshz3 | 2.36E-24 | 0.504554 | 0.605 | 0.246 | 4.75E-20 | 0 M1 _ | Tshz3 |
| Maob | 2.25E-24 | 0.270108 | 0.216 | 0.038 | 4.52E-20 | 0 M1 _ | Maob |
| Ank2 | 1.97E-24 | 0.529205 | 0.689 | 0.306 | 3.96E-20 | 0 M1 _ | Ank2 |
| Ccdc71l | 1.76E-24 | 0.509146 | 0.575 | 0.231 | 3.54E-20 | 0 M1 _ | Ccdc71l |
| Nav1 | 1.13E-24 | 0.750722 | 0.934 | 0.597 | 2.27E-20 | 0 M1 _ | Nav1 |
| Lpl | 1.01E-24 | 0.891074 | 0.982 | 0.662 | 2.03E-20 | 0 M1 _ | Lpl |
| Pcdh7 | 7.51E-25 | 0.655289 | 0.725 | 0.349 | 1.51E-20 | 0 M1 _ | Pcdh7 |
| Itm2a | 4.69E-25 | 0.846233 | 0.94 | 0.609 | 9.45E-21 | 0 M1 _ | Itm2a |
| Auts2 | 4.53E-25 | 0.716802 | 0.844 | 0.461 | 9.12E-21 | 0 M1 _ | Auts2 |
| AY036118 | 4.35E-25 | 1.356401 | 0.88 | 0.543 | 8.76E-21 | 0 M1 _ | AY036118 |
| G0s2 | 4.27E-25 | 0.689154 | 0.886 | 0.495 | 8.61E-21 | 0 M1 _ | G0s2 |
| Arl5a | 3.94E-25 | 0.609838 | 0.713 | 0.346 | 7.94E-21 | 0 M1 _ | Arl5a |
| Pdxk | 3.74E-25 | 0.357442 | 0.353 | 0.092 | 7.54E-21 | 0 M1 _ | Pdxk |
| Vcan | 2.96E-25 | 0.75262 | 0.659 | 0.281 | 5.95E-21 | 0 M1 _ | Vcan |
| St3gal1 | 2.57E-25 | 0.597696 | 0.551 | 0.211 | 5.19E-21 | 0 M1 _ | St3gal1 |
| Stt3b | 2.26E-25 | 0.633897 | 0.689 | 0.329 | 4.55E-21 | 0 M1 _ | Stt3b |
| Ifi204 | 2.05E-25 | 0.717766 | 0.707 | 0.332 | 4.13E-21 | 0 M1 _ | Ifi204 |
| Adcyap1r1 | 1.95E-25 | 0.489608 | 0.635 | 0.258 | 3.92E-21 | 0 M1 _ | Adcyap1r1 |
| Cnmd | 1.89E-25 | 0.628366 | 0.138 | 0.014 | 3.80E-21 | 0 M1 _ | Cnmd |
| Gm42418 | 1.66E-25 | 0.99898 | 1 | 0.999 | 3.35E-21 | 0 M1 _ | Gm42418 |

| | | | | | | | |
|---|---|---|---|---|---|---|---|
| Cyp2f2 | 1.59E-25 | 0.630694 | 0.18 | 0.025 | 3.19E-21 | 0 M1 _ | Cyp2f2 |
| Glipr2 | 1.41E-25 | 0.494936 | 0.437 | 0.137 | 2.85E-21 | 0 M1 _ | Glipr2 |
| F3 | 1.40E-25 | 0.643631 | 0.647 | 0.257 | 2.81E-21 | 0 M1 _ | F3 |
| Fbln2 | 1.39E-25 | 1.198837 | 0.838 | 0.524 | 2.80E-21 | 0 M1 _ | Fbln2 |
| Nfasc | 7.27E-26 | 0.351495 | 0.311 | 0.072 | 1.46E-21 | 0 M1 _ | Nfasc |
| Fcgr2b | 5.11E-26 | 0.797713 | 0.437 | 0.143 | 1.03E-21 | 0 M1 _ | Fcgr2b |
| Ctsh | 4.88E-26 | 0.7373 | 0.916 | 0.545 | 9.82E-22 | 0 M1 _ | Ctsh |
| Ramp2 | 3.78E-26 | 0.569597 | 0.874 | 0.469 | 7.61E-22 | 0 M1 _ | Ramp2 |
| Ltbp4 | 3.74E-26 | 0.862388 | 0.928 | 0.652 | 7.54E-22 | 0 M1 _ | Ltbp4 |
| Zim1 | 1.69E-26 | 0.341704 | 0.18 | 0.024 | 3.41E-22 | 0 M1 _ | Zim1 |
| Ptgfrn | 1.54E-26 | 0.720568 | 0.766 | 0.378 | 3.10E-22 | 0 M1 _ | Ptgfrn |
| Fez1 | 1.27E-26 | 0.331511 | 0.269 | 0.053 | 2.57E-22 | 0 M1 _ | Fez1 |
| Serpinh1 | 1.01E-26 | 0.76085 | 0.994 | 0.798 | 2.03E-22 | 0 M1 _ | Serpinh1 |
| Col6a2 | 5.34E-27 | 0.817646 | 0.976 | 0.643 | 1.07E-22 | 0 M1 _ | Col6a2 |
| Serpinb6a | 5.04E-27 | 0.750541 | 0.952 | 0.695 | 1.01E-22 | 0 M1 _ | Serpinb6a |
| Ggt5 | 4.89E-27 | 0.761635 | 0.713 | 0.337 | 9.84E-23 | 0 M1 _ | Ggt5 |
| Ctsl | 4.22E-27 | 0.745146 | 0.994 | 0.798 | 8.50E-23 | 0 M1 _ | Ctsl |
| Fkbp5 | 6.84E-28 | 0.622991 | 0.599 | 0.236 | 1.38E-23 | 0 M1 _ | Fkbp5 |
| Twsg1 | 6.64E-28 | 0.637399 | 0.743 | 0.352 | 1.34E-23 | 0 M1 _ | Twsg1 |
| Fhl1 | 5.35E-28 | 0.464144 | 0.293 | 0.059 | 1.08E-23 | 0 M1 _ | Fhl1 |
| Penk | 3.98E-28 | 0.477748 | 0.605 | 0.224 | 8.01E-24 | 0 M1 _ | Penk |
| Pde10a | 3.78E-28 | 0.555115 | 0.443 | 0.132 | 7.61E-24 | 0 M1 _ | Pde10a |
| Adam12 | 3.57E-28 | 0.275691 | 0.257 | 0.046 | 7.19E-24 | 0 M1 _ | Adam12 |
| Evc | 3.38E-28 | 0.40928 | 0.425 | 0.118 | 6.80E-24 | 0 M1 _ | Evc |
| Mme | 2.89E-28 | 0.593096 | 0.443 | 0.136 | 5.83E-24 | 0 M1 _ | Mme |
| Gpx3 | 2.54E-28 | 1.096251 | 0.76 | 0.364 | 5.11E-24 | 0 M1 _ | Gpx3 |
| Entpd2 | 1.88E-28 | 0.779327 | 0.934 | 0.517 | 3.80E-24 | 0 M1 _ | Entpd2 |
| Add3 | 1.61E-28 | 0.778424 | 0.844 | 0.483 | 3.24E-24 | 0 M1 _ | Add3 |
| Nedd4 | 1.42E-28 | 0.627193 | 0.988 | 0.882 | 2.86E-24 | 0 M1 _ | Nedd4 |
| Cd248 | 1.37E-28 | 0.715653 | 0.808 | 0.396 | 2.77E-24 | 0 M1 _ | Cd248 |
| Fbn1 | 1.32E-28 | 0.709245 | 0.97 | 0.596 | 2.66E-24 | 0 M1 _ | Fbn1 |
| Robo1 | 1.31E-28 | 0.51059 | 0.575 | 0.204 | 2.65E-24 | 0 M1 _ | Robo1 |
| Plekha6 | 8.07E-29 | 0.729359 | 0.731 | 0.336 | 1.62E-24 | 0 M1 _ | Plekha6 |
| Adam9 | 5.78E-29 | 0.315521 | 0.461 | 0.128 | 1.16E-24 | 0 M1 _ | Adam9 |
| Atf5 | 3.79E-29 | 0.927442 | 0.856 | 0.468 | 7.63E-25 | 0 M1 _ | Atf5 |
| Cp | 2.11E-29 | 0.872611 | 0.922 | 0.578 | 4.24E-25 | 0 M1 _ | Cp |
| Ccdc50 | 1.44E-29 | 0.803453 | 0.88 | 0.556 | 2.91E-25 | 0 M1 _ | Ccdc50 |
| Scn7a | 1.38E-29 | 0.639288 | 0.934 | 0.487 | 2.78E-25 | 0 M1 _ | Scn7a |
| Gstt1 | 1.15E-29 | 0.769922 | 0.778 | 0.375 | 2.32E-25 | 0 M1 _ | Gstt1 |
| Prg4 | 1.15E-29 | 0.541306 | 0.228 | 0.034 | 2.31E-25 | 0 M1 _ | Prg4 |
| Sned1 | 9.40E-30 | 1.037595 | 0.707 | 0.328 | 1.89E-25 | 0 M1 _ | Sned1 |

| | | | | | | | |
|---|---|---|---|---|---|---|---|
| Vit | 9.36E-30 | 0.464826 | 0.132 | 0.01 | 1.89E-25 | 0 M1 _ | Vit |
| Synm | 8.67E-30 | 0.44159 | 0.407 | 0.107 | 1.75E-25 | 0 M1 _ | Synm |
| Csrp2 | 6.22E-30 | 1.019866 | 0.659 | 0.263 | 1.25E-25 | 0 M1 _ | Csrp2 |
| Klf9 | 5.87E-30 | 0.874899 | 0.97 | 0.763 | 1.18E-25 | 0 M1 _ | Klf9 |
| Adm | 3.70E-30 | 0.497645 | 0.353 | 0.081 | 7.46E-26 | 0 M1 _ | Adm |
| Gpr153 | 3.19E-30 | 0.673641 | 0.772 | 0.355 | 6.42E-26 | 0 M1 _ | Gpr153 |
| Igfbp5 | 3.07E-30 | 1.460679 | 0.856 | 0.492 | 6.19E-26 | 0 M1 _ | Igfbp5 |
| Oaf | 1.84E-30 | 0.81903 | 0.88 | 0.475 | 3.71E-26 | 0 M1 _ | Oaf |
| Col6a3 | 1.26E-30 | 0.833353 | 0.982 | 0.568 | 2.54E-26 | 0 M1 _ | Col6a3 |
| Nsg1 | 1.12E-30 | 0.446901 | 0.455 | 0.122 | 2.26E-26 | 0 M1 _ | Nsg1 |
| Adamts2 | 1.03E-30 | 0.723868 | 0.689 | 0.28 | 2.08E-26 | 0 M1 _ | Adamts2 |
| Adamts12 | 7.28E-31 | 0.960935 | 0.856 | 0.46 | 1.47E-26 | 0 M1 _ | Adamts12 |
| Calml4 | 7.11E-31 | 0.450338 | 0.347 | 0.077 | 1.43E-26 | 0 M1 _ | Calml4 |
| Sdc4 | 6.76E-31 | 0.648909 | 0.784 | 0.333 | 1.36E-26 | 0 M1 _ | Sdc4 |
| Lars2 | 6.22E-31 | 1.046437 | 0.976 | 0.802 | 1.25E-26 | 0 M1 _ | Lars2 |
| Ctsb | 3.09E-31 | 0.712529 | 0.982 | 0.759 | 6.22E-27 | 0 M1 _ | Ctsb |
| Tpst1 | 2.81E-31 | 0.643616 | 0.707 | 0.294 | 5.67E-27 | 0 M1 _ | Tpst1 |
| Sox9 | 2.60E-31 | 0.381331 | 0.257 | 0.042 | 5.23E-27 | 0 M1 _ | Sox9 |
| Scd2 | 1.81E-31 | 0.842255 | 0.88 | 0.494 | 3.65E-27 | 0 M1 _ | Scd2 |
| 2900097C17Rik | 1.65E-31 | 0.712966 | 0.904 | 0.529 | 3.33E-27 | 0 M1 _ | 2900097C17Rik |
| Selenop | 1.03E-31 | 0.840676 | 0.97 | 0.75 | 2.08E-27 | 0 M1 _ | Selenop |
| Man2a1 | 3.24E-32 | 0.961973 | 0.898 | 0.537 | 6.52E-28 | 0 M1 _ | Man2a1 |
| Tpbgl | 2.14E-32 | 0.39458 | 0.353 | 0.074 | 4.31E-28 | 0 M1 _ | Tpbgl |
| Htra1 | 2.00E-32 | 1.014743 | 0.934 | 0.632 | 4.04E-28 | 0 M1 _ | Htra1 |
| Loxl2 | 1.97E-32 | 0.764064 | 0.707 | 0.302 | 3.96E-28 | 0 M1 _ | Loxl2 |
| Steap3 | 1.27E-32 | 0.916319 | 0.737 | 0.335 | 2.57E-28 | 0 M1 _ | Steap3 |
| Osr1 | 1.20E-32 | 0.658553 | 0.533 | 0.167 | 2.42E-28 | 0 M1 _ | Osr1 |
| Antxr1 | 1.09E-32 | 0.66661 | 0.701 | 0.277 | 2.19E-28 | 0 M1 _ | Antxr1 |
| Slc7a8 | 8.64E-33 | 0.380315 | 0.311 | 0.059 | 1.74E-28 | 0 M1 _ | Slc7a8 |
| Creb3l1 | 7.49E-33 | 0.749571 | 0.749 | 0.325 | 1.51E-28 | 0 M1 _ | Creb3l1 |
| Calu | 7.16E-33 | 0.805362 | 0.946 | 0.609 | 1.44E-28 | 0 M1 _ | Calu |
| Fndc3b | 5.90E-33 | 0.860676 | 0.874 | 0.508 | 1.19E-28 | 0 M1 _ | Fndc3b |
| Neo1 | 2.56E-33 | 0.609846 | 0.635 | 0.228 | 5.15E-29 | 0 M1 _ | Neo1 |
| Cebpd | 2.51E-33 | 1.181512 | 0.85 | 0.491 | 5.06E-29 | 0 M1 _ | Cebpd |
| Ar | 2.20E-33 | 0.904535 | 0.922 | 0.548 | 4.44E-29 | 0 M1 _ | Ar |
| Tnfaip6 | 2.19E-33 | 0.540259 | 0.467 | 0.128 | 4.42E-29 | 0 M1 _ | Tnfaip6 |
| Pkd2 | 1.77E-33 | 0.809372 | 0.904 | 0.493 | 3.56E-29 | 0 M1 _ | Pkd2 |
| Egfr | 1.68E-33 | 0.741999 | 0.701 | 0.284 | 3.38E-29 | 0 M1 _ | Egfr |
| Cd302 | 6.08E-34 | 0.98267 | 0.946 | 0.599 | 1.22E-29 | 0 M1 _ | Cd302 |
| Il17d | 2.70E-34 | 0.443295 | 0.419 | 0.098 | 5.44E-30 | 0 M1 _ | Il17d |
| Astn2 | 2.67E-34 | 0.403292 | 0.377 | 0.082 | 5.38E-30 | 0 M1 _ | Astn2 |

| | | | | | | | |
|---|---|---|---|---|---|---|---|
| Ndn | 2.22E-34 | 0.752395 | 0.892 | 0.438 | 4.47E-30 | 0 M1 _ | Ndn |
| S1pr3 | 1.18E-34 | 0.68103 | 0.689 | 0.261 | 2.37E-30 | 0 M1 _ | S1pr3 |
| Tril | 9.91E-35 | 0.95269 | 0.82 | 0.425 | 2.00E-30 | 0 M1 _ | Tril |
| Eda | 7.46E-35 | 0.48937 | 0.479 | 0.127 | 1.50E-30 | 0 M1 _ | Eda |
| Lpar1 | 5.23E-35 | 0.825444 | 0.814 | 0.386 | 1.05E-30 | 0 M1 _ | Lpar1 |
| Ltbp1 | 2.77E-35 | 0.955688 | 0.82 | 0.385 | 5.57E-31 | 0 M1 _ | Ltbp1 |
| Clcn5 | 1.86E-35 | 0.654222 | 0.623 | 0.218 | 3.74E-31 | 0 M1 _ | Clcn5 |
| Col6a5 | 7.63E-36 | 0.518182 | 0.281 | 0.044 | 1.54E-31 | 0 M1 _ | Col6a5 |
| Axl | 6.19E-36 | 0.905567 | 0.928 | 0.553 | 1.25E-31 | 0 M1 _ | Axl |
| Cd55 | 5.68E-36 | 0.536839 | 0.419 | 0.098 | 1.14E-31 | 0 M1 _ | Cd55 |
| Plau | 4.41E-36 | 0.787867 | 0.743 | 0.295 | 8.88E-32 | 0 M1 _ | Plau |
| Igsf3 | 4.35E-36 | 0.663078 | 0.599 | 0.206 | 8.75E-32 | 0 M1 _ | Igsf3 |
| Rhoj | 2.37E-36 | 0.941042 | 0.97 | 0.725 | 4.77E-32 | 0 M1 _ | Rhoj |
| Epb41l3 | 1.58E-36 | 0.549666 | 0.521 | 0.143 | 3.19E-32 | 0 M1 _ | Epb41l3 |
| Fbln5 | 1.25E-36 | 0.769356 | 0.814 | 0.358 | 2.52E-32 | 0 M1 _ | Fbln5 |
| Adamts15 | 9.46E-37 | 0.752956 | 0.641 | 0.217 | 1.90E-32 | 0 M1 _ | Adamts15 |
| Fam102b | 8.33E-37 | 0.802468 | 0.713 | 0.286 | 1.68E-32 | 0 M1 _ | Fam102b |
| C1ra | 4.72E-37 | 0.889062 | 0.946 | 0.522 | 9.50E-33 | 0 M1 _ | C1ra |
| Il33 | 4.12E-37 | 1.131653 | 0.826 | 0.414 | 8.30E-33 | 0 M1 _ | Il33 |
| Celf2 | 3.87E-37 | 0.950837 | 0.916 | 0.552 | 7.80E-33 | 0 M1 _ | Celf2 |
| Rhob | 3.84E-37 | 1.060755 | 0.934 | 0.593 | 7.73E-33 | 0 M1 _ | Rhob |
| Nid2 | 2.84E-37 | 0.945916 | 0.754 | 0.337 | 5.72E-33 | 0 M1 _ | Nid2 |
| Il11ra1 | 2.66E-37 | 0.996814 | 0.97 | 0.586 | 5.35E-33 | 0 M1 _ | Il11ra1 |
| Tgfbr3 | 2.39E-37 | 0.782486 | 0.862 | 0.432 | 4.81E-33 | 0 M1 _ | Tgfbr3 |
| Tsc22d3 | 8.73E-38 | 0.942228 | 0.886 | 0.448 | 1.76E-33 | 0 M1 _ | Tsc22d3 |
| Nrxn2 | 6.63E-38 | 0.98284 | 0.677 | 0.251 | 1.34E-33 | 0 M1 _ | Nrxn2 |
| Islr | 5.68E-38 | 1.005321 | 0.886 | 0.441 | 1.14E-33 | 0 M1 _ | Islr |
| Fndc5 | 4.26E-38 | 0.493571 | 0.389 | 0.08 | 8.59E-34 | 0 M1 _ | Fndc5 |
| Inmt | 3.93E-38 | 1.051001 | 0.743 | 0.271 | 7.91E-34 | 0 M1 _ | Inmt |
| Xpnpep2 | 3.52E-38 | 0.449708 | 0.311 | 0.051 | 7.09E-34 | 0 M1 _ | Xpnpep2 |
| Lpar4 | 1.57E-38 | 0.759976 | 0.527 | 0.151 | 3.15E-34 | 0 M1 _ | Lpar4 |
| Lsamp | 7.81E-39 | 1.022105 | 0.892 | 0.474 | 1.57E-34 | 0 M1 _ | Lsamp |
| Zcchc24 | 2.43E-39 | 0.908213 | 0.838 | 0.397 | 4.90E-35 | 0 M1 _ | Zcchc24 |
| Ddr2 | 1.54E-39 | 0.884815 | 0.85 | 0.413 | 3.09E-35 | 0 M1 _ | Ddr2 |
| Igsf10 | 1.44E-39 | 0.940663 | 0.91 | 0.452 | 2.89E-35 | 0 M1 _ | Igsf10 |
| Plac8 | 1.32E-39 | 0.887952 | 0.826 | 0.331 | 2.66E-35 | 0 M1 _ | Plac8 |
| Fmo1 | 1.19E-39 | 1.072256 | 0.868 | 0.463 | 2.41E-35 | 0 M1 _ | Fmo1 |
| Ppp1r3c | 5.71E-40 | 0.322824 | 0.305 | 0.045 | 1.15E-35 | 0 M1 _ | Ppp1r3c |
| Cmklr1 | 5.30E-40 | 0.754403 | 0.731 | 0.272 | 1.07E-35 | 0 M1 _ | Cmklr1 |
| Adam23 | 4.38E-40 | 0.373955 | 0.317 | 0.049 | 8.82E-36 | 0 M1 _ | Adam23 |
| Cyp1b1 | 1.17E-40 | 1.061242 | 0.754 | 0.312 | 2.36E-36 | 0 M1 _ | Cyp1b1 |

| | | | | | | | |
|---|---|---|---|---|---|---|---|
| Hspg2 | 1.11E-40 | 1.106963 | 0.976 | 0.678 | 2.24E-36 | 0 M1 _ | Hspg2 |
| Rassf2 | 7.21E-41 | 0.727416 | 0.593 | 0.183 | 1.45E-36 | 0 M1 _ | Rassf2 |
| Col5a3 | 4.67E-41 | 1.179126 | 0.946 | 0.536 | 9.41E-37 | 0 M1 _ | Col5a3 |
| Plpp3 | 3.31E-41 | 0.967307 | 0.994 | 0.711 | 6.66E-37 | 0 M1 _ | Plpp3 |
| B4galt1 | 2.60E-41 | 0.759026 | 0.808 | 0.331 | 5.24E-37 | 0 M1 _ | B4galt1 |
| AU019823 | 2.00E-41 | 0.288899 | 0.246 | 0.026 | 4.02E-37 | 0 M1 _ | AU019823 |
| Ddit4 | 1.02E-41 | 0.721458 | 0.527 | 0.133 | 2.05E-37 | 0 M1 _ | Ddit4 |
| 4632415L05Rik | 9.34E-42 | 0.255607 | 0.222 | 0.02 | 1.88E-37 | 0 M1 _ | 4632415L05Rik |
| Col5a1 | 5.26E-42 | 0.972596 | 0.94 | 0.483 | 1.06E-37 | 0 M1 _ | Col5a1 |
| Dopey2 | 4.02E-42 | 0.291747 | 0.18 | 0.012 | 8.10E-38 | 0 M1 _ | Dopey2 |
| Lrig1 | 3.48E-42 | 0.614114 | 0.605 | 0.179 | 7.01E-38 | 0 M1 _ | Lrig1 |
| Dnm1 | 2.58E-42 | 0.639364 | 0.629 | 0.19 | 5.19E-38 | 0 M1 _ | Dnm1 |
| Lama2 | 2.35E-42 | 1.08589 | 0.982 | 0.599 | 4.73E-38 | 0 M1 _ | Lama2 |
| Lamc1 | 1.79E-42 | 1.143563 | 0.928 | 0.59 | 3.60E-38 | 0 M1 _ | Lamc1 |
| Htra3 | 1.33E-42 | 1.216033 | 0.964 | 0.538 | 2.67E-38 | 0 M1 _ | Htra3 |
| Fetub | 1.31E-42 | 0.315828 | 0.192 | 0.014 | 2.63E-38 | 0 M1 _ | Fetub |
| Fam208b | 9.10E-43 | 0.279117 | 0.234 | 0.023 | 1.83E-38 | 0 M1 _ | Fam208b |
| Zbtb16 | 8.33E-43 | 0.461637 | 0.413 | 0.076 | 1.68E-38 | 0 M1 _ | Zbtb16 |
| Scd1 | 5.73E-43 | 1.195369 | 0.731 | 0.287 | 1.15E-38 | 0 M1 _ | Scd1 |
| Pdgfra | 5.72E-43 | 1.100675 | 0.982 | 0.652 | 1.15E-38 | 0 M1 _ | Pdgfra |
| Taf6l | 4.44E-43 | 0.405308 | 0.389 | 0.069 | 8.95E-39 | 0 M1 _ | Taf6l |
| Fgfr1 | 3.08E-43 | 0.926459 | 0.946 | 0.456 | 6.19E-39 | 0 M1 _ | Fgfr1 |
| Sphkap | 2.18E-43 | 0.409935 | 0.293 | 0.038 | 4.39E-39 | 0 M1 _ | Sphkap |
| P2ry1 | 1.83E-43 | 0.945033 | 0.695 | 0.235 | 3.69E-39 | 0 M1 _ | P2ry1 |
| Cyr61 | 1.39E-43 | 0.627245 | 0.234 | 0.022 | 2.80E-39 | 0 M1 _ | Cyr61 |
| Igfbp4 | 1.02E-43 | 1.153269 | 0.994 | 0.715 | 2.05E-39 | 0 M1 _ | Igfbp4 |
| Igf1 | 9.10E-44 | 1.454797 | 0.91 | 0.479 | 1.83E-39 | 0 M1 _ | Igf1 |
| Lamb1 | 5.15E-44 | 1.120808 | 0.958 | 0.635 | 1.04E-39 | 0 M1 _ | Lamb1 |
| Mmp14 | 4.60E-44 | 1.117548 | 0.946 | 0.51 | 9.26E-40 | 0 M1 _ | Mmp14 |
| Fam49a | 2.25E-44 | 0.587022 | 0.611 | 0.165 | 4.54E-40 | 0 M1 _ | Fam49a |
| Aldh2 | 2.06E-44 | 0.964148 | 0.982 | 0.741 | 4.14E-40 | 0 M1 _ | Aldh2 |
| Atp1a2 | 5.42E-45 | 0.811577 | 0.671 | 0.213 | 1.09E-40 | 0 M1 _ | Atp1a2 |
| Zfp395 | 3.31E-45 | 0.459225 | 0.377 | 0.064 | 6.67E-41 | 0 M1 _ | Zfp395 |
| Chtf8 | 2.96E-45 | 0.412298 | 0.401 | 0.069 | 5.97E-41 | 0 M1 _ | Chtf8 |
| Plxdc2 | 2.68E-45 | 1.158985 | 0.982 | 0.557 | 5.39E-41 | 0 M1 _ | Plxdc2 |
| Hsp90b1 | 1.97E-45 | 0.90123 | 0.994 | 0.886 | 3.96E-41 | 0 M1 _ | Hsp90b1 |
| Col14a1 | 1.93E-45 | 1.240033 | 0.88 | 0.428 | 3.88E-41 | 0 M1 _ | Col14a1 |
| Medag | 1.75E-45 | 1.080347 | 0.778 | 0.299 | 3.53E-41 | 0 M1 _ | Medag |
| Kcna2 | 4.59E-46 | 0.470752 | 0.287 | 0.034 | 9.25E-42 | 0 M1 _ | Kcna2 |
| 1500011K16Rik | 3.60E-46 | 0.349784 | 0.287 | 0.032 | 7.25E-42 | 0 M1 _ | 1500011K16Rik |
| Emilin2 | 2.14E-46 | 0.713539 | 0.563 | 0.145 | 4.32E-42 | 0 M1 _ | Emilin2 |

| | | | | | | | |
|---|---|---|---|---|---|---|---|
| Ifi207 | 2.05E-46 | 1.136341 | 0.778 | 0.302 | 4.13E-42 | 0 M1 _ | Ifi207 |
| Itih5 | 1.58E-46 | 1.3831 | 0.982 | 0.655 | 3.19E-42 | 0 M1 _ | Itih5 |
| Fam96b | 9.22E-47 | 0.364629 | 0.359 | 0.05 | 1.86E-42 | 0 M1 _ | Fam96b |
| 0610009O20Rik | 6.26E-47 | 0.347035 | 0.246 | 0.023 | 1.26E-42 | 0 M1 _ | 0610009O20Rik |
| Lrp1 | 5.86E-47 | 1.120168 | 0.976 | 0.678 | 1.18E-42 | 0 M1 _ | Lrp1 |
| Ret | 5.34E-47 | 0.282573 | 0.21 | 0.015 | 1.07E-42 | 0 M1 _ | Ret |
| Ptges | 5.04E-47 | 0.783178 | 0.719 | 0.23 | 1.01E-42 | 0 M1 _ | Ptges |
| Pyurf | 3.67E-47 | 0.294464 | 0.281 | 0.03 | 7.39E-43 | 0 M1 _ | Pyurf |
| Camk2n1 | 3.49E-47 | 1.177396 | 0.922 | 0.516 | 7.02E-43 | 0 M1 _ | Camk2n1 |
| Nid1 | 1.75E-47 | 1.145733 | 0.982 | 0.716 | 3.53E-43 | 0 M1 _ | Nid1 |
| Masp1 | 9.50E-48 | 0.642787 | 0.269 | 0.029 | 1.91E-43 | 0 M1 _ | Masp1 |
| Scara3 | 5.88E-48 | 0.866266 | 0.671 | 0.204 | 1.18E-43 | 0 M1 _ | Scara3 |
| Nfix | 3.93E-48 | 1.092298 | 0.976 | 0.668 | 7.92E-44 | 0 M1 _ | Nfix |
| Crispld2 | 3.13E-48 | 1.390828 | 0.946 | 0.56 | 6.31E-44 | 0 M1 _ | Crispld2 |
| C1s1 | 2.31E-48 | 1.137403 | 0.982 | 0.702 | 4.66E-44 | 0 M1 _ | C1s1 |
| Sh3pxd2b | 1.62E-48 | 0.84527 | 0.784 | 0.293 | 3.27E-44 | 0 M1 _ | Sh3pxd2b |
| Frmd6 | 9.91E-49 | 0.998146 | 0.886 | 0.354 | 2.00E-44 | 0 M1 _ | Frmd6 |
| Tspan11 | 7.50E-49 | 0.986915 | 0.868 | 0.353 | 1.51E-44 | 0 M1 _ | Tspan11 |
| Mrc2 | 6.98E-49 | 0.949865 | 0.892 | 0.39 | 1.41E-44 | 0 M1 _ | Mrc2 |
| Ms4a4d | 4.36E-49 | 1.190745 | 0.892 | 0.415 | 8.78E-45 | 0 M1 _ | Ms4a4d |
| Cd34 | 3.17E-49 | 1.42013 | 0.892 | 0.415 | 6.39E-45 | 0 M1 _ | Cd34 |
| Dapk1 | 1.84E-49 | 1.008476 | 0.826 | 0.33 | 3.71E-45 | 0 M1 _ | Dapk1 |
| Fam212a | 1.81E-49 | 0.285018 | 0.24 | 0.019 | 3.64E-45 | 0 M1 _ | Fam212a |
| Lrrtm3 | 7.65E-50 | 0.323548 | 0.198 | 0.012 | 1.54E-45 | 0 M1 _ | Lrrtm3 |
| C3 | 4.30E-50 | 2.083386 | 0.85 | 0.365 | 8.65E-46 | 0 M1 _ | C3 |
| Thsd7a | 3.88E-50 | 0.623905 | 0.419 | 0.071 | 7.82E-46 | 0 M1 _ | Thsd7a |
| Sssca1 | 3.41E-50 | 0.289439 | 0.24 | 0.019 | 6.86E-46 | 0 M1 _ | Sssca1 |
| Zbtb20 | 2.51E-50 | 1.181689 | 0.988 | 0.893 | 5.05E-46 | 0 M1 _ | Zbtb20 |
| Rspo1 | 1.25E-50 | 0.71843 | 0.413 | 0.071 | 2.52E-46 | 0 M1 _ | Rspo1 |
| Nfia | 9.71E-51 | 1.197289 | 0.928 | 0.534 | 1.96E-46 | 0 M1 _ | Nfia |
| Gxylt2 | 6.59E-51 | 0.80029 | 0.581 | 0.146 | 1.33E-46 | 0 M1 _ | Gxylt2 |
| Lum | 1.49E-51 | 1.332408 | 0.982 | 0.58 | 3.00E-47 | 0 M1 _ | Lum |
| Anpep | 1.27E-51 | 0.96266 | 0.725 | 0.238 | 2.56E-47 | 0 M1 _ | Anpep |
| Ppl | 8.82E-53 | 0.406916 | 0.269 | 0.024 | 1.78E-48 | 0 M1 _ | Ppl |
| Ces2g | 8.77E-53 | 0.836855 | 0.521 | 0.112 | 1.77E-48 | 0 M1 _ | Ces2g |
| Adam19 | 5.02E-53 | 1.142004 | 0.725 | 0.242 | 1.01E-48 | 0 M1 _ | Adam19 |
| Maf | 4.76E-53 | 1.44239 | 0.922 | 0.505 | 9.58E-49 | 0 M1 _ | Maf |
| Pcsk6 | 2.33E-53 | 0.502099 | 0.347 | 0.043 | 4.68E-49 | 0 M1 _ | Pcsk6 |
| Gpm6b | 1.35E-53 | 1.292049 | 0.952 | 0.491 | 2.71E-49 | 0 M1 _ | Gpm6b |
| Gcn1l1 | 1.34E-53 | 0.339542 | 0.293 | 0.028 | 2.69E-49 | 0 M1 _ | Gcn1l1 |
| Zufsp | 5.34E-54 | 0.347412 | 0.257 | 0.02 | 1.08E-49 | 0 M1 _ | Zufsp |

| | | | | | | | |
|---|---|---|---|---|---|---|---|
| Abca8b | 3.39E-54 | 1.0542 | 0.629 | 0.165 | 6.83E-50 | 0 M1 _ | Abca8b |
| Lrrc17 | 7.35E-55 | 0.647891 | 0.365 | 0.047 | 1.48E-50 | 0 M1 _ | Lrrc17 |
| Col15a1 | 4.99E-55 | 1.493697 | 0.97 | 0.609 | 1.00E-50 | 0 M1 _ | Col15a1 |
| mt-Atp8 | 1.19E-55 | 1.301682 | 0.946 | 0.581 | 2.40E-51 | 0 M1 _ | mt-Atp8 |
| Fam58b | 1.17E-55 | 0.326755 | 0.275 | 0.023 | 2.36E-51 | 0 M1 _ | Fam58b |
| Crabp1 | 6.28E-56 | 1.829339 | 0.623 | 0.164 | 1.26E-51 | 0 M1 _ | Crabp1 |
| Mgst1 | 1.89E-56 | 1.226461 | 0.934 | 0.469 | 3.81E-52 | 0 M1 _ | Mgst1 |
| Rbms3 | 5.13E-57 | 1.394069 | 0.964 | 0.543 | 1.03E-52 | 0 M1 _ | Rbms3 |
| Timp2 | 3.99E-57 | 1.240202 | 0.988 | 0.685 | 8.04E-53 | 0 M1 _ | Timp2 |
| Clmp | 1.40E-57 | 0.98522 | 0.79 | 0.267 | 2.81E-53 | 0 M1 _ | Clmp |
| Clec3b | 1.34E-57 | 1.56311 | 0.916 | 0.401 | 2.70E-53 | 0 M1 _ | Clec3b |
| Efemp1 | 6.53E-58 | 1.527102 | 0.838 | 0.32 | 1.31E-53 | 0 M1 _ | Efemp1 |
| Abca8a | 1.25E-58 | 1.559474 | 0.958 | 0.498 | 2.52E-54 | 0 M1 _ | Abca8a |
| Papd7 | 2.07E-59 | 0.292941 | 0.222 | 0.012 | 4.18E-55 | 0 M1 _ | Papd7 |
| D3Ertd254e | 1.86E-59 | 0.258995 | 0.204 | 0.009 | 3.74E-55 | 0 M1 _ | D3Ertd254e |
| Gsta3 | 1.37E-59 | 0.583267 | 0.347 | 0.039 | 2.76E-55 | 0 M1 _ | Gsta3 |
| D10Jhu81e | 1.80E-60 | 0.353479 | 0.311 | 0.027 | 3.63E-56 | 0 M1 _ | D10Jhu81e |
| Lppos | 1.63E-60 | 0.265254 | 0.216 | 0.01 | 3.28E-56 | 0 M1 _ | Lppos |
| Rspo3 | 9.26E-61 | 1.283672 | 0.766 | 0.237 | 1.87E-56 | 0 M1 _ | Rspo3 |
| Aox1 | 7.82E-61 | 0.6219 | 0.497 | 0.084 | 1.58E-56 | 0 M1 _ | Aox1 |
| Tnxb | 3.89E-61 | 1.317981 | 0.725 | 0.204 | 7.84E-57 | 0 M1 _ | Tnxb |
| Ikbkap | 4.49E-62 | 0.267338 | 0.204 | 0.008 | 9.03E-58 | 0 M1 _ | Ikbkap |
| A230050P20Rik | 1.07E-62 | 0.314755 | 0.246 | 0.014 | 2.16E-58 | 0 M1 _ | A230050P20Rik |
| Mest | 5.83E-64 | 0.570149 | 0.299 | 0.024 | 1.17E-59 | 0 M1 _ | Mest |
| Bmp4 | 2.25E-64 | 1.314265 | 0.731 | 0.207 | 4.53E-60 | 0 M1 _ | Bmp4 |
| Tmem57 | 5.11E-65 | 0.347911 | 0.305 | 0.024 | 1.03E-60 | 0 M1 _ | Tmem57 |
| Scara5 | 4.82E-65 | 1.497965 | 0.916 | 0.371 | 9.71E-61 | 0 M1 _ | Scara5 |
| Adh1 | 3.30E-65 | 1.037855 | 0.641 | 0.135 | 6.66E-61 | 0 M1 _ | Adh1 |
| Mmp2 | 2.72E-65 | 1.715286 | 0.982 | 0.531 | 5.48E-61 | 0 M1 _ | Mmp2 |
| 9030624J02Rik | 1.16E-65 | 0.343812 | 0.317 | 0.025 | 2.33E-61 | 0 M1 _ | 9030624J02Rik |
| Gas1 | 1.10E-65 | 1.789474 | 0.976 | 0.592 | 2.21E-61 | 0 M1 _ | Gas1 |
| 2410089E03Rik | 9.75E-66 | 0.301423 | 0.287 | 0.019 | 1.96E-61 | 0 M1 _ | 2410089E03Rik |
| Mum1 | 2.51E-66 | 0.460373 | 0.365 | 0.036 | 5.06E-62 | 0 M1 _ | Mum1 |
| Coro1b | 1.90E-66 | 0.495047 | 0.503 | 0.074 | 3.83E-62 | 0 M1 _ | Coro1b |
| Fam96a | 8.86E-67 | 0.364455 | 0.317 | 0.025 | 1.78E-62 | 0 M1 _ | Fam96a |
| Rbm12 | 1.40E-67 | 0.2836 | 0.24 | 0.011 | 2.81E-63 | 0 M1 _ | Rbm12 |
| Fam26e | 9.59E-68 | 0.318151 | 0.234 | 0.01 | 1.93E-63 | 0 M1 _ | Fam26e |
| Ptgfr | 7.07E-68 | 0.671536 | 0.359 | 0.035 | 1.42E-63 | 0 M1 _ | Ptgfr |
| Spock2 | 2.26E-68 | 0.958393 | 0.665 | 0.144 | 4.54E-64 | 0 M1 _ | Spock2 |
| Capn6 | 2.07E-68 | 0.876311 | 0.593 | 0.116 | 4.17E-64 | 0 M1 _ | Capn6 |
| 2010111I01Rik | 5.01E-69 | 0.448359 | 0.449 | 0.052 | 1.01E-64 | 0 M1 _ | 2010111I01Rik |

| | | | | | | | |
|---|---|---|---|---|---|---|---|
| Ccl11 | 6.15E-70 | 1.677639 | 0.946 | 0.402 | 1.24E-65 | 0 M1 _ | Ccl11 |
| Antxr2 | 1.46E-70 | 1.49731 | 0.892 | 0.358 | 2.94E-66 | 0 M1 _ | Antxr2 |
| Dcn | 3.25E-71 | 1.614986 | 0.994 | 0.652 | 6.55E-67 | 0 M1 _ | Dcn |
| D17Wsu92e | 3.08E-71 | 0.41894 | 0.389 | 0.037 | 6.19E-67 | 0 M1 _ | D17Wsu92e |
| Pla2g16 | 2.84E-71 | 0.419435 | 0.323 | 0.024 | 5.71E-67 | 0 M1 _ | Pla2g16 |
| 2310035C23Rik | 2.53E-71 | 0.338901 | 0.299 | 0.019 | 5.09E-67 | 0 M1 _ | 2310035C23Rik |
| Fam69b | 2.23E-71 | 0.30907 | 0.222 | 0.007 | 4.49E-67 | 0 M1 _ | Fam69b |
| Angptl1 | 8.22E-72 | 1.055621 | 0.281 | 0.017 | 1.66E-67 | 0 M1 _ | Angptl1 |
| Srpx | 4.25E-72 | 1.810232 | 0.94 | 0.438 | 8.55E-68 | 0 M1 _ | Srpx |
| BC017643 | 3.09E-72 | 0.367075 | 0.263 | 0.013 | 6.23E-68 | 0 M1 _ | BC017643 |
| Rnase4 | 2.04E-72 | 1.632341 | 0.97 | 0.574 | 4.11E-68 | 0 M1 _ | Rnase4 |
| Spon2 | 5.97E-73 | 1.673353 | 0.743 | 0.179 | 1.20E-68 | 0 M1 _ | Spon2 |
| Col18a1 | 2.00E-73 | 1.285684 | 0.754 | 0.187 | 4.03E-69 | 0 M1 _ | Col18a1 |
| Tmem55b | 3.01E-74 | 0.423016 | 0.353 | 0.028 | 6.06E-70 | 0 M1 _ | Tmem55b |
| Fst | 9.71E-75 | 2.120007 | 0.826 | 0.259 | 1.96E-70 | 0 M1 _ | Fst |
| Papd4 | 3.37E-75 | 0.362856 | 0.323 | 0.022 | 6.79E-71 | 0 M1 _ | Papd4 |
| Ifi205 | 7.01E-77 | 1.821281 | 0.838 | 0.257 | 1.41E-72 | 0 M1 _ | Ifi205 |
| Gsn | 3.59E-77 | 2.243742 | 0.994 | 0.788 | 7.24E-73 | 0 M1 _ | Gsn |
| Dpep1 | 1.39E-77 | 2.009672 | 0.976 | 0.47 | 2.79E-73 | 0 M1 _ | Dpep1 |
| Fam208a | 3.73E-79 | 0.682268 | 0.491 | 0.056 | 7.51E-75 | 0 M1 _ | Fam208a |
| Mmp19 | 1.30E-79 | 1.381187 | 0.868 | 0.266 | 2.62E-75 | 0 M1 _ | Mmp19 |
| Ntrk2 | 5.76E-80 | 1.582131 | 0.814 | 0.226 | 1.16E-75 | 0 M1 _ | Ntrk2 |
| Fam103a1 | 1.71E-81 | 0.581345 | 0.431 | 0.04 | 3.44E-77 | 0 M1 _ | Fam103a1 |
| 1190002N15Rik | 9.65E-82 | 0.345144 | 0.269 | 0.011 | 1.94E-77 | 0 M1 _ | 1190002N15Rik |
| Dirc2 | 3.80E-82 | 0.447247 | 0.395 | 0.031 | 7.65E-78 | 0 M1 _ | Dirc2 |
| Gatsl2 | 2.74E-82 | 0.378024 | 0.293 | 0.014 | 5.51E-78 | 0 M1 _ | Gatsl2 |
| 0610037L13Rik | 1.70E-82 | 0.481 | 0.383 | 0.029 | 3.43E-78 | 0 M1 _ | 0610037L13Rik |
| Fbxo18 | 1.06E-82 | 0.415286 | 0.371 | 0.027 | 2.14E-78 | 0 M1 _ | Fbxo18 |
| Sdccag3 | 5.75E-83 | 0.452865 | 0.359 | 0.024 | 1.16E-78 | 0 M1 _ | Sdccag3 |
| Fam220a | 7.39E-84 | 0.461644 | 0.407 | 0.033 | 1.49E-79 | 0 M1 _ | Fam220a |
| Wisp1 | 9.98E-85 | 0.341919 | 0.192 | 0.001 | 2.01E-80 | 0 M1 _ | Wisp1 |
| 1110008F13Rik | 6.14E-85 | 0.594663 | 0.545 | 0.062 | 1.24E-80 | 0 M1 _ | 1110008F13Rik |
| Kdelc1 | 7.16E-86 | 0.35777 | 0.287 | 0.012 | 1.44E-81 | 0 M1 _ | Kdelc1 |
| Enpp2 | 2.76E-88 | 2.601172 | 0.958 | 0.43 | 5.57E-84 | 0 M1 _ | Enpp2 |
| Pappa | 1.73E-89 | 1.72344 | 0.778 | 0.177 | 3.48E-85 | 0 M1 _ | Pappa |
| Skiv2l2 | 1.15E-89 | 0.546309 | 0.425 | 0.033 | 2.31E-85 | 0 M1 _ | Skiv2l2 |
| Ptp4a1 | 7.70E-91 | 0.661302 | 0.467 | 0.043 | 1.55E-86 | 0 M1 _ | Ptp4a1 |
| Myoc | 1.10E-91 | 1.028965 | 0.335 | 0.017 | 2.21E-87 | 0 M1 _ | Myoc |
| Aox3 | 1.88E-92 | 1.479643 | 0.599 | 0.084 | 3.78E-88 | 0 M1 _ | Aox3 |
| Fgf10 | 1.00E-93 | 2.022837 | 0.85 | 0.229 | 2.02E-89 | 0 M1 _ | Fgf10 |
| Fam19a5 | 2.93E-94 | 0.37864 | 0.228 | 0.003 | 5.91E-90 | 0 M1 _ | Fam19a5 |

| | | | | | | | |
|---|---|---|---|---|---|---|---|
| CT025619.1 | 7.05E-98 | 0.446878 | 0.311 | 0.011 | 1.42E-93 | 0 M1 _ | CT025619.1 |
| Zcchc11 | 8.57E-99 | 0.681321 | 0.491 | 0.042 | 1.73E-94 | 0 M1 _ | Zcchc11 |
| Dopey1 | 2.70E-100 | 0.525355 | 0.419 | 0.027 | 5.44E-96 | 0 M1 _ | Dopey1 |
| Tmem110 | 1.39E-105 | 0.448563 | 0.353 | 0.014 | 2.80E-101 | 0 M1 _ | Tmem110 |
| Mut | 1.83E-107 | 0.516489 | 0.413 | 0.023 | 3.70E-103 | 0 M1 _ | Mut |
| Zcchc6 | 1.63E-109 | 0.85791 | 0.611 | 0.061 | 3.28E-105 | 0 M1 _ | Zcchc6 |
| 1700020I14Rik | 8.79E-113 | 0.797352 | 0.653 | 0.067 | 1.77E-108 | 0 M1 _ | 1700020I14Rik |
| 2810474O19Rik | 6.23E-115 | 0.787035 | 0.551 | 0.044 | 1.25E-110 | 0 M1 _ | 2810474O19Rik |
| Mgea5 | 2.84E-115 | 0.587365 | 0.461 | 0.027 | 5.72E-111 | 0 M1 _ | Mgea5 |
| BC003331 | 3.20E-119 | 0.705796 | 0.509 | 0.035 | 6.44E-115 | 0 M1 _ | BC003331 |
| Tmem5 | 1.36E-125 | 0.762448 | 0.605 | 0.049 | 2.74E-121 | 0 M1 _ | Tmem5 |
| 1810022K09Rik | 1.03E-125 | 0.872009 | 0.659 | 0.062 | 2.08E-121 | 0 M1 _ | 1810022K09Rik |
| Trove2 | 3.26E-129 | 0.844335 | 0.599 | 0.046 | 6.56E-125 | 0 M1 _ | Trove2 |
| Ctgf | 3.07E-134 | 1.800601 | 0.449 | 0.019 | 6.19E-130 | 0 M1 _ | Ctgf |
| Fam69a | 1.79E-134 | 0.701563 | 0.527 | 0.031 | 3.61E-130 | 0 M1 _ | Fam69a |
| 2410015M20Rik | 6.22E-142 | 0.977578 | 0.743 | 0.071 | 1.25E-137 | 0 M1 _ | 2410015M20Rik |
| Usmg5 | 2.43E-142 | 1.066346 | 0.808 | 0.082 | 4.89E-138 | 0 M1 _ | Usmg5 |
| Aaed1 | 1.63E-145 | 0.80122 | 0.581 | 0.036 | 3.29E-141 | 0 M1 _ | Aaed1 |
| Dpt | 1.26E-145 | 2.342407 | 0.85 | 0.118 | 2.53E-141 | 0 M1 _ | Dpt |
| AI314180 | 5.96E-146 | 0.794313 | 0.581 | 0.035 | 1.20E-141 | 0 M1 _ | AI314180 |
| 2010107E04Rik | 3.77E-162 | 1.184718 | 0.832 | 0.077 | 7.59E-158 | 0 M1 _ | 2010107E04Rik |
| Atp5o.1 | 9.90E-166 | 1.123091 | 0.82 | 0.072 | 1.99E-161 | 0 M1 _ | Atp5o.1 |
| Minos1 | 3.62E-167 | 1.269847 | 0.832 | 0.075 | 7.29E-163 | 0 M1 _ | Minos1 |
| Kdelc2 | 1.04E-181 | 0.95725 | 0.641 | 0.031 | 2.09E-177 | 0 M1 _ | Kdelc2 |
| Aes | 1.99E-185 | 1.491024 | 0.904 | 0.079 | 4.02E-181 | 0 M1 _ | Aes |
| Fam46a | 4.20E-230 | 1.386155 | 0.617 | 0.014 | 8.45E-226 | 0 M1 _ | Fam46a |
| Fam198b | 0 | 2.324246 | 0.94 | 0.034 | 0 | 0 M1 _ | Fam198b |
| Anp32e | 2.48E-06 | 0.262143 | 0.43 | 0.313 | 0.049851 | 1 L1 _ | Anp32e |
| Uqcrc1 | 2.33E-06 | 0.258574 | 0.602 | 0.483 | 0.046835 | 1 L1 _ | Uqcrc1 |
| Idnk | 2.21E-06 | 0.269327 | 0.508 | 0.387 | 0.044421 | 1 L1 _ | Idnk |
| Cycs1 | 2.02E-06 | 0.251392 | 0.621 | 0.518 | 0.040619 | 1 L1 _ | Cycs |
| Jak1 | 1.82E-06 | 0.287048 | 0.793 | 0.765 | 0.036557 | 1 L1 _ | Jak1 |
| Kif13b | 1.72E-06 | 0.275396 | 0.501 | 0.391 | 0.034666 | 1 L1 _ | Kif13b |
| Rbm25 | 1.67E-06 | 0.269886 | 0.726 | 0.679 | 0.033584 | 1 L1 _ | Rbm25 |
| Ccdc12 | 1.60E-06 | 0.321056 | 0.694 | 0.617 | 0.032243 | 1 L1 _ | Ccdc12 |
| Pgs1 | 1.57E-06 | 0.259785 | 0.248 | 0.152 | 0.031627 | 1 L1 _ | Pgs1 |
| Lst1 | 1.52E-06 | 0.779046 | 0.248 | 0.162 | 0.030713 | 1 L1 _ | Lst1 |
| Eif4a1 | 7.99E-07 | 0.31805 | 0.786 | 0.764 | 0.016084 | 1 L1 _ | Eif4a1 |
| Wipf11 | 6.68E-07 | 0.27241 | 0.552 | 0.431 | 0.013456 | 1 L1 _ | Wipf1 |
| Hist1h4d | 6.44E-07 | 0.287601 | 0.529 | 0.407 | 0.012975 | 1 L1 _ | Hist1h4d |
| Snrpd2 | 5.80E-07 | 0.265798 | 0.759 | 0.679 | 0.011679 | 1 L1 _ | Snrpd2 |

| | | | | | | | |
|---|---|---|---|---|---|---|---|
| Srpk1 | 5.76E-07 | 0.261274 | 0.476 | 0.358 | 0.011604 | 1 L1 _ | Srpk1 |
| Nfkb1 | 3.43E-07 | 0.253469 | 0.517 | 0.402 | 0.006898 | 1 L1 _ | Nfkb1 |
| Ifnar1 | 3.28E-07 | 0.269214 | 0.423 | 0.297 | 0.006608 | 1 L1 _ | Ifnar1 |
| Rgs10 | 2.64E-07 | 0.495063 | 0.322 | 0.224 | 0.005313 | 1 L1 _ | Rgs10 |
| Ggnbp2 | 2.50E-07 | 0.287439 | 0.589 | 0.459 | 0.005033 | 1 L1 _ | Ggnbp2 |
| Phb2 | 2.27E-07 | 0.270872 | 0.644 | 0.53 | 0.004567 | 1 L1 _ | Phb2 |
| Pitpna | 2.15E-07 | 0.323533 | 0.609 | 0.485 | 0.004328 | 1 L1 _ | Pitpna |
| Os9 | 9.31E-08 | 0.308398 | 0.648 | 0.523 | 0.001875 | 1 L1 _ | Os9 |
| Uqcc2 | 9.17E-08 | 0.318413 | 0.683 | 0.572 | 0.001847 | 1 L1 _ | Uqcc2 |
| Gas71 | 8.46E-08 | 0.320139 | 0.462 | 0.332 | 0.001703 | 1 L1 _ | Gas7 |
| Man2b1 | 7.40E-08 | 0.457365 | 0.444 | 0.328 | 0.00149 | 1 L1 _ | Man2b1 |
| Dennd1b | 7.04E-08 | 0.252121 | 0.343 | 0.217 | 0.001419 | 1 L1 _ | Dennd1b |
| Mrpl58 | 6.72E-08 | 0.257375 | 0.517 | 0.378 | 0.001353 | 1 L1 _ | Mrpl58 |
| Hpcal11 | 6.55E-08 | 0.286881 | 0.478 | 0.341 | 0.00132 | 1 L1 _ | Hpcal1 |
| Fam173a | 6.49E-08 | 0.298468 | 0.556 | 0.43 | 0.001307 | 1 L1 _ | Fam173a |
| Atp5o | 6.03E-08 | 0.254228 | 0.761 | 0.587 | 0.001214 | 1 L1 _ | Atp5o |
| Grb2 | 5.07E-08 | 0.303337 | 0.51 | 0.385 | 0.001022 | 1 L1 _ | Grb2 |
| Cnot6l | 4.81E-08 | 0.274904 | 0.492 | 0.363 | 0.000969 | 1 L1 _ | Cnot6l |
| Sema6d1 | 4.80E-08 | 0.258154 | 0.48 | 0.331 | 0.000966 | 1 L1 _ | Sema6d |
| Ldlrad4 | 4.15E-08 | 0.273771 | 0.278 | 0.163 | 0.000836 | 1 L1 _ | Ldlrad4 |
| Srsf51 | 4.03E-08 | 0.296609 | 0.807 | 0.752 | 0.000811 | 1 L1 _ | Srsf5 |
| Crot | 4.00E-08 | 0.258593 | 0.407 | 0.278 | 0.000806 | 1 L1 _ | Crot |
| Tle2 | 3.99E-08 | 0.250019 | 0.179 | 0.089 | 0.000803 | 1 L1 _ | Tle2 |
| Actr2 | 3.95E-08 | 0.276889 | 0.641 | 0.505 | 0.000795 | 1 L1 _ | Actr2 |
| Akr1b3 | 3.54E-08 | 0.285558 | 0.552 | 0.422 | 0.000712 | 1 L1 _ | Akr1b3 |
| Klf61 | 3.04E-08 | 0.320938 | 0.662 | 0.536 | 0.000612 | 1 L1 _ | Klf6 |
| Trp53 | 3.00E-08 | 0.291972 | 0.529 | 0.398 | 0.000604 | 1 L1 _ | Trp53 |
| Pole4 | 2.95E-08 | 0.254609 | 0.421 | 0.29 | 0.000595 | 1 L1 _ | Pole4 |
| Borcs8 | 2.84E-08 | 0.291234 | 0.322 | 0.201 | 0.000571 | 1 L1 _ | Borcs8 |
| Izumo4 | 2.63E-08 | 0.274054 | 0.361 | 0.234 | 0.000529 | 1 L1 _ | Izumo4 |
| Mark2 | 1.88E-08 | 0.315262 | 0.446 | 0.304 | 0.00038 | 1 L1 _ | Mark2 |
| Swap701 | 1.37E-08 | 0.340935 | 0.492 | 0.359 | 0.000276 | 1 L1 _ | Swap70 |
| Spcs2 | 1.04E-08 | 0.276228 | 0.818 | 0.736 | 0.000209 | 1 L1 _ | Spcs2 |
| Mrpl54 | 6.94E-09 | 0.270727 | 0.533 | 0.404 | 0.00014 | 1 L1 _ | Mrpl54 |
| Odc1 | 6.17E-09 | 0.302378 | 0.379 | 0.245 | 0.000124 | 1 L1 _ | Odc1 |
| Ndufb7 | 6.16E-09 | 0.269224 | 0.768 | 0.692 | 0.000124 | 1 L1 _ | Ndufb7 |
| Sin3b | 5.28E-09 | 0.287521 | 0.579 | 0.441 | 0.000106 | 1 L1 _ | Sin3b |
| Cdc37 | 5.09E-09 | 0.271782 | 0.754 | 0.67 | 0.000103 | 1 L1 _ | Cdc37 |
| Gm10076 | 4.79E-09 | 0.253129 | 0.391 | 0.249 | 9.64E-05 | 1 L1 _ | Gm10076 |
| Pabpn1 | 4.66E-09 | 0.279395 | 0.687 | 0.538 | 9.39E-05 | 1 L1 _ | Pabpn1 |
| 1110008P14Rik | 4.31E-09 | 0.310512 | 0.405 | 0.27 | 8.69E-05 | 1 L1 _ | 1110008P14Rik |

| | | | | | | | |
|---|---|---|---|---|---|---|---|
| Tln11 | 3.89E-09 | 0.274693 | 0.837 | 0.81 | 7.84E-05 | 1 L1 _ | Tln1 |
| Ran | 3.24E-09 | 0.363852 | 0.775 | 0.682 | 6.53E-05 | 1 L1 _ | Ran |
| H2-Eb1 | 3.20E-09 | 1.747341 | 0.313 | 0.2 | 6.44E-05 | 1 L1 _ | H2-Eb1 |
| Arl6ip5 | 2.97E-09 | 0.335864 | 0.625 | 0.481 | 5.98E-05 | 1 L1 _ | Arl6ip5 |
| Med10 | 2.69E-09 | 0.262206 | 0.501 | 0.358 | 5.42E-05 | 1 L1 _ | Med10 |
| Cnot3 | 2.45E-09 | 0.29317 | 0.591 | 0.446 | 4.94E-05 | 1 L1 _ | Cnot3 |
| Hspe1 | 2.17E-09 | 0.376937 | 0.791 | 0.713 | 4.36E-05 | 1 L1 _ | Hspe1 |
| Cdk2ap2 | 2.08E-09 | 0.326948 | 0.621 | 0.477 | 4.18E-05 | 1 L1 _ | Cdk2ap2 |
| Cbl | 2.02E-09 | 0.272052 | 0.462 | 0.308 | 4.07E-05 | 1 L1 _ | Cbl |
| Tkt | 1.46E-09 | 0.336148 | 0.57 | 0.441 | 2.94E-05 | 1 L1 _ | Tkt |
| Use1 | 1.44E-09 | 0.364781 | 0.611 | 0.491 | 2.90E-05 | 1 L1 _ | Use1 |
| Xrn2 | 1.28E-09 | 0.318579 | 0.602 | 0.462 | 2.58E-05 | 1 L1 _ | Xrn2 |
| Mitd1 | 1.13E-09 | 0.312253 | 0.377 | 0.24 | 2.27E-05 | 1 L1 _ | Mitd1 |
| Ndfip1 | 1.10E-09 | 0.356698 | 0.825 | 0.703 | 2.21E-05 | 1 L1 _ | Ndfip1 |
| Znrf1 | 8.76E-10 | 0.266752 | 0.384 | 0.235 | 1.76E-05 | 1 L1 _ | Znrf1 |
| Ccnd3 | 8.33E-10 | 0.357524 | 0.63 | 0.51 | 1.68E-05 | 1 L1 _ | Ccnd3 |
| Serbp1 | 5.93E-10 | 0.305894 | 0.906 | 0.897 | 1.19E-05 | 1 L1 _ | Serbp1 |
| Rpa2 | 5.93E-10 | 0.250805 | 0.205 | 0.098 | 1.19E-05 | 1 L1 _ | Rpa2 |
| Eif5a | 4.61E-10 | 0.34861 | 0.871 | 0.841 | 9.29E-06 | 1 L1 _ | Eif5a |
| Cetn2 | 3.64E-10 | 0.367837 | 0.559 | 0.39 | 7.34E-06 | 1 L1 _ | Cetn2 |
| Ppm1g | 3.09E-10 | 0.267984 | 0.543 | 0.389 | 6.23E-06 | 1 L1 _ | Ppm1g |
| Cct5 | 2.86E-10 | 0.313012 | 0.713 | 0.59 | 5.77E-06 | 1 L1 _ | Cct5 |
| Sat1 | 2.72E-10 | 0.542933 | 0.752 | 0.679 | 5.47E-06 | 1 L1 _ | Sat1 |
| Vgll4 | 2.65E-10 | 0.304905 | 0.586 | 0.443 | 5.33E-06 | 1 L1 _ | Vgll4 |
| Tcirg1 | 2.58E-10 | 0.295293 | 0.274 | 0.146 | 5.21E-06 | 1 L1 _ | Tcirg1 |
| Myo1e | 2.55E-10 | 0.253529 | 0.432 | 0.279 | 5.13E-06 | 1 L1 _ | Myo1e |
| Nt5c | 1.71E-10 | 0.334423 | 0.522 | 0.361 | 3.44E-06 | 1 L1 _ | Nt5c |
| Aif1 | 1.30E-10 | 0.976766 | 0.159 | 0.069 | 2.62E-06 | 1 L1 _ | Aif1 |
| Cct7 | 1.25E-10 | 0.328334 | 0.692 | 0.555 | 2.53E-06 | 1 L1 _ | Cct7 |
| Npc1 | 1.16E-10 | 0.252288 | 0.326 | 0.187 | 2.34E-06 | 1 L1 _ | Npc1 |
| Morc3 | 1.16E-10 | 0.284964 | 0.393 | 0.235 | 2.33E-06 | 1 L1 _ | Morc3 |
| Rhog | 1.14E-10 | 0.371561 | 0.469 | 0.312 | 2.30E-06 | 1 L1 _ | Rhog |
| Sec61b | 1.13E-10 | 0.300252 | 0.851 | 0.783 | 2.27E-06 | 1 L1 _ | Sec61b |
| H2-Aa | 1.09E-10 | 1.730697 | 0.322 | 0.196 | 2.19E-06 | 1 L1 _ | H2-Aa |
| Mfsd10 | 9.20E-11 | 0.281163 | 0.437 | 0.28 | 1.85E-06 | 1 L1 _ | Mfsd10 |
| Mef2a2 | 8.88E-11 | 0.294574 | 0.754 | 0.64 | 1.79E-06 | 1 L1 _ | Mef2a |
| Ptbp31 | 8.77E-11 | 0.296919 | 0.697 | 0.566 | 1.77E-06 | 1 L1 _ | Ptbp3 |
| Vps13a | 8.64E-11 | 0.292738 | 0.425 | 0.258 | 1.74E-06 | 1 L1 _ | Vps13a |
| Mndal1 | 8.53E-11 | 0.308049 | 0.754 | 0.649 | 1.72E-06 | 1 L1 _ | Mndal |
| Mrps23 | 8.12E-11 | 0.26649 | 0.389 | 0.236 | 1.64E-06 | 1 L1 _ | Mrps23 |
| Tmem189 | 7.89E-11 | 0.257176 | 0.251 | 0.125 | 1.59E-06 | 1 L1 _ | Tmem189 |

| | | | | | | | |
|---|---|---|---|---|---|---|---|
| Ncl | 7.37E-11 | 0.350326 | 0.876 | 0.843 | 1.48E-06 | 1 L1 _ | Ncl |
| Tmem160 | 7.24E-11 | 0.373942 | 0.651 | 0.513 | 1.46E-06 | 1 L1 _ | Tmem160 |
| Mcl11 | 6.64E-11 | 0.370126 | 0.738 | 0.605 | 1.34E-06 | 1 L1 _ | Mcl1 |
| Cap1 | 6.25E-11 | 0.302057 | 0.478 | 0.316 | 1.26E-06 | 1 L1 _ | Cap1 |
| Chd3 | 6.08E-11 | 0.350577 | 0.595 | 0.423 | 1.22E-06 | 1 L1 _ | Chd3 |
| Pet100 | 4.97E-11 | 0.293334 | 0.536 | 0.376 | 1.00E-06 | 1 L1 _ | Pet100 |
| Lbr | 4.45E-11 | 0.252358 | 0.34 | 0.191 | 8.95E-07 | 1 L1 _ | Lbr |
| Eif4b | 3.91E-11 | 0.322623 | 0.598 | 0.438 | 7.87E-07 | 1 L1 _ | Eif4b |
| H2-Ke6 | 3.86E-11 | 0.315282 | 0.359 | 0.21 | 7.78E-07 | 1 L1 _ | H2-Ke6 |
| 2510002D24Rik | 3.68E-11 | 0.257758 | 0.271 | 0.139 | 7.42E-07 | 1 L1 _ | 2510002D24Rik |
| Pcna | 3.35E-11 | 0.256016 | 0.425 | 0.261 | 6.74E-07 | 1 L1 _ | Pcna |
| Cirbp | 2.81E-11 | 0.363536 | 0.595 | 0.438 | 5.66E-07 | 1 L1 _ | Cirbp |
| Itpr2 | 2.72E-11 | 0.287668 | 0.501 | 0.323 | 5.47E-07 | 1 L1 _ | Itpr2 |
| Zfas1 | 2.47E-11 | 0.282206 | 0.338 | 0.19 | 4.97E-07 | 1 L1 _ | Zfas1 |
| Dnajc9 | 2.42E-11 | 0.258015 | 0.278 | 0.143 | 4.88E-07 | 1 L1 _ | Dnajc9 |
| Fkbp31 | 2.35E-11 | 0.362056 | 0.699 | 0.573 | 4.73E-07 | 1 L1 _ | Fkbp3 |
| Ptpn2 | 2.22E-11 | 0.297324 | 0.331 | 0.189 | 4.48E-07 | 1 L1 _ | Ptpn2 |
| Tpp2 | 1.77E-11 | 0.356947 | 0.52 | 0.367 | 3.57E-07 | 1 L1 _ | Tpp2 |
| Tpst21 | 1.68E-11 | 0.268931 | 0.446 | 0.275 | 3.37E-07 | 1 L1 _ | Tpst2 |
| Igbp1 | 1.36E-11 | 0.332276 | 0.441 | 0.279 | 2.73E-07 | 1 L1 _ | Igbp1 |
| Stk24 | 1.32E-11 | 0.341411 | 0.559 | 0.388 | 2.66E-07 | 1 L1 _ | Stk24 |
| 1700017B05Rik | 1.11E-11 | 0.348736 | 0.267 | 0.132 | 2.23E-07 | 1 L1 _ | 1700017B05Rik |
| Fbl | 1.06E-11 | 0.350187 | 0.457 | 0.301 | 2.14E-07 | 1 L1 _ | Fbl |
| Rbl2 | 1.01E-11 | 0.328823 | 0.407 | 0.255 | 2.04E-07 | 1 L1 _ | Rbl2 |
| Clta | 9.90E-12 | 0.340412 | 0.862 | 0.791 | 1.99E-07 | 1 L1 _ | Clta |
| Nans | 9.21E-12 | 0.304076 | 0.395 | 0.245 | 1.85E-07 | 1 L1 _ | Nans |
| Tap1 | 6.94E-12 | 0.306895 | 0.464 | 0.291 | 1.40E-07 | 1 L1 _ | Tap1 |
| Ankrd11 | 6.89E-12 | 0.329981 | 0.807 | 0.717 | 1.39E-07 | 1 L1 _ | Ankrd11 |
| Atp5l | 6.88E-12 | 0.282524 | 0.91 | 0.897 | 1.39E-07 | 1 L1 _ | Atp5l |
| Plgrkt | 6.27E-12 | 0.340381 | 0.611 | 0.457 | 1.26E-07 | 1 L1 _ | Plgrkt |
| Cx3cr1 | 5.16E-12 | 0.704471 | 0.115 | 0.037 | 1.04E-07 | 1 L1 _ | Cx3cr1 |
| Arhgef31 | 4.84E-12 | 0.297536 | 0.209 | 0.092 | 9.75E-08 | 1 L1 _ | Arhgef3 |
| Ntan1 | 4.49E-12 | 0.425055 | 0.662 | 0.543 | 9.04E-08 | 1 L1 _ | Ntan1 |
| Slbp | 4.13E-12 | 0.306766 | 0.474 | 0.303 | 8.32E-08 | 1 L1 _ | Slbp |
| Capza1 | 4.08E-12 | 0.349382 | 0.545 | 0.38 | 8.21E-08 | 1 L1 _ | Capza1 |
| Klk1 | 4.02E-12 | 0.440498 | 0.529 | 0.349 | 8.10E-08 | 1 L1 _ | Klk1 |
| Hdgfl2 | 3.93E-12 | 0.375915 | 0.4 | 0.247 | 7.91E-08 | 1 L1 _ | Hdgfl2 |
| Ikbkb | 3.85E-12 | 0.276846 | 0.448 | 0.281 | 7.75E-08 | 1 L1 _ | Ikbkb |
| Ndufa3 | 3.44E-12 | 0.33818 | 0.779 | 0.682 | 6.94E-08 | 1 L1 _ | Ndufa3 |
| Skap2 | 3.39E-12 | 0.309916 | 0.262 | 0.132 | 6.83E-08 | 1 L1 _ | Skap2 |
| Cox7c1 | 2.81E-12 | 0.262024 | 0.913 | 0.91 | 5.66E-08 | 1 L1 _ | Cox7c |

| | | | | | | | |
|---|---|---|---|---|---|---|---|
| Mrpl57 | 2.59E-12 | 0.41094 | 0.616 | 0.444 | 5.21E-08 | 1 L1 _ | Mrpl57 |
| Higd1a | 2.43E-12 | 0.352871 | 0.602 | 0.437 | 4.89E-08 | 1 L1 _ | Higd1a |
| Eif3e1 | 2.13E-12 | 0.378405 | 0.749 | 0.641 | 4.28E-08 | 1 L1 _ | Eif3e |
| Fnbp1 | 2.06E-12 | 0.371825 | 0.671 | 0.499 | 4.15E-08 | 1 L1 _ | Fnbp1 |
| H2-Ab1 | 2.05E-12 | 1.762007 | 0.306 | 0.176 | 4.13E-08 | 1 L1 _ | H2-Ab1 |
| Exosc8 | 1.96E-12 | 0.295297 | 0.411 | 0.245 | 3.95E-08 | 1 L1 _ | Exosc8 |
| Tpm31 | 1.85E-12 | 0.345642 | 0.823 | 0.736 | 3.72E-08 | 1 L1 _ | Tpm3 |
| Arpc3 | 9.47E-13 | 0.304403 | 0.867 | 0.785 | 1.91E-08 | 1 L1 _ | Arpc3 |
| Clcn3 | 8.36E-13 | 0.261597 | 0.306 | 0.155 | 1.68E-08 | 1 L1 _ | Clcn3 |
| Ndufa41 | 8.21E-13 | 0.334336 | 0.837 | 0.714 | 1.65E-08 | 1 L1 _ | Ndufa4 |
| Psmb1 | 6.98E-13 | 0.37428 | 0.832 | 0.742 | 1.41E-08 | 1 L1 _ | Psmb1 |
| Reep51 | 6.74E-13 | 0.369305 | 0.814 | 0.729 | 1.36E-08 | 1 L1 _ | Reep5 |
| Ddx39 | 6.52E-13 | 0.252485 | 0.308 | 0.159 | 1.31E-08 | 1 L1 _ | Ddx39 |
| Oaz1 | 6.28E-13 | 0.265713 | 0.954 | 0.936 | 1.26E-08 | 1 L1 _ | Oaz1 |
| Gtf3a | 5.39E-13 | 0.281221 | 0.421 | 0.248 | 1.09E-08 | 1 L1 _ | Gtf3a |
| Ntpcr | 5.03E-13 | 0.406859 | 0.218 | 0.097 | 1.01E-08 | 1 L1 _ | Ntpcr |
| Trmt112 | 3.87E-13 | 0.389478 | 0.687 | 0.544 | 7.80E-09 | 1 L1 _ | Trmt112 |
| Lamtor4 | 3.53E-13 | 0.415051 | 0.69 | 0.554 | 7.10E-09 | 1 L1 _ | Lamtor4 |
| Map3k8 | 3.08E-13 | 0.258796 | 0.239 | 0.108 | 6.19E-09 | 1 L1 _ | Map3k8 |
| Hist1h1c | 2.76E-13 | 0.345303 | 0.559 | 0.37 | 5.56E-09 | 1 L1 _ | Hist1h1c |
| Psmb3 | 2.72E-13 | 0.38364 | 0.8 | 0.735 | 5.48E-09 | 1 L1 _ | Psmb3 |
| Gm17018 | 2.65E-13 | 0.262823 | 0.317 | 0.164 | 5.34E-09 | 1 L1 _ | Gm17018 |
| Snrpf | 2.43E-13 | 0.362171 | 0.708 | 0.557 | 4.89E-09 | 1 L1 _ | Snrpf |
| Fam214a | 1.76E-13 | 0.297257 | 0.274 | 0.132 | 3.55E-09 | 1 L1 _ | Fam214a |
| Rpl9-ps6 | 1.76E-13 | 0.310801 | 0.313 | 0.163 | 3.54E-09 | 1 L1 _ | Rpl9-ps6 |
| Rsbn1l1 | 1.56E-13 | 0.34102 | 0.648 | 0.465 | 3.14E-09 | 1 L1 _ | Rsbn1l |
| Nme1 | 1.50E-13 | 0.509091 | 0.701 | 0.571 | 3.03E-09 | 1 L1 _ | Nme1 |
| Nedd91 | 1.20E-13 | 0.342322 | 0.506 | 0.317 | 2.42E-09 | 1 L1 _ | Nedd9 |
| Hist1h1e | 1.19E-13 | 0.364404 | 0.554 | 0.365 | 2.39E-09 | 1 L1 _ | Hist1h1e |
| Eef1g | 7.82E-14 | 0.367633 | 0.814 | 0.752 | 1.57E-09 | 1 L1 _ | Eef1g |
| Tuba1c | 7.40E-14 | 0.374439 | 0.308 | 0.155 | 1.49E-09 | 1 L1 _ | Tuba1c |
| Birc2 | 7.27E-14 | 0.413065 | 0.462 | 0.279 | 1.46E-09 | 1 L1 _ | Birc2 |
| Magohb | 6.96E-14 | 0.267264 | 0.271 | 0.128 | 1.40E-09 | 1 L1 _ | Magohb |
| Zmiz2 | 6.32E-14 | 0.337611 | 0.292 | 0.14 | 1.27E-09 | 1 L1 _ | Zmiz2 |
| Arpc5 | 5.81E-14 | 0.372725 | 0.775 | 0.665 | 1.17E-09 | 1 L1 _ | Arpc5 |
| Tgfb1 | 5.40E-14 | 0.329054 | 0.639 | 0.441 | 1.09E-09 | 1 L1 _ | Tgfb1 |
| 1-Jun | 4.36E-14 | 0.34885 | 0.832 | 0.666 | 8.78E-10 | 1 L1 _ | Jun |
| Ier5 | 3.56E-14 | 0.423418 | 0.662 | 0.474 | 7.16E-10 | 1 L1 _ | Ier5 |
| Flt3l | 3.50E-14 | 0.309265 | 0.349 | 0.18 | 7.05E-10 | 1 L1 _ | Flt3l |
| Fryl | 3.13E-14 | 0.330293 | 0.407 | 0.224 | 6.31E-10 | 1 L1 _ | Fryl |
| Phgdh | 3.10E-14 | 0.335432 | 0.287 | 0.138 | 6.24E-10 | 1 L1 _ | Phgdh |

| | | | | | | | |
|---|---|---|---|---|---|---|---|
| Mien1 | 2.85E-14 | 0.367964 | 0.572 | 0.399 | 5.73E-10 | 1 L1 _ | Mien1 |
| Commd4 | 2.39E-14 | 0.429972 | 0.46 | 0.286 | 4.81E-10 | 1 L1 _ | Commd4 |
| Wdr1 | 2.17E-14 | 0.445167 | 0.522 | 0.348 | 4.36E-10 | 1 L1 _ | Wdr1 |
| Kxd1 | 1.92E-14 | 0.299227 | 0.499 | 0.307 | 3.87E-10 | 1 L1 _ | Kxd1 |
| Hnrnpa2b1 | 1.77E-14 | 0.391892 | 0.899 | 0.876 | 3.56E-10 | 1 L1 _ | Hnrnpa2b1 |
| Ostf12 | 1.70E-14 | 0.344226 | 0.736 | 0.583 | 3.43E-10 | 1 L1 _ | Ostf1 |
| Atp5d | 1.62E-14 | 0.35706 | 0.851 | 0.794 | 3.25E-10 | 1 L1 _ | Atp5d |
| Dut | 1.19E-14 | 0.307769 | 0.375 | 0.201 | 2.39E-10 | 1 L1 _ | Dut |
| Ubl5 | 1.18E-14 | 0.375891 | 0.846 | 0.77 | 2.38E-10 | 1 L1 _ | Ubl5 |
| Gapdh2 | 1.15E-14 | 0.392107 | 0.92 | 0.905 | 2.31E-10 | 1 L1 _ | Gapdh |
| Mpeg1 | 1.13E-14 | 0.518139 | 0.14 | 0.044 | 2.28E-10 | 1 L1 _ | Mpeg1 |
| Snhg1 | 8.20E-15 | 0.303279 | 0.428 | 0.237 | 1.65E-10 | 1 L1 _ | Snhg1 |
| Rab8b1 | 6.25E-15 | 0.29321 | 0.531 | 0.322 | 1.26E-10 | 1 L1 _ | Rab8b |
| Card19 | 6.01E-15 | 0.291556 | 0.448 | 0.262 | 1.21E-10 | 1 L1 _ | Card19 |
| Etv3 | 5.63E-15 | 0.356732 | 0.301 | 0.144 | 1.13E-10 | 1 L1 _ | Etv3 |
| Nosip | 5.32E-15 | 0.341373 | 0.411 | 0.231 | 1.07E-10 | 1 L1 _ | Nosip |
| Sun2 | 4.99E-15 | 0.266057 | 0.384 | 0.202 | 1.00E-10 | 1 L1 _ | Sun2 |
| Ube2d3 | 4.73E-15 | 0.413708 | 0.841 | 0.751 | 9.52E-11 | 1 L1 _ | Ube2d3 |
| Crlf2 | 4.71E-15 | 0.412279 | 0.559 | 0.369 | 9.50E-11 | 1 L1 _ | Crlf2 |
| Mpp7 | 3.89E-15 | 0.321839 | 0.276 | 0.126 | 7.84E-11 | 1 L1 _ | Mpp7 |
| Slc25a5 | 3.74E-15 | 0.407138 | 0.816 | 0.679 | 7.53E-11 | 1 L1 _ | Slc25a5 |
| Ciapin1 | 3.59E-15 | 0.263383 | 0.329 | 0.159 | 7.24E-11 | 1 L1 _ | Ciapin1 |
| Tut7 | 3.13E-15 | 0.262403 | 0.607 | 0.385 | 6.31E-11 | 1 L1 _ | Tut7 |
| C1qa | 2.93E-15 | 2.456217 | 0.338 | 0.182 | 5.91E-11 | 1 L1 _ | C1qa |
| Irf11 | 2.66E-15 | 0.3541 | 0.529 | 0.319 | 5.36E-11 | 1 L1 _ | Irf1 |
| Ubac2 | 2.65E-15 | 0.342046 | 0.411 | 0.23 | 5.34E-11 | 1 L1 _ | Ubac2 |
| Zfp318 | 2.44E-15 | 0.405137 | 0.409 | 0.237 | 4.91E-11 | 1 L1 _ | Zfp318 |
| Prkcb | 2.41E-15 | 0.286516 | 0.267 | 0.117 | 4.86E-11 | 1 L1 _ | Prkcb |
| Arhgap17 | 2.09E-15 | 0.361212 | 0.506 | 0.308 | 4.21E-11 | 1 L1 _ | Arhgap17 |
| Rpl10-ps3 | 2.07E-15 | 0.271794 | 0.322 | 0.156 | 4.17E-11 | 1 L1 _ | Rpl10-ps3 |
| Sdf2l1 | 1.76E-15 | 0.370411 | 0.368 | 0.197 | 3.54E-11 | 1 L1 _ | Sdf2l1 |
| Asxl2 | 1.53E-15 | 0.313901 | 0.503 | 0.304 | 3.08E-11 | 1 L1 _ | Asxl2 |
| Rpap2 | 1.52E-15 | 0.259577 | 0.257 | 0.11 | 3.06E-11 | 1 L1 _ | Rpap2 |
| Abr | 1.49E-15 | 0.288905 | 0.345 | 0.169 | 3.00E-11 | 1 L1 _ | Abr |
| Tpd52 | 1.38E-15 | 0.260058 | 0.186 | 0.067 | 2.78E-11 | 1 L1 _ | Tpd52 |
| Arl6ip1 | 1.27E-15 | 0.499136 | 0.821 | 0.741 | 2.55E-11 | 1 L1 _ | Arl6ip1 |
| Commd7 | 1.24E-15 | 0.469681 | 0.559 | 0.371 | 2.49E-11 | 1 L1 _ | Commd7 |
| Ptger4 | 1.19E-15 | 0.264549 | 0.29 | 0.133 | 2.40E-11 | 1 L1 _ | Ptger4 |
| Cox7b | 1.19E-15 | 0.362863 | 0.839 | 0.768 | 2.39E-11 | 1 L1 _ | Cox7b |
| Lsm5 | 1.12E-15 | 0.334069 | 0.492 | 0.301 | 2.25E-11 | 1 L1 _ | Lsm5 |
| Rpgrip1 | 1.07E-15 | 0.266693 | 0.255 | 0.109 | 2.16E-11 | 1 L1 _ | Rpgrip1 |

| | | | | | | | |
|---|---|---|---|---|---|---|---|
| Snhg8 | 1.05E-15 | 0.351151 | 0.623 | 0.418 | 2.11E-11 | 1 L1 _ | Snhg8 |
| Crlf3 | 9.30E-16 | 0.320728 | 0.423 | 0.229 | 1.87E-11 | 1 L1 _ | Crlf3 |
| Cd74 | 8.97E-16 | 2.678858 | 0.497 | 0.33 | 1.81E-11 | 1 L1 _ | Cd74 |
| Smc4 | 8.51E-16 | 0.331242 | 0.398 | 0.215 | 1.71E-11 | 1 L1 _ | Smc4 |
| Nhp2 | 8.20E-16 | 0.352307 | 0.49 | 0.307 | 1.65E-11 | 1 L1 _ | Nhp2 |
| Ak2 | 8.13E-16 | 0.409402 | 0.437 | 0.26 | 1.64E-11 | 1 L1 _ | Ak2 |
| Coq10b | 7.60E-16 | 0.38602 | 0.476 | 0.281 | 1.53E-11 | 1 L1 _ | Coq10b |
| Mettl23 | 6.05E-16 | 0.44679 | 0.522 | 0.328 | 1.22E-11 | 1 L1 _ | Mettl23 |
| Zfp512 | 4.68E-16 | 0.328305 | 0.405 | 0.22 | 9.43E-12 | 1 L1 _ | Zfp512 |
| Supt4a | 4.53E-16 | 0.434125 | 0.68 | 0.513 | 9.12E-12 | 1 L1 _ | Supt4a |
| Git2 | 3.98E-16 | 0.395974 | 0.575 | 0.371 | 8.01E-12 | 1 L1 _ | Git2 |
| Hnrnpf1 | 3.66E-16 | 0.466514 | 0.807 | 0.715 | 7.38E-12 | 1 L1 _ | Hnrnpf |
| Cox5b | 3.44E-16 | 0.441414 | 0.862 | 0.807 | 6.93E-12 | 1 L1 _ | Cox5b |
| Taldo1 | 3.43E-16 | 0.420559 | 0.71 | 0.563 | 6.90E-12 | 1 L1 _ | Taldo1 |
| Tapbpl | 3.36E-16 | 0.350143 | 0.347 | 0.169 | 6.78E-12 | 1 L1 _ | Tapbpl |
| Gas5 | 3.14E-16 | 0.410498 | 0.89 | 0.675 | 6.33E-12 | 1 L1 _ | Gas5 |
| Spi1 | 2.37E-16 | 0.625844 | 0.14 | 0.04 | 4.77E-12 | 1 L1 _ | Spi1 |
| Scly | 2.31E-16 | 0.278107 | 0.246 | 0.099 | 4.66E-12 | 1 L1 _ | Scly |
| Csnk2b | 2.22E-16 | 0.408465 | 0.674 | 0.503 | 4.48E-12 | 1 L1 _ | Csnk2b |
| Sipa11 | 2.11E-16 | 0.329059 | 0.607 | 0.391 | 4.24E-12 | 1 L1 _ | Sipa1 |
| Tex9 | 2.02E-16 | 0.289954 | 0.202 | 0.074 | 4.06E-12 | 1 L1 _ | Tex9 |
| Lyz2 | 1.94E-16 | 1.402654 | 0.262 | 0.117 | 3.90E-12 | 1 L1 _ | Lyz2 |
| Sp100 | 1.91E-16 | 0.474622 | 0.703 | 0.515 | 3.85E-12 | 1 L1 _ | Sp100 |
| Fgl2 | 1.61E-16 | 0.352428 | 0.579 | 0.372 | 3.23E-12 | 1 L1 _ | Fgl2 |
| Sipa1l1 | 1.32E-16 | 0.295604 | 0.317 | 0.147 | 2.65E-12 | 1 L1 _ | Sipa1l1 |
| Srrm2 | 1.05E-16 | 0.396239 | 0.906 | 0.889 | 2.11E-12 | 1 L1 _ | Srrm2 |
| Rab8a | 8.45E-17 | 0.425938 | 0.598 | 0.389 | 1.70E-12 | 1 L1 _ | Rab8a |
| Bola2 | 7.66E-17 | 0.446257 | 0.669 | 0.49 | 1.54E-12 | 1 L1 _ | Bola2 |
| Zfp869 | 7.58E-17 | 0.356218 | 0.402 | 0.214 | 1.53E-12 | 1 L1 _ | Zfp869 |
| Tnik | 7.49E-17 | 0.29089 | 0.338 | 0.163 | 1.51E-12 | 1 L1 _ | Tnik |
| Atp6v1f | 7.46E-17 | 0.474761 | 0.814 | 0.725 | 1.50E-12 | 1 L1 _ | Atp6v1f |
| Tmem173 | 5.56E-17 | 0.307501 | 0.393 | 0.207 | 1.12E-12 | 1 L1 _ | Tmem173 |
| Selenoh | 4.22E-17 | 0.386866 | 0.425 | 0.226 | 8.49E-13 | 1 L1 _ | Selenoh |
| Ppp6r1 | 4.12E-17 | 0.358509 | 0.382 | 0.196 | 8.30E-13 | 1 L1 _ | Ppp6r1 |
| Btf3 | 2.85E-17 | 0.385275 | 0.887 | 0.871 | 5.74E-13 | 1 L1 _ | Btf3 |
| Dcps | 2.70E-17 | 0.270286 | 0.324 | 0.146 | 5.43E-13 | 1 L1 _ | Dcps |
| Taf15 | 2.46E-17 | 0.331278 | 0.577 | 0.362 | 4.95E-13 | 1 L1 _ | Taf15 |
| Gbp41 | 2.38E-17 | 0.344321 | 0.336 | 0.156 | 4.80E-13 | 1 L1 _ | Gbp4 |
| Tecpr1 | 2.00E-17 | 0.415933 | 0.448 | 0.245 | 4.03E-13 | 1 L1 _ | Tecpr1 |
| Fosb | 1.74E-17 | 0.483401 | 0.584 | 0.372 | 3.51E-13 | 1 L1 _ | Fosb |
| Tmem134 | 1.72E-17 | 0.380608 | 0.586 | 0.373 | 3.46E-13 | 1 L1 _ | Tmem134 |

| | | | | | | | |
|---|---|---|---|---|---|---|---|
| Psmb10 | 1.49E-17 | 0.39173 | 0.745 | 0.546 | 3.00E-13 | 1 L1 _ | Psmb10 |
| Rnf114 | 1.41E-17 | 0.250401 | 0.331 | 0.15 | 2.84E-13 | 1 L1 _ | Rnf114 |
| Lypla2 | 1.14E-17 | 0.326565 | 0.379 | 0.191 | 2.29E-13 | 1 L1 _ | Lypla2 |
| Tut4 | 1.11E-17 | 0.388037 | 0.549 | 0.332 | 2.24E-13 | 1 L1 _ | Tut4 |
| Nol7 | 1.08E-17 | 0.389525 | 0.726 | 0.539 | 2.17E-13 | 1 L1 _ | Nol7 |
| Cox7a2l | 1.07E-17 | 0.440329 | 0.814 | 0.736 | 2.16E-13 | 1 L1 _ | Cox7a2l |
| Vamp8 | 9.44E-18 | 0.43601 | 0.676 | 0.493 | 1.90E-13 | 1 L1 _ | Vamp8 |
| Ppil2 | 7.81E-18 | 0.4707 | 0.517 | 0.318 | 1.57E-13 | 1 L1 _ | Ppil2 |
| Ciao2a | 7.27E-18 | 0.331567 | 0.407 | 0.213 | 1.46E-13 | 1 L1 _ | Ciao2a |
| Arpc1b | 6.00E-18 | 0.441936 | 0.89 | 0.806 | 1.21E-13 | 1 L1 _ | Arpc1b |
| Vars | 5.80E-18 | 0.33722 | 0.363 | 0.178 | 1.17E-13 | 1 L1 _ | Vars |
| Dnajc2 | 5.28E-18 | 0.339807 | 0.428 | 0.228 | 1.06E-13 | 1 L1 _ | Dnajc2 |
| Snrnp70 | 4.59E-18 | 0.401324 | 0.848 | 0.725 | 9.24E-14 | 1 L1 _ | Snrnp70 |
| Sigirr1 | 2.80E-18 | 0.323116 | 0.207 | 0.07 | 5.64E-14 | 1 L1 _ | Sigirr |
| Bin1 | 2.64E-18 | 0.396256 | 0.503 | 0.295 | 5.31E-14 | 1 L1 _ | Bin1 |
| Rnf166 | 2.02E-18 | 0.444889 | 0.476 | 0.27 | 4.07E-14 | 1 L1 _ | Rnf166 |
| Plekhj1 | 1.53E-18 | 0.457857 | 0.586 | 0.385 | 3.07E-14 | 1 L1 _ | Plekhj1 |
| Ramp1 | 1.48E-18 | 0.329556 | 0.255 | 0.097 | 2.97E-14 | 1 L1 _ | Ramp1 |
| 1810026B05Rik | 1.23E-18 | 0.478072 | 0.513 | 0.292 | 2.48E-14 | 1 L1 _ | 1810026B05Rik |
| Snx5 | 1.14E-18 | 0.518008 | 0.503 | 0.318 | 2.31E-14 | 1 L1 _ | Snx5 |
| Tomm7 | 1.05E-18 | 0.418059 | 0.867 | 0.815 | 2.12E-14 | 1 L1 _ | Tomm7 |
| Nfkbiz | 8.36E-19 | 0.378244 | 0.253 | 0.098 | 1.68E-14 | 1 L1 _ | Nfkbiz |
| Alox5ap | 7.71E-19 | 0.466559 | 0.131 | 0.031 | 1.55E-14 | 1 L1 _ | Alox5ap |
| Mllt3 | 6.10E-19 | 0.335696 | 0.329 | 0.146 | 1.23E-14 | 1 L1 _ | Mllt3 |
| Cnn22 | 4.91E-19 | 0.454645 | 0.625 | 0.403 | 9.88E-15 | 1 L1 _ | Cnn2 |
| Rp9 | 4.61E-19 | 0.481696 | 0.743 | 0.562 | 9.29E-15 | 1 L1 _ | Rp9 |
| Snrpe | 4.13E-19 | 0.478813 | 0.784 | 0.676 | 8.31E-15 | 1 L1 _ | Snrpe |
| Syngr2 | 4.06E-19 | 0.501687 | 0.538 | 0.327 | 8.18E-15 | 1 L1 _ | Syngr2 |
| Zfpm11 | 3.63E-19 | 0.357443 | 0.336 | 0.152 | 7.32E-15 | 1 L1 _ | Zfpm1 |
| Eif3m | 3.49E-19 | 0.508579 | 0.713 | 0.544 | 7.03E-15 | 1 L1 _ | Eif3m |
| Gatm | 3.25E-19 | 0.274588 | 0.108 | 0.019 | 6.55E-15 | 1 L1 _ | Gatm |
| Cbx4 | 3.24E-19 | 0.312557 | 0.301 | 0.126 | 6.53E-15 | 1 L1 _ | Cbx4 |
| Traf2 | 2.69E-19 | 0.324346 | 0.313 | 0.136 | 5.42E-15 | 1 L1 _ | Traf2 |
| Haus3 | 1.99E-19 | 0.366586 | 0.382 | 0.186 | 4.02E-15 | 1 L1 _ | Haus3 |
| Arpc4 | 1.92E-19 | 0.465094 | 0.717 | 0.55 | 3.86E-15 | 1 L1 _ | Arpc4 |
| Cox17 | 1.11E-19 | 0.433497 | 0.685 | 0.488 | 2.24E-15 | 1 L1 _ | Cox17 |
| Grap1 | 1.00E-19 | 0.39675 | 0.29 | 0.116 | 2.02E-15 | 1 L1 _ | Grap |
| Rnaseh2c | 8.44E-20 | 0.469078 | 0.625 | 0.43 | 1.70E-15 | 1 L1 _ | Rnaseh2c |
| Rpl42 | 8.08E-20 | 0.407779 | 0.88 | 0.832 | 1.63E-15 | 1 L1 _ | Rpl4 |
| Zbp1 | 6.73E-20 | 0.442923 | 0.271 | 0.107 | 1.36E-15 | 1 L1 _ | Zbp1 |
| Lsm4 | 6.58E-20 | 0.47316 | 0.68 | 0.466 | 1.33E-15 | 1 L1 _ | Lsm4 |

| | | | | | | | |
|---|---|---|---|---|---|---|---|
| Cep250 | 6.50E-20 | 0.314427 | 0.292 | 0.119 | 1.31E-15 | 1 L1 _ | Cep250 |
| Rab4b | 5.37E-20 | 0.436882 | 0.513 | 0.29 | 1.08E-15 | 1 L1 _ | Rab4b |
| Smap2 | 5.37E-20 | 0.495362 | 0.469 | 0.261 | 1.08E-15 | 1 L1 _ | Smap2 |
| Cdkn2aipnl | 5.28E-20 | 0.26446 | 0.257 | 0.094 | 1.06E-15 | 1 L1 _ | Cdkn2aipnl |
| Mast3 | 4.60E-20 | 0.329726 | 0.239 | 0.085 | 9.27E-16 | 1 L1 _ | Mast3 |
| Rnaset2a | 3.68E-20 | 0.492322 | 0.628 | 0.426 | 7.41E-16 | 1 L1 _ | Rnaset2a |
| Cyld | 3.42E-20 | 0.432887 | 0.451 | 0.24 | 6.90E-16 | 1 L1 _ | Cyld |
| Dock5 | 1.89E-20 | 0.315141 | 0.306 | 0.126 | 3.81E-16 | 1 L1 _ | Dock5 |
| Rnasek | 1.78E-20 | 0.421507 | 0.779 | 0.554 | 3.58E-16 | 1 L1 _ | Rnasek |
| Ndufa6 | 1.35E-20 | 0.495384 | 0.839 | 0.706 | 2.72E-16 | 1 L1 _ | Ndufa6 |
| Cmc1 | 1.31E-20 | 0.346558 | 0.372 | 0.173 | 2.65E-16 | 1 L1 _ | Cmc1 |
| Bhlhe40 | 1.18E-20 | 0.444141 | 0.423 | 0.216 | 2.37E-16 | 1 L1 _ | Bhlhe40 |
| Elf11 | 9.08E-21 | 0.424246 | 0.657 | 0.423 | 1.83E-16 | 1 L1 _ | Elf1 |
| Card6 | 8.49E-21 | 0.260896 | 0.257 | 0.093 | 1.71E-16 | 1 L1 _ | Card6 |
| Cks2 | 6.52E-21 | 0.30433 | 0.154 | 0.037 | 1.31E-16 | 1 L1 _ | Cks2 |
| Gm10138 | 6.42E-21 | 0.283644 | 0.177 | 0.048 | 1.29E-16 | 1 L1 _ | Gm10138 |
| Csk | 6.22E-21 | 0.45036 | 0.478 | 0.267 | 1.25E-16 | 1 L1 _ | Csk |
| Exoc6 | 4.59E-21 | 0.27082 | 0.223 | 0.071 | 9.25E-17 | 1 L1 _ | Exoc6 |
| Slfn8 | 3.98E-21 | 0.328513 | 0.29 | 0.113 | 8.01E-17 | 1 L1 _ | Slfn8 |
| C1qc | 3.73E-21 | 1.945643 | 0.269 | 0.108 | 7.52E-17 | 1 L1 _ | C1qc |
| A630072M18Rik | 3.49E-21 | 0.421081 | 0.336 | 0.145 | 7.03E-17 | 1 L1 _ | A630072M18Rik |
| Dgkd | 2.48E-21 | 0.412172 | 0.416 | 0.2 | 4.99E-17 | 1 L1 _ | Dgkd |
| Ubash3b | 2.08E-21 | 0.316512 | 0.322 | 0.133 | 4.19E-17 | 1 L1 _ | Ubash3b |
| Csf1r | 1.94E-21 | 0.629152 | 0.129 | 0.026 | 3.90E-17 | 1 L1 _ | Csf1r |
| Wfdc17 | 1.52E-21 | 0.460004 | 0.117 | 0.021 | 3.07E-17 | 1 L1 _ | Wfdc17 |
| Ms4a7 | 1.18E-21 | 1.464015 | 0.193 | 0.058 | 2.37E-17 | 1 L1 _ | Ms4a7 |
| Me2 | 1.14E-21 | 0.307083 | 0.28 | 0.105 | 2.29E-17 | 1 L1 _ | Me2 |
| C1qb | 8.85E-22 | 2.333123 | 0.372 | 0.183 | 1.78E-17 | 1 L1 _ | C1qb |
| Ctsc | 8.59E-22 | 0.713078 | 0.382 | 0.183 | 1.73E-17 | 1 L1 _ | Ctsc |
| Dcxr | 7.97E-22 | 0.318617 | 0.26 | 0.093 | 1.60E-17 | 1 L1 _ | Dcxr |
| Ppp1r10 | 6.83E-22 | 0.445506 | 0.572 | 0.332 | 1.38E-17 | 1 L1 _ | Ppp1r10 |
| Cdk6 | 6.13E-22 | 0.388214 | 0.366 | 0.16 | 1.24E-17 | 1 L1 _ | Cdk6 |
| Vsir | 5.21E-22 | 0.42019 | 0.359 | 0.161 | 1.05E-17 | 1 L1 _ | Vsir |
| B930036N10Rik1 | 4.78E-22 | 0.357903 | 0.234 | 0.077 | 9.62E-18 | 1 L1 _ | B930036N10Rik |
| Sh2d3c1 | 4.32E-22 | 0.267641 | 0.308 | 0.116 | 8.70E-18 | 1 L1 _ | Sh2d3c |
| Irf8 | 3.98E-22 | 0.556147 | 0.382 | 0.179 | 8.02E-18 | 1 L1 _ | Irf8 |
| Vasp1 | 3.70E-22 | 0.461435 | 0.639 | 0.414 | 7.46E-18 | 1 L1 _ | Vasp |
| D16Ertd472e | 3.65E-22 | 0.378306 | 0.331 | 0.137 | 7.35E-18 | 1 L1 _ | D16Ertd472e |
| Gm47664 | 3.55E-22 | 0.25628 | 0.154 | 0.035 | 7.16E-18 | 1 L1 _ | Gm47664 |
| Fuca2 | 3.09E-22 | 0.2732 | 0.244 | 0.081 | 6.22E-18 | 1 L1 _ | Fuca2 |
| Hvcn11 | 2.89E-22 | 0.329403 | 0.31 | 0.125 | 5.82E-18 | 1 L1 _ | Hvcn1 |

| | | | | | | | |
|---|---|---|---|---|---|---|---|
| Rpl31 | 2.23E-22 | 0.424672 | 0.917 | 0.871 | 4.50E-18 | 1 L1 _ | Rpl31 |
| Capzb | 1.66E-22 | 0.526611 | 0.777 | 0.632 | 3.34E-18 | 1 L1 _ | Capzb |
| Ppp1r15a1 | 1.54E-22 | 0.490989 | 0.471 | 0.247 | 3.11E-18 | 1 L1 _ | Ppp1r15a |
| Slc3a21 | 1.27E-22 | 0.518148 | 0.752 | 0.55 | 2.57E-18 | 1 L1 _ | Slc3a2 |
| Cep192 | 9.65E-23 | 0.343322 | 0.271 | 0.097 | 1.94E-18 | 1 L1 _ | Cep192 |
| Cd300c2 | 9.32E-23 | 0.413729 | 0.115 | 0.018 | 1.88E-18 | 1 L1 _ | Cd300c2 |
| Evi2a | 9.21E-23 | 0.250628 | 0.14 | 0.028 | 1.85E-18 | 1 L1 _ | Evi2a |
| Chsy1 | 8.74E-23 | 0.395756 | 0.366 | 0.164 | 1.76E-18 | 1 L1 _ | Chsy1 |
| Cdc42se2 | 8.08E-23 | 0.43933 | 0.448 | 0.23 | 1.63E-18 | 1 L1 _ | Cdc42se2 |
| Saraf1 | 7.01E-23 | 0.525262 | 0.759 | 0.594 | 1.41E-18 | 1 L1 _ | Saraf |
| Atp5h | 6.65E-23 | 0.472348 | 0.908 | 0.871 | 1.34E-18 | 1 L1 _ | Atp5h |
| Rnaset2b | 6.48E-23 | 0.522777 | 0.593 | 0.375 | 1.30E-18 | 1 L1 _ | Rnaset2b |
| Nsa2 | 4.62E-23 | 0.513951 | 0.846 | 0.76 | 9.31E-19 | 1 L1 _ | Nsa2 |
| Gm2000 | 3.99E-23 | 0.406459 | 0.556 | 0.318 | 8.05E-19 | 1 L1 _ | Gm2000 |
| Nr4a2 | 3.20E-23 | 0.353374 | 0.251 | 0.083 | 6.44E-19 | 1 L1 _ | Nr4a2 |
| Hspa8 | 2.85E-23 | 0.4804 | 0.959 | 0.933 | 5.73E-19 | 1 L1 _ | Hspa8 |
| Dbnl | 2.08E-23 | 0.441028 | 0.545 | 0.307 | 4.20E-19 | 1 L1 _ | Dbnl |
| Lrrfip1 | 1.43E-23 | 0.615683 | 0.641 | 0.444 | 2.89E-19 | 1 L1 _ | Lrrfip1 |
| Phf11b | 9.96E-24 | 0.337614 | 0.189 | 0.049 | 2.01E-19 | 1 L1 _ | Phf11b |
| Il7r | 9.24E-24 | 0.312919 | 0.101 | 0.012 | 1.86E-19 | 1 L1 _ | Il7r |
| Elmo11 | 8.42E-24 | 0.313128 | 0.308 | 0.116 | 1.70E-19 | 1 L1 _ | Elmo1 |
| Gpi1 | 8.17E-24 | 0.571425 | 0.749 | 0.584 | 1.64E-19 | 1 L1 _ | Gpi1 |
| Atp5mpl | 6.59E-24 | 0.499954 | 0.798 | 0.59 | 1.33E-19 | 1 L1 _ | Atp5mpl |
| Tmem179b | 6.35E-24 | 0.410594 | 0.485 | 0.246 | 1.28E-19 | 1 L1 _ | Tmem179b |
| Tsc22d4 | 4.46E-24 | 0.584518 | 0.754 | 0.607 | 8.99E-20 | 1 L1 _ | Tsc22d4 |
| Nfatc1 | 4.36E-24 | 0.282454 | 0.276 | 0.095 | 8.78E-20 | 1 L1 _ | Nfatc1 |
| Mgat4a1 | 3.80E-24 | 0.251968 | 0.248 | 0.077 | 7.65E-20 | 1 L1 _ | Mgat4a |
| Cox4i1 | 2.82E-24 | 0.415794 | 0.947 | 0.933 | 5.68E-20 | 1 L1 _ | Cox4i1 |
| Slc38a2 | 2.47E-24 | 0.708381 | 0.83 | 0.726 | 4.98E-20 | 1 L1 _ | Slc38a2 |
| Tra2b | 2.12E-24 | 0.474506 | 0.782 | 0.59 | 4.26E-20 | 1 L1 _ | Tra2b |
| Nlrc5 | 1.99E-24 | 0.260196 | 0.177 | 0.042 | 4.01E-20 | 1 L1 _ | Nlrc5 |
| Strbp | 1.73E-24 | 0.298077 | 0.306 | 0.113 | 3.49E-20 | 1 L1 _ | Strbp |
| Atp5md | 1.20E-24 | 0.480817 | 0.805 | 0.555 | 2.43E-20 | 1 L1 _ | Atp5md |
| Ppp1ca | 1.19E-24 | 0.524431 | 0.834 | 0.715 | 2.40E-20 | 1 L1 _ | Ppp1ca |
| Ube2i | 1.07E-24 | 0.532587 | 0.775 | 0.619 | 2.15E-20 | 1 L1 _ | Ube2i |
| Cln31 | 9.45E-25 | 0.374286 | 0.297 | 0.107 | 1.90E-20 | 1 L1 _ | Cln3 |
| Adgre1 | 9.15E-25 | 0.500539 | 0.108 | 0.014 | 1.84E-20 | 1 L1 _ | Adgre1 |
| Lta4h | 5.52E-25 | 0.522069 | 0.517 | 0.284 | 1.11E-20 | 1 L1 _ | Lta4h |
| Elf4 | 1.85E-25 | 0.263425 | 0.234 | 0.068 | 3.73E-21 | 1 L1 _ | Elf4 |
| Slamf9 | 1.80E-25 | 0.600882 | 0.133 | 0.023 | 3.63E-21 | 1 L1 _ | Slamf9 |
| Cntrl | 1.32E-25 | 0.426812 | 0.402 | 0.176 | 2.66E-21 | 1 L1 _ | Cntrl |

| | | | | | | | |
|---|---|---|---|---|---|---|---|
| Rabgap1l | 9.48E-26 | 0.450868 | 0.474 | 0.234 | 1.91E-21 | 1 L1 _ | Rabgap1l |
| Atp11b | 6.81E-26 | 0.511184 | 0.501 | 0.26 | 1.37E-21 | 1 L1 _ | Atp11b |
| Psme2 | 5.56E-26 | 0.592977 | 0.766 | 0.593 | 1.12E-21 | 1 L1 _ | Psme2 |
| Vps28 | 5.14E-26 | 0.541386 | 0.731 | 0.508 | 1.04E-21 | 1 L1 _ | Vps28 |
| Rfc2 | 4.50E-26 | 0.407827 | 0.393 | 0.165 | 9.06E-22 | 1 L1 _ | Rfc2 |
| Unc119 | 3.15E-26 | 0.421412 | 0.407 | 0.181 | 6.35E-22 | 1 L1 _ | Unc119 |
| Eif3k | 2.84E-26 | 0.496808 | 0.853 | 0.759 | 5.71E-22 | 1 L1 _ | Eif3k |
| Ctsd | 2.64E-26 | 0.585935 | 0.922 | 0.82 | 5.32E-22 | 1 L1 _ | Ctsd |
| Chd2 | 2.29E-26 | 0.58414 | 0.628 | 0.369 | 4.61E-22 | 1 L1 _ | Chd2 |
| Casp8 | 2.21E-26 | 0.349943 | 0.418 | 0.178 | 4.45E-22 | 1 L1 _ | Casp8 |
| Tle4 | 1.89E-26 | 0.471518 | 0.533 | 0.275 | 3.80E-22 | 1 L1 _ | Tle4 |
| Prdx6 | 1.03E-26 | 0.624222 | 0.724 | 0.527 | 2.08E-22 | 1 L1 _ | Prdx6 |
| 2310001H17Rik | 9.71E-27 | 0.339169 | 0.234 | 0.066 | 1.96E-22 | 1 L1 _ | 2310001H17Rik |
| Stk101 | 7.69E-27 | 0.397314 | 0.326 | 0.117 | 1.55E-22 | 1 L1 _ | Stk10 |
| Itpr31 | 4.32E-27 | 0.321619 | 0.225 | 0.06 | 8.69E-23 | 1 L1 _ | Itpr3 |
| Eef1d | 4.04E-27 | 0.526725 | 0.878 | 0.809 | 8.14E-23 | 1 L1 _ | Eef1d |
| 4930581F22Rik | 3.76E-27 | 0.379339 | 0.361 | 0.137 | 7.58E-23 | 1 L1 _ | 4930581F22Rik |
| Ly86 | 3.65E-27 | 0.55741 | 0.122 | 0.016 | 7.34E-23 | 1 L1 _ | Ly86 |
| Ifi47 | 3.20E-27 | 0.398881 | 0.497 | 0.234 | 6.45E-23 | 1 L1 _ | Ifi47 |
| Camk1d1 | 2.82E-27 | 0.263467 | 0.189 | 0.042 | 5.68E-23 | 1 L1 _ | Camk1d |
| Leprotl1 | 2.65E-27 | 0.630542 | 0.662 | 0.435 | 5.33E-23 | 1 L1 _ | Leprotl1 |
| 2410006H16Rik | 1.96E-27 | 0.585687 | 0.731 | 0.477 | 3.95E-23 | 1 L1 _ | 2410006H16Rik |
| Sulf21 | 1.77E-27 | 0.475167 | 0.386 | 0.163 | 3.56E-23 | 1 L1 _ | Sulf2 |
| Ppm1j | 8.65E-28 | 0.250691 | 0.182 | 0.038 | 1.74E-23 | 1 L1 _ | Ppm1j |
| Iqgap2 | 8.49E-28 | 0.285034 | 0.156 | 0.028 | 1.71E-23 | 1 L1 _ | Iqgap2 |
| St3gal61 | 7.93E-28 | 0.37999 | 0.444 | 0.186 | 1.60E-23 | 1 L1 _ | St3gal6 |
| Cd68 | 6.98E-28 | 0.692973 | 0.149 | 0.026 | 1.40E-23 | 1 L1 _ | Cd68 |
| Cfl1 | 4.55E-28 | 0.529158 | 0.929 | 0.901 | 9.16E-24 | 1 L1 _ | Cfl1 |
| Rbm3 | 1.59E-28 | 0.622653 | 0.834 | 0.712 | 3.21E-24 | 1 L1 _ | Rbm3 |
| Lpcat2 | 1.30E-28 | 0.381113 | 0.117 | 0.013 | 2.63E-24 | 1 L1 _ | Lpcat2 |
| Apbb1ip | 9.61E-29 | 0.557659 | 0.662 | 0.398 | 1.94E-24 | 1 L1 _ | Apbb1ip |
| Map11 | 8.22E-29 | 0.30532 | 0.179 | 0.036 | 1.66E-24 | 1 L1 _ | Map11 |
| Cox6a2 | 5.64E-29 | 0.89923 | 0.108 | 0.01 | 1.14E-24 | 1 L1 _ | Cox6a2 |
| Arf6 | 5.51E-29 | 0.554304 | 0.556 | 0.302 | 1.11E-24 | 1 L1 _ | Arf6 |
| Micos13 | 4.77E-29 | 0.573569 | 0.726 | 0.486 | 9.60E-25 | 1 L1 _ | Micos13 |
| Gmnn | 3.97E-29 | 0.260666 | 0.193 | 0.041 | 7.99E-25 | 1 L1 _ | Gmnn |
| Dusp5 | 3.48E-29 | 0.312551 | 0.216 | 0.053 | 7.01E-25 | 1 L1 _ | Dusp5 |
| Tspan131 | 2.93E-29 | 0.259249 | 0.402 | 0.144 | 5.90E-25 | 1 L1 _ | Tspan13 |
| Ddx5 | 2.82E-29 | 0.436826 | 0.97 | 0.964 | 5.68E-25 | 1 L1 _ | Ddx5 |
| Ggct | 2.57E-29 | 0.438075 | 0.287 | 0.092 | 5.18E-25 | 1 L1 _ | Ggct |
| Ndufb1-ps | 2.18E-29 | 0.569197 | 0.749 | 0.53 | 4.40E-25 | 1 L1 _ | Ndufb1-ps |

| | | | | | | | |
|---|---|---|---|---|---|---|---|
| Satb1 | 1.76E-29 | 0.27826 | 0.211 | 0.049 | 3.53E-25 | 1 L1 _ | Satb1 |
| Serpinb6b1 | 8.56E-30 | 0.297053 | 0.191 | 0.039 | 1.72E-25 | 1 L1 _ | Serpinb6b |
| Plbd1 | 6.11E-30 | 0.368499 | 0.101 | 0.007 | 1.23E-25 | 1 L1 _ | Plbd1 |
| Diaph1 | 5.75E-30 | 0.570301 | 0.575 | 0.308 | 1.16E-25 | 1 L1 _ | Diaph1 |
| Abracl | 4.83E-30 | 0.572816 | 0.651 | 0.405 | 9.74E-26 | 1 L1 _ | Abracl |
| Rps17 | 4.48E-30 | 0.487352 | 0.936 | 0.906 | 9.03E-26 | 1 L1 _ | Rps17 |
| Tma7 | 3.96E-30 | 0.556744 | 0.848 | 0.722 | 7.97E-26 | 1 L1 _ | Tma7 |
| Cnp | 2.14E-30 | 0.479391 | 0.343 | 0.123 | 4.32E-26 | 1 L1 _ | Cnp |
| Pkn1 | 1.89E-30 | 0.497682 | 0.517 | 0.245 | 3.81E-26 | 1 L1 _ | Pkn1 |
| Atad2 | 1.59E-30 | 0.508317 | 0.389 | 0.151 | 3.20E-26 | 1 L1 _ | Atad2 |
| C730034F03Rik | 1.33E-30 | 0.292457 | 0.172 | 0.031 | 2.68E-26 | 1 L1 _ | C730034F03Rik |
| Hivep3 | 1.01E-30 | 0.553882 | 0.492 | 0.234 | 2.04E-26 | 1 L1 _ | Hivep3 |
| Ablim11 | 6.58E-31 | 0.609825 | 0.618 | 0.349 | 1.33E-26 | 1 L1 _ | Ablim1 |
| Rps27rt | 4.50E-31 | 0.650476 | 0.533 | 0.274 | 9.07E-27 | 1 L1 _ | Rps27rt |
| Glrx | 3.06E-31 | 0.457627 | 0.347 | 0.122 | 6.15E-27 | 1 L1 _ | Glrx |
| Trdc1 | 1.90E-31 | 0.272069 | 0.115 | 0.01 | 3.82E-27 | 1 L1 _ | Trdc |
| Socs1 | 1.87E-31 | 0.498205 | 0.349 | 0.126 | 3.76E-27 | 1 L1 _ | Socs1 |
| Irf5 | 1.11E-31 | 0.296285 | 0.161 | 0.025 | 2.24E-27 | 1 L1 _ | Irf5 |
| Tab2 | 1.11E-31 | 0.663968 | 0.623 | 0.36 | 2.23E-27 | 1 L1 _ | Tab2 |
| Pld4 | 9.03E-32 | 0.449687 | 0.129 | 0.014 | 1.82E-27 | 1 L1 _ | Pld4 |
| Mvb12b | 8.12E-32 | 0.35853 | 0.34 | 0.113 | 1.64E-27 | 1 L1 _ | Mvb12b |
| Sub1 | 5.36E-32 | 0.65463 | 0.874 | 0.777 | 1.08E-27 | 1 L1 _ | Sub1 |
| Uqcrh | 3.03E-32 | 0.526996 | 0.915 | 0.882 | 6.10E-28 | 1 L1 _ | Uqcrh |
| Gpsm3 | 2.47E-32 | 0.774509 | 0.687 | 0.423 | 4.98E-28 | 1 L1 _ | Gpsm3 |
| Cybb | 2.29E-32 | 0.561421 | 0.113 | 0.009 | 4.60E-28 | 1 L1 _ | Cybb |
| Gzma | 2.23E-32 | 2.273683 | 0.218 | 0.051 | 4.49E-28 | 1 L1 _ | Gzma |
| Rapgef6 | 8.25E-33 | 0.669212 | 0.651 | 0.39 | 1.66E-28 | 1 L1 _ | Rapgef6 |
| Lsp1 | 5.10E-33 | 0.688815 | 0.72 | 0.462 | 1.03E-28 | 1 L1 _ | Lsp1 |
| Spata131 | 4.72E-33 | 0.369643 | 0.271 | 0.072 | 9.51E-29 | 1 L1 _ | Spata13 |
| Slfn2 | 4.11E-33 | 0.691905 | 0.552 | 0.272 | 8.27E-29 | 1 L1 _ | Slfn2 |
| Zdhhc181 | 2.90E-33 | 0.482728 | 0.423 | 0.163 | 5.84E-29 | 1 L1 _ | Zdhhc18 |
| Chl1 | 1.92E-33 | 0.486876 | 0.149 | 0.019 | 3.86E-29 | 1 L1 _ | Chl1 |
| Hpgds | 1.58E-33 | 0.372982 | 0.136 | 0.015 | 3.18E-29 | 1 L1 _ | Hpgds |
| Rbpj | 1.03E-33 | 0.649462 | 0.655 | 0.398 | 2.08E-29 | 1 L1 _ | Rbpj |
| Grk2 | 1.02E-33 | 0.488193 | 0.462 | 0.193 | 2.04E-29 | 1 L1 _ | Grk2 |
| Eif3h | 3.09E-34 | 0.648832 | 0.839 | 0.714 | 6.23E-30 | 1 L1 _ | Eif3h |
| Jund | 2.94E-34 | 0.653139 | 0.908 | 0.833 | 5.92E-30 | 1 L1 _ | Jund |
| Ripor2 | 2.28E-34 | 0.281444 | 0.211 | 0.041 | 4.59E-30 | 1 L1 _ | Ripor2 |
| Rel | 2.10E-34 | 0.407994 | 0.271 | 0.07 | 4.23E-30 | 1 L1 _ | Rel |
| Ppp1r16b1 | 1.72E-34 | 0.358797 | 0.285 | 0.075 | 3.46E-30 | 1 L1 _ | Ppp1r16b |
| Nbeal2 | 1.64E-34 | 0.267454 | 0.202 | 0.038 | 3.31E-30 | 1 L1 _ | Nbeal2 |

| | | | | | | | |
|---|---|---|---|---|---|---|---|
| Zfp36l2 | 1.63E-34 | 0.755736 | 0.809 | 0.622 | 3.28E-30 | 1 L1 _ | Zfp36l2 |
| Pou2f2 | 1.19E-34 | 0.440915 | 0.154 | 0.02 | 2.40E-30 | 1 L1 _ | Pou2f2 |
| Akap13 | 1.19E-34 | 0.684801 | 0.844 | 0.662 | 2.39E-30 | 1 L1 _ | Akap13 |
| Klhl61 | 1.06E-34 | 0.35541 | 0.225 | 0.048 | 2.14E-30 | 1 L1 _ | Klhl6 |
| Peli1 | 9.92E-35 | 0.591966 | 0.563 | 0.281 | 2.00E-30 | 1 L1 _ | Peli1 |
| Ttc7 | 9.29E-35 | 0.431638 | 0.306 | 0.089 | 1.87E-30 | 1 L1 _ | Ttc7 |
| Dleu2 | 4.26E-35 | 0.443114 | 0.515 | 0.212 | 8.59E-31 | 1 L1 _ | Dleu2 |
| Taf10 | 4.11E-35 | 0.714494 | 0.738 | 0.518 | 8.28E-31 | 1 L1 _ | Taf10 |
| Gpr183 | 3.95E-35 | 0.310933 | 0.124 | 0.01 | 7.96E-31 | 1 L1 _ | Gpr183 |
| 6-Sep | 1.99E-35 | 0.522705 | 0.439 | 0.167 | 4.01E-31 | 1 L1 _ | 6-Sep |
| Rin32 | 1.32E-35 | 0.384644 | 0.329 | 0.1 | 2.65E-31 | 1 L1 _ | Rin3 |
| Cox8a1 | 1.26E-35 | 0.442961 | 0.949 | 0.945 | 2.53E-31 | 1 L1 _ | Cox8a |
| H3f3b1 | 8.68E-36 | 0.490349 | 0.984 | 0.983 | 1.75E-31 | 1 L1 _ | H3f3b |
| Tnfaip3 | 5.79E-36 | 0.368629 | 0.223 | 0.045 | 1.17E-31 | 1 L1 _ | Tnfaip3 |
| Sirt7 | 4.81E-36 | 0.539248 | 0.497 | 0.209 | 9.68E-32 | 1 L1 _ | Sirt7 |
| Lair1 | 4.38E-36 | 0.463437 | 0.126 | 0.01 | 8.83E-32 | 1 L1 _ | Lair1 |
| H2-DMa | 4.09E-36 | 0.439058 | 0.205 | 0.037 | 8.23E-32 | 1 L1 _ | H2-DMa |
| Rab19 | 3.72E-36 | 0.304436 | 0.17 | 0.024 | 7.49E-32 | 1 L1 _ | Rab19 |
| Baz1a1 | 3.35E-36 | 0.420492 | 0.372 | 0.123 | 6.75E-32 | 1 L1 _ | Baz1a |
| Gzmk | 3.05E-36 | 0.366849 | 0.101 | 0.004 | 6.13E-32 | 1 L1 _ | Gzmk |
| Ms4a6c | 3.04E-36 | 0.524781 | 0.163 | 0.022 | 6.12E-32 | 1 L1 _ | Ms4a6c |
| Otulin | 2.16E-36 | 0.666725 | 0.563 | 0.279 | 4.35E-32 | 1 L1 _ | Otulin |
| AI504432 | 1.67E-36 | 0.346543 | 0.198 | 0.033 | 3.36E-32 | 1 L1 _ | AI504432 |
| H2afj | 1.55E-36 | 0.700068 | 0.832 | 0.65 | 3.12E-32 | 1 L1 _ | H2afj |
| H2afz | 1.54E-36 | 0.731347 | 0.862 | 0.716 | 3.10E-32 | 1 L1 _ | H2afz |
| Actg11 | 8.56E-37 | 0.577374 | 0.966 | 0.941 | 1.72E-32 | 1 L1 _ | Actg1 |
| Armc7 | 6.84E-37 | 0.284469 | 0.177 | 0.025 | 1.38E-32 | 1 L1 _ | Armc7 |
| Rpl35 | 5.67E-37 | 0.594603 | 0.894 | 0.851 | 1.14E-32 | 1 L1 _ | Rpl35 |
| Arpc2 | 4.81E-37 | 0.601339 | 0.894 | 0.825 | 9.68E-33 | 1 L1 _ | Arpc2 |
| Cox5a | 3.29E-37 | 0.687394 | 0.897 | 0.74 | 6.64E-33 | 1 L1 _ | Cox5a |
| Znrf2 | 1.22E-37 | 0.459189 | 0.391 | 0.133 | 2.46E-33 | 1 L1 _ | Znrf2 |
| Rpl22l1 | 7.28E-38 | 0.645253 | 0.903 | 0.839 | 1.47E-33 | 1 L1 _ | Rpl22l1 |
| 1700028K03Rik | 6.72E-38 | 0.276694 | 0.11 | 0.005 | 1.35E-33 | 1 L1 _ | 1700028K03Rik |
| C330011M18Rik | 5.72E-38 | 0.326814 | 0.166 | 0.021 | 1.15E-33 | 1 L1 _ | C330011M18Rik |
| Cd86 | 5.72E-38 | 0.253863 | 0.113 | 0.005 | 1.15E-33 | 1 L1 _ | Cd86 |
| B2m1 | 5.25E-38 | 0.47959 | 0.989 | 0.942 | 1.06E-33 | 1 L1 _ | B2m |
| Cd28 | 4.31E-38 | 0.332555 | 0.106 | 0.004 | 8.68E-34 | 1 L1 _ | Cd28 |
| Smad71 | 3.12E-38 | 0.764063 | 0.667 | 0.355 | 6.29E-34 | 1 L1 _ | Smad7 |
| Fam102a1 | 1.43E-38 | 0.463976 | 0.368 | 0.117 | 2.88E-34 | 1 L1 _ | Fam102a |
| Atp1b11 | 8.05E-39 | 0.312763 | 0.32 | 0.082 | 1.62E-34 | 1 L1 _ | Atp1b1 |
| St8sia41 | 7.20E-39 | 0.457381 | 0.474 | 0.165 | 1.45E-34 | 1 L1 _ | St8sia4 |

| | | | | | | | |
|---|---|---|---|---|---|---|---|
| Fam107b1 | 4.90E-39 | 0.330791 | 0.29 | 0.067 | 9.88E-35 | 1 L1 _ | Fam107b |
| Def6 | 4.25E-39 | 0.395495 | 0.294 | 0.076 | 8.56E-35 | 1 L1 _ | Def6 |
| Ppia | 2.36E-39 | 0.433282 | 0.984 | 0.984 | 4.74E-35 | 1 L1 _ | Ppia |
| Afmid | 2.06E-39 | 0.340578 | 0.211 | 0.036 | 4.15E-35 | 1 L1 _ | Afmid |
| Acot7 | 1.75E-39 | 0.595731 | 0.462 | 0.187 | 3.53E-35 | 1 L1 _ | Acot7 |
| Ifi206 | 1.56E-39 | 0.404562 | 0.191 | 0.028 | 3.15E-35 | 1 L1 _ | Ifi206 |
| Zyx1 | 1.21E-39 | 0.762389 | 0.674 | 0.383 | 2.44E-35 | 1 L1 _ | Zyx |
| Isg20 | 1.14E-39 | 0.532277 | 0.363 | 0.113 | 2.29E-35 | 1 L1 _ | Isg20 |
| Atp5g2 | 2.32E-40 | 0.657777 | 0.897 | 0.813 | 4.68E-36 | 1 L1 _ | Atp5g2 |
| Cd8b1 | 2.05E-40 | 1.194014 | 0.17 | 0.021 | 4.13E-36 | 1 L1 _ | Cd8b1 |
| Lgals3 | 1.40E-40 | 0.99961 | 0.63 | 0.344 | 2.83E-36 | 1 L1 _ | Lgals3 |
| Abi31 | 4.64E-41 | 0.546599 | 0.432 | 0.142 | 9.34E-37 | 1 L1 _ | Abi3 |
| Scimp | 4.56E-41 | 0.331406 | 0.143 | 0.011 | 9.18E-37 | 1 L1 _ | Scimp |
| Mfng | 3.36E-41 | 0.38575 | 0.276 | 0.059 | 6.76E-37 | 1 L1 _ | Mfng |
| Sytl22 | 3.13E-41 | 0.358615 | 0.237 | 0.043 | 6.30E-37 | 1 L1 _ | Sytl2 |
| Gmip | 1.99E-41 | 0.456166 | 0.264 | 0.055 | 4.01E-37 | 1 L1 _ | Gmip |
| Spin2c | 1.86E-41 | 0.274065 | 0.156 | 0.015 | 3.75E-37 | 1 L1 _ | Spin2c |
| Prkcz | 1.41E-41 | 0.283741 | 0.207 | 0.031 | 2.85E-37 | 1 L1 _ | Prkcz |
| 4930520O04Rik | 1.36E-41 | 0.273716 | 0.136 | 0.008 | 2.74E-37 | 1 L1 _ | 4930520O04Rik |
| Ly6c2 | 9.19E-42 | 1.18532 | 0.189 | 0.025 | 1.85E-37 | 1 L1 _ | Ly6c2 |
| Vps37b | 6.35E-42 | 0.583879 | 0.405 | 0.132 | 1.28E-37 | 1 L1 _ | Vps37b |
| Itgam | 4.95E-42 | 0.255644 | 0.131 | 0.007 | 9.97E-38 | 1 L1 _ | Itgam |
| Pim1 | 3.84E-42 | 0.765631 | 0.467 | 0.18 | 7.73E-38 | 1 L1 _ | Pim1 |
| Eef2 | 3.16E-42 | 0.593767 | 0.954 | 0.886 | 6.35E-38 | 1 L1 _ | Eef2 |
| Arhgef1 | 2.38E-42 | 0.721015 | 0.745 | 0.462 | 4.79E-38 | 1 L1 _ | Arhgef1 |
| Gm4070 | 2.32E-42 | 0.358014 | 0.29 | 0.064 | 4.68E-38 | 1 L1 _ | Gm4070 |
| Npm3 | 1.23E-42 | 0.612765 | 0.561 | 0.252 | 2.47E-38 | 1 L1 _ | Npm3 |
| Npm1 | 4.89E-43 | 0.674485 | 0.91 | 0.868 | 9.84E-39 | 1 L1 _ | Npm1 |
| Smpd51 | 9.42E-44 | 0.354576 | 0.202 | 0.027 | 1.90E-39 | 1 L1 _ | Smpd5 |
| Tec | 2.82E-44 | 0.327552 | 0.223 | 0.035 | 5.67E-40 | 1 L1 _ | Tec |
| Atp5e | 1.62E-44 | 0.605059 | 0.94 | 0.903 | 3.26E-40 | 1 L1 _ | Atp5e |
| Tpt1 | 1.17E-44 | 0.446098 | 0.991 | 0.986 | 2.36E-40 | 1 L1 _ | Tpt1 |
| Smyd3 | 7.28E-45 | 0.437637 | 0.331 | 0.082 | 1.47E-40 | 1 L1 _ | Smyd3 |
| Lilr4b | 6.06E-45 | 0.348353 | 0.138 | 0.007 | 1.22E-40 | 1 L1 _ | Lilr4b |
| Gem | 3.16E-45 | 0.666371 | 0.366 | 0.103 | 6.36E-41 | 1 L1 _ | Gem |
| H2afy | 1.95E-45 | 0.751773 | 0.784 | 0.563 | 3.94E-41 | 1 L1 _ | H2afy |
| Rpl36a | 1.56E-45 | 0.611838 | 0.938 | 0.915 | 3.14E-41 | 1 L1 _ | Rpl36a |
| Prr13 | 1.06E-45 | 0.781474 | 0.733 | 0.446 | 2.13E-41 | 1 L1 _ | Prr13 |
| Naca | 8.93E-46 | 0.594037 | 0.94 | 0.922 | 1.80E-41 | 1 L1 _ | Naca |
| Bcl2l11 | 8.50E-46 | 0.686947 | 0.395 | 0.118 | 1.71E-41 | 1 L1 _ | Bcl2l11 |
| Prkch1 | 4.82E-46 | 0.528178 | 0.428 | 0.127 | 9.71E-42 | 1 L1 _ | Prkch |

| | | | | | | | |
|---|---|---|---|---|---|---|---|
| Casp1 | 4.07E-46 | 0.386151 | 0.267 | 0.05 | 8.20E-42 | 1 L1 _ | Casp1 |
| Plcg2 | 1.67E-46 | 0.266636 | 0.175 | 0.016 | 3.36E-42 | 1 L1 _ | Plcg2 |
| Pabpc1 | 7.45E-47 | 0.754099 | 0.876 | 0.731 | 1.50E-42 | 1 L1 _ | Pabpc1 |
| Pdcd4 | 6.00E-47 | 0.873094 | 0.766 | 0.496 | 1.21E-42 | 1 L1 _ | Pdcd4 |
| Twf2 | 5.04E-47 | 0.588762 | 0.453 | 0.155 | 1.02E-42 | 1 L1 _ | Twf2 |
| AB1246111 | 5.01E-47 | 0.32926 | 0.154 | 0.01 | 1.01E-42 | 1 L1 _ | AB124611 |
| Apobr | 4.85E-47 | 0.413143 | 0.292 | 0.059 | 9.77E-43 | 1 L1 _ | Apobr |
| Pfdn5 | 4.50E-47 | 0.662112 | 0.938 | 0.893 | 9.07E-43 | 1 L1 _ | Pfdn5 |
| Plaat3 | 4.39E-47 | 0.729555 | 0.676 | 0.327 | 8.83E-43 | 1 L1 _ | Plaat3 |
| Retreg1 | 1.88E-47 | 0.322612 | 0.191 | 0.02 | 3.79E-43 | 1 L1 _ | Retreg1 |
| Tnfsf101 | 8.65E-48 | 0.592418 | 0.503 | 0.167 | 1.74E-43 | 1 L1 _ | Tnfsf10 |
| Ppp1r18 | 5.35E-48 | 0.782458 | 0.625 | 0.304 | 1.08E-43 | 1 L1 _ | Ppp1r18 |
| Trac | 2.25E-48 | 0.868141 | 0.241 | 0.038 | 4.52E-44 | 1 L1 _ | Trac |
| Cysltr2 | 1.53E-48 | 0.259288 | 0.145 | 0.007 | 3.08E-44 | 1 L1 _ | Cysltr2 |
| Cd8a | 1.47E-48 | 0.761658 | 0.161 | 0.011 | 2.96E-44 | 1 L1 _ | Cd8a |
| Mxd1 | 1.28E-48 | 0.523242 | 0.343 | 0.083 | 2.59E-44 | 1 L1 _ | Mxd1 |
| Mapkapk3 | 8.42E-49 | 0.41635 | 0.241 | 0.037 | 1.70E-44 | 1 L1 _ | Mapkapk3 |
| Arid5a | 7.73E-49 | 0.407823 | 0.329 | 0.072 | 1.56E-44 | 1 L1 _ | Arid5a |
| Gimap9 | 6.71E-49 | 0.697421 | 0.499 | 0.18 | 1.35E-44 | 1 L1 _ | Gimap9 |
| Syk | 6.52E-49 | 0.267362 | 0.172 | 0.014 | 1.31E-44 | 1 L1 _ | Syk |
| Dapp11 | 6.23E-49 | 0.428762 | 0.26 | 0.045 | 1.26E-44 | 1 L1 _ | Dapp1 |
| Stk4 | 5.98E-49 | 0.62141 | 0.471 | 0.156 | 1.21E-44 | 1 L1 _ | Stk4 |
| Sh3bp1 | 4.49E-49 | 0.458193 | 0.329 | 0.075 | 9.03E-45 | 1 L1 _ | Sh3bp1 |
| Sms | 2.57E-49 | 0.739688 | 0.52 | 0.197 | 5.18E-45 | 1 L1 _ | Sms |
| Psmb91 | 1.58E-49 | 0.756869 | 0.667 | 0.319 | 3.19E-45 | 1 L1 _ | Psmb9 |
| Snrpg | 1.18E-49 | 0.731574 | 0.901 | 0.813 | 2.38E-45 | 1 L1 _ | Snrpg |
| Sp110 | 1.05E-49 | 0.734909 | 0.568 | 0.235 | 2.12E-45 | 1 L1 _ | Sp110 |
| Nt5e | 8.10E-50 | 0.351771 | 0.159 | 0.009 | 1.63E-45 | 1 L1 _ | Nt5e |
| AI467606 | 4.89E-50 | 0.336503 | 0.189 | 0.017 | 9.84E-46 | 1 L1 _ | AI467606 |
| Gsap | 4.16E-50 | 0.298676 | 0.221 | 0.027 | 8.38E-46 | 1 L1 _ | Gsap |
| P2ry10 | 1.69E-50 | 0.281121 | 0.149 | 0.007 | 3.39E-46 | 1 L1 _ | P2ry10 |
| D830025C05Rik | 2.44E-51 | 0.475755 | 0.317 | 0.065 | 4.92E-47 | 1 L1 _ | D830025C05Rik |
| Glcci1 | 2.05E-51 | 0.478406 | 0.274 | 0.046 | 4.13E-47 | 1 L1 _ | Glcci1 |
| Cd72 | 1.55E-51 | 0.778398 | 0.278 | 0.05 | 3.11E-47 | 1 L1 _ | Cd72 |
| N4bp2l1 | 7.25E-52 | 0.455809 | 0.257 | 0.039 | 1.46E-47 | 1 L1 _ | N4bp2l1 |
| Il10ra1 | 6.85E-52 | 0.287451 | 0.168 | 0.011 | 1.38E-47 | 1 L1 _ | Il10ra |
| Eif3f | 5.08E-52 | 0.762122 | 0.926 | 0.8 | 1.02E-47 | 1 L1 _ | Eif3f |
| Grcc10 | 1.19E-52 | 0.860623 | 0.869 | 0.659 | 2.40E-48 | 1 L1 _ | Grcc10 |
| Arhgap251 | 5.85E-53 | 0.345009 | 0.232 | 0.029 | 1.18E-48 | 1 L1 _ | Arhgap25 |
| Rpl7a | 5.13E-53 | 0.622656 | 0.938 | 0.914 | 1.03E-48 | 1 L1 _ | Rpl7a |
| 2810013P06Rik | 3.29E-53 | 0.654104 | 0.503 | 0.168 | 6.62E-49 | 1 L1 _ | 2810013P06Rik |

| | | | | | | | |
|---|---|---|---|---|---|---|---|
| Sh3bp2 | 2.71E-53 | 0.352341 | 0.182 | 0.013 | 5.45E-49 | 1 L1 _ | Sh3bp2 |
| Jaml1 | 6.77E-54 | 0.647149 | 0.338 | 0.074 | 1.36E-49 | 1 L1 _ | Jaml |
| Inpp4b1 | 6.02E-54 | 0.635196 | 0.614 | 0.213 | 1.21E-49 | 1 L1 _ | Inpp4b |
| Vipr21 | 3.98E-54 | 0.382734 | 0.179 | 0.012 | 8.01E-50 | 1 L1 _ | Vipr2 |
| Lamb3 | 2.16E-54 | 0.259682 | 0.168 | 0.009 | 4.35E-50 | 1 L1 _ | Lamb3 |
| Dennd2d | 8.71E-55 | 0.252404 | 0.145 | 0.004 | 1.75E-50 | 1 L1 _ | Dennd2d |
| Srgap3 | 5.45E-55 | 0.417821 | 0.274 | 0.043 | 1.10E-50 | 1 L1 _ | Srgap3 |
| Zcchc18 | 3.57E-55 | 0.258332 | 0.168 | 0.008 | 7.19E-51 | 1 L1 _ | Zcchc18 |
| Tnfaip8l2 | 1.22E-55 | 0.421506 | 0.216 | 0.022 | 2.45E-51 | 1 L1 _ | Tnfaip8l2 |
| Gramd1a | 5.71E-56 | 0.782263 | 0.513 | 0.181 | 1.15E-51 | 1 L1 _ | Gramd1a |
| Pde3b | 5.80E-57 | 0.438607 | 0.269 | 0.039 | 1.17E-52 | 1 L1 _ | Pde3b |
| Fam189b | 3.10E-57 | 0.701217 | 0.418 | 0.112 | 6.25E-53 | 1 L1 _ | Fam189b |
| Il16 | 2.75E-57 | 0.485865 | 0.336 | 0.066 | 5.54E-53 | 1 L1 _ | Il16 |
| 5830428M24Rik | 2.66E-57 | 0.326514 | 0.166 | 0.007 | 5.37E-53 | 1 L1 _ | 5830428M24Rik |
| Junb | 2.37E-57 | 1.003709 | 0.897 | 0.61 | 4.78E-53 | 1 L1 _ | Junb |
| Ifng | 2.33E-57 | 0.418873 | 0.166 | 0.007 | 4.68E-53 | 1 L1 _ | Ifng |
| Gm36738 | 6.55E-58 | 0.410944 | 0.287 | 0.044 | 1.32E-53 | 1 L1 _ | Gm36738 |
| Ppp3cc | 6.31E-58 | 0.531194 | 0.315 | 0.057 | 1.27E-53 | 1 L1 _ | Ppp3cc |
| Plcxd2 | 3.53E-58 | 0.342422 | 0.184 | 0.011 | 7.10E-54 | 1 L1 _ | Plcxd2 |
| Gimap81 | 2.53E-58 | 0.514529 | 0.366 | 0.076 | 5.09E-54 | 1 L1 _ | Gimap8 |
| Asb2 | 7.85E-59 | 0.355731 | 0.232 | 0.023 | 1.58E-54 | 1 L1 _ | Asb2 |
| H2-D1 | 6.02E-59 | 0.650883 | 0.982 | 0.948 | 1.21E-54 | 1 L1 _ | H2-D1 |
| Ankrd44 | 3.93E-59 | 0.824143 | 0.641 | 0.266 | 7.92E-55 | 1 L1 _ | Ankrd44 |
| Dock10 | 2.44E-59 | 0.890773 | 0.715 | 0.348 | 4.91E-55 | 1 L1 _ | Dock10 |
| S100a101 | 2.37E-59 | 0.996973 | 0.86 | 0.654 | 4.77E-55 | 1 L1 _ | S100a10 |
| Itgad | 8.50E-60 | 0.305811 | 0.156 | 0.004 | 1.71E-55 | 1 L1 _ | Itgad |
| Rpl24 | 3.87E-60 | 0.594283 | 0.954 | 0.974 | 7.80E-56 | 1 L1 _ | Rpl24 |
| Lyrm9 | 3.55E-60 | 0.345415 | 0.205 | 0.015 | 7.16E-56 | 1 L1 _ | Lyrm9 |
| Bcl11b | 2.83E-60 | 0.502456 | 0.186 | 0.01 | 5.70E-56 | 1 L1 _ | Bcl11b |
| Faah | 7.01E-61 | 0.426038 | 0.209 | 0.016 | 1.41E-56 | 1 L1 _ | Faah |
| Cd247 | 4.97E-61 | 0.307474 | 0.184 | 0.009 | 1.00E-56 | 1 L1 _ | Cd247 |
| Actr3 | 1.25E-61 | 0.975029 | 0.83 | 0.54 | 2.53E-57 | 1 L1 _ | Actr3 |
| Rps6 | 9.37E-62 | 0.728568 | 0.959 | 0.888 | 1.89E-57 | 1 L1 _ | Rps6 |
| Cish | 7.25E-62 | 0.563692 | 0.31 | 0.049 | 1.46E-57 | 1 L1 _ | Cish |
| Mbnl1 | 5.81E-62 | 0.952815 | 0.897 | 0.791 | 1.17E-57 | 1 L1 _ | Mbnl1 |
| Jakmip11 | 4.75E-62 | 0.309285 | 0.182 | 0.008 | 9.56E-58 | 1 L1 _ | Jakmip1 |
| Ptpre1 | 3.52E-62 | 0.65627 | 0.462 | 0.124 | 7.09E-58 | 1 L1 _ | Ptpre |
| Mob3a | 3.41E-62 | 0.70781 | 0.423 | 0.104 | 6.86E-58 | 1 L1 _ | Mob3a |
| Mbp | 1.24E-62 | 0.532592 | 0.241 | 0.024 | 2.50E-58 | 1 L1 _ | Mbp |
| Ugcg | 8.03E-63 | 1.053049 | 0.621 | 0.267 | 1.62E-58 | 1 L1 _ | Ugcg |
| Gm43065 | 7.56E-63 | 0.278298 | 0.149 | 0.001 | 1.52E-58 | 1 L1 _ | Gm43065 |

| | | | | | | | |
|---|---|---|---|---|---|---|---|
| Map4k1 | 4.24E-63 | 0.292902 | 0.182 | 0.007 | 8.55E-59 | 1 L1 _ | Map4k1 |
| Psme11 | 3.38E-63 | 1.000318 | 0.906 | 0.69 | 6.81E-59 | 1 L1 _ | Psme1 |
| Gata3 | 2.83E-63 | 0.38123 | 0.2 | 0.012 | 5.71E-59 | 1 L1 _ | Gata3 |
| Pag1 | 9.96E-64 | 0.537041 | 0.352 | 0.063 | 2.00E-59 | 1 L1 _ | Pag1 |
| Susd3 | 5.25E-64 | 0.340283 | 0.182 | 0.007 | 1.06E-59 | 1 L1 _ | Susd3 |
| Tnf | 5.07E-64 | 0.393463 | 0.184 | 0.007 | 1.02E-59 | 1 L1 _ | Tnf |
| Tmsb15b2 | 1.07E-64 | 0.26282 | 0.189 | 0.008 | 2.15E-60 | 1 L1 _ | Tmsb15b2 |
| Rbm38 | 7.80E-65 | 0.338533 | 0.241 | 0.022 | 1.57E-60 | 1 L1 _ | Rbm38 |
| Cd82 | 7.18E-65 | 0.80261 | 0.494 | 0.142 | 1.45E-60 | 1 L1 _ | Cd82 |
| Rpl28 | 3.57E-65 | 0.633461 | 0.977 | 0.973 | 7.19E-61 | 1 L1 _ | Rpl28 |
| Klrb1b1 | 1.94E-65 | 0.303733 | 0.191 | 0.008 | 3.90E-61 | 1 L1 _ | Klrb1b |
| Gbp81 | 9.20E-66 | 0.398386 | 0.26 | 0.027 | 1.85E-61 | 1 L1 _ | Gbp8 |
| Rps8 | 6.17E-66 | 0.57867 | 0.995 | 0.992 | 1.24E-61 | 1 L1 _ | Rps8 |
| AC149090.11 | 5.44E-66 | 0.985536 | 0.793 | 0.401 | 1.10E-61 | 1 L1 _ | AC149090.1 |
| Rpl5 | 4.09E-66 | 0.830887 | 0.945 | 0.892 | 8.24E-62 | 1 L1 _ | Rpl5 |
| Klrc1 | 3.32E-66 | 0.546336 | 0.189 | 0.007 | 6.68E-62 | 1 L1 _ | Klrc1 |
| Parvg1 | 7.22E-67 | 0.297172 | 0.191 | 0.007 | 1.45E-62 | 1 L1 _ | Parvg |
| Ctss | 7.02E-67 | 2.256347 | 0.46 | 0.121 | 1.41E-62 | 1 L1 _ | Ctss |
| Stk26 | 6.66E-67 | 0.350628 | 0.2 | 0.009 | 1.34E-62 | 1 L1 _ | Stk26 |
| Cd160 | 5.23E-67 | 0.552874 | 0.214 | 0.013 | 1.05E-62 | 1 L1 _ | Cd160 |
| H2-Q6 | 8.99E-68 | 0.826896 | 0.625 | 0.217 | 1.81E-63 | 1 L1 _ | H2-Q6 |
| Efhd21 | 5.63E-68 | 0.849316 | 0.66 | 0.252 | 1.13E-63 | 1 L1 _ | Efhd2 |
| Fmnl11 | 4.88E-68 | 0.492543 | 0.315 | 0.045 | 9.83E-64 | 1 L1 _ | Fmnl1 |
| Mir142hg | 3.97E-68 | 0.394296 | 0.228 | 0.016 | 8.00E-64 | 1 L1 _ | Mir142hg |
| Gm11808 | 2.80E-68 | 0.896527 | 0.722 | 0.356 | 5.64E-64 | 1 L1 _ | Gm11808 |
| Cyba | 2.65E-68 | 1.036445 | 0.92 | 0.806 | 5.34E-64 | 1 L1 _ | Cyba |
| Arhgap151 | 2.37E-68 | 0.518206 | 0.407 | 0.079 | 4.77E-64 | 1 L1 _ | Arhgap15 |
| Tspan32 | 5.87E-69 | 0.433083 | 0.218 | 0.013 | 1.18E-64 | 1 L1 _ | Tspan32 |
| Sytl3 | 3.04E-69 | 0.35732 | 0.228 | 0.015 | 6.12E-65 | 1 L1 _ | Sytl3 |
| Klra1 | 2.03E-69 | 0.867978 | 0.223 | 0.014 | 4.09E-65 | 1 L1 _ | Klra1 |
| Rab37 | 1.04E-69 | 0.38982 | 0.274 | 0.027 | 2.09E-65 | 1 L1 _ | Rab37 |
| Tmsb10 | 8.26E-70 | 1.030199 | 0.97 | 0.913 | 1.66E-65 | 1 L1 _ | Tmsb10 |
| Ezr | 4.05E-71 | 0.646909 | 0.411 | 0.083 | 8.16E-67 | 1 L1 _ | Ezr |
| Otulinl | 3.86E-71 | 0.572921 | 0.331 | 0.047 | 7.77E-67 | 1 L1 _ | Otulinl |
| Nckap1l | 1.67E-71 | 0.422088 | 0.239 | 0.017 | 3.36E-67 | 1 L1 _ | Nckap1l |
| Stxbp2 | 1.64E-71 | 0.388729 | 0.232 | 0.015 | 3.30E-67 | 1 L1 _ | Stxbp2 |
| Pde7a | 1.42E-71 | 0.661004 | 0.428 | 0.092 | 2.86E-67 | 1 L1 _ | Pde7a |
| Dnajc15 | 1.37E-71 | 1.034116 | 0.66 | 0.275 | 2.76E-67 | 1 L1 _ | Dnajc15 |
| Gng21 | 1.20E-71 | 0.864798 | 0.57 | 0.185 | 2.41E-67 | 1 L1 _ | Gng2 |
| Rnf43 | 9.35E-72 | 0.344504 | 0.184 | 0.004 | 1.88E-67 | 1 L1 _ | Rnf43 |
| Kif21b1 | 3.98E-72 | 0.399724 | 0.264 | 0.023 | 8.01E-68 | 1 L1 _ | Kif21b |

| | | | | | | | |
|---|---|---|---|---|---|---|---|
| Sh2d1a | 3.83E-72 | 0.388704 | 0.198 | 0.006 | 7.70E-68 | 1 L1 _ | Sh2d1a |
| Dennd4a | 2.41E-72 | 1.042252 | 0.607 | 0.224 | 4.85E-68 | 1 L1 _ | Dennd4a |
| Hist1h1d1 | 8.02E-73 | 0.519019 | 0.239 | 0.016 | 1.61E-68 | 1 L1 _ | Hist1h1d |
| Clec2i1 | 4.78E-73 | 0.380935 | 0.207 | 0.008 | 9.64E-69 | 1 L1 _ | Clec2i |
| Hmgb21 | 2.14E-73 | 1.142611 | 0.717 | 0.319 | 4.30E-69 | 1 L1 _ | Hmgb2 |
| Rpl10 | 6.42E-74 | 0.801636 | 0.968 | 0.937 | 1.29E-69 | 1 L1 _ | Rpl10 |
| Cd27 | 4.54E-74 | 0.54288 | 0.32 | 0.04 | 9.14E-70 | 1 L1 _ | Cd27 |
| Eef1b2 | 2.96E-74 | 0.929088 | 0.933 | 0.857 | 5.96E-70 | 1 L1 _ | Eef1b2 |
| Sec11c | 2.08E-74 | 1.043566 | 0.733 | 0.356 | 4.19E-70 | 1 L1 _ | Sec11c |
| Bcl2a1b | 1.03E-74 | 0.811022 | 0.271 | 0.024 | 2.08E-70 | 1 L1 _ | Bcl2a1b |
| Fxyd5 | 5.69E-75 | 0.998308 | 0.917 | 0.71 | 1.15E-70 | 1 L1 _ | Fxyd5 |
| Chd7 | 5.63E-75 | 0.523647 | 0.363 | 0.055 | 1.13E-70 | 1 L1 _ | Chd7 |
| Mctp2 | 2.39E-75 | 0.392823 | 0.202 | 0.006 | 4.81E-71 | 1 L1 _ | Mctp2 |
| Gprin3 | 1.99E-75 | 0.484339 | 0.244 | 0.016 | 4.00E-71 | 1 L1 _ | Gprin3 |
| Slc16a6 | 1.36E-75 | 0.36094 | 0.237 | 0.014 | 2.74E-71 | 1 L1 _ | Slc16a6 |
| Gpr1741 | 7.95E-76 | 0.396446 | 0.218 | 0.009 | 1.60E-71 | 1 L1 _ | Gpr174 |
| Rpl36al | 6.99E-76 | 0.870616 | 0.929 | 0.853 | 1.41E-71 | 1 L1 _ | Rpl36al |
| Pik3ap1 | 2.30E-76 | 0.431767 | 0.234 | 0.013 | 4.64E-72 | 1 L1 _ | Pik3ap1 |
| Card11 | 1.79E-76 | 0.436242 | 0.232 | 0.012 | 3.61E-72 | 1 L1 _ | Card11 |
| Was | 2.20E-77 | 0.449767 | 0.308 | 0.032 | 4.44E-73 | 1 L1 _ | Was |
| I730030J21Rik | 1.33E-77 | 0.371773 | 0.186 | 0.002 | 2.68E-73 | 1 L1 _ | I730030J21Rik |
| Rpl7 | 4.49E-78 | 0.712974 | 0.972 | 0.951 | 9.05E-74 | 1 L1 _ | Rpl7 |
| Gm2682 | 1.34E-78 | 0.523926 | 0.239 | 0.013 | 2.69E-74 | 1 L1 _ | Gm2682 |
| Pstpip1 | 1.26E-78 | 0.647007 | 0.395 | 0.067 | 2.54E-74 | 1 L1 _ | Pstpip1 |
| Tox | 7.18E-79 | 0.693162 | 0.333 | 0.043 | 1.45E-74 | 1 L1 _ | Tox |
| Arl4c | 2.62E-79 | 0.710371 | 0.359 | 0.052 | 5.28E-75 | 1 L1 _ | Arl4c |
| Rps2 | 2.21E-79 | 0.755711 | 0.977 | 0.973 | 4.46E-75 | 1 L1 _ | Rps2 |
| Actb2 | 1.79E-79 | 0.773124 | 0.995 | 0.992 | 3.61E-75 | 1 L1 _ | Actb |
| Aim21 | 1.20E-79 | 0.49674 | 0.292 | 0.026 | 2.43E-75 | 1 L1 _ | Aim2 |
| Spint2 | 8.91E-80 | 0.401556 | 0.253 | 0.015 | 1.79E-75 | 1 L1 _ | Spint2 |
| Snx20 | 6.39E-80 | 0.447807 | 0.262 | 0.018 | 1.29E-75 | 1 L1 _ | Snx20 |
| Gpr34 | 2.50E-80 | 0.412324 | 0.225 | 0.008 | 5.03E-76 | 1 L1 _ | Gpr34 |
| Pfn12 | 1.88E-80 | 1.134612 | 0.885 | 0.774 | 3.79E-76 | 1 L1 _ | Pfn1 |
| Itgb72 | 1.57E-80 | 0.691973 | 0.315 | 0.034 | 3.17E-76 | 1 L1 _ | Itgb7 |
| Osbpl3 | 8.31E-81 | 0.701379 | 0.418 | 0.071 | 1.67E-76 | 1 L1 _ | Osbpl3 |
| Vav11 | 4.84E-81 | 0.539553 | 0.274 | 0.02 | 9.74E-77 | 1 L1 _ | Vav1 |
| Tle5 | 3.33E-81 | 1.147379 | 0.883 | 0.599 | 6.70E-77 | 1 L1 _ | Tle5 |
| Gimap61 | 2.37E-81 | 1.150011 | 0.768 | 0.324 | 4.77E-77 | 1 L1 _ | Gimap6 |
| Nabp1 | 2.28E-81 | 0.621441 | 0.31 | 0.032 | 4.60E-77 | 1 L1 _ | Nabp1 |
| Inpp5d | 7.78E-82 | 0.596807 | 0.379 | 0.055 | 1.57E-77 | 1 L1 _ | Inpp5d |
| Rpl14 | 6.96E-82 | 0.792566 | 0.952 | 0.938 | 1.40E-77 | 1 L1 _ | Rpl14 |

| | | | | | | | |
|---|---|---|---|---|---|---|---|
| Pik3cd | 6.89E-82 | 0.671437 | 0.402 | 0.064 | 1.39E-77 | 1 L1 _ | Pik3cd |
| Akna1 | 1.83E-82 | 0.688388 | 0.395 | 0.062 | 3.68E-78 | 1 L1 _ | Akna |
| Fam78a | 5.24E-83 | 0.401497 | 0.234 | 0.009 | 1.06E-78 | 1 L1 _ | Fam78a |
| Lilrb4a | 2.25E-83 | 0.561359 | 0.244 | 0.011 | 4.53E-79 | 1 L1 _ | Lilrb4a |
| Lpxn1 | 1.58E-83 | 0.607305 | 0.285 | 0.022 | 3.17E-79 | 1 L1 _ | Lpxn |
| Rasgrp1 | 1.67E-84 | 0.470284 | 0.255 | 0.013 | 3.37E-80 | 1 L1 _ | Rasgrp1 |
| Rpl15 | 5.05E-85 | 0.749555 | 0.975 | 0.964 | 1.02E-80 | 1 L1 _ | Rpl15 |
| Eomes | 4.22E-85 | 0.540925 | 0.234 | 0.008 | 8.50E-81 | 1 L1 _ | Eomes |
| Arhgap41 | 3.88E-85 | 0.435737 | 0.257 | 0.013 | 7.82E-81 | 1 L1 _ | Arhgap4 |
| Rpl23a | 2.50E-85 | 0.877161 | 0.961 | 0.916 | 5.03E-81 | 1 L1 _ | Rpl23a |
| Clic1 | 2.01E-85 | 1.067521 | 0.929 | 0.783 | 4.04E-81 | 1 L1 _ | Clic1 |
| Tmem273 | 1.01E-85 | 0.580018 | 0.382 | 0.048 | 2.04E-81 | 1 L1 _ | Tmem273 |
| Lrmp | 6.07E-86 | 0.379331 | 0.262 | 0.014 | 1.22E-81 | 1 L1 _ | Lrmp |
| Rps19 | 4.15E-86 | 0.818431 | 0.975 | 0.965 | 8.35E-82 | 1 L1 _ | Rps19 |
| Uba52 | 3.04E-86 | 1.198087 | 0.855 | 0.553 | 6.12E-82 | 1 L1 _ | Uba52 |
| Jpt1 | 2.45E-86 | 1.127072 | 0.722 | 0.305 | 4.93E-82 | 1 L1 _ | Jpt1 |
| Cd3d | 2.31E-86 | 1.521391 | 0.287 | 0.022 | 4.65E-82 | 1 L1 _ | Cd3d |
| Rps25 | 2.18E-86 | 0.80598 | 0.966 | 0.937 | 4.38E-82 | 1 L1 _ | Rps25 |
| Rack1 | 9.67E-87 | 0.949364 | 0.972 | 0.918 | 1.95E-82 | 1 L1 _ | Rack1 |
| Lax11 | 5.90E-87 | 0.482992 | 0.244 | 0.009 | 1.19E-82 | 1 L1 _ | Lax1 |
| Nlrc3 | 4.05E-87 | 0.561602 | 0.276 | 0.017 | 8.15E-83 | 1 L1 _ | Nlrc3 |
| Rassf51 | 2.97E-87 | 0.588605 | 0.375 | 0.049 | 5.97E-83 | 1 L1 _ | Rassf5 |
| Psmb81 | 1.07E-87 | 1.129752 | 0.906 | 0.571 | 2.15E-83 | 1 L1 _ | Psmb8 |
| Tm6sf11 | 8.85E-88 | 0.623853 | 0.402 | 0.058 | 1.78E-83 | 1 L1 _ | Tm6sf1 |
| Samsn11 | 8.36E-88 | 0.554364 | 0.297 | 0.023 | 1.68E-83 | 1 L1 _ | Samsn1 |
| Napsa1 | 4.94E-88 | 0.440917 | 0.246 | 0.009 | 9.94E-84 | 1 L1 _ | Napsa |
| Dok2 | 2.22E-88 | 0.495509 | 0.253 | 0.011 | 4.46E-84 | 1 L1 _ | Dok2 |
| Cdh12 | 1.50E-88 | 0.488673 | 0.276 | 0.016 | 3.02E-84 | 1 L1 _ | Cdh1 |
| Ubash3a | 2.32E-89 | 0.406281 | 0.237 | 0.006 | 4.67E-85 | 1 L1 _ | Ubash3a |
| Fcgr3 | 1.52E-89 | 0.597989 | 0.294 | 0.021 | 3.05E-85 | 1 L1 _ | Fcgr3 |
| Sla2 | 8.47E-90 | 0.461468 | 0.26 | 0.011 | 1.71E-85 | 1 L1 _ | Sla2 |
| Gm19585 | 8.38E-90 | 0.55063 | 0.239 | 0.007 | 1.69E-85 | 1 L1 _ | Gm19585 |
| Nup2101 | 3.55E-90 | 0.50965 | 0.269 | 0.013 | 7.15E-86 | 1 L1 _ | Nup210 |
| Cd3e | 2.34E-90 | 1.222298 | 0.292 | 0.02 | 4.71E-86 | 1 L1 _ | Cd3e |
| Arrb2 | 1.92E-90 | 0.734469 | 0.448 | 0.076 | 3.86E-86 | 1 L1 _ | Arrb2 |
| Gpr68 | 7.51E-91 | 0.584994 | 0.315 | 0.026 | 1.51E-86 | 1 L1 _ | Gpr68 |
| Plek | 3.31E-91 | 0.739433 | 0.363 | 0.042 | 6.66E-87 | 1 L1 _ | Plek |
| Psd41 | 1.02E-91 | 0.485959 | 0.301 | 0.021 | 2.05E-87 | 1 L1 _ | Psd4 |
| Rplp0 | 4.75E-92 | 0.900855 | 0.989 | 0.955 | 9.56E-88 | 1 L1 _ | Rplp0 |
| AU0202062 | 4.26E-92 | 1.154987 | 0.582 | 0.146 | 8.57E-88 | 1 L1 _ | AU020206 |
| Apol7e | 1.64E-92 | 0.461551 | 0.244 | 0.006 | 3.31E-88 | 1 L1 _ | Apol7e |

| | | | | | | | |
|---|---|---|---|---|---|---|---|
| Sh2d1b11 | 3.76E-93 | 0.460968 | 0.29 | 0.017 | 7.57E-89 | 1 L1 _ | Sh2d1b1 |
| Tbx211 | 2.54E-93 | 0.457009 | 0.264 | 0.011 | 5.11E-89 | 1 L1 _ | Tbx21 |
| Rpl29 | 4.60E-94 | 0.935049 | 0.959 | 0.932 | 9.26E-90 | 1 L1 _ | Rpl29 |
| Gimap1os | 1.81E-94 | 0.453393 | 0.232 | 0.003 | 3.64E-90 | 1 L1 _ | Gimap1os |
| Tnni1 | 1.43E-94 | 0.689887 | 0.264 | 0.01 | 2.87E-90 | 1 L1 _ | Tnni1 |
| Rpl26 | 4.82E-95 | 0.75346 | 0.984 | 0.961 | 9.72E-91 | 1 L1 _ | Rpl26 |
| Klra31 | 3.16E-95 | 1.091387 | 0.317 | 0.024 | 6.37E-91 | 1 L1 _ | Klra3 |
| Prkcq | 5.88E-96 | 0.505675 | 0.267 | 0.009 | 1.19E-91 | 1 L1 _ | Prkcq |
| Matk | 4.87E-96 | 0.582284 | 0.287 | 0.015 | 9.81E-92 | 1 L1 _ | Matk |
| Klk8 | 2.25E-96 | 0.757194 | 0.37 | 0.039 | 4.52E-92 | 1 L1 _ | Klk8 |
| Rps281 | 3.52E-97 | 0.968115 | 0.966 | 0.962 | 7.09E-93 | 1 L1 _ | Rps28 |
| Gm14029 | 9.05E-98 | 0.595031 | 0.255 | 0.006 | 1.82E-93 | 1 L1 _ | Gm14029 |
| Rpl22 | 4.04E-98 | 0.908667 | 0.954 | 0.931 | 8.13E-94 | 1 L1 _ | Rpl22 |
| Rpl41 | 1.58E-98 | 0.841035 | 0.986 | 0.989 | 3.17E-94 | 1 L1 _ | Rpl41 |
| Ccdc88c | 1.52E-98 | 0.673579 | 0.423 | 0.054 | 3.07E-94 | 1 L1 _ | Ccdc88c |
| Atp8b4 | 1.10E-98 | 0.564061 | 0.285 | 0.013 | 2.22E-94 | 1 L1 _ | Atp8b4 |
| Bcl2 | 7.76E-99 | 1.621738 | 0.747 | 0.322 | 1.56E-94 | 1 L1 _ | Bcl2 |
| Tmem1631 | 3.68E-99 | 0.427595 | 0.257 | 0.006 | 7.42E-95 | 1 L1 _ | Tmem163 |
| Ifitm10 | 3.12E-99 | 0.722134 | 0.4 | 0.049 | 6.28E-95 | 1 L1 _ | Ifitm10 |
| H2-K1 | 2.52E-99 | 1.06585 | 0.986 | 0.923 | 5.08E-95 | 1 L1 _ | H2-K1 |
| Cotl1 | 1.73E-99 | 1.378443 | 0.795 | 0.37 | 3.48E-95 | 1 L1 _ | Cotl1 |
| Rps15 | 1.51E-99 | 0.958989 | 0.975 | 0.946 | 3.03E-95 | 1 L1 _ | Rps15 |
| Klrb1a | 1.21E-99 | 0.693617 | 0.329 | 0.024 | 2.45E-95 | 1 L1 _ | Klrb1a |
| Grap2 | 1.11E-99 | 0.691133 | 0.264 | 0.007 | 2.24E-95 | 1 L1 _ | Grap2 |
| Chn2 | 1.01E-99 | 0.760308 | 0.409 | 0.05 | 2.03E-95 | 1 L1 _ | Chn2 |
| Ctla2b | 5.17E-100 | 0.713366 | 0.366 | 0.035 | 1.04E-95 | 1 L1 _ | Ctla2b |
| Rps4x | 2.55E-100 | 0.871975 | 0.979 | 0.973 | 5.14E-96 | 1 L1 _ | Rps4x |
| Rpl21 | 1.45E-100 | 0.840749 | 0.986 | 0.978 | 2.92E-96 | 1 L1 _ | Rpl21 |
| Arhgdib1 | 2.20E-101 | 1.322253 | 0.901 | 0.518 | 4.43E-97 | 1 L1 _ | Arhgdib |
| Glipr11 | 1.39E-101 | 0.595071 | 0.324 | 0.021 | 2.81E-97 | 1 L1 _ | Glipr1 |
| Ctla2a1 | 1.06E-101 | 1.182877 | 0.795 | 0.257 | 2.13E-97 | 1 L1 _ | Ctla2a |
| Myo1g | 5.19E-102 | 0.500602 | 0.278 | 0.009 | 1.05E-97 | 1 L1 _ | Myo1g |
| Ifi209 | 8.86E-104 | 0.635237 | 0.308 | 0.016 | 1.78E-99 | 1 L1 _ | Ifi209 |
| S100a4 | 1.31E-105 | 2.094793 | 0.779 | 0.358 | 2.64E-101 | 1 L1 _ | S100a4 |
| Sema4a | 1.20E-105 | 0.635547 | 0.361 | 0.028 | 2.42E-101 | 1 L1 _ | Sema4a |
| Itk | 2.98E-106 | 0.5756 | 0.292 | 0.01 | 6.00E-102 | 1 L1 _ | Itk |
| Malt1 | 1.95E-106 | 1.049123 | 0.561 | 0.114 | 3.93E-102 | 1 L1 _ | Malt1 |
| Sla1 | 1.80E-106 | 0.585624 | 0.333 | 0.021 | 3.63E-102 | 1 L1 _ | Sla |
| H2-Q7 | 1.15E-106 | 1.526932 | 0.869 | 0.439 | 2.31E-102 | 1 L1 _ | H2-Q7 |
| Rhof1 | 6.51E-107 | 0.597178 | 0.31 | 0.014 | 1.31E-102 | 1 L1 _ | Rhof |
| Cd84 | 1.36E-108 | 0.634391 | 0.303 | 0.012 | 2.75E-104 | 1 L1 _ | Cd84 |

| | | | | | | | |
|---|---|---|---|---|---|---|---|
| Rps3a1 | 9.69E-109 | 1.054344 | 0.975 | 0.963 | 1.95E-104 | 1 L1 _ | Rps3a1 |
| Rps26 | 5.34E-109 | 1.085731 | 0.972 | 0.955 | 1.08E-104 | 1 L1 _ | Rps26 |
| Rps21 | 2.23E-110 | 0.996087 | 0.982 | 0.981 | 4.48E-106 | 1 L1 _ | Rps21 |
| Rps23 | 2.19E-110 | 0.958525 | 0.966 | 0.969 | 4.41E-106 | 1 L1 _ | Rps23 |
| Rpl8 | 1.15E-110 | 0.99764 | 0.982 | 0.967 | 2.31E-106 | 1 L1 _ | Rpl8 |
| Fasl | 5.24E-111 | 0.535374 | 0.29 | 0.007 | 1.05E-106 | 1 L1 _ | Fasl |
| Cited41 | 1.78E-111 | 0.806745 | 0.379 | 0.031 | 3.59E-107 | 1 L1 _ | Cited4 |
| AI662270 | 9.37E-112 | 0.598945 | 0.317 | 0.014 | 1.89E-107 | 1 L1 _ | AI662270 |
| Rps27 | 9.20E-112 | 1.100557 | 0.986 | 0.975 | 1.85E-107 | 1 L1 _ | Rps27 |
| Rpl9 | 1.95E-112 | 0.977021 | 0.982 | 0.965 | 3.92E-108 | 1 L1 _ | Rpl9 |
| Rplp1 | 1.33E-112 | 0.878248 | 0.995 | 0.983 | 2.69E-108 | 1 L1 _ | Rplp1 |
| Lcp2 | 4.24E-113 | 0.55858 | 0.34 | 0.018 | 8.54E-109 | 1 L1 _ | Lcp2 |
| Rpl6 | 2.54E-113 | 0.945694 | 0.982 | 0.983 | 5.12E-109 | 1 L1 _ | Rpl6 |
| Gmfg | 5.94E-115 | 0.880187 | 0.411 | 0.039 | 1.20E-110 | 1 L1 _ | Gmfg |
| Btg2 | 3.05E-115 | 1.758217 | 0.802 | 0.318 | 6.15E-111 | 1 L1 _ | Btg2 |
| Rps12 | 1.76E-115 | 1.039063 | 0.986 | 0.971 | 3.54E-111 | 1 L1 _ | Rps12 |
| Rpl35a | 1.25E-115 | 0.877438 | 0.984 | 0.983 | 2.52E-111 | 1 L1 _ | Rpl35a |
| Spn1 | 1.24E-115 | 0.603167 | 0.32 | 0.012 | 2.49E-111 | 1 L1 _ | Spn |
| Rpl17 | 1.14E-115 | 0.997193 | 0.982 | 0.954 | 2.30E-111 | 1 L1 _ | Rpl17 |
| Cyfip21 | 4.70E-116 | 0.683286 | 0.37 | 0.025 | 9.46E-112 | 1 L1 _ | Cyfip2 |
| Ptpn7 | 4.06E-116 | 0.592498 | 0.326 | 0.013 | 8.18E-112 | 1 L1 _ | Ptpn7 |
| Rps291 | 1.80E-116 | 0.908731 | 0.986 | 0.986 | 3.63E-112 | 1 L1 _ | Rps29 |
| Clnk1 | 6.53E-118 | 0.660507 | 0.322 | 0.011 | 1.31E-113 | 1 L1 _ | Clnk |
| Evl1 | 4.87E-118 | 1.230926 | 0.713 | 0.207 | 9.81E-114 | 1 L1 _ | Evl |
| Tnfrsf9 | 4.58E-118 | 0.782853 | 0.343 | 0.016 | 9.22E-114 | 1 L1 _ | Tnfrsf9 |
| Gpr651 | 1.66E-118 | 0.759299 | 0.356 | 0.019 | 3.35E-114 | 1 L1 _ | Gpr65 |
| Rpl13 | 1.43E-118 | 1.000095 | 0.995 | 0.988 | 2.87E-114 | 1 L1 _ | Rpl13 |
| Igflr1 | 3.06E-119 | 0.770272 | 0.379 | 0.026 | 6.16E-115 | 1 L1 _ | Igflr1 |
| Gm36723 | 1.67E-119 | 0.79274 | 0.31 | 0.008 | 3.35E-115 | 1 L1 _ | Gm36723 |
| Eef1a1 | 9.74E-120 | 0.931909 | 0.998 | 0.987 | 1.96E-115 | 1 L1 _ | Eef1a1 |
| Shisa5 | 9.09E-120 | 1.627566 | 0.931 | 0.735 | 1.83E-115 | 1 L1 _ | Shisa5 |
| Gimap51 | 4.84E-120 | 1.120067 | 0.618 | 0.124 | 9.76E-116 | 1 L1 _ | Gimap5 |
| Fam49b | 2.45E-120 | 1.060181 | 0.602 | 0.121 | 4.93E-116 | 1 L1 _ | Fam49b |
| Rpl3 | 1.88E-120 | 1.104788 | 0.979 | 0.947 | 3.79E-116 | 1 L1 _ | Rpl3 |
| Coro2a | 1.11E-120 | 0.683477 | 0.363 | 0.02 | 2.23E-116 | 1 L1 _ | Coro2a |
| Pycard | 4.39E-121 | 1.187209 | 0.667 | 0.172 | 8.83E-117 | 1 L1 _ | Pycard |
| Pkp31 | 2.76E-121 | 0.704694 | 0.379 | 0.024 | 5.55E-117 | 1 L1 _ | Pkp3 |
| Itga4 | 6.88E-122 | 0.870217 | 0.361 | 0.018 | 1.39E-117 | 1 L1 _ | Itga4 |
| Rps6ka1 | 6.60E-123 | 0.825945 | 0.439 | 0.042 | 1.33E-118 | 1 L1 _ | Rps6ka1 |
| Rpl39 | 5.64E-123 | 1.123194 | 0.984 | 0.971 | 1.14E-118 | 1 L1 _ | Rpl39 |
| Rpl37a1 | 5.59E-123 | 1.073466 | 0.989 | 0.984 | 1.13E-118 | 1 L1 _ | Rpl37a |

| | | | | | | | |
|---|---|---|---|---|---|---|---|
| Stk17b1 | 5.39E-123 | 1.422879 | 0.766 | 0.249 | 1.09E-118 | 1 L1 _ | Stk17b |
| Gpr171 | 4.98E-123 | 0.622265 | 0.324 | 0.009 | 1.00E-118 | 1 L1 _ | Gpr171 |
| Cxcr3 | 4.76E-123 | 0.819494 | 0.356 | 0.017 | 9.59E-119 | 1 L1 _ | Cxcr3 |
| Rpl37 | 4.55E-123 | 1.14044 | 0.982 | 0.977 | 9.16E-119 | 1 L1 _ | Rpl37 |
| Rpl10a | 3.13E-123 | 1.148375 | 0.979 | 0.931 | 6.30E-119 | 1 L1 _ | Rpl10a |
| Limd2 | 2.36E-124 | 1.456918 | 0.752 | 0.243 | 4.75E-120 | 1 L1 _ | Limd2 |
| Apobec3 | 2.13E-124 | 1.243714 | 0.699 | 0.179 | 4.30E-120 | 1 L1 _ | Apobec3 |
| Klrb1f | 1.28E-124 | 0.756304 | 0.338 | 0.012 | 2.58E-120 | 1 L1 _ | Klrb1f |
| Sash3 | 5.10E-125 | 0.659137 | 0.354 | 0.015 | 1.03E-120 | 1 L1 _ | Sash3 |
| Pglyrp2 | 3.59E-125 | 0.644224 | 0.326 | 0.009 | 7.24E-121 | 1 L1 _ | Pglyrp2 |
| Ccr21 | 1.40E-125 | 0.989625 | 0.379 | 0.022 | 2.81E-121 | 1 L1 _ | Ccr2 |
| Rps16 | 1.19E-125 | 1.109895 | 0.979 | 0.982 | 2.39E-121 | 1 L1 _ | Rps16 |
| Klri21 | 2.72E-126 | 0.901519 | 0.375 | 0.02 | 5.47E-122 | 1 L1 _ | Klri2 |
| Slc9a3r1 | 1.03E-126 | 0.849082 | 0.48 | 0.052 | 2.07E-122 | 1 L1 _ | Slc9a3r1 |
| Rps27a | 5.08E-127 | 1.009015 | 0.989 | 0.978 | 1.02E-122 | 1 L1 _ | Rps27a |
| Il21r | 3.88E-127 | 0.702902 | 0.361 | 0.016 | 7.82E-123 | 1 L1 _ | Il21r |
| Srgn2 | 2.81E-127 | 1.38863 | 0.897 | 0.312 | 5.66E-123 | 1 L1 _ | Srgn |
| Gzmc | 1.50E-127 | 3.790966 | 0.692 | 0.204 | 3.01E-123 | 1 L1 _ | Gzmc |
| Rps3 | 1.15E-127 | 1.174925 | 0.977 | 0.943 | 2.32E-123 | 1 L1 _ | Rps3 |
| Rpl30 | 9.09E-128 | 1.073236 | 0.984 | 0.974 | 1.83E-123 | 1 L1 _ | Rpl30 |
| Rpl361 | 8.65E-128 | 1.198454 | 0.975 | 0.971 | 1.74E-123 | 1 L1 _ | Rpl36 |
| Cytip | 2.99E-128 | 0.774978 | 0.379 | 0.02 | 6.02E-124 | 1 L1 _ | Cytip |
| Rps9 | 1.32E-128 | 1.11913 | 0.977 | 0.968 | 2.66E-124 | 1 L1 _ | Rps9 |
| Rpl11 | 1.06E-128 | 1.054848 | 0.972 | 0.963 | 2.13E-124 | 1 L1 _ | Rpl11 |
| Traf3ip3 | 9.96E-129 | 0.652911 | 0.377 | 0.019 | 2.00E-124 | 1 L1 _ | Traf3ip3 |
| Klra7 | 2.22E-129 | 1.532629 | 0.455 | 0.045 | 4.47E-125 | 1 L1 _ | Klra7 |
| Rpl18 | 2.05E-129 | 1.140711 | 0.975 | 0.967 | 4.13E-125 | 1 L1 _ | Rpl18 |
| 12-Sep | 6.95E-130 | 1.322089 | 0.68 | 0.163 | 1.40E-125 | 1 L1 _ | 1-Sep |
| Rpl13a | 3.69E-130 | 1.359861 | 0.982 | 0.862 | 7.43E-126 | 1 L1 _ | Rpl13a |
| Rps20 | 4.32E-131 | 1.088654 | 0.995 | 0.972 | 8.70E-127 | 1 L1 _ | Rps20 |
| Sh2d2a | 6.82E-132 | 0.830326 | 0.405 | 0.025 | 1.37E-127 | 1 L1 _ | Sh2d2a |
| Ccl3 | 2.86E-132 | 1.287426 | 0.409 | 0.026 | 5.77E-128 | 1 L1 _ | Ccl3 |
| Rpl34 | 2.30E-132 | 1.212443 | 0.984 | 0.968 | 4.63E-128 | 1 L1 _ | Rpl34 |
| Rgs2 | 1.86E-132 | 1.370728 | 0.575 | 0.096 | 3.74E-128 | 1 L1 _ | Rgs2 |
| Tes | 7.35E-133 | 0.831811 | 0.487 | 0.05 | 1.48E-128 | 1 L1 _ | Tes |
| B4galnt11 | 4.10E-133 | 1.175877 | 0.641 | 0.132 | 8.26E-129 | 1 L1 _ | B4galnt1 |
| Rplp2 | 1.81E-133 | 1.146083 | 0.961 | 0.955 | 3.64E-129 | 1 L1 _ | Rplp2 |
| Tnfrsf181 | 1.76E-133 | 0.841526 | 0.423 | 0.029 | 3.54E-129 | 1 L1 _ | Tnfrsf18 |
| Itgal1 | 2.42E-134 | 0.780133 | 0.414 | 0.026 | 4.88E-130 | 1 L1 _ | Itgal |
| Rpsa | 6.48E-135 | 1.290134 | 0.986 | 0.974 | 1.31E-130 | 1 L1 _ | Rpsa |
| Btg1 | 3.59E-135 | 1.999736 | 0.949 | 0.728 | 7.24E-131 | 1 L1 _ | Btg1 |

| | | | | | | | |
|---|---|---|---|---|---|---|---|
| Samd31 | 2.15E-135 | 0.792872 | 0.389 | 0.018 | 4.34E-131 | 1 L1 _ | Samd3 |
| Rps7 | 1.86E-135 | 1.289639 | 0.977 | 0.963 | 3.75E-131 | 1 L1 _ | Rps7 |
| Rps5 | 4.69E-136 | 1.172259 | 0.982 | 0.964 | 9.45E-132 | 1 L1 _ | Rps5 |
| Id21 | 2.33E-136 | 1.91556 | 0.869 | 0.419 | 4.69E-132 | 1 L1 _ | Id2 |
| Hcls1 | 2.18E-136 | 1.035124 | 0.522 | 0.062 | 4.38E-132 | 1 L1 _ | Hcls1 |
| Rpl27 | 9.94E-137 | 1.39239 | 0.956 | 0.922 | 2.00E-132 | 1 L1 _ | Rpl27 |
| Gimap11 | 6.16E-137 | 1.476396 | 0.724 | 0.17 | 1.24E-132 | 1 L1 _ | Gimap1 |
| Rps10 | 3.54E-137 | 1.210156 | 0.979 | 0.974 | 7.12E-133 | 1 L1 _ | Rps10 |
| Rps14 | 2.04E-137 | 1.136779 | 0.989 | 0.97 | 4.10E-133 | 1 L1 _ | Rps14 |
| Padi2 | 1.38E-137 | 0.828853 | 0.414 | 0.024 | 2.79E-133 | 1 L1 _ | Padi2 |
| Rpl381 | 2.89E-138 | 1.319031 | 0.977 | 0.967 | 5.83E-134 | 1 L1 _ | Rpl38 |
| Cd3g | 2.23E-138 | 1.873649 | 0.434 | 0.031 | 4.49E-134 | 1 L1 _ | Cd3g |
| Dennd1c | 1.65E-138 | 0.813076 | 0.414 | 0.023 | 3.33E-134 | 1 L1 _ | Dennd1c |
| Car21 | 1.10E-138 | 1.325667 | 0.533 | 0.064 | 2.22E-134 | 1 L1 _ | Car2 |
| Klra41 | 2.13E-139 | 1.675176 | 0.444 | 0.033 | 4.29E-135 | 1 L1 _ | Klra4 |
| Rpl19 | 3.88E-140 | 1.13681 | 0.989 | 0.985 | 7.81E-136 | 1 L1 _ | Rpl19 |
| Rpl12 | 2.91E-140 | 1.311752 | 0.979 | 0.932 | 5.85E-136 | 1 L1 _ | Rpl12 |
| Gm15472 | 1.13E-140 | 0.708038 | 0.356 | 0.008 | 2.28E-136 | 1 L1 _ | Gm15472 |
| Acap11 | 6.13E-142 | 0.862013 | 0.439 | 0.028 | 1.23E-137 | 1 L1 _ | Acap1 |
| Prf11 | 1.99E-142 | 0.860533 | 0.411 | 0.02 | 4.01E-138 | 1 L1 _ | Prf1 |
| Klrc21 | 7.47E-143 | 1.012533 | 0.398 | 0.016 | 1.50E-138 | 1 L1 _ | Klrc2 |
| Rps18 | 2.41E-144 | 1.269967 | 0.989 | 0.958 | 4.86E-140 | 1 L1 _ | Rps18 |
| Rpl23 | 2.07E-144 | 1.158231 | 0.991 | 0.978 | 4.17E-140 | 1 L1 _ | Rpl23 |
| Ms4a6b1 | 8.41E-145 | 1.167393 | 0.43 | 0.024 | 1.69E-140 | 1 L1 _ | Ms4a6b |
| Lat | 1.80E-145 | 0.93813 | 0.405 | 0.017 | 3.63E-141 | 1 L1 _ | Lat |
| Trbc21 | 5.58E-146 | 2.045105 | 0.513 | 0.053 | 1.12E-141 | 1 L1 _ | Trbc2 |
| Tespa11 | 4.79E-146 | 0.826397 | 0.393 | 0.014 | 9.65E-142 | 1 L1 _ | Tespa1 |
| Traf1 | 1.14E-146 | 0.98632 | 0.428 | 0.023 | 2.30E-142 | 1 L1 _ | Traf1 |
| Gimap41 | 1.08E-146 | 1.751386 | 0.772 | 0.201 | 2.17E-142 | 1 L1 _ | Gimap4 |
| Myo1f1 | 1.71E-147 | 0.73097 | 0.425 | 0.021 | 3.45E-143 | 1 L1 _ | Myo1f |
| Stap11 | 1.13E-147 | 0.876977 | 0.444 | 0.026 | 2.28E-143 | 1 L1 _ | Stap1 |
| Rpl27a | 5.37E-148 | 1.177531 | 0.995 | 0.982 | 1.08E-143 | 1 L1 _ | Rpl27a |
| Slamf7 | 2.89E-148 | 0.857185 | 0.4 | 0.015 | 5.82E-144 | 1 L1 _ | Slamf7 |
| Rps24 | 1.33E-150 | 1.339749 | 0.998 | 0.976 | 2.68E-146 | 1 L1 _ | Rps24 |
| Prex11 | 2.93E-151 | 1.072593 | 0.577 | 0.071 | 5.91E-147 | 1 L1 _ | Prex1 |
| Rpl18a | 2.77E-152 | 1.285438 | 0.991 | 0.976 | 5.57E-148 | 1 L1 _ | Rpl18a |
| Cd96 | 1.02E-152 | 0.775077 | 0.418 | 0.016 | 2.06E-148 | 1 L1 _ | Cd96 |
| Sh3bgrl3 | 2.72E-156 | 1.849754 | 0.963 | 0.752 | 5.47E-152 | 1 L1 _ | Sh3bgrl3 |
| Rps13 | 5.23E-157 | 1.310696 | 0.991 | 0.964 | 1.05E-152 | 1 L1 _ | Rps13 |
| Klra8 | 3.91E-157 | 1.905978 | 0.526 | 0.05 | 7.88E-153 | 1 L1 _ | Klra8 |
| Tmsb4x2 | 2.15E-157 | 1.662345 | 0.998 | 0.994 | 4.34E-153 | 1 L1 _ | Tmsb4x |

| | | | | | | | |
|---|---|---|---|---|---|---|---|
| Cd531 | 1.23E-157 | 0.871043 | 0.462 | 0.025 | 2.47E-153 | 1 L1 _ | Cd53 |
| Lat21 | 5.31E-158 | 1.087723 | 0.515 | 0.043 | 1.07E-153 | 1 L1 _ | Lat2 |
| AW112010 | 2.66E-158 | 2.472802 | 0.959 | 0.668 | 5.36E-154 | 1 L1 _ | AW112010 |
| Rps15a | 9.77E-160 | 1.669017 | 0.989 | 0.975 | 1.97E-155 | 1 L1 _ | Rps15a |
| Rasal31 | 5.74E-160 | 0.857574 | 0.448 | 0.02 | 1.16E-155 | 1 L1 _ | Rasal3 |
| Stat41 | 4.82E-160 | 0.919258 | 0.434 | 0.017 | 9.71E-156 | 1 L1 _ | Stat4 |
| Rpl32 | 3.45E-160 | 1.416499 | 0.991 | 0.967 | 6.94E-156 | 1 L1 _ | Rpl32 |
| Dock8 | 6.53E-161 | 0.946226 | 0.554 | 0.051 | 1.32E-156 | 1 L1 _ | Dock8 |
| Ikzf11 | 5.13E-161 | 0.82041 | 0.453 | 0.021 | 1.03E-156 | 1 L1 _ | Ikzf1 |
| Klra91 | 4.44E-161 | 1.917673 | 0.531 | 0.048 | 8.93E-157 | 1 L1 _ | Klra9 |
| Zap701 | 4.11E-161 | 0.910469 | 0.467 | 0.025 | 8.27E-157 | 1 L1 _ | Zap70 |
| Ptk2b1 | 7.79E-163 | 0.963291 | 0.51 | 0.037 | 1.57E-158 | 1 L1 _ | Ptk2b |
| Arhgap9 | 5.76E-164 | 1.010375 | 0.533 | 0.043 | 1.16E-159 | 1 L1 _ | Arhgap9 |
| Fau | 7.99E-165 | 1.405987 | 0.991 | 0.984 | 1.61E-160 | 1 L1 _ | Fau |
| Rps11 | 4.22E-166 | 1.580156 | 0.989 | 0.97 | 8.50E-162 | 1 L1 _ | Rps11 |
| Ncf4 | 3.74E-167 | 0.890401 | 0.483 | 0.025 | 7.53E-163 | 1 L1 _ | Ncf4 |
| Ipcef11 | 2.39E-167 | 0.923468 | 0.474 | 0.023 | 4.82E-163 | 1 L1 _ | Ipcef1 |
| Sorl11 | 7.03E-169 | 0.963894 | 0.485 | 0.025 | 1.42E-164 | 1 L1 _ | Sorl1 |
| Sema4d | 2.70E-169 | 0.849324 | 0.434 | 0.012 | 5.44E-165 | 1 L1 _ | Sema4d |
| Ucp22 | 6.76E-170 | 1.691182 | 0.834 | 0.224 | 1.36E-165 | 1 L1 _ | Ucp2 |
| Ikzf31 | 1.78E-171 | 1.010533 | 0.467 | 0.019 | 3.58E-167 | 1 L1 _ | Ikzf3 |
| Itgae | 1.60E-171 | 1.369375 | 0.48 | 0.024 | 3.23E-167 | 1 L1 _ | Itgae |
| Cxcr61 | 1.88E-172 | 1.300298 | 0.483 | 0.023 | 3.79E-168 | 1 L1 _ | Cxcr6 |
| Styk11 | 3.45E-173 | 1.236668 | 0.492 | 0.025 | 6.94E-169 | 1 L1 _ | Styk1 |
| Ptpn6 | 1.71E-173 | 1.473456 | 0.586 | 0.059 | 3.44E-169 | 1 L1 _ | Ptpn6 |
| Slc38a12 | 1.28E-175 | 0.932774 | 0.48 | 0.02 | 2.58E-171 | 1 L1 _ | Slc38a1 |
| Ccl4 | 1.98E-176 | 2.050028 | 0.589 | 0.056 | 4.00E-172 | 1 L1 _ | Ccl4 |
| Gzmb | 5.99E-178 | 3.071863 | 0.694 | 0.116 | 1.21E-173 | 1 L1 _ | Gzmb |
| Itgax | 1.27E-179 | 1.36112 | 0.586 | 0.053 | 2.56E-175 | 1 L1 _ | Itgax |
| Bin21 | 1.07E-179 | 0.943083 | 0.501 | 0.024 | 2.15E-175 | 1 L1 _ | Bin2 |
| Pglyrp1 | 8.56E-180 | 1.226362 | 0.545 | 0.037 | 1.72E-175 | 1 L1 _ | Pglyrp1 |
| Cst71 | 7.60E-181 | 1.175427 | 0.515 | 0.027 | 1.53E-176 | 1 L1 _ | Cst7 |
| Cd481 | 3.86E-181 | 0.955474 | 0.494 | 0.021 | 7.77E-177 | 1 L1 _ | Cd48 |
| Cd21 | 9.45E-182 | 1.193684 | 0.531 | 0.032 | 1.90E-177 | 1 L1 _ | Cd2 |
| Serpina3g | 9.48E-188 | 1.452779 | 0.536 | 0.029 | 1.91E-183 | 1 L1 _ | Serpina3g |
| Ncr11 | 5.82E-188 | 1.367458 | 0.533 | 0.028 | 1.17E-183 | 1 L1 _ | Ncr1 |
| Itgb21 | 1.94E-188 | 1.022791 | 0.529 | 0.025 | 3.91E-184 | 1 L1 _ | Itgb2 |
| Gimap7 | 1.38E-190 | 1.260154 | 0.51 | 0.02 | 2.78E-186 | 1 L1 _ | Gimap7 |
| Ccl5 | 4.67E-191 | 4.414308 | 0.952 | 0.456 | 9.40E-187 | 1 L1 _ | Ccl5 |
| Klrk11 | 1.04E-191 | 1.759574 | 0.575 | 0.039 | 2.10E-187 | 1 L1 _ | Klrk1 |
| Tbc1d10c1 | 2.70E-192 | 1.185904 | 0.554 | 0.033 | 5.43E-188 | 1 L1 _ | Tbc1d10c |

| | | | | | | | |
|---|---|---|---|---|---|---|---|
| Tyrobp | 1.05E-196 | 3.61655 | 0.867 | 0.272 | 2.12E-192 | 1 L1 _ | Tyrobp |
| Ccr51 | 6.30E-197 | 1.24715 | 0.556 | 0.029 | 1.27E-192 | 1 L1 _ | Ccr5 |
| Fcer1g | 7.32E-198 | 3.712658 | 0.88 | 0.299 | 1.47E-193 | 1 L1 _ | Fcer1g |
| Trbc1 | 1.03E-198 | 1.681974 | 0.591 | 0.039 | 2.07E-194 | 1 L1 _ | Trbc1 |
| Fermt31 | 1.37E-200 | 1.060488 | 0.572 | 0.032 | 2.76E-196 | 1 L1 _ | Fermt3 |
| Cd71 | 6.76E-203 | 3.160685 | 0.733 | 0.116 | 1.36E-198 | 1 L1 _ | Cd7 |
| Cyth4 | 1.33E-203 | 1.331989 | 0.607 | 0.043 | 2.68E-199 | 1 L1 _ | Cyth4 |
| Arhgap451 | 2.43E-204 | 1.207649 | 0.621 | 0.045 | 4.90E-200 | 1 L1 _ | Arhgap45 |
| Neurl31 | 1.73E-204 | 1.264835 | 0.593 | 0.035 | 3.48E-200 | 1 L1 _ | Neurl3 |
| Arhgap30 | 7.47E-205 | 1.071798 | 0.563 | 0.027 | 1.50E-200 | 1 L1 _ | Arhgap30 |
| Cd244a | 7.21E-205 | 1.239233 | 0.522 | 0.016 | 1.45E-200 | 1 L1 _ | Cd244a |
| Ptpn221 | 1.10E-206 | 1.193445 | 0.533 | 0.018 | 2.21E-202 | 1 L1 _ | Ptpn22 |
| Klre11 | 7.72E-207 | 1.469215 | 0.568 | 0.027 | 1.56E-202 | 1 L1 _ | Klre1 |
| Xcl11 | 4.89E-208 | 3.266548 | 0.77 | 0.129 | 9.84E-204 | 1 L1 _ | Xcl1 |
| Rhoh | 1.03E-208 | 1.317037 | 0.568 | 0.027 | 2.07E-204 | 1 L1 _ | Rhoh |
| Cd226 | 2.43E-210 | 1.526177 | 0.556 | 0.023 | 4.90E-206 | 1 L1 _ | Cd226 |
| Ighm | 6.53E-211 | 1.457304 | 0.623 | 0.044 | 1.32E-206 | 1 L1 _ | Ighm |
| Dusp21 | 5.52E-214 | 1.987951 | 0.683 | 0.066 | 1.11E-209 | 1 L1 _ | Dusp2 |
| Skap11 | 1.06E-214 | 1.557444 | 0.687 | 0.067 | 2.14E-210 | 1 L1 _ | Skap1 |
| Rinl1 | 3.64E-222 | 1.551875 | 0.697 | 0.062 | 7.34E-218 | 1 L1 _ | Rinl |
| Klrb1c1 | 2.40E-231 | 1.92162 | 0.644 | 0.037 | 4.83E-227 | 1 L1 _ | Klrb1c |
| Dock21 | 1.21E-232 | 1.288423 | 0.628 | 0.03 | 2.43E-228 | 1 L1 _ | Dock2 |
| Ctsw | 2.55E-235 | 2.853686 | 0.867 | 0.173 | 5.14E-231 | 1 L1 _ | Ctsw |
| Il2rg1 | 2.07E-236 | 1.737611 | 0.733 | 0.068 | 4.18E-232 | 1 L1 _ | Il2rg |
| Ptpn181 | 9.86E-237 | 2.337889 | 0.876 | 0.168 | 1.98E-232 | 1 L1 _ | Ptpn18 |
| Epsti11 | 2.87E-237 | 1.501812 | 0.662 | 0.038 | 5.78E-233 | 1 L1 _ | Epsti1 |
| Fyb | 7.78E-241 | 1.904926 | 0.74 | 0.069 | 1.57E-236 | 1 L1 _ | Fyb |
| Cd521 | 2.97E-243 | 3.973661 | 0.963 | 0.332 | 5.98E-239 | 1 L1 _ | Cd52 |
| Txk1 | 4.65E-245 | 2.025372 | 0.667 | 0.036 | 9.36E-241 | 1 L1 _ | Txk |
| Klrd11 | 9.10E-248 | 2.21851 | 0.722 | 0.056 | 1.83E-243 | 1 L1 _ | Klrd1 |
| Cd371 | 8.48E-249 | 1.619183 | 0.733 | 0.058 | 1.71E-244 | 1 L1 _ | Cd37 |
| Nkg71 | 1.29E-249 | 3.332095 | 0.846 | 0.138 | 2.60E-245 | 1 L1 _ | Nkg7 |
| Lck | 7.51E-252 | 1.720016 | 0.706 | 0.046 | 1.51E-247 | 1 L1 _ | Lck |
| Gimap31 | 1.78E-254 | 1.947685 | 0.761 | 0.066 | 3.59E-250 | 1 L1 _ | Gimap3 |
| Il2rb1 | 9.39E-260 | 2.104634 | 0.761 | 0.064 | 1.89E-255 | 1 L1 _ | Il2rb |
| Runx31 | 9.65E-261 | 1.567855 | 0.685 | 0.03 | 1.94E-256 | 1 L1 _ | Runx3 |
| Hcst1 | 8.33E-262 | 2.337366 | 0.777 | 0.069 | 1.68E-257 | 1 L1 _ | Hcst |
| Lcp11 | 1.24E-262 | 2.23028 | 0.86 | 0.115 | 2.50E-258 | 1 L1 _ | Lcp1 |
| Cd691 | 1.53E-264 | 2.155166 | 0.726 | 0.043 | 3.09E-260 | 1 L1 _ | Cd69 |
| Ms4a4b1 | 3.68E-268 | 2.55112 | 0.768 | 0.06 | 7.42E-264 | 1 L1 _ | Ms4a4b |
| Coro1a1 | 1.45E-275 | 2.475978 | 0.883 | 0.122 | 2.91E-271 | 1 L1 _ | Coro1a |

| | | | | | | | |
|---|---|---|---|---|---|---|---|
| Rgs11 | 1.36E-276 | 2.798272 | 0.83 | 0.084 | 2.73E-272 | 1 L1 _ | Rgs1 |
| Ltb1 | 2.21E-278 | 2.693025 | 0.8 | 0.066 | 4.44E-274 | 1 L1 _ | Ltb |
| Ptprcap1 | 1.67E-289 | 2.250279 | 0.793 | 0.054 | 3.36E-285 | 1 L1 _ | Ptprcap |
| Laptm51 | 1.70E-291 | 2.40607 | 0.871 | 0.095 | 3.43E-287 | 1 L1 _ | Laptm5 |
| Rac21 | 3.05E-294 | 2.736721 | 0.906 | 0.119 | 6.15E-290 | 1 L1 _ | Rac2 |
| Selplg1 | 2.01E-299 | 2.035129 | 0.821 | 0.057 | 4.04E-295 | 1 L1 _ | Selplg |
| Ptprc1 | 0 | 2.922558 | 0.899 | 0.089 | 0 | 1 L1 _ | Ptprc |
| Hist1h1d | 2.34E-06 | 0.311672 | 0.357 | 0.056 | 0.047117 | 1 M1 _ | Hist1h1d |
| Il10ra | 2.31E-06 | 0.436435 | 0.286 | 0.038 | 0.046523 | 1 M1 _ | Il10ra |
| Eif3e | 2.16E-06 | 1.202077 | 0.929 | 0.66 | 0.043577 | 1 M1 _ | Eif3e |
| Arhgap15 | 2.01E-06 | 0.64443 | 0.571 | 0.138 | 0.040528 | 1 M1 _ | Arhgap15 |
| Rassf5 | 2.00E-06 | 0.51132 | 0.5 | 0.107 | 0.040376 | 1 M1 _ | Rassf5 |
| Ablim1 | 1.91E-06 | 1.188526 | 0.929 | 0.396 | 0.038532 | 1 M1 _ | Ablim1 |
| Apol9b | 1.83E-06 | 0.423694 | 0.357 | 0.057 | 0.036947 | 1 M1 _ | Apol9b |
| Cd48 | 1.77E-06 | 0.795157 | 0.5 | 0.106 | 0.03573 | 1 M1 _ | Cd48 |
| Cd5 | 1.76E-06 | 0.368183 | 0.214 | 0.022 | 0.035497 | 1 M1 _ | Cd5 |
| Tbc1d10c | 1.71E-06 | 0.513958 | 0.571 | 0.127 | 0.034369 | 1 M1 _ | Tbc1d10c |
| Dusp10 | 1.70E-06 | 0.291957 | 0.214 | 0.022 | 0.034174 | 1 M1 _ | Dusp10 |
| Mgea52 | 1.63E-06 | 0.476683 | 0.357 | 0.056 | 0.032781 | 1 M1 _ | Mgea5 |
| Ucp2 | 1.57E-06 | 1.167827 | 0.857 | 0.335 | 0.031656 | 1 M1 _ | Ucp2 |
| Fam69b1 | 1.54E-06 | 0.314229 | 0.214 | 0.022 | 0.030925 | 1 M1 _ | Fam69b |
| Gnpda1 | 1.48E-06 | 0.442552 | 0.5 | 0.101 | 0.029798 | 1 M1 _ | Gnpda1 |
| Gm17745 | 1.48E-06 | 0.389054 | 0.214 | 0.022 | 0.029792 | 1 M1 _ | Gm17745 |
| Trbc2 | 1.47E-06 | 1.209687 | 0.571 | 0.136 | 0.02951 | 1 M1 _ | Trbc2 |
| D3Ertd254e1 | 1.38E-06 | 0.268792 | 0.214 | 0.022 | 0.027757 | 1 M1 _ | D3Ertd254e |
| Sla | 1.34E-06 | 0.44743 | 0.429 | 0.077 | 0.026924 | 1 M1 _ | Sla |
| Wdr95 | 1.33E-06 | 0.368418 | 0.143 | 0.01 | 0.026721 | 1 M1 _ | Wdr95 |
| Klra9 | 1.28E-06 | 1.286366 | 0.571 | 0.136 | 0.025834 | 1 M1 _ | Klra9 |
| Mum12 | 1.26E-06 | 0.767519 | 0.357 | 0.057 | 0.02528 | 1 M1 _ | Mum1 |
| Rpl37a | 1.25E-06 | 1.182129 | 1 | 0.985 | 0.025203 | 1 M1 _ | Rpl37a |
| Sytl1 | 1.25E-06 | 0.66831 | 0.214 | 0.022 | 0.025166 | 1 M1 _ | Sytl1 |
| Fam208b1 | 1.21E-06 | 0.49014 | 0.286 | 0.036 | 0.024402 | 1 M1 _ | Fam208b |
| Zfp873 | 1.21E-06 | 0.33764 | 0.214 | 0.022 | 0.024329 | 1 M1 _ | Zfp873 |
| Zap70 | 1.15E-06 | 0.710975 | 0.5 | 0.105 | 0.023084 | 1 M1 _ | Zap70 |
| Saraf | 1.10E-06 | 0.94988 | 1 | 0.623 | 0.022158 | 1 M1 _ | Saraf |
| 3110062M04Rik1 | 1.03E-06 | 0.336004 | 0.286 | 0.036 | 0.020795 | 1 M1 _ | 3110062M04Rik |
| Zufsp2 | 1.02E-06 | 0.328232 | 0.286 | 0.036 | 0.020541 | 1 M1 _ | Zufsp |
| Rpl36 | 9.91E-07 | 1.053331 | 1 | 0.972 | 0.019953 | 1 M1 _ | Rpl36 |
| Rps28 | 9.51E-07 | 1.308448 | 1 | 0.963 | 0.019155 | 1 M1 _ | Rps28 |
| Cln3 | 8.97E-07 | 0.724166 | 0.571 | 0.14 | 0.018069 | 1 M1 _ | Cln3 |
| Klra3 | 8.92E-07 | 0.987027 | 0.429 | 0.077 | 0.017956 | 1 M1 _ | Klra3 |

| | | | | | | | |
|---|---|---|---|---|---|---|---|
| Cd53 | 8.27E-07 | 0.881132 | 0.5 | 0.104 | 0.016649 | 1 M1 _ | Cd53 |
| Xcl1 | 8.19E-07 | 0.962252 | 0.786 | 0.245 | 0.016495 | 1 M1 _ | Xcl1 |
| Gng2 | 8.02E-07 | 0.818679 | 0.786 | 0.254 | 0.016158 | 1 M1 _ | Gng2 |
| AB124611 | 7.98E-07 | 0.295108 | 0.286 | 0.035 | 0.016082 | 1 M1 _ | AB124611 |
| Jaml | 7.96E-07 | 0.46065 | 0.571 | 0.12 | 0.016026 | 1 M1 _ | Jaml |
| Atp5s | 7.95E-07 | 0.287741 | 0.143 | 0.009 | 0.016008 | 1 M1 _ | Atp5s |
| Foxm1 | 7.67E-07 | 0.25327 | 0.143 | 0.009 | 0.015457 | 1 M1 _ | Foxm1 |
| Tmsb4x | 7.52E-07 | 1.610477 | 1 | 0.994 | 0.015142 | 1 M1 _ | Tmsb4x |
| Lat2 | 7.31E-07 | 0.812017 | 0.571 | 0.128 | 0.014714 | 1 M1 _ | Lat2 |
| Prc1 | 6.91E-07 | 0.573809 | 0.143 | 0.009 | 0.013912 | 1 M1 _ | Prc1 |
| Slc38a1 | 6.29E-07 | 1.014246 | 0.5 | 0.103 | 0.012665 | 1 M1 _ | Slc38a1 |
| Rin3 | 6.17E-07 | 0.7755 | 0.571 | 0.14 | 0.012429 | 1 M1 _ | Rin3 |
| Grip2 | 5.92E-07 | 0.257994 | 0.214 | 0.02 | 0.011923 | 1 M1 _ | Grip2 |
| AU020206 | 5.76E-07 | 0.67142 | 0.786 | 0.224 | 0.011608 | 1 M1 _ | AU020206 |
| Samsn1 | 5.76E-07 | 0.348166 | 0.429 | 0.071 | 0.011599 | 1 M1 _ | Samsn1 |
| Id2 | 4.66E-07 | 1.552989 | 0.929 | 0.5 | 0.009381 | 1 M1 _ | Id2 |
| Narfl | 4.48E-07 | 0.306182 | 0.214 | 0.02 | 0.009019 | 1 M1 _ | Narfl |
| Rps29 | 4.43E-07 | 1.318314 | 1 | 0.986 | 0.008916 | 1 M1 _ | Rps29 |
| Cd40lg | 4.19E-07 | 0.343894 | 0.143 | 0.009 | 0.00843 | 1 M1 _ | Cd40lg |
| Ikzf1 | 4.02E-07 | 0.931127 | 0.5 | 0.099 | 0.008088 | 1 M1 _ | Ikzf1 |
| 0610037L13Rik2 | 3.92E-07 | 0.89243 | 0.357 | 0.053 | 0.007886 | 1 M1 _ | 0610037L13Rik |
| AA467197 | 3.45E-07 | 0.809241 | 0.214 | 0.02 | 0.006948 | 1 M1 _ | AA467197 |
| Cd69 | 3.26E-07 | 1.181668 | 0.714 | 0.167 | 0.006568 | 1 M1 _ | Cd69 |
| Hvcn1 | 3.21E-07 | 0.694539 | 0.643 | 0.157 | 0.00647 | 1 M1 _ | Hvcn1 |
| Cd7 | 2.89E-07 | 1.367656 | 0.786 | 0.227 | 0.005812 | 1 M1 _ | Cd7 |
| Crybg1 | 2.80E-07 | 0.404075 | 0.286 | 0.033 | 0.005631 | 1 M1 _ | Crybg1 |
| Rapsn | 2.68E-07 | 0.386336 | 0.214 | 0.019 | 0.005396 | 1 M1 _ | Rapsn |
| Ptpn22 | 2.48E-07 | 0.47912 | 0.571 | 0.112 | 0.004993 | 1 M1 _ | Ptpn22 |
| Klra4 | 2.46E-07 | 2.352667 | 0.5 | 0.107 | 0.004962 | 1 M1 _ | Klra4 |
| Stat4 | 2.35E-07 | 0.743259 | 0.5 | 0.092 | 0.004726 | 1 M1 _ | Stat4 |
| Prex1 | 2.34E-07 | 1.036567 | 0.643 | 0.162 | 0.004707 | 1 M1 _ | Prex1 |
| Gfi1 | 2.31E-07 | 0.569924 | 0.214 | 0.019 | 0.004659 | 1 M1 _ | Gfi1 |
| Rpl38 | 2.25E-07 | 1.322965 | 1 | 0.969 | 0.004525 | 1 M1 _ | Rpl38 |
| Gm26882 | 2.22E-07 | 0.267723 | 0.143 | 0.009 | 0.004467 | 1 M1 _ | Gm26882 |
| Ms4a6b | 2.02E-07 | 1.686675 | 0.5 | 0.097 | 0.004077 | 1 M1 _ | Ms4a6b |
| Lax1 | 1.92E-07 | 0.580527 | 0.357 | 0.051 | 0.003869 | 1 M1 _ | Lax1 |
| Cenpq | 1.87E-07 | 0.474886 | 0.357 | 0.05 | 0.003761 | 1 M1 _ | Cenpq |
| Pfn11 | 1.74E-07 | 1.597674 | 1 | 0.794 | 0.003504 | 1 M1 _ | Pfn1 |
| Il2rb | 1.66E-07 | 1.261233 | 0.714 | 0.191 | 0.003338 | 1 M1 _ | Il2rb |
| Car2 | 1.61E-07 | 1.247872 | 0.643 | 0.149 | 0.00325 | 1 M1 _ | Car2 |
| Ncr1 | 1.61E-07 | 1.311824 | 0.571 | 0.12 | 0.003248 | 1 M1 _ | Ncr1 |

| | | | | | | | |
|---|---|---|---|---|---|---|---|
| Ikbkg | 1.57E-07 | 0.427069 | 0.5 | 0.089 | 0.003163 | 1 M1 _ | Ikbkg |
| Napsa | 1.42E-07 | 1.146743 | 0.357 | 0.051 | 0.002868 | 1 M1 _ | Napsa |
| Camk1d | 1.41E-07 | 0.406279 | 0.429 | 0.067 | 0.002847 | 1 M1 _ | Camk1d |
| Fgr | 1.25E-07 | 0.277071 | 0.286 | 0.031 | 0.002522 | 1 M1 _ | Fgr |
| Hmgb2 | 9.24E-08 | 1.950274 | 0.929 | 0.39 | 0.001861 | 1 M1 _ | Hmgb2 |
| Pyurf2 | 8.76E-08 | 0.414219 | 0.357 | 0.046 | 0.001765 | 1 M1 _ | Pyurf |
| Dock2 | 8.61E-08 | 0.767083 | 0.643 | 0.138 | 0.001734 | 1 M1 _ | Dock2 |
| Fam103a12 | 8.34E-08 | 0.53999 | 0.429 | 0.065 | 0.001679 | 1 M1 _ | Fam103a1 |
| Acsbg1 | 5.98E-08 | 0.376427 | 0.214 | 0.018 | 0.001204 | 1 M1 _ | Acsbg1 |
| Fam102a | 5.57E-08 | 0.933095 | 0.643 | 0.161 | 0.001122 | 1 M1 _ | Fam102a |
| Rhof | 5.40E-08 | 0.957671 | 0.429 | 0.067 | 0.001087 | 1 M1 _ | Rhof |
| Arhgap25 | 5.05E-08 | 0.497547 | 0.429 | 0.065 | 0.001018 | 1 M1 _ | Arhgap25 |
| Klrc2 | 4.83E-08 | 0.741863 | 0.5 | 0.085 | 0.000973 | 1 M1 _ | Klrc2 |
| Srgn | 4.78E-08 | 1.596926 | 1 | 0.418 | 0.000962 | 1 M1 _ | Srgn |
| Klrb1c | 4.67E-08 | 1.637588 | 0.643 | 0.147 | 0.00094 | 1 M1 _ | Klrb1c |
| Clnk | 4.63E-08 | 0.880036 | 0.429 | 0.067 | 0.000932 | 1 M1 _ | Clnk |
| Naip2 | 4.37E-08 | 0.77204 | 0.286 | 0.03 | 0.000879 | 1 M1 _ | Naip2 |
| Trat1 | 4.18E-08 | 0.305355 | 0.143 | 0.008 | 0.000843 | 1 M1 _ | Trat1 |
| Mirt1 | 3.90E-08 | 0.564105 | 0.214 | 0.017 | 0.000785 | 1 M1 _ | Mirt1 |
| Vav1 | 3.88E-08 | 0.61378 | 0.429 | 0.065 | 0.00078 | 1 M1 _ | Vav1 |
| Tcf7 | 3.36E-08 | 0.591859 | 0.286 | 0.03 | 0.000677 | 1 M1 _ | Tcf7 |
| Gpr174 | 3.22E-08 | 0.581214 | 0.357 | 0.046 | 0.000649 | 1 M1 _ | Gpr174 |
| Tespa1 | 2.81E-08 | 0.545047 | 0.5 | 0.082 | 0.000566 | 1 M1 _ | Tespa1 |
| Rgs1 | 2.66E-08 | 2.110985 | 0.786 | 0.219 | 0.000536 | 1 M1 _ | Rgs1 |
| Prf1 | 2.62E-08 | 1.349123 | 0.5 | 0.09 | 0.000528 | 1 M1 _ | Prf1 |
| Klri2 | 2.54E-08 | 0.719601 | 0.5 | 0.084 | 0.000511 | 1 M1 _ | Klri2 |
| B4galnt1 | 2.28E-08 | 1.034694 | 0.786 | 0.223 | 0.000458 | 1 M1 _ | B4galnt1 |
| Nkg7 | 2.25E-08 | 1.929975 | 0.857 | 0.267 | 0.000452 | 1 M1 _ | Nkg7 |
| Rinl | 2.17E-08 | 1.323613 | 0.714 | 0.177 | 0.000438 | 1 M1 _ | Rinl |
| Neurl3 | 1.99E-08 | 1.035838 | 0.643 | 0.136 | 0.000401 | 1 M1 _ | Neurl3 |
| Cxcr6 | 1.80E-08 | 0.883002 | 0.571 | 0.106 | 0.000362 | 1 M1 _ | Cxcr6 |
| Selp | 1.68E-08 | 0.726261 | 0.143 | 0.007 | 0.000339 | 1 M1 _ | Selp |
| Ptpn18 | 1.44E-08 | 1.452926 | 0.929 | 0.296 | 0.00029 | 1 M1 _ | Ptpn18 |
| Cd37 | 1.41E-08 | 0.805701 | 0.786 | 0.18 | 0.000284 | 1 M1 _ | Cd37 |
| Fcho1 | 1.36E-08 | 0.646315 | 0.286 | 0.029 | 0.000275 | 1 M1 _ | Fcho1 |
| Fermt3 | 1.36E-08 | 1.355109 | 0.643 | 0.13 | 0.000274 | 1 M1 _ | Fermt3 |
| Gm265261 | 1.27E-08 | 0.349544 | 0.286 | 0.028 | 0.000256 | 1 M1 _ | Gm26526 |
| Gpr65 | 1.23E-08 | 0.62979 | 0.5 | 0.08 | 0.000249 | 1 M1 _ | Gpr65 |
| Stk17b | 1.12E-08 | 1.411062 | 0.929 | 0.342 | 0.000225 | 1 M1 _ | Stk17b |
| Vipr2 | 1.09E-08 | 0.311708 | 0.357 | 0.041 | 0.000219 | 1 M1 _ | Vipr2 |
| Samd3 | 1.08E-08 | 1.025523 | 0.5 | 0.085 | 0.000217 | 1 M1 _ | Samd3 |

| | | | | | | | |
|---|---|---|---|---|---|---|---|
| Clec2i | 1.07E-08 | 0.8894 | 0.357 | 0.043 | 0.000216 | 1 M1 _ | Clec2i |
| Cd52 | 9.67E-09 | 2.058759 | 1 | 0.446 | 0.000195 | 1 M1 _ | Cd52 |
| Klhl6 | 9.19E-09 | 0.650985 | 0.5 | 0.078 | 0.000185 | 1 M1 _ | Klhl6 |
| Selplg | 7.65E-09 | 1.279135 | 0.786 | 0.196 | 0.000154 | 1 M1 _ | Selplg |
| Dapp1 | 7.19E-09 | 0.645047 | 0.5 | 0.082 | 0.000145 | 1 M1 _ | Dapp1 |
| Arnt2 | 5.88E-09 | 0.363569 | 0.357 | 0.041 | 0.000118 | 1 M1 _ | Arnt2 |
| Skap1 | 5.45E-09 | 0.991509 | 0.786 | 0.179 | 0.00011 | 1 M1 _ | Skap1 |
| Evl | 5.40E-09 | 1.343079 | 0.929 | 0.297 | 0.000109 | 1 M1 _ | Evl |
| Styk1 | 5.02E-09 | 1.648429 | 0.571 | 0.109 | 0.000101 | 1 M1 _ | Styk1 |
| Ptk2b | 5.01E-09 | 0.752783 | 0.643 | 0.122 | 0.000101 | 1 M1 _ | Ptk2b |
| Rab27a | 4.58E-09 | 0.601278 | 0.286 | 0.027 | 9.22E-05 | 1 M1 _ | Rab27a |
| Tmem2 | 4.58E-09 | 0.459856 | 0.286 | 0.027 | 9.22E-05 | 1 M1 _ | Tmem2 |
| Cd2 | 4.33E-09 | 0.836404 | 0.643 | 0.121 | 8.72E-05 | 1 M1 _ | Cd2 |
| Rac2 | 4.21E-09 | 1.430177 | 0.929 | 0.262 | 8.48E-05 | 1 M1 _ | Rac2 |
| Klre1 | 4.03E-09 | 1.261233 | 0.643 | 0.124 | 8.11E-05 | 1 M1 _ | Klre1 |
| Acap1 | 3.81E-09 | 1.154066 | 0.571 | 0.102 | 7.68E-05 | 1 M1 _ | Acap1 |
| Cst7 | 3.64E-09 | 0.669243 | 0.643 | 0.115 | 7.34E-05 | 1 M1 _ | Cst7 |
| Arhgap4 | 3.46E-09 | 0.518254 | 0.429 | 0.056 | 6.97E-05 | 1 M1 _ | Arhgap4 |
| Rbm121 | 3.07E-09 | 0.316554 | 0.286 | 0.026 | 6.19E-05 | 1 M1 _ | Rbm12 |
| Ccr5 | 3.00E-09 | 1.261309 | 0.643 | 0.124 | 6.03E-05 | 1 M1 _ | Ccr5 |
| Smpd5 | 2.93E-09 | 0.621751 | 0.429 | 0.058 | 5.91E-05 | 1 M1 _ | Smpd5 |
| Gm9493 | 2.35E-09 | 0.389152 | 0.214 | 0.015 | 4.73E-05 | 1 M1 _ | Gm9493 |
| Parvg | 2.20E-09 | 0.779701 | 0.357 | 0.04 | 4.44E-05 | 1 M1 _ | Parvg |
| Tbx21 | 1.97E-09 | 0.509688 | 0.429 | 0.056 | 3.97E-05 | 1 M1 _ | Tbx21 |
| Ms4a4b | 1.77E-09 | 1.799564 | 0.786 | 0.188 | 3.56E-05 | 1 M1 _ | Ms4a4b |
| Txk | 1.59E-09 | 1.837168 | 0.714 | 0.15 | 3.20E-05 | 1 M1 _ | Txk |
| Mkl1 | 1.41E-09 | 0.440277 | 0.286 | 0.025 | 2.85E-05 | 1 M1 _ | Mkl1 |
| Ptp4a12 | 1.29E-09 | 0.536682 | 0.5 | 0.071 | 2.60E-05 | 1 M1 _ | Ptp4a1 |
| 1-Sep | 1.21E-09 | 1.049045 | 0.929 | 0.255 | 2.43E-05 | 1 M1 _ | 1-Sep |
| Fam46a1 | 1.12E-09 | 1.207785 | 0.429 | 0.055 | 2.25E-05 | 1 M1 _ | Fam46a |
| Gab3 | 1.09E-09 | 0.412014 | 0.357 | 0.037 | 2.20E-05 | 1 M1 _ | Gab3 |
| B3gnt5 | 1.07E-09 | 0.413827 | 0.214 | 0.014 | 2.16E-05 | 1 M1 _ | B3gnt5 |
| Myo1f | 9.62E-10 | 0.671238 | 0.571 | 0.093 | 1.94E-05 | 1 M1 _ | Myo1f |
| Hist1h2bb | 5.45E-10 | 0.286971 | 0.143 | 0.006 | 1.10E-05 | 1 M1 _ | Hist1h2bb |
| Fam96b2 | 4.18E-10 | 0.824863 | 0.5 | 0.07 | 8.43E-06 | 1 M1 _ | Fam96b |
| Lpxn | 4.00E-10 | 0.658605 | 0.5 | 0.068 | 8.06E-06 | 1 M1 _ | Lpxn |
| Tmem1101 | 3.79E-10 | 0.603516 | 0.357 | 0.037 | 7.64E-06 | 1 M1 _ | Tmem110 |
| Fam208a2 | 3.58E-10 | 0.770982 | 0.571 | 0.084 | 7.21E-06 | 1 M1 _ | Fam208a |
| Pkp3 | 3.19E-10 | 0.704966 | 0.571 | 0.087 | 6.42E-06 | 1 M1 _ | Pkp3 |
| Ptpre | 2.39E-10 | 0.904188 | 0.857 | 0.183 | 4.82E-06 | 1 M1 _ | Ptpre |
| Ptprcap | 1.83E-10 | 1.361737 | 0.857 | 0.188 | 3.69E-06 | 1 M1 _ | Ptprcap |

| | | | | | | | |
|---|---|---|---|---|---|---|---|
| Lppos1 | 1.74E-10 | 0.489261 | 0.286 | 0.023 | 3.51E-06 | 1 M1 _ | Lppos |
| Psd4 | 1.70E-10 | 0.736296 | 0.5 | 0.07 | 3.42E-06 | 1 M1 _ | Psd4 |
| Bin2 | 1.67E-10 | 0.969817 | 0.643 | 0.109 | 3.37E-06 | 1 M1 _ | Bin2 |
| Sit1 | 1.50E-10 | 0.552632 | 0.214 | 0.013 | 3.02E-06 | 1 M1 _ | Sit1 |
| Gbp8 | 1.04E-10 | 0.830915 | 0.5 | 0.068 | 2.09E-06 | 1 M1 _ | Gbp8 |
| Tmem163 | 8.44E-11 | 0.788093 | 0.429 | 0.05 | 1.70E-06 | 1 M1 _ | Tmem163 |
| Stap1 | 8.38E-11 | 0.61677 | 0.643 | 0.1 | 1.69E-06 | 1 M1 _ | Stap1 |
| Cyfip2 | 8.12E-11 | 0.854449 | 0.571 | 0.086 | 1.63E-06 | 1 M1 _ | Cyfip2 |
| Klrk1 | 7.69E-11 | 1.516261 | 0.714 | 0.136 | 1.55E-06 | 1 M1 _ | Klrk1 |
| Ikzf3 | 7.00E-11 | 0.717783 | 0.643 | 0.099 | 1.41E-06 | 1 M1 _ | Ikzf3 |
| Sh2d1b1 | 6.09E-11 | 0.754995 | 0.5 | 0.065 | 1.23E-06 | 1 M1 _ | Sh2d1b1 |
| Ptprc | 4.14E-11 | 2.000281 | 0.929 | 0.236 | 8.34E-07 | 1 M1 _ | Ptprc |
| Xlr4a | 3.64E-11 | 0.477568 | 0.214 | 0.012 | 7.33E-07 | 1 M1 _ | Xlr4a |
| Gimap3 | 3.46E-11 | 0.91703 | 0.929 | 0.191 | 6.96E-07 | 1 M1 _ | Gimap3 |
| Zcchc62 | 2.69E-11 | 0.993212 | 0.643 | 0.097 | 5.41E-07 | 1 M1 _ | Zcchc6 |
| A430046D13Rik | 2.62E-11 | 0.261359 | 0.143 | 0.005 | 5.27E-07 | 1 M1 _ | A430046D13Rik |
| 1110008F13Rik2 | 2.60E-11 | 0.68454 | 0.643 | 0.093 | 5.23E-07 | 1 M1 _ | 1110008F13Rik |
| Gm5547 | 2.01E-11 | 0.412063 | 0.357 | 0.032 | 4.04E-07 | 1 M1 _ | Gm5547 |
| Serpinb6b | 1.73E-11 | 1.097059 | 0.5 | 0.065 | 3.49E-07 | 1 M1 _ | Serpinb6b |
| Gcn1l12 | 1.19E-11 | 0.552424 | 0.429 | 0.045 | 2.40E-07 | 1 M1 _ | Gcn1l1 |
| Dusp2 | 1.16E-11 | 1.462799 | 0.857 | 0.177 | 2.33E-07 | 1 M1 _ | Dusp2 |
| Laptm5 | 1.13E-11 | 1.779198 | 1 | 0.235 | 2.28E-07 | 1 M1 _ | Laptm5 |
| Cdh1 | 1.04E-11 | 0.79553 | 0.5 | 0.062 | 2.10E-07 | 1 M1 _ | Cdh1 |
| Lcp1 | 9.47E-12 | 1.632119 | 1 | 0.249 | 1.91E-07 | 1 M1 _ | Lcp1 |
| Klrd1 | 8.88E-12 | 1.84121 | 0.857 | 0.176 | 1.79E-07 | 1 M1 _ | Klrd1 |
| Hcst | 5.68E-12 | 1.607154 | 0.929 | 0.196 | 1.14E-07 | 1 M1 _ | Hcst |
| Cited4 | 4.44E-12 | 1.182158 | 0.643 | 0.093 | 8.94E-08 | 1 M1 _ | Cited4 |
| Tnfrsf18 | 4.42E-12 | 0.99364 | 0.643 | 0.099 | 8.90E-08 | 1 M1 _ | Tnfrsf18 |
| Coro1a | 3.82E-12 | 1.711574 | 1 | 0.259 | 7.69E-08 | 1 M1 _ | Coro1a |
| Ltb | 3.33E-12 | 1.548587 | 0.929 | 0.198 | 6.70E-08 | 1 M1 _ | Ltb |
| Akna | 3.30E-12 | 1.035619 | 0.714 | 0.12 | 6.64E-08 | 1 M1 _ | Akna |
| Il2rg | 3.09E-12 | 1.17648 | 0.929 | 0.188 | 6.22E-08 | 1 M1 _ | Il2rg |
| B930036N10Rik | 2.89E-12 | 1.134861 | 0.643 | 0.103 | 5.83E-08 | 1 M1 _ | B930036N10Rik |
| Runx3 | 2.67E-12 | 1.853393 | 0.786 | 0.148 | 5.39E-08 | 1 M1 _ | Runx3 |
| Nup210 | 2.61E-12 | 0.827655 | 0.5 | 0.058 | 5.26E-08 | 1 M1 _ | Nup210 |
| Minos12 | 1.12E-12 | 1.007759 | 0.786 | 0.125 | 2.26E-08 | 1 M1 _ | Minos1 |
| AI606181 | 9.56E-13 | 0.35974 | 0.214 | 0.01 | 1.92E-08 | 1 M1 _ | AI606181 |
| Aim2 | 8.92E-13 | 0.901531 | 0.571 | 0.072 | 1.80E-08 | 1 M1 _ | Aim2 |
| Rpl27-ps3 | 8.57E-13 | 0.430382 | 0.214 | 0.01 | 1.73E-08 | 1 M1 _ | Rpl27-ps3 |
| Gm15232 | 6.43E-13 | 0.350264 | 0.286 | 0.018 | 1.30E-08 | 1 M1 _ | Gm15232 |
| Pla2g161 | 5.88E-13 | 0.938539 | 0.429 | 0.043 | 1.18E-08 | 1 M1 _ | Pla2g16 |

| | | | | | | | |
|---|---|---|---|---|---|---|---|
| AU0198232 | 5.40E-13 | 0.579692 | 0.429 | 0.04 | 1.09E-08 | 1 M1 _ | AU019823 |
| Usmg52 | 4.87E-13 | 1.296972 | 0.786 | 0.13 | 9.81E-09 | 1 M1 _ | Usmg5 |
| Trdc | 3.08E-13 | 1.133676 | 0.357 | 0.028 | 6.20E-09 | 1 M1 _ | Trdc |
| Ccr2 | 3.03E-13 | 0.885193 | 0.643 | 0.085 | 6.10E-09 | 1 M1 _ | Ccr2 |
| Sorl1 | 2.76E-13 | 0.90881 | 0.714 | 0.107 | 5.56E-09 | 1 M1 _ | Sorl1 |
| Arhgap45 | 2.55E-13 | 0.910495 | 0.857 | 0.148 | 5.14E-09 | 1 M1 _ | Arhgap45 |
| 2010107E04Rik2 | 2.12E-13 | 1.151208 | 0.786 | 0.127 | 4.26E-09 | 1 M1 _ | 2010107E04Rik |
| Klrb1b | 1.69E-13 | 1.091136 | 0.429 | 0.04 | 3.40E-09 | 1 M1 _ | Klrb1b |
| Lpcat4 | 8.63E-14 | 0.851811 | 0.214 | 0.009 | 1.74E-09 | 1 M1 _ | Lpcat4 |
| Itgb7 | 6.30E-14 | 0.903802 | 0.643 | 0.083 | 1.27E-09 | 1 M1 _ | Itgb7 |
| Spn | 6.10E-14 | 0.828541 | 0.571 | 0.066 | 1.23E-09 | 1 M1 _ | Spn |
| Ly9 | 5.99E-14 | 0.471982 | 0.357 | 0.026 | 1.21E-09 | 1 M1 _ | Ly9 |
| Kif21b | 2.91E-14 | 0.749965 | 0.571 | 0.065 | 5.85E-10 | 1 M1 _ | Kif21b |
| 1810022K09Rik2 | 2.62E-14 | 1.009577 | 0.714 | 0.101 | 5.28E-10 | 1 M1 _ | 1810022K09Rik |
| Rasal3 | 2.01E-14 | 0.776452 | 0.714 | 0.096 | 4.05E-10 | 1 M1 _ | Rasal3 |
| Itgb2 | 1.35E-14 | 1.008499 | 0.786 | 0.115 | 2.72E-10 | 1 M1 _ | Itgb2 |
| Itgal | 1.20E-14 | 1.046629 | 0.714 | 0.094 | 2.42E-10 | 1 M1 _ | Itgal |
| C920006O11Rik | 8.92E-15 | 0.312038 | 0.214 | 0.009 | 1.80E-10 | 1 M1 _ | C920006O11Rik |
| Ipcef1 | 8.79E-15 | 1.492504 | 0.714 | 0.103 | 1.77E-10 | 1 M1 _ | Ipcef1 |
| D17Wsu92e2 | 5.88E-15 | 0.580862 | 0.571 | 0.059 | 1.18E-10 | 1 M1 _ | D17Wsu92e |
| Ccr9 | 1.69E-15 | 0.435168 | 0.143 | 0.003 | 3.40E-11 | 1 M1 _ | Ccr9 |
| Gm26722 | 1.35E-15 | 0.353229 | 0.143 | 0.003 | 2.71E-11 | 1 M1 _ | Gm26722 |
| Glipr1 | 1.22E-15 | 1.413765 | 0.643 | 0.074 | 2.46E-11 | 1 M1 _ | Glipr1 |
| Epsti1 | 4.56E-16 | 1.455547 | 0.929 | 0.15 | 9.19E-12 | 1 M1 _ | Epsti1 |
| Fmnl1 | 4.42E-16 | 0.931418 | 0.714 | 0.091 | 8.89E-12 | 1 M1 _ | Fmnl1 |
| Atp5o.12 | 3.27E-16 | 1.287291 | 0.857 | 0.121 | 6.58E-12 | 1 M1 _ | Atp5o.1 |
| 2410015M20Rik2 | 1.41E-16 | 0.971657 | 0.857 | 0.114 | 2.83E-12 | 1 M1 _ | 2410015M20Rik |
| Zcchc112 | 5.85E-17 | 1.211646 | 0.643 | 0.071 | 1.18E-12 | 1 M1 _ | Zcchc11 |
| Il18rap | 5.68E-17 | 0.475507 | 0.286 | 0.013 | 1.14E-12 | 1 M1 _ | Il18rap |
| Jakmip1 | 1.45E-18 | 0.60855 | 0.5 | 0.037 | 2.93E-14 | 1 M1 _ | Jakmip1 |
| Haspin | 3.75E-19 | 0.332743 | 0.286 | 0.012 | 7.56E-15 | 1 M1 _ | Haspin |
| Tnfrsf13c | 3.33E-19 | 0.281364 | 0.143 | 0.003 | 6.71E-15 | 1 M1 _ | Tnfrsf13c |
| Aes2 | 2.68E-19 | 2.00284 | 0.929 | 0.134 | 5.40E-15 | 1 M1 _ | Aes |
| Gpr132 | 2.35E-19 | 0.6958 | 0.357 | 0.019 | 4.73E-15 | 1 M1 _ | Gpr132 |
| Prr36 | 4.05E-21 | 0.308881 | 0.214 | 0.006 | 8.16E-17 | 1 M1 _ | Prr36 |
| Gm26853 | 1.51E-22 | 0.347901 | 0.357 | 0.015 | 3.05E-18 | 1 M1 _ | Gm26853 |
| Il18r1 | 1.25E-22 | 1.12364 | 0.357 | 0.016 | 2.52E-18 | 1 M1 _ | Il18r1 |
| 2810474O19Rik2 | 1.40E-23 | 1.308218 | 0.786 | 0.076 | 2.81E-19 | 1 M1 _ | 2810474O19Rik |
| 1810011H11Rik1 | 2.12E-27 | 0.651687 | 0.357 | 0.012 | 4.28E-23 | 1 M1 _ | 1810011H11Rik |
| A730071L15Rik | 1.30E-30 | 0.367248 | 0.214 | 0.003 | 2.61E-26 | 1 M1 _ | A730071L15Rik |
| AL590144.2 | 5.80E-33 | 0.579788 | 0.286 | 0.006 | 1.17E-28 | 1 M1 _ | AL590144.2 |

| | | | | | | | |
|---|---|---|---|---|---|---|---|
| Fam60a | 3.48E-38 | 0.542845 | 0.429 | 0.012 | 7.01E-34 | 1 M1 _ | Fam60a |
| Fam105a | 4.60E-46 | 0.861878 | 0.357 | 0.006 | 9.26E-42 | 1 M1 _ | Fam105a |
| Gm26884 | 2.01E-66 | 0.272701 | 0.214 | 0.001 | 4.04E-62 | 1 M1 _ | Gm26884 |
| Gm28053 | 4.07E-74 | 0.303334 | 0.143 | 0 | 8.20E-70 | 1 M1 _ | Gm28053 |
| Dmrta1 | 4.07E-74 | 0.261138 | 0.143 | 0 | 8.20E-70 | 1 M1 _ | Dmrta1 |
| B130006D01Rik | 3.24E-91 | 0.417404 | 0.357 | 0.002 | 6.53E-87 | 1 M1 _ | B130006D01Rik |
| Gm26541 | 8.44E-102 | 0.575875 | 0.357 | 0.002 | 1.70E-97 | 1 M1 _ | Gm26541 |
| AC163354.1 | 2.78E-110 | 0.429426 | 0.214 | 0 | 5.61E-106 | 1 M1 _ | AC163354.1 |
| Cd244 | 1.95E-130 | 0.783626 | 0.357 | 0.001 | 3.93E-126 | 1 M1 _ | Cd244 |
| Ptp4a31 | 2.43E-06 | 0.900777 | 0.383 | 0.32 | 0.04884 | 2 L1 _ | Ptp4a3 |
| G0s22 | 2.31E-06 | 0.636234 | 0.54 | 0.52 | 0.046516 | 2 L1 _ | G0s2 |
| Gnai11 | 2.13E-06 | 0.397178 | 0.128 | 0.058 | 0.042993 | 2 L1 _ | Gnai1 |
| Tmem941 | 2.13E-06 | 0.537286 | 0.204 | 0.12 | 0.042852 | 2 L1 _ | Tmem94 |
| P2ry14 | 2.11E-06 | 0.746971 | 0.409 | 0.364 | 0.042472 | 2 L1 _ | P2ry14 |
| Map22 | 1.89E-06 | 0.665839 | 0.23 | 0.145 | 0.038012 | 2 L1 _ | Map2 |
| Pla2r1 | 1.66E-06 | 0.78061 | 0.358 | 0.294 | 0.033437 | 2 L1 _ | Pla2r1 |
| Tmem2731 | 1.66E-06 | 0.666036 | 0.182 | 0.1 | 0.033382 | 2 L1 _ | Tmem273 |
| Hnrnpk | 1.38E-06 | 0.373178 | 0.763 | 0.88 | 0.027712 | 2 L1 _ | Hnrnpk |
| Pmp222 | 1.05E-06 | 0.546019 | 0.664 | 0.713 | 0.021074 | 2 L1 _ | Pmp22 |
| Tmem511 | 9.85E-07 | 0.503044 | 0.161 | 0.081 | 0.019848 | 2 L1 _ | Tmem51 |
| Hopx1 | 9.27E-07 | 0.757695 | 0.365 | 0.29 | 0.01866 | 2 L1 _ | Hopx |
| Fam53b | 9.11E-07 | 0.710305 | 0.358 | 0.291 | 0.018345 | 2 L1 _ | Fam53b |
| Galnt162 | 8.94E-07 | 0.502177 | 0.208 | 0.12 | 0.018013 | 2 L1 _ | Galnt16 |
| Ifitm31 | 8.90E-07 | 0.371157 | 0.964 | 0.936 | 0.01793 | 2 L1 _ | Ifitm3 |
| Prpf4b | 7.80E-07 | 0.529931 | 0.664 | 0.753 | 0.01571 | 2 L1 _ | Prpf4b |
| Prkar1b1 | 7.18E-07 | 0.392635 | 0.117 | 0.048 | 0.014467 | 2 L1 _ | Prkar1b |
| Nr1h3 | 6.55E-07 | 0.700559 | 0.358 | 0.291 | 0.0132 | 2 L1 _ | Nr1h3 |
| Gm26632 | 6.06E-07 | 0.452484 | 0.153 | 0.074 | 0.012211 | 2 L1 _ | Gm26632 |
| Spaar1 | 5.69E-07 | 0.354619 | 0.113 | 0.045 | 0.011463 | 2 L1 _ | Spaar |
| Ncf2 | 5.55E-07 | 0.436608 | 0.131 | 0.057 | 0.011181 | 2 L1 _ | Ncf2 |
| Synpo21 | 5.32E-07 | 0.589888 | 0.223 | 0.13 | 0.010718 | 2 L1 _ | Synpo2 |
| Rabac1 | 5.13E-07 | 0.297231 | 0.821 | 0.912 | 0.010335 | 2 L1 _ | Rabac1 |
| Sorbs11 | 3.92E-07 | 0.553277 | 0.255 | 0.16 | 0.007892 | 2 L1 _ | Sorbs1 |
| 4930523C07Rik2 | 3.60E-07 | 0.762661 | 0.46 | 0.433 | 0.007248 | 2 L1 _ | 4930523C07Rik |
| Pacsin31 | 3.53E-07 | 0.588395 | 0.197 | 0.109 | 0.007106 | 2 L1 _ | Pacsin3 |
| Smpdl3a2 | 3.51E-07 | 0.72406 | 0.485 | 0.48 | 0.00707 | 2 L1 _ | Smpdl3a |
| Gm14964 | 3.08E-07 | 0.450031 | 0.139 | 0.061 | 0.006197 | 2 L1 _ | Gm14964 |
| Arglu1 | 2.84E-07 | 0.52988 | 0.682 | 0.796 | 0.00572 | 2 L1 _ | Arglu1 |
| Ndufa5 | 2.76E-07 | 0.472749 | 0.624 | 0.74 | 0.005559 | 2 L1 _ | Ndufa5 |
| Snhg181 | 2.49E-07 | 0.669305 | 0.613 | 0.627 | 0.005014 | 2 L1 _ | Snhg18 |
| Prrx12 | 2.31E-07 | 0.456914 | 0.745 | 0.715 | 0.004647 | 2 L1 _ | Prrx1 |

| | | | | | | | |
|---|---|---|---|---|---|---|---|
| Avpr1a | 2.29E-07 | 0.655529 | 0.23 | 0.139 | 0.004607 | 2 L1 _ | Avpr1a |
| Bcas3 | 2.08E-07 | 0.793038 | 0.245 | 0.152 | 0.004186 | 2 L1 _ | Bcas3 |
| Il342 | 1.85E-07 | 0.570314 | 0.551 | 0.491 | 0.003719 | 2 L1 _ | Il34 |
| Lpp2 | 1.69E-07 | 0.604896 | 0.591 | 0.636 | 0.003412 | 2 L1 _ | Lpp |
| Hnrnpa2b11 | 1.69E-07 | 0.400189 | 0.781 | 0.894 | 0.003407 | 2 L1 _ | Hnrnpa2b1 |
| Fam107b2 | 1.52E-07 | 0.556414 | 0.19 | 0.098 | 0.003061 | 2 L1 _ | Fam107b |
| Tnfrsf1a | 1.51E-07 | 0.648054 | 0.635 | 0.729 | 0.003044 | 2 L1 _ | Tnfrsf1a |
| Kitl2 | 1.46E-07 | 0.508073 | 0.482 | 0.4 | 0.002934 | 2 L1 _ | Kitl |
| Cnbp | 1.44E-07 | 0.464868 | 0.69 | 0.82 | 0.002891 | 2 L1 _ | Cnbp |
| Mark11 | 1.33E-07 | 0.699514 | 0.296 | 0.206 | 0.002682 | 2 L1 _ | Mark1 |
| Ndufa7 | 1.13E-07 | 0.440648 | 0.712 | 0.845 | 0.002274 | 2 L1 _ | Ndufa7 |
| Atp6v0c1 | 1.12E-07 | 0.411204 | 0.679 | 0.681 | 0.002256 | 2 L1 _ | Atp6v0c |
| Zfhx31 | 9.79E-08 | 0.782048 | 0.354 | 0.268 | 0.001972 | 2 L1 _ | Zfhx3 |
| Sem1 | 9.64E-08 | 0.406685 | 0.766 | 0.898 | 0.001942 | 2 L1 _ | Sem1 |
| Nrarp1 | 8.81E-08 | 0.537099 | 0.201 | 0.105 | 0.001774 | 2 L1 _ | Nrarp |
| Rsu11 | 7.90E-08 | 0.648443 | 0.551 | 0.602 | 0.001592 | 2 L1 _ | Rsu1 |
| Nedd42 | 7.80E-08 | 0.330773 | 0.88 | 0.89 | 0.001571 | 2 L1 _ | Nedd4 |
| Kcne41 | 7.35E-08 | 0.487535 | 0.142 | 0.061 | 0.001481 | 2 L1 _ | Kcne4 |
| Mcam1 | 6.77E-08 | 0.535212 | 0.245 | 0.137 | 0.001363 | 2 L1 _ | Mcam |
| Srrm21 | 6.43E-08 | 0.395779 | 0.818 | 0.902 | 0.001296 | 2 L1 _ | Srrm2 |
| Prkar1a1 | 5.28E-08 | 0.48243 | 0.686 | 0.804 | 0.001064 | 2 L1 _ | Prkar1a |
| Gapdh3 | 5.00E-08 | 0.35873 | 0.825 | 0.919 | 0.001008 | 2 L1 _ | Gapdh |
| Flna1 | 3.65E-08 | 0.543717 | 0.595 | 0.651 | 0.000736 | 2 L1 _ | Flna |
| Atp5mpl1 | 3.57E-08 | 0.502021 | 0.62 | 0.63 | 0.000718 | 2 L1 _ | Atp5mpl |
| Rsrp1 | 3.52E-08 | 0.446995 | 0.81 | 0.903 | 0.000708 | 2 L1 _ | Rsrp1 |
| Snrnp701 | 1.81E-08 | 0.611512 | 0.686 | 0.756 | 0.000365 | 2 L1 _ | Snrnp70 |
| Sema6d2 | 1.67E-08 | 0.689059 | 0.434 | 0.349 | 0.000337 | 2 L1 _ | Sema6d |
| Arntl2 | 1.42E-08 | 0.38543 | 0.12 | 0.044 | 0.000287 | 2 L1 _ | Arntl2 |
| Jag11 | 1.35E-08 | 0.817523 | 0.347 | 0.25 | 0.000272 | 2 L1 _ | Jag1 |
| Ift431 | 1.20E-08 | 0.770904 | 0.416 | 0.343 | 0.000241 | 2 L1 _ | Ift43 |
| Slc11a11 | 1.16E-08 | 0.424028 | 0.106 | 0.034 | 0.000233 | 2 L1 _ | Slc11a1 |
| Tsc22d11 | 9.16E-09 | 0.491482 | 0.807 | 0.839 | 0.000184 | 2 L1 _ | Tsc22d1 |
| Fez22 | 8.36E-09 | 0.69074 | 0.449 | 0.371 | 0.000168 | 2 L1 _ | Fez2 |
| Uqcr111 | 6.48E-09 | 0.447853 | 0.723 | 0.82 | 0.000131 | 2 L1 _ | Uqcr11 |
| Srsf7 | 6.33E-09 | 0.760648 | 0.588 | 0.635 | 0.000127 | 2 L1 _ | Srsf7 |
| Gpx4 | 5.32E-09 | 0.4302 | 0.803 | 0.903 | 0.000107 | 2 L1 _ | Gpx4 |
| B3gnt2 | 5.21E-09 | 0.750585 | 0.442 | 0.372 | 0.000105 | 2 L1 _ | B3gnt2 |
| Jam32 | 4.90E-09 | 0.729727 | 0.471 | 0.398 | 9.87E-05 | 2 L1 _ | Jam3 |
| Pip5k1b1 | 4.57E-09 | 0.393135 | 0.113 | 0.037 | 9.21E-05 | 2 L1 _ | Pip5k1b |
| Calm21 | 3.87E-09 | 0.420526 | 0.81 | 0.915 | 7.79E-05 | 2 L1 _ | Calm2 |
| Dstn1 | 3.67E-09 | 0.555698 | 0.748 | 0.83 | 7.40E-05 | 2 L1 _ | Dstn |

| | | | | | | | |
|---|---|---|---|---|---|---|---|
| Ptma1 | 2.92E-09 | 0.26864 | 0.971 | 0.989 | 5.88E-05 | 2 L1 _ | Ptma |
| Chchd21 | 2.84E-09 | 0.328638 | 0.934 | 0.965 | 5.72E-05 | 2 L1 _ | Chchd2 |
| Jph21 | 2.36E-09 | 0.343355 | 0.106 | 0.032 | 4.75E-05 | 2 L1 _ | Jph2 |
| Pitx11 | 1.50E-09 | 0.836209 | 0.372 | 0.274 | 3.02E-05 | 2 L1 _ | Pitx1 |
| Trpv21 | 1.47E-09 | 0.46836 | 0.161 | 0.065 | 2.97E-05 | 2 L1 _ | Trpv2 |
| Ccnd2 | 1.01E-09 | 0.755837 | 0.679 | 0.717 | 2.04E-05 | 2 L1 _ | Ccnd2 |
| Stard82 | 7.75E-10 | 0.647063 | 0.325 | 0.201 | 1.56E-05 | 2 L1 _ | Stard8 |
| Ggt52 | 6.20E-10 | 0.854538 | 0.442 | 0.353 | 1.25E-05 | 2 L1 _ | Ggt5 |
| Tshz21 | 5.60E-10 | 0.615383 | 0.639 | 0.642 | 1.13E-05 | 2 L1 _ | Tshz2 |
| Ecm21 | 5.24E-10 | 0.803245 | 0.526 | 0.505 | 1.06E-05 | 2 L1 _ | Ecm2 |
| Dleu21 | 4.15E-10 | 0.839057 | 0.369 | 0.255 | 8.36E-06 | 2 L1 _ | Dleu2 |
| Nckap5 | 3.62E-10 | 0.46735 | 0.135 | 0.047 | 7.28E-06 | 2 L1 _ | Nckap5 |
| Procr1 | 2.77E-10 | 0.760901 | 0.23 | 0.114 | 5.57E-06 | 2 L1 _ | Procr |
| Pde1b1 | 1.91E-10 | 0.380682 | 0.131 | 0.043 | 3.85E-06 | 2 L1 _ | Pde1b |
| Atp5b1 | 1.62E-10 | 0.450476 | 0.752 | 0.843 | 3.26E-06 | 2 L1 _ | Atp5b |
| Plcl11 | 1.33E-10 | 0.759206 | 0.255 | 0.137 | 2.68E-06 | 2 L1 _ | Plcl1 |
| Gm15270 | 1.29E-10 | 0.577289 | 0.128 | 0.041 | 2.59E-06 | 2 L1 _ | Gm15270 |
| Dab2ip2 | 1.05E-10 | 0.724797 | 0.536 | 0.492 | 2.12E-06 | 2 L1 _ | Dab2ip |
| Rnd21 | 5.83E-11 | 0.53743 | 0.153 | 0.055 | 1.17E-06 | 2 L1 _ | Rnd2 |
| Dgkg1 | 5.67E-11 | 0.353383 | 0.109 | 0.03 | 1.14E-06 | 2 L1 _ | Dgkg |
| Dmd1 | 2.39E-11 | 0.908186 | 0.365 | 0.243 | 4.82E-07 | 2 L1 _ | Dmd |
| Lbh | 2.05E-11 | 0.809922 | 0.54 | 0.5 | 4.13E-07 | 2 L1 _ | Lbh |
| Wtip1 | 1.87E-11 | 1.065573 | 0.427 | 0.334 | 3.77E-07 | 2 L1 _ | Wtip |
| Cav12 | 1.38E-11 | 0.44128 | 0.478 | 0.29 | 2.77E-07 | 2 L1 _ | Cav1 |
| Ppargc1b1 | 1.38E-11 | 0.506726 | 0.131 | 0.04 | 2.77E-07 | 2 L1 _ | Ppargc1b |
| Pawr1 | 1.18E-11 | 0.689732 | 0.252 | 0.126 | 2.38E-07 | 2 L1 _ | Pawr |
| Afap1l21 | 7.12E-12 | 0.780017 | 0.314 | 0.186 | 1.43E-07 | 2 L1 _ | Afap1l2 |
| Rgs16 | 6.35E-12 | 0.612713 | 0.179 | 0.068 | 1.28E-07 | 2 L1 _ | Rgs16 |
| Fads31 | 5.21E-12 | 0.8069 | 0.325 | 0.194 | 1.05E-07 | 2 L1 _ | Fads3 |
| Id31 | 4.68E-12 | 0.683862 | 0.788 | 0.758 | 9.42E-08 | 2 L1 _ | Id3 |
| Eif1b1 | 2.61E-12 | 0.830355 | 0.599 | 0.615 | 5.26E-08 | 2 L1 _ | Eif1b |
| Utrn2 | 2.61E-12 | 0.916251 | 0.558 | 0.482 | 5.25E-08 | 2 L1 _ | Utrn |
| Mtus11 | 2.25E-12 | 0.870785 | 0.416 | 0.301 | 4.53E-08 | 2 L1 _ | Mtus1 |
| Khdrbs31 | 2.16E-12 | 0.691412 | 0.285 | 0.147 | 4.34E-08 | 2 L1 _ | Khdrbs3 |
| Pakap.1 | 1.85E-12 | 0.760083 | 0.526 | 0.43 | 3.72E-08 | 2 L1 _ | Pakap.1 |
| Ywhaq2 | 1.78E-12 | 0.575087 | 0.726 | 0.807 | 3.58E-08 | 2 L1 _ | Ywhaq |
| Rbm39 | 1.12E-12 | 0.519149 | 0.858 | 0.927 | 2.25E-08 | 2 L1 _ | Rbm39 |
| Myl62 | 8.19E-13 | 0.496459 | 0.923 | 0.973 | 1.65E-08 | 2 L1 _ | Myl6 |
| Arap21 | 6.71E-13 | 0.908249 | 0.485 | 0.391 | 1.35E-08 | 2 L1 _ | Arap2 |
| Nudt4 | 6.47E-13 | 0.858524 | 0.584 | 0.574 | 1.30E-08 | 2 L1 _ | Nudt4 |
| Klf92 | 4.32E-13 | 0.526957 | 0.803 | 0.774 | 8.71E-09 | 2 L1 _ | Klf9 |

| | | | | | | | |
|---|---|---|---|---|---|---|---|
| Trpc31 | 4.30E-13 | 0.531556 | 0.124 | 0.032 | 8.66E-09 | 2 L1 _ | Trpc3 |
| Gucy1a21 | 3.82E-13 | 0.686624 | 0.248 | 0.112 | 7.69E-09 | 2 L1 _ | Gucy1a2 |
| Ccdc1601 | 3.63E-13 | 0.388092 | 0.106 | 0.024 | 7.31E-09 | 2 L1 _ | Ccdc160 |
| Herc11 | 3.62E-13 | 1.222529 | 0.566 | 0.54 | 7.29E-09 | 2 L1 _ | Herc1 |
| Dusp41 | 3.06E-13 | 0.419997 | 0.131 | 0.035 | 6.16E-09 | 2 L1 _ | Dusp4 |
| Sdc11 | 3.02E-13 | 0.906244 | 0.347 | 0.199 | 6.08E-09 | 2 L1 _ | Sdc1 |
| Cox6c1 | 2.45E-13 | 0.408405 | 0.85 | 0.936 | 4.93E-09 | 2 L1 _ | Cox6c |
| Mapt1 | 2.03E-13 | 0.59288 | 0.175 | 0.059 | 4.08E-09 | 2 L1 _ | Mapt |
| Gm106871 | 2.02E-13 | 0.414425 | 0.128 | 0.033 | 4.08E-09 | 2 L1 _ | Gm10687 |
| Cd362 | 1.78E-13 | 0.516038 | 0.361 | 0.184 | 3.58E-09 | 2 L1 _ | Cd36 |
| Uba21 | 1.64E-13 | 0.958164 | 0.5 | 0.425 | 3.31E-09 | 2 L1 _ | Uba2 |
| Cyp4b11 | 9.94E-14 | 0.52691 | 0.15 | 0.044 | 2.00E-09 | 2 L1 _ | Cyp4b1 |
| Cacna1c1 | 9.86E-14 | 0.93018 | 0.296 | 0.154 | 1.99E-09 | 2 L1 _ | Cacna1c |
| Gm134701 | 7.49E-14 | 0.38761 | 0.128 | 0.032 | 1.51E-09 | 2 L1 _ | Gm13470 |
| Hspb12 | 5.72E-14 | 0.712979 | 0.661 | 0.596 | 1.15E-09 | 2 L1 _ | Hspb1 |
| Zfp4671 | 4.63E-14 | 0.710196 | 0.321 | 0.169 | 9.33E-10 | 2 L1 _ | Zfp467 |
| Nt5dc21 | 4.52E-14 | 0.786483 | 0.274 | 0.13 | 9.10E-10 | 2 L1 _ | Nt5dc2 |
| Csnk1e1 | 3.82E-14 | 0.91483 | 0.474 | 0.368 | 7.69E-10 | 2 L1 _ | Csnk1e |
| Prkg11 | 2.88E-14 | 0.84805 | 0.328 | 0.182 | 5.80E-10 | 2 L1 _ | Prkg1 |
| Cox8a2 | 2.63E-14 | 0.447436 | 0.891 | 0.953 | 5.30E-10 | 2 L1 _ | Cox8a |
| Txnip | 2.24E-14 | 0.727736 | 0.803 | 0.862 | 4.52E-10 | 2 L1 _ | Txnip |
| Raph11 | 1.77E-14 | 1.109355 | 0.471 | 0.339 | 3.56E-10 | 2 L1 _ | Raph1 |
| Vcl1 | 1.16E-14 | 0.952749 | 0.544 | 0.472 | 2.35E-10 | 2 L1 _ | Vcl |
| Id41 | 7.07E-15 | 0.989813 | 0.321 | 0.16 | 1.42E-10 | 2 L1 _ | Id4 |
| Tns12 | 5.73E-15 | 0.854083 | 0.595 | 0.526 | 1.15E-10 | 2 L1 _ | Tns1 |
| Selenow1 | 5.68E-15 | 0.575746 | 0.839 | 0.901 | 1.14E-10 | 2 L1 _ | Selenow |
| Timp32 | 4.23E-15 | 0.420651 | 0.88 | 0.78 | 8.51E-11 | 2 L1 _ | Timp3 |
| Elmod11 | 3.61E-15 | 0.419682 | 0.168 | 0.047 | 7.27E-11 | 2 L1 _ | Elmod1 |
| Bok1 | 2.93E-15 | 0.796475 | 0.281 | 0.131 | 5.90E-11 | 2 L1 _ | Bok |
| Rap2a | 2.59E-15 | 1.144548 | 0.467 | 0.352 | 5.22E-11 | 2 L1 _ | Rap2a |
| Pnrc1 | 2.21E-15 | 0.683199 | 0.708 | 0.749 | 4.45E-11 | 2 L1 _ | Pnrc1 |
| E130102H24Rik | 2.07E-15 | 1.030236 | 0.438 | 0.304 | 4.16E-11 | 2 L1 _ | E130102H24Rik |
| Meox21 | 1.63E-15 | 0.737859 | 0.307 | 0.148 | 3.29E-11 | 2 L1 _ | Meox2 |
| Ajuba1 | 1.26E-15 | 0.625794 | 0.237 | 0.09 | 2.53E-11 | 2 L1 _ | Ajuba |
| Sort11 | 9.63E-16 | 0.708869 | 0.252 | 0.102 | 1.94E-11 | 2 L1 _ | Sort1 |
| Pde8b1 | 9.55E-16 | 0.596517 | 0.19 | 0.059 | 1.92E-11 | 2 L1 _ | Pde8b |
| Ptms1 | 6.16E-16 | 0.515333 | 0.894 | 0.907 | 1.24E-11 | 2 L1 _ | Ptms |
| Cd1512 | 4.35E-16 | 0.789427 | 0.613 | 0.554 | 8.75E-12 | 2 L1 _ | Cd151 |
| Ppp1r1a1 | 3.75E-16 | 0.405905 | 0.109 | 0.02 | 7.54E-12 | 2 L1 _ | Ppp1r1a |
| Rcan21 | 3.57E-16 | 0.60763 | 0.175 | 0.051 | 7.18E-12 | 2 L1 _ | Rcan2 |
| Syne22 | 3.48E-16 | 0.897061 | 0.544 | 0.406 | 7.01E-12 | 2 L1 _ | Syne2 |

| | | | | | | | |
|---|---|---|---|---|---|---|---|
| B130024G19Rik | 2.53E-16 | 0.896535 | 0.358 | 0.195 | 5.09E-12 | 2 L1 _ | B130024G19Rik |
| Hmgb1 | 2.52E-16 | 0.445738 | 0.916 | 0.956 | 5.07E-12 | 2 L1 _ | Hmgb1 |
| Pik3r12 | 1.53E-16 | 0.808368 | 0.675 | 0.665 | 3.09E-12 | 2 L1 _ | Pik3r1 |
| Slc25a41 | 1.51E-16 | 0.591745 | 0.818 | 0.866 | 3.05E-12 | 2 L1 _ | Slc25a4 |
| Il2ra1 | 1.51E-16 | 0.633549 | 0.175 | 0.05 | 3.04E-12 | 2 L1 _ | Il2ra |
| Chst12 | 1.03E-16 | 0.786639 | 0.248 | 0.097 | 2.07E-12 | 2 L1 _ | Chst1 |
| Gnb41 | 4.80E-17 | 0.89966 | 0.686 | 0.668 | 9.68E-13 | 2 L1 _ | Gnb4 |
| Laptm4a2 | 3.97E-17 | 0.496806 | 0.956 | 0.922 | 7.99E-13 | 2 L1 _ | Laptm4a |
| mt-Co3 | 3.70E-17 | 0.28163 | 0.993 | 0.993 | 7.44E-13 | 2 L1 _ | mt-Co3 |
| Arhgdib2 | 3.62E-17 | 0.84854 | 0.693 | 0.575 | 7.30E-13 | 2 L1 _ | Arhgdib |
| Htra4 | 3.56E-17 | 0.438492 | 0.157 | 0.04 | 7.17E-13 | 2 L1 _ | Htra4 |
| Sox51 | 3.34E-17 | 0.62144 | 0.201 | 0.064 | 6.74E-13 | 2 L1 _ | Sox5 |
| Mef2a3 | 2.72E-17 | 0.82883 | 0.679 | 0.659 | 5.48E-13 | 2 L1 _ | Mef2a |
| Ech11 | 2.52E-17 | 0.814743 | 0.642 | 0.629 | 5.07E-13 | 2 L1 _ | Ech1 |
| Tbx22 | 6.95E-18 | 1.103158 | 0.442 | 0.277 | 1.40E-13 | 2 L1 _ | Tbx2 |
| H2-M91 | 5.75E-18 | 0.628659 | 0.106 | 0.016 | 1.16E-13 | 2 L1 _ | H2-M9 |
| Slc38a111 | 4.85E-18 | 0.469167 | 0.128 | 0.025 | 9.78E-14 | 2 L1 _ | Slc38a11 |
| Cobll11 | 4.69E-18 | 0.643056 | 0.252 | 0.09 | 9.44E-14 | 2 L1 _ | Cobll1 |
| Ktn12 | 4.43E-18 | 0.929463 | 0.653 | 0.623 | 8.92E-14 | 2 L1 _ | Ktn1 |
| Dgkb1 | 3.02E-18 | 0.511642 | 0.168 | 0.042 | 6.09E-14 | 2 L1 _ | Dgkb |
| Pdlim11 | 1.51E-18 | 0.923779 | 0.423 | 0.24 | 3.03E-14 | 2 L1 _ | Pdlim1 |
| Ptprk1 | 1.43E-18 | 1.006045 | 0.401 | 0.227 | 2.89E-14 | 2 L1 _ | Ptprk |
| Pcdh191 | 1.30E-18 | 0.968464 | 0.573 | 0.449 | 2.63E-14 | 2 L1 _ | Pcdh19 |
| AW011738 | 1.30E-18 | 0.790931 | 0.263 | 0.099 | 2.61E-14 | 2 L1 _ | AW011738 |
| Angpt21 | 1.19E-18 | 0.700509 | 0.223 | 0.073 | 2.39E-14 | 2 L1 _ | Angpt2 |
| Fxyd12 | 5.41E-19 | 0.75688 | 0.723 | 0.673 | 1.09E-14 | 2 L1 _ | Fxyd1 |
| Rgs61 | 4.57E-19 | 0.53524 | 0.142 | 0.029 | 9.21E-15 | 2 L1 _ | Rgs6 |
| Rock11 | 2.89E-19 | 0.801517 | 0.723 | 0.741 | 5.81E-15 | 2 L1 _ | Rock1 |
| Scn3a1 | 1.47E-19 | 0.576919 | 0.142 | 0.029 | 2.96E-15 | 2 L1 _ | Scn3a |
| Tmem1003 | 1.00E-19 | 1.033748 | 0.456 | 0.264 | 2.02E-15 | 2 L1 _ | Tmem100 |
| Myom11 | 7.06E-20 | 0.47088 | 0.142 | 0.028 | 1.42E-15 | 2 L1 _ | Myom1 |
| Tbx32 | 5.13E-20 | 0.819539 | 0.31 | 0.127 | 1.03E-15 | 2 L1 _ | Tbx3 |
| Plxdc11 | 2.80E-20 | 0.490338 | 0.135 | 0.025 | 5.63E-16 | 2 L1 _ | Plxdc1 |
| Emid11 | 1.41E-20 | 0.952583 | 0.416 | 0.218 | 2.83E-16 | 2 L1 _ | Emid1 |
| Ace21 | 1.40E-20 | 0.769059 | 0.226 | 0.064 | 2.83E-16 | 2 L1 _ | Ace2 |
| Cspg41 | 1.18E-20 | 1.040449 | 0.5 | 0.314 | 2.38E-16 | 2 L1 _ | Cspg4 |
| Nes2 | 1.09E-20 | 0.981199 | 0.412 | 0.209 | 2.19E-16 | 2 L1 _ | Nes |
| Itga12 | 8.23E-21 | 0.632867 | 0.832 | 0.796 | 1.66E-16 | 2 L1 _ | Itga1 |
| Trpc11 | 6.80E-21 | 0.815891 | 0.274 | 0.1 | 1.37E-16 | 2 L1 _ | Trpc1 |
| Ndrg21 | 6.65E-21 | 0.982256 | 0.595 | 0.479 | 1.34E-16 | 2 L1 _ | Ndrg2 |
| Hey21 | 5.25E-21 | 0.436459 | 0.131 | 0.022 | 1.06E-16 | 2 L1 _ | Hey2 |

| | | | | | | | |
|---|---|---|---|---|---|---|---|
| Arhgap101 | 4.06E-21 | 0.957633 | 0.526 | 0.369 | 8.18E-17 | 2 L1 _ | Arhgap10 |
| Bcam2 | 2.80E-21 | 0.791906 | 0.54 | 0.304 | 5.63E-17 | 2 L1 _ | Bcam |
| Trmt9b | 2.60E-21 | 0.474554 | 0.124 | 0.019 | 5.24E-17 | 2 L1 _ | Trmt9b |
| Tagln1 | 2.49E-21 | 1.441774 | 0.391 | 0.182 | 5.01E-17 | 2 L1 _ | Tagln |
| Heyl1 | 2.45E-21 | 1.05867 | 0.511 | 0.331 | 4.94E-17 | 2 L1 _ | Heyl |
| Olfml2b2 | 8.18E-22 | 0.855638 | 0.296 | 0.11 | 1.65E-17 | 2 L1 _ | Olfml2b |
| Rdh51 | 7.94E-22 | 0.816367 | 0.266 | 0.089 | 1.60E-17 | 2 L1 _ | Rdh5 |
| Aoc32 | 7.19E-22 | 0.837052 | 0.391 | 0.178 | 1.45E-17 | 2 L1 _ | Aoc3 |
| Son | 6.03E-22 | 0.616459 | 0.945 | 0.95 | 1.21E-17 | 2 L1 _ | Son |
| Atpif11 | 2.64E-22 | 0.807477 | 0.745 | 0.782 | 5.31E-18 | 2 L1 _ | Atpif1 |
| Arhgef171 | 1.81E-22 | 1.101876 | 0.474 | 0.284 | 3.65E-18 | 2 L1 _ | Arhgef17 |
| Ddx51 | 1.71E-22 | 0.574071 | 0.956 | 0.967 | 3.45E-18 | 2 L1 _ | Ddx5 |
| Mrvi11 | 1.17E-22 | 0.978151 | 0.412 | 0.207 | 2.36E-18 | 2 L1 _ | Mrvi1 |
| Nrip21 | 3.93E-23 | 0.660657 | 0.164 | 0.032 | 7.92E-19 | 2 L1 _ | Nrip2 |
| Cbfa2t31 | 3.31E-23 | 0.975574 | 0.449 | 0.225 | 6.66E-19 | 2 L1 _ | Cbfa2t3 |
| Hrct11 | 2.00E-23 | 0.896715 | 0.339 | 0.135 | 4.02E-19 | 2 L1 _ | Hrct1 |
| Tesc1 | 1.92E-23 | 1.090247 | 0.259 | 0.078 | 3.87E-19 | 2 L1 _ | Tesc |
| Snrk2 | 1.38E-23 | 0.872191 | 0.723 | 0.595 | 2.77E-19 | 2 L1 _ | Snrk |
| Pcp4l12 | 1.17E-23 | 0.972626 | 0.438 | 0.198 | 2.35E-19 | 2 L1 _ | Pcp4l1 |
| Rcsd11 | 9.26E-24 | 1.041559 | 0.412 | 0.201 | 1.87E-19 | 2 L1 _ | Rcsd1 |
| Cdh21 | 4.10E-24 | 0.715141 | 0.172 | 0.034 | 8.25E-20 | 2 L1 _ | Cdh2 |
| Hrc1 | 1.36E-24 | 0.493071 | 0.128 | 0.017 | 2.74E-20 | 2 L1 _ | Hrc |
| B430010I23Rik | 1.08E-24 | 0.933829 | 0.237 | 0.065 | 2.17E-20 | 2 L1 _ | B430010I23Rik |
| Pde1a1 | 9.51E-25 | 1.025852 | 0.372 | 0.163 | 1.92E-20 | 2 L1 _ | Pde1a |
| Slc4a81 | 5.35E-25 | 0.662656 | 0.212 | 0.049 | 1.08E-20 | 2 L1 _ | Slc4a8 |
| Dtx31 | 4.79E-25 | 1.094827 | 0.58 | 0.423 | 9.65E-21 | 2 L1 _ | Dtx3 |
| Map7d21 | 2.18E-25 | 0.506028 | 0.15 | 0.023 | 4.39E-21 | 2 L1 _ | Map7d2 |
| Pakap | 1.85E-25 | 0.80016 | 0.248 | 0.07 | 3.73E-21 | 2 L1 _ | Pakap |
| Map3k201 | 1.78E-25 | 1.081079 | 0.489 | 0.28 | 3.58E-21 | 2 L1 _ | Map3k20 |
| Cog71 | 1.54E-25 | 1.078325 | 0.394 | 0.177 | 3.11E-21 | 2 L1 _ | Cog7 |
| Adra2a | 7.87E-26 | 0.504145 | 0.113 | 0.012 | 1.58E-21 | 2 L1 _ | Adra2a |
| Mustn12 | 5.73E-26 | 1.210983 | 0.299 | 0.098 | 1.15E-21 | 2 L1 _ | Mustn1 |
| Tspan152 | 3.88E-26 | 0.920202 | 0.372 | 0.146 | 7.81E-22 | 2 L1 _ | Tspan15 |
| Gprc5c1 | 2.76E-26 | 0.975979 | 0.369 | 0.152 | 5.56E-22 | 2 L1 _ | Gprc5c |
| Arhgap421 | 2.12E-26 | 1.051122 | 0.401 | 0.177 | 4.28E-22 | 2 L1 _ | Arhgap42 |
| Map1lc3a1 | 2.01E-26 | 0.890956 | 0.774 | 0.747 | 4.05E-22 | 2 L1 _ | Map1lc3a |
| 1500009L16Rik | 1.35E-26 | 0.695705 | 0.223 | 0.053 | 2.72E-22 | 2 L1 _ | 1500009L16Rik |
| Ppp1r14a1 | 1.13E-26 | 0.735171 | 0.354 | 0.121 | 2.28E-22 | 2 L1 _ | Ppp1r14a |
| Tbc1d11 | 7.61E-27 | 1.105962 | 0.507 | 0.288 | 1.53E-22 | 2 L1 _ | Tbc1d1 |
| Gamt1 | 6.32E-27 | 1.143847 | 0.46 | 0.246 | 1.27E-22 | 2 L1 _ | Gamt |
| Ccdc681 | 3.91E-27 | 0.610562 | 0.175 | 0.031 | 7.88E-23 | 2 L1 _ | Ccdc68 |

| | | | | | | | |
|---|---|---|---|---|---|---|---|
| Kcnk31 | 2.80E-27 | 0.780693 | 0.204 | 0.043 | 5.63E-23 | 2 L1 _ | Kcnk3 |
| Vstm41 | 1.12E-27 | 1.238885 | 0.511 | 0.296 | 2.25E-23 | 2 L1 _ | Vstm4 |
| Ednra1 | 6.96E-28 | 1.275125 | 0.606 | 0.435 | 1.40E-23 | 2 L1 _ | Ednra |
| Sema5a | 5.87E-28 | 1.17345 | 0.474 | 0.249 | 1.18E-23 | 2 L1 _ | Sema5a |
| Klhl231 | 5.38E-28 | 0.960624 | 0.299 | 0.093 | 1.08E-23 | 2 L1 _ | Klhl23 |
| Epas12 | 2.85E-28 | 0.841806 | 0.726 | 0.504 | 5.74E-24 | 2 L1 _ | Epas1 |
| Slco3a11 | 2.75E-28 | 1.123747 | 0.526 | 0.296 | 5.55E-24 | 2 L1 _ | Slco3a1 |
| Enpep1 | 2.10E-28 | 1.114747 | 0.358 | 0.135 | 4.23E-24 | 2 L1 _ | Enpep |
| Mef2c2 | 1.21E-28 | 0.787411 | 0.792 | 0.633 | 2.43E-24 | 2 L1 _ | Mef2c |
| Pdgfa1 | 1.08E-28 | 1.053958 | 0.474 | 0.228 | 2.18E-24 | 2 L1 _ | Pdgfa |
| Crip22 | 3.44E-29 | 0.860588 | 0.839 | 0.763 | 6.93E-25 | 2 L1 _ | Crip2 |
| Mgst31 | 1.02E-29 | 1.104328 | 0.445 | 0.207 | 2.06E-25 | 2 L1 _ | Mgst3 |
| Esam2 | 7.61E-30 | 0.77294 | 0.631 | 0.297 | 1.53E-25 | 2 L1 _ | Esam |
| Casq21 | 5.17E-30 | 0.940362 | 0.212 | 0.041 | 1.04E-25 | 2 L1 _ | Casq2 |
| Ubb1 | 2.45E-30 | 0.510962 | 0.971 | 0.98 | 4.94E-26 | 2 L1 _ | Ubb |
| Limd11 | 1.92E-30 | 1.227917 | 0.591 | 0.378 | 3.86E-26 | 2 L1 _ | Limd1 |
| Adcy61 | 8.65E-31 | 1.204447 | 0.427 | 0.185 | 1.74E-26 | 2 L1 _ | Adcy6 |
| Acta21 | 7.55E-31 | 2.047854 | 0.453 | 0.198 | 1.52E-26 | 2 L1 _ | Acta2 |
| Ebf22 | 4.36E-31 | 1.02455 | 0.401 | 0.15 | 8.79E-27 | 2 L1 _ | Ebf2 |
| Chn11 | 2.53E-31 | 0.805592 | 0.255 | 0.059 | 5.10E-27 | 2 L1 _ | Chn1 |
| Lin7a1 | 4.73E-32 | 1.29611 | 0.529 | 0.288 | 9.52E-28 | 2 L1 _ | Lin7a |
| Fzd42 | 2.77E-32 | 1.04994 | 0.595 | 0.316 | 5.57E-28 | 2 L1 _ | Fzd4 |
| Stk38l1 | 2.22E-33 | 0.977675 | 0.35 | 0.113 | 4.46E-29 | 2 L1 _ | Stk38l |
| Agtr1a1 | 1.05E-33 | 1.013423 | 0.328 | 0.092 | 2.12E-29 | 2 L1 _ | Agtr1a |
| Ccdc1411 | 1.00E-33 | 0.941305 | 0.288 | 0.072 | 2.02E-29 | 2 L1 _ | Ccdc141 |
| Tnfrsf21 | 8.31E-34 | 1.287228 | 0.529 | 0.281 | 1.67E-29 | 2 L1 _ | Tnfrsf21 |
| Rrad | 7.75E-35 | 1.024502 | 0.328 | 0.094 | 1.56E-30 | 2 L1 _ | Rrad |
| Pten1 | 7.03E-35 | 1.074515 | 0.788 | 0.719 | 1.42E-30 | 2 L1 _ | Pten |
| Ppp1r12a1 | 5.70E-35 | 1.186444 | 0.73 | 0.635 | 1.15E-30 | 2 L1 _ | Ppp1r12a |
| Npy1r1 | 4.04E-35 | 1.019429 | 0.332 | 0.092 | 8.14E-31 | 2 L1 _ | Npy1r |
| Sgip11 | 2.25E-35 | 0.899916 | 0.281 | 0.065 | 4.53E-31 | 2 L1 _ | Sgip1 |
| Tbx3os11 | 5.04E-36 | 0.735019 | 0.219 | 0.036 | 1.01E-31 | 2 L1 _ | Tbx3os1 |
| Mylk1 | 3.86E-36 | 1.021842 | 0.788 | 0.619 | 7.77E-32 | 2 L1 _ | Mylk |
| Daam21 | 1.15E-36 | 1.06811 | 0.398 | 0.13 | 2.31E-32 | 2 L1 _ | Daam2 |
| Arhgef161 | 1.13E-36 | 0.567736 | 0.168 | 0.018 | 2.28E-32 | 2 L1 _ | Arhgef16 |
| Btbd31 | 9.46E-37 | 1.297004 | 0.533 | 0.262 | 1.90E-32 | 2 L1 _ | Btbd3 |
| Adap2os1 | 6.35E-37 | 1.082621 | 0.405 | 0.133 | 1.28E-32 | 2 L1 _ | Adap2os |
| Rbpms21 | 3.83E-37 | 1.084199 | 0.31 | 0.077 | 7.71E-33 | 2 L1 _ | Rbpms2 |
| Dlc12 | 3.76E-37 | 1.139478 | 0.792 | 0.694 | 7.56E-33 | 2 L1 _ | Dlc1 |
| Pdlim31 | 2.85E-37 | 0.748636 | 0.237 | 0.042 | 5.74E-33 | 2 L1 _ | Pdlim3 |
| Cst31 | 1.61E-37 | 0.521758 | 0.982 | 0.95 | 3.24E-33 | 2 L1 _ | Cst3 |

| | | | | | | | |
|---|---|---|---|---|---|---|---|
| Fam162a1 | 1.22E-37 | 1.335555 | 0.73 | 0.6 | 2.45E-33 | 2 L1 _ | Fam162a |
| Mprip1 | 4.44E-38 | 1.117017 | 0.777 | 0.71 | 8.94E-34 | 2 L1 _ | Mprip |
| Adap21 | 4.59E-39 | 1.300829 | 0.507 | 0.218 | 9.25E-35 | 2 L1 _ | Adap2 |
| Cryab1 | 4.21E-39 | 1.487451 | 0.69 | 0.472 | 8.49E-35 | 2 L1 _ | Cryab |
| Gcnt22 | 3.76E-39 | 1.213208 | 0.464 | 0.172 | 7.57E-35 | 2 L1 _ | Gcnt2 |
| Fry2 | 3.43E-39 | 1.245781 | 0.445 | 0.158 | 6.91E-35 | 2 L1 _ | Fry |
| Trpc61 | 4.00E-40 | 0.840194 | 0.248 | 0.043 | 8.06E-36 | 2 L1 _ | Trpc6 |
| Inka2 | 1.67E-40 | 0.700112 | 0.186 | 0.021 | 3.36E-36 | 2 L1 _ | Inka2 |
| Cpm1 | 9.07E-41 | 0.947509 | 0.354 | 0.084 | 1.83E-36 | 2 L1 _ | Cpm |
| Slc2a41 | 1.97E-41 | 0.872719 | 0.31 | 0.067 | 3.96E-37 | 2 L1 _ | Slc2a4 |
| Aspn1 | 1.12E-41 | 1.327849 | 0.737 | 0.492 | 2.26E-37 | 2 L1 _ | Aspn |
| Cpe1 | 4.76E-42 | 1.219135 | 0.69 | 0.415 | 9.59E-38 | 2 L1 _ | Cpe |
| Pde3a1 | 4.29E-43 | 1.081932 | 0.409 | 0.117 | 8.63E-39 | 2 L1 _ | Pde3a |
| Crip11 | 3.13E-43 | 1.093175 | 0.956 | 0.95 | 6.31E-39 | 2 L1 _ | Crip1 |
| Parm11 | 1.71E-43 | 1.130064 | 0.365 | 0.094 | 3.45E-39 | 2 L1 _ | Parm1 |
| Nr2f21 | 2.60E-44 | 1.18306 | 0.821 | 0.664 | 5.23E-40 | 2 L1 _ | Nr2f2 |
| Atp1b21 | 1.28E-44 | 1.372569 | 0.573 | 0.259 | 2.59E-40 | 2 L1 _ | Atp1b2 |
| Asap21 | 8.87E-45 | 1.400089 | 0.551 | 0.251 | 1.79E-40 | 2 L1 _ | Asap2 |
| Mnd11 | 3.71E-45 | 1.096717 | 0.383 | 0.096 | 7.47E-41 | 2 L1 _ | Mnd1 |
| Colec111 | 1.82E-46 | 1.123533 | 0.252 | 0.036 | 3.67E-42 | 2 L1 _ | Colec11 |
| Meg31 | 6.02E-47 | 1.484149 | 0.858 | 0.623 | 1.21E-42 | 2 L1 _ | Meg3 |
| Inpp4b2 | 2.69E-47 | 1.316196 | 0.595 | 0.247 | 5.42E-43 | 2 L1 _ | Inpp4b |
| Sep-71 | 1.11E-47 | 0.980769 | 0.894 | 0.901 | 2.24E-43 | 2 L1 _ | 7-Sep |
| Rarres22 | 2.52E-48 | 1.113634 | 0.869 | 0.667 | 5.08E-44 | 2 L1 _ | Rarres2 |
| Hspb21 | 1.61E-48 | 1.030651 | 0.35 | 0.076 | 3.25E-44 | 2 L1 _ | Hspb2 |
| Serpine21 | 1.27E-48 | 1.41022 | 0.788 | 0.509 | 2.56E-44 | 2 L1 _ | Serpine2 |
| Gm138611 | 5.37E-49 | 0.853735 | 0.255 | 0.034 | 1.08E-44 | 2 L1 _ | Gm13861 |
| Foxs11 | 1.63E-49 | 1.422809 | 0.672 | 0.376 | 3.28E-45 | 2 L1 _ | Foxs1 |
| Tm4sf12 | 1.05E-50 | 1.247917 | 0.876 | 0.672 | 2.11E-46 | 2 L1 _ | Tm4sf1 |
| Myh111 | 1.77E-51 | 1.351531 | 0.387 | 0.082 | 3.56E-47 | 2 L1 _ | Myh11 |
| H3f3b2 | 4.16E-53 | 0.793504 | 0.978 | 0.984 | 8.38E-49 | 2 L1 _ | H3f3b |
| Ephx31 | 1.21E-54 | 1.148026 | 0.347 | 0.061 | 2.44E-50 | 2 L1 _ | Ephx3 |
| Filip1l1 | 1.08E-56 | 1.60921 | 0.639 | 0.268 | 2.17E-52 | 2 L1 _ | Filip1l |
| Lgi11 | 5.03E-57 | 1.21579 | 0.398 | 0.078 | 1.01E-52 | 2 L1 _ | Lgi1 |
| Fermt21 | 1.11E-57 | 1.175775 | 0.891 | 0.76 | 2.24E-53 | 2 L1 _ | Fermt2 |
| Stac1 | 4.11E-58 | 0.899385 | 0.255 | 0.027 | 8.28E-54 | 2 L1 _ | Stac |
| Dbndd2 | 2.78E-58 | 1.497064 | 0.591 | 0.239 | 5.61E-54 | 2 L1 _ | Dbndd2 |
| Trarg1 | 5.07E-60 | 0.6519 | 0.19 | 0.01 | 1.02E-55 | 2 L1 _ | Trarg1 |
| Rbpms1 | 4.04E-61 | 1.463106 | 0.839 | 0.704 | 8.14E-57 | 2 L1 _ | Rbpms |
| Gjc11 | 2.52E-62 | 1.658049 | 0.737 | 0.41 | 5.08E-58 | 2 L1 _ | Gjc1 |
| Cald11 | 7.67E-63 | 1.136737 | 0.964 | 0.862 | 1.55E-58 | 2 L1 _ | Cald1 |

| | | | | | | | |
|---|---|---|---|---|---|---|---|
| Rasgrp21 | 1.58E-63 | 1.318788 | 0.62 | 0.21 | 3.19E-59 | 2 L1 _ | Rasgrp2 |
| Art31 | 7.93E-64 | 1.421863 | 0.474 | 0.107 | 1.60E-59 | 2 L1 _ | Art3 |
| Lmod11 | 3.86E-66 | 1.184108 | 0.383 | 0.062 | 7.77E-62 | 2 L1 _ | Lmod1 |
| Tpm11 | 3.96E-69 | 1.821895 | 0.872 | 0.703 | 7.97E-65 | 2 L1 _ | Tpm1 |
| Gng112 | 4.34E-70 | 1.371329 | 0.931 | 0.806 | 8.74E-66 | 2 L1 _ | Gng11 |
| Tpm21 | 6.18E-71 | 1.419255 | 0.77 | 0.357 | 1.25E-66 | 2 L1 _ | Tpm2 |
| Ano11 | 1.07E-71 | 1.253552 | 0.401 | 0.063 | 2.15E-67 | 2 L1 _ | Ano1 |
| Cystm11 | 1.58E-73 | 1.554703 | 0.697 | 0.27 | 3.17E-69 | 2 L1 _ | Cystm1 |
| Pde5a1 | 2.53E-75 | 1.537095 | 0.602 | 0.17 | 5.10E-71 | 2 L1 _ | Pde5a |
| Phlda11 | 1.39E-75 | 2.006173 | 0.755 | 0.376 | 2.81E-71 | 2 L1 _ | Phlda1 |
| Slc12a21 | 6.04E-76 | 1.687003 | 0.679 | 0.226 | 1.22E-71 | 2 L1 _ | Slc12a2 |
| Serpini11 | 7.07E-78 | 1.477489 | 0.526 | 0.11 | 1.42E-73 | 2 L1 _ | Serpini1 |
| Zeb21 | 5.77E-80 | 1.816584 | 0.898 | 0.675 | 1.16E-75 | 2 L1 _ | Zeb2 |
| Ifitm11 | 4.09E-80 | 2.320677 | 0.599 | 0.169 | 8.24E-76 | 2 L1 _ | Ifitm1 |
| Ntm1 | 5.68E-81 | 1.126883 | 0.31 | 0.026 | 1.14E-76 | 2 L1 _ | Ntm |
| Ddit4l1 | 4.13E-83 | 1.317621 | 0.383 | 0.047 | 8.32E-79 | 2 L1 _ | Ddit4l |
| Rem11 | 1.31E-84 | 1.660565 | 0.547 | 0.119 | 2.64E-80 | 2 L1 _ | Rem1 |
| Mfge81 | 2.21E-86 | 1.594787 | 0.953 | 0.784 | 4.46E-82 | 2 L1 _ | Mfge8 |
| Rasl11a1 | 5.94E-87 | 1.705126 | 0.642 | 0.178 | 1.20E-82 | 2 L1 _ | Rasl11a |
| Des1 | 1.74E-89 | 1.740653 | 0.657 | 0.179 | 3.51E-85 | 2 L1 _ | Des |
| Sparcl1 | 1.19E-91 | 1.41712 | 0.978 | 0.861 | 2.39E-87 | 2 L1 _ | Sparcl1 |
| Sep-41 | 9.14E-92 | 1.82456 | 0.912 | 0.693 | 1.84E-87 | 2 L1 _ | 4-Sep |
| Steap42 | 1.65E-92 | 1.909487 | 0.861 | 0.397 | 3.32E-88 | 2 L1 _ | Steap4 |
| Itga71 | 3.47E-93 | 1.541433 | 0.511 | 0.085 | 6.98E-89 | 2 L1 _ | Itga7 |
| Ebf11 | 4.95E-95 | 1.803943 | 0.967 | 0.767 | 9.98E-91 | 2 L1 _ | Ebf1 |
| Malat11 | 5.82E-101 | 1.275969 | 1 | 0.997 | 1.17E-96 | 2 L1 _ | Malat1 |
| Ednrb1 | 3.47E-101 | 1.6892 | 0.836 | 0.248 | 7.00E-97 | 2 L1 _ | Ednrb |
| Abcc91 | 3.44E-101 | 1.974554 | 0.81 | 0.272 | 6.93E-97 | 2 L1 _ | Abcc9 |
| Myo1b1 | 1.59E-101 | 2.077292 | 0.872 | 0.452 | 3.20E-97 | 2 L1 _ | Myo1b |
| Ndufa4l21 | 7.72E-106 | 2.143224 | 0.942 | 0.578 | 1.55E-101 | 2 L1 _ | Ndufa4l2 |
| Sncg1 | 8.80E-107 | 2.044359 | 0.704 | 0.166 | 1.77E-102 | 2 L1 _ | Sncg |
| Gm138891 | 2.95E-108 | 2.48647 | 0.905 | 0.446 | 5.95E-104 | 2 L1 _ | Gm13889 |
| Pdgfrb1 | 8.74E-110 | 1.821479 | 0.996 | 0.814 | 1.76E-105 | 2 L1 _ | Pdgfrb |
| Vtn1 | 2.83E-112 | 2.163241 | 0.701 | 0.164 | 5.70E-108 | 2 L1 _ | Vtn |
| Rasl121 | 1.08E-113 | 1.536218 | 0.54 | 0.074 | 2.17E-109 | 2 L1 _ | Rasl12 |
| Myl91 | 5.62E-118 | 2.319601 | 0.88 | 0.307 | 1.13E-113 | 2 L1 _ | Myl9 |
| Tinagl12 | 1.21E-122 | 1.950837 | 0.865 | 0.236 | 2.44E-118 | 2 L1 _ | Tinagl1 |
| Rgs41 | 6.15E-123 | 1.861772 | 0.624 | 0.093 | 1.24E-118 | 2 L1 _ | Rgs4 |
| Gucy1b11 | 5.61E-125 | 2.274866 | 0.865 | 0.3 | 1.13E-120 | 2 L1 _ | Gucy1b1 |
| Cox4i21 | 8.46E-129 | 1.69644 | 0.69 | 0.107 | 1.70E-124 | 2 L1 _ | Cox4i2 |
| Gja41 | 9.46E-134 | 2.210867 | 0.803 | 0.182 | 1.91E-129 | 2 L1 _ | Gja4 |

| | | | | | | | |
|---|---|---|---|---|---|---|---|
| Kcnj81 | 9.28E-141 | 2.518904 | 0.814 | 0.177 | 1.87E-136 | 2 L1 _ | Kcnj8 |
| Olfr5581 | 2.09E-146 | 1.92119 | 0.635 | 0.077 | 4.20E-142 | 2 L1 _ | Olfr558 |
| Carmn1 | 1.61E-146 | 2.219002 | 0.748 | 0.139 | 3.25E-142 | 2 L1 _ | Carmn |
| Gucy1a11 | 3.94E-148 | 2.502234 | 0.938 | 0.312 | 7.93E-144 | 2 L1 _ | Gucy1a1 |
| Notch31 | 4.76E-159 | 2.18987 | 0.865 | 0.17 | 9.59E-155 | 2 L1 _ | Notch3 |
| Higd1b1 | 1.89E-194 | 2.417761 | 0.85 | 0.108 | 3.80E-190 | 2 L1 _ | Higd1b |
| Rgs51 | 6.84E-221 | 3.184942 | 0.993 | 0.162 | 1.38E-216 | 2 L1 _ | Rgs5 |
| St5 | 2.50E-06 | 0.468634 | 0.445 | 0.308 | 0.050281 | 2 M1 _ | St5 |
| Nptn | 2.36E-06 | 0.543915 | 0.68 | 0.601 | 0.047626 | 2 M1 _ | Nptn |
| Sipa1 | 2.27E-06 | 0.514971 | 0.539 | 0.425 | 0.045703 | 2 M1 _ | Sipa1 |
| Uqcrq | 2.11E-06 | 0.400118 | 0.836 | 0.77 | 0.042407 | 2 M1 _ | Uqcrq |
| Rhoj1 | 1.81E-06 | 0.379553 | 0.852 | 0.736 | 0.036503 | 2 M1 _ | Rhoj |
| Cox7a2 | 1.73E-06 | 0.385774 | 0.875 | 0.852 | 0.034774 | 2 M1 _ | Cox7a2 |
| Nbr1 | 1.62E-06 | 0.539349 | 0.461 | 0.333 | 0.032656 | 2 M1 _ | Nbr1 |
| Armcx3 | 1.54E-06 | 0.621644 | 0.492 | 0.368 | 0.031023 | 2 M1 _ | Armcx3 |
| Smtn | 1.51E-06 | 0.641734 | 0.43 | 0.306 | 0.030384 | 2 M1 _ | Smtn |
| Ndufs7 | 1.38E-06 | 0.389612 | 0.734 | 0.681 | 0.027823 | 2 M1 _ | Ndufs7 |
| Fam110b | 1.35E-06 | 0.351871 | 0.188 | 0.075 | 0.027165 | 2 M1 _ | Fam110b |
| 9930111J21Rik2 | 1.22E-06 | 0.574695 | 0.562 | 0.435 | 0.024533 | 2 M1 _ | 9930111J21Rik2 |
| Mphosph6 | 1.12E-06 | 0.553222 | 0.352 | 0.207 | 0.022579 | 2 M1 _ | Mphosph6 |
| Maged21 | 1.04E-06 | 0.636805 | 0.555 | 0.43 | 0.020994 | 2 M1 _ | Maged2 |
| Limk1 | 9.86E-07 | 0.543677 | 0.281 | 0.146 | 0.019851 | 2 M1 _ | Limk1 |
| Taf6l1 | 9.75E-07 | 0.368524 | 0.203 | 0.085 | 0.019628 | 2 M1 _ | Taf6l |
| Maml3 | 9.66E-07 | 0.547692 | 0.438 | 0.292 | 0.019455 | 2 M1 _ | Maml3 |
| Sytl2 | 9.41E-07 | 0.343301 | 0.188 | 0.072 | 0.018957 | 2 M1 _ | Sytl2 |
| Bcl9l | 9.40E-07 | 0.491532 | 0.453 | 0.307 | 0.018939 | 2 M1 _ | Bcl9l |
| Ndufc1 | 8.95E-07 | 0.443289 | 0.75 | 0.707 | 0.018021 | 2 M1 _ | Ndufc1 |
| C2cd2 | 8.66E-07 | 0.501761 | 0.344 | 0.194 | 0.01744 | 2 M1 _ | C2cd2 |
| Olfr1033 | 8.64E-07 | 0.409125 | 0.188 | 0.075 | 0.017409 | 2 M1 _ | Olfr1033 |
| Arhgap26 | 8.57E-07 | 0.265992 | 0.18 | 0.067 | 0.017263 | 2 M1 _ | Arhgap26 |
| Ttll7 | 7.97E-07 | 0.372612 | 0.219 | 0.091 | 0.016051 | 2 M1 _ | Ttll7 |
| Cox7c | 7.76E-07 | 0.321721 | 0.945 | 0.908 | 0.015619 | 2 M1 _ | Cox7c |
| Pygm | 7.30E-07 | 0.435649 | 0.305 | 0.157 | 0.01471 | 2 M1 _ | Pygm |
| Pcdh18 | 7.29E-07 | 0.577001 | 0.586 | 0.44 | 0.014679 | 2 M1 _ | Pcdh18 |
| Nr2f11 | 7.15E-07 | 0.497689 | 0.734 | 0.593 | 0.014407 | 2 M1 _ | Nr2f1 |
| Timm17a | 6.54E-07 | 0.490886 | 0.523 | 0.392 | 0.013164 | 2 M1 _ | Timm17a |
| Plxna1 | 6.19E-07 | 0.381158 | 0.352 | 0.195 | 0.012466 | 2 M1 _ | Plxna1 |
| Phlda3 | 6.19E-07 | 0.426136 | 0.719 | 0.547 | 0.012458 | 2 M1 _ | Phlda3 |
| Dynlt3 | 6.17E-07 | 0.55347 | 0.633 | 0.526 | 0.012431 | 2 M1 _ | Dynlt3 |
| Soga1 | 5.95E-07 | 0.427698 | 0.211 | 0.09 | 0.011987 | 2 M1 _ | Soga1 |
| Rsbn1l | 5.28E-07 | 0.456038 | 0.617 | 0.492 | 0.01064 | 2 M1 _ | Rsbn1l |

| | | | | | | | |
|---|---|---|---|---|---|---|---|
| Cnnm2 | 5.08E-07 | 0.352802 | 0.18 | 0.067 | 0.010233 | 2 M1 _ | Cnnm2 |
| Ybey | 4.84E-07 | 0.293224 | 0.133 | 0.04 | 0.009752 | 2 M1 _ | Ybey |
| Gm12840 | 4.79E-07 | 0.752991 | 0.18 | 0.069 | 0.009656 | 2 M1 _ | Gm12840 |
| Phtf2 | 4.67E-07 | 0.335928 | 0.188 | 0.072 | 0.009407 | 2 M1 _ | Phtf2 |
| Fxyd2 | 4.54E-07 | 0.583645 | 0.375 | 0.216 | 0.009151 | 2 M1 _ | Fxyd2 |
| Specc1 | 3.94E-07 | 0.446996 | 0.328 | 0.174 | 0.007941 | 2 M1 _ | Specc1 |
| Rheb | 3.54E-07 | 0.52087 | 0.711 | 0.639 | 0.007132 | 2 M1 _ | Rheb |
| Slc25a3 | 3.44E-07 | 0.475373 | 0.867 | 0.84 | 0.006933 | 2 M1 _ | Slc25a3 |
| Camk1 | 3.39E-07 | 0.516367 | 0.586 | 0.467 | 0.006829 | 2 M1 _ | Camk1 |
| Mark1 | 3.29E-07 | 0.522849 | 0.359 | 0.208 | 0.006633 | 2 M1 _ | Mark1 |
| Cyp4b1 | 2.88E-07 | 0.331861 | 0.156 | 0.05 | 0.00581 | 2 M1 _ | Cyp4b1 |
| Trp53inp2 | 2.88E-07 | 0.544137 | 0.312 | 0.167 | 0.005795 | 2 M1 _ | Trp53inp2 |
| Spry21 | 2.20E-07 | 0.659384 | 0.375 | 0.225 | 0.004425 | 2 M1 _ | Spry2 |
| Tmem571 | 2.13E-07 | 0.269056 | 0.133 | 0.038 | 0.004294 | 2 M1 _ | Tmem57 |
| Dag11 | 2.08E-07 | 0.534258 | 0.68 | 0.587 | 0.004197 | 2 M1 _ | Dag1 |
| Rsu1 | 1.86E-07 | 0.558599 | 0.688 | 0.591 | 0.003752 | 2 M1 _ | Rsu1 |
| mt-Atp6 | 1.85E-07 | 0.294823 | 1 | 0.993 | 0.003736 | 2 M1 _ | mt-Atp6 |
| Ankrd50 | 1.65E-07 | 0.476523 | 0.391 | 0.231 | 0.003322 | 2 M1 _ | Ankrd50 |
| Dab2ip | 1.37E-07 | 0.541405 | 0.625 | 0.49 | 0.002755 | 2 M1 _ | Dab2ip |
| Sparc | 1.29E-07 | 0.346003 | 1 | 0.918 | 0.002595 | 2 M1 _ | Sparc |
| Ppp1r12c | 1.16E-07 | 0.875905 | 0.422 | 0.286 | 0.002344 | 2 M1 _ | Ppp1r12c |
| Cdc42bpa | 1.16E-07 | 0.65829 | 0.562 | 0.423 | 0.002329 | 2 M1 _ | Cdc42bpa |
| Grk5 | 1.13E-07 | 0.408292 | 0.391 | 0.219 | 0.002281 | 2 M1 _ | Grk5 |
| Map1b | 1.09E-07 | 0.77192 | 0.414 | 0.262 | 0.002199 | 2 M1 _ | Map1b |
| Uqcrb | 1.05E-07 | 0.383568 | 0.867 | 0.822 | 0.00212 | 2 M1 _ | Uqcrb |
| Jade11 | 9.83E-08 | 0.720555 | 0.492 | 0.343 | 0.00198 | 2 M1 _ | Jade1 |
| Ppp2r1a | 9.58E-08 | 0.578419 | 0.578 | 0.47 | 0.00193 | 2 M1 _ | Ppp2r1a |
| Anxa11 | 9.45E-08 | 0.551818 | 0.336 | 0.182 | 0.001904 | 2 M1 _ | Anxa11 |
| Unc45a | 8.97E-08 | 0.472368 | 0.328 | 0.172 | 0.001807 | 2 M1 _ | Unc45a |
| Cyp7b1 | 8.80E-08 | 0.450849 | 0.312 | 0.152 | 0.001772 | 2 M1 _ | Cyp7b1 |
| Rab14 | 8.71E-08 | 0.465184 | 0.844 | 0.817 | 0.001754 | 2 M1 _ | Rab14 |
| Rnf152 | 8.12E-08 | 0.265921 | 0.211 | 0.077 | 0.001636 | 2 M1 _ | Rnf152 |
| Fbxo30 | 8.12E-08 | 0.410308 | 0.273 | 0.125 | 0.001636 | 2 M1 _ | Fbxo30 |
| Tgfb1i1 | 7.93E-08 | 0.602903 | 0.547 | 0.411 | 0.001597 | 2 M1 _ | Tgfb1i1 |
| Enpep | 7.51E-08 | 0.380229 | 0.32 | 0.152 | 0.001512 | 2 M1 _ | Enpep |
| Crip2 | 7.10E-08 | 0.42464 | 0.867 | 0.766 | 0.00143 | 2 M1 _ | Crip2 |
| Rapgef2 | 7.09E-08 | 0.633501 | 0.391 | 0.22 | 0.001428 | 2 M1 _ | Rapgef2 |
| Pdha1 | 6.93E-08 | 0.562153 | 0.578 | 0.432 | 0.001396 | 2 M1 _ | Pdha1 |
| Vdac1 | 6.81E-08 | 0.539186 | 0.625 | 0.54 | 0.001371 | 2 M1 _ | Vdac1 |
| Fxr1 | 6.58E-08 | 0.600488 | 0.656 | 0.537 | 0.001326 | 2 M1 _ | Fxr1 |
| Hbb-bs | 6.42E-08 | 1.712328 | 0.398 | 0.22 | 0.001293 | 2 M1 _ | Hbb-bs |

| | | | | | | | |
|---|---|---|---|---|---|---|---|
| Ncor1 | 6.37E-08 | 0.428683 | 0.797 | 0.753 | 0.001283 | 2 M1 _ | Ncor1 |
| Fgf1 | 4.86E-08 | 0.494234 | 0.242 | 0.101 | 0.000978 | 2 M1 _ | Fgf1 |
| Ntm | 4.85E-08 | 0.337065 | 0.172 | 0.053 | 0.000976 | 2 M1 _ | Ntm |
| Tnfaip1 | 4.83E-08 | 0.460415 | 0.555 | 0.372 | 0.000973 | 2 M1 _ | Tnfaip1 |
| Pmp221 | 4.58E-08 | 0.518272 | 0.844 | 0.7 | 0.000923 | 2 M1 _ | Pmp22 |
| Fam174a | 4.54E-08 | 0.692959 | 0.523 | 0.399 | 0.000915 | 2 M1 _ | Fam174a |
| Camk2n11 | 4.53E-08 | 0.72616 | 0.664 | 0.538 | 0.000913 | 2 M1 _ | Camk2n1 |
| Itpr1 | 4.16E-08 | 0.648083 | 0.594 | 0.472 | 0.000837 | 2 M1 _ | Itpr1 |
| Gpc6 | 4.16E-08 | 0.542353 | 0.578 | 0.415 | 0.000837 | 2 M1 _ | Gpc6 |
| Cnn2 | 3.94E-08 | 0.584155 | 0.586 | 0.437 | 0.000793 | 2 M1 _ | Cnn2 |
| Cavin1 | 3.83E-08 | 0.998158 | 0.586 | 0.478 | 0.000772 | 2 M1 _ | Cavin1 |
| Eif4e3 | 3.07E-08 | 0.508789 | 0.383 | 0.215 | 0.000619 | 2 M1 _ | Eif4e3 |
| Stx8 | 2.93E-08 | 0.551418 | 0.516 | 0.36 | 0.00059 | 2 M1 _ | Stx8 |
| Pacsin2 | 2.90E-08 | 0.604588 | 0.438 | 0.278 | 0.000583 | 2 M1 _ | Pacsin2 |
| Ggta1 | 2.78E-08 | 0.372834 | 0.281 | 0.12 | 0.00056 | 2 M1 _ | Ggta1 |
| Angpt1 | 2.69E-08 | 0.559185 | 0.32 | 0.157 | 0.000542 | 2 M1 _ | Angpt1 |
| Kank4 | 2.65E-08 | 0.256832 | 0.148 | 0.043 | 0.000534 | 2 M1 _ | Kank4 |
| Tprgl | 2.55E-08 | 0.526886 | 0.766 | 0.696 | 0.000514 | 2 M1 _ | Tprgl |
| Cux1 | 2.49E-08 | 0.61384 | 0.547 | 0.393 | 0.000501 | 2 M1 _ | Cux1 |
| Selenom | 2.43E-08 | 0.405848 | 0.953 | 0.816 | 0.000488 | 2 M1 _ | Selenom |
| Vdac2 | 2.39E-08 | 0.470074 | 0.766 | 0.673 | 0.000482 | 2 M1 _ | Vdac2 |
| Utrn | 2.22E-08 | 0.829355 | 0.617 | 0.484 | 0.000446 | 2 M1 _ | Utrn |
| Hnrnpa0 | 2.17E-08 | 0.502442 | 0.828 | 0.733 | 0.000436 | 2 M1 _ | Hnrnpa0 |
| Spry4 | 2.13E-08 | 0.571282 | 0.43 | 0.258 | 0.000429 | 2 M1 _ | Spry4 |
| Gnb4 | 2.03E-08 | 0.566536 | 0.797 | 0.663 | 0.000408 | 2 M1 _ | Gnb4 |
| Myh9 | 1.96E-08 | 0.548546 | 0.844 | 0.828 | 0.000395 | 2 M1 _ | Myh9 |
| Sult1a1 | 1.86E-08 | 0.652329 | 0.5 | 0.328 | 0.000375 | 2 M1 _ | Sult1a1 |
| Plxnb2 | 1.84E-08 | 0.564604 | 0.531 | 0.374 | 0.000371 | 2 M1 _ | Plxnb2 |
| C1qtnf11 | 1.60E-08 | 0.459548 | 0.328 | 0.158 | 0.000323 | 2 M1 _ | C1qtnf1 |
| Mgat4a | 1.59E-08 | 0.454941 | 0.25 | 0.101 | 0.000321 | 2 M1 _ | Mgat4a |
| Ptp4a3 | 1.54E-08 | 0.996253 | 0.469 | 0.319 | 0.000309 | 2 M1 _ | Ptp4a3 |
| 4930523C07Rik | 1.47E-08 | 0.629584 | 0.578 | 0.428 | 0.000297 | 2 M1 _ | 4930523C07Rik |
| Ppargc1b | 1.40E-08 | 0.303726 | 0.156 | 0.045 | 0.000282 | 2 M1 _ | Ppargc1b |
| Hbegf | 1.36E-08 | 0.339973 | 0.25 | 0.097 | 0.000275 | 2 M1 _ | Hbegf |
| Ostf1 | 1.26E-08 | 0.523472 | 0.727 | 0.605 | 0.000255 | 2 M1 _ | Ostf1 |
| Ilk | 1.12E-08 | 0.563603 | 0.664 | 0.576 | 0.000225 | 2 M1 _ | Ilk |
| Amotl11 | 1.01E-08 | 0.577959 | 0.742 | 0.633 | 0.000203 | 2 M1 _ | Amotl1 |
| Anxa51 | 1.01E-08 | 0.463824 | 0.977 | 0.882 | 0.000203 | 2 M1 _ | Anxa5 |
| Ubr1 | 8.42E-09 | 0.746829 | 0.531 | 0.381 | 0.00017 | 2 M1 _ | Ubr1 |
| Kitl | 8.27E-09 | 0.473099 | 0.609 | 0.398 | 0.000167 | 2 M1 _ | Kitl |
| Sox5 | 7.54E-09 | 0.383489 | 0.211 | 0.072 | 0.000152 | 2 M1 _ | Sox5 |

| | | | | | | | |
|---|---|---|---|---|---|---|---|
| Marveld1 | 7.00E-09 | 0.715896 | 0.5 | 0.351 | 0.000141 | 2 M1 _ | Marveld1 |
| Hipk1 | 6.91E-09 | 0.698579 | 0.523 | 0.367 | 0.000139 | 2 M1 _ | Hipk1 |
| Rasd1 | 6.83E-09 | 0.342498 | 0.133 | 0.033 | 0.000138 | 2 M1 _ | Rasd1 |
| Pfn1 | 6.74E-09 | 0.42568 | 0.883 | 0.79 | 0.000136 | 2 M1 _ | Pfn1 |
| Ecm2 | 6.67E-09 | 0.706256 | 0.648 | 0.5 | 0.000134 | 2 M1 _ | Ecm2 |
| Adam22 | 6.66E-09 | 0.27095 | 0.141 | 0.037 | 0.000134 | 2 M1 _ | Adam22 |
| Fhl11 | 6.35E-09 | 0.402074 | 0.203 | 0.068 | 0.000128 | 2 M1 _ | Fhl1 |
| Adcy3 | 5.54E-09 | 0.71186 | 0.391 | 0.225 | 0.000112 | 2 M1 _ | Adcy3 |
| Mustn1 | 5.45E-09 | 1.095518 | 0.273 | 0.113 | 0.00011 | 2 M1 _ | Mustn1 |
| Eif4ebp2 | 5.08E-09 | 0.528325 | 0.625 | 0.483 | 0.000102 | 2 M1 _ | Eif4ebp2 |
| Herc1 | 4.93E-09 | 0.6602 | 0.664 | 0.536 | 9.94E-05 | 2 M1 _ | Herc1 |
| Sox13 | 4.90E-09 | 0.330504 | 0.234 | 0.085 | 9.88E-05 | 2 M1 _ | Sox13 |
| Eif1b | 4.00E-09 | 0.610261 | 0.727 | 0.606 | 8.05E-05 | 2 M1 _ | Eif1b |
| Ppp2r3a | 3.78E-09 | 0.553977 | 0.398 | 0.216 | 7.62E-05 | 2 M1 _ | Ppp2r3a |
| Trib2 | 3.71E-09 | 0.815498 | 0.516 | 0.346 | 7.48E-05 | 2 M1 _ | Trib2 |
| Dapk2 | 3.57E-09 | 0.594969 | 0.547 | 0.359 | 7.19E-05 | 2 M1 _ | Dapk2 |
| Srsf5 | 2.72E-09 | 0.478684 | 0.836 | 0.758 | 5.48E-05 | 2 M1 _ | Srsf5 |
| Msrb1 | 2.69E-09 | 0.624863 | 0.484 | 0.313 | 5.42E-05 | 2 M1 _ | Msrb1 |
| Arrdc31 | 2.49E-09 | 0.775613 | 0.398 | 0.221 | 5.02E-05 | 2 M1 _ | Arrdc3 |
| Cycs | 2.46E-09 | 0.638536 | 0.648 | 0.531 | 4.96E-05 | 2 M1 _ | Cycs |
| Atp5k | 2.36E-09 | 0.568271 | 0.781 | 0.752 | 4.74E-05 | 2 M1 _ | Atp5k |
| Stim1 | 2.13E-09 | 0.589514 | 0.578 | 0.401 | 4.30E-05 | 2 M1 _ | Stim1 |
| Fam107b | 1.89E-09 | 0.354072 | 0.266 | 0.1 | 3.81E-05 | 2 M1 _ | Fam107b |
| Lpcat3 | 1.72E-09 | 0.642493 | 0.492 | 0.316 | 3.46E-05 | 2 M1 _ | Lpcat3 |
| Pear1 | 1.65E-09 | 0.671022 | 0.578 | 0.425 | 3.33E-05 | 2 M1 _ | Pear1 |
| Map2 | 1.52E-09 | 0.482181 | 0.328 | 0.145 | 3.06E-05 | 2 M1 _ | Map2 |
| Dpysl21 | 1.37E-09 | 0.724781 | 0.617 | 0.471 | 2.76E-05 | 2 M1 _ | Dpysl2 |
| Aig1 | 1.35E-09 | 0.66667 | 0.406 | 0.219 | 2.72E-05 | 2 M1 _ | Aig1 |
| Sdccag31 | 1.20E-09 | 0.334167 | 0.156 | 0.042 | 2.41E-05 | 2 M1 _ | Sdccag3 |
| Cox6c | 1.16E-09 | 0.453576 | 0.977 | 0.923 | 2.34E-05 | 2 M1 _ | Cox6c |
| Ppp1r3d | 1.10E-09 | 0.297511 | 0.148 | 0.037 | 2.22E-05 | 2 M1 _ | Ppp1r3d |
| Ralgps1 | 9.94E-10 | 0.313226 | 0.164 | 0.044 | 2.00E-05 | 2 M1 _ | Ralgps1 |
| Suco | 9.16E-10 | 0.810698 | 0.461 | 0.272 | 1.85E-05 | 2 M1 _ | Suco |
| Gpcpd1 | 9.16E-10 | 0.462984 | 0.352 | 0.167 | 1.84E-05 | 2 M1 _ | Gpcpd1 |
| Epb41 | 8.95E-10 | 0.500348 | 0.359 | 0.169 | 1.80E-05 | 2 M1 _ | Epb41 |
| Tnfaip8 | 8.21E-10 | 0.630682 | 0.625 | 0.441 | 1.65E-05 | 2 M1 _ | Tnfaip8 |
| Klhdc8b | 8.19E-10 | 0.479522 | 0.375 | 0.18 | 1.65E-05 | 2 M1 _ | Klhdc8b |
| Calm1 | 6.42E-10 | 0.427623 | 0.977 | 0.95 | 1.29E-05 | 2 M1 _ | Calm1 |
| Tead1 | 6.37E-10 | 0.65964 | 0.531 | 0.357 | 1.28E-05 | 2 M1 _ | Tead1 |
| Ywhab | 6.21E-10 | 0.526862 | 0.867 | 0.832 | 1.25E-05 | 2 M1 _ | Ywhab |
| Gcn1l11 | 6.05E-10 | 0.381237 | 0.156 | 0.041 | 1.22E-05 | 2 M1 _ | Gcn1l1 |

| | | | | | | | |
|---|---|---|---|---|---|---|---|
| Map3k20 | 5.89E-10 | 1.038977 | 0.469 | 0.295 | 1.19E-05 | 2 M1 _ | Map3k20 |
| Nmt1 | 5.87E-10 | 0.650625 | 0.75 | 0.659 | 1.18E-05 | 2 M1 _ | Nmt1 |
| Snrpn | 5.84E-10 | 0.325501 | 0.141 | 0.034 | 1.18E-05 | 2 M1 _ | Snrpn |
| Ccnd1 | 5.08E-10 | 1.127652 | 0.359 | 0.167 | 1.02E-05 | 2 M1 _ | Ccnd1 |
| Dyrk2 | 4.44E-10 | 0.582768 | 0.391 | 0.202 | 8.95E-06 | 2 M1 _ | Dyrk2 |
| Sema6d | 4.27E-10 | 0.607882 | 0.562 | 0.347 | 8.60E-06 | 2 M1 _ | Sema6d |
| Ppm1l | 4.21E-10 | 0.489914 | 0.164 | 0.044 | 8.49E-06 | 2 M1 _ | Ppm1l |
| Uqcr11 | 4.20E-10 | 0.514505 | 0.82 | 0.808 | 8.46E-06 | 2 M1 _ | Uqcr11 |
| Lamb2 | 2.97E-10 | 0.747207 | 0.625 | 0.511 | 5.97E-06 | 2 M1 _ | Lamb2 |
| Tuba1b1 | 2.88E-10 | 0.521374 | 0.867 | 0.82 | 5.80E-06 | 2 M1 _ | Tuba1b |
| Filip1 | 2.86E-10 | 0.618033 | 0.383 | 0.184 | 5.77E-06 | 2 M1 _ | Filip1 |
| Pcp4l1 | 2.67E-10 | 0.70433 | 0.445 | 0.214 | 5.38E-06 | 2 M1 _ | Pcp4l1 |
| Dmpk1 | 2.57E-10 | 0.538741 | 0.273 | 0.106 | 5.17E-06 | 2 M1 _ | Dmpk |
| Lims1 | 2.43E-10 | 0.662419 | 0.656 | 0.548 | 4.90E-06 | 2 M1 _ | Lims1 |
| Ptp4a11 | 2.11E-10 | 0.324467 | 0.211 | 0.065 | 4.24E-06 | 2 M1 _ | Ptp4a1 |
| Gapdh | 2.03E-10 | 0.497748 | 0.953 | 0.905 | 4.09E-06 | 2 M1 _ | Gapdh |
| Bok | 1.78E-10 | 0.418461 | 0.336 | 0.138 | 3.58E-06 | 2 M1 _ | Bok |
| Stk38l | 1.67E-10 | 0.452875 | 0.32 | 0.13 | 3.37E-06 | 2 M1 _ | Stk38l |
| Prdx1 | 1.64E-10 | 0.450863 | 0.977 | 0.906 | 3.31E-06 | 2 M1 _ | Prdx1 |
| Tmco1 | 1.37E-10 | 0.66276 | 0.797 | 0.721 | 2.76E-06 | 2 M1 _ | Tmco1 |
| Dlc1 | 1.32E-10 | 0.478515 | 0.836 | 0.698 | 2.67E-06 | 2 M1 _ | Dlc1 |
| Rcsd1 | 1.26E-10 | 0.447358 | 0.438 | 0.214 | 2.54E-06 | 2 M1 _ | Rcsd1 |
| Cd811 | 1.25E-10 | 0.574721 | 0.992 | 0.891 | 2.51E-06 | 2 M1 _ | Cd81 |
| Smpdl3a | 1.17E-10 | 0.795064 | 0.633 | 0.472 | 2.36E-06 | 2 M1 _ | Smpdl3a |
| Etfb | 9.82E-11 | 0.582395 | 0.703 | 0.562 | 1.98E-06 | 2 M1 _ | Etfb |
| Ktn1 | 7.38E-11 | 0.663149 | 0.734 | 0.62 | 1.49E-06 | 2 M1 _ | Ktn1 |
| Ndufs2 | 7.16E-11 | 0.821965 | 0.68 | 0.556 | 1.44E-06 | 2 M1 _ | Ndufs2 |
| Smagp | 5.47E-11 | 0.593746 | 0.344 | 0.142 | 1.10E-06 | 2 M1 _ | Smagp |
| A230050P20Rik1 | 5.25E-11 | 0.33289 | 0.125 | 0.025 | 1.06E-06 | 2 M1 _ | A230050P20Rik |
| Gnas1 | 5.04E-11 | 0.39407 | 1 | 0.959 | 1.01E-06 | 2 M1 _ | Gnas |
| Rap1a | 4.59E-11 | 0.640751 | 0.734 | 0.649 | 9.24E-07 | 2 M1 _ | Rap1a |
| S1pr31 | 4.49E-11 | 0.785937 | 0.469 | 0.281 | 9.05E-07 | 2 M1 _ | S1pr3 |
| Kcnab1 | 4.47E-11 | 0.382482 | 0.109 | 0.019 | 9.01E-07 | 2 M1 _ | Kcnab1 |
| Ephx2 | 4.12E-11 | 0.284814 | 0.164 | 0.04 | 8.31E-07 | 2 M1 _ | Ephx2 |
| Mgea51 | 4.03E-11 | 0.338736 | 0.188 | 0.051 | 8.12E-07 | 2 M1 _ | Mgea5 |
| Slc11a1 | 3.89E-11 | 0.426891 | 0.156 | 0.036 | 7.82E-07 | 2 M1 _ | Slc11a1 |
| Nedd41 | 3.81E-11 | 0.486813 | 0.977 | 0.884 | 7.67E-07 | 2 M1 _ | Nedd4 |
| Asap2 | 3.68E-11 | 0.52387 | 0.508 | 0.274 | 7.42E-07 | 2 M1 _ | Asap2 |
| Tns3 | 3.44E-11 | 0.613546 | 0.461 | 0.257 | 6.92E-07 | 2 M1 _ | Tns3 |
| Papd41 | 3.02E-11 | 0.279337 | 0.156 | 0.037 | 6.08E-07 | 2 M1 _ | Papd4 |
| Ate1 | 2.93E-11 | 0.751362 | 0.586 | 0.414 | 5.90E-07 | 2 M1 _ | Ate1 |

| | | | | | | | |
|---|---|---|---|---|---|---|---|
| Nab1 | 2.83E-11 | 0.889189 | 0.531 | 0.348 | 5.71E-07 | 2 M1 _ | Nab1 |
| Ptk2 | 2.64E-11 | 0.640595 | 0.555 | 0.34 | 5.32E-07 | 2 M1 _ | Ptk2 |
| Celf1 | 2.62E-11 | 0.637815 | 0.594 | 0.423 | 5.28E-07 | 2 M1 _ | Celf1 |
| 4933431E20Rik | 2.50E-11 | 0.43872 | 0.133 | 0.027 | 5.04E-07 | 2 M1 _ | 4933431E20Rik |
| Ptpn9 | 2.46E-11 | 0.793761 | 0.594 | 0.435 | 4.95E-07 | 2 M1 _ | Ptpn9 |
| Peli2 | 2.43E-11 | 0.617975 | 0.359 | 0.16 | 4.89E-07 | 2 M1 _ | Peli2 |
| Mbnl2 | 2.42E-11 | 0.644131 | 0.805 | 0.697 | 4.86E-07 | 2 M1 _ | Mbnl2 |
| Pik3r11 | 2.40E-11 | 0.634372 | 0.781 | 0.659 | 4.84E-07 | 2 M1 _ | Pik3r1 |
| Cfl2 | 2.21E-11 | 0.71484 | 0.602 | 0.428 | 4.44E-07 | 2 M1 _ | Cfl2 |
| Chtf81 | 2.20E-11 | 0.409582 | 0.25 | 0.083 | 4.43E-07 | 2 M1 _ | Chtf8 |
| Rhou | 2.02E-11 | 0.826078 | 0.312 | 0.133 | 4.06E-07 | 2 M1 _ | Rhou |
| Mef2c | 1.85E-11 | 0.436417 | 0.852 | 0.64 | 3.72E-07 | 2 M1 _ | Mef2c |
| Il34 | 1.76E-11 | 0.570851 | 0.766 | 0.482 | 3.54E-07 | 2 M1 _ | Il34 |
| 9030624J02Rik1 | 1.69E-11 | 0.394857 | 0.164 | 0.039 | 3.41E-07 | 2 M1 _ | 9030624J02Rik |
| Pgrmc1 | 1.65E-11 | 0.660137 | 0.711 | 0.54 | 3.32E-07 | 2 M1 _ | Pgrmc1 |
| Slc4a3 | 1.60E-11 | 0.474725 | 0.234 | 0.075 | 3.22E-07 | 2 M1 _ | Slc4a3 |
| Pde4d | 1.51E-11 | 0.393202 | 0.203 | 0.057 | 3.03E-07 | 2 M1 _ | Pde4d |
| Fry | 1.36E-11 | 0.600541 | 0.406 | 0.179 | 2.75E-07 | 2 M1 _ | Fry |
| Epb41l21 | 1.36E-11 | 0.611475 | 0.797 | 0.673 | 2.73E-07 | 2 M1 _ | Epb41l2 |
| Ndufb9 | 1.06E-11 | 0.573541 | 0.906 | 0.822 | 2.14E-07 | 2 M1 _ | Ndufb9 |
| Prkar1b | 1.03E-11 | 0.444703 | 0.188 | 0.048 | 2.07E-07 | 2 M1 _ | Prkar1b |
| Hsp90aa1 | 9.47E-12 | 0.644593 | 0.828 | 0.736 | 1.91E-07 | 2 M1 _ | Hsp90aa1 |
| Cdc42ep4 | 9.35E-12 | 0.633793 | 0.469 | 0.261 | 1.88E-07 | 2 M1 _ | Cdc42ep4 |
| Epha2 | 7.87E-12 | 0.313338 | 0.188 | 0.048 | 1.59E-07 | 2 M1 _ | Epha2 |
| Naf1 | 7.80E-12 | 0.636758 | 0.391 | 0.184 | 1.57E-07 | 2 M1 _ | Naf1 |
| Fam124a | 7.78E-12 | 0.323124 | 0.18 | 0.044 | 1.57E-07 | 2 M1 _ | Fam124a |
| Stbd1 | 6.51E-12 | 0.70434 | 0.234 | 0.073 | 1.31E-07 | 2 M1 _ | Stbd1 |
| Syn3 | 5.53E-12 | 0.497598 | 0.289 | 0.101 | 1.11E-07 | 2 M1 _ | Syn3 |
| Rbm8a | 5.01E-12 | 0.685589 | 0.734 | 0.574 | 1.01E-07 | 2 M1 _ | Rbm8a |
| Cyr611 | 4.64E-12 | 0.332079 | 0.148 | 0.031 | 9.35E-08 | 2 M1 _ | Cyr61 |
| Svopl | 4.40E-12 | 0.53578 | 0.172 | 0.041 | 8.86E-08 | 2 M1 _ | Svopl |
| Slc25a33 | 4.24E-12 | 0.389254 | 0.219 | 0.062 | 8.54E-08 | 2 M1 _ | Slc25a33 |
| Pik3ca | 3.92E-12 | 0.817988 | 0.562 | 0.365 | 7.90E-08 | 2 M1 _ | Pik3ca |
| Trpc3 | 3.61E-12 | 0.337253 | 0.164 | 0.036 | 7.27E-08 | 2 M1 _ | Trpc3 |
| Anxa3 | 3.37E-12 | 0.46027 | 0.578 | 0.301 | 6.80E-08 | 2 M1 _ | Anxa3 |
| Arhgef16 | 3.11E-12 | 0.272867 | 0.148 | 0.029 | 6.26E-08 | 2 M1 _ | Arhgef16 |
| Rab8b | 2.94E-12 | 0.706986 | 0.57 | 0.349 | 5.92E-08 | 2 M1 _ | Rab8b |
| Rras | 2.89E-12 | 0.724435 | 0.773 | 0.673 | 5.81E-08 | 2 M1 _ | Rras |
| Arap2 | 2.87E-12 | 0.766147 | 0.602 | 0.39 | 5.78E-08 | 2 M1 _ | Arap2 |
| Adcyap1r11 | 2.81E-12 | 0.886086 | 0.484 | 0.274 | 5.65E-08 | 2 M1 _ | Adcyap1r1 |
| Gnai1 | 2.50E-12 | 0.461464 | 0.211 | 0.058 | 5.04E-08 | 2 M1 _ | Gnai1 |

| | | | | | | | |
|---|---|---|---|---|---|---|---|
| Trpc1 | 2.45E-12 | 0.540451 | 0.305 | 0.11 | 4.93E-08 | 2 M1 _ | Trpc1 |
| Pdcd10 | 2.26E-12 | 0.707175 | 0.719 | 0.57 | 4.54E-08 | 2 M1 _ | Pdcd10 |
| Suclg1 | 2.07E-12 | 0.652917 | 0.688 | 0.526 | 4.16E-08 | 2 M1 _ | Suclg1 |
| Sh3bgrl | 1.70E-12 | 0.805705 | 0.75 | 0.683 | 3.43E-08 | 2 M1 _ | Sh3bgrl |
| Papd5 | 1.64E-12 | 0.337796 | 0.133 | 0.024 | 3.30E-08 | 2 M1 _ | Papd5 |
| 1110004E09Rik | 1.22E-12 | 0.312119 | 0.148 | 0.03 | 2.46E-08 | 2 M1 _ | 1110004E09Rik |
| Tpm4 | 1.18E-12 | 0.791759 | 0.844 | 0.751 | 2.39E-08 | 2 M1 _ | Tpm4 |
| Meox2 | 1.11E-12 | 0.499548 | 0.383 | 0.154 | 2.23E-08 | 2 M1 _ | Meox2 |
| Dusp11 | 1.09E-12 | 0.755547 | 0.562 | 0.326 | 2.19E-08 | 2 M1 _ | Dusp1 |
| Tmem51 | 1.02E-12 | 0.495142 | 0.258 | 0.08 | 2.05E-08 | 2 M1 _ | Tmem51 |
| Col4a11 | 7.69E-13 | 0.634426 | 0.992 | 0.866 | 1.55E-08 | 2 M1 _ | Col4a1 |
| Stac | 7.22E-13 | 0.325923 | 0.195 | 0.046 | 1.45E-08 | 2 M1 _ | Stac |
| Tspan3 | 7.11E-13 | 0.659639 | 0.828 | 0.659 | 1.43E-08 | 2 M1 _ | Tspan3 |
| Rock2 | 6.37E-13 | 0.811271 | 0.766 | 0.665 | 1.28E-08 | 2 M1 _ | Rock2 |
| Atp2b4 | 6.09E-13 | 0.802786 | 0.531 | 0.301 | 1.23E-08 | 2 M1 _ | Atp2b4 |
| Fam110a | 5.67E-13 | 0.438091 | 0.258 | 0.079 | 1.14E-08 | 2 M1 _ | Fam110a |
| 0610009O20Rik1 | 5.30E-13 | 0.255544 | 0.156 | 0.032 | 1.07E-08 | 2 M1 _ | 0610009O20Rik |
| Daam1 | 5.25E-13 | 0.702346 | 0.594 | 0.367 | 1.06E-08 | 2 M1 _ | Daam1 |
| Snrnp27 | 5.22E-13 | 0.764444 | 0.695 | 0.517 | 1.05E-08 | 2 M1 _ | Snrnp27 |
| Nbeal1 | 4.84E-13 | 0.696045 | 0.758 | 0.624 | 9.74E-09 | 2 M1 _ | Nbeal1 |
| Kdelc21 | 4.81E-13 | 0.418497 | 0.234 | 0.066 | 9.68E-09 | 2 M1 _ | Kdelc2 |
| Cav2 | 4.52E-13 | 0.501916 | 0.453 | 0.194 | 9.10E-09 | 2 M1 _ | Cav2 |
| Ech1 | 4.26E-13 | 0.689106 | 0.781 | 0.621 | 8.58E-09 | 2 M1 _ | Ech1 |
| Hgf | 3.85E-13 | 0.689151 | 0.328 | 0.125 | 7.76E-09 | 2 M1 _ | Hgf |
| Kank1 | 3.16E-13 | 0.552348 | 0.312 | 0.114 | 6.37E-09 | 2 M1 _ | Kank1 |
| Ptprf | 2.66E-13 | 0.364156 | 0.25 | 0.073 | 5.36E-09 | 2 M1 _ | Ptprf |
| Zfhx3 | 1.85E-13 | 1.055904 | 0.492 | 0.265 | 3.73E-09 | 2 M1 _ | Zfhx3 |
| Spag71 | 1.78E-13 | 0.61887 | 0.391 | 0.172 | 3.59E-09 | 2 M1 _ | Spag7 |
| Clstn1 | 1.76E-13 | 0.744922 | 0.742 | 0.56 | 3.54E-09 | 2 M1 _ | Clstn1 |
| Akap2 | 1.64E-13 | 0.30909 | 0.125 | 0.02 | 3.30E-09 | 2 M1 _ | Akap2 |
| Atp9a | 1.23E-13 | 0.573735 | 0.281 | 0.091 | 2.47E-09 | 2 M1 _ | Atp9a |
| Sntb1 | 1.19E-13 | 0.380863 | 0.172 | 0.036 | 2.40E-09 | 2 M1 _ | Sntb1 |
| Pkig | 1.08E-13 | 0.668494 | 0.789 | 0.611 | 2.18E-09 | 2 M1 _ | Pkig |
| Vasp | 7.23E-14 | 0.824294 | 0.648 | 0.445 | 1.46E-09 | 2 M1 _ | Vasp |
| Ubb | 6.88E-14 | 0.422361 | 1 | 0.978 | 1.38E-09 | 2 M1 _ | Ubb |
| Hrc | 6.39E-14 | 0.271885 | 0.141 | 0.024 | 1.29E-09 | 2 M1 _ | Hrc |
| Afap1l2 | 5.39E-14 | 0.567276 | 0.43 | 0.188 | 1.09E-09 | 2 M1 _ | Afap1l2 |
| Pacsin3 | 4.43E-14 | 0.684487 | 0.312 | 0.108 | 8.93E-10 | 2 M1 _ | Pacsin3 |
| Plekha2 | 3.80E-14 | 0.758696 | 0.445 | 0.212 | 7.66E-10 | 2 M1 _ | Plekha2 |
| Adora2a | 3.60E-14 | 0.420812 | 0.234 | 0.062 | 7.25E-10 | 2 M1 _ | Adora2a |
| Apold1 | 3.53E-14 | 0.42421 | 0.258 | 0.072 | 7.12E-10 | 2 M1 _ | Apold1 |

| | | | | | | | |
|---|---|---|---|---|---|---|---|
| Fabp5 | 3.49E-14 | 0.850912 | 0.5 | 0.239 | 7.03E-10 | 2 M1 _ | Fabp5 |
| Chchd2 | 3.29E-14 | 0.436797 | 0.984 | 0.96 | 6.63E-10 | 2 M1 _ | Chchd2 |
| Ggt51 | 3.10E-14 | 0.808192 | 0.609 | 0.349 | 6.24E-10 | 2 M1 _ | Ggt5 |
| Wtip | 2.86E-14 | 1.230742 | 0.539 | 0.334 | 5.76E-10 | 2 M1 _ | Wtip |
| Atp5g3 | 2.24E-14 | 0.759621 | 0.797 | 0.689 | 4.51E-10 | 2 M1 _ | Atp5g3 |
| Selenow | 1.85E-14 | 0.601273 | 0.969 | 0.889 | 3.73E-10 | 2 M1 _ | Selenow |
| Rgs7bp1 | 1.45E-14 | 0.928509 | 0.375 | 0.151 | 2.92E-10 | 2 M1 _ | Rgs7bp |
| Bdnf | 1.44E-14 | 0.326003 | 0.109 | 0.014 | 2.90E-10 | 2 M1 _ | Bdnf |
| Cds2 | 1.35E-14 | 0.664039 | 0.547 | 0.292 | 2.73E-10 | 2 M1 _ | Cds2 |
| Plxdc1 | 1.26E-14 | 0.334808 | 0.164 | 0.03 | 2.54E-10 | 2 M1 _ | Plxdc1 |
| Hspb1 | 1.20E-14 | 0.843266 | 0.797 | 0.592 | 2.41E-10 | 2 M1 _ | Hspb1 |
| Dopey11 | 1.09E-14 | 0.382292 | 0.203 | 0.046 | 2.20E-10 | 2 M1 _ | Dopey1 |
| 11-Sep | 7.38E-15 | 0.699197 | 0.859 | 0.735 | 1.49E-10 | 2 M1 _ | 11-Sep |
| Plce1 | 5.55E-15 | 0.681001 | 0.445 | 0.194 | 1.12E-10 | 2 M1 _ | Plce1 |
| Grk3 | 5.47E-15 | 0.460183 | 0.242 | 0.062 | 1.10E-10 | 2 M1 _ | Grk3 |
| Gm13470 | 4.64E-15 | 0.394126 | 0.18 | 0.036 | 9.34E-11 | 2 M1 _ | Gm13470 |
| Rbm24 | 4.14E-15 | 0.261212 | 0.102 | 0.011 | 8.34E-11 | 2 M1 _ | Rbm24 |
| D17Wsu92e1 | 3.83E-15 | 0.471783 | 0.219 | 0.053 | 7.70E-11 | 2 M1 _ | D17Wsu92e |
| Fam26e1 | 3.68E-15 | 0.26073 | 0.133 | 0.02 | 7.42E-11 | 2 M1 _ | Fam26e |
| Ptges3l | 3.31E-15 | 0.367352 | 0.172 | 0.034 | 6.66E-11 | 2 M1 _ | Ptges3l |
| Prkg1 | 3.27E-15 | 0.781952 | 0.438 | 0.185 | 6.58E-11 | 2 M1 _ | Prkg1 |
| Anxa1 | 2.25E-15 | 0.908912 | 0.75 | 0.564 | 4.53E-11 | 2 M1 _ | Anxa1 |
| Mkl2 | 2.17E-15 | 0.320011 | 0.211 | 0.047 | 4.37E-11 | 2 M1 _ | Mkl2 |
| Ifitm1 | 2.04E-15 | 1.037288 | 0.484 | 0.204 | 4.11E-11 | 2 M1 _ | Ifitm1 |
| Aldh21 | 1.99E-15 | 0.698509 | 0.891 | 0.751 | 4.01E-11 | 2 M1 _ | Aldh2 |
| Prune2 | 1.98E-15 | 0.376634 | 0.109 | 0.013 | 4.00E-11 | 2 M1 _ | Prune2 |
| Sep-81 | 1.76E-15 | 0.962585 | 0.594 | 0.381 | 3.55E-11 | 2 M1 _ | 8-Sep |
| Ndufs5 | 1.63E-15 | 0.619141 | 0.875 | 0.778 | 3.28E-11 | 2 M1 _ | Ndufs5 |
| Tm4sf1 | 9.35E-16 | 0.552553 | 0.891 | 0.685 | 1.88E-11 | 2 M1 _ | Tm4sf1 |
| Msn | 7.66E-16 | 0.574318 | 0.945 | 0.789 | 1.54E-11 | 2 M1 _ | Msn |
| Nr2f2 | 7.39E-16 | 0.73299 | 0.852 | 0.673 | 1.49E-11 | 2 M1 _ | Nr2f2 |
| Tspan2 | 5.74E-16 | 0.447079 | 0.227 | 0.053 | 1.16E-11 | 2 M1 _ | Tspan2 |
| Snx181 | 5.67E-16 | 0.84221 | 0.648 | 0.404 | 1.14E-11 | 2 M1 _ | Snx18 |
| Angpt2 | 4.11E-16 | 0.484292 | 0.289 | 0.079 | 8.28E-12 | 2 M1 _ | Angpt2 |
| AU0198231 | 3.92E-16 | 0.445615 | 0.18 | 0.034 | 7.90E-12 | 2 M1 _ | AU019823 |
| Arhgef25 | 3.29E-16 | 0.683658 | 0.625 | 0.367 | 6.63E-12 | 2 M1 _ | Arhgef25 |
| Epb41l1 | 3.05E-16 | 0.759277 | 0.414 | 0.169 | 6.15E-12 | 2 M1 _ | Epb41l1 |
| Cited2 | 2.71E-16 | 0.819245 | 0.555 | 0.271 | 5.46E-12 | 2 M1 _ | Cited2 |
| Jag1 | 2.59E-16 | 0.741342 | 0.516 | 0.247 | 5.21E-12 | 2 M1 _ | Jag1 |
| Kcnq4 | 2.39E-16 | 0.439459 | 0.234 | 0.055 | 4.81E-12 | 2 M1 _ | Kcnq4 |
| Cend1 | 2.00E-16 | 0.32819 | 0.117 | 0.014 | 4.04E-12 | 2 M1 _ | Cend1 |

| | | | | | | | |
|---|---|---|---|---|---|---|---|
| Tln1 | 1.72E-16 | 0.676666 | 0.867 | 0.812 | 3.47E-12 | 2 M1 _ | Tln1 |
| Plcl1 | 1.66E-16 | 0.660464 | 0.383 | 0.137 | 3.33E-12 | 2 M1 _ | Plcl1 |
| 0610037L13Rik1 | 1.38E-16 | 0.416831 | 0.211 | 0.045 | 2.78E-12 | 2 M1 _ | 0610037L13Rik |
| 2410089E03Rik1 | 1.09E-16 | 0.494566 | 0.172 | 0.031 | 2.19E-12 | 2 M1 _ | 2410089E03Rik |
| Olfr78 | 1.01E-16 | 0.347171 | 0.117 | 0.014 | 2.03E-12 | 2 M1 _ | Olfr78 |
| Smarcd3 | 8.83E-17 | 0.560849 | 0.289 | 0.081 | 1.78E-12 | 2 M1 _ | Smarcd3 |
| Stard8 | 8.74E-17 | 0.845035 | 0.477 | 0.201 | 1.76E-12 | 2 M1 _ | Stard8 |
| Ppp1r12b | 7.12E-17 | 1.010283 | 0.445 | 0.188 | 1.43E-12 | 2 M1 _ | Ppp1r12b |
| Fahd1 | 6.76E-17 | 0.723711 | 0.375 | 0.134 | 1.36E-12 | 2 M1 _ | Fahd1 |
| Pip5k1b | 6.35E-17 | 0.47003 | 0.195 | 0.038 | 1.28E-12 | 2 M1 _ | Pip5k1b |
| Cadm3 | 6.14E-17 | 0.564195 | 0.227 | 0.051 | 1.24E-12 | 2 M1 _ | Cadm3 |
| Il2ra | 4.91E-17 | 0.444933 | 0.242 | 0.055 | 9.88E-13 | 2 M1 _ | Il2ra |
| 4931406P16Rik | 4.26E-17 | 0.977189 | 0.688 | 0.433 | 8.57E-13 | 2 M1 _ | 4931406P16Rik |
| Tubb4b | 4.10E-17 | 0.766312 | 0.867 | 0.693 | 8.25E-13 | 2 M1 _ | Tubb4b |
| Nes | 3.98E-17 | 0.78215 | 0.516 | 0.216 | 8.02E-13 | 2 M1 _ | Nes |
| Ldlrap1 | 3.78E-17 | 0.738898 | 0.453 | 0.191 | 7.62E-13 | 2 M1 _ | Ldlrap1 |
| Fam58b1 | 3.44E-17 | 0.428603 | 0.18 | 0.033 | 6.93E-13 | 2 M1 _ | Fam58b |
| Pawr | 3.17E-17 | 0.690793 | 0.375 | 0.127 | 6.37E-13 | 2 M1 _ | Pawr |
| Zufsp1 | 2.78E-17 | 0.437447 | 0.172 | 0.029 | 5.59E-13 | 2 M1 _ | Zufsp |
| Gstm11 | 2.50E-17 | 1.055422 | 0.859 | 0.655 | 5.03E-13 | 2 M1 _ | Gstm1 |
| Laptm4a | 2.11E-17 | 0.706803 | 1 | 0.922 | 4.24E-13 | 2 M1 _ | Laptm4a |
| Gstm5 | 1.97E-17 | 1.314971 | 0.609 | 0.37 | 3.97E-13 | 2 M1 _ | Gstm5 |
| Gm26699 | 1.80E-17 | 0.373136 | 0.109 | 0.011 | 3.62E-13 | 2 M1 _ | Gm26699 |
| Gamt | 1.77E-17 | 0.715816 | 0.547 | 0.255 | 3.56E-13 | 2 M1 _ | Gamt |
| Grm7 | 1.14E-17 | 0.312988 | 0.133 | 0.016 | 2.30E-13 | 2 M1 _ | Grm7 |
| Zfp467 | 9.96E-18 | 0.698487 | 0.453 | 0.171 | 2.01E-13 | 2 M1 _ | Zfp467 |
| Uba2 | 8.95E-18 | 0.971246 | 0.648 | 0.422 | 1.80E-13 | 2 M1 _ | Uba2 |
| Bmp2 | 8.25E-18 | 0.690368 | 0.289 | 0.079 | 1.66E-13 | 2 M1 _ | Bmp2 |
| Cdh2 | 8.12E-18 | 0.499157 | 0.211 | 0.041 | 1.63E-13 | 2 M1 _ | Cdh2 |
| Bmp5 | 7.44E-18 | 0.365032 | 0.141 | 0.019 | 1.50E-13 | 2 M1 _ | Bmp5 |
| Ddit4l | 6.59E-18 | 0.486948 | 0.297 | 0.074 | 1.33E-13 | 2 M1 _ | Ddit4l |
| Nxf11 | 4.92E-18 | 0.860135 | 0.539 | 0.261 | 9.91E-14 | 2 M1 _ | Nxf1 |
| mt-Nd2 | 4.91E-18 | 0.518629 | 0.992 | 0.98 | 9.88E-14 | 2 M1 _ | mt-Nd2 |
| 2310035C23Rik1 | 4.77E-18 | 0.384111 | 0.18 | 0.031 | 9.60E-14 | 2 M1 _ | 2310035C23Rik |
| Gpx8 | 3.21E-18 | 1.008116 | 0.875 | 0.64 | 6.46E-14 | 2 M1 _ | Gpx8 |
| Col4a2 | 3.10E-18 | 0.761792 | 0.977 | 0.85 | 6.25E-14 | 2 M1 _ | Col4a2 |
| Tnks1bp1 | 2.17E-18 | 0.913032 | 0.602 | 0.331 | 4.37E-14 | 2 M1 _ | Tnks1bp1 |
| Trabd2b1 | 1.98E-18 | 0.70772 | 0.344 | 0.104 | 3.99E-14 | 2 M1 _ | Trabd2b |
| Cox8a | 1.95E-18 | 0.656166 | 0.969 | 0.944 | 3.92E-14 | 2 M1 _ | Cox8a |
| Mut1 | 1.73E-18 | 0.461288 | 0.211 | 0.041 | 3.48E-14 | 2 M1 _ | Mut |
| Fam129a1 | 1.71E-18 | 0.916473 | 0.664 | 0.381 | 3.45E-14 | 2 M1 _ | Fam129a |

| | | | | | | | |
|---|---|---|---|---|---|---|---|
| Ywhaq | 1.35E-18 | 0.707637 | 0.883 | 0.793 | 2.72E-14 | 2 M1 _ | Ywhaq |
| Tbx3os1 | 1.34E-18 | 0.445413 | 0.234 | 0.048 | 2.71E-14 | 2 M1 _ | Tbx3os1 |
| Plcb4 | 1.24E-18 | 0.975718 | 0.523 | 0.243 | 2.50E-14 | 2 M1 _ | Plcb4 |
| Olfml2b1 | 1.10E-18 | 0.583579 | 0.383 | 0.117 | 2.22E-14 | 2 M1 _ | Olfml2b |
| Kcnb1 | 9.30E-19 | 0.457827 | 0.258 | 0.058 | 1.87E-14 | 2 M1 _ | Kcnb1 |
| Ehd1 | 8.53E-19 | 0.859248 | 0.555 | 0.275 | 1.72E-14 | 2 M1 _ | Ehd1 |
| Ctgf1 | 8.07E-19 | 0.320321 | 0.219 | 0.04 | 1.63E-14 | 2 M1 _ | Ctgf |
| Csnk1e | 4.86E-19 | 0.9387 | 0.641 | 0.365 | 9.79E-15 | 2 M1 _ | Csnk1e |
| Mtus1 | 2.98E-19 | 0.862331 | 0.602 | 0.298 | 6.01E-15 | 2 M1 _ | Mtus1 |
| Phlda1 | 2.87E-19 | 0.957375 | 0.688 | 0.405 | 5.78E-15 | 2 M1 _ | Phlda1 |
| Sort1 | 2.75E-19 | 0.617848 | 0.359 | 0.106 | 5.55E-15 | 2 M1 _ | Sort1 |
| Tmem38b | 2.06E-19 | 0.723238 | 0.391 | 0.131 | 4.15E-15 | 2 M1 _ | Tmem38b |
| Postn | 1.76E-19 | 0.868678 | 0.836 | 0.45 | 3.55E-15 | 2 M1 _ | Postn |
| Cacna1c | 1.25E-19 | 0.675658 | 0.438 | 0.155 | 2.53E-15 | 2 M1 _ | Cacna1c |
| Gprc5c | 1.17E-19 | 0.719943 | 0.453 | 0.162 | 2.36E-15 | 2 M1 _ | Gprc5c |
| Ppp1r12a | 1.07E-19 | 1.056533 | 0.781 | 0.639 | 2.16E-15 | 2 M1 _ | Ppp1r12a |
| Dgkb | 8.49E-20 | 0.651106 | 0.234 | 0.047 | 1.71E-15 | 2 M1 _ | Dgkb |
| Map1lc3a | 6.25E-20 | 0.772149 | 0.883 | 0.742 | 1.26E-15 | 2 M1 _ | Map1lc3a |
| Esyt1 | 5.23E-20 | 0.858984 | 0.688 | 0.412 | 1.05E-15 | 2 M1 _ | Esyt1 |
| Atp5b | 2.79E-20 | 0.801881 | 0.922 | 0.827 | 5.62E-16 | 2 M1 _ | Atp5b |
| Fblim1 | 2.66E-20 | 0.853634 | 0.445 | 0.162 | 5.36E-16 | 2 M1 _ | Fblim1 |
| Thsd4 | 2.62E-20 | 0.329778 | 0.156 | 0.02 | 5.28E-16 | 2 M1 _ | Thsd4 |
| Mef2a | 2.52E-20 | 0.823569 | 0.844 | 0.651 | 5.08E-16 | 2 M1 _ | Mef2a |
| Fbxl22 | 1.98E-20 | 0.550715 | 0.133 | 0.014 | 3.99E-16 | 2 M1 _ | Fbxl22 |
| Rasal2 | 1.64E-20 | 0.869674 | 0.633 | 0.32 | 3.29E-16 | 2 M1 _ | Rasal2 |
| Lrrc8c | 1.57E-20 | 0.843216 | 0.57 | 0.231 | 3.17E-16 | 2 M1 _ | Lrrc8c |
| Myof | 1.49E-20 | 0.983929 | 0.711 | 0.435 | 3.00E-16 | 2 M1 _ | Myof |
| Speg | 1.42E-20 | 0.457217 | 0.234 | 0.045 | 2.86E-16 | 2 M1 _ | Speg |
| Nrarp | 1.38E-20 | 0.764652 | 0.359 | 0.102 | 2.77E-16 | 2 M1 _ | Nrarp |
| AC149090.1 | 1.10E-20 | 0.973424 | 0.773 | 0.457 | 2.21E-16 | 2 M1 _ | AC149090.1 |
| Jam3 | 9.04E-21 | 0.876718 | 0.711 | 0.389 | 1.82E-16 | 2 M1 _ | Jam3 |
| Myadm1 | 7.57E-21 | 0.921299 | 0.609 | 0.322 | 1.52E-16 | 2 M1 _ | Myadm |
| Pitx1 | 6.98E-21 | 0.97631 | 0.57 | 0.269 | 1.41E-16 | 2 M1 _ | Pitx1 |
| Aaed11 | 6.40E-21 | 0.494291 | 0.281 | 0.062 | 1.29E-16 | 2 M1 _ | Aaed1 |
| Arvcf | 5.86E-21 | 0.662757 | 0.305 | 0.075 | 1.18E-16 | 2 M1 _ | Arvcf |
| Rnd2 | 5.63E-21 | 0.5079 | 0.266 | 0.055 | 1.13E-16 | 2 M1 _ | Rnd2 |
| Lurap1l | 5.34E-21 | 0.604024 | 0.273 | 0.059 | 1.08E-16 | 2 M1 _ | Lurap1l |
| Sssca11 | 4.28E-21 | 0.445642 | 0.18 | 0.026 | 8.63E-17 | 2 M1 _ | Sssca1 |
| Ndrg2 | 4.03E-21 | 0.855456 | 0.766 | 0.476 | 8.11E-17 | 2 M1 _ | Ndrg2 |
| B230219D22Rik | 2.29E-21 | 0.753774 | 0.891 | 0.699 | 4.61E-17 | 2 M1 _ | B230219D22Rik |
| Atp2a21 | 2.01E-21 | 1.227636 | 0.828 | 0.676 | 4.05E-17 | 2 M1 _ | Atp2a2 |

| | | | | | | | |
|---|---|---|---|---|---|---|---|
| AI3141801 | 1.77E-21 | 0.630819 | 0.281 | 0.062 | 3.57E-17 | 2 M1 _ | AI314180 |
| Pparg | 1.71E-21 | 0.565883 | 0.234 | 0.043 | 3.45E-17 | 2 M1 _ | Pparg |
| Ldhb | 1.58E-21 | 0.939394 | 0.594 | 0.279 | 3.18E-17 | 2 M1 _ | Ldhb |
| Fbxo181 | 1.14E-21 | 0.478461 | 0.227 | 0.041 | 2.29E-17 | 2 M1 _ | Fbxo18 |
| Rock1 | 9.72E-22 | 1.047364 | 0.852 | 0.732 | 1.96E-17 | 2 M1 _ | Rock1 |
| Hes1 | 9.49E-22 | 0.98647 | 0.883 | 0.585 | 1.91E-17 | 2 M1 _ | Hes1 |
| Klf91 | 7.38E-22 | 0.888387 | 0.953 | 0.767 | 1.49E-17 | 2 M1 _ | Klf9 |
| Rcan2 | 6.41E-22 | 0.71741 | 0.266 | 0.054 | 1.29E-17 | 2 M1 _ | Rcan2 |
| Fads3 | 6.06E-22 | 0.911933 | 0.5 | 0.192 | 1.22E-17 | 2 M1 _ | Fads3 |
| Tsc22d1 | 5.42E-22 | 0.814544 | 0.969 | 0.827 | 1.09E-17 | 2 M1 _ | Tsc22d1 |
| Csrp1 | 5.22E-22 | 1.522849 | 0.703 | 0.455 | 1.05E-17 | 2 M1 _ | Csrp1 |
| Ntn4 | 4.17E-22 | 1.039817 | 0.414 | 0.139 | 8.40E-18 | 2 M1 _ | Ntn4 |
| Oaz2 | 4.06E-22 | 0.901501 | 0.727 | 0.431 | 8.18E-18 | 2 M1 _ | Oaz2 |
| Arhgdib | 3.71E-22 | 0.936652 | 0.852 | 0.574 | 7.47E-18 | 2 M1 _ | Arhgdib |
| Fam69a1 | 3.37E-22 | 0.582697 | 0.266 | 0.055 | 6.80E-18 | 2 M1 _ | Fam69a |
| Heyl | 3.19E-22 | 1.017439 | 0.664 | 0.334 | 6.43E-18 | 2 M1 _ | Heyl |
| Iqsec3 | 3.01E-22 | 0.260174 | 0.133 | 0.012 | 6.07E-18 | 2 M1 _ | Iqsec3 |
| Bzw2 | 2.10E-22 | 0.876352 | 0.523 | 0.21 | 4.22E-18 | 2 M1 _ | Bzw2 |
| Dync1i2 | 1.93E-22 | 1.148966 | 0.891 | 0.78 | 3.88E-18 | 2 M1 _ | Dync1i2 |
| Rab3a | 1.41E-22 | 0.909939 | 0.43 | 0.143 | 2.85E-18 | 2 M1 _ | Rab3a |
| H3f3b | 1.23E-22 | 0.586794 | 1 | 0.982 | 2.48E-18 | 2 M1 _ | H3f3b |
| Myom1 | 1.17E-22 | 0.342619 | 0.211 | 0.032 | 2.35E-18 | 2 M1 _ | Myom1 |
| Neurl1b | 1.08E-22 | 0.53506 | 0.305 | 0.069 | 2.18E-18 | 2 M1 _ | Neurl1b |
| Ssfa2 | 1.06E-22 | 0.371148 | 0.195 | 0.028 | 2.14E-18 | 2 M1 _ | Ssfa2 |
| H2-M9 | 7.33E-23 | 0.731675 | 0.164 | 0.019 | 1.48E-18 | 2 M1 _ | H2-M9 |
| Ywhaz | 6.11E-23 | 0.918135 | 0.906 | 0.804 | 1.23E-18 | 2 M1 _ | Ywhaz |
| Syne2 | 5.99E-23 | 0.928155 | 0.727 | 0.404 | 1.21E-18 | 2 M1 _ | Syne2 |
| Gm424181 | 4.69E-23 | 0.735081 | 1 | 0.999 | 9.45E-19 | 2 M1 _ | Gm42418 |
| Tubb4a | 4.54E-23 | 0.341579 | 0.164 | 0.019 | 9.14E-19 | 2 M1 _ | Tubb4a |
| Cd151 | 4.38E-23 | 0.962174 | 0.828 | 0.546 | 8.81E-19 | 2 M1 _ | Cd151 |
| Zbtb161 | 3.78E-23 | 0.754359 | 0.344 | 0.086 | 7.62E-19 | 2 M1 _ | Zbtb16 |
| Calm2 | 3.38E-23 | 0.839137 | 0.961 | 0.899 | 6.80E-19 | 2 M1 _ | Calm2 |
| Edil3 | 3.09E-23 | 0.630872 | 0.164 | 0.019 | 6.22E-19 | 2 M1 _ | Edil3 |
| 3110062M04Rik | 2.37E-23 | 0.520177 | 0.195 | 0.028 | 4.78E-19 | 2 M1 _ | 3110062M04Rik |
| Tmem55b1 | 2.07E-23 | 0.420494 | 0.234 | 0.04 | 4.17E-19 | 2 M1 _ | Tmem55b |
| Pdgfc | 1.52E-23 | 0.275373 | 0.117 | 0.009 | 3.07E-19 | 2 M1 _ | Pdgfc |
| Ednra | 1.43E-23 | 0.917024 | 0.758 | 0.438 | 2.88E-19 | 2 M1 _ | Ednra |
| Them6 | 7.52E-24 | 0.650543 | 0.281 | 0.058 | 1.51E-19 | 2 M1 _ | Them6 |
| Ehd2 | 6.61E-24 | 1.116394 | 0.734 | 0.473 | 1.33E-19 | 2 M1 _ | Ehd2 |
| Hey2 | 6.20E-24 | 0.405789 | 0.195 | 0.026 | 1.25E-19 | 2 M1 _ | Hey2 |
| Etv1 | 5.65E-24 | 1.104913 | 0.602 | 0.283 | 1.14E-19 | 2 M1 _ | Etv1 |

| | | | | | | | |
|---|---|---|---|---|---|---|---|
| Fez2 | 5.12E-24 | 0.995564 | 0.695 | 0.362 | 1.03E-19 | 2 M1 _ | Fez2 |
| Vstm4 | 3.80E-24 | 1.104088 | 0.633 | 0.303 | 7.65E-20 | 2 M1 _ | Vstm4 |
| Pkia | 3.63E-24 | 0.58443 | 0.258 | 0.047 | 7.31E-20 | 2 M1 _ | Pkia |
| Fabp7 | 3.60E-24 | 0.270792 | 0.117 | 0.008 | 7.25E-20 | 2 M1 _ | Fabp7 |
| Efhd2 | 3.54E-24 | 0.96251 | 0.664 | 0.309 | 7.12E-20 | 2 M1 _ | Efhd2 |
| Nt5dc2 | 3.20E-24 | 0.84352 | 0.43 | 0.13 | 6.44E-20 | 2 M1 _ | Nt5dc2 |
| Sorbs2 | 2.95E-24 | 1.120658 | 0.531 | 0.174 | 5.93E-20 | 2 M1 _ | Sorbs2 |
| Epn2 | 2.87E-24 | 1.133664 | 0.664 | 0.364 | 5.78E-20 | 2 M1 _ | Epn2 |
| Selenof | 2.68E-24 | 0.87148 | 0.953 | 0.842 | 5.39E-20 | 2 M1 _ | Selenof |
| Kcne4 | 2.12E-24 | 0.689172 | 0.289 | 0.058 | 4.27E-20 | 2 M1 _ | Kcne4 |
| Crim1 | 1.78E-24 | 1.002026 | 0.648 | 0.277 | 3.59E-20 | 2 M1 _ | Crim1 |
| Ehd3 | 1.75E-24 | 0.766696 | 0.43 | 0.13 | 3.52E-20 | 2 M1 _ | Ehd3 |
| Scn3a | 1.31E-24 | 0.495174 | 0.219 | 0.032 | 2.64E-20 | 2 M1 _ | Scn3a |
| Alkal2 | 6.27E-25 | 0.672313 | 0.273 | 0.051 | 1.26E-20 | 2 M1 _ | Alkal2 |
| Stom | 4.72E-25 | 0.758542 | 0.477 | 0.151 | 9.50E-21 | 2 M1 _ | Stom |
| Tspan12 | 4.63E-25 | 0.819723 | 0.469 | 0.139 | 9.32E-21 | 2 M1 _ | Tspan12 |
| Procr | 4.45E-25 | 0.819533 | 0.406 | 0.112 | 8.96E-21 | 2 M1 _ | Procr |
| Pdlim3 | 3.85E-25 | 0.680984 | 0.281 | 0.052 | 7.75E-21 | 2 M1 _ | Pdlim3 |
| Meg3 | 1.89E-25 | 1.236532 | 0.898 | 0.636 | 3.81E-21 | 2 M1 _ | Meg3 |
| BC0033311 | 1.55E-25 | 0.606634 | 0.289 | 0.056 | 3.13E-21 | 2 M1 _ | BC003331 |
| Adcy6 | 1.47E-25 | 0.889404 | 0.539 | 0.194 | 2.97E-21 | 2 M1 _ | Adcy6 |
| Atp1b1 | 1.21E-25 | 1.232862 | 0.398 | 0.11 | 2.44E-21 | 2 M1 _ | Atp1b1 |
| Mum11 | 1.20E-25 | 0.574643 | 0.266 | 0.047 | 2.42E-21 | 2 M1 _ | Mum1 |
| Myl6 | 1.17E-25 | 1.053 | 0.984 | 0.966 | 2.35E-21 | 2 M1 _ | Myl6 |
| Pten | 9.26E-26 | 0.956577 | 0.914 | 0.716 | 1.87E-21 | 2 M1 _ | Pten |
| Lpp1 | 7.52E-26 | 1.348739 | 0.797 | 0.621 | 1.51E-21 | 2 M1 _ | Lpp |
| Ppp1r1a | 6.28E-26 | 0.463633 | 0.188 | 0.022 | 1.26E-21 | 2 M1 _ | Ppp1r1a |
| Prkar1a | 6.08E-26 | 0.91399 | 0.906 | 0.783 | 1.22E-21 | 2 M1 _ | Prkar1a |
| Fam213b | 5.10E-26 | 0.257349 | 0.148 | 0.013 | 1.03E-21 | 2 M1 _ | Fam213b |
| Mgst3 | 4.49E-26 | 0.992226 | 0.562 | 0.216 | 9.04E-22 | 2 M1 _ | Mgst3 |
| Ptprk | 4.10E-26 | 0.839892 | 0.594 | 0.227 | 8.27E-22 | 2 M1 _ | Ptprk |
| Prkar2b | 3.44E-26 | 0.676997 | 0.32 | 0.067 | 6.94E-22 | 2 M1 _ | Prkar2b |
| G0s21 | 3.11E-26 | 1.473209 | 0.797 | 0.507 | 6.25E-22 | 2 M1 _ | G0s2 |
| Acta2 | 2.86E-26 | 3.239459 | 0.539 | 0.21 | 5.76E-22 | 2 M1 _ | Acta2 |
| Actb | 2.82E-26 | 0.657782 | 1 | 0.992 | 5.69E-22 | 2 M1 _ | Actb |
| Lgalsl | 2.45E-26 | 1.267549 | 0.672 | 0.351 | 4.93E-22 | 2 M1 _ | Lgalsl |
| 7-Sep | 1.96E-26 | 0.879213 | 0.969 | 0.896 | 3.96E-22 | 2 M1 _ | 7-Sep |
| Pde1a | 1.91E-26 | 0.853752 | 0.516 | 0.169 | 3.85E-22 | 2 M1 _ | Pde1a |
| Zeb2 | 1.28E-26 | 1.080025 | 0.898 | 0.69 | 2.58E-22 | 2 M1 _ | Zeb2 |
| Pde1b | 1.19E-26 | 0.508044 | 0.258 | 0.042 | 2.39E-22 | 2 M1 _ | Pde1b |
| Mrvi1 | 1.17E-26 | 1.035748 | 0.562 | 0.212 | 2.36E-22 | 2 M1 _ | Mrvi1 |

| | | | | | | | |
|---|---|---|---|---|---|---|---|
| Gm10687 | 1.05E-26 | 0.430406 | 0.234 | 0.034 | 2.10E-22 | 2 M1 _ | Gm10687 |
| Kcnmb1 | 5.21E-27 | 0.357619 | 0.125 | 0.008 | 1.05E-22 | 2 M1 _ | Kcnmb1 |
| Lars21 | 4.01E-27 | 0.86389 | 0.969 | 0.805 | 8.08E-23 | 2 M1 _ | Lars2 |
| Ccl111 | 3.13E-27 | 1.806674 | 0.734 | 0.424 | 6.31E-23 | 2 M1 _ | Ccl11 |
| Galnt16 | 2.80E-27 | 0.805958 | 0.422 | 0.114 | 5.64E-23 | 2 M1 _ | Galnt16 |
| Chst1 | 2.18E-27 | 0.772908 | 0.398 | 0.099 | 4.38E-23 | 2 M1 _ | Chst1 |
| Pcdh1 | 2.05E-27 | 0.791692 | 0.516 | 0.161 | 4.12E-23 | 2 M1 _ | Pcdh1 |
| Malat1 | 1.98E-27 | 0.834549 | 0.984 | 0.998 | 3.98E-23 | 2 M1 _ | Malat1 |
| Sod3 | 1.94E-27 | 1.110127 | 0.641 | 0.279 | 3.91E-23 | 2 M1 _ | Sod3 |
| Stum | 1.67E-27 | 0.315224 | 0.156 | 0.014 | 3.37E-23 | 2 M1 _ | Stum |
| Aatk | 1.12E-27 | 0.426275 | 0.227 | 0.03 | 2.25E-23 | 2 M1 _ | Aatk |
| Klhl23 | 1.06E-27 | 0.767669 | 0.406 | 0.101 | 2.13E-23 | 2 M1 _ | Klhl23 |
| Oraov1 | 8.36E-28 | 0.4945 | 0.195 | 0.023 | 1.68E-23 | 2 M1 _ | Oraov1 |
| Crip1 | 8.17E-28 | 1.571276 | 0.992 | 0.948 | 1.65E-23 | 2 M1 _ | Crip1 |
| Atp2a3 | 6.63E-28 | 0.792075 | 0.359 | 0.081 | 1.33E-23 | 2 M1 _ | Atp2a3 |
| Ift43 | 4.21E-28 | 1.140894 | 0.688 | 0.332 | 8.47E-24 | 2 M1 _ | Ift43 |
| Btbd3 | 3.63E-28 | 1.147943 | 0.648 | 0.273 | 7.31E-24 | 2 M1 _ | Btbd3 |
| Eps81 | 2.59E-28 | 1.173237 | 0.75 | 0.409 | 5.22E-24 | 2 M1 _ | Eps8 |
| Arhgap10 | 1.90E-28 | 1.115156 | 0.727 | 0.368 | 3.82E-24 | 2 M1 _ | Arhgap10 |
| Dstn | 1.54E-28 | 1.473502 | 0.969 | 0.812 | 3.09E-24 | 2 M1 _ | Dstn |
| Cog7 | 1.36E-28 | 1.046216 | 0.539 | 0.183 | 2.73E-24 | 2 M1 _ | Cog7 |
| Aldoa | 7.24E-29 | 0.998356 | 0.961 | 0.841 | 1.46E-24 | 2 M1 _ | Aldoa |
| Mamdc2 | 6.31E-29 | 0.399164 | 0.219 | 0.027 | 1.27E-24 | 2 M1 _ | Mamdc2 |
| Slc38a11 | 2.04E-29 | 0.545744 | 0.219 | 0.026 | 4.11E-25 | 2 M1 _ | Slc38a11 |
| Ppm1e | 1.63E-29 | 0.467939 | 0.203 | 0.022 | 3.27E-25 | 2 M1 _ | Ppm1e |
| 2810474O19Rik1 | 1.53E-29 | 0.661513 | 0.336 | 0.065 | 3.08E-25 | 2 M1 _ | 2810474O19Rik |
| Fam220a1 | 1.41E-29 | 0.659189 | 0.281 | 0.047 | 2.83E-25 | 2 M1 _ | Fam220a |
| Inpp4b | 1.00E-29 | 1.237421 | 0.672 | 0.265 | 2.02E-25 | 2 M1 _ | Inpp4b |
| Coro1b1 | 8.13E-30 | 0.770939 | 0.383 | 0.089 | 1.64E-25 | 2 M1 _ | Coro1b |
| Mpp2 | 7.53E-30 | 0.394106 | 0.203 | 0.022 | 1.52E-25 | 2 M1 _ | Mpp2 |
| Map3k7cl | 7.49E-30 | 0.586353 | 0.133 | 0.008 | 1.51E-25 | 2 M1 _ | Map3k7cl |
| Vcl | 4.44E-30 | 1.215865 | 0.781 | 0.463 | 8.95E-26 | 2 M1 _ | Vcl |
| mt-Atp81 | 2.09E-30 | 1.162232 | 0.852 | 0.593 | 4.21E-26 | 2 M1 _ | mt-Atp8 |
| Cd36 | 1.97E-30 | 0.907701 | 0.602 | 0.182 | 3.97E-26 | 2 M1 _ | Cd36 |
| Ccdc160 | 1.78E-30 | 0.413215 | 0.211 | 0.023 | 3.58E-26 | 2 M1 _ | Ccdc160 |
| Cdkn1a | 1.33E-30 | 0.941564 | 0.555 | 0.168 | 2.68E-26 | 2 M1 _ | Cdkn1a |
| Pde4b | 1.05E-30 | 0.976253 | 0.453 | 0.12 | 2.11E-26 | 2 M1 _ | Pde4b |
| Tmem1001 | 8.92E-31 | 1.180328 | 0.672 | 0.264 | 1.80E-26 | 2 M1 _ | Tmem100 |
| Mylk | 7.90E-31 | 1.65274 | 0.891 | 0.624 | 1.59E-26 | 2 M1 _ | Mylk |
| Errfi11 | 6.47E-31 | 1.444423 | 0.898 | 0.614 | 1.30E-26 | 2 M1 _ | Errfi1 |
| Snrk | 6.46E-31 | 0.98975 | 0.938 | 0.591 | 1.30E-26 | 2 M1 _ | Snrk |

| | | | | | | | |
|---|---|---|---|---|---|---|---|
| Emid1 | 3.20E-31 | 1.140546 | 0.617 | 0.219 | 6.45E-27 | 2 M1 _ | Emid1 |
| Plekhg3 | 2.67E-31 | 0.914134 | 0.453 | 0.12 | 5.39E-27 | 2 M1 _ | Plekhg3 |
| Ptms | 2.29E-31 | 0.990743 | 0.969 | 0.902 | 4.61E-27 | 2 M1 _ | Ptms |
| Flna | 1.77E-31 | 1.773493 | 0.844 | 0.633 | 3.57E-27 | 2 M1 _ | Flna |
| Hcn2 | 6.53E-32 | 0.497362 | 0.188 | 0.017 | 1.32E-27 | 2 M1 _ | Hcn2 |
| Ccdc68 | 4.65E-32 | 0.490954 | 0.266 | 0.035 | 9.36E-28 | 2 M1 _ | Ccdc68 |
| Insc | 4.22E-32 | 0.537401 | 0.195 | 0.019 | 8.50E-28 | 2 M1 _ | Insc |
| Mprip | 3.12E-32 | 1.222945 | 0.875 | 0.708 | 6.28E-28 | 2 M1 _ | Mprip |
| Rtn4rl1 | 9.16E-33 | 0.812961 | 0.344 | 0.063 | 1.84E-28 | 2 M1 _ | Rtn4rl1 |
| Zcchc111 | 3.92E-33 | 0.721943 | 0.336 | 0.059 | 7.89E-29 | 2 M1 _ | Zcchc11 |
| Foxs1 | 3.38E-33 | 1.158727 | 0.789 | 0.389 | 6.81E-29 | 2 M1 _ | Foxs1 |
| Nrip2 | 2.18E-33 | 0.856172 | 0.266 | 0.034 | 4.39E-29 | 2 M1 _ | Nrip2 |
| Dmd | 1.99E-33 | 1.227925 | 0.633 | 0.236 | 4.00E-29 | 2 M1 _ | Dmd |
| Cacna2d1 | 1.63E-33 | 1.249383 | 0.75 | 0.372 | 3.28E-29 | 2 M1 _ | Cacna2d1 |
| Adgrv1 | 1.51E-33 | 0.39922 | 0.141 | 0.008 | 3.05E-29 | 2 M1 _ | Adgrv1 |
| Zcchc61 | 1.35E-33 | 0.805527 | 0.406 | 0.082 | 2.71E-29 | 2 M1 _ | Zcchc6 |
| Fermt2 | 1.30E-33 | 1.113236 | 0.977 | 0.764 | 2.62E-29 | 2 M1 _ | Fermt2 |
| Pde4c | 1.18E-33 | 0.402068 | 0.172 | 0.013 | 2.38E-29 | 2 M1 _ | Pde4c |
| Skiv2l21 | 8.98E-34 | 0.643734 | 0.305 | 0.047 | 1.81E-29 | 2 M1 _ | Skiv2l2 |
| Tmem35a | 7.51E-34 | 0.445218 | 0.18 | 0.014 | 1.51E-29 | 2 M1 _ | Tmem35a |
| Ebf1 | 4.67E-34 | 1.172025 | 0.961 | 0.78 | 9.40E-30 | 2 M1 _ | Ebf1 |
| Synpo2 | 3.26E-34 | 1.170751 | 0.477 | 0.122 | 6.57E-30 | 2 M1 _ | Synpo2 |
| Atpif1 | 2.18E-34 | 1.035382 | 0.961 | 0.767 | 4.40E-30 | 2 M1 _ | Atpif1 |
| Cbfa2t3 | 1.64E-34 | 1.24156 | 0.656 | 0.228 | 3.30E-30 | 2 M1 _ | Cbfa2t3 |
| Trpv2 | 1.38E-34 | 0.670413 | 0.352 | 0.06 | 2.79E-30 | 2 M1 _ | Trpv2 |
| Slc25a4 | 1.18E-34 | 0.980632 | 0.977 | 0.854 | 2.37E-30 | 2 M1 _ | Slc25a4 |
| Trove21 | 9.28E-35 | 0.831406 | 0.375 | 0.069 | 1.87E-30 | 2 M1 _ | Trove2 |
| 4632415L05Rik1 | 1.91E-35 | 0.550284 | 0.227 | 0.024 | 3.84E-31 | 2 M1 _ | 4632415L05Rik |
| Adap2 | 5.26E-36 | 1.161319 | 0.656 | 0.229 | 1.06E-31 | 2 M1 _ | Adap2 |
| Tbx2 | 4.57E-36 | 1.365148 | 0.688 | 0.274 | 9.21E-32 | 2 M1 _ | Tbx2 |
| Pyurf1 | 4.08E-36 | 0.568963 | 0.273 | 0.035 | 8.22E-32 | 2 M1 _ | Pyurf |
| Lin7a | 3.50E-36 | 1.100281 | 0.734 | 0.292 | 7.04E-32 | 2 M1 _ | Lin7a |
| Fam81a | 2.71E-36 | 0.633114 | 0.219 | 0.021 | 5.45E-32 | 2 M1 _ | Fam81a |
| Colec11 | 1.99E-36 | 1.172251 | 0.32 | 0.046 | 4.00E-32 | 2 M1 _ | Colec11 |
| Tbx3 | 6.50E-37 | 1.023181 | 0.516 | 0.127 | 1.31E-32 | 2 M1 _ | Tbx3 |
| Gm26526 | 3.24E-37 | 0.439713 | 0.211 | 0.019 | 6.52E-33 | 2 M1 _ | Gm26526 |
| Angptl2 | 1.91E-37 | 1.582118 | 0.75 | 0.339 | 3.85E-33 | 2 M1 _ | Angptl2 |
| Bcam | 7.89E-38 | 1.218828 | 0.805 | 0.304 | 1.59E-33 | 2 M1 _ | Bcam |
| Ngf | 6.16E-38 | 0.935224 | 0.516 | 0.128 | 1.24E-33 | 2 M1 _ | Ngf |
| Tns1 | 5.69E-38 | 1.446716 | 0.852 | 0.516 | 1.15E-33 | 2 M1 _ | Tns1 |
| Khdrbs3 | 5.03E-38 | 1.05663 | 0.539 | 0.142 | 1.01E-33 | 2 M1 _ | Khdrbs3 |

| | | | | | | | |
|---|---|---|---|---|---|---|---|
| Raph1 | 4.91E-38 | 1.454774 | 0.758 | 0.332 | 9.89E-34 | 2 M1 _ | Raph1 |
| Hspb2 | 1.95E-38 | 1.014713 | 0.445 | 0.088 | 3.93E-34 | 2 M1 _ | Hspb2 |
| Rbpms2 | 1.55E-38 | 0.88904 | 0.438 | 0.085 | 3.12E-34 | 2 M1 _ | Rbpms2 |
| Slc24a3 | 8.66E-39 | 0.871233 | 0.375 | 0.063 | 1.74E-34 | 2 M1 _ | Slc24a3 |
| Parm1 | 3.93E-39 | 1.069676 | 0.484 | 0.105 | 7.91E-35 | 2 M1 _ | Parm1 |
| Pdgfrb | 2.35E-39 | 1.381282 | 1 | 0.826 | 4.74E-35 | 2 M1 _ | Pdgfrb |
| Cap2 | 2.09E-39 | 0.742337 | 0.305 | 0.04 | 4.21E-35 | 2 M1 _ | Cap2 |
| Fam103a11 | 1.08E-39 | 0.680041 | 0.344 | 0.052 | 2.18E-35 | 2 M1 _ | Fam103a1 |
| Tmem94 | 7.97E-40 | 1.084108 | 0.484 | 0.109 | 1.61E-35 | 2 M1 _ | Tmem94 |
| Rarres21 | 3.66E-40 | 1.424075 | 0.969 | 0.675 | 7.38E-36 | 2 M1 _ | Rarres2 |
| Cspg4 | 3.09E-40 | 1.35627 | 0.766 | 0.311 | 6.22E-36 | 2 M1 _ | Cspg4 |
| Map7d2 | 1.91E-40 | 0.590921 | 0.258 | 0.025 | 3.85E-36 | 2 M1 _ | Map7d2 |
| Epas1 | 9.21E-41 | 1.513304 | 0.906 | 0.509 | 1.86E-36 | 2 M1 _ | Epas1 |
| Rgs6 | 9.16E-41 | 0.689862 | 0.273 | 0.029 | 1.84E-36 | 2 M1 _ | Rgs6 |
| Rbpms | 4.17E-41 | 1.289905 | 0.953 | 0.706 | 8.40E-37 | 2 M1 _ | Rbpms |
| Ccdc141 | 1.89E-41 | 0.780415 | 0.438 | 0.078 | 3.80E-37 | 2 M1 _ | Ccdc141 |
| Dtx3 | 1.63E-41 | 1.379761 | 0.844 | 0.418 | 3.29E-37 | 2 M1 _ | Dtx3 |
| P2rx1 | 1.44E-41 | 0.574161 | 0.141 | 0.005 | 2.90E-37 | 2 M1 _ | P2rx1 |
| Actn4 | 1.03E-41 | 1.436316 | 0.938 | 0.707 | 2.08E-37 | 2 M1 _ | Actn4 |
| Timp3 | 6.41E-42 | 1.438321 | 0.984 | 0.781 | 1.29E-37 | 2 M1 _ | Timp3 |
| Wfdc1 | 6.14E-42 | 1.042753 | 0.414 | 0.074 | 1.24E-37 | 2 M1 _ | Wfdc1 |
| D10Jhu81e1 | 5.17E-42 | 0.647206 | 0.289 | 0.034 | 1.04E-37 | 2 M1 _ | D10Jhu81e |
| Limd1 | 1.96E-42 | 1.459066 | 0.812 | 0.38 | 3.94E-38 | 2 M1 _ | Limd1 |
| 4-Sep | 8.13E-43 | 1.513959 | 0.961 | 0.705 | 1.64E-38 | 2 M1 _ | 4-Sep |
| Cfap69 | 6.51E-43 | 0.63289 | 0.203 | 0.014 | 1.31E-38 | 2 M1 _ | Cfap69 |
| Mtss1l | 4.33E-43 | 0.488509 | 0.195 | 0.012 | 8.72E-39 | 2 M1 _ | Mtss1l |
| Gcnt2 | 2.55E-43 | 1.203672 | 0.664 | 0.18 | 5.14E-39 | 2 M1 _ | Gcnt2 |
| 1500011K16Rik1 | 1.67E-43 | 0.643623 | 0.305 | 0.036 | 3.36E-39 | 2 M1 _ | 1500011K16Rik |
| Sgip1 | 1.43E-43 | 0.830152 | 0.43 | 0.07 | 2.88E-39 | 2 M1 _ | Sgip1 |
| Olfr558 | 1.06E-43 | 1.380104 | 0.555 | 0.118 | 2.14E-39 | 2 M1 _ | Olfr558 |
| Tesc | 8.00E-44 | 1.499826 | 0.445 | 0.079 | 1.61E-39 | 2 M1 _ | Tesc |
| Itga1 | 6.58E-44 | 1.510149 | 0.977 | 0.79 | 1.32E-39 | 2 M1 _ | Itga1 |
| Slco3a1 | 5.60E-44 | 1.336493 | 0.789 | 0.296 | 1.13E-39 | 2 M1 _ | Slco3a1 |
| Tpm1 | 2.34E-44 | 2.172516 | 0.938 | 0.711 | 4.72E-40 | 2 M1 _ | Tpm1 |
| Arhgap42 | 2.33E-44 | 1.215594 | 0.641 | 0.178 | 4.69E-40 | 2 M1 _ | Arhgap42 |
| Pcdh19 | 1.55E-44 | 1.551265 | 0.883 | 0.439 | 3.13E-40 | 2 M1 _ | Pcdh19 |
| Aspn | 1.18E-44 | 1.572581 | 0.922 | 0.498 | 2.38E-40 | 2 M1 _ | Aspn |
| Gm13889 | 1.15E-44 | 1.448109 | 0.93 | 0.475 | 2.31E-40 | 2 M1 _ | Gm13889 |
| Gm13861 | 2.18E-45 | 0.823417 | 0.352 | 0.043 | 4.39E-41 | 2 M1 _ | Gm13861 |
| Lrrc8b | 9.68E-46 | 0.745588 | 0.328 | 0.038 | 1.95E-41 | 2 M1 _ | Lrrc8b |
| Ccdc3 | 8.83E-46 | 0.641633 | 0.312 | 0.034 | 1.78E-41 | 2 M1 _ | Ccdc3 |

| | | | | | | | |
|---|---|---|---|---|---|---|---|
| Cav1 | 7.86E-46 | 1.371781 | 0.82 | 0.283 | 1.58E-41 | 2 M1 _ | Cav1 |
| Trpc6 | 6.42E-46 | 0.703711 | 0.375 | 0.049 | 1.29E-41 | 2 M1 _ | Trpc6 |
| Mtcl1 | 2.56E-46 | 0.548067 | 0.242 | 0.019 | 5.16E-42 | 2 M1 _ | Mtcl1 |
| Ano4 | 1.40E-46 | 0.484588 | 0.172 | 0.007 | 2.83E-42 | 2 M1 _ | Ano4 |
| Id4 | 5.40E-47 | 1.661954 | 0.609 | 0.154 | 1.09E-42 | 2 M1 _ | Id4 |
| Rhob1 | 3.59E-47 | 1.787128 | 0.938 | 0.599 | 7.23E-43 | 2 M1 _ | Rhob |
| Arhgef17 | 1.97E-47 | 1.415795 | 0.773 | 0.279 | 3.96E-43 | 2 M1 _ | Arhgef17 |
| Gucy1a2 | 9.23E-48 | 0.970839 | 0.523 | 0.105 | 1.86E-43 | 2 M1 _ | Gucy1a2 |
| Cryab | 3.75E-48 | 1.510929 | 0.914 | 0.474 | 7.54E-44 | 2 M1 _ | Cryab |
| Myh11 | 3.68E-48 | 3.195112 | 0.5 | 0.096 | 7.41E-44 | 2 M1 _ | Myh11 |
| Esam | 2.34E-48 | 1.276176 | 0.914 | 0.303 | 4.72E-44 | 2 M1 _ | Esam |
| Cald1 | 2.09E-48 | 1.453937 | 1 | 0.866 | 4.20E-44 | 2 M1 _ | Cald1 |
| Kcnh1 | 4.41E-49 | 0.685765 | 0.211 | 0.012 | 8.89E-45 | 2 M1 _ | Kcnh1 |
| Hrct1 | 8.26E-50 | 1.286075 | 0.594 | 0.134 | 1.66E-45 | 2 M1 _ | Hrct1 |
| Fzd4 | 7.52E-51 | 1.462537 | 0.859 | 0.319 | 1.51E-46 | 2 M1 _ | Fzd4 |
| Casq2 | 5.11E-51 | 1.177449 | 0.367 | 0.043 | 1.03E-46 | 2 M1 _ | Casq2 |
| Tspan15 | 1.91E-51 | 1.32374 | 0.633 | 0.146 | 3.84E-47 | 2 M1 _ | Tspan15 |
| Fam162a | 7.40E-52 | 2.065189 | 0.93 | 0.597 | 1.49E-47 | 2 M1 _ | Fam162a |
| Jph2 | 7.36E-52 | 0.685646 | 0.297 | 0.026 | 1.48E-47 | 2 M1 _ | Jph2 |
| Rdh5 | 3.49E-53 | 1.142679 | 0.508 | 0.087 | 7.02E-49 | 2 M1 _ | Rdh5 |
| Myo1b | 5.62E-54 | 1.700577 | 0.938 | 0.476 | 1.13E-49 | 2 M1 _ | Myo1b |
| Rasl12 | 3.05E-54 | 1.012489 | 0.578 | 0.103 | 6.15E-50 | 2 M1 _ | Rasl12 |
| Fam208a1 | 2.45E-54 | 0.982657 | 0.453 | 0.066 | 4.94E-50 | 2 M1 _ | Fam208a |
| Kcna5 | 1.62E-54 | 0.598335 | 0.25 | 0.016 | 3.27E-50 | 2 M1 _ | Kcna5 |
| Sdc1 | 3.91E-55 | 1.748314 | 0.688 | 0.189 | 7.86E-51 | 2 M1 _ | Sdc1 |
| Dirc21 | 2.48E-55 | 0.821306 | 0.359 | 0.039 | 4.99E-51 | 2 M1 _ | Dirc2 |
| Ebf2 | 2.04E-56 | 1.296978 | 0.672 | 0.151 | 4.12E-52 | 2 M1 _ | Ebf2 |
| Kcnk3 | 1.18E-56 | 0.806341 | 0.391 | 0.043 | 2.37E-52 | 2 M1 _ | Kcnk3 |
| Itgb1 | 1.11E-56 | 1.458128 | 0.984 | 0.895 | 2.23E-52 | 2 M1 _ | Itgb1 |
| Npy1r | 8.98E-57 | 1.309514 | 0.547 | 0.096 | 1.81E-52 | 2 M1 _ | Npy1r |
| Ndufa4l2 | 3.99E-57 | 1.96895 | 0.977 | 0.6 | 8.04E-53 | 2 M1 _ | Ndufa4l2 |
| Carmn | 3.68E-57 | 1.291576 | 0.75 | 0.179 | 7.41E-53 | 2 M1 _ | Carmn |
| AY0361181 | 3.53E-57 | 2.21072 | 1 | 0.542 | 7.11E-53 | 2 M1 _ | AY036118 |
| Pdlim1 | 3.27E-57 | 1.280796 | 0.789 | 0.231 | 6.59E-53 | 2 M1 _ | Pdlim1 |
| Dusp4 | 2.09E-57 | 0.686418 | 0.336 | 0.03 | 4.22E-53 | 2 M1 _ | Dusp4 |
| Rem1 | 1.71E-57 | 1.287915 | 0.664 | 0.141 | 3.43E-53 | 2 M1 _ | Rem1 |
| Vtn | 1.28E-57 | 1.64695 | 0.758 | 0.197 | 2.58E-53 | 2 M1 _ | Vtn |
| Tagln | 1.18E-57 | 3.570739 | 0.695 | 0.178 | 2.38E-53 | 2 M1 _ | Tagln |
| Hopx | 6.89E-58 | 1.917124 | 0.812 | 0.269 | 1.39E-53 | 2 M1 _ | Hopx |
| Tbc1d1 | 1.53E-58 | 1.431149 | 0.852 | 0.283 | 3.08E-54 | 2 M1 _ | Tbc1d1 |
| Cpe | 1.31E-58 | 1.903571 | 0.906 | 0.42 | 2.64E-54 | 2 M1 _ | Cpe |

| | | | | | | | |
|---|---|---|---|---|---|---|---|
| Spaar | 1.01E-58 | 0.854414 | 0.359 | 0.035 | 2.04E-54 | 2 M1 _ | Spaar |
| Palm2 | 9.02E-59 | 0.358574 | 0.133 | 0.001 | 1.82E-54 | 2 M1 _ | Palm2 |
| Gjc1 | 9.00E-59 | 1.846177 | 0.922 | 0.421 | 1.81E-54 | 2 M1 _ | Gjc1 |
| 1700020I14Rik1 | 7.08E-59 | 1.190925 | 0.523 | 0.085 | 1.43E-54 | 2 M1 _ | 1700020I14Rik |
| Steap41 | 4.15E-59 | 2.154556 | 0.938 | 0.423 | 8.35E-55 | 2 M1 _ | Steap4 |
| Cdk15 | 5.59E-61 | 0.508628 | 0.211 | 0.008 | 1.12E-56 | 2 M1 _ | Cdk15 |
| Pdgfa | 2.41E-62 | 1.621745 | 0.789 | 0.226 | 4.86E-58 | 2 M1 _ | Pdgfa |
| Gng11 | 1.12E-62 | 1.900109 | 1 | 0.81 | 2.25E-58 | 2 M1 _ | Gng11 |
| Rasl11a | 4.66E-63 | 1.415821 | 0.797 | 0.2 | 9.39E-59 | 2 M1 _ | Rasl11a |
| Kcnj8 | 1.59E-64 | 2.73249 | 0.805 | 0.22 | 3.20E-60 | 2 M1 _ | Kcnj8 |
| Cobll1 | 1.41E-64 | 1.218083 | 0.547 | 0.084 | 2.84E-60 | 2 M1 _ | Cobll1 |
| Tmem52 | 3.12E-65 | 0.968366 | 0.492 | 0.065 | 6.28E-61 | 2 M1 _ | Tmem5 |
| 1810011H11Rik | 7.03E-66 | 0.663417 | 0.188 | 0.004 | 1.42E-61 | 2 M1 _ | 1810011H11Rik |
| Mfge8 | 5.18E-66 | 2.266503 | 0.984 | 0.793 | 1.04E-61 | 2 M1 _ | Mfge8 |
| Fam96b1 | 1.63E-66 | 0.919315 | 0.445 | 0.051 | 3.29E-62 | 2 M1 _ | Fam96b |
| Filip1l | 1.20E-66 | 2.095781 | 0.859 | 0.279 | 2.42E-62 | 2 M1 _ | Filip1l |
| Gucy1a1 | 1.77E-67 | 1.984181 | 0.953 | 0.353 | 3.57E-63 | 2 M1 _ | Gucy1a1 |
| Mcam | 9.88E-68 | 1.51007 | 0.648 | 0.121 | 1.99E-63 | 2 M1 _ | Mcam |
| 1810022K09Rik1 | 6.83E-68 | 1.054988 | 0.547 | 0.079 | 1.38E-63 | 2 M1 _ | 1810022K09Rik |
| Gucy1b1 | 2.61E-68 | 1.923912 | 0.945 | 0.333 | 5.26E-64 | 2 M1 _ | Gucy1b1 |
| Bcar3 | 3.50E-69 | 0.870365 | 0.414 | 0.039 | 7.04E-65 | 2 M1 _ | Bcar3 |
| Tinagl1 | 2.35E-69 | 1.79119 | 0.945 | 0.273 | 4.74E-65 | 2 M1 _ | Tinagl1 |
| Plxnb1 | 3.03E-70 | 0.730129 | 0.336 | 0.023 | 6.10E-66 | 2 M1 _ | Plxnb1 |
| Dgkg | 2.77E-71 | 0.798869 | 0.336 | 0.022 | 5.59E-67 | 2 M1 _ | Dgkg |
| Slc12a2 | 1.94E-71 | 2.290456 | 0.852 | 0.245 | 3.91E-67 | 2 M1 _ | Slc12a2 |
| Aoc31 | 1.58E-71 | 1.707271 | 0.75 | 0.172 | 3.19E-67 | 2 M1 _ | Aoc3 |
| Atp1b2 | 1.49E-71 | 1.722282 | 0.875 | 0.263 | 3.00E-67 | 2 M1 _ | Atp1b2 |
| Lmod1 | 6.63E-72 | 1.432205 | 0.555 | 0.073 | 1.33E-67 | 2 M1 _ | Lmod1 |
| Adap2os | 1.09E-72 | 1.486627 | 0.703 | 0.134 | 2.19E-68 | 2 M1 _ | Adap2os |
| Aes1 | 3.03E-73 | 1.4382 | 0.664 | 0.108 | 6.11E-69 | 2 M1 _ | Aes |
| Cystm1 | 1.63E-73 | 1.807844 | 0.914 | 0.286 | 3.29E-69 | 2 M1 _ | Cystm1 |
| Abcc9 | 9.17E-74 | 2.891062 | 0.906 | 0.302 | 1.85E-69 | 2 M1 _ | Abcc9 |
| 2410015M20Rik1 | 9.09E-74 | 1.191395 | 0.602 | 0.091 | 1.83E-69 | 2 M1 _ | 2410015M20Rik |
| Chn1 | 3.27E-74 | 1.110782 | 0.508 | 0.057 | 6.60E-70 | 2 M1 _ | Chn1 |
| Atp5o.11 | 2.19E-74 | 1.301331 | 0.625 | 0.097 | 4.41E-70 | 2 M1 _ | Atp5o.1 |
| Mnd1 | 1.65E-74 | 1.668482 | 0.633 | 0.101 | 3.33E-70 | 2 M1 _ | Mnd1 |
| 2010111I01Rik1 | 1.03E-74 | 0.982252 | 0.492 | 0.056 | 2.07E-70 | 2 M1 _ | 2010111I01Rik |
| Des | 3.85E-76 | 1.917845 | 0.844 | 0.2 | 7.76E-72 | 2 M1 _ | Des |
| Pde3a | 5.89E-77 | 1.661412 | 0.688 | 0.12 | 1.19E-72 | 2 M1 _ | Pde3a |
| Gja4 | 6.38E-78 | 1.685003 | 0.93 | 0.216 | 1.28E-73 | 2 M1 _ | Gja4 |
| Mapt | 8.25E-79 | 1.150495 | 0.484 | 0.048 | 1.66E-74 | 2 M1 _ | Mapt |

| | | | | | | | |
|---|---|---|---|---|---|---|---|
| Pde5a | 4.93E-82 | 1.817907 | 0.836 | 0.185 | 9.94E-78 | 2 M1 _ | Pde5a |
| Slc4a8 | 1.81E-83 | 1.235719 | 0.484 | 0.044 | 3.65E-79 | 2 M1 _ | Slc4a8 |
| Agtr1a | 5.19E-84 | 1.629851 | 0.641 | 0.09 | 1.05E-79 | 2 M1 _ | Agtr1a |
| Slc2a4 | 4.89E-85 | 1.18378 | 0.586 | 0.067 | 9.85E-81 | 2 M1 _ | Slc2a4 |
| Ednrb | 3.20E-85 | 3.056014 | 0.938 | 0.281 | 6.43E-81 | 2 M1 _ | Ednrb |
| Rasgrp2 | 4.33E-86 | 2.114367 | 0.891 | 0.221 | 8.72E-82 | 2 M1 _ | Rasgrp2 |
| Daam2 | 2.83E-87 | 1.526932 | 0.75 | 0.127 | 5.69E-83 | 2 M1 _ | Daam2 |
| Tpm2 | 2.64E-87 | 3.315021 | 0.984 | 0.372 | 5.31E-83 | 2 M1 _ | Tpm2 |
| Itga7 | 6.00E-88 | 1.277585 | 0.719 | 0.101 | 1.21E-83 | 2 M1 _ | Itga7 |
| Sncg | 1.76E-88 | 2.141263 | 0.891 | 0.191 | 3.55E-84 | 2 M1 _ | Sncg |
| Ephx3 | 1.36E-89 | 1.627182 | 0.594 | 0.066 | 2.73E-85 | 2 M1 _ | Ephx3 |
| Ano1 | 5.26E-91 | 1.282782 | 0.633 | 0.072 | 1.06E-86 | 2 M1 _ | Ano1 |
| Myl9 | 4.81E-91 | 3.052524 | 1 | 0.338 | 9.69E-87 | 2 M1 _ | Myl9 |
| 2010107E04Rik1 | 1.89E-91 | 1.48712 | 0.688 | 0.098 | 3.80E-87 | 2 M1 _ | 2010107E04Rik |
| Minos11 | 2.39E-98 | 1.461851 | 0.711 | 0.095 | 4.81E-94 | 2 M1 _ | Minos1 |
| Art3 | 1.15E-98 | 2.047923 | 0.766 | 0.115 | 2.32E-94 | 2 M1 _ | Art3 |
| Rgs5 | 1.02E-98 | 2.838544 | 0.984 | 0.217 | 2.06E-94 | 2 M1 _ | Rgs5 |
| 6430573F11Rik | 1.34E-99 | 0.657038 | 0.266 | 0.005 | 2.69E-95 | 2 M1 _ | 6430573F11Rik |
| Ajuba | 2.30E-106 | 1.371846 | 0.672 | 0.075 | 4.64E-102 | 2 M1 _ | Ajuba |
| 1110008F13Rik1 | 6.55E-107 | 1.336702 | 0.633 | 0.066 | 1.32E-102 | 2 M1 _ | 1110008F13Rik |
| Notch3 | 1.35E-108 | 2.5046 | 0.977 | 0.21 | 2.72E-104 | 2 M1 _ | Notch3 |
| Lgi1 | 4.46E-110 | 2.011954 | 0.711 | 0.081 | 8.99E-106 | 2 M1 _ | Lgi1 |
| Serpini1 | 1.45E-110 | 2.009952 | 0.828 | 0.12 | 2.92E-106 | 2 M1 _ | Serpini1 |
| Rgs4 | 4.77E-114 | 1.955958 | 0.852 | 0.115 | 9.60E-110 | 2 M1 _ | Rgs4 |
| Higd1b | 7.34E-119 | 2.420674 | 0.945 | 0.152 | 1.48E-114 | 2 M1 _ | Higd1b |
| Fam212b | 2.21E-121 | 0.830621 | 0.328 | 0.006 | 4.44E-117 | 2 M1 _ | Fam212b |
| Usmg51 | 1.55E-123 | 1.748283 | 0.789 | 0.096 | 3.12E-119 | 2 M1 _ | Usmg5 |
| Pde8b | 2.81E-127 | 1.490355 | 0.609 | 0.043 | 5.67E-123 | 2 M1 _ | Pde8b |
| Cpm | 3.82E-129 | 2.193179 | 0.758 | 0.078 | 7.70E-125 | 2 M1 _ | Cpm |
| Cox4i2 | 1.08E-142 | 2.590002 | 0.953 | 0.13 | 2.18E-138 | 2 M1 _ | Cox4i2 |
| Gm14005 | 7.44E-149 | 1.112219 | 0.453 | 0.013 | 1.50E-144 | 2 M1 _ | Gm14005 |
| Elmod1 | 1.34E-158 | 1.841322 | 0.609 | 0.029 | 2.71E-154 | 2 M1 _ | Elmod1 |
| Ppp1r14a | 6.04E-162 | 2.511993 | 0.922 | 0.103 | 1.22E-157 | 2 M1 _ | Ppp1r14a |
| Ace2 | 9.49E-164 | 2.255285 | 0.719 | 0.046 | 1.91E-159 | 2 M1 _ | Ace2 |
| Tusc5 | 1.21E-207 | 1.19777 | 0.469 | 0.003 | 2.44E-203 | 2 M1 _ | Tusc5 |
| Inpp5k | 2.42E-06 | 0.303866 | 0.273 | 0.162 | 0.048746 | 3 L1 _ | Inpp5k |
| Cct6a | 2.33E-06 | 0.276628 | 0.596 | 0.446 | 0.046923 | 3 L1 _ | Cct6a |
| Sun1 | 2.25E-06 | 0.253589 | 0.319 | 0.202 | 0.045307 | 3 L1 _ | Sun1 |
| Purb1 | 2.14E-06 | 0.274604 | 0.75 | 0.667 | 0.043071 | 3 L1 _ | Purb |
| Eif4ebp22 | 2.08E-06 | 0.320421 | 0.596 | 0.477 | 0.041811 | 3 L1 _ | Eif4ebp2 |
| Sbno2 | 2.02E-06 | 0.288267 | 0.3 | 0.185 | 0.040739 | 3 L1 _ | Sbno2 |

| | | | | | | | |
|---|---|---|---|---|---|---|---|
| Snai21 | 1.82E-06 | 0.307758 | 0.327 | 0.204 | 0.036614 | 3 L1 _ | Snai2 |
| Pcgf5 | 1.79E-06 | 0.310148 | 0.238 | 0.132 | 0.036087 | 3 L1 _ | Pcgf5 |
| Znrf11 | 1.53E-06 | 0.275235 | 0.381 | 0.248 | 0.030714 | 3 L1 _ | Znrf1 |
| Fosb1 | 1.36E-06 | 0.505995 | 0.523 | 0.397 | 0.027386 | 3 L1 _ | Fosb |
| Prrc2c | 1.32E-06 | 0.309153 | 0.85 | 0.799 | 0.026539 | 3 L1 _ | Prrc2c |
| Zkscan3 | 1.27E-06 | 0.365322 | 0.492 | 0.365 | 0.025493 | 3 L1 _ | Zkscan3 |
| Slc44a1 | 1.22E-06 | 0.292172 | 0.415 | 0.282 | 0.024659 | 3 L1 _ | Slc44a1 |
| Ccser2 | 1.21E-06 | 0.296579 | 0.665 | 0.555 | 0.02428 | 3 L1 _ | Ccser2 |
| Idnk1 | 1.16E-06 | 0.281856 | 0.527 | 0.395 | 0.023428 | 3 L1 _ | Idnk |
| Spag9 | 1.14E-06 | 0.277349 | 0.762 | 0.666 | 0.023008 | 3 L1 _ | Spag9 |
| Fas1 | 1.07E-06 | 0.271764 | 0.273 | 0.159 | 0.021527 | 3 L1 _ | Fas |
| Phf14 | 1.03E-06 | 0.353619 | 0.469 | 0.333 | 0.020671 | 3 L1 _ | Phf14 |
| Polg | 1.02E-06 | 0.287214 | 0.296 | 0.181 | 0.020631 | 3 L1 _ | Polg |
| Spaca6 | 9.74E-07 | 0.269785 | 0.373 | 0.244 | 0.019614 | 3 L1 _ | Spaca6 |
| Lrrfip11 | 9.21E-07 | 0.301326 | 0.612 | 0.464 | 0.018549 | 3 L1 _ | Lrrfip1 |
| Rnf215 | 8.95E-07 | 0.287691 | 0.4 | 0.27 | 0.018029 | 3 L1 _ | Rnf215 |
| Ppp1r9b | 8.80E-07 | 0.275779 | 0.423 | 0.294 | 0.017729 | 3 L1 _ | Ppp1r9b |
| Sft2d1 | 8.63E-07 | 0.367886 | 0.346 | 0.232 | 0.017381 | 3 L1 _ | Sft2d1 |
| Ash1l | 8.46E-07 | 0.319955 | 0.723 | 0.624 | 0.017047 | 3 L1 _ | Ash1l |
| Epn1 | 8.30E-07 | 0.288239 | 0.481 | 0.343 | 0.016714 | 3 L1 _ | Epn1 |
| Stx12 | 6.86E-07 | 0.346546 | 0.477 | 0.349 | 0.013808 | 3 L1 _ | Stx12 |
| Zfp644 | 6.85E-07 | 0.321526 | 0.665 | 0.546 | 0.01379 | 3 L1 _ | Zfp644 |
| Sash11 | 6.80E-07 | 0.261143 | 0.708 | 0.563 | 0.013689 | 3 L1 _ | Sash1 |
| Fcho2 | 6.39E-07 | 0.403956 | 0.612 | 0.5 | 0.012875 | 3 L1 _ | Fcho2 |
| Herc6 | 6.14E-07 | 0.377721 | 0.327 | 0.201 | 0.01236 | 3 L1 _ | Herc6 |
| Eef21 | 6.13E-07 | 0.250586 | 0.931 | 0.895 | 0.012353 | 3 L1 _ | Eef2 |
| Mtm1 | 5.29E-07 | 0.272734 | 0.215 | 0.11 | 0.010657 | 3 L1 _ | Mtm1 |
| Elovl1 | 5.26E-07 | 0.260783 | 0.5 | 0.361 | 0.010586 | 3 L1 _ | Elovl1 |
| Dpp8 | 5.21E-07 | 0.362193 | 0.496 | 0.36 | 0.010497 | 3 L1 _ | Dpp8 |
| Pcnx1 | 4.97E-07 | 0.302685 | 0.377 | 0.237 | 0.010012 | 3 L1 _ | Pcnx |
| Tra2a | 4.97E-07 | 0.276933 | 0.719 | 0.63 | 0.010003 | 3 L1 _ | Tra2a |
| Hspa1a | 4.76E-07 | 0.428954 | 0.342 | 0.207 | 0.009585 | 3 L1 _ | Hspa1a |
| Ocel1 | 4.48E-07 | 0.254474 | 0.227 | 0.12 | 0.00903 | 3 L1 _ | Ocel1 |
| Tspo | 4.28E-07 | 0.333982 | 0.915 | 0.857 | 0.008611 | 3 L1 _ | Tspo |
| Myh92 | 4.05E-07 | 0.315058 | 0.881 | 0.822 | 0.008154 | 3 L1 _ | Myh9 |
| Tsc22d32 | 3.99E-07 | 0.261352 | 0.627 | 0.461 | 0.008026 | 3 L1 _ | Tsc22d3 |
| Tmem234 | 3.87E-07 | 0.264814 | 0.8 | 0.707 | 0.007791 | 3 L1 _ | Tmem234 |
| Pak21 | 3.78E-07 | 0.336728 | 0.746 | 0.65 | 0.007623 | 3 L1 _ | Pak2 |
| Btbd9 | 3.75E-07 | 0.263579 | 0.342 | 0.211 | 0.007555 | 3 L1 _ | Btbd9 |
| Ifnar2 | 3.48E-07 | 0.378133 | 0.762 | 0.669 | 0.007001 | 3 L1 _ | Ifnar2 |
| Osmr | 3.46E-07 | 0.296207 | 0.558 | 0.42 | 0.006973 | 3 L1 _ | Osmr |

| | | | | | | | |
|---|---|---|---|---|---|---|---|
| Dip2b1 | 3.44E-07 | 0.290762 | 0.385 | 0.246 | 0.006926 | 3 L1 _ | Dip2b |
| Csnk2a1 | 3.35E-07 | 0.276145 | 0.785 | 0.699 | 0.006752 | 3 L1 _ | Csnk2a1 |
| Cactin1 | 3.33E-07 | 0.296707 | 0.227 | 0.119 | 0.006708 | 3 L1 _ | Cactin |
| Gpd21 | 3.30E-07 | 0.305598 | 0.219 | 0.115 | 0.006648 | 3 L1 _ | Gpd2 |
| Tmbim6 | 3.30E-07 | 0.329547 | 0.854 | 0.784 | 0.006636 | 3 L1 _ | Tmbim6 |
| Ifit2 | 3.26E-07 | 0.416211 | 0.312 | 0.188 | 0.006572 | 3 L1 _ | Ifit2 |
| Nrbp11 | 3.19E-07 | 0.294817 | 0.615 | 0.483 | 0.006422 | 3 L1 _ | Nrbp1 |
| Serpinb9 | 2.89E-07 | 0.295801 | 0.462 | 0.326 | 0.005819 | 3 L1 _ | Serpinb9 |
| Nbea | 2.83E-07 | 0.284098 | 0.396 | 0.254 | 0.005697 | 3 L1 _ | Nbea |
| 2700081O15Rik1 | 2.72E-07 | 0.278111 | 0.415 | 0.268 | 0.005483 | 3 L1 _ | 2700081O15Rik |
| Cttnbp2nl1 | 2.66E-07 | 0.287752 | 0.258 | 0.144 | 0.005359 | 3 L1 _ | Cttnbp2nl |
| Hpcal12 | 2.56E-07 | 0.315239 | 0.488 | 0.352 | 0.005157 | 3 L1 _ | Hpcal1 |
| Tapbpl1 | 2.51E-07 | 0.272274 | 0.315 | 0.188 | 0.005062 | 3 L1 _ | Tapbpl |
| Dram21 | 2.30E-07 | 0.278482 | 0.342 | 0.21 | 0.004633 | 3 L1 _ | Dram2 |
| Ralgds | 2.23E-07 | 0.318882 | 0.3 | 0.177 | 0.004483 | 3 L1 _ | Ralgds |
| Skap21 | 2.07E-07 | 0.250095 | 0.265 | 0.143 | 0.004162 | 3 L1 _ | Skap2 |
| Fam129b1 | 1.95E-07 | 0.311903 | 0.346 | 0.211 | 0.003931 | 3 L1 _ | Fam129b |
| Raly | 1.75E-07 | 0.32452 | 0.665 | 0.57 | 0.003531 | 3 L1 _ | Raly |
| Gatad2a1 | 1.49E-07 | 0.251715 | 0.465 | 0.315 | 0.002996 | 3 L1 _ | Gatad2a |
| Mcmbp | 1.44E-07 | 0.305829 | 0.385 | 0.249 | 0.0029 | 3 L1 _ | Mcmbp |
| Tap11 | 1.37E-07 | 0.28557 | 0.458 | 0.307 | 0.002756 | 3 L1 _ | Tap1 |
| R3hdm1 | 1.34E-07 | 0.287381 | 0.569 | 0.424 | 0.002704 | 3 L1 _ | R3hdm1 |
| Smarca41 | 1.34E-07 | 0.321312 | 0.396 | 0.258 | 0.002694 | 3 L1 _ | Smarca4 |
| Stim21 | 1.27E-07 | 0.324597 | 0.385 | 0.244 | 0.002566 | 3 L1 _ | Stim2 |
| Mgat11 | 1.18E-07 | 0.33823 | 0.45 | 0.319 | 0.002382 | 3 L1 _ | Mgat1 |
| Sdcbp1 | 1.14E-07 | 0.35774 | 0.712 | 0.608 | 0.002289 | 3 L1 _ | Sdcbp |
| Btbd7 | 1.11E-07 | 0.358735 | 0.692 | 0.561 | 0.002227 | 3 L1 _ | Btbd7 |
| Rarg | 1.09E-07 | 0.300886 | 0.431 | 0.277 | 0.002205 | 3 L1 _ | Rarg |
| Frmd4b | 1.06E-07 | 0.289809 | 0.262 | 0.141 | 0.002136 | 3 L1 _ | Frmd4b |
| Fam117b1 | 1.00E-07 | 0.28561 | 0.362 | 0.219 | 0.002023 | 3 L1 _ | Fam117b |
| Map4k31 | 9.46E-08 | 0.35819 | 0.358 | 0.216 | 0.001906 | 3 L1 _ | Map4k3 |
| Fam53b1 | 9.46E-08 | 0.297162 | 0.438 | 0.282 | 0.001905 | 3 L1 _ | Fam53b |
| Tet3 | 9.07E-08 | 0.378621 | 0.396 | 0.261 | 0.001827 | 3 L1 _ | Tet3 |
| Afap11 | 8.95E-08 | 0.317983 | 0.531 | 0.38 | 0.001802 | 3 L1 _ | Afap1 |
| Pomp | 8.84E-08 | 0.373376 | 0.827 | 0.756 | 0.00178 | 3 L1 _ | Pomp |
| Tax1bp31 | 8.40E-08 | 0.290451 | 0.542 | 0.395 | 0.001691 | 3 L1 _ | Tax1bp3 |
| Ube2v1 | 8.35E-08 | 0.289477 | 0.546 | 0.403 | 0.001682 | 3 L1 _ | Ube2v1 |
| Cdk19 | 8.15E-08 | 0.274862 | 0.304 | 0.173 | 0.001641 | 3 L1 _ | Cdk19 |
| Gm26802 | 7.11E-08 | 0.347648 | 0.177 | 0.079 | 0.001431 | 3 L1 _ | Gm26802 |
| Mcl12 | 6.74E-08 | 0.345602 | 0.719 | 0.619 | 0.001357 | 3 L1 _ | Mcl1 |
| Rock21 | 6.67E-08 | 0.287042 | 0.773 | 0.658 | 0.001343 | 3 L1 _ | Rock2 |

| | | | | | | | |
|---|---|---|---|---|---|---|---|
| Gcn1 | 6.63E-08 | 0.26268 | 0.381 | 0.23 | 0.001335 | 3 L1 _ | Gcn1 |
| Rbm391 | 6.40E-08 | 0.250339 | 0.931 | 0.917 | 0.001289 | 3 L1 _ | Rbm39 |
| Bloc1s6 | 6.33E-08 | 0.275846 | 0.281 | 0.154 | 0.001275 | 3 L1 _ | Bloc1s6 |
| Socs5 | 5.89E-08 | 0.277931 | 0.454 | 0.296 | 0.001186 | 3 L1 _ | Socs5 |
| Fmo22 | 5.88E-08 | 0.30992 | 0.623 | 0.447 | 0.001185 | 3 L1 _ | Fmo2 |
| Nr1h2 | 5.59E-08 | 0.284914 | 0.454 | 0.304 | 0.001127 | 3 L1 _ | Nr1h2 |
| Nsd1 | 5.26E-08 | 0.34423 | 0.723 | 0.602 | 0.001059 | 3 L1 _ | Nsd1 |
| Osbpl9 | 5.16E-08 | 0.353632 | 0.542 | 0.395 | 0.00104 | 3 L1 _ | Osbpl9 |
| Mid2 | 5.05E-08 | 0.263113 | 0.231 | 0.113 | 0.001016 | 3 L1 _ | Mid2 |
| Atp6v1e1 | 5.04E-08 | 0.350914 | 0.685 | 0.549 | 0.001015 | 3 L1 _ | Atp6v1e1 |
| Map3k12 | 4.66E-08 | 0.339883 | 0.269 | 0.147 | 0.000938 | 3 L1 _ | Map3k1 |
| Trim2 | 4.48E-08 | 0.446788 | 0.377 | 0.231 | 0.000903 | 3 L1 _ | Trim2 |
| Rnf19a1 | 4.44E-08 | 0.275839 | 0.338 | 0.196 | 0.000893 | 3 L1 _ | Rnf19a |
| Rap1a2 | 4.40E-08 | 0.3315 | 0.765 | 0.639 | 0.000886 | 3 L1 _ | Rap1a |
| Pea15a1 | 4.31E-08 | 0.290876 | 0.696 | 0.553 | 0.000869 | 3 L1 _ | Pea15a |
| Pik3c2a1 | 4.00E-08 | 0.336531 | 0.431 | 0.28 | 0.000805 | 3 L1 _ | Pik3c2a |
| Baz2b1 | 3.71E-08 | 0.429817 | 0.519 | 0.386 | 0.000746 | 3 L1 _ | Baz2b |
| Nkapd1 | 3.34E-08 | 0.275265 | 0.304 | 0.167 | 0.000672 | 3 L1 _ | Nkapd1 |
| Tasor | 3.23E-08 | 0.382526 | 0.5 | 0.336 | 0.00065 | 3 L1 _ | Tasor |
| Rilpl1 | 3.17E-08 | 0.297696 | 0.315 | 0.18 | 0.000638 | 3 L1 _ | Rilpl1 |
| Larp1b1 | 2.91E-08 | 0.250754 | 0.246 | 0.123 | 0.000585 | 3 L1 _ | Larp1b |
| Etnk11 | 2.86E-08 | 0.314555 | 0.592 | 0.423 | 0.000575 | 3 L1 _ | Etnk1 |
| Hspg23 | 2.68E-08 | 0.270386 | 0.846 | 0.681 | 0.000539 | 3 L1 _ | Hspg2 |
| Klc11 | 2.64E-08 | 0.316894 | 0.512 | 0.35 | 0.000532 | 3 L1 _ | Klc1 |
| Ccnl2 | 2.59E-08 | 0.368381 | 0.565 | 0.406 | 0.000522 | 3 L1 _ | Ccnl2 |
| Dstyk | 2.34E-08 | 0.37681 | 0.438 | 0.291 | 0.000472 | 3 L1 _ | Dstyk |
| Nod11 | 2.34E-08 | 0.274292 | 0.265 | 0.139 | 0.000472 | 3 L1 _ | Nod1 |
| 3222401L13Rik | 2.29E-08 | 0.269633 | 0.173 | 0.073 | 0.000462 | 3 L1 _ | 3222401L13Rik |
| Eif4g22 | 2.10E-08 | 0.30568 | 0.838 | 0.773 | 0.000423 | 3 L1 _ | Eif4g2 |
| Cdipt1 | 2.10E-08 | 0.404955 | 0.431 | 0.289 | 0.000423 | 3 L1 _ | Cdipt |
| Kctd101 | 2.01E-08 | 0.291603 | 0.469 | 0.307 | 0.000404 | 3 L1 _ | Kctd10 |
| Ciao2b | 1.84E-08 | 0.281239 | 0.454 | 0.293 | 0.00037 | 3 L1 _ | Ciao2b |
| Tmem30a1 | 1.73E-08 | 0.36358 | 0.715 | 0.598 | 0.000348 | 3 L1 _ | Tmem30a |
| Map3k3 | 1.60E-08 | 0.274072 | 0.312 | 0.174 | 0.000322 | 3 L1 _ | Map3k3 |
| Cebpg1 | 1.39E-08 | 0.380734 | 0.527 | 0.391 | 0.000281 | 3 L1 _ | Cebpg |
| Tsc22d12 | 1.32E-08 | 0.303218 | 0.946 | 0.821 | 0.000265 | 3 L1 _ | Tsc22d1 |
| Pdzd2 | 1.31E-08 | 0.301435 | 0.338 | 0.187 | 0.000264 | 3 L1 _ | Pdzd2 |
| Micall1 | 1.30E-08 | 0.261874 | 0.231 | 0.113 | 0.000261 | 3 L1 _ | Micall1 |
| St6galnac2 | 1.17E-08 | 0.272102 | 0.281 | 0.149 | 0.000236 | 3 L1 _ | St6galnac2 |
| Tmtc11 | 1.11E-08 | 0.295687 | 0.192 | 0.082 | 0.000224 | 3 L1 _ | Tmtc1 |
| Xdh2 | 9.58E-09 | 0.392158 | 0.573 | 0.416 | 0.000193 | 3 L1 _ | Xdh |

| | | | | | | | |
|---|---|---|---|---|---|---|---|
| Diaph21 | 9.49E-09 | 0.330272 | 0.542 | 0.387 | 0.000191 | 3 L1 _ | Diaph2 |
| Plekha1 | 8.75E-09 | 0.370843 | 0.404 | 0.249 | 0.000176 | 3 L1 _ | Plekha1 |
| Ints91 | 7.36E-09 | 0.277938 | 0.246 | 0.12 | 0.000148 | 3 L1 _ | Ints9 |
| Baiap21 | 6.80E-09 | 0.266404 | 0.269 | 0.138 | 0.000137 | 3 L1 _ | Baiap2 |
| Nmi1 | 6.74E-09 | 0.364881 | 0.454 | 0.301 | 0.000136 | 3 L1 _ | Nmi |
| Rab13 | 6.65E-09 | 0.410193 | 0.523 | 0.371 | 0.000134 | 3 L1 _ | Rab13 |
| St3gal41 | 6.39E-09 | 0.338152 | 0.362 | 0.208 | 0.000129 | 3 L1 _ | St3gal4 |
| Ifi203 | 6.28E-09 | 0.36345 | 0.838 | 0.711 | 0.000126 | 3 L1 _ | Ifi203 |
| Tpm42 | 6.11E-09 | 0.356819 | 0.854 | 0.744 | 0.000123 | 3 L1 _ | Tpm4 |
| Tgfb11 | 6.03E-09 | 0.31322 | 0.642 | 0.458 | 0.000121 | 3 L1 _ | Tgfb1 |
| Fbxo221 | 5.68E-09 | 0.330378 | 0.427 | 0.266 | 0.000114 | 3 L1 _ | Fbxo22 |
| Magi3 | 5.65E-09 | 0.281948 | 0.254 | 0.125 | 0.000114 | 3 L1 _ | Magi3 |
| Fos1 | 5.35E-09 | 0.759581 | 0.627 | 0.487 | 0.000108 | 3 L1 _ | Fos |
| Meis2 | 4.81E-09 | 0.451504 | 0.527 | 0.369 | 9.68E-05 | 3 L1 _ | Meis2 |
| Dgkz | 4.35E-09 | 0.292536 | 0.358 | 0.208 | 8.76E-05 | 3 L1 _ | Dgkz |
| Slc39a10 | 4.34E-09 | 0.372699 | 0.377 | 0.223 | 8.75E-05 | 3 L1 _ | Slc39a10 |
| Gnb2 | 4.14E-09 | 0.333649 | 0.877 | 0.854 | 8.33E-05 | 3 L1 _ | Gnb2 |
| Fam171a11 | 4.11E-09 | 0.293454 | 0.285 | 0.147 | 8.28E-05 | 3 L1 _ | Fam171a1 |
| Bag3 | 4.00E-09 | 0.337938 | 0.408 | 0.244 | 8.07E-05 | 3 L1 _ | Bag3 |
| Exoc61 | 3.92E-09 | 0.275934 | 0.2 | 0.087 | 7.90E-05 | 3 L1 _ | Exoc6 |
| Ywhae1 | 3.90E-09 | 0.328919 | 0.869 | 0.775 | 7.86E-05 | 3 L1 _ | Ywhae |
| Arhgap171 | 3.83E-09 | 0.362648 | 0.485 | 0.328 | 7.71E-05 | 3 L1 _ | Arhgap17 |
| Hprt1 | 3.27E-09 | 0.39608 | 0.473 | 0.316 | 6.59E-05 | 3 L1 _ | Hprt |
| Ccdc88c1 | 3.26E-09 | 0.327409 | 0.235 | 0.109 | 6.57E-05 | 3 L1 _ | Ccdc88c |
| Pdlim12 | 3.17E-09 | 0.329415 | 0.412 | 0.243 | 6.39E-05 | 3 L1 _ | Pdlim1 |
| Rbms11 | 3.13E-09 | 0.331173 | 0.858 | 0.77 | 6.30E-05 | 3 L1 _ | Rbms1 |
| Phf20l1 | 3.04E-09 | 0.414167 | 0.65 | 0.509 | 6.13E-05 | 3 L1 _ | Phf20l1 |
| Atp5j | 2.95E-09 | 0.326625 | 0.927 | 0.882 | 5.95E-05 | 3 L1 _ | Atp5j |
| Ehd22 | 2.80E-09 | 0.368121 | 0.619 | 0.471 | 5.63E-05 | 3 L1 _ | Ehd2 |
| B230219D22Rik2 | 2.72E-09 | 0.358935 | 0.785 | 0.7 | 5.47E-05 | 3 L1 _ | B230219D22Rik |
| Slk1 | 2.71E-09 | 0.412458 | 0.608 | 0.464 | 5.46E-05 | 3 L1 _ | Slk |
| Gbp2 | 2.62E-09 | 0.353844 | 0.508 | 0.344 | 5.28E-05 | 3 L1 _ | Gbp2 |
| Card191 | 2.11E-09 | 0.406247 | 0.438 | 0.279 | 4.25E-05 | 3 L1 _ | Card19 |
| Palm1 | 2.07E-09 | 0.385347 | 0.388 | 0.234 | 4.16E-05 | 3 L1 _ | Palm |
| Rab5c1 | 1.82E-09 | 0.39062 | 0.696 | 0.572 | 3.66E-05 | 3 L1 _ | Rab5c |
| Klhl24 | 1.79E-09 | 0.431948 | 0.642 | 0.499 | 3.61E-05 | 3 L1 _ | Klhl24 |
| Zfp36l1 | 1.68E-09 | 0.38101 | 0.881 | 0.823 | 3.39E-05 | 3 L1 _ | Zfp36l1 |
| Dnm2 | 1.66E-09 | 0.43049 | 0.531 | 0.365 | 3.34E-05 | 3 L1 _ | Dnm2 |
| Dazap2 | 1.62E-09 | 0.430323 | 0.6 | 0.452 | 3.27E-05 | 3 L1 _ | Dazap2 |
| Trim56 | 1.61E-09 | 0.449493 | 0.481 | 0.321 | 3.23E-05 | 3 L1 _ | Trim56 |
| Fut81 | 1.56E-09 | 0.472609 | 0.465 | 0.314 | 3.14E-05 | 3 L1 _ | Fut8 |

| | | | | | | | |
|---|---|---|---|---|---|---|---|
| Igtp | 1.49E-09 | 0.36521 | 0.354 | 0.201 | 3.00E-05 | 3 L1 _ | Igtp |
| Arhgef2 | 1.42E-09 | 0.384998 | 0.485 | 0.328 | 2.85E-05 | 3 L1 _ | Arhgef2 |
| Tmpo1 | 1.31E-09 | 0.291729 | 0.527 | 0.338 | 2.63E-05 | 3 L1 _ | Tmpo |
| Arpc51 | 1.29E-09 | 0.32129 | 0.808 | 0.67 | 2.60E-05 | 3 L1 _ | Arpc5 |
| Cds22 | 1.24E-09 | 0.35057 | 0.458 | 0.287 | 2.50E-05 | 3 L1 _ | Cds2 |
| S100a11 | 1.11E-09 | 0.34336 | 0.608 | 0.425 | 2.23E-05 | 3 L1 _ | S100a1 |
| Pnkd | 1.07E-09 | 0.385583 | 0.612 | 0.452 | 2.16E-05 | 3 L1 _ | Pnkd |
| Casp81 | 1.05E-09 | 0.370274 | 0.358 | 0.206 | 2.11E-05 | 3 L1 _ | Casp8 |
| Kmt2e | 1.00E-09 | 0.397605 | 0.823 | 0.76 | 2.02E-05 | 3 L1 _ | Kmt2e |
| Map2k3 | 9.99E-10 | 0.295642 | 0.377 | 0.212 | 2.01E-05 | 3 L1 _ | Map2k3 |
| Trib22 | 9.63E-10 | 0.42052 | 0.512 | 0.336 | 1.94E-05 | 3 L1 _ | Trib2 |
| Stat1 | 9.28E-10 | 0.404453 | 0.558 | 0.394 | 1.87E-05 | 3 L1 _ | Stat1 |
| Stard9 | 9.05E-10 | 0.381271 | 0.408 | 0.248 | 1.82E-05 | 3 L1 _ | Stard9 |
| Atp6v0c2 | 8.44E-10 | 0.289779 | 0.846 | 0.66 | 1.70E-05 | 3 L1 _ | Atp6v0c |
| Ndufc21 | 8.30E-10 | 0.358185 | 0.812 | 0.699 | 1.67E-05 | 3 L1 _ | Ndufc2 |
| Cenpb | 7.65E-10 | 0.432521 | 0.669 | 0.527 | 1.54E-05 | 3 L1 _ | Cenpb |
| Stx21 | 7.25E-10 | 0.421847 | 0.412 | 0.252 | 1.46E-05 | 3 L1 _ | Stx2 |
| Mfap3 | 7.11E-10 | 0.335099 | 0.338 | 0.182 | 1.43E-05 | 3 L1 _ | Mfap3 |
| Midn1 | 6.63E-10 | 0.357397 | 0.512 | 0.343 | 1.34E-05 | 3 L1 _ | Midn |
| Apc1 | 6.43E-10 | 0.375559 | 0.5 | 0.337 | 1.30E-05 | 3 L1 _ | Apc |
| Mrpl17 | 6.38E-10 | 0.427574 | 0.592 | 0.439 | 1.28E-05 | 3 L1 _ | Mrpl17 |
| Clstn12 | 6.21E-10 | 0.331197 | 0.7 | 0.553 | 1.25E-05 | 3 L1 _ | Clstn1 |
| Sh3glb12 | 5.88E-10 | 0.35926 | 0.892 | 0.804 | 1.18E-05 | 3 L1 _ | Sh3glb1 |
| Ahnak3 | 5.80E-10 | 0.368287 | 0.973 | 0.885 | 1.17E-05 | 3 L1 _ | Ahnak |
| Zswim6 | 5.63E-10 | 0.285732 | 0.223 | 0.097 | 1.13E-05 | 3 L1 _ | Zswim6 |
| Tmem63b1 | 4.69E-10 | 0.270163 | 0.258 | 0.123 | 9.45E-06 | 3 L1 _ | Tmem63b |
| Gna111 | 4.53E-10 | 0.408731 | 0.723 | 0.603 | 9.13E-06 | 3 L1 _ | Gna11 |
| Taf7 | 4.49E-10 | 0.27588 | 0.419 | 0.239 | 9.04E-06 | 3 L1 _ | Taf7 |
| Prrg21 | 4.46E-10 | 0.301985 | 0.223 | 0.097 | 8.98E-06 | 3 L1 _ | Prrg2 |
| Txnip1 | 4.38E-10 | 0.33352 | 0.927 | 0.846 | 8.83E-06 | 3 L1 _ | Txnip |
| Tecpr11 | 4.01E-10 | 0.318954 | 0.435 | 0.264 | 8.07E-06 | 3 L1 _ | Tecpr1 |
| Cdc42ep3 | 3.55E-10 | 0.397458 | 0.388 | 0.231 | 7.15E-06 | 3 L1 _ | Cdc42ep3 |
| Snn | 2.56E-10 | 0.259806 | 0.15 | 0.052 | 5.16E-06 | 3 L1 _ | Snn |
| Psme21 | 2.48E-10 | 0.365548 | 0.738 | 0.611 | 5.00E-06 | 3 L1 _ | Psme2 |
| Unc119b1 | 2.36E-10 | 0.349779 | 0.385 | 0.222 | 4.75E-06 | 3 L1 _ | Unc119b |
| Phc2 | 2.02E-10 | 0.371597 | 0.454 | 0.29 | 4.06E-06 | 3 L1 _ | Phc2 |
| Ifi351 | 1.98E-10 | 0.42801 | 0.554 | 0.383 | 3.98E-06 | 3 L1 _ | Ifi35 |
| Pelo | 1.86E-10 | 0.306038 | 0.358 | 0.191 | 3.75E-06 | 3 L1 _ | Pelo |
| Ddit41 | 1.78E-10 | 0.343421 | 0.292 | 0.145 | 3.59E-06 | 3 L1 _ | Ddit4 |
| Eml1 | 1.64E-10 | 0.398301 | 0.312 | 0.161 | 3.30E-06 | 3 L1 _ | Eml1 |
| Emp21 | 1.29E-10 | 0.49564 | 0.692 | 0.546 | 2.59E-06 | 3 L1 _ | Emp2 |

| | | | | | | | |
|---|---|---|---|---|---|---|---|
| Pfkfb31 | 1.25E-10 | 0.258224 | 0.135 | 0.042 | 2.51E-06 | 3 L1 _ | Pfkfb3 |
| Mtss1 | 1.19E-10 | 0.370331 | 0.312 | 0.159 | 2.39E-06 | 3 L1 _ | Mtss1 |
| Map4k4 | 1.13E-10 | 0.484359 | 0.742 | 0.627 | 2.28E-06 | 3 L1 _ | Map4k4 |
| Stmn11 | 1.08E-10 | 0.258878 | 0.227 | 0.096 | 2.18E-06 | 3 L1 _ | Stmn1 |
| Asap12 | 1.07E-10 | 0.3741 | 0.769 | 0.6 | 2.16E-06 | 3 L1 _ | Asap1 |
| Ndst1 | 1.02E-10 | 0.459828 | 0.446 | 0.285 | 2.06E-06 | 3 L1 _ | Ndst1 |
| Actg12 | 9.14E-11 | 0.290428 | 0.969 | 0.942 | 1.84E-06 | 3 L1 _ | Actg1 |
| Slc50a1 | 8.10E-11 | 0.465903 | 0.738 | 0.632 | 1.63E-06 | 3 L1 _ | Slc50a1 |
| Rtp4 | 7.59E-11 | 0.362951 | 0.519 | 0.318 | 1.53E-06 | 3 L1 _ | Rtp4 |
| Tcn21 | 7.54E-11 | 0.544235 | 0.654 | 0.505 | 1.52E-06 | 3 L1 _ | Tcn2 |
| Ptk22 | 7.36E-11 | 0.468677 | 0.512 | 0.332 | 1.48E-06 | 3 L1 _ | Ptk2 |
| Rps6ka31 | 7.21E-11 | 0.460739 | 0.462 | 0.286 | 1.45E-06 | 3 L1 _ | Rps6ka3 |
| Rell1 | 6.08E-11 | 0.365668 | 0.358 | 0.189 | 1.22E-06 | 3 L1 _ | Rell1 |
| Prickle3 | 6.00E-11 | 0.322099 | 0.242 | 0.107 | 1.21E-06 | 3 L1 _ | Prickle3 |
| Triobp | 5.90E-11 | 0.370737 | 0.431 | 0.257 | 1.19E-06 | 3 L1 _ | Triobp |
| Atp2b42 | 5.74E-11 | 0.32246 | 0.496 | 0.291 | 1.16E-06 | 3 L1 _ | Atp2b4 |
| Sod12 | 5.55E-11 | 0.421546 | 0.688 | 0.515 | 1.12E-06 | 3 L1 _ | Sod1 |
| Pdlim51 | 5.16E-11 | 0.439845 | 0.485 | 0.304 | 1.04E-06 | 3 L1 _ | Pdlim5 |
| Maco1 | 4.69E-11 | 0.361055 | 0.331 | 0.172 | 9.44E-07 | 3 L1 _ | Maco1 |
| Tubb4b2 | 4.64E-11 | 0.491988 | 0.804 | 0.69 | 9.35E-07 | 3 L1 _ | Tubb4b |
| Ptpn121 | 4.56E-11 | 0.472659 | 0.492 | 0.315 | 9.19E-07 | 3 L1 _ | Ptpn12 |
| Rac11 | 4.46E-11 | 0.383496 | 0.835 | 0.751 | 8.97E-07 | 3 L1 _ | Rac1 |
| Abhd17a | 4.19E-11 | 0.46177 | 0.642 | 0.496 | 8.43E-07 | 3 L1 _ | Abhd17a |
| Itsn21 | 4.00E-11 | 0.454199 | 0.481 | 0.296 | 8.05E-07 | 3 L1 _ | Itsn2 |
| Colgalt21 | 3.88E-11 | 0.307275 | 0.25 | 0.108 | 7.81E-07 | 3 L1 _ | Colgalt2 |
| Snx31 | 3.66E-11 | 0.430438 | 0.788 | 0.683 | 7.37E-07 | 3 L1 _ | Snx3 |
| Trim30a1 | 3.55E-11 | 0.461081 | 0.465 | 0.285 | 7.15E-07 | 3 L1 _ | Trim30a |
| Map4 | 2.49E-11 | 0.545965 | 0.638 | 0.494 | 5.01E-07 | 3 L1 _ | Map4 |
| Ier21 | 2.48E-11 | 0.61162 | 0.608 | 0.441 | 4.99E-07 | 3 L1 _ | Ier2 |
| Fyn | 2.39E-11 | 0.407667 | 0.573 | 0.38 | 4.81E-07 | 3 L1 _ | Fyn |
| Rap1b1 | 2.20E-11 | 0.440944 | 0.785 | 0.674 | 4.43E-07 | 3 L1 _ | Rap1b |
| 4930581F22Rik1 | 2.05E-11 | 0.328027 | 0.323 | 0.16 | 4.12E-07 | 3 L1 _ | 4930581F22Rik |
| Kif5b1 | 2.00E-11 | 0.465938 | 0.785 | 0.668 | 4.02E-07 | 3 L1 _ | Kif5b |
| Mmd1 | 1.90E-11 | 0.292408 | 0.208 | 0.082 | 3.83E-07 | 3 L1 _ | Mmd |
| Hip1r | 1.88E-11 | 0.277872 | 0.212 | 0.083 | 3.78E-07 | 3 L1 _ | Hip1r |
| Nop53 | 1.69E-11 | 0.54326 | 0.642 | 0.48 | 3.40E-07 | 3 L1 _ | Nop53 |
| Atp8b1 | 1.62E-11 | 0.428936 | 0.369 | 0.206 | 3.25E-07 | 3 L1 _ | Atp8b1 |
| Ccrl2 | 1.54E-11 | 0.318043 | 0.292 | 0.135 | 3.11E-07 | 3 L1 _ | Ccrl2 |
| Tnfaip21 | 1.40E-11 | 0.442526 | 0.6 | 0.409 | 2.83E-07 | 3 L1 _ | Tnfaip2 |
| Rasa4 | 1.08E-11 | 0.25462 | 0.188 | 0.066 | 2.17E-07 | 3 L1 _ | Rasa4 |
| Cox7a2l1 | 1.04E-11 | 0.43629 | 0.85 | 0.738 | 2.09E-07 | 3 L1 _ | Cox7a2l |

| | | | | | | | |
|---|---|---|---|---|---|---|---|
| Cyb5r31 | 8.89E-12 | 0.456752 | 0.719 | 0.566 | 1.79E-07 | 3 L1 _ | Cyb5r3 |
| Kmt2a | 8.73E-12 | 0.378616 | 0.742 | 0.551 | 1.76E-07 | 3 L1 _ | Kmt2a |
| Myl12b1 | 8.27E-12 | 0.402873 | 0.869 | 0.787 | 1.67E-07 | 3 L1 _ | Myl12b |
| Aopep | 8.13E-12 | 0.342618 | 0.523 | 0.325 | 1.64E-07 | 3 L1 _ | Aopep |
| Cd472 | 8.05E-12 | 0.39911 | 0.919 | 0.856 | 1.62E-07 | 3 L1 _ | Cd47 |
| Zhx32 | 7.65E-12 | 0.474642 | 0.492 | 0.311 | 1.54E-07 | 3 L1 _ | Zhx3 |
| Pdgfd1 | 7.04E-12 | 0.322611 | 0.242 | 0.101 | 1.42E-07 | 3 L1 _ | Pdgfd |
| Gnb11 | 5.82E-12 | 0.419702 | 0.827 | 0.678 | 1.17E-07 | 3 L1 _ | Gnb1 |
| Mndal2 | 5.65E-12 | 0.443681 | 0.815 | 0.65 | 1.14E-07 | 3 L1 _ | Mndal |
| Gle1 | 5.62E-12 | 0.34945 | 0.4 | 0.217 | 1.13E-07 | 3 L1 _ | Gle1 |
| Bok2 | 5.46E-12 | 0.261977 | 0.292 | 0.131 | 1.10E-07 | 3 L1 _ | Bok |
| Ndufa81 | 4.94E-12 | 0.436656 | 0.735 | 0.589 | 9.95E-08 | 3 L1 _ | Ndufa8 |
| Elf12 | 4.91E-12 | 0.494578 | 0.619 | 0.447 | 9.89E-08 | 3 L1 _ | Elf1 |
| Cr1l | 4.61E-12 | 0.528651 | 0.688 | 0.521 | 9.28E-08 | 3 L1 _ | Cr1l |
| Tapbp1 | 4.37E-12 | 0.451368 | 0.765 | 0.624 | 8.80E-08 | 3 L1 _ | Tapbp |
| Zeb11 | 4.25E-12 | 0.427636 | 0.738 | 0.592 | 8.56E-08 | 3 L1 _ | Zeb1 |
| Cyb5611 | 4.22E-12 | 0.294631 | 0.227 | 0.091 | 8.50E-08 | 3 L1 _ | Cyb561 |
| Rnasek1 | 3.71E-12 | 0.425039 | 0.796 | 0.571 | 7.47E-08 | 3 L1 _ | Rnasek |
| Atp1a11 | 3.68E-12 | 0.453657 | 0.596 | 0.422 | 7.40E-08 | 3 L1 _ | Atp1a1 |
| Lats21 | 3.31E-12 | 0.423998 | 0.5 | 0.317 | 6.66E-08 | 3 L1 _ | Lats2 |
| Ninj1 | 3.22E-12 | 0.348858 | 0.488 | 0.292 | 6.48E-08 | 3 L1 _ | Ninj1 |
| Isg15 | 2.81E-12 | 0.508441 | 0.531 | 0.321 | 5.66E-08 | 3 L1 _ | Isg15 |
| Sgk31 | 2.81E-12 | 0.297622 | 0.273 | 0.119 | 5.65E-08 | 3 L1 _ | Sgk3 |
| Itm2b1 | 2.64E-12 | 0.345752 | 1 | 0.989 | 5.31E-08 | 3 L1 _ | Itm2b |
| Chd71 | 2.53E-12 | 0.316964 | 0.242 | 0.096 | 5.10E-08 | 3 L1 _ | Chd7 |
| Sema3c2 | 2.28E-12 | 0.414106 | 0.254 | 0.105 | 4.60E-08 | 3 L1 _ | Sema3c |
| Ostf13 | 2.14E-12 | 0.472957 | 0.746 | 0.594 | 4.32E-08 | 3 L1 _ | Ostf1 |
| Arl6ip51 | 2.08E-12 | 0.444268 | 0.669 | 0.487 | 4.19E-08 | 3 L1 _ | Arl6ip5 |
| Hes11 | 2.00E-12 | 0.454049 | 0.762 | 0.581 | 4.04E-08 | 3 L1 _ | Hes1 |
| Znfx11 | 1.98E-12 | 0.456008 | 0.423 | 0.241 | 3.98E-08 | 3 L1 _ | Znfx1 |
| Hras1 | 1.85E-12 | 0.394192 | 0.615 | 0.42 | 3.72E-08 | 3 L1 _ | Hras |
| Ipo111 | 1.85E-12 | 0.366348 | 0.262 | 0.113 | 3.72E-08 | 3 L1 _ | Ipo11 |
| Epha41 | 1.83E-12 | 0.379387 | 0.354 | 0.176 | 3.69E-08 | 3 L1 _ | Epha4 |
| Sntb21 | 1.68E-12 | 0.458523 | 0.708 | 0.532 | 3.39E-08 | 3 L1 _ | Sntb2 |
| Nuak1 | 1.67E-12 | 0.379214 | 0.285 | 0.127 | 3.35E-08 | 3 L1 _ | Nuak1 |
| Zdhhc182 | 1.54E-12 | 0.465817 | 0.373 | 0.191 | 3.09E-08 | 3 L1 _ | Zdhhc18 |
| Pkn11 | 1.50E-12 | 0.447713 | 0.473 | 0.274 | 3.02E-08 | 3 L1 _ | Pkn1 |
| Tmem1401 | 1.38E-12 | 0.47414 | 0.585 | 0.397 | 2.78E-08 | 3 L1 _ | Tmem140 |
| Ro601 | 1.35E-12 | 0.375661 | 0.469 | 0.27 | 2.73E-08 | 3 L1 _ | Ro60 |
| Mrpl53 | 1.35E-12 | 0.40374 | 0.45 | 0.256 | 2.71E-08 | 3 L1 _ | Mrpl53 |
| Cyp26b12 | 1.19E-12 | 0.507047 | 0.242 | 0.094 | 2.39E-08 | 3 L1 _ | Cyp26b1 |

| | | | | | | | |
|---|---|---|---|---|---|---|---|
| Cyp2d22 | 1.14E-12 | 0.440528 | 0.55 | 0.361 | 2.30E-08 | 3 L1 _ | Cyp2d22 |
| Adipor11 | 1.13E-12 | 0.429587 | 0.592 | 0.414 | 2.28E-08 | 3 L1 _ | Adipor1 |
| Wwc21 | 1.10E-12 | 0.404413 | 0.638 | 0.448 | 2.22E-08 | 3 L1 _ | Wwc2 |
| Gm40701 | 1.10E-12 | 0.400639 | 0.231 | 0.091 | 2.21E-08 | 3 L1 _ | Gm4070 |
| Trim472 | 1.06E-12 | 0.452063 | 0.581 | 0.385 | 2.13E-08 | 3 L1 _ | Trim47 |
| Peak11 | 8.05E-13 | 0.458633 | 0.754 | 0.616 | 1.62E-08 | 3 L1 _ | Peak1 |
| Dab21 | 6.76E-13 | 0.453117 | 0.604 | 0.404 | 1.36E-08 | 3 L1 _ | Dab2 |
| Ncoa71 | 6.71E-13 | 0.403869 | 0.558 | 0.348 | 1.35E-08 | 3 L1 _ | Ncoa7 |
| Tmem2042 | 6.53E-13 | 0.444269 | 0.758 | 0.558 | 1.32E-08 | 3 L1 _ | Tmem204 |
| Golm11 | 6.38E-13 | 0.455694 | 0.362 | 0.185 | 1.29E-08 | 3 L1 _ | Golm1 |
| Ccnyl11 | 5.89E-13 | 0.301943 | 0.2 | 0.07 | 1.19E-08 | 3 L1 _ | Ccnyl1 |
| Irf2bpl1 | 5.69E-13 | 0.491204 | 0.658 | 0.492 | 1.15E-08 | 3 L1 _ | Irf2bpl |
| Pwwp3a | 5.40E-13 | 0.398436 | 0.431 | 0.23 | 1.09E-08 | 3 L1 _ | Pwwp3a |
| Slfn21 | 5.09E-13 | 0.602995 | 0.504 | 0.302 | 1.02E-08 | 3 L1 _ | Slfn2 |
| Cdc42ep2 | 5.06E-13 | 0.493455 | 0.542 | 0.357 | 1.02E-08 | 3 L1 _ | Cdc42ep2 |
| Ppfibp1 | 4.43E-13 | 0.509674 | 0.592 | 0.402 | 8.93E-09 | 3 L1 _ | Ppfibp1 |
| H2-T22 | 4.25E-13 | 0.570246 | 0.692 | 0.501 | 8.56E-09 | 3 L1 _ | H2-T22 |
| Timeless1 | 3.71E-13 | 0.310305 | 0.246 | 0.098 | 7.47E-09 | 3 L1 _ | Timeless |
| A930005H10Rik | 3.69E-13 | 0.306787 | 0.2 | 0.069 | 7.42E-09 | 3 L1 _ | A930005H10Rik |
| Dchs1 | 3.47E-13 | 0.311382 | 0.315 | 0.145 | 6.98E-09 | 3 L1 _ | Dchs1 |
| Vamp31 | 3.37E-13 | 0.427858 | 0.492 | 0.302 | 6.78E-09 | 3 L1 _ | Vamp3 |
| Ier51 | 2.99E-13 | 0.509093 | 0.669 | 0.489 | 6.02E-09 | 3 L1 _ | Ier5 |
| Sp1001 | 2.98E-13 | 0.597596 | 0.696 | 0.532 | 6.01E-09 | 3 L1 _ | Sp100 |
| Tnfsf13 | 2.86E-13 | 0.292408 | 0.338 | 0.156 | 5.75E-09 | 3 L1 _ | Tnfsf13 |
| Cox5a1 | 2.77E-13 | 0.422096 | 0.869 | 0.756 | 5.57E-09 | 3 L1 _ | Cox5a |
| Prrg1 | 2.04E-13 | 0.402417 | 0.288 | 0.126 | 4.11E-09 | 3 L1 _ | Prrg1 |
| Dusp13 | 1.85E-13 | 0.593369 | 0.523 | 0.316 | 3.73E-09 | 3 L1 _ | Dusp1 |
| Enpp4 | 1.85E-13 | 0.415864 | 0.281 | 0.12 | 3.72E-09 | 3 L1 _ | Enpp4 |
| Stard41 | 1.57E-13 | 0.364561 | 0.231 | 0.088 | 3.16E-09 | 3 L1 _ | Stard4 |
| Endod11 | 1.55E-13 | 0.495383 | 0.554 | 0.359 | 3.12E-09 | 3 L1 _ | Endod1 |
| Zfyve9 | 1.40E-13 | 0.350721 | 0.404 | 0.204 | 2.82E-09 | 3 L1 _ | Zfyve9 |
| Nedd92 | 1.31E-13 | 0.581465 | 0.527 | 0.33 | 2.63E-09 | 3 L1 _ | Nedd9 |
| Tmod31 | 1.02E-13 | 0.507363 | 0.75 | 0.594 | 2.06E-09 | 3 L1 _ | Tmod3 |
| Oaz22 | 8.41E-14 | 0.568961 | 0.619 | 0.426 | 1.69E-09 | 3 L1 _ | Oaz2 |
| Gja11 | 7.14E-14 | 0.727819 | 0.519 | 0.333 | 1.44E-09 | 3 L1 _ | Gja1 |
| Flnb | 7.08E-14 | 0.429321 | 0.438 | 0.246 | 1.42E-09 | 3 L1 _ | Flnb |
| Sorbs31 | 7.06E-14 | 0.340482 | 0.323 | 0.147 | 1.42E-09 | 3 L1 _ | Sorbs3 |
| Rgs3 | 6.89E-14 | 0.563195 | 0.577 | 0.384 | 1.39E-09 | 3 L1 _ | Rgs3 |
| Zmiz11 | 6.71E-14 | 0.499134 | 0.673 | 0.493 | 1.35E-09 | 3 L1 _ | Zmiz1 |
| Cavin12 | 6.05E-14 | 0.364121 | 0.696 | 0.458 | 1.22E-09 | 3 L1 _ | Cavin1 |
| Cdc42ep11 | 6.00E-14 | 0.358035 | 0.458 | 0.247 | 1.21E-09 | 3 L1 _ | Cdc42ep1 |

| | | | | | | | |
|---|---|---|---|---|---|---|---|
| Phactr41 | 5.71E-14 | 0.449926 | 0.423 | 0.221 | 1.15E-09 | 3 L1 _ | Phactr4 |
| Samhd1 | 4.96E-14 | 0.522587 | 0.608 | 0.419 | 1.00E-09 | 3 L1 _ | Samhd1 |
| Lgals9 | 3.84E-14 | 0.491709 | 0.692 | 0.498 | 7.73E-10 | 3 L1 _ | Lgals9 |
| Rgl21 | 3.73E-14 | 0.33847 | 0.342 | 0.16 | 7.52E-10 | 3 L1 _ | Rgl2 |
| Ctnnd11 | 3.30E-14 | 0.464163 | 0.519 | 0.32 | 6.64E-10 | 3 L1 _ | Ctnnd1 |
| Agrn1 | 3.10E-14 | 0.369115 | 0.381 | 0.185 | 6.24E-10 | 3 L1 _ | Agrn |
| Peg13 | 3.09E-14 | 0.382562 | 0.335 | 0.149 | 6.22E-10 | 3 L1 _ | Peg13 |
| Psmb101 | 2.67E-14 | 0.479091 | 0.742 | 0.563 | 5.37E-10 | 3 L1 _ | Psmb10 |
| Sppl2a1 | 2.01E-14 | 0.514071 | 0.696 | 0.505 | 4.05E-10 | 3 L1 _ | Sppl2a |
| Atrx | 1.99E-14 | 0.55143 | 0.877 | 0.769 | 4.00E-10 | 3 L1 _ | Atrx |
| Ier31 | 1.98E-14 | 0.507687 | 0.327 | 0.145 | 3.98E-10 | 3 L1 _ | Ier3 |
| Dock11 | 1.86E-14 | 0.498037 | 0.562 | 0.362 | 3.75E-10 | 3 L1 _ | Dock1 |
| Rps141 | 1.62E-14 | 0.299021 | 0.985 | 0.972 | 3.26E-10 | 3 L1 _ | Rps14 |
| Ripor11 | 1.38E-14 | 0.444669 | 0.458 | 0.25 | 2.78E-10 | 3 L1 _ | Ripor1 |
| Cmip1 | 1.35E-14 | 0.462109 | 0.45 | 0.245 | 2.72E-10 | 3 L1 _ | Cmip |
| Exoc3l4 | 1.30E-14 | 0.483961 | 0.431 | 0.233 | 2.61E-10 | 3 L1 _ | Exoc3l4 |
| Dlc13 | 1.16E-14 | 0.440322 | 0.823 | 0.691 | 2.34E-10 | 3 L1 _ | Dlc1 |
| Dgkh1 | 1.05E-14 | 0.490906 | 0.404 | 0.203 | 2.11E-10 | 3 L1 _ | Dgkh |
| Cep85l1 | 8.60E-15 | 0.324375 | 0.292 | 0.118 | 1.73E-10 | 3 L1 _ | Cep85l |
| Dock61 | 8.24E-15 | 0.415573 | 0.373 | 0.176 | 1.66E-10 | 3 L1 _ | Dock6 |
| Slco2a11 | 7.86E-15 | 0.541528 | 0.308 | 0.134 | 1.58E-10 | 3 L1 _ | Slco2a1 |
| Gch11 | 7.28E-15 | 0.311484 | 0.196 | 0.063 | 1.47E-10 | 3 L1 _ | Gch1 |
| Sh3bgrl4 | 6.93E-15 | 0.520238 | 0.819 | 0.671 | 1.40E-10 | 3 L1 _ | Sh3bgrl |
| Slc3a22 | 6.62E-15 | 0.543179 | 0.723 | 0.57 | 1.33E-10 | 3 L1 _ | Slc3a2 |
| Iqgap11 | 6.12E-15 | 0.441628 | 0.854 | 0.746 | 1.23E-10 | 3 L1 _ | Iqgap1 |
| 0610010K14Rik | 5.96E-15 | 0.47777 | 0.531 | 0.314 | 1.20E-10 | 3 L1 _ | 0610010K14Rik |
| Plec | 5.89E-15 | 0.498795 | 0.731 | 0.548 | 1.19E-10 | 3 L1 _ | Plec |
| Swap702 | 5.82E-15 | 0.43334 | 0.585 | 0.358 | 1.17E-10 | 3 L1 _ | Swap70 |
| Ubald2 | 5.44E-15 | 0.487763 | 0.615 | 0.414 | 1.09E-10 | 3 L1 _ | Ubald2 |
| Reep32 | 4.73E-15 | 0.481405 | 0.823 | 0.682 | 9.52E-11 | 3 L1 _ | Reep3 |
| Trim25 | 4.42E-15 | 0.497635 | 0.442 | 0.238 | 8.89E-11 | 3 L1 _ | Trim25 |
| Gbp3 | 4.20E-15 | 0.391122 | 0.377 | 0.177 | 8.47E-11 | 3 L1 _ | Gbp3 |
| Cxcl121 | 3.89E-15 | 0.885087 | 0.854 | 0.786 | 7.84E-11 | 3 L1 _ | Cxcl12 |
| Dleu22 | 3.09E-15 | 0.473697 | 0.465 | 0.244 | 6.23E-11 | 3 L1 _ | Dleu2 |
| Hsp90ab1 | 2.96E-15 | 0.331416 | 0.992 | 0.979 | 5.95E-11 | 3 L1 _ | Hsp90ab1 |
| Tns21 | 2.53E-15 | 0.460186 | 0.55 | 0.327 | 5.09E-11 | 3 L1 _ | Tns2 |
| Phlpp11 | 2.42E-15 | 0.34441 | 0.342 | 0.151 | 4.87E-11 | 3 L1 _ | Phlpp1 |
| Tspan141 | 2.38E-15 | 0.353368 | 0.238 | 0.086 | 4.80E-11 | 3 L1 _ | Tspan14 |
| Eogt1 | 2.21E-15 | 0.329868 | 0.2 | 0.062 | 4.45E-11 | 3 L1 _ | Eogt |
| Ifitm32 | 2.21E-15 | 0.36825 | 0.996 | 0.933 | 4.45E-11 | 3 L1 _ | Ifitm3 |
| Spata132 | 1.93E-15 | 0.315909 | 0.25 | 0.091 | 3.88E-11 | 3 L1 _ | Spata13 |

| | | | | | | | |
|---|---|---|---|---|---|---|---|
| H2-T231 | 1.84E-15 | 0.593093 | 0.715 | 0.513 | 3.71E-11 | 3 L1 _ | H2-T23 |
| Tap2 | 1.70E-15 | 0.527638 | 0.508 | 0.29 | 3.42E-11 | 3 L1 _ | Tap2 |
| Cavin3 | 1.61E-15 | 0.459101 | 0.881 | 0.716 | 3.24E-11 | 3 L1 _ | Cavin3 |
| Sox41 | 1.56E-15 | 0.674258 | 0.615 | 0.419 | 3.15E-11 | 3 L1 _ | Sox4 |
| Dennd5b1 | 1.54E-15 | 0.471016 | 0.415 | 0.204 | 3.09E-11 | 3 L1 _ | Dennd5b |
| Efhd12 | 1.32E-15 | 0.256675 | 0.138 | 0.031 | 2.65E-11 | 3 L1 _ | Efhd1 |
| Tagln2 | 1.23E-15 | 0.462539 | 0.877 | 0.789 | 2.48E-11 | 3 L1 _ | Tagln2 |
| Cpne81 | 1.20E-15 | 0.58864 | 0.519 | 0.315 | 2.41E-11 | 3 L1 _ | Cpne8 |
| Phldb21 | 1.07E-15 | 0.488722 | 0.496 | 0.28 | 2.15E-11 | 3 L1 _ | Phldb2 |
| Pola2 | 9.21E-16 | 0.400715 | 0.304 | 0.126 | 1.85E-11 | 3 L1 _ | Pola2 |
| Rnf213 | 6.86E-16 | 0.677032 | 0.5 | 0.284 | 1.38E-11 | 3 L1 _ | Rnf213 |
| Ptbp32 | 4.83E-16 | 0.563648 | 0.75 | 0.57 | 9.74E-12 | 3 L1 _ | Ptbp3 |
| Ralgapa11 | 4.06E-16 | 0.50753 | 0.519 | 0.305 | 8.17E-12 | 3 L1 _ | Ralgapa1 |
| Cables2 | 3.92E-16 | 0.307743 | 0.227 | 0.075 | 7.90E-12 | 3 L1 _ | Cables2 |
| St3gal51 | 3.74E-16 | 0.258175 | 0.165 | 0.043 | 7.53E-12 | 3 L1 _ | St3gal5 |
| Fam174b | 3.55E-16 | 0.375971 | 0.215 | 0.069 | 7.16E-12 | 3 L1 _ | Fam174b |
| Slco2b11 | 3.31E-16 | 0.389893 | 0.358 | 0.159 | 6.67E-12 | 3 L1 _ | Slco2b1 |
| Wwtr11 | 3.10E-16 | 0.498563 | 0.765 | 0.572 | 6.25E-12 | 3 L1 _ | Wwtr1 |
| Ppm1f1 | 2.44E-16 | 0.260936 | 0.231 | 0.076 | 4.91E-12 | 3 L1 _ | Ppm1f |
| Dcbld11 | 2.18E-16 | 0.351775 | 0.204 | 0.061 | 4.39E-12 | 3 L1 _ | Dcbld1 |
| Plekhg51 | 2.03E-16 | 0.304071 | 0.258 | 0.093 | 4.09E-12 | 3 L1 _ | Plekhg5 |
| Hmcn1 | 1.93E-16 | 0.607594 | 0.523 | 0.303 | 3.90E-12 | 3 L1 _ | Hmcn1 |
| Ifi471 | 1.85E-16 | 0.514889 | 0.496 | 0.256 | 3.73E-12 | 3 L1 _ | Ifi47 |
| Vgll41 | 1.80E-16 | 0.575117 | 0.677 | 0.444 | 3.63E-12 | 3 L1 _ | Vgll4 |
| Cpd2 | 1.73E-16 | 0.56692 | 0.673 | 0.461 | 3.48E-12 | 3 L1 _ | Cpd |
| Ace4 | 1.64E-16 | 0.563039 | 0.608 | 0.372 | 3.29E-12 | 3 L1 _ | Ace |
| Xaf1 | 1.43E-16 | 0.575361 | 0.596 | 0.363 | 2.87E-12 | 3 L1 _ | Xaf1 |
| Nes3 | 8.43E-17 | 0.398385 | 0.446 | 0.206 | 1.70E-12 | 3 L1 _ | Nes |
| Chst7 | 8.27E-17 | 0.297041 | 0.188 | 0.053 | 1.67E-12 | 3 L1 _ | Chst7 |
| Abcc42 | 7.64E-17 | 0.348595 | 0.246 | 0.083 | 1.54E-12 | 3 L1 _ | Abcc4 |
| Prkce1 | 7.51E-17 | 0.330294 | 0.269 | 0.097 | 1.51E-12 | 3 L1 _ | Prkce |
| Mxd41 | 6.24E-17 | 0.537009 | 0.873 | 0.763 | 1.26E-12 | 3 L1 _ | Mxd4 |
| Gpr182 | 6.16E-17 | 0.272677 | 0.15 | 0.034 | 1.24E-12 | 3 L1 _ | Gpr182 |
| Arhgef7 | 5.54E-17 | 0.507478 | 0.512 | 0.281 | 1.12E-12 | 3 L1 _ | Arhgef7 |
| Slc27a31 | 4.73E-17 | 0.331058 | 0.177 | 0.047 | 9.53E-13 | 3 L1 _ | Slc27a3 |
| Tut71 | 4.65E-17 | 0.58411 | 0.627 | 0.401 | 9.36E-13 | 3 L1 _ | Tut7 |
| Ralb | 3.99E-17 | 0.489039 | 0.519 | 0.291 | 8.04E-13 | 3 L1 _ | Ralb |
| Sorbs12 | 3.75E-17 | 0.273425 | 0.362 | 0.147 | 7.55E-13 | 3 L1 _ | Sorbs1 |
| Gm20559 | 3.53E-17 | 0.448248 | 0.496 | 0.259 | 7.10E-13 | 3 L1 _ | Gm20559 |
| Klf10 | 3.48E-17 | 0.45408 | 0.446 | 0.22 | 7.02E-13 | 3 L1 _ | Klf10 |
| Slc28a21 | 3.25E-17 | 0.300588 | 0.165 | 0.041 | 6.55E-13 | 3 L1 _ | Slc28a2 |

| | | | | | | | |
|---|---|---|---|---|---|---|---|
| Arhgap18 | 3.03E-17 | 0.671691 | 0.554 | 0.329 | 6.10E-13 | 3 L1 _ | Arhgap18 |
| Dennd31 | 2.85E-17 | 0.304459 | 0.192 | 0.053 | 5.73E-13 | 3 L1 _ | Dennd3 |
| Xiap1 | 2.72E-17 | 0.542794 | 0.715 | 0.504 | 5.48E-13 | 3 L1 _ | Xiap |
| Irf7 | 2.18E-17 | 0.626969 | 0.612 | 0.369 | 4.40E-13 | 3 L1 _ | Irf7 |
| Rnf1522 | 1.84E-17 | 0.30105 | 0.223 | 0.066 | 3.70E-13 | 3 L1 _ | Rnf152 |
| Lbr1 | 1.52E-17 | 0.450234 | 0.419 | 0.194 | 3.07E-13 | 3 L1 _ | Lbr |
| Myo102 | 1.48E-17 | 0.564408 | 0.65 | 0.429 | 2.98E-13 | 3 L1 _ | Myo10 |
| Fnbp1l1 | 1.43E-17 | 0.461735 | 0.392 | 0.182 | 2.89E-13 | 3 L1 _ | Fnbp1l |
| Foxo11 | 1.40E-17 | 0.524671 | 0.488 | 0.263 | 2.83E-13 | 3 L1 _ | Foxo1 |
| Hey11 | 1.16E-17 | 0.340915 | 0.15 | 0.033 | 2.34E-13 | 3 L1 _ | Hey1 |
| Arhgef32 | 1.11E-17 | 0.395971 | 0.269 | 0.094 | 2.24E-13 | 3 L1 _ | Arhgef3 |
| Il10rb1 | 8.33E-18 | 0.522212 | 0.581 | 0.33 | 1.68E-13 | 3 L1 _ | Il10rb |
| Vwa11 | 6.17E-18 | 0.55588 | 0.562 | 0.311 | 1.24E-13 | 3 L1 _ | Vwa1 |
| Btg3 | 4.81E-18 | 0.474275 | 0.477 | 0.242 | 9.68E-14 | 3 L1 _ | Btg3 |
| Sos11 | 4.18E-18 | 0.529746 | 0.388 | 0.177 | 8.42E-14 | 3 L1 _ | Sos1 |
| Cemip2 | 2.91E-18 | 0.43747 | 0.438 | 0.202 | 5.85E-14 | 3 L1 _ | Cemip2 |
| Esm11 | 2.88E-18 | 0.706663 | 0.146 | 0.03 | 5.81E-14 | 3 L1 _ | Esm1 |
| Zfp6641 | 2.59E-18 | 0.460322 | 0.523 | 0.282 | 5.22E-14 | 3 L1 _ | Zfp664 |
| Samd9l1 | 2.53E-18 | 0.690527 | 0.585 | 0.347 | 5.10E-14 | 3 L1 _ | Samd9l |
| Rhoc | 2.13E-18 | 0.593767 | 0.831 | 0.702 | 4.28E-14 | 3 L1 _ | Rhoc |
| Ktn13 | 2.11E-18 | 0.485164 | 0.792 | 0.605 | 4.26E-14 | 3 L1 _ | Ktn1 |
| Fat41 | 1.99E-18 | 0.438464 | 0.258 | 0.086 | 4.00E-14 | 3 L1 _ | Fat4 |
| Nfe2l11 | 1.63E-18 | 0.599422 | 0.8 | 0.622 | 3.27E-14 | 3 L1 _ | Nfe2l1 |
| Psme12 | 1.43E-18 | 0.574822 | 0.854 | 0.714 | 2.88E-14 | 3 L1 _ | Psme1 |
| Ifi213 | 9.05E-19 | 0.358929 | 0.165 | 0.038 | 1.82E-14 | 3 L1 _ | Ifi213 |
| Actn42 | 8.15E-19 | 0.484077 | 0.838 | 0.705 | 1.64E-14 | 3 L1 _ | Actn4 |
| Sh3bgrl21 | 7.89E-19 | 0.303194 | 0.162 | 0.036 | 1.59E-14 | 3 L1 _ | Sh3bgrl2 |
| Gng113 | 7.53E-19 | 0.345787 | 0.935 | 0.806 | 1.52E-14 | 3 L1 _ | Gng11 |
| Pggt1b | 5.86E-19 | 0.539807 | 0.477 | 0.249 | 1.18E-14 | 3 L1 _ | Pggt1b |
| Kctd12 | 5.46E-19 | 0.610954 | 0.742 | 0.531 | 1.10E-14 | 3 L1 _ | Kctd12 |
| Gnaq2 | 5.34E-19 | 0.566541 | 0.665 | 0.419 | 1.07E-14 | 3 L1 _ | Gnaq |
| Prpf40b | 5.18E-19 | 0.475823 | 0.281 | 0.1 | 1.04E-14 | 3 L1 _ | Prpf40b |
| Ier5l | 3.45E-19 | 0.583974 | 0.55 | 0.306 | 6.95E-15 | 3 L1 _ | Ier5l |
| Serinc31 | 3.40E-19 | 0.497123 | 0.931 | 0.851 | 6.85E-15 | 3 L1 _ | Serinc3 |
| Hhex1 | 3.18E-19 | 0.38684 | 0.227 | 0.066 | 6.40E-15 | 3 L1 _ | Hhex |
| Oip5os1 | 2.41E-19 | 0.499712 | 0.681 | 0.426 | 4.86E-15 | 3 L1 _ | Oip5os1 |
| N4bp1 | 2.25E-19 | 0.546743 | 0.508 | 0.27 | 4.53E-15 | 3 L1 _ | N4bp1 |
| Hdac72 | 1.85E-19 | 0.550153 | 0.662 | 0.426 | 3.72E-15 | 3 L1 _ | Hdac7 |
| Fry3 | 1.74E-19 | 0.343036 | 0.415 | 0.164 | 3.51E-15 | 3 L1 _ | Fry |
| Ptp4a32 | 1.45E-19 | 0.556546 | 0.554 | 0.299 | 2.92E-15 | 3 L1 _ | Ptp4a3 |
| Cd813 | 1.38E-19 | 0.45361 | 0.981 | 0.886 | 2.79E-15 | 3 L1 _ | Cd81 |

| | | | | | | | |
|---|---|---|---|---|---|---|---|
| Mcam2 | 1.30E-19 | 0.361158 | 0.346 | 0.125 | 2.63E-15 | 3 L1 _ | Mcam |
| Parp141 | 1.24E-19 | 0.6567 | 0.623 | 0.363 | 2.49E-15 | 3 L1 _ | Parp14 |
| Gimap82 | 1.13E-19 | 0.496664 | 0.304 | 0.108 | 2.27E-15 | 3 L1 _ | Gimap8 |
| Arl151 | 8.63E-20 | 0.445721 | 0.304 | 0.108 | 1.74E-15 | 3 L1 _ | Arl15 |
| Tmcc31 | 6.82E-20 | 0.507698 | 0.362 | 0.143 | 1.37E-15 | 3 L1 _ | Tmcc3 |
| Myo1c1 | 6.78E-20 | 0.600429 | 0.627 | 0.391 | 1.37E-15 | 3 L1 _ | Myo1c |
| Gbp71 | 5.97E-20 | 0.709728 | 0.665 | 0.428 | 1.20E-15 | 3 L1 _ | Gbp7 |
| Git11 | 5.43E-20 | 0.440457 | 0.408 | 0.176 | 1.09E-15 | 3 L1 _ | Git1 |
| Phf11d | 5.03E-20 | 0.54015 | 0.488 | 0.236 | 1.01E-15 | 3 L1 _ | Phf11d |
| Lyn1 | 3.39E-20 | 0.46676 | 0.4 | 0.167 | 6.83E-16 | 3 L1 _ | Lyn |
| Dhrs32 | 3.25E-20 | 0.625129 | 0.635 | 0.395 | 6.55E-16 | 3 L1 _ | Dhrs3 |
| Gnas3 | 2.75E-20 | 0.36395 | 0.981 | 0.959 | 5.54E-16 | 3 L1 _ | Gnas |
| Rapgef11 | 2.63E-20 | 0.495048 | 0.431 | 0.192 | 5.30E-16 | 3 L1 _ | Rapgef1 |
| Lnx21 | 2.44E-20 | 0.376145 | 0.227 | 0.062 | 4.92E-16 | 3 L1 _ | Lnx2 |
| Taok2 | 2.32E-20 | 0.627515 | 0.488 | 0.258 | 4.67E-16 | 3 L1 _ | Taok2 |
| Gda2 | 2.21E-20 | 0.368295 | 0.231 | 0.065 | 4.45E-16 | 3 L1 _ | Gda |
| Fryl1 | 2.08E-20 | 0.62382 | 0.469 | 0.231 | 4.18E-16 | 3 L1 _ | Fryl |
| Mast4 | 1.94E-20 | 0.706082 | 0.708 | 0.486 | 3.92E-16 | 3 L1 _ | Mast4 |
| Wars1 | 1.43E-20 | 0.520144 | 0.4 | 0.174 | 2.89E-16 | 3 L1 _ | Wars |
| Zbtb46 | 1.35E-20 | 0.390502 | 0.369 | 0.146 | 2.72E-16 | 3 L1 _ | Zbtb46 |
| Cd1513 | 1.30E-20 | 0.631816 | 0.758 | 0.537 | 2.61E-16 | 3 L1 _ | Cd151 |
| Fxyd51 | 1.22E-20 | 0.684979 | 0.892 | 0.73 | 2.46E-16 | 3 L1 _ | Fxyd5 |
| Piezo22 | 1.21E-20 | 0.580234 | 0.596 | 0.331 | 2.43E-16 | 3 L1 _ | Piezo2 |
| 4833403J16Rik1 | 1.18E-20 | 0.285851 | 0.192 | 0.045 | 2.38E-16 | 3 L1 _ | 4833403J16Rik |
| B2m2 | 1.18E-20 | 0.462734 | 0.985 | 0.947 | 2.38E-16 | 3 L1 _ | B2m |
| Tgtp2 | 1.16E-20 | 0.392844 | 0.181 | 0.041 | 2.34E-16 | 3 L1 _ | Tgtp2 |
| Crybg31 | 1.09E-20 | 0.597903 | 0.481 | 0.236 | 2.20E-16 | 3 L1 _ | Crybg3 |
| Syne23 | 1.03E-20 | 0.640398 | 0.658 | 0.393 | 2.07E-16 | 3 L1 _ | Syne2 |
| Zfpm12 | 9.79E-21 | 0.427648 | 0.392 | 0.16 | 1.97E-16 | 3 L1 _ | Zfpm1 |
| Nck11 | 9.13E-21 | 0.662186 | 0.735 | 0.498 | 1.84E-16 | 3 L1 _ | Nck1 |
| Ctnna11 | 8.75E-21 | 0.503056 | 0.815 | 0.654 | 1.76E-16 | 3 L1 _ | Ctnna1 |
| Nfkbia1 | 8.66E-21 | 0.69599 | 0.762 | 0.516 | 1.74E-16 | 3 L1 _ | Nfkbia |
| Qk1 | 8.24E-21 | 0.6713 | 0.808 | 0.667 | 1.66E-16 | 3 L1 _ | Qk |
| Dusp31 | 8.06E-21 | 0.585605 | 0.592 | 0.332 | 1.62E-16 | 3 L1 _ | Dusp3 |
| Tmed51 | 7.88E-21 | 0.640896 | 0.638 | 0.387 | 1.59E-16 | 3 L1 _ | Tmed5 |
| Gbp5 | 7.46E-21 | 0.450569 | 0.281 | 0.09 | 1.50E-16 | 3 L1 _ | Gbp5 |
| Dnm31 | 7.34E-21 | 0.682003 | 0.477 | 0.238 | 1.48E-16 | 3 L1 _ | Dnm3 |
| Cmah1 | 5.44E-21 | 0.385308 | 0.181 | 0.039 | 1.10E-16 | 3 L1 _ | Cmah |
| Calcrl1 | 5.18E-21 | 0.593732 | 0.55 | 0.287 | 1.04E-16 | 3 L1 _ | Calcrl |
| Yes1 | 3.26E-21 | 0.538871 | 0.404 | 0.171 | 6.56E-17 | 3 L1 _ | Yes1 |
| Sypl1 | 2.88E-21 | 0.661259 | 0.685 | 0.446 | 5.80E-17 | 3 L1 _ | Sypl |

| | | | | | | | |
|---|---|---|---|---|---|---|---|
| Ccnd12 | 2.88E-21 | 0.351716 | 0.4 | 0.15 | 5.79E-17 | 3 L1 _ | Ccnd1 |
| Ankrd291 | 2.86E-21 | 0.254254 | 0.188 | 0.042 | 5.75E-17 | 3 L1 _ | Ankrd29 |
| Tspan153 | 2.25E-21 | 0.313759 | 0.4 | 0.144 | 4.52E-17 | 3 L1 _ | Tspan15 |
| Tinagl13 | 1.81E-21 | 0.251409 | 0.635 | 0.269 | 3.65E-17 | 3 L1 _ | Tinagl1 |
| Trpv41 | 1.73E-21 | 0.298976 | 0.131 | 0.02 | 3.48E-17 | 3 L1 _ | Trpv4 |
| Sema6d3 | 1.32E-21 | 0.629488 | 0.612 | 0.328 | 2.66E-17 | 3 L1 _ | Sema6d |
| Ptma2 | 1.15E-21 | 0.379125 | 0.996 | 0.986 | 2.32E-17 | 3 L1 _ | Ptma |
| Gcnt23 | 1.08E-21 | 0.338774 | 0.45 | 0.176 | 2.18E-17 | 3 L1 _ | Gcnt2 |
| Rapgef22 | 9.08E-22 | 0.424131 | 0.469 | 0.199 | 1.83E-17 | 3 L1 _ | Rapgef2 |
| Tmsb4x3 | 9.08E-22 | 0.352099 | 1 | 0.994 | 1.83E-17 | 3 L1 _ | Tmsb4x |
| Ppp1r16b2 | 7.71E-22 | 0.417581 | 0.292 | 0.092 | 1.55E-17 | 3 L1 _ | Ppp1r16b |
| 2-Jun | 6.77E-22 | 0.9215 | 0.865 | 0.676 | 1.36E-17 | 3 L1 _ | Jun |
| Myo61 | 6.58E-22 | 0.874373 | 0.558 | 0.33 | 1.33E-17 | 3 L1 _ | Myo6 |
| Prdm161 | 6.50E-22 | 0.345375 | 0.181 | 0.039 | 1.31E-17 | 3 L1 _ | Prdm16 |
| Sh2d3c2 | 6.39E-22 | 0.486027 | 0.35 | 0.127 | 1.29E-17 | 3 L1 _ | Sh2d3c |
| Tns13 | 5.63E-22 | 0.530782 | 0.781 | 0.503 | 1.13E-17 | 3 L1 _ | Tns1 |
| Pde8a2 | 4.99E-22 | 0.546427 | 0.423 | 0.175 | 1.00E-17 | 3 L1 _ | Pde8a |
| Stard83 | 4.74E-22 | 0.466306 | 0.45 | 0.186 | 9.54E-18 | 3 L1 _ | Stard8 |
| Tnfaip8l1 | 4.35E-22 | 0.264359 | 0.127 | 0.018 | 8.75E-18 | 3 L1 _ | Tnfaip8l1 |
| Adamts91 | 4.10E-22 | 0.579422 | 0.404 | 0.166 | 8.26E-18 | 3 L1 _ | Adamts9 |
| Sipa1l21 | 3.83E-22 | 0.3647 | 0.269 | 0.079 | 7.71E-18 | 3 L1 _ | Sipa1l2 |
| Glul1 | 3.69E-22 | 1.161216 | 0.512 | 0.273 | 7.44E-18 | 3 L1 _ | Glul |
| Ifi2081 | 3.63E-22 | 0.470298 | 0.219 | 0.055 | 7.32E-18 | 3 L1 _ | Ifi208 |
| Adam101 | 3.24E-22 | 0.622445 | 0.554 | 0.296 | 6.52E-18 | 3 L1 _ | Adam10 |
| Plcg11 | 3.20E-22 | 0.637093 | 0.477 | 0.23 | 6.44E-18 | 3 L1 _ | Plcg1 |
| Ppia1 | 2.77E-22 | 0.404726 | 1 | 0.982 | 5.59E-18 | 3 L1 _ | Ppia |
| Tbkbp1 | 2.04E-22 | 0.418308 | 0.262 | 0.076 | 4.10E-18 | 3 L1 _ | Tbkbp1 |
| Castor1 | 1.71E-22 | 0.446421 | 0.315 | 0.105 | 3.44E-18 | 3 L1 _ | Castor1 |
| Als2cl1 | 1.70E-22 | 0.310282 | 0.196 | 0.043 | 3.43E-18 | 3 L1 _ | Als2cl |
| Fez23 | 1.58E-22 | 0.568264 | 0.642 | 0.348 | 3.17E-18 | 3 L1 _ | Fez2 |
| Lmo71 | 1.40E-22 | 0.384239 | 0.196 | 0.044 | 2.83E-18 | 3 L1 _ | Lmo7 |
| Arhgef121 | 1.36E-22 | 0.601549 | 0.838 | 0.65 | 2.75E-18 | 3 L1 _ | Arhgef12 |
| Elmo12 | 1.22E-22 | 0.552093 | 0.35 | 0.127 | 2.45E-18 | 3 L1 _ | Elmo1 |
| Ccdc28b1 | 1.10E-22 | 0.567748 | 0.585 | 0.314 | 2.22E-18 | 3 L1 _ | Ccdc28b |
| Lrrc3b1 | 7.39E-23 | 0.303633 | 0.115 | 0.014 | 1.49E-18 | 3 L1 _ | Lrrc3b |
| Hs3st11 | 6.18E-23 | 0.324293 | 0.165 | 0.03 | 1.25E-18 | 3 L1 _ | Hs3st1 |
| Pcdh121 | 5.05E-23 | 0.5768 | 0.365 | 0.139 | 1.02E-18 | 3 L1 _ | Pcdh12 |
| Cdc42bpb1 | 4.52E-23 | 0.603795 | 0.546 | 0.283 | 9.10E-19 | 3 L1 _ | Cdc42bpb |
| Bst21 | 4.28E-23 | 0.784054 | 0.896 | 0.693 | 8.62E-19 | 3 L1 _ | Bst2 |
| Plekhg11 | 3.63E-23 | 0.485366 | 0.4 | 0.161 | 7.32E-19 | 3 L1 _ | Plekhg1 |
| H2-Q71 | 3.12E-23 | 0.642166 | 0.773 | 0.487 | 6.29E-19 | 3 L1 _ | H2-Q7 |

| | | | | | | | |
|---|---|---|---|---|---|---|---|
| Bend71 | 2.99E-23 | 0.281745 | 0.165 | 0.03 | 6.02E-19 | 3 L1 _ | Bend7 |
| Bambi | 2.97E-23 | 0.31386 | 0.181 | 0.037 | 5.97E-19 | 3 L1 _ | Bambi |
| H2-Q4 | 2.36E-23 | 0.67499 | 0.531 | 0.263 | 4.76E-19 | 3 L1 _ | H2-Q4 |
| Rab11a1 | 2.18E-23 | 0.672322 | 0.673 | 0.415 | 4.40E-19 | 3 L1 _ | Rab11a |
| S100a162 | 2.18E-23 | 0.617899 | 0.842 | 0.567 | 4.39E-19 | 3 L1 _ | S100a16 |
| Fmnl31 | 2.01E-23 | 0.580728 | 0.535 | 0.27 | 4.05E-19 | 3 L1 _ | Fmnl3 |
| Git21 | 1.09E-23 | 0.70084 | 0.638 | 0.381 | 2.20E-19 | 3 L1 _ | Git2 |
| Hmg20b | 1.05E-23 | 0.647247 | 0.731 | 0.498 | 2.11E-19 | 3 L1 _ | Hmg20b |
| Piezo11 | 8.35E-24 | 0.490501 | 0.323 | 0.109 | 1.68E-19 | 3 L1 _ | Piezo1 |
| Cldn15 | 7.72E-24 | 0.372132 | 0.146 | 0.023 | 1.55E-19 | 3 L1 _ | Cldn15 |
| Macf11 | 6.80E-24 | 0.700189 | 0.923 | 0.798 | 1.37E-19 | 3 L1 _ | Macf1 |
| Ablim12 | 6.55E-24 | 0.724873 | 0.638 | 0.369 | 1.32E-19 | 3 L1 _ | Ablim1 |
| Tpm32 | 5.87E-24 | 0.630862 | 0.885 | 0.736 | 1.18E-19 | 3 L1 _ | Tpm3 |
| Ndufa42 | 4.59E-24 | 0.633442 | 0.892 | 0.717 | 9.25E-20 | 3 L1 _ | Ndufa4 |
| H2-Q61 | 3.82E-24 | 0.773807 | 0.535 | 0.263 | 7.69E-20 | 3 L1 _ | H2-Q6 |
| Smad1 | 3.28E-24 | 0.689175 | 0.45 | 0.198 | 6.62E-20 | 3 L1 _ | Smad1 |
| Prkd21 | 2.94E-24 | 0.438761 | 0.342 | 0.117 | 5.93E-20 | 3 L1 _ | Prkd2 |
| Flywch11 | 2.46E-24 | 0.674589 | 0.535 | 0.274 | 4.96E-20 | 3 L1 _ | Flywch1 |
| Aig12 | 2.30E-24 | 0.606545 | 0.469 | 0.199 | 4.63E-20 | 3 L1 _ | Aig1 |
| Plcb42 | 2.24E-24 | 0.565047 | 0.512 | 0.227 | 4.52E-20 | 3 L1 _ | Plcb4 |
| St3gal62 | 1.31E-24 | 0.617791 | 0.473 | 0.204 | 2.63E-20 | 3 L1 _ | St3gal6 |
| Arhgef51 | 9.59E-25 | 0.564823 | 0.404 | 0.157 | 1.93E-20 | 3 L1 _ | Arhgef5 |
| Hyal21 | 9.52E-25 | 0.46092 | 0.331 | 0.107 | 1.92E-20 | 3 L1 _ | Hyal2 |
| Nrp21 | 5.34E-25 | 0.635305 | 0.608 | 0.317 | 1.08E-20 | 3 L1 _ | Nrp2 |
| 9930111J21Rik22 | 3.83E-25 | 0.657186 | 0.696 | 0.41 | 7.71E-21 | 3 L1 _ | 9930111J21Rik2 |
| Map3k111 | 3.80E-25 | 0.437946 | 0.296 | 0.087 | 7.66E-21 | 3 L1 _ | Map3k11 |
| Psmb92 | 3.43E-25 | 0.59974 | 0.662 | 0.349 | 6.91E-21 | 3 L1 _ | Psmb9 |
| Thsd11 | 2.66E-25 | 0.407883 | 0.215 | 0.048 | 5.35E-21 | 3 L1 _ | Thsd1 |
| S100a131 | 2.46E-25 | 0.621497 | 0.896 | 0.741 | 4.95E-21 | 3 L1 _ | S100a13 |
| Abi32 | 2.02E-25 | 0.621196 | 0.431 | 0.167 | 4.07E-21 | 3 L1 _ | Abi3 |
| Nhsl21 | 1.80E-25 | 0.619915 | 0.438 | 0.177 | 3.62E-21 | 3 L1 _ | Nhsl2 |
| Rasal22 | 1.79E-25 | 0.61453 | 0.604 | 0.303 | 3.60E-21 | 3 L1 _ | Rasal2 |
| Pcdh13 | 1.69E-25 | 0.465555 | 0.415 | 0.151 | 3.40E-21 | 3 L1 _ | Pcdh1 |
| Pitpnc11 | 1.68E-25 | 0.741171 | 0.631 | 0.342 | 3.39E-21 | 3 L1 _ | Pitpnc1 |
| Acer21 | 1.46E-25 | 0.266947 | 0.192 | 0.036 | 2.94E-21 | 3 L1 _ | Acer2 |
| Atp8a11 | 1.46E-25 | 0.534591 | 0.331 | 0.109 | 2.94E-21 | 3 L1 _ | Atp8a1 |
| 2-Mar | 1.43E-25 | 0.813416 | 0.762 | 0.535 | 2.88E-21 | 3 L1 _ | 2-Mar |
| Plscr11 | 1.07E-25 | 0.720119 | 0.423 | 0.172 | 2.16E-21 | 3 L1 _ | Plscr1 |
| Myzap | 1.03E-25 | 0.259258 | 0.138 | 0.018 | 2.08E-21 | 3 L1 _ | Myzap |
| Pik3r6 | 4.17E-26 | 0.254783 | 0.142 | 0.019 | 8.39E-22 | 3 L1 _ | Pik3r6 |
| Lpcat11 | 4.02E-26 | 0.658059 | 0.438 | 0.177 | 8.10E-22 | 3 L1 _ | Lpcat1 |

| | | | | | | | |
|---|---|---|---|---|---|---|---|
| Gimap52 | 3.04E-26 | 0.475964 | 0.477 | 0.183 | 6.12E-22 | 3 L1 _ | Gimap5 |
| Numb1 | 2.05E-26 | 0.666926 | 0.523 | 0.238 | 4.12E-22 | 3 L1 _ | Numb |
| Bnip21 | 2.02E-26 | 0.705873 | 0.773 | 0.567 | 4.07E-22 | 3 L1 _ | Bnip2 |
| Iigp12 | 1.83E-26 | 0.737146 | 0.696 | 0.391 | 3.69E-22 | 3 L1 _ | Iigp1 |
| Lysmd2 | 1.66E-26 | 0.638994 | 0.585 | 0.287 | 3.34E-22 | 3 L1 _ | Lysmd2 |
| Csgalnact11 | 1.23E-26 | 0.519415 | 0.419 | 0.154 | 2.47E-22 | 3 L1 _ | Csgalnact1 |
| Galnt151 | 8.90E-27 | 0.45012 | 0.346 | 0.108 | 1.79E-22 | 3 L1 _ | Galnt15 |
| Inka1 | 7.63E-27 | 0.538243 | 0.419 | 0.159 | 1.54E-22 | 3 L1 _ | Inka1 |
| Grap3 | 7.22E-27 | 0.39215 | 0.369 | 0.12 | 1.45E-22 | 3 L1 _ | Grap |
| Exoc3l1 | 7.16E-27 | 0.262498 | 0.154 | 0.022 | 1.44E-22 | 3 L1 _ | Exoc3l |
| Irf12 | 6.81E-27 | 0.653751 | 0.619 | 0.325 | 1.37E-22 | 3 L1 _ | Irf1 |
| Mapk121 | 5.73E-27 | 0.473678 | 0.365 | 0.121 | 1.15E-22 | 3 L1 _ | Mapk12 |
| Timp33 | 4.18E-27 | 0.653792 | 0.965 | 0.77 | 8.41E-23 | 3 L1 _ | Timp3 |
| Apold12 | 2.47E-27 | 0.442844 | 0.258 | 0.06 | 4.98E-23 | 3 L1 _ | Apold1 |
| Luzp11 | 2.35E-27 | 0.709793 | 0.8 | 0.575 | 4.73E-23 | 3 L1 _ | Luzp1 |
| Tcf43 | 1.82E-27 | 0.576539 | 0.969 | 0.879 | 3.67E-23 | 3 L1 _ | Tcf4 |
| Sptan11 | 1.70E-27 | 0.747377 | 0.838 | 0.647 | 3.43E-23 | 3 L1 _ | Sptan1 |
| Anp32a1 | 1.44E-27 | 0.739368 | 0.831 | 0.644 | 2.89E-23 | 3 L1 _ | Anp32a |
| Clic41 | 1.08E-27 | 0.705377 | 0.812 | 0.557 | 2.17E-23 | 3 L1 _ | Clic4 |
| H2-D11 | 9.64E-28 | 0.610173 | 0.981 | 0.951 | 1.94E-23 | 3 L1 _ | H2-D1 |
| Gbp91 | 9.44E-28 | 0.592091 | 0.415 | 0.155 | 1.90E-23 | 3 L1 _ | Gbp9 |
| Dab2ip3 | 8.20E-28 | 0.726351 | 0.75 | 0.466 | 1.65E-23 | 3 L1 _ | Dab2ip |
| Car71 | 7.22E-28 | 0.321751 | 0.162 | 0.023 | 1.45E-23 | 3 L1 _ | Car7 |
| Slc30a11 | 6.43E-28 | 0.536718 | 0.331 | 0.099 | 1.30E-23 | 3 L1 _ | Slc30a1 |
| Bmpr21 | 5.69E-28 | 0.755427 | 0.846 | 0.624 | 1.15E-23 | 3 L1 _ | Bmpr2 |
| Bik | 5.33E-28 | 0.330607 | 0.208 | 0.039 | 1.07E-23 | 3 L1 _ | Bik |
| Fam13c | 3.63E-28 | 0.607624 | 0.492 | 0.204 | 7.31E-24 | 3 L1 _ | Fam13c |
| Gipc31 | 2.29E-28 | 0.26852 | 0.15 | 0.019 | 4.62E-24 | 3 L1 _ | Gipc3 |
| Pxn1 | 2.17E-28 | 0.572778 | 0.462 | 0.179 | 4.36E-24 | 3 L1 _ | Pxn |
| Fgd61 | 1.68E-28 | 0.326978 | 0.2 | 0.036 | 3.39E-24 | 3 L1 _ | Fgd6 |
| Foxf11 | 1.55E-28 | 0.315521 | 0.162 | 0.023 | 3.13E-24 | 3 L1 _ | Foxf1 |
| Alpl1 | 1.48E-28 | 0.659157 | 0.246 | 0.056 | 2.97E-24 | 3 L1 _ | Alpl |
| Cd2ap1 | 1.19E-28 | 0.757309 | 0.8 | 0.548 | 2.40E-24 | 3 L1 _ | Cd2ap |
| Tpst22 | 8.73E-29 | 0.750568 | 0.569 | 0.274 | 1.76E-24 | 3 L1 _ | Tpst2 |
| Klhl51 | 8.56E-29 | 0.599477 | 0.504 | 0.21 | 1.72E-24 | 3 L1 _ | Klhl5 |
| Gimap42 | 5.90E-29 | 0.533345 | 0.623 | 0.268 | 1.19E-24 | 3 L1 _ | Gimap4 |
| St8sia42 | 5.07E-29 | 0.630916 | 0.496 | 0.188 | 1.02E-24 | 3 L1 _ | St8sia4 |
| Rgs121 | 4.04E-29 | 0.494316 | 0.365 | 0.117 | 8.13E-25 | 3 L1 _ | Rgs12 |
| Casz11 | 2.91E-29 | 0.283563 | 0.146 | 0.017 | 5.85E-25 | 3 L1 _ | Casz1 |
| Klhl4 | 2.51E-29 | 0.26938 | 0.131 | 0.013 | 5.06E-25 | 3 L1 _ | Klhl4 |
| Magi11 | 1.51E-29 | 0.436849 | 0.219 | 0.042 | 3.04E-25 | 3 L1 _ | Magi1 |

| | | | | | | | |
|---|---|---|---|---|---|---|---|
| Tspan91 | 8.50E-30 | 0.678486 | 0.565 | 0.263 | 1.71E-25 | 3 L1 _ | Tspan9 |
| Col4a13 | 6.13E-30 | 0.777864 | 0.981 | 0.86 | 1.23E-25 | 3 L1 _ | Col4a1 |
| Col4a22 | 4.27E-30 | 0.755172 | 0.954 | 0.844 | 8.61E-26 | 3 L1 _ | Col4a2 |
| Clec2d1 | 2.90E-30 | 0.82268 | 0.858 | 0.625 | 5.85E-26 | 3 L1 _ | Clec2d |
| Pim31 | 2.87E-30 | 0.595291 | 0.412 | 0.139 | 5.78E-26 | 3 L1 _ | Pim3 |
| Tmem441 | 2.20E-30 | 0.396181 | 0.192 | 0.031 | 4.44E-26 | 3 L1 _ | Tmem44 |
| H2-K11 | 9.77E-31 | 0.621424 | 0.996 | 0.927 | 1.97E-26 | 3 L1 _ | H2-K1 |
| Lama51 | 8.92E-31 | 0.391172 | 0.242 | 0.049 | 1.80E-26 | 3 L1 _ | Lama5 |
| Arpc31 | 6.92E-31 | 0.704379 | 0.923 | 0.785 | 1.39E-26 | 3 L1 _ | Arpc3 |
| Tanc11 | 4.24E-31 | 0.683962 | 0.596 | 0.287 | 8.54E-27 | 3 L1 _ | Tanc1 |
| Sec14l11 | 4.01E-31 | 0.736257 | 0.623 | 0.314 | 8.07E-27 | 3 L1 _ | Sec14l1 |
| Mcc1 | 2.88E-31 | 0.564247 | 0.419 | 0.143 | 5.79E-27 | 3 L1 _ | Mcc |
| Jcad1 | 2.55E-31 | 0.712669 | 0.496 | 0.204 | 5.14E-27 | 3 L1 _ | Jcad |
| Rgl11 | 2.20E-31 | 0.850763 | 0.715 | 0.433 | 4.43E-27 | 3 L1 _ | Rgl1 |
| Tal11 | 2.15E-31 | 0.263243 | 0.131 | 0.012 | 4.33E-27 | 3 L1 _ | Tal1 |
| Klf71 | 2.10E-31 | 0.831934 | 0.777 | 0.522 | 4.23E-27 | 3 L1 _ | Klf7 |
| Efnb11 | 9.36E-32 | 0.737685 | 0.515 | 0.207 | 1.89E-27 | 3 L1 _ | Efnb1 |
| Gm34455 | 8.00E-32 | 0.549005 | 0.273 | 0.061 | 1.61E-27 | 3 L1 _ | Gm34455 |
| Hoxd81 | 7.80E-32 | 0.400472 | 0.254 | 0.053 | 1.57E-27 | 3 L1 _ | Hoxd8 |
| Slfn31 | 7.01E-32 | 0.306247 | 0.138 | 0.013 | 1.41E-27 | 3 L1 _ | Slfn3 |
| Sybu1 | 5.25E-32 | 0.422387 | 0.2 | 0.032 | 1.06E-27 | 3 L1 _ | Sybu |
| Ttll72 | 4.43E-32 | 0.474687 | 0.304 | 0.072 | 8.93E-28 | 3 L1 _ | Ttll7 |
| B3gnt31 | 3.22E-32 | 0.385627 | 0.25 | 0.051 | 6.49E-28 | 3 L1 _ | B3gnt3 |
| Dusp71 | 2.83E-32 | 0.643306 | 0.462 | 0.171 | 5.70E-28 | 3 L1 _ | Dusp7 |
| Enah1 | 2.59E-32 | 0.581869 | 0.296 | 0.069 | 5.22E-28 | 3 L1 _ | Enah |
| Fzd43 | 1.57E-32 | 0.571744 | 0.708 | 0.304 | 3.16E-28 | 3 L1 _ | Fzd4 |
| Sgk11 | 1.46E-32 | 0.92132 | 0.581 | 0.256 | 2.95E-28 | 3 L1 _ | Sgk1 |
| Slc26a101 | 1.37E-32 | 0.528543 | 0.196 | 0.029 | 2.76E-28 | 3 L1 _ | Slc26a10 |
| Foxp11 | 1.25E-32 | 0.840813 | 0.804 | 0.526 | 2.51E-28 | 3 L1 _ | Foxp1 |
| Ybx11 | 1.05E-32 | 0.674979 | 0.985 | 0.899 | 2.11E-28 | 3 L1 _ | Ybx1 |
| Klf3 | 1.03E-32 | 0.816944 | 0.715 | 0.432 | 2.07E-28 | 3 L1 _ | Klf3 |
| Gm326881 | 9.88E-33 | 0.308807 | 0.177 | 0.023 | 1.99E-28 | 3 L1 _ | Gm32688 |
| Sox132 | 9.74E-33 | 0.41097 | 0.296 | 0.067 | 1.96E-28 | 3 L1 _ | Sox13 |
| Psmb82 | 8.75E-33 | 0.815095 | 0.869 | 0.604 | 1.76E-28 | 3 L1 _ | Psmb8 |
| Tdrp1 | 7.80E-33 | 0.376836 | 0.235 | 0.043 | 1.57E-28 | 3 L1 _ | Tdrp |
| 2900026A02Rik1 | 5.43E-33 | 0.797683 | 0.585 | 0.27 | 1.09E-28 | 3 L1 _ | 2900026A02Rik |
| Utrn3 | 4.60E-33 | 0.858421 | 0.769 | 0.456 | 9.26E-29 | 3 L1 _ | Utrn |
| Nrp11 | 3.68E-33 | 0.825451 | 0.942 | 0.783 | 7.41E-29 | 3 L1 _ | Nrp1 |
| Smad72 | 2.37E-33 | 0.847812 | 0.723 | 0.374 | 4.78E-29 | 3 L1 _ | Smad7 |
| Epb41l4a | 1.49E-33 | 0.276275 | 0.138 | 0.012 | 3.01E-29 | 3 L1 _ | Epb41l4a |
| Wasf21 | 1.49E-33 | 0.745053 | 0.865 | 0.71 | 3.01E-29 | 3 L1 _ | Wasf2 |

| | | | | | | | |
|---|---|---|---|---|---|---|---|
| Ptpn141 | 9.47E-34 | 0.594228 | 0.354 | 0.098 | 1.91E-29 | 3 L1 _ | Ptpn14 |
| Rtl8b1 | 8.79E-34 | 0.758486 | 0.573 | 0.26 | 1.77E-29 | 3 L1 _ | Rtl8b |
| Tjp11 | 8.43E-34 | 0.807372 | 0.681 | 0.364 | 1.70E-29 | 3 L1 _ | Tjp1 |
| Stox21 | 7.42E-34 | 0.709925 | 0.481 | 0.184 | 1.49E-29 | 3 L1 _ | Stox2 |
| Fkbp1a1 | 2.91E-34 | 0.748618 | 0.915 | 0.77 | 5.85E-30 | 3 L1 _ | Fkbp1a |
| Stc11 | 1.65E-34 | 0.670517 | 0.396 | 0.115 | 3.32E-30 | 3 L1 _ | Stc1 |
| Adarb11 | 1.56E-34 | 0.774066 | 0.431 | 0.146 | 3.13E-30 | 3 L1 _ | Adarb1 |
| Sigirr2 | 1.41E-34 | 0.421786 | 0.304 | 0.069 | 2.84E-30 | 3 L1 _ | Sigirr |
| Shroom41 | 1.25E-34 | 0.455113 | 0.223 | 0.038 | 2.51E-30 | 3 L1 _ | Shroom4 |
| Gas62 | 1.21E-34 | 0.818468 | 0.842 | 0.536 | 2.44E-30 | 3 L1 _ | Gas6 |
| 6430548M08Rik1 | 4.99E-35 | 0.502846 | 0.3 | 0.068 | 1.00E-30 | 3 L1 _ | 6430548M08Rik |
| Hoxd4 | 4.92E-35 | 0.32073 | 0.142 | 0.012 | 9.91E-31 | 3 L1 _ | Hoxd4 |
| Slc12a71 | 4.87E-35 | 0.46485 | 0.273 | 0.056 | 9.81E-31 | 3 L1 _ | Slc12a7 |
| Ggta12 | 2.94E-35 | 0.572302 | 0.369 | 0.098 | 5.93E-31 | 3 L1 _ | Ggta1 |
| Hip11 | 2.79E-35 | 0.807686 | 0.662 | 0.318 | 5.62E-31 | 3 L1 _ | Hip1 |
| Ivns1abp1 | 2.75E-35 | 1.095433 | 0.6 | 0.287 | 5.53E-31 | 3 L1 _ | Ivns1abp |
| App3 | 2.20E-35 | 0.72015 | 0.965 | 0.824 | 4.43E-31 | 3 L1 _ | App |
| Fabp52 | 2.18E-35 | 0.833811 | 0.562 | 0.215 | 4.39E-31 | 3 L1 _ | Fabp5 |
| Sncaip1 | 2.04E-35 | 0.346524 | 0.173 | 0.02 | 4.10E-31 | 3 L1 _ | Sncaip |
| Rnf1251 | 1.98E-35 | 0.422655 | 0.277 | 0.056 | 3.99E-31 | 3 L1 _ | Rnf125 |
| Caskin21 | 1.59E-35 | 0.565662 | 0.412 | 0.12 | 3.21E-31 | 3 L1 _ | Caskin2 |
| Jam33 | 1.55E-35 | 0.864614 | 0.708 | 0.368 | 3.12E-31 | 3 L1 _ | Jam3 |
| Ctnnb11 | 1.32E-35 | 0.835968 | 0.869 | 0.682 | 2.65E-31 | 3 L1 _ | Ctnnb1 |
| Stom2 | 1.12E-35 | 0.648009 | 0.438 | 0.135 | 2.25E-31 | 3 L1 _ | Stom |
| Tox31 | 8.33E-36 | 0.335576 | 0.142 | 0.012 | 1.68E-31 | 3 L1 _ | Tox3 |
| Thbd2 | 7.44E-36 | 0.790638 | 0.715 | 0.379 | 1.50E-31 | 3 L1 _ | Thbd |
| Eps8l2 | 7.15E-36 | 0.306525 | 0.138 | 0.011 | 1.44E-31 | 3 L1 _ | Eps8l2 |
| Grasp1 | 2.68E-36 | 0.744923 | 0.515 | 0.201 | 5.40E-32 | 3 L1 _ | Grasp |
| Tnfrsf10b1 | 1.52E-36 | 0.425259 | 0.219 | 0.033 | 3.07E-32 | 3 L1 _ | Tnfrsf10b |
| Tbxa2r1 | 1.44E-36 | 0.425379 | 0.25 | 0.044 | 2.90E-32 | 3 L1 _ | Tbxa2r |
| Efr3b1 | 1.24E-36 | 0.400606 | 0.231 | 0.037 | 2.49E-32 | 3 L1 _ | Efr3b |
| F11r1 | 1.13E-36 | 0.487343 | 0.412 | 0.115 | 2.27E-32 | 3 L1 _ | F11r |
| Hspa12b1 | 7.42E-37 | 0.572215 | 0.281 | 0.057 | 1.49E-32 | 3 L1 _ | Hspa12b |
| N4bp3 | 7.37E-37 | 0.462085 | 0.258 | 0.047 | 1.48E-32 | 3 L1 _ | N4bp3 |
| Ptprg1 | 7.31E-37 | 0.840816 | 0.727 | 0.385 | 1.47E-32 | 3 L1 _ | Ptprg |
| Arhgap271 | 5.68E-37 | 0.498325 | 0.338 | 0.081 | 1.14E-32 | 3 L1 _ | Arhgap27 |
| Plxnd11 | 5.37E-37 | 0.81936 | 0.685 | 0.346 | 1.08E-32 | 3 L1 _ | Plxnd1 |
| Mapk31 | 5.07E-37 | 0.798672 | 0.808 | 0.517 | 1.02E-32 | 3 L1 _ | Mapk3 |
| Abcb1a1 | 3.06E-37 | 0.606902 | 0.331 | 0.079 | 6.17E-33 | 3 L1 _ | Abcb1a |
| Inhbb | 2.19E-37 | 0.639222 | 0.273 | 0.053 | 4.41E-33 | 3 L1 _ | Inhbb |
| Lrrc8c2 | 1.74E-37 | 0.705138 | 0.573 | 0.209 | 3.50E-33 | 3 L1 _ | Lrrc8c |

| | | | | | | | |
|---|---|---|---|---|---|---|---|
| Hecw21 | 1.63E-37 | 0.726517 | 0.469 | 0.153 | 3.29E-33 | 3 L1 _ | Hecw2 |
| Prdm11 | 3.72E-38 | 0.536246 | 0.319 | 0.068 | 7.49E-34 | 3 L1 _ | Prdm1 |
| Tgm21 | 3.68E-38 | 0.905963 | 0.608 | 0.273 | 7.42E-34 | 3 L1 _ | Tgm2 |
| Tspan71 | 3.28E-38 | 0.587216 | 0.25 | 0.043 | 6.61E-34 | 3 L1 _ | Tspan7 |
| Hmbox11 | 2.49E-38 | 0.795539 | 0.585 | 0.236 | 5.01E-34 | 3 L1 _ | Hmbox1 |
| Stab1 | 1.35E-38 | 0.48793 | 0.296 | 0.059 | 2.72E-34 | 3 L1 _ | Stab1 |
| Dock91 | 1.09E-38 | 0.861163 | 0.527 | 0.193 | 2.19E-34 | 3 L1 _ | Dock9 |
| Mpzl1 | 9.34E-39 | 0.788754 | 0.623 | 0.275 | 1.88E-34 | 3 L1 _ | Mpzl1 |
| Nxn | 6.66E-39 | 0.827636 | 0.538 | 0.211 | 1.34E-34 | 3 L1 _ | Nxn |
| Sema3g1 | 4.78E-39 | 0.718394 | 0.196 | 0.024 | 9.62E-35 | 3 L1 _ | Sema3g |
| 9530082P21Rik1 | 3.33E-39 | 0.581737 | 0.346 | 0.081 | 6.71E-35 | 3 L1 _ | 9530082P21Rik |
| Gbp42 | 2.85E-39 | 0.79751 | 0.481 | 0.153 | 5.73E-35 | 3 L1 _ | Gbp4 |
| Syt151 | 1.65E-39 | 0.486257 | 0.212 | 0.027 | 3.33E-35 | 3 L1 _ | Syt15 |
| Ephb11 | 8.88E-40 | 0.327344 | 0.165 | 0.014 | 1.79E-35 | 3 L1 _ | Ephb1 |
| Endou1 | 5.42E-40 | 0.334029 | 0.204 | 0.025 | 1.09E-35 | 3 L1 _ | Endou |
| Slfn52 | 3.04E-40 | 0.960364 | 0.923 | 0.691 | 6.12E-36 | 3 L1 _ | Slfn5 |
| St8sia6 | 2.96E-40 | 0.322604 | 0.138 | 0.008 | 5.95E-36 | 3 L1 _ | St8sia6 |
| Pkp41 | 2.67E-40 | 0.627941 | 0.431 | 0.125 | 5.37E-36 | 3 L1 _ | Pkp4 |
| Klf131 | 2.18E-40 | 0.941877 | 0.831 | 0.519 | 4.40E-36 | 3 L1 _ | Klf13 |
| Heg12 | 1.95E-40 | 0.845776 | 0.681 | 0.297 | 3.93E-36 | 3 L1 _ | Heg1 |
| Rai141 | 1.18E-40 | 0.865247 | 0.681 | 0.318 | 2.37E-36 | 3 L1 _ | Rai14 |
| Pik3c2b | 9.98E-41 | 0.480446 | 0.246 | 0.038 | 2.01E-36 | 3 L1 _ | Pik3c2b |
| Mfng1 | 9.32E-41 | 0.579044 | 0.331 | 0.071 | 1.88E-36 | 3 L1 _ | Mfng |
| Ctnnbip11 | 7.64E-41 | 0.869761 | 0.588 | 0.244 | 1.54E-36 | 3 L1 _ | Ctnnbip1 |
| Slc43a32 | 5.26E-41 | 1.07024 | 0.8 | 0.473 | 1.06E-36 | 3 L1 _ | Slc43a3 |
| Tnfaip12 | 3.35E-41 | 0.954152 | 0.692 | 0.343 | 6.75E-37 | 3 L1 _ | Tnfaip1 |
| Slc6a61 | 2.84E-41 | 1.235762 | 0.673 | 0.327 | 5.73E-37 | 3 L1 _ | Slc6a6 |
| Ptgs11 | 2.20E-41 | 0.88614 | 0.588 | 0.238 | 4.44E-37 | 3 L1 _ | Ptgs1 |
| Rtp31 | 1.57E-41 | 0.507898 | 0.223 | 0.029 | 3.16E-37 | 3 L1 _ | Rtp3 |
| Shank31 | 1.24E-41 | 0.413359 | 0.25 | 0.038 | 2.50E-37 | 3 L1 _ | Shank3 |
| Ephb41 | 1.12E-41 | 0.67657 | 0.435 | 0.125 | 2.26E-37 | 3 L1 _ | Ephb4 |
| Ltbp42 | 4.82E-42 | 0.946622 | 0.9 | 0.644 | 9.70E-38 | 3 L1 _ | Ltbp4 |
| Impdh11 | 3.92E-42 | 0.708553 | 0.458 | 0.138 | 7.89E-38 | 3 L1 _ | Impdh1 |
| Card101 | 3.17E-42 | 0.593523 | 0.392 | 0.099 | 6.38E-38 | 3 L1 _ | Card10 |
| Ppp1r13b1 | 2.90E-42 | 0.65856 | 0.392 | 0.099 | 5.85E-38 | 3 L1 _ | Ppp1r13b |
| Galnt18 | 1.93E-42 | 0.503734 | 0.281 | 0.047 | 3.90E-38 | 3 L1 _ | Galnt18 |
| Rtl8a1 | 1.07E-42 | 1.043428 | 0.781 | 0.452 | 2.16E-38 | 3 L1 _ | Rtl8a |
| Cbfa2t32 | 9.34E-43 | 0.738722 | 0.604 | 0.207 | 1.88E-38 | 3 L1 _ | Cbfa2t3 |
| Acvrl11 | 7.41E-43 | 0.880887 | 0.723 | 0.368 | 1.49E-38 | 3 L1 _ | Acvrl1 |
| Msn2 | 7.07E-43 | 0.989224 | 0.935 | 0.78 | 1.42E-38 | 3 L1 _ | Msn |
| Spns21 | 5.54E-43 | 0.413893 | 0.242 | 0.034 | 1.12E-38 | 3 L1 _ | Spns2 |

| | | | | | | | |
|---|---|---|---|---|---|---|---|
| Stmn2 | 5.51E-43 | 0.982882 | 0.212 | 0.025 | 1.11E-38 | 3 L1 _ | Stmn2 |
| Itga13 | 3.52E-43 | 0.905432 | 0.962 | 0.78 | 7.10E-39 | 3 L1 _ | Itga1 |
| Filip12 | 3.43E-43 | 0.881675 | 0.5 | 0.157 | 6.91E-39 | 3 L1 _ | Filip1 |
| Kitl3 | 3.16E-43 | 1.123323 | 0.75 | 0.367 | 6.37E-39 | 3 L1 _ | Kitl |
| Ifi441 | 2.35E-43 | 1.062504 | 0.473 | 0.143 | 4.74E-39 | 3 L1 _ | Ifi44 |
| Gngt21 | 2.20E-43 | 0.812125 | 0.6 | 0.22 | 4.44E-39 | 3 L1 _ | Gngt2 |
| St6galnac31 | 1.80E-43 | 0.461208 | 0.242 | 0.033 | 3.63E-39 | 3 L1 _ | St6galnac3 |
| Proser21 | 1.57E-43 | 0.49973 | 0.258 | 0.038 | 3.16E-39 | 3 L1 _ | Proser2 |
| Itpkb1 | 1.29E-43 | 1.01999 | 0.723 | 0.364 | 2.60E-39 | 3 L1 _ | Itpkb |
| Ackr32 | 1.18E-43 | 0.620891 | 0.396 | 0.092 | 2.38E-39 | 3 L1 _ | Ackr3 |
| B3gnt21 | 1.15E-43 | 0.969529 | 0.704 | 0.34 | 2.32E-39 | 3 L1 _ | B3gnt2 |
| Notch11 | 1.13E-43 | 0.892647 | 0.692 | 0.306 | 2.28E-39 | 3 L1 _ | Notch1 |
| Lpar61 | 6.78E-44 | 0.952892 | 0.5 | 0.158 | 1.37E-39 | 3 L1 _ | Lpar6 |
| Pitpnm21 | 3.04E-44 | 0.624807 | 0.415 | 0.106 | 6.12E-40 | 3 L1 _ | Pitpnm2 |
| Rassf31 | 2.96E-44 | 0.635856 | 0.404 | 0.097 | 5.95E-40 | 3 L1 _ | Rassf3 |
| Ptprm1 | 1.79E-44 | 0.707527 | 0.462 | 0.132 | 3.60E-40 | 3 L1 _ | Ptprm |
| Prex21 | 9.38E-45 | 0.917459 | 0.758 | 0.377 | 1.89E-40 | 3 L1 _ | Prex2 |
| Insr | 6.91E-45 | 1.097194 | 0.531 | 0.182 | 1.39E-40 | 3 L1 _ | Insr |
| Pkn31 | 5.15E-45 | 0.446945 | 0.227 | 0.027 | 1.04E-40 | 3 L1 _ | Pkn3 |
| Abcg21 | 3.39E-45 | 1.033868 | 0.681 | 0.308 | 6.82E-41 | 3 L1 _ | Abcg2 |
| Mmp151 | 2.85E-45 | 0.46968 | 0.319 | 0.057 | 5.74E-41 | 3 L1 _ | Mmp15 |
| Cldn5 | 1.71E-45 | 0.994721 | 0.212 | 0.023 | 3.45E-41 | 3 L1 _ | Cldn5 |
| Sh3bp51 | 2.75E-46 | 0.943042 | 0.577 | 0.218 | 5.54E-42 | 3 L1 _ | Sh3bp5 |
| Hspb13 | 8.29E-47 | 1.123523 | 0.888 | 0.568 | 1.67E-42 | 3 L1 _ | Hspb1 |
| Car82 | 2.87E-47 | 0.54415 | 0.281 | 0.041 | 5.77E-43 | 3 L1 _ | Car8 |
| Ablim31 | 2.29E-47 | 0.392955 | 0.181 | 0.013 | 4.62E-43 | 3 L1 _ | Ablim3 |
| Apbb21 | 2.15E-47 | 0.999127 | 0.792 | 0.434 | 4.33E-43 | 3 L1 _ | Apbb2 |
| Bvht1 | 1.71E-47 | 0.717288 | 0.477 | 0.134 | 3.44E-43 | 3 L1 _ | Bvht |
| Crim12 | 1.29E-47 | 0.852373 | 0.673 | 0.25 | 2.60E-43 | 3 L1 _ | Crim1 |
| Ets11 | 1.01E-47 | 1.095669 | 0.869 | 0.577 | 2.03E-43 | 3 L1 _ | Ets1 |
| Gimap12 | 6.36E-48 | 0.688692 | 0.677 | 0.223 | 1.28E-43 | 3 L1 _ | Gimap1 |
| Lmo21 | 3.47E-48 | 0.82447 | 0.462 | 0.124 | 6.99E-44 | 3 L1 _ | Lmo2 |
| Myo18a1 | 3.29E-48 | 0.861816 | 0.608 | 0.226 | 6.63E-44 | 3 L1 _ | Myo18a |
| Id32 | 2.43E-48 | 1.036455 | 0.985 | 0.733 | 4.89E-44 | 3 L1 _ | Id3 |
| Kcnq11 | 2.19E-48 | 0.40278 | 0.212 | 0.02 | 4.41E-44 | 3 L1 _ | Kcnq1 |
| Tshz22 | 1.80E-48 | 1.11328 | 0.888 | 0.611 | 3.62E-44 | 3 L1 _ | Tshz2 |
| Dipk2b | 6.80E-49 | 0.30553 | 0.142 | 0.005 | 1.37E-44 | 3 L1 _ | Dipk2b |
| Meox11 | 6.38E-49 | 0.97644 | 0.577 | 0.2 | 1.28E-44 | 3 L1 _ | Meox1 |
| Ssu21 | 4.47E-49 | 0.503227 | 0.223 | 0.023 | 9.00E-45 | 3 L1 _ | Ssu2 |
| Sox71 | 3.92E-49 | 0.452346 | 0.231 | 0.025 | 7.90E-45 | 3 L1 _ | Sox7 |
| Scarf11 | 3.51E-49 | 0.477802 | 0.265 | 0.034 | 7.06E-45 | 3 L1 _ | Scarf1 |

| | | | | | | | |
|---|---|---|---|---|---|---|---|
| Igfbp72 | 3.31E-49 | 0.874353 | 1 | 0.937 | 6.66E-45 | 3 L1 _ | Igfbp7 |
| Ahr1 | 2.07E-49 | 0.879252 | 0.535 | 0.161 | 4.17E-45 | 3 L1 _ | Ahr |
| Nos31 | 1.76E-49 | 0.4844 | 0.262 | 0.034 | 3.54E-45 | 3 L1 _ | Nos3 |
| 4931406P16Rik2 | 6.44E-50 | 1.025341 | 0.796 | 0.404 | 1.30E-45 | 3 L1 _ | 4931406P16Rik |
| Tm4sf13 | 5.49E-50 | 1.301005 | 0.938 | 0.666 | 1.11E-45 | 3 L1 _ | Tm4sf1 |
| Atox11 | 2.86E-50 | 1.097419 | 0.942 | 0.764 | 5.76E-46 | 3 L1 _ | Atox1 |
| Ndrg11 | 1.19E-50 | 1.049479 | 0.662 | 0.275 | 2.39E-46 | 3 L1 _ | Ndrg1 |
| Clu1 | 9.46E-51 | 1.491896 | 0.415 | 0.099 | 1.91E-46 | 3 L1 _ | Clu |
| Adam151 | 7.21E-51 | 1.086909 | 0.7 | 0.316 | 1.45E-46 | 3 L1 _ | Adam15 |
| Gimap62 | 4.84E-51 | 1.071903 | 0.792 | 0.358 | 9.74E-47 | 3 L1 _ | Gimap6 |
| Hoxd3os11 | 4.67E-51 | 0.406744 | 0.246 | 0.027 | 9.40E-47 | 3 L1 _ | Hoxd3os1 |
| Chst21 | 4.56E-51 | 0.919648 | 0.488 | 0.136 | 9.18E-47 | 3 L1 _ | Chst2 |
| Pakap.11 | 4.42E-51 | 1.098788 | 0.804 | 0.395 | 8.91E-47 | 3 L1 _ | Pakap.1 |
| Ednrb2 | 4.05E-51 | 0.498754 | 0.777 | 0.259 | 8.15E-47 | 3 L1 _ | Ednrb |
| Fzd6 | 2.70E-51 | 0.414426 | 0.2 | 0.016 | 5.44E-47 | 3 L1 _ | Fzd6 |
| Cd363 | 2.56E-51 | 1.061033 | 0.558 | 0.16 | 5.16E-47 | 3 L1 _ | Cd36 |
| Zfp9791 | 2.51E-51 | 0.52194 | 0.288 | 0.04 | 5.06E-47 | 3 L1 _ | Zfp979 |
| Eng1 | 1.60E-51 | 1.062793 | 0.942 | 0.641 | 3.22E-47 | 3 L1 _ | Eng |
| Prkch2 | 3.57E-52 | 0.926787 | 0.515 | 0.141 | 7.19E-48 | 3 L1 _ | Prkch |
| Fbxl71 | 7.09E-53 | 1.03916 | 0.585 | 0.195 | 1.43E-48 | 3 L1 _ | Fbxl7 |
| Nxpe41 | 5.75E-53 | 0.481837 | 0.277 | 0.035 | 1.16E-48 | 3 L1 _ | Nxpe4 |
| Notch41 | 3.16E-53 | 0.755735 | 0.465 | 0.113 | 6.36E-49 | 3 L1 _ | Notch4 |
| Pik3r31 | 2.11E-53 | 0.647957 | 0.342 | 0.057 | 4.26E-49 | 3 L1 _ | Pik3r3 |
| Lipe1 | 1.52E-53 | 0.692229 | 0.396 | 0.08 | 3.05E-49 | 3 L1 _ | Lipe |
| Arhgap291 | 1.20E-53 | 1.269611 | 0.885 | 0.59 | 2.42E-49 | 3 L1 _ | Arhgap29 |
| Bcl6b | 6.54E-54 | 0.441958 | 0.212 | 0.017 | 1.32E-49 | 3 L1 _ | Bcl6b |
| Nav21 | 4.85E-54 | 0.502539 | 0.327 | 0.05 | 9.76E-50 | 3 L1 _ | Nav2 |
| Npr12 | 2.15E-54 | 0.717449 | 0.415 | 0.088 | 4.34E-50 | 3 L1 _ | Npr1 |
| Rhbdl21 | 8.24E-55 | 0.430608 | 0.2 | 0.013 | 1.66E-50 | 3 L1 _ | Rhbdl2 |
| Adcy41 | 5.73E-55 | 0.746083 | 0.488 | 0.12 | 1.15E-50 | 3 L1 _ | Adcy4 |
| Alox121 | 4.68E-55 | 0.752081 | 0.292 | 0.038 | 9.43E-51 | 3 L1 _ | Alox12 |
| Parvb1 | 2.35E-55 | 0.553791 | 0.327 | 0.048 | 4.73E-51 | 3 L1 _ | Parvb |
| Irx31 | 5.23E-56 | 0.593882 | 0.292 | 0.037 | 1.05E-51 | 3 L1 _ | Irx3 |
| Afdn1 | 1.85E-56 | 1.104627 | 0.758 | 0.366 | 3.72E-52 | 3 L1 _ | Afdn |
| Dok41 | 1.81E-56 | 0.68546 | 0.412 | 0.081 | 3.64E-52 | 3 L1 _ | Dok4 |
| Gpr41 | 1.25E-56 | 0.479747 | 0.254 | 0.025 | 2.52E-52 | 3 L1 _ | Gpr4 |
| Crip23 | 1.12E-56 | 1.063426 | 0.969 | 0.747 | 2.26E-52 | 3 L1 _ | Crip2 |
| Tspan122 | 8.09E-57 | 0.73807 | 0.5 | 0.115 | 1.63E-52 | 3 L1 _ | Tspan12 |
| Wscd1 | 1.37E-57 | 0.543627 | 0.219 | 0.016 | 2.76E-53 | 3 L1 _ | Wscd1 |
| Stap21 | 8.91E-58 | 0.468854 | 0.246 | 0.023 | 1.79E-53 | 3 L1 _ | Stap2 |
| Fam43a1 | 7.13E-58 | 1.001776 | 0.515 | 0.137 | 1.44E-53 | 3 L1 _ | Fam43a |

| | | | | | | | |
|---|---|---|---|---|---|---|---|
| Rdx1 | 1.39E-58 | 1.103743 | 0.946 | 0.785 | 2.80E-54 | 3 L1 _ | Rdx |
| Jup1 | 7.33E-59 | 1.043484 | 0.769 | 0.348 | 1.48E-54 | 3 L1 _ | Jup |
| Nxpe21 | 2.00E-59 | 0.442013 | 0.231 | 0.018 | 4.03E-55 | 3 L1 _ | Nxpe2 |
| Elk31 | 1.90E-59 | 1.12947 | 0.842 | 0.457 | 3.82E-55 | 3 L1 _ | Elk3 |
| Snrk3 | 1.41E-59 | 1.252229 | 0.923 | 0.57 | 2.84E-55 | 3 L1 _ | Snrk |
| Etl41 | 1.12E-59 | 0.981585 | 0.531 | 0.139 | 2.25E-55 | 3 L1 _ | Etl4 |
| Fbln22 | 6.42E-60 | 1.270676 | 0.896 | 0.502 | 1.29E-55 | 3 L1 _ | Fbln2 |
| Tmem255b1 | 5.76E-60 | 0.620846 | 0.308 | 0.039 | 1.16E-55 | 3 L1 _ | Tmem255b |
| Hlx1 | 5.48E-60 | 0.798491 | 0.373 | 0.062 | 1.10E-55 | 3 L1 _ | Hlx |
| Ica11 | 4.53E-60 | 0.558773 | 0.285 | 0.031 | 9.12E-56 | 3 L1 _ | Ica1 |
| Gpr1461 | 3.67E-60 | 0.709334 | 0.404 | 0.074 | 7.39E-56 | 3 L1 _ | Gpr146 |
| Ldb21 | 2.17E-60 | 1.155603 | 0.762 | 0.336 | 4.38E-56 | 3 L1 _ | Ldb2 |
| Uaca1 | 2.02E-60 | 1.155316 | 0.65 | 0.231 | 4.07E-56 | 3 L1 _ | Uaca |
| Lama31 | 1.01E-60 | 0.62258 | 0.335 | 0.046 | 2.03E-56 | 3 L1 _ | Lama3 |
| Ptprr1 | 4.40E-61 | 0.673679 | 0.246 | 0.02 | 8.86E-57 | 3 L1 _ | Ptprr |
| Rapgef31 | 3.95E-61 | 0.828532 | 0.485 | 0.111 | 7.95E-57 | 3 L1 _ | Rapgef3 |
| Pde2a | 1.28E-61 | 1.089286 | 0.515 | 0.134 | 2.59E-57 | 3 L1 _ | Pde2a |
| Epas13 | 8.89E-62 | 1.144162 | 0.919 | 0.482 | 1.79E-57 | 3 L1 _ | Epas1 |
| Tspan181 | 2.67E-62 | 0.761369 | 0.408 | 0.074 | 5.37E-58 | 3 L1 _ | Tspan18 |
| Cd381 | 2.18E-62 | 1.038687 | 0.612 | 0.186 | 4.39E-58 | 3 L1 _ | Cd38 |
| B4galt41 | 7.65E-63 | 0.782706 | 0.504 | 0.116 | 1.54E-58 | 3 L1 _ | B4galt4 |
| Sptbn11 | 2.82E-63 | 1.116647 | 0.992 | 0.89 | 5.67E-59 | 3 L1 _ | Sptbn1 |
| Kctd12b1 | 2.55E-63 | 1.216956 | 0.723 | 0.279 | 5.14E-59 | 3 L1 _ | Kctd12b |
| Plpp32 | 6.71E-64 | 1.35453 | 0.958 | 0.703 | 1.35E-59 | 3 L1 _ | Plpp3 |
| Lrg1 | 4.03E-64 | 0.6265 | 0.262 | 0.022 | 8.12E-60 | 3 L1 _ | Lrg1 |
| Adgrl21 | 2.33E-64 | 1.140499 | 0.688 | 0.234 | 4.69E-60 | 3 L1 _ | Adgrl2 |
| Ces2e1 | 1.53E-64 | 0.680938 | 0.377 | 0.054 | 3.08E-60 | 3 L1 _ | Ces2e |
| Ankrd33b1 | 1.04E-65 | 0.56607 | 0.269 | 0.024 | 2.09E-61 | 3 L1 _ | Ankrd33b |
| Vamp51 | 5.19E-66 | 1.197671 | 0.842 | 0.427 | 1.04E-61 | 3 L1 _ | Vamp5 |
| Plxna21 | 3.38E-66 | 0.839455 | 0.508 | 0.109 | 6.81E-62 | 3 L1 _ | Plxna2 |
| Ccdc85b | 1.79E-66 | 1.31702 | 0.835 | 0.458 | 3.61E-62 | 3 L1 _ | Ccdc85b |
| Plpp11 | 1.25E-66 | 1.501949 | 0.908 | 0.563 | 2.52E-62 | 3 L1 _ | Plpp1 |
| BC0285281 | 4.93E-67 | 1.042336 | 0.585 | 0.156 | 9.93E-63 | 3 L1 _ | BC028528 |
| Rassf91 | 4.36E-67 | 0.946224 | 0.438 | 0.079 | 8.78E-63 | 3 L1 _ | Rassf9 |
| Cdh131 | 1.50E-67 | 0.645495 | 0.338 | 0.041 | 3.02E-63 | 3 L1 _ | Cdh13 |
| Sema7a1 | 9.80E-68 | 1.110086 | 0.642 | 0.189 | 1.97E-63 | 3 L1 _ | Sema7a |
| Depp1 | 4.82E-68 | 0.60643 | 0.2 | 0.008 | 9.70E-64 | 3 L1 _ | Depp1 |
| Tnfsf102 | 1.06E-68 | 1.06503 | 0.654 | 0.177 | 2.14E-64 | 3 L1 _ | Tnfsf10 |
| Klf42 | 9.19E-69 | 1.558457 | 0.742 | 0.263 | 1.85E-64 | 3 L1 _ | Klf4 |
| Mef2c3 | 9.45E-70 | 1.388562 | 0.95 | 0.615 | 1.90E-65 | 3 L1 _ | Mef2c |
| Aplp21 | 4.10E-70 | 1.406379 | 0.954 | 0.673 | 8.26E-66 | 3 L1 _ | Aplp2 |

| | | | | | | | |
|---|---|---|---|---|---|---|---|
| Smagp2 | 3.78E-70 | 0.87656 | 0.523 | 0.106 | 7.61E-66 | 3 L1 _ | Smagp |
| Ccdc85a1 | 3.49E-70 | 0.643047 | 0.335 | 0.038 | 7.04E-66 | 3 L1 _ | Ccdc85a |
| Anxa33 | 2.99E-70 | 1.070288 | 0.773 | 0.259 | 6.02E-66 | 3 L1 _ | Anxa3 |
| Mrtfb | 1.32E-71 | 1.011474 | 0.562 | 0.137 | 2.65E-67 | 3 L1 _ | Mrtfb |
| Trp53i11 | 8.10E-72 | 1.369964 | 0.623 | 0.175 | 1.63E-67 | 3 L1 _ | Trp53i11 |
| Cav22 | 5.60E-72 | 0.995511 | 0.627 | 0.156 | 1.13E-67 | 3 L1 _ | Cav2 |
| Ly6e1 | 3.41E-72 | 1.19529 | 0.992 | 0.865 | 6.87E-68 | 3 L1 _ | Ly6e |
| Mecom1 | 1.54E-72 | 1.024528 | 0.362 | 0.045 | 3.10E-68 | 3 L1 _ | Mecom |
| Tspan132 | 1.37E-72 | 1.098766 | 0.588 | 0.143 | 2.76E-68 | 3 L1 _ | Tspan13 |
| Plscr21 | 5.88E-73 | 1.327931 | 0.808 | 0.33 | 1.18E-68 | 3 L1 _ | Plscr2 |
| Ccm2l1 | 5.88E-73 | 0.758549 | 0.358 | 0.042 | 1.18E-68 | 3 L1 _ | Ccm2l |
| Kif26a1 | 4.48E-73 | 0.590577 | 0.308 | 0.028 | 9.01E-69 | 3 L1 _ | Kif26a |
| Chst151 | 1.87E-73 | 0.928704 | 0.512 | 0.1 | 3.77E-69 | 3 L1 _ | Chst15 |
| Gnai21 | 1.54E-73 | 1.041292 | 0.973 | 0.919 | 3.09E-69 | 3 L1 _ | Gnai2 |
| Mall1 | 9.04E-74 | 0.617859 | 0.296 | 0.025 | 1.82E-69 | 3 L1 _ | Mall |
| Ripply31 | 3.35E-74 | 0.783874 | 0.458 | 0.077 | 6.74E-70 | 3 L1 _ | Ripply3 |
| Calm12 | 2.81E-74 | 1.094692 | 0.992 | 0.947 | 5.66E-70 | 3 L1 _ | Calm1 |
| Rasgrf21 | 6.98E-75 | 0.593406 | 0.323 | 0.031 | 1.41E-70 | 3 L1 _ | Rasgrf2 |
| Lmcd11 | 5.58E-75 | 1.208931 | 0.577 | 0.138 | 1.12E-70 | 3 L1 _ | Lmcd1 |
| Syne11 | 2.53E-75 | 1.448675 | 0.762 | 0.289 | 5.09E-71 | 3 L1 _ | Syne1 |
| Klf21 | 5.14E-76 | 1.811151 | 0.877 | 0.416 | 1.03E-71 | 3 L1 _ | Klf2 |
| Pcp4l13 | 3.09E-76 | 1.121929 | 0.688 | 0.169 | 6.23E-72 | 3 L1 _ | Pcp4l1 |
| Bcam3 | 2.44E-76 | 1.266926 | 0.815 | 0.271 | 4.91E-72 | 3 L1 _ | Bcam |
| C430049B03Rik | 2.35E-76 | 0.468828 | 0.262 | 0.015 | 4.72E-72 | 3 L1 _ | C430049B03Rik |
| Srgn3 | 1.54E-76 | 1.348922 | 0.915 | 0.359 | 3.09E-72 | 3 L1 _ | Srgn |
| Nfib2 | 6.83E-77 | 1.303151 | 0.981 | 0.734 | 1.38E-72 | 3 L1 _ | Nfib |
| Cav13 | 1.66E-77 | 1.247946 | 0.804 | 0.25 | 3.35E-73 | 3 L1 _ | Cav1 |
| Pdgfb1 | 1.51E-77 | 1.130293 | 0.658 | 0.164 | 3.04E-73 | 3 L1 _ | Pdgfb |
| Limch11 | 1.12E-79 | 0.86305 | 0.392 | 0.048 | 2.26E-75 | 3 L1 _ | Limch1 |
| Unc45b1 | 4.88E-80 | 0.822768 | 0.469 | 0.074 | 9.83E-76 | 3 L1 _ | Unc45b |
| Mfsd61 | 4.79E-80 | 0.841028 | 0.465 | 0.073 | 9.65E-76 | 3 L1 _ | Mfsd6 |
| Sema3f1 | 3.62E-80 | 0.848527 | 0.431 | 0.06 | 7.30E-76 | 3 L1 _ | Sema3f |
| Jag21 | 1.94E-80 | 0.827085 | 0.388 | 0.046 | 3.91E-76 | 3 L1 _ | Jag2 |
| Nostrin1 | 1.83E-80 | 1.173584 | 0.623 | 0.139 | 3.69E-76 | 3 L1 _ | Nostrin |
| Flt41 | 2.01E-81 | 0.745915 | 0.381 | 0.042 | 4.06E-77 | 3 L1 _ | Flt4 |
| Zfp3661 | 5.04E-82 | 0.675119 | 0.35 | 0.033 | 1.01E-77 | 3 L1 _ | Zfp366 |
| Rsad21 | 4.61E-82 | 1.391144 | 0.677 | 0.166 | 9.29E-78 | 3 L1 _ | Rsad2 |
| Pltp1 | 1.85E-82 | 1.406833 | 0.792 | 0.268 | 3.72E-78 | 3 L1 _ | Pltp |
| Sema6a1 | 1.74E-82 | 0.946673 | 0.569 | 0.117 | 3.50E-78 | 3 L1 _ | Sema6a |
| Ece11 | 1.88E-83 | 1.39463 | 0.819 | 0.312 | 3.79E-79 | 3 L1 _ | Ece1 |
| Afap1l11 | 2.45E-84 | 0.672995 | 0.404 | 0.046 | 4.94E-80 | 3 L1 _ | Afap1l1 |

| | | | | | | | |
|---|---|---|---|---|---|---|---|
| Fli11 | 1.39E-84 | 1.236053 | 0.692 | 0.187 | 2.79E-80 | 3 L1 _ | Fli1 |
| Dysf1 | 1.09E-85 | 0.942097 | 0.535 | 0.091 | 2.19E-81 | 3 L1 _ | Dysf |
| Ehd41 | 4.97E-86 | 1.427332 | 0.823 | 0.294 | 1.00E-81 | 3 L1 _ | Ehd4 |
| Aplnr1 | 3.85E-86 | 1.090759 | 0.373 | 0.037 | 7.76E-82 | 3 L1 _ | Aplnr |
| Nova21 | 1.74E-86 | 0.680637 | 0.369 | 0.035 | 3.51E-82 | 3 L1 _ | Nova2 |
| Lims21 | 4.15E-87 | 1.122498 | 0.565 | 0.105 | 8.35E-83 | 3 L1 _ | Lims2 |
| Tcim1 | 4.61E-88 | 1.545132 | 0.762 | 0.224 | 9.29E-84 | 3 L1 _ | Tcim |
| Gm120021 | 1.41E-88 | 0.799027 | 0.362 | 0.031 | 2.83E-84 | 3 L1 _ | Gm12002 |
| Ly6a1 | 8.02E-89 | 1.856265 | 0.996 | 0.7 | 1.62E-84 | 3 L1 _ | Ly6a |
| Samd121 | 2.23E-91 | 0.719643 | 0.377 | 0.034 | 4.50E-87 | 3 L1 _ | Samd12 |
| Clec14a1 | 2.44E-92 | 1.639557 | 0.865 | 0.324 | 4.92E-88 | 3 L1 _ | Clec14a |
| Plekho11 | 2.18E-93 | 1.13055 | 0.608 | 0.116 | 4.39E-89 | 3 L1 _ | Plekho1 |
| Mcf2l1 | 1.27E-93 | 1.085698 | 0.531 | 0.08 | 2.55E-89 | 3 L1 _ | Mcf2l |
| Id11 | 9.14E-94 | 1.533782 | 0.792 | 0.224 | 1.84E-89 | 3 L1 _ | Id1 |
| Gata21 | 1.88E-94 | 0.73672 | 0.388 | 0.034 | 3.80E-90 | 3 L1 _ | Gata2 |
| Ctla2a2 | 1.20E-94 | 1.910712 | 0.869 | 0.293 | 2.42E-90 | 3 L1 _ | Ctla2a |
| Car41 | 1.53E-95 | 1.066499 | 0.442 | 0.048 | 3.08E-91 | 3 L1 _ | Car4 |
| Exoc3l21 | 5.96E-96 | 1.005729 | 0.508 | 0.068 | 1.20E-91 | 3 L1 _ | Exoc3l2 |
| Smco41 | 3.34E-96 | 1.272001 | 0.758 | 0.204 | 6.73E-92 | 3 L1 _ | Smco4 |
| Cracr2b1 | 1.29E-96 | 1.068712 | 0.542 | 0.087 | 2.60E-92 | 3 L1 _ | Cracr2b |
| Tcf151 | 1.43E-97 | 1.189558 | 0.527 | 0.074 | 2.87E-93 | 3 L1 _ | Tcf15 |
| Dll4 | 8.97E-99 | 0.764431 | 0.331 | 0.019 | 1.81E-94 | 3 L1 _ | Dll4 |
| Ets21 | 2.44E-99 | 1.519684 | 0.738 | 0.201 | 4.91E-95 | 3 L1 _ | Ets2 |
| Erg1 | 2.38E-99 | 1.064415 | 0.508 | 0.072 | 4.80E-95 | 3 L1 _ | Erg |
| Cd343 | 7.71E-100 | 1.725247 | 0.938 | 0.388 | 1.55E-95 | 3 L1 _ | Cd34 |
| Adgrg11 | 5.46E-101 | 0.838689 | 0.512 | 0.064 | 1.10E-96 | 3 L1 _ | Adgrg1 |
| Plk21 | 2.23E-101 | 1.230307 | 0.558 | 0.087 | 4.50E-97 | 3 L1 _ | Plk2 |
| Efnb21 | 1.57E-101 | 1.181683 | 0.55 | 0.083 | 3.16E-97 | 3 L1 _ | Efnb2 |
| Slc9a3r21 | 6.15E-103 | 1.847641 | 0.931 | 0.415 | 1.24E-98 | 3 L1 _ | Slc9a3r2 |
| Clec1a1 | 5.05E-103 | 1.280176 | 0.631 | 0.117 | 1.02E-98 | 3 L1 _ | Clec1a |
| Fgd51 | 2.85E-103 | 1.178187 | 0.658 | 0.132 | 5.74E-99 | 3 L1 _ | Fgd5 |
| Arap31 | 1.62E-103 | 1.148422 | 0.658 | 0.129 | 3.26E-99 | 3 L1 _ | Arap3 |
| Ramp22 | 1.10E-104 | 1.859265 | 0.95 | 0.442 | 2.23E-100 | 3 L1 _ | Ramp2 |
| Jam21 | 3.80E-107 | 1.784196 | 0.838 | 0.275 | 7.65E-103 | 3 L1 _ | Jam2 |
| She1 | 1.64E-107 | 1.046552 | 0.546 | 0.074 | 3.30E-103 | 3 L1 _ | She |
| Rapgef41 | 9.26E-108 | 0.922837 | 0.488 | 0.054 | 1.86E-103 | 3 L1 _ | Rapgef4 |
| Robo41 | 3.67E-110 | 1.019484 | 0.492 | 0.053 | 7.39E-106 | 3 L1 _ | Robo4 |
| Esam3 | 7.49E-111 | 1.698911 | 0.908 | 0.265 | 1.51E-106 | 3 L1 _ | Esam |
| Upp11 | 4.06E-111 | 0.912644 | 0.427 | 0.034 | 8.18E-107 | 3 L1 _ | Upp1 |
| Cd24a1 | 1.89E-112 | 1.085693 | 0.604 | 0.085 | 3.80E-108 | 3 L1 _ | Cd24a |
| Sox181 | 4.45E-114 | 0.978804 | 0.462 | 0.042 | 8.96E-110 | 3 L1 _ | Sox18 |

| | | | | | | | |
|---|---|---|---|---|---|---|---|
| Dock41 | 7.81E-115 | 1.054494 | 0.558 | 0.073 | 1.57E-110 | 3 L1 _ | Dock4 |
| Sorbs22 | 6.66E-116 | 1.417044 | 0.712 | 0.129 | 1.34E-111 | 3 L1 _ | Sorbs2 |
| Entpd11 | 4.36E-116 | 1.372157 | 0.715 | 0.141 | 8.78E-112 | 3 L1 _ | Entpd1 |
| Sox171 | 2.87E-116 | 1.359258 | 0.485 | 0.046 | 5.79E-112 | 3 L1 _ | Sox17 |
| Col13a11 | 2.31E-116 | 1.495998 | 0.677 | 0.118 | 4.66E-112 | 3 L1 _ | Col13a1 |
| Arhgef151 | 2.27E-117 | 1.034834 | 0.573 | 0.071 | 4.56E-113 | 3 L1 _ | Arhgef15 |
| Arhgap311 | 2.17E-122 | 1.883795 | 0.9 | 0.295 | 4.37E-118 | 3 L1 _ | Arhgap31 |
| Fabp41 | 1.56E-122 | 2.227095 | 0.973 | 0.39 | 3.15E-118 | 3 L1 _ | Fabp4 |
| Plvap1 | 6.06E-123 | 2.646672 | 0.973 | 0.471 | 1.22E-118 | 3 L1 _ | Plvap |
| Rflnb1 | 9.29E-126 | 1.97734 | 0.885 | 0.252 | 1.87E-121 | 3 L1 _ | Rflnb |
| Rgcc1 | 3.07E-127 | 1.746869 | 0.842 | 0.197 | 6.18E-123 | 3 L1 _ | Rgcc |
| Kank31 | 7.81E-129 | 1.225005 | 0.665 | 0.098 | 1.57E-124 | 3 L1 _ | Kank3 |
| Ecscr1 | 1.61E-131 | 1.650773 | 0.785 | 0.166 | 3.23E-127 | 3 L1 _ | Ecscr |
| C130074G19Rik1 | 2.36E-132 | 1.141514 | 0.55 | 0.053 | 4.74E-128 | 3 L1 _ | C130074G19Rik |
| Cavin21 | 5.91E-133 | 1.904657 | 0.9 | 0.214 | 1.19E-128 | 3 L1 _ | Cavin2 |
| S1pr11 | 3.26E-133 | 1.582394 | 0.738 | 0.129 | 6.57E-129 | 3 L1 _ | S1pr1 |
| Plcb11 | 3.26E-134 | 1.436807 | 0.673 | 0.092 | 6.57E-130 | 3 L1 _ | Plcb1 |
| Ceacam11 | 6.24E-135 | 1.162365 | 0.554 | 0.052 | 1.26E-130 | 3 L1 _ | Ceacam1 |
| Ramp31 | 1.18E-138 | 1.699742 | 0.669 | 0.089 | 2.37E-134 | 3 L1 _ | Ramp3 |
| Rbp71 | 1.19E-141 | 2.220824 | 0.8 | 0.149 | 2.40E-137 | 3 L1 _ | Rbp7 |
| Cmtm81 | 1.59E-142 | 1.285629 | 0.619 | 0.065 | 3.20E-138 | 3 L1 _ | Cmtm8 |
| Palmd1 | 2.27E-147 | 1.263781 | 0.6 | 0.057 | 4.57E-143 | 3 L1 _ | Palmd |
| Fam167b1 | 2.05E-153 | 1.469743 | 0.65 | 0.067 | 4.13E-149 | 3 L1 _ | Fam167b |
| Efna11 | 4.20E-154 | 1.469633 | 0.635 | 0.067 | 8.46E-150 | 3 L1 _ | Efna1 |
| Ly6c11 | 3.42E-154 | 2.937014 | 1 | 0.355 | 6.88E-150 | 3 L1 _ | Ly6c1 |
| Tmem881 | 2.13E-154 | 1.284048 | 0.619 | 0.057 | 4.29E-150 | 3 L1 _ | Tmem88 |
| Btnl91 | 1.57E-154 | 1.67197 | 0.712 | 0.082 | 3.16E-150 | 3 L1 _ | Btnl9 |
| Rapgef51 | 2.35E-155 | 1.595312 | 0.677 | 0.078 | 4.74E-151 | 3 L1 _ | Rapgef5 |
| Cd2001 | 2.15E-155 | 2.13356 | 0.915 | 0.194 | 4.33E-151 | 3 L1 _ | Cd200 |
| Cd300lg1 | 1.81E-156 | 1.796513 | 0.769 | 0.11 | 3.64E-152 | 3 L1 _ | Cd300lg |
| Itga61 | 3.38E-158 | 1.538434 | 0.773 | 0.106 | 6.81E-154 | 3 L1 _ | Itga6 |
| Aqp1 | 2.89E-159 | 2.173589 | 0.727 | 0.101 | 5.81E-155 | 3 L1 _ | Aqp1 |
| Adgrf51 | 2.61E-159 | 1.895289 | 0.881 | 0.16 | 5.26E-155 | 3 L1 _ | Adgrf5 |
| Tek2 | 5.22E-160 | 1.485466 | 0.738 | 0.094 | 1.05E-155 | 3 L1 _ | Tek |
| Rasgrp31 | 4.20E-160 | 1.408151 | 0.688 | 0.074 | 8.46E-156 | 3 L1 _ | Rasgrp3 |
| Grrp11 | 8.91E-161 | 1.470478 | 0.654 | 0.066 | 1.79E-156 | 3 L1 _ | Grrp1 |
| Igfbp31 | 2.72E-161 | 3.235897 | 0.888 | 0.196 | 5.47E-157 | 3 L1 _ | Igfbp3 |
| Thrsp1 | 2.66E-161 | 1.561309 | 0.762 | 0.095 | 5.37E-157 | 3 L1 _ | Thrsp |
| Mgll1 | 1.23E-161 | 2.030258 | 0.923 | 0.199 | 2.47E-157 | 3 L1 _ | Mgll |
| Ushbp11 | 1.34E-162 | 1.606179 | 0.735 | 0.092 | 2.69E-158 | 3 L1 _ | Ushbp1 |
| Egfl71 | 1.12E-165 | 2.901588 | 0.996 | 0.318 | 2.26E-161 | 3 L1 _ | Egfl7 |

| | | | | | | | |
|---|---|---|---|---|---|---|---|
| Tie11 | 4.66E-169 | 1.643916 | 0.815 | 0.111 | 9.39E-165 | 3 L1 _ | Tie1 |
| Emcn1 | 2.92E-172 | 2.431317 | 0.946 | 0.197 | 5.89E-168 | 3 L1 _ | Emcn |
| Icam21 | 4.11E-173 | 1.60089 | 0.75 | 0.09 | 8.28E-169 | 3 L1 _ | Icam2 |
| Cd931 | 1.46E-175 | 1.960879 | 0.808 | 0.108 | 2.93E-171 | 3 L1 _ | Cd93 |
| Rasip11 | 6.90E-180 | 1.525299 | 0.742 | 0.078 | 1.39E-175 | 3 L1 _ | Rasip1 |
| Pcdh171 | 1.28E-185 | 1.756564 | 0.735 | 0.07 | 2.58E-181 | 3 L1 _ | Pcdh17 |
| Gpihbp11 | 8.10E-186 | 2.8208 | 0.881 | 0.144 | 1.63E-181 | 3 L1 _ | Gpihbp1 |
| Flt11 | 6.20E-188 | 2.770948 | 0.985 | 0.206 | 1.25E-183 | 3 L1 _ | Flt1 |
| Kdr1 | 4.06E-188 | 2.237426 | 0.931 | 0.152 | 8.19E-184 | 3 L1 _ | Kdr |
| Scarb11 | 1.66E-191 | 2.217317 | 0.819 | 0.105 | 3.35E-187 | 3 L1 _ | Scarb1 |
| Myct11 | 4.48E-192 | 1.609764 | 0.731 | 0.067 | 9.03E-188 | 3 L1 _ | Myct1 |
| Ptprb1 | 1.01E-192 | 2.73324 | 0.981 | 0.198 | 2.03E-188 | 3 L1 _ | Ptprb |
| Podxl1 | 1.69E-195 | 1.977278 | 0.904 | 0.121 | 3.40E-191 | 3 L1 _ | Podxl |
| Pecam11 | 1.00E-197 | 2.713041 | 0.988 | 0.194 | 2.02E-193 | 3 L1 _ | Pecam1 |
| Clic51 | 1.12E-198 | 1.87346 | 0.8 | 0.079 | 2.25E-194 | 3 L1 _ | Clic5 |
| Adgrl41 | 3.78E-202 | 2.274273 | 0.9 | 0.121 | 7.60E-198 | 3 L1 _ | Adgrl4 |
| Mmrn21 | 3.54E-205 | 2.024343 | 0.831 | 0.092 | 7.12E-201 | 3 L1 _ | Mmrn2 |
| Cyyr11 | 5.57E-220 | 2.101367 | 0.869 | 0.087 | 1.12E-215 | 3 L1 _ | Cyyr1 |
| Cdh51 | 3.50E-231 | 2.369626 | 0.938 | 0.109 | 7.05E-227 | 3 L1 _ | Cdh5 |
| Rab7 | 2.49E-06 | 0.349418 | 0.829 | 0.642 | 0.050181 | 3 M1 _ | Rab7 |
| Tax1bp3 | 2.44E-06 | 0.391532 | 0.634 | 0.404 | 0.049081 | 3 M1 _ | Tax1bp3 |
| Usp25 | 2.36E-06 | 0.417775 | 0.598 | 0.379 | 0.047598 | 3 M1 _ | Usp25 |
| Mmd | 2.21E-06 | 0.277472 | 0.244 | 0.091 | 0.044495 | 3 M1 _ | Mmd |
| Lin7c | 2.16E-06 | 0.28423 | 0.439 | 0.22 | 0.043449 | 3 M1 _ | Lin7c |
| Socs2 | 2.14E-06 | 0.373606 | 0.744 | 0.488 | 0.043177 | 3 M1 _ | Socs2 |
| Capza2 | 2.11E-06 | 0.427399 | 0.854 | 0.689 | 0.042458 | 3 M1 _ | Capza2 |
| Myl61 | 2.08E-06 | 0.331547 | 1 | 0.966 | 0.041825 | 3 M1 _ | Myl6 |
| Trak2 | 2.03E-06 | 0.353681 | 0.366 | 0.167 | 0.040923 | 3 M1 _ | Trak2 |
| Otud4 | 2.01E-06 | 0.456293 | 0.427 | 0.22 | 0.040484 | 3 M1 _ | Otud4 |
| Metap2 | 1.84E-06 | 0.389509 | 0.854 | 0.692 | 0.037051 | 3 M1 _ | Metap2 |
| Erf | 1.78E-06 | 0.279129 | 0.305 | 0.126 | 0.035877 | 3 M1 _ | Erf |
| Ass1 | 1.78E-06 | 0.312907 | 0.329 | 0.143 | 0.035778 | 3 M1 _ | Ass1 |
| Fam126a | 1.77E-06 | 0.349877 | 0.354 | 0.159 | 0.035731 | 3 M1 _ | Fam126a |
| Map21 | 1.75E-06 | 0.34877 | 0.341 | 0.149 | 0.035151 | 3 M1 _ | Map2 |
| Ep400 | 1.74E-06 | 0.367279 | 0.573 | 0.343 | 0.035093 | 3 M1 _ | Ep400 |
| Emc2 | 1.73E-06 | 0.426884 | 0.61 | 0.386 | 0.034804 | 3 M1 _ | Emc2 |
| Kbtbd2 | 1.69E-06 | 0.347162 | 0.378 | 0.179 | 0.033999 | 3 M1 _ | Kbtbd2 |
| Plcb41 | 1.63E-06 | 0.371264 | 0.488 | 0.25 | 0.032825 | 3 M1 _ | Plcb4 |
| Mindy1 | 1.57E-06 | 0.39678 | 0.402 | 0.197 | 0.031685 | 3 M1 _ | Mindy1 |
| Kmt2d | 1.53E-06 | 0.380298 | 0.402 | 0.194 | 0.030791 | 3 M1 _ | Kmt2d |
| Ubc | 1.46E-06 | 0.33134 | 0.988 | 0.928 | 0.029439 | 3 M1 _ | Ubc |

| | | | | | | | |
|---|---|---|---|---|---|---|---|
| Iigp1 | 1.42E-06 | 0.561256 | 0.634 | 0.417 | 0.028523 | 3 M1 _ | Iigp1 |
| Atxn3 | 1.36E-06 | 0.267687 | 0.305 | 0.126 | 0.027374 | 3 M1 _ | Atxn3 |
| Ddx54 | 1.34E-06 | 0.373565 | 0.5 | 0.274 | 0.027005 | 3 M1 _ | Ddx54 |
| Fam220a2 | 1.31E-06 | 0.298119 | 0.183 | 0.055 | 0.026477 | 3 M1 _ | Fam220a |
| Nexn | 1.24E-06 | 0.384936 | 0.366 | 0.167 | 0.024979 | 3 M1 _ | Nexn |
| Arhgap5 | 1.22E-06 | 0.500464 | 0.78 | 0.589 | 0.024562 | 3 M1 _ | Arhgap5 |
| AU0202061 | 1.21E-06 | 0.27354 | 0.451 | 0.219 | 0.024289 | 3 M1 _ | AU020206 |
| G3bp1 | 1.19E-06 | 0.54656 | 0.72 | 0.491 | 0.024003 | 3 M1 _ | G3bp1 |
| Ywhae | 1.19E-06 | 0.450865 | 0.878 | 0.782 | 0.023964 | 3 M1 _ | Ywhae |
| Cpne8 | 1.13E-06 | 0.383866 | 0.561 | 0.33 | 0.022782 | 3 M1 _ | Cpne8 |
| Tsnax | 1.12E-06 | 0.346535 | 0.512 | 0.278 | 0.022613 | 3 M1 _ | Tsnax |
| Fam210b | 1.10E-06 | 0.284203 | 0.28 | 0.106 | 0.022158 | 3 M1 _ | Fam210b |
| Hnrnpf | 1.08E-06 | 0.36244 | 0.878 | 0.727 | 0.021768 | 3 M1 _ | Hnrnpf |
| Stox2 | 1.05E-06 | 0.448697 | 0.415 | 0.21 | 0.021095 | 3 M1 _ | Stox2 |
| Wdfy1 | 1.01E-06 | 0.315493 | 0.329 | 0.14 | 0.020266 | 3 M1 _ | Wdfy1 |
| Tcn2 | 9.77E-07 | 0.561866 | 0.695 | 0.515 | 0.019668 | 3 M1 _ | Tcn2 |
| Pdcl3 | 9.58E-07 | 0.49924 | 0.622 | 0.401 | 0.019292 | 3 M1 _ | Pdcl3 |
| Hdac7 | 8.91E-07 | 0.488597 | 0.671 | 0.444 | 0.017947 | 3 M1 _ | Hdac7 |
| Mef2a1 | 8.34E-07 | 0.414476 | 0.866 | 0.654 | 0.016789 | 3 M1 _ | Mef2a |
| Nomo1 | 8.09E-07 | 0.336475 | 0.341 | 0.148 | 0.016295 | 3 M1 _ | Nomo1 |
| Cyth1 | 7.91E-07 | 0.262712 | 0.268 | 0.101 | 0.015934 | 3 M1 _ | Cyth1 |
| Wbp4 | 7.84E-07 | 0.515952 | 0.659 | 0.467 | 0.015799 | 3 M1 _ | Wbp4 |
| Nras | 7.39E-07 | 0.414743 | 0.573 | 0.338 | 0.01488 | 3 M1 _ | Nras |
| Mef2d | 7.17E-07 | 0.422202 | 0.451 | 0.236 | 0.01445 | 3 M1 _ | Mef2d |
| Zmynd11 | 7.07E-07 | 0.463126 | 0.756 | 0.545 | 0.01424 | 3 M1 _ | Zmynd11 |
| Fkbp1a | 6.75E-07 | 0.444231 | 0.854 | 0.784 | 0.013594 | 3 M1 _ | Fkbp1a |
| Nhsl2 | 6.73E-07 | 0.367487 | 0.415 | 0.198 | 0.013549 | 3 M1 _ | Nhsl2 |
| Hras | 6.70E-07 | 0.492525 | 0.634 | 0.435 | 0.013502 | 3 M1 _ | Hras |
| Smurf2 | 6.64E-07 | 0.401843 | 0.549 | 0.307 | 0.013364 | 3 M1 _ | Smurf2 |
| Mndal | 6.56E-07 | 0.477909 | 0.866 | 0.661 | 0.013222 | 3 M1 _ | Mndal |
| Zfp462 | 6.56E-07 | 0.499497 | 0.354 | 0.158 | 0.013211 | 3 M1 _ | Zfp462 |
| Afap1 | 6.51E-07 | 0.467301 | 0.61 | 0.389 | 0.013117 | 3 M1 _ | Afap1 |
| Peak1 | 6.45E-07 | 0.42669 | 0.805 | 0.625 | 0.012991 | 3 M1 _ | Peak1 |
| Eno1 | 6.28E-07 | 0.536225 | 0.585 | 0.377 | 0.012651 | 3 M1 _ | Eno1 |
| Wdr37 | 6.21E-07 | 0.351382 | 0.329 | 0.141 | 0.012497 | 3 M1 _ | Wdr37 |
| Baz2b | 6.04E-07 | 0.532164 | 0.61 | 0.394 | 0.012167 | 3 M1 _ | Baz2b |
| Bmi1 | 5.73E-07 | 0.320622 | 0.341 | 0.151 | 0.011535 | 3 M1 _ | Bmi1 |
| B2m | 5.67E-07 | 0.409489 | 1 | 0.949 | 0.011424 | 3 M1 _ | B2m |
| Ctbp1 | 5.65E-07 | 0.457936 | 0.732 | 0.518 | 0.011376 | 3 M1 _ | Ctbp1 |
| Sin3a | 5.44E-07 | 0.350979 | 0.402 | 0.189 | 0.010962 | 3 M1 _ | Sin3a |
| Taok11 | 5.23E-07 | 0.424462 | 0.707 | 0.516 | 0.010528 | 3 M1 _ | Taok1 |

| | | | | | | | |
|---|---|---|---|---|---|---|---|
| Tmtc4 | 5.08E-07 | 0.258895 | 0.171 | 0.047 | 0.010241 | 3 M1 _ | Tmtc4 |
| Hif1a | 5.08E-07 | 0.533788 | 0.646 | 0.422 | 0.010225 | 3 M1 _ | Hif1a |
| Apc | 4.90E-07 | 0.441002 | 0.573 | 0.347 | 0.009872 | 3 M1 _ | Apc |
| Tns11 | 4.61E-07 | 0.527914 | 0.744 | 0.527 | 0.009276 | 3 M1 _ | Tns1 |
| Tmpo | 4.57E-07 | 0.396788 | 0.61 | 0.35 | 0.0092 | 3 M1 _ | Tmpo |
| Aplnr | 4.50E-07 | 0.289909 | 0.22 | 0.069 | 0.009067 | 3 M1 _ | Aplnr |
| Nqo2 | 4.50E-07 | 0.315046 | 0.329 | 0.134 | 0.00906 | 3 M1 _ | Nqo2 |
| Cnn21 | 4.33E-07 | 0.467813 | 0.695 | 0.436 | 0.008717 | 3 M1 _ | Cnn2 |
| Cic | 4.27E-07 | 0.339699 | 0.476 | 0.245 | 0.008591 | 3 M1 _ | Cic |
| Shoc2 | 4.19E-07 | 0.509617 | 0.61 | 0.365 | 0.008441 | 3 M1 _ | Shoc2 |
| Hbb-bs1 | 4.02E-07 | 3.356176 | 0.451 | 0.221 | 0.008099 | 3 M1 _ | Hbb-bs |
| Myl12b | 3.92E-07 | 0.430483 | 0.902 | 0.792 | 0.007886 | 3 M1 _ | Myl12b |
| Parp14 | 3.79E-07 | 0.405253 | 0.634 | 0.383 | 0.007624 | 3 M1 _ | Parp14 |
| Dab2 | 3.77E-07 | 0.536077 | 0.646 | 0.418 | 0.0076 | 3 M1 _ | Dab2 |
| Palm | 3.66E-07 | 0.492339 | 0.463 | 0.244 | 0.007374 | 3 M1 _ | Palm |
| Swap70 | 3.64E-07 | 0.484159 | 0.598 | 0.376 | 0.007339 | 3 M1 _ | Swap70 |
| Kifc3 | 3.64E-07 | 0.314083 | 0.305 | 0.116 | 0.007333 | 3 M1 _ | Kifc3 |
| Slco2b1 | 3.51E-07 | 0.53998 | 0.378 | 0.174 | 0.007074 | 3 M1 _ | Slco2b1 |
| Sh3bgrl1 | 3.32E-07 | 0.521376 | 0.817 | 0.682 | 0.006677 | 3 M1 _ | Sh3bgrl |
| Selenot1 | 3.18E-07 | 0.483157 | 0.622 | 0.384 | 0.006395 | 3 M1 _ | Selenot |
| Tollip | 3.00E-07 | 0.285433 | 0.329 | 0.131 | 0.006048 | 3 M1 _ | Tollip |
| Cnot6 | 2.86E-07 | 0.376321 | 0.61 | 0.36 | 0.005758 | 3 M1 _ | Cnot6 |
| Golm1 | 2.79E-07 | 0.472157 | 0.415 | 0.197 | 0.005629 | 3 M1 _ | Golm1 |
| Gtf2i | 2.76E-07 | 0.485971 | 0.659 | 0.422 | 0.005551 | 3 M1 _ | Gtf2i |
| Gapdh1 | 2.61E-07 | 0.395936 | 0.951 | 0.906 | 0.005262 | 3 M1 _ | Gapdh |
| Trappc6b | 2.50E-07 | 0.45896 | 0.707 | 0.499 | 0.005043 | 3 M1 _ | Trappc6b |
| Nmi | 2.39E-07 | 0.545241 | 0.561 | 0.309 | 0.004814 | 3 M1 _ | Nmi |
| Inpp5a | 2.36E-07 | 0.389926 | 0.451 | 0.221 | 0.004754 | 3 M1 _ | Inpp5a |
| Il6st1 | 2.32E-07 | 0.552152 | 0.78 | 0.572 | 0.004665 | 3 M1 _ | Il6st |
| Snrnp200 | 2.31E-07 | 0.447015 | 0.561 | 0.319 | 0.004651 | 3 M1 _ | Snrnp200 |
| Ppp1r15a | 2.25E-07 | 0.34542 | 0.549 | 0.279 | 0.004537 | 3 M1 _ | Ppp1r15a |
| Rgl2 | 2.16E-07 | 0.444585 | 0.378 | 0.173 | 0.004357 | 3 M1 _ | Rgl2 |
| Rab12 | 2.13E-07 | 0.466067 | 0.732 | 0.503 | 0.00429 | 3 M1 _ | Rab12 |
| Tm6sf1 | 2.12E-07 | 0.363333 | 0.305 | 0.116 | 0.004275 | 3 M1 _ | Tm6sf1 |
| 2700081O15Rik | 2.12E-07 | 0.37183 | 0.524 | 0.276 | 0.004271 | 3 M1 _ | 2700081O15Rik |
| Prdx4 | 2.12E-07 | 0.48574 | 0.732 | 0.516 | 0.00427 | 3 M1 _ | Prdx4 |
| Stim2 | 2.04E-07 | 0.478843 | 0.488 | 0.251 | 0.004103 | 3 M1 _ | Stim2 |
| Lrp5 | 1.85E-07 | 0.31869 | 0.341 | 0.138 | 0.003732 | 3 M1 _ | Lrp5 |
| Kat6a | 1.84E-07 | 0.485205 | 0.634 | 0.4 | 0.003712 | 3 M1 _ | Kat6a |
| Lifr1 | 1.82E-07 | 0.642673 | 0.634 | 0.403 | 0.003666 | 3 M1 _ | Lifr |
| Fbxo7 | 1.80E-07 | 0.258246 | 0.329 | 0.129 | 0.003631 | 3 M1 _ | Fbxo7 |

| | | | | | | | |
|---|---|---|---|---|---|---|---|
| Sptlc2 | 1.75E-07 | 0.48284 | 0.549 | 0.303 | 0.003518 | 3 M1 _ | Sptlc2 |
| Nmt11 | 1.69E-07 | 0.444513 | 0.902 | 0.655 | 0.003405 | 3 M1 _ | Nmt1 |
| Slk | 1.58E-07 | 0.543441 | 0.683 | 0.472 | 0.00319 | 3 M1 _ | Slk |
| Scoc1 | 1.55E-07 | 0.385008 | 0.61 | 0.341 | 0.003122 | 3 M1 _ | Scoc |
| Ptpn12 | 1.53E-07 | 0.4591 | 0.573 | 0.326 | 0.003091 | 3 M1 _ | Ptpn12 |
| Mapre1 | 1.52E-07 | 0.52158 | 0.756 | 0.583 | 0.00307 | 3 M1 _ | Mapre1 |
| Jade2 | 1.51E-07 | 0.389405 | 0.366 | 0.153 | 0.003045 | 3 M1 _ | Jade2 |
| Actb1 | 1.50E-07 | 0.330553 | 1 | 0.992 | 0.003018 | 3 M1 _ | Actb |
| H2-T23 | 1.48E-07 | 0.486151 | 0.78 | 0.527 | 0.002991 | 3 M1 _ | H2-T23 |
| Prkce | 1.48E-07 | 0.371123 | 0.293 | 0.11 | 0.002983 | 3 M1 _ | Prkce |
| Elovl5 | 1.41E-07 | 0.493158 | 0.463 | 0.233 | 0.002845 | 3 M1 _ | Elovl5 |
| Nploc4 | 1.36E-07 | 0.329136 | 0.305 | 0.118 | 0.00273 | 3 M1 _ | Nploc4 |
| Atxn1 | 1.32E-07 | 0.494572 | 0.451 | 0.231 | 0.002666 | 3 M1 _ | Atxn1 |
| Tapbp | 1.30E-07 | 0.643543 | 0.817 | 0.634 | 0.00261 | 3 M1 _ | Tapbp |
| Kctd10 | 1.28E-07 | 0.509975 | 0.573 | 0.316 | 0.002587 | 3 M1 _ | Kctd10 |
| Aldh22 | 1.27E-07 | 0.570659 | 0.915 | 0.753 | 0.002554 | 3 M1 _ | Aldh2 |
| Arhgdia | 1.26E-07 | 0.449307 | 0.902 | 0.764 | 0.002532 | 3 M1 _ | Arhgdia |
| Asap1 | 1.25E-07 | 0.554294 | 0.841 | 0.61 | 0.002519 | 3 M1 _ | Asap1 |
| Crk | 1.21E-07 | 0.62061 | 0.585 | 0.366 | 0.00244 | 3 M1 _ | Crk |
| Mtor | 1.20E-07 | 0.314557 | 0.317 | 0.123 | 0.002421 | 3 M1 _ | Mtor |
| Xylt2 | 1.16E-07 | 0.324665 | 0.329 | 0.13 | 0.002342 | 3 M1 _ | Xylt2 |
| Pdcl | 1.14E-07 | 0.371521 | 0.451 | 0.213 | 0.002305 | 3 M1 _ | Pdcl |
| Klc1 | 1.11E-07 | 0.508322 | 0.622 | 0.359 | 0.002245 | 3 M1 _ | Klc1 |
| Chd1 | 1.09E-07 | 0.510185 | 0.549 | 0.308 | 0.002201 | 3 M1 _ | Chd1 |
| Psmb9 | 1.07E-07 | 0.551177 | 0.634 | 0.374 | 0.002152 | 3 M1 _ | Psmb9 |
| Gosr2 | 1.01E-07 | 0.416175 | 0.439 | 0.209 | 0.002041 | 3 M1 _ | Gosr2 |
| Map3k1 | 9.45E-08 | 0.337358 | 0.366 | 0.153 | 0.001902 | 3 M1 _ | Map3k1 |
| Nrp2 | 8.81E-08 | 0.506683 | 0.622 | 0.339 | 0.001775 | 3 M1 _ | Nrp2 |
| Hnrnpa1 | 8.69E-08 | 0.502695 | 0.756 | 0.54 | 0.00175 | 3 M1 _ | Hnrnpa1 |
| Hipk11 | 8.35E-08 | 0.462284 | 0.61 | 0.367 | 0.001682 | 3 M1 _ | Hipk1 |
| Herc2 | 8.12E-08 | 0.603308 | 0.646 | 0.437 | 0.001635 | 3 M1 _ | Herc2 |
| Suz12 | 7.71E-08 | 0.411941 | 0.5 | 0.251 | 0.001552 | 3 M1 _ | Suz12 |
| Top1 | 7.38E-08 | 0.550426 | 0.854 | 0.703 | 0.001485 | 3 M1 _ | Top1 |
| Eogt | 7.31E-08 | 0.276682 | 0.232 | 0.072 | 0.001473 | 3 M1 _ | Eogt |
| Rbm8a1 | 7.27E-08 | 0.483719 | 0.817 | 0.574 | 0.001464 | 3 M1 _ | Rbm8a |
| Ddx21 | 6.67E-08 | 0.623422 | 0.695 | 0.445 | 0.001343 | 3 M1 _ | Ddx21 |
| Tmod3 | 6.47E-08 | 0.559866 | 0.78 | 0.606 | 0.001303 | 3 M1 _ | Tmod3 |
| Elmo1 | 6.24E-08 | 0.260763 | 0.366 | 0.144 | 0.001257 | 3 M1 _ | Elmo1 |
| Sertad2 | 5.94E-08 | 0.377305 | 0.561 | 0.287 | 0.001196 | 3 M1 _ | Sertad2 |
| Jun | 5.54E-08 | 0.914576 | 0.915 | 0.689 | 0.001116 | 3 M1 _ | Jun |
| Ywhaz1 | 5.33E-08 | 0.459694 | 0.902 | 0.806 | 0.001073 | 3 M1 _ | Ywhaz |

| | | | | | | | |
|---|---|---|---|---|---|---|---|
| Psmc1 | 5.30E-08 | 0.524611 | 0.756 | 0.503 | 0.001068 | 3 M1 _ | Psmc1 |
| Vamp3 | 4.77E-08 | 0.440685 | 0.573 | 0.314 | 0.00096 | 3 M1 _ | Vamp3 |
| Ivns1abp | 4.43E-08 | 0.571158 | 0.561 | 0.313 | 0.000891 | 3 M1 _ | Ivns1abp |
| Coro2b | 4.40E-08 | 0.275936 | 0.244 | 0.077 | 0.000885 | 3 M1 _ | Coro2b |
| Itga5 | 4.27E-08 | 0.35577 | 0.415 | 0.18 | 0.00086 | 3 M1 _ | Itga5 |
| Pfkfb3 | 4.25E-08 | 0.300511 | 0.183 | 0.047 | 0.000856 | 3 M1 _ | Pfkfb3 |
| Smim10l1 | 3.84E-08 | 0.536003 | 0.768 | 0.537 | 0.000773 | 3 M1 _ | Smim10l1 |
| Ifi35 | 3.79E-08 | 0.51731 | 0.659 | 0.393 | 0.000764 | 3 M1 _ | Ifi35 |
| Rab5c | 3.55E-08 | 0.407642 | 0.805 | 0.578 | 0.000715 | 3 M1 _ | Rab5c |
| Rapgef1 | 3.32E-08 | 0.481086 | 0.451 | 0.21 | 0.000668 | 3 M1 _ | Rapgef1 |
| Nod1 | 3.32E-08 | 0.363726 | 0.366 | 0.145 | 0.000668 | 3 M1 _ | Nod1 |
| Abca3 | 3.28E-08 | 0.345459 | 0.317 | 0.117 | 0.00066 | 3 M1 _ | Abca3 |
| Gimap5 | 3.21E-08 | 0.422617 | 0.476 | 0.206 | 0.000646 | 3 M1 _ | Gimap5 |
| Egln1 | 3.06E-08 | 0.49034 | 0.659 | 0.418 | 0.000616 | 3 M1 _ | Egln1 |
| Ucp21 | 2.97E-08 | 0.49671 | 0.622 | 0.327 | 0.000598 | 3 M1 _ | Ucp2 |
| Gpd2 | 2.87E-08 | 0.35252 | 0.317 | 0.119 | 0.000577 | 3 M1 _ | Gpd2 |
| Rb1 | 2.85E-08 | 0.530499 | 0.451 | 0.205 | 0.000574 | 3 M1 _ | Rb1 |
| Pogk | 2.61E-08 | 0.467434 | 0.366 | 0.149 | 0.000525 | 3 M1 _ | Pogk |
| Tmem30a | 2.50E-08 | 0.506115 | 0.793 | 0.604 | 0.000504 | 3 M1 _ | Tmem30a |
| Tlr3 | 2.44E-08 | 0.515411 | 0.378 | 0.156 | 0.000491 | 3 M1 _ | Tlr3 |
| Baiap2 | 2.33E-08 | 0.310632 | 0.366 | 0.145 | 0.00047 | 3 M1 _ | Baiap2 |
| Spop | 2.32E-08 | 0.548386 | 0.744 | 0.549 | 0.000467 | 3 M1 _ | Spop |
| Chmp5 | 2.30E-08 | 0.433602 | 0.817 | 0.543 | 0.000463 | 3 M1 _ | Chmp5 |
| Rnf19a | 2.26E-08 | 0.452589 | 0.439 | 0.204 | 0.000455 | 3 M1 _ | Rnf19a |
| Aig11 | 2.16E-08 | 0.331201 | 0.488 | 0.22 | 0.000435 | 3 M1 _ | Aig1 |
| Mxd4 | 2.16E-08 | 0.532879 | 0.866 | 0.772 | 0.000434 | 3 M1 _ | Mxd4 |
| Irf1 | 2.03E-08 | 0.643285 | 0.598 | 0.349 | 0.000409 | 3 M1 _ | Irf1 |
| Cdc42ep41 | 2.01E-08 | 0.464042 | 0.524 | 0.264 | 0.000405 | 3 M1 _ | Cdc42ep4 |
| Snx3 | 2.00E-08 | 0.481138 | 0.878 | 0.688 | 0.000403 | 3 M1 _ | Snx3 |
| Rras1 | 2.00E-08 | 0.542875 | 0.841 | 0.673 | 0.000402 | 3 M1 _ | Rras |
| Hey1 | 1.97E-08 | 0.442644 | 0.171 | 0.041 | 0.000397 | 3 M1 _ | Hey1 |
| Adarb1 | 1.94E-08 | 0.450674 | 0.402 | 0.169 | 0.000392 | 3 M1 _ | Adarb1 |
| Bend7 | 1.79E-08 | 0.30105 | 0.171 | 0.041 | 0.00036 | 3 M1 _ | Bend7 |
| Smtn1 | 1.76E-08 | 0.485775 | 0.561 | 0.303 | 0.000354 | 3 M1 _ | Smtn |
| Dock4 | 1.76E-08 | 0.385939 | 0.329 | 0.119 | 0.000353 | 3 M1 _ | Dock4 |
| B230219D22Rik1 | 1.74E-08 | 0.51611 | 0.866 | 0.704 | 0.000351 | 3 M1 _ | B230219D22Rik |
| Agrn | 1.74E-08 | 0.39384 | 0.439 | 0.198 | 0.000351 | 3 M1 _ | Agrn |
| Mink1 | 1.62E-08 | 0.382856 | 0.378 | 0.154 | 0.000326 | 3 M1 _ | Mink1 |
| Hprt | 1.54E-08 | 0.446983 | 0.585 | 0.324 | 0.00031 | 3 M1 _ | Hprt |
| Timeless | 1.28E-08 | 0.380465 | 0.305 | 0.107 | 0.000258 | 3 M1 _ | Timeless |
| Phldb2 | 1.08E-08 | 0.47617 | 0.573 | 0.295 | 0.000218 | 3 M1 _ | Phldb2 |

| | | | | | | | |
|---|---|---|---|---|---|---|---|
| Trim65 | 1.06E-08 | 0.295803 | 0.256 | 0.078 | 0.000213 | 3 M1 _ | Trim65 |
| Bst2 | 1.04E-08 | 0.578352 | 0.939 | 0.708 | 0.000209 | 3 M1 _ | Bst2 |
| Ddx3x | 1.04E-08 | 0.435451 | 0.854 | 0.661 | 0.000209 | 3 M1 _ | Ddx3x |
| Txn11 | 9.88E-09 | 0.502622 | 0.939 | 0.83 | 0.000199 | 3 M1 _ | Txn1 |
| Ace1 | 9.39E-09 | 0.644025 | 0.646 | 0.39 | 0.000189 | 3 M1 _ | Ace |
| Zhx31 | 9.28E-09 | 0.53838 | 0.585 | 0.322 | 0.000187 | 3 M1 _ | Zhx3 |
| Pfkl | 9.22E-09 | 0.418075 | 0.378 | 0.149 | 0.000186 | 3 M1 _ | Pfkl |
| Rnf44 | 9.02E-09 | 0.341574 | 0.402 | 0.165 | 0.000182 | 3 M1 _ | Rnf44 |
| Tmem63b | 8.63E-09 | 0.343241 | 0.354 | 0.13 | 0.000174 | 3 M1 _ | Tmem63b |
| Csgalnact1 | 8.52E-09 | 0.452359 | 0.415 | 0.175 | 0.000172 | 3 M1 _ | Csgalnact1 |
| Rin31 | 8.50E-09 | 0.259452 | 0.366 | 0.134 | 0.000171 | 3 M1 _ | Rin3 |
| Prrg2 | 7.49E-09 | 0.355191 | 0.305 | 0.104 | 0.000151 | 3 M1 _ | Prrg2 |
| Hhex | 7.38E-09 | 0.321276 | 0.256 | 0.077 | 0.000149 | 3 M1 _ | Hhex |
| Grap | 7.08E-09 | 0.256773 | 0.378 | 0.14 | 0.000143 | 3 M1 _ | Grap |
| Card10 | 6.87E-09 | 0.356757 | 0.341 | 0.124 | 0.000138 | 3 M1 _ | Card10 |
| Fbxl7 | 6.85E-09 | 0.319886 | 0.512 | 0.228 | 0.000138 | 3 M1 _ | Fbxl7 |
| Pak2 | 6.72E-09 | 0.544644 | 0.841 | 0.654 | 0.000135 | 3 M1 _ | Pak2 |
| Plekhg5 | 6.58E-09 | 0.357396 | 0.305 | 0.105 | 0.000132 | 3 M1 _ | Plekhg5 |
| Znfx1 | 6.42E-09 | 0.484301 | 0.512 | 0.252 | 0.000129 | 3 M1 _ | Znfx1 |
| Prkd2 | 6.24E-09 | 0.348432 | 0.354 | 0.134 | 0.000126 | 3 M1 _ | Prkd2 |
| Dusp6 | 6.19E-09 | 0.683074 | 0.561 | 0.309 | 0.000125 | 3 M1 _ | Dusp6 |
| Setx | 5.99E-09 | 0.501559 | 0.476 | 0.223 | 0.000121 | 3 M1 _ | Setx |
| Atp11c | 5.93E-09 | 0.356887 | 0.378 | 0.146 | 0.00012 | 3 M1 _ | Atp11c |
| Smarca4 | 5.48E-09 | 0.646442 | 0.512 | 0.265 | 0.00011 | 3 M1 _ | Smarca4 |
| Cab39 | 5.38E-09 | 0.521483 | 0.573 | 0.315 | 0.000108 | 3 M1 _ | Cab39 |
| Gng111 | 5.33E-09 | 0.447123 | 0.939 | 0.816 | 0.000107 | 3 M1 _ | Gng11 |
| Ttc7b | 5.24E-09 | 0.38612 | 0.244 | 0.072 | 0.000105 | 3 M1 _ | Ttc7b |
| Myh91 | 5.07E-09 | 0.652404 | 0.915 | 0.826 | 0.000102 | 3 M1 _ | Myh9 |
| Wasf2 | 5.02E-09 | 0.592651 | 0.817 | 0.724 | 0.000101 | 3 M1 _ | Wasf2 |
| Pcnx | 4.99E-09 | 0.403914 | 0.512 | 0.243 | 0.0001 | 3 M1 _ | Pcnx |
| Gna11 | 4.58E-09 | 0.625773 | 0.78 | 0.61 | 9.23E-05 | 3 M1 _ | Gna11 |
| Ybx1 | 4.46E-09 | 0.449504 | 0.963 | 0.906 | 8.99E-05 | 3 M1 _ | Ybx1 |
| Fes | 4.46E-09 | 0.388723 | 0.329 | 0.116 | 8.98E-05 | 3 M1 _ | Fes |
| Ppp1r21 | 4.43E-09 | 0.576475 | 0.878 | 0.681 | 8.92E-05 | 3 M1 _ | Ppp1r2 |
| Hk1 | 4.27E-09 | 0.488563 | 0.585 | 0.306 | 8.60E-05 | 3 M1 _ | Hk1 |
| Tmem50b | 4.21E-09 | 0.358806 | 0.39 | 0.155 | 8.47E-05 | 3 M1 _ | Tmem50b |
| Picalm | 4.18E-09 | 0.549458 | 0.793 | 0.583 | 8.43E-05 | 3 M1 _ | Picalm |
| Birc6 | 4.17E-09 | 0.641134 | 0.768 | 0.563 | 8.39E-05 | 3 M1 _ | Birc6 |
| Cd2ap | 3.85E-09 | 0.591544 | 0.78 | 0.569 | 7.75E-05 | 3 M1 _ | Cd2ap |
| Fbxo22 | 3.80E-09 | 0.520791 | 0.549 | 0.275 | 7.66E-05 | 3 M1 _ | Fbxo22 |
| Trim30a | 3.66E-09 | 0.564835 | 0.573 | 0.295 | 7.37E-05 | 3 M1 _ | Trim30a |

| | | | | | | | |
|---|---|---|---|---|---|---|---|
| Pdlim5 | 3.50E-09 | 0.448079 | 0.585 | 0.315 | 7.04E-05 | 3 M1 _ | Pdlim5 |
| Iqsec2 | 3.46E-09 | 0.372407 | 0.317 | 0.109 | 6.97E-05 | 3 M1 _ | Iqsec2 |
| Smpdl3a1 | 3.34E-09 | 0.623336 | 0.744 | 0.471 | 6.72E-05 | 3 M1 _ | Smpdl3a |
| Xpo1 | 3.30E-09 | 0.325943 | 0.5 | 0.225 | 6.65E-05 | 3 M1 _ | Xpo1 |
| Ralgapa1 | 3.11E-09 | 0.607909 | 0.61 | 0.319 | 6.26E-05 | 3 M1 _ | Ralgapa1 |
| Cd812 | 3.10E-09 | 0.455266 | 0.963 | 0.894 | 6.24E-05 | 3 M1 _ | Cd81 |
| Slc35b1 | 2.93E-09 | 0.552916 | 0.573 | 0.296 | 5.89E-05 | 3 M1 _ | Slc35b1 |
| Stx2 | 2.87E-09 | 0.526005 | 0.524 | 0.261 | 5.78E-05 | 3 M1 _ | Stx2 |
| Slc3a2 | 2.83E-09 | 0.640805 | 0.805 | 0.579 | 5.70E-05 | 3 M1 _ | Slc3a2 |
| Itm2b | 2.79E-09 | 0.473966 | 1 | 0.99 | 5.61E-05 | 3 M1 _ | Itm2b |
| Dusp12 | 2.78E-09 | 0.95082 | 0.598 | 0.329 | 5.59E-05 | 3 M1 _ | Dusp1 |
| Kcnb11 | 2.65E-09 | 0.346512 | 0.232 | 0.063 | 5.33E-05 | 3 M1 _ | Kcnb1 |
| Flywch1 | 2.63E-09 | 0.631988 | 0.573 | 0.293 | 5.29E-05 | 3 M1 _ | Flywch1 |
| Prex2 | 2.58E-09 | 0.624567 | 0.695 | 0.409 | 5.20E-05 | 3 M1 _ | Prex2 |
| Piezo1 | 2.28E-09 | 0.422995 | 0.341 | 0.126 | 4.59E-05 | 3 M1 _ | Piezo1 |
| Kcnq1 | 2.21E-09 | 0.309341 | 0.171 | 0.037 | 4.45E-05 | 3 M1 _ | Kcnq1 |
| Il10rb | 2.19E-09 | 0.597438 | 0.622 | 0.348 | 4.42E-05 | 3 M1 _ | Il10rb |
| Reep31 | 2.15E-09 | 0.591079 | 0.878 | 0.691 | 4.33E-05 | 3 M1 _ | Reep3 |
| Pip5k1c | 2.12E-09 | 0.388814 | 0.451 | 0.186 | 4.26E-05 | 3 M1 _ | Pip5k1c |
| Dram2 | 2.01E-09 | 0.459194 | 0.488 | 0.215 | 4.05E-05 | 3 M1 _ | Dram2 |
| Ablim3 | 1.92E-09 | 0.283815 | 0.146 | 0.028 | 3.86E-05 | 3 M1 _ | Ablim3 |
| BC0055371 | 1.89E-09 | 0.537672 | 0.72 | 0.442 | 3.80E-05 | 3 M1 _ | BC005537 |
| Irf2bpl | 1.81E-09 | 0.606263 | 0.744 | 0.502 | 3.65E-05 | 3 M1 _ | Irf2bpl |
| Nrbp1 | 1.69E-09 | 0.650811 | 0.744 | 0.488 | 3.41E-05 | 3 M1 _ | Nrbp1 |
| Ctnnd1 | 1.59E-09 | 0.546168 | 0.61 | 0.332 | 3.20E-05 | 3 M1 _ | Ctnnd1 |
| 4833403J16Rik | 1.58E-09 | 0.314798 | 0.22 | 0.056 | 3.18E-05 | 3 M1 _ | 4833403J16Rik |
| Arhgef10 | 1.56E-09 | 0.26032 | 0.378 | 0.134 | 3.14E-05 | 3 M1 _ | Arhgef10 |
| Tspan151 | 1.43E-09 | 0.340929 | 0.439 | 0.163 | 2.89E-05 | 3 M1 _ | Tspan15 |
| Csnk1a1 | 1.32E-09 | 0.508275 | 0.951 | 0.803 | 2.66E-05 | 3 M1 _ | Csnk1a1 |
| Casz1 | 1.14E-09 | 0.338086 | 0.146 | 0.028 | 2.30E-05 | 3 M1 _ | Casz1 |
| Ktn11 | 1.11E-09 | 0.54973 | 0.829 | 0.619 | 2.24E-05 | 3 M1 _ | Ktn1 |
| Mark4 | 1.02E-09 | 0.359747 | 0.378 | 0.14 | 2.05E-05 | 3 M1 _ | Mark4 |
| Hs3st1 | 1.01E-09 | 0.251617 | 0.183 | 0.04 | 2.03E-05 | 3 M1 _ | Hs3st1 |
| Kif13a | 1.00E-09 | 0.3575 | 0.378 | 0.136 | 2.02E-05 | 3 M1 _ | Kif13a |
| Adamts9 | 9.94E-10 | 0.538638 | 0.451 | 0.183 | 2.00E-05 | 3 M1 _ | Adamts9 |
| Psme1 | 9.78E-10 | 0.563337 | 0.902 | 0.724 | 1.97E-05 | 3 M1 _ | Psme1 |
| Ier2 | 9.55E-10 | 0.745917 | 0.744 | 0.449 | 1.92E-05 | 3 M1 _ | Ier2 |
| Mgat1 | 9.35E-10 | 0.539778 | 0.598 | 0.324 | 1.88E-05 | 3 M1 _ | Mgat1 |
| Sorbs1 | 8.60E-10 | 0.377115 | 0.427 | 0.161 | 1.73E-05 | 3 M1 _ | Sorbs1 |
| Npr2 | 8.50E-10 | 0.493324 | 0.378 | 0.149 | 1.71E-05 | 3 M1 _ | Npr2 |
| Git1 | 8.05E-10 | 0.383134 | 0.463 | 0.193 | 1.62E-05 | 3 M1 _ | Git1 |

| | | | | | | | |
|---|---|---|---|---|---|---|---|
| Myo10 | 7.69E-10 | 0.851677 | 0.695 | 0.445 | 1.55E-05 | 3 M1 _ | Myo10 |
| Fos | 7.29E-10 | 1.546778 | 0.744 | 0.494 | 1.47E-05 | 3 M1 _ | Fos |
| Tbx31 | 6.61E-10 | 0.437111 | 0.39 | 0.14 | 1.33E-05 | 3 M1 _ | Tbx3 |
| Zfp36 | 6.33E-10 | 0.898027 | 0.524 | 0.249 | 1.27E-05 | 3 M1 _ | Zfp36 |
| Gemin7 | 5.91E-10 | 0.518364 | 0.598 | 0.315 | 1.19E-05 | 3 M1 _ | Gemin7 |
| Tspan7 | 5.83E-10 | 0.369565 | 0.232 | 0.06 | 1.17E-05 | 3 M1 _ | Tspan7 |
| Sh3bgrl2 | 5.79E-10 | 0.275214 | 0.195 | 0.045 | 1.17E-05 | 3 M1 _ | Sh3bgrl2 |
| Abcc41 | 5.54E-10 | 0.277627 | 0.305 | 0.094 | 1.12E-05 | 3 M1 _ | Abcc4 |
| Per1 | 5.35E-10 | 0.350123 | 0.329 | 0.107 | 1.08E-05 | 3 M1 _ | Per1 |
| Stard4 | 5.32E-10 | 0.460587 | 0.305 | 0.097 | 1.07E-05 | 3 M1 _ | Stard4 |
| Itpr3 | 5.28E-10 | 0.416836 | 0.28 | 0.084 | 1.06E-05 | 3 M1 _ | Itpr3 |
| Dock1 | 5.19E-10 | 0.793184 | 0.622 | 0.375 | 1.05E-05 | 3 M1 _ | Dock1 |
| Hbegf1 | 4.02E-10 | 0.341594 | 0.317 | 0.098 | 8.09E-06 | 3 M1 _ | Hbegf |
| Tjp2 | 3.97E-10 | 0.380033 | 0.293 | 0.089 | 7.99E-06 | 3 M1 _ | Tjp2 |
| Ttc39c | 3.78E-10 | 0.387694 | 0.232 | 0.062 | 7.62E-06 | 3 M1 _ | Ttc39c |
| Sppl2a | 3.78E-10 | 0.657471 | 0.756 | 0.518 | 7.62E-06 | 3 M1 _ | Sppl2a |
| Dennd5a | 3.69E-10 | 0.492232 | 0.646 | 0.35 | 7.43E-06 | 3 M1 _ | Dennd5a |
| Grb101 | 3.65E-10 | 0.665354 | 0.634 | 0.347 | 7.35E-06 | 3 M1 _ | Grb10 |
| S100a1 | 3.58E-10 | 0.595397 | 0.707 | 0.436 | 7.22E-06 | 3 M1 _ | S100a1 |
| 4930523C07Rik1 | 3.14E-10 | 0.564475 | 0.732 | 0.425 | 6.33E-06 | 3 M1 _ | 4930523C07Rik |
| Gas6 | 3.04E-10 | 0.581227 | 0.89 | 0.559 | 6.13E-06 | 3 M1 _ | Gas6 |
| S100a13 | 2.84E-10 | 0.595373 | 0.939 | 0.752 | 5.71E-06 | 3 M1 _ | S100a13 |
| Nedd9 | 2.63E-10 | 0.681458 | 0.622 | 0.343 | 5.30E-06 | 3 M1 _ | Nedd9 |
| Rgl1 | 2.46E-10 | 0.631469 | 0.72 | 0.455 | 4.95E-06 | 3 M1 _ | Rgl1 |
| Msi2 | 2.16E-10 | 0.667896 | 0.61 | 0.311 | 4.36E-06 | 3 M1 _ | Msi2 |
| Fam219a | 1.83E-10 | 0.284759 | 0.317 | 0.098 | 3.68E-06 | 3 M1 _ | Fam219a |
| Ifi208 | 1.78E-10 | 0.308253 | 0.256 | 0.067 | 3.58E-06 | 3 M1 _ | Ifi208 |
| Foxf1 | 1.74E-10 | 0.263618 | 0.171 | 0.033 | 3.50E-06 | 3 M1 _ | Foxf1 |
| Psmb8 | 1.69E-10 | 0.604835 | 0.902 | 0.623 | 3.41E-06 | 3 M1 _ | Psmb8 |
| Mcl1 | 1.69E-10 | 0.629093 | 0.866 | 0.622 | 3.40E-06 | 3 M1 _ | Mcl1 |
| Arid5b | 1.68E-10 | 0.760517 | 0.829 | 0.602 | 3.38E-06 | 3 M1 _ | Arid5b |
| Samd9l | 1.47E-10 | 0.64051 | 0.695 | 0.362 | 2.96E-06 | 3 M1 _ | Samd9l |
| Fbln21 | 1.39E-10 | 0.917473 | 0.805 | 0.537 | 2.80E-06 | 3 M1 _ | Fbln2 |
| Tshz2 | 1.35E-10 | 0.65655 | 0.89 | 0.633 | 2.71E-06 | 3 M1 _ | Tshz2 |
| Xdh1 | 1.27E-10 | 0.673199 | 0.744 | 0.422 | 2.56E-06 | 3 M1 _ | Xdh |
| Map3k5 | 1.22E-10 | 0.375555 | 0.415 | 0.15 | 2.46E-06 | 3 M1 _ | Map3k5 |
| Stap2 | 1.17E-10 | 0.285539 | 0.195 | 0.042 | 2.37E-06 | 3 M1 _ | Stap2 |
| Dock6 | 1.14E-10 | 0.40785 | 0.476 | 0.188 | 2.29E-06 | 3 M1 _ | Dock6 |
| Dip2b | 1.12E-10 | 0.545082 | 0.549 | 0.251 | 2.25E-06 | 3 M1 _ | Dip2b |
| Plxna4 | 1.11E-10 | 0.337808 | 0.159 | 0.029 | 2.23E-06 | 3 M1 _ | Plxna4 |
| Zfp715 | 1.08E-10 | 0.350006 | 0.305 | 0.09 | 2.18E-06 | 3 M1 _ | Zfp715 |

| | | | | | | | |
|---|---|---|---|---|---|---|---|
| Grasp | 1.05E-10 | 0.545465 | 0.512 | 0.226 | 2.12E-06 | 3 M1 _ | Grasp |
| Arhgef5 | 9.43E-11 | 0.535811 | 0.439 | 0.175 | 1.90E-06 | 3 M1 _ | Arhgef5 |
| Ptbp3 | 9.16E-11 | 0.72214 | 0.793 | 0.583 | 1.85E-06 | 3 M1 _ | Ptbp3 |
| Etnk1 | 8.63E-11 | 0.707373 | 0.72 | 0.432 | 1.74E-06 | 3 M1 _ | Etnk1 |
| Sgms1 | 8.14E-11 | 0.398642 | 0.5 | 0.201 | 1.64E-06 | 3 M1 _ | Sgms1 |
| Gbp9 | 7.79E-11 | 0.477897 | 0.451 | 0.174 | 1.57E-06 | 3 M1 _ | Gbp9 |
| Zmiz1 | 7.70E-11 | 0.636664 | 0.78 | 0.503 | 1.55E-06 | 3 M1 _ | Zmiz1 |
| B3gnt3 | 7.49E-11 | 0.35004 | 0.256 | 0.066 | 1.51E-06 | 3 M1 _ | B3gnt3 |
| Gatad2a | 7.37E-11 | 0.635438 | 0.622 | 0.321 | 1.48E-06 | 3 M1 _ | Gatad2a |
| Bag5 | 7.12E-11 | 0.534768 | 0.707 | 0.393 | 1.43E-06 | 3 M1 _ | Bag5 |
| Lyn | 6.88E-11 | 0.476228 | 0.463 | 0.183 | 1.39E-06 | 3 M1 _ | Lyn |
| Zfp664 | 6.02E-11 | 0.519197 | 0.598 | 0.298 | 1.21E-06 | 3 M1 _ | Zfp664 |
| Gbp7 | 5.96E-11 | 0.820253 | 0.732 | 0.444 | 1.20E-06 | 3 M1 _ | Gbp7 |
| Trim47 | 5.92E-11 | 0.648606 | 0.695 | 0.397 | 1.19E-06 | 3 M1 _ | Trim47 |
| Cit | 5.91E-11 | 0.303915 | 0.146 | 0.024 | 1.19E-06 | 3 M1 _ | Cit |
| Rap1b | 5.81E-11 | 0.707846 | 0.854 | 0.681 | 1.17E-06 | 3 M1 _ | Rap1b |
| Elf1 | 5.04E-11 | 0.653793 | 0.756 | 0.456 | 1.02E-06 | 3 M1 _ | Elf1 |
| St3gal4 | 4.93E-11 | 0.413169 | 0.512 | 0.214 | 9.92E-07 | 3 M1 _ | St3gal4 |
| Sh3bp5 | 4.77E-11 | 0.624448 | 0.549 | 0.247 | 9.60E-07 | 3 M1 _ | Sh3bp5 |
| Ywhaq1 | 4.76E-11 | 0.543547 | 0.878 | 0.795 | 9.58E-07 | 3 M1 _ | Ywhaq |
| Fnbp1l | 4.26E-11 | 0.582755 | 0.476 | 0.195 | 8.59E-07 | 3 M1 _ | Fnbp1l |
| Tlnrd1 | 4.20E-11 | 0.370516 | 0.427 | 0.157 | 8.46E-07 | 3 M1 _ | Tlnrd1 |
| Cactin | 4.11E-11 | 0.400872 | 0.366 | 0.123 | 8.27E-07 | 3 M1 _ | Cactin |
| Nfe2l1 | 4.06E-11 | 0.803198 | 0.817 | 0.636 | 8.18E-07 | 3 M1 _ | Nfe2l1 |
| Stk10 | 4.02E-11 | 0.423078 | 0.415 | 0.146 | 8.09E-07 | 3 M1 _ | Stk10 |
| Syt7 | 3.89E-11 | 0.273782 | 0.11 | 0.014 | 7.84E-07 | 3 M1 _ | Syt7 |
| Cttnbp2nl | 3.84E-11 | 0.50269 | 0.402 | 0.147 | 7.74E-07 | 3 M1 _ | Cttnbp2nl |
| Erg | 3.83E-11 | 0.30228 | 0.354 | 0.112 | 7.72E-07 | 3 M1 _ | Erg |
| Atp8a1 | 3.54E-11 | 0.389662 | 0.378 | 0.125 | 7.13E-07 | 3 M1 _ | Atp8a1 |
| Ada | 3.35E-11 | 0.710264 | 0.293 | 0.088 | 6.74E-07 | 3 M1 _ | Ada |
| Spry41 | 3.28E-11 | 0.587332 | 0.549 | 0.257 | 6.60E-07 | 3 M1 _ | Spry4 |
| Atp1a1 | 3.26E-11 | 0.720154 | 0.732 | 0.431 | 6.57E-07 | 3 M1 _ | Atp1a1 |
| Cep85l | 3.02E-11 | 0.530878 | 0.378 | 0.129 | 6.08E-07 | 3 M1 _ | Cep85l |
| Atox1 | 3.01E-11 | 0.612751 | 0.902 | 0.78 | 6.06E-07 | 3 M1 _ | Atox1 |
| Tmem184b | 2.50E-11 | 0.490307 | 0.488 | 0.201 | 5.03E-07 | 3 M1 _ | Tmem184b |
| H2afy2 | 2.26E-11 | 0.455352 | 0.378 | 0.128 | 4.56E-07 | 3 M1 _ | H2afy2 |
| Tpm3 | 2.25E-11 | 0.696982 | 0.915 | 0.746 | 4.54E-07 | 3 M1 _ | Tpm3 |
| Arhgef28 | 2.05E-11 | 0.253523 | 0.195 | 0.039 | 4.13E-07 | 3 M1 _ | Arhgef28 |
| Nedd4l | 2.03E-11 | 0.430819 | 0.305 | 0.089 | 4.09E-07 | 3 M1 _ | Nedd4l |
| Sos1 | 1.93E-11 | 0.576781 | 0.476 | 0.191 | 3.90E-07 | 3 M1 _ | Sos1 |
| Galnt7 | 1.83E-11 | 0.418818 | 0.463 | 0.175 | 3.68E-07 | 3 M1 _ | Galnt7 |

| | | | | | | | |
|---|---|---|---|---|---|---|---|
| Cdc14a | 1.82E-11 | 0.369506 | 0.329 | 0.098 | 3.66E-07 | 3 M1 _ | Cdc14a |
| Csde11 | 1.70E-11 | 0.589352 | 0.866 | 0.697 | 3.43E-07 | 3 M1 _ | Csde1 |
| Rtl8b | 1.67E-11 | 0.65404 | 0.598 | 0.284 | 3.37E-07 | 3 M1 _ | Rtl8b |
| Edn1 | 1.60E-11 | 1.158525 | 0.159 | 0.028 | 3.22E-07 | 3 M1 _ | Edn1 |
| Cd274 | 1.36E-11 | 0.337958 | 0.256 | 0.063 | 2.74E-07 | 3 M1 _ | Cd274 |
| Fgd6 | 1.32E-11 | 0.293604 | 0.22 | 0.048 | 2.66E-07 | 3 M1 _ | Fgd6 |
| Arl15 | 1.14E-11 | 0.490492 | 0.366 | 0.122 | 2.29E-07 | 3 M1 _ | Arl15 |
| Cyb561 | 1.09E-11 | 0.388385 | 0.329 | 0.098 | 2.19E-07 | 3 M1 _ | Cyb561 |
| Sh3glb1 | 9.46E-12 | 0.61629 | 0.915 | 0.81 | 1.91E-07 | 3 M1 _ | Sh3glb1 |
| Rala | 9.06E-12 | 0.587627 | 0.744 | 0.438 | 1.82E-07 | 3 M1 _ | Rala |
| Mecom | 8.82E-12 | 0.568641 | 0.28 | 0.073 | 1.78E-07 | 3 M1 _ | Mecom |
| Rac1 | 7.99E-12 | 0.646475 | 0.939 | 0.754 | 1.61E-07 | 3 M1 _ | Rac1 |
| Pik3cb | 7.39E-12 | 0.273776 | 0.22 | 0.047 | 1.49E-07 | 3 M1 _ | Pik3cb |
| Serinc3 | 6.63E-12 | 0.718485 | 0.902 | 0.858 | 1.33E-07 | 3 M1 _ | Serinc3 |
| Ndufa4 | 6.47E-12 | 0.670282 | 0.915 | 0.73 | 1.30E-07 | 3 M1 _ | Ndufa4 |
| Midn | 5.56E-12 | 0.664749 | 0.659 | 0.351 | 1.12E-07 | 3 M1 _ | Midn |
| Larp1b | 5.13E-12 | 0.544463 | 0.39 | 0.128 | 1.03E-07 | 3 M1 _ | Larp1b |
| Ggta11 | 4.74E-12 | 0.303122 | 0.39 | 0.119 | 9.54E-08 | 3 M1 _ | Ggta1 |
| Kras | 4.23E-12 | 0.566317 | 0.756 | 0.436 | 8.53E-08 | 3 M1 _ | Kras |
| Ptk21 | 3.91E-12 | 0.525426 | 0.671 | 0.34 | 7.87E-08 | 3 M1 _ | Ptk2 |
| St3gal11 | 3.71E-12 | 0.553306 | 0.524 | 0.225 | 7.48E-08 | 3 M1 _ | St3gal1 |
| Fmnl3 | 3.44E-12 | 0.644506 | 0.598 | 0.288 | 6.93E-08 | 3 M1 _ | Fmnl3 |
| Adipor1 | 3.21E-12 | 0.78695 | 0.695 | 0.424 | 6.47E-08 | 3 M1 _ | Adipor1 |
| Baz1a | 3.10E-12 | 0.52563 | 0.439 | 0.159 | 6.24E-08 | 3 M1 _ | Baz1a |
| Magi1 | 2.98E-12 | 0.369009 | 0.244 | 0.055 | 6.00E-08 | 3 M1 _ | Magi1 |
| Stmn1 | 2.65E-12 | 0.504512 | 0.341 | 0.102 | 5.34E-08 | 3 M1 _ | Stmn1 |
| Fam181b | 2.57E-12 | 0.373614 | 0.207 | 0.042 | 5.18E-08 | 3 M1 _ | Fam181b |
| Rtn3 | 2.35E-12 | 0.688385 | 0.829 | 0.604 | 4.74E-08 | 3 M1 _ | Rtn3 |
| Gda | 2.25E-12 | 0.469293 | 0.293 | 0.075 | 4.53E-08 | 3 M1 _ | Gda |
| Tmed5 | 2.05E-12 | 0.832382 | 0.72 | 0.404 | 4.12E-08 | 3 M1 _ | Tmed5 |
| Col4a3bp | 2.01E-12 | 0.652231 | 0.671 | 0.358 | 4.05E-08 | 3 M1 _ | Col4a3bp |
| Nxf12 | 1.99E-12 | 0.547336 | 0.598 | 0.264 | 4.00E-08 | 3 M1 _ | Nxf1 |
| Sh3rf1 | 1.95E-12 | 0.347377 | 0.354 | 0.104 | 3.93E-08 | 3 M1 _ | Sh3rf1 |
| Gse1 | 1.54E-12 | 0.444494 | 0.402 | 0.128 | 3.10E-08 | 3 M1 _ | Gse1 |
| Tnk2 | 1.54E-12 | 0.550859 | 0.476 | 0.181 | 3.09E-08 | 3 M1 _ | Tnk2 |
| Ift172 | 1.47E-12 | 0.49909 | 0.354 | 0.107 | 2.96E-08 | 3 M1 _ | Ift172 |
| Coro1b2 | 1.28E-12 | 0.394808 | 0.341 | 0.096 | 2.57E-08 | 3 M1 _ | Coro1b |
| Pxn | 1.26E-12 | 0.544896 | 0.512 | 0.2 | 2.55E-08 | 3 M1 _ | Pxn |
| Foxo1 | 1.01E-12 | 0.570789 | 0.61 | 0.277 | 2.03E-08 | 3 M1 _ | Foxo1 |
| Fam57a | 9.86E-13 | 0.362729 | 0.195 | 0.036 | 1.99E-08 | 3 M1 _ | Fam57a |
| Srf | 9.01E-13 | 0.344088 | 0.244 | 0.054 | 1.81E-08 | 3 M1 _ | Srf |

| | | | | | | | |
|---|---|---|---|---|---|---|---|
| Ywhab1 | 8.69E-13 | 0.609412 | 0.963 | 0.829 | 1.75E-08 | 3 M1 _ | Ywhab |
| Zfp46 | 6.39E-13 | 0.411075 | 0.378 | 0.118 | 1.29E-08 | 3 M1 _ | Zfp46 |
| Dusp16 | 6.30E-13 | 0.378771 | 0.256 | 0.058 | 1.27E-08 | 3 M1 _ | Dusp16 |
| Chmp3 | 5.45E-13 | 0.666397 | 0.841 | 0.559 | 1.10E-08 | 3 M1 _ | Chmp3 |
| Tanc1 | 5.18E-13 | 0.61368 | 0.659 | 0.309 | 1.04E-08 | 3 M1 _ | Tanc1 |
| Dusp7 | 4.97E-13 | 0.491217 | 0.512 | 0.193 | 1.00E-08 | 3 M1 _ | Dusp7 |
| Tns2 | 4.93E-13 | 0.689741 | 0.671 | 0.34 | 9.93E-09 | 3 M1 _ | Tns2 |
| Rbms1 | 4.67E-13 | 0.758353 | 0.927 | 0.774 | 9.40E-09 | 3 M1 _ | Rbms1 |
| Arhgap261 | 4.00E-13 | 0.309686 | 0.28 | 0.066 | 8.05E-09 | 3 M1 _ | Arhgap26 |
| Dock9 | 3.77E-13 | 0.482222 | 0.549 | 0.218 | 7.59E-09 | 3 M1 _ | Dock9 |
| Mapk12 | 3.54E-13 | 0.449937 | 0.427 | 0.138 | 7.13E-09 | 3 M1 _ | Mapk12 |
| Sigirr | 3.40E-13 | 0.372986 | 0.329 | 0.087 | 6.85E-09 | 3 M1 _ | Sigirr |
| Fam129b | 3.27E-13 | 0.521349 | 0.537 | 0.215 | 6.58E-09 | 3 M1 _ | Fam129b |
| Sgk3 | 3.12E-13 | 0.480494 | 0.402 | 0.126 | 6.28E-09 | 3 M1 _ | Sgk3 |
| Cd59a | 3.07E-13 | 0.330792 | 0.39 | 0.116 | 6.17E-09 | 3 M1 _ | Cd59a |
| Capn2 | 2.90E-13 | 0.692838 | 0.768 | 0.45 | 5.83E-09 | 3 M1 _ | Capn2 |
| Dync1i21 | 2.73E-13 | 0.642934 | 0.915 | 0.782 | 5.50E-09 | 3 M1 _ | Dync1i2 |
| Pls3 | 2.70E-13 | 0.834723 | 0.854 | 0.586 | 5.43E-09 | 3 M1 _ | Pls3 |
| Ptpn14 | 2.57E-13 | 0.499513 | 0.378 | 0.117 | 5.17E-09 | 3 M1 _ | Ptpn14 |
| Crybg3 | 2.17E-13 | 0.637652 | 0.585 | 0.252 | 4.38E-09 | 3 M1 _ | Crybg3 |
| Lipa | 2.17E-13 | 0.865056 | 0.768 | 0.481 | 4.37E-09 | 3 M1 _ | Lipa |
| Purb | 2.04E-13 | 0.751044 | 0.902 | 0.668 | 4.11E-09 | 3 M1 _ | Purb |
| Gm424182 | 1.72E-13 | 0.943592 | 1 | 0.999 | 3.46E-09 | 3 M1 _ | Gm42418 |
| Wwc2 | 1.64E-13 | 0.68531 | 0.768 | 0.458 | 3.31E-09 | 3 M1 _ | Wwc2 |
| Ints9 | 1.42E-13 | 0.411823 | 0.402 | 0.125 | 2.86E-09 | 3 M1 _ | Ints9 |
| Fam212b1 | 1.31E-13 | 0.259148 | 0.146 | 0.019 | 2.64E-09 | 3 M1 _ | Fam212b |
| Amotl2 | 1.18E-13 | 0.6155 | 0.537 | 0.217 | 2.38E-09 | 3 M1 _ | Amotl2 |
| Actg1 | 1.08E-13 | 0.680657 | 0.988 | 0.944 | 2.17E-09 | 3 M1 _ | Actg1 |
| Tmem140 | 1.06E-13 | 0.693134 | 0.756 | 0.406 | 2.13E-09 | 3 M1 _ | Tmem140 |
| Lmo7 | 1.01E-13 | 0.404459 | 0.256 | 0.054 | 2.03E-09 | 3 M1 _ | Lmo7 |
| Oaz21 | 9.33E-14 | 0.874412 | 0.732 | 0.437 | 1.88E-09 | 3 M1 _ | Oaz2 |
| Tmsb4x1 | 9.23E-14 | 0.529063 | 1 | 0.994 | 1.86E-09 | 3 M1 _ | Tmsb4x |
| Nov | 7.66E-14 | 0.309448 | 0.11 | 0.011 | 1.54E-09 | 3 M1 _ | Nov |
| Diaph2 | 7.56E-14 | 0.868089 | 0.695 | 0.394 | 1.52E-09 | 3 M1 _ | Diaph2 |
| Macf1 | 7.00E-14 | 0.775673 | 0.939 | 0.807 | 1.41E-09 | 3 M1 _ | Macf1 |
| Hoxd3os1 | 6.91E-14 | 0.363879 | 0.232 | 0.045 | 1.39E-09 | 3 M1 _ | Hoxd3os1 |
| Ankrd33b | 6.49E-14 | 0.291665 | 0.232 | 0.044 | 1.31E-09 | 3 M1 _ | Ankrd33b |
| Hpcal1 | 6.29E-14 | 0.6901 | 0.707 | 0.354 | 1.27E-09 | 3 M1 _ | Hpcal1 |
| Ncoa3 | 6.22E-14 | 0.748167 | 0.634 | 0.307 | 1.25E-09 | 3 M1 _ | Ncoa3 |
| Lmcd1 | 5.89E-14 | 0.391111 | 0.512 | 0.175 | 1.19E-09 | 3 M1 _ | Lmcd1 |
| Nr4a1 | 4.95E-14 | 0.404237 | 0.293 | 0.068 | 9.97E-10 | 3 M1 _ | Nr4a1 |

| | | | | | | | |
|---|---|---|---|---|---|---|---|
| Cdc42ep1 | 4.78E-14 | 0.647377 | 0.61 | 0.258 | 9.63E-10 | 3 M1 _ | Cdc42ep1 |
| Stom1 | 4.33E-14 | 0.511136 | 0.476 | 0.157 | 8.73E-10 | 3 M1 _ | Stom |
| Cyp26b11 | 2.41E-14 | 0.57901 | 0.366 | 0.102 | 4.86E-10 | 3 M1 _ | Cyp26b1 |
| Vwa1 | 2.21E-14 | 0.913421 | 0.659 | 0.327 | 4.45E-10 | 3 M1 _ | Vwa1 |
| Etv5 | 2.21E-14 | 0.574794 | 0.451 | 0.157 | 4.44E-10 | 3 M1 _ | Etv5 |
| Zufsp3 | 1.92E-14 | 0.271948 | 0.195 | 0.031 | 3.86E-10 | 3 M1 _ | Zufsp |
| Zfpm1 | 1.79E-14 | 0.609255 | 0.488 | 0.175 | 3.60E-10 | 3 M1 _ | Zfpm1 |
| 9530082P21Rik | 1.69E-14 | 0.480523 | 0.366 | 0.101 | 3.40E-10 | 3 M1 _ | 9530082P21Rik |
| Pitpnc1 | 1.58E-14 | 0.702464 | 0.72 | 0.362 | 3.19E-10 | 3 M1 _ | Pitpnc1 |
| Sec14l1 | 1.49E-14 | 0.676928 | 0.683 | 0.336 | 3.00E-10 | 3 M1 _ | Sec14l1 |
| Xiap | 1.41E-14 | 0.830366 | 0.817 | 0.517 | 2.84E-10 | 3 M1 _ | Xiap |
| Cyb5r3 | 1.23E-14 | 0.681667 | 0.866 | 0.573 | 2.48E-10 | 3 M1 _ | Cyb5r3 |
| Pqlc1 | 1.21E-14 | 0.589676 | 0.524 | 0.197 | 2.43E-10 | 3 M1 _ | Pqlc1 |
| Fkbp3 | 1.16E-14 | 0.905507 | 0.805 | 0.589 | 2.33E-10 | 3 M1 _ | Fkbp3 |
| Jdp2 | 1.04E-14 | 0.606421 | 0.585 | 0.237 | 2.09E-10 | 3 M1 _ | Jdp2 |
| Fbxw2 | 9.23E-15 | 0.728407 | 0.78 | 0.425 | 1.86E-10 | 3 M1 _ | Fbxw2 |
| Sypl | 8.87E-15 | 0.700558 | 0.817 | 0.46 | 1.79E-10 | 3 M1 _ | Sypl |
| Tjp1 | 7.53E-15 | 0.695581 | 0.744 | 0.386 | 1.52E-10 | 3 M1 _ | Tjp1 |
| Nes1 | 6.60E-15 | 0.762913 | 0.573 | 0.22 | 1.33E-10 | 3 M1 _ | Nes |
| Rhoa | 6.22E-15 | 0.587989 | 0.963 | 0.901 | 1.25E-10 | 3 M1 _ | Rhoa |
| Rassf9 | 6.04E-15 | 0.677514 | 0.39 | 0.11 | 1.22E-10 | 3 M1 _ | Rassf9 |
| Fam171a1 | 5.67E-15 | 0.627629 | 0.463 | 0.151 | 1.14E-10 | 3 M1 _ | Fam171a1 |
| Apbb2 | 5.39E-15 | 0.743863 | 0.805 | 0.461 | 1.09E-10 | 3 M1 _ | Apbb2 |
| Papss2 | 5.03E-15 | 0.499724 | 0.366 | 0.102 | 1.01E-10 | 3 M1 _ | Papss2 |
| Fmo2 | 4.97E-15 | 0.979778 | 0.793 | 0.455 | 1.00E-10 | 3 M1 _ | Fmo2 |
| Tspan14 | 4.56E-15 | 0.318871 | 0.366 | 0.093 | 9.19E-11 | 3 M1 _ | Tspan14 |
| Plekhg1 | 4.11E-15 | 0.542496 | 0.5 | 0.176 | 8.28E-11 | 3 M1 _ | Plekhg1 |
| Dcbld1 | 4.05E-15 | 0.388714 | 0.305 | 0.068 | 8.17E-11 | 3 M1 _ | Dcbld1 |
| Acvr2a1 | 3.91E-15 | 0.606843 | 0.488 | 0.164 | 7.87E-11 | 3 M1 _ | Acvr2a |
| Cds21 | 3.36E-15 | 0.758087 | 0.659 | 0.293 | 6.77E-11 | 3 M1 _ | Cds2 |
| Glul | 3.17E-15 | 1.349189 | 0.61 | 0.288 | 6.39E-11 | 3 M1 _ | Glul |
| Gimap1 | 2.87E-15 | 0.536632 | 0.683 | 0.258 | 5.79E-11 | 3 M1 _ | Gimap1 |
| Trib21 | 2.59E-15 | 0.889281 | 0.695 | 0.343 | 5.21E-11 | 3 M1 _ | Trib2 |
| Rab11a | 2.33E-15 | 0.748311 | 0.756 | 0.433 | 4.70E-11 | 3 M1 _ | Rab11a |
| Arl4d | 2.31E-15 | 0.453303 | 0.244 | 0.046 | 4.65E-11 | 3 M1 _ | Arl4d |
| Eif4ebp21 | 2.30E-15 | 0.851998 | 0.793 | 0.48 | 4.63E-11 | 3 M1 _ | Eif4ebp2 |
| Cavin11 | 2.27E-15 | 0.870422 | 0.793 | 0.473 | 4.57E-11 | 3 M1 _ | Cavin1 |
| Fnip2 | 1.94E-15 | 0.456102 | 0.354 | 0.091 | 3.90E-11 | 3 M1 _ | Fnip2 |
| Zdhhc18 | 1.93E-15 | 0.555464 | 0.549 | 0.199 | 3.88E-11 | 3 M1 _ | Zdhhc18 |
| Pde8a | 1.87E-15 | 0.546568 | 0.537 | 0.191 | 3.76E-11 | 3 M1 _ | Pde8a |
| Mlec1 | 1.69E-15 | 0.893517 | 0.841 | 0.528 | 3.41E-11 | 3 M1 _ | Mlec |

| | | | | | | | |
|---|---|---|---|---|---|---|---|
| Limch1 | 1.65E-15 | 0.479411 | 0.329 | 0.077 | 3.33E-11 | 3 M1 _ | Limch1 |
| Sorbs3 | 1.65E-15 | 0.609051 | 0.463 | 0.156 | 3.32E-11 | 3 M1 _ | Sorbs3 |
| Numb | 1.63E-15 | 0.686802 | 0.61 | 0.258 | 3.29E-11 | 3 M1 _ | Numb |
| Tcf41 | 1.58E-15 | 0.741978 | 1 | 0.885 | 3.17E-11 | 3 M1 _ | Tcf4 |
| Tubb4b1 | 1.47E-15 | 0.911767 | 0.915 | 0.695 | 2.95E-11 | 3 M1 _ | Tubb4b |
| Itsn2 | 1.43E-15 | 0.713706 | 0.671 | 0.304 | 2.89E-11 | 3 M1 _ | Itsn2 |
| Lpar6 | 1.23E-15 | 0.713708 | 0.524 | 0.184 | 2.49E-11 | 3 M1 _ | Lpar6 |
| Colgalt2 | 1.18E-15 | 0.545113 | 0.402 | 0.114 | 2.38E-11 | 3 M1 _ | Colgalt2 |
| Gimap6 | 1.14E-15 | 0.682397 | 0.817 | 0.391 | 2.30E-11 | 3 M1 _ | Gimap6 |
| Ipo11 | 1.10E-15 | 0.544778 | 0.415 | 0.119 | 2.21E-11 | 3 M1 _ | Ipo11 |
| Tsc22d31 | 1.05E-15 | 0.80086 | 0.817 | 0.467 | 2.12E-11 | 3 M1 _ | Tsc22d3 |
| Lfng | 1.01E-15 | 0.82578 | 0.5 | 0.179 | 2.04E-11 | 3 M1 _ | Lfng |
| AU021092 | 1.01E-15 | 0.289877 | 0.159 | 0.02 | 2.03E-11 | 3 M1 _ | AU021092 |
| Prdm1 | 1.01E-15 | 0.498114 | 0.354 | 0.087 | 2.03E-11 | 3 M1 _ | Prdm1 |
| Klf12 | 9.99E-16 | 0.396534 | 0.341 | 0.083 | 2.01E-11 | 3 M1 _ | Klf12 |
| Epha4 | 9.72E-16 | 0.698276 | 0.512 | 0.184 | 1.96E-11 | 3 M1 _ | Epha4 |
| Ptp4a13 | 8.20E-16 | 0.301906 | 0.305 | 0.065 | 1.65E-11 | 3 M1 _ | Ptp4a1 |
| Rnf125 | 7.17E-16 | 0.400406 | 0.317 | 0.072 | 1.44E-11 | 3 M1 _ | Rnf125 |
| Itga11 | 6.55E-16 | 0.730884 | 0.976 | 0.794 | 1.32E-11 | 3 M1 _ | Itga1 |
| Gimap4 | 6.16E-16 | 0.548559 | 0.744 | 0.291 | 1.24E-11 | 3 M1 _ | Gimap4 |
| Ctnnb1 | 5.01E-16 | 0.814351 | 0.878 | 0.696 | 1.01E-11 | 3 M1 _ | Ctnnb1 |
| Tmem204 | 4.92E-16 | 0.806114 | 0.902 | 0.569 | 9.92E-12 | 3 M1 _ | Tmem204 |
| Atoh8 | 4.88E-16 | 0.38237 | 0.195 | 0.029 | 9.82E-12 | 3 M1 _ | Atoh8 |
| Ostf11 | 4.15E-16 | 0.77471 | 0.878 | 0.602 | 8.35E-12 | 3 M1 _ | Ostf1 |
| Gnaq1 | 3.90E-16 | 0.739489 | 0.78 | 0.434 | 7.85E-12 | 3 M1 _ | Gnaq |
| Adam10 | 3.81E-16 | 0.719547 | 0.695 | 0.311 | 7.68E-12 | 3 M1 _ | Adam10 |
| Ehd21 | 3.22E-16 | 0.973922 | 0.768 | 0.477 | 6.49E-12 | 3 M1 _ | Ehd2 |
| Upp1 | 3.20E-16 | 0.44814 | 0.317 | 0.069 | 6.44E-12 | 3 M1 _ | Upp1 |
| Ica1 | 2.83E-16 | 0.431859 | 0.268 | 0.052 | 5.70E-12 | 3 M1 _ | Ica1 |
| Tpst2 | 2.74E-16 | 0.572767 | 0.695 | 0.292 | 5.52E-12 | 3 M1 _ | Tpst2 |
| Rhob2 | 2.48E-16 | 1.147425 | 0.854 | 0.609 | 4.99E-12 | 3 M1 _ | Rhob |
| Tgm2 | 2.36E-16 | 0.835224 | 0.671 | 0.297 | 4.75E-12 | 3 M1 _ | Tgm2 |
| Cd471 | 2.33E-16 | 0.737883 | 0.976 | 0.859 | 4.68E-12 | 3 M1 _ | Cd47 |
| Slc28a2 | 2.30E-16 | 0.459565 | 0.256 | 0.047 | 4.63E-12 | 3 M1 _ | Slc28a2 |
| Endod1 | 2.22E-16 | 0.840297 | 0.72 | 0.368 | 4.47E-12 | 3 M1 _ | Endod1 |
| Snai2 | 2.21E-16 | 0.646825 | 0.561 | 0.205 | 4.45E-12 | 3 M1 _ | Snai2 |
| Col4a21 | 2.16E-16 | 0.819182 | 0.988 | 0.852 | 4.36E-12 | 3 M1 _ | Col4a2 |
| Bnip2 | 2.05E-16 | 0.861815 | 0.854 | 0.58 | 4.13E-12 | 3 M1 _ | Bnip2 |
| Tmem44 | 2.04E-16 | 0.315688 | 0.244 | 0.042 | 4.11E-12 | 3 M1 _ | Tmem44 |
| Rap1a1 | 2.04E-16 | 0.78853 | 0.89 | 0.645 | 4.11E-12 | 3 M1 _ | Rap1a |
| Notch1 | 1.88E-16 | 0.940114 | 0.683 | 0.337 | 3.79E-12 | 3 M1 _ | Notch1 |

| | | | | | | | |
|---|---|---|---|---|---|---|---|
| St3gal6 | 1.85E-16 | 0.703155 | 0.61 | 0.22 | 3.73E-12 | 3 M1 _ | St3gal6 |
| Dab2ip1 | 1.81E-16 | 0.816545 | 0.829 | 0.485 | 3.65E-12 | 3 M1 _ | Dab2ip |
| Slc30a1 | 1.63E-16 | 0.524911 | 0.415 | 0.114 | 3.29E-12 | 3 M1 _ | Slc30a1 |
| Slfn3 | 1.58E-16 | 0.413704 | 0.171 | 0.022 | 3.18E-12 | 3 M1 _ | Slfn3 |
| Tnfrsf10b | 1.51E-16 | 0.312935 | 0.256 | 0.047 | 3.04E-12 | 3 M1 _ | Tnfrsf10b |
| Map4k3 | 1.36E-16 | 0.629213 | 0.585 | 0.219 | 2.73E-12 | 3 M1 _ | Map4k3 |
| Myo6 | 1.30E-16 | 0.838941 | 0.72 | 0.343 | 2.61E-12 | 3 M1 _ | Myo6 |
| Ets2 | 1.24E-16 | 0.588527 | 0.646 | 0.247 | 2.50E-12 | 3 M1 _ | Ets2 |
| Cd1511 | 1.18E-16 | 0.706882 | 0.866 | 0.55 | 2.38E-12 | 3 M1 _ | Cd151 |
| Sncaip | 1.13E-16 | 0.271868 | 0.207 | 0.031 | 2.28E-12 | 3 M1 _ | Sncaip |
| Nxpe2 | 1.11E-16 | 0.487998 | 0.22 | 0.035 | 2.24E-12 | 3 M1 _ | Nxpe2 |
| Kdelc22 | 1.03E-16 | 0.385042 | 0.317 | 0.066 | 2.07E-12 | 3 M1 _ | Kdelc2 |
| Pcdh12 | 6.12E-17 | 0.565478 | 0.488 | 0.152 | 1.23E-12 | 3 M1 _ | Pcdh12 |
| Adamts11 | 5.75E-17 | 1.077478 | 0.707 | 0.33 | 1.16E-12 | 3 M1 _ | Adamts1 |
| Ptma | 5.07E-17 | 0.610598 | 1 | 0.986 | 1.02E-12 | 3 M1 _ | Ptma |
| Gimap8 | 5.01E-17 | 0.598009 | 0.439 | 0.119 | 1.01E-12 | 3 M1 _ | Gimap8 |
| Cdipt | 4.65E-17 | 0.750422 | 0.671 | 0.291 | 9.36E-13 | 3 M1 _ | Cdipt |
| Ldb2 | 4.63E-17 | 0.902037 | 0.732 | 0.37 | 9.32E-13 | 3 M1 _ | Ldb2 |
| Kif5b | 4.25E-17 | 0.85518 | 0.878 | 0.674 | 8.57E-13 | 3 M1 _ | Kif5b |
| Sema6b | 3.43E-17 | 0.386056 | 0.256 | 0.046 | 6.91E-13 | 3 M1 _ | Sema6b |
| Dhrs31 | 2.79E-17 | 0.841118 | 0.817 | 0.407 | 5.61E-13 | 3 M1 _ | Dhrs3 |
| Pik3c2a | 2.13E-17 | 0.734435 | 0.671 | 0.283 | 4.29E-13 | 3 M1 _ | Pik3c2a |
| Mlf1 | 1.90E-17 | 0.358204 | 0.22 | 0.033 | 3.83E-13 | 3 M1 _ | Mlf1 |
| Rps6ka3 | 1.83E-17 | 0.822195 | 0.671 | 0.292 | 3.69E-13 | 3 M1 _ | Rps6ka3 |
| Sod11 | 1.28E-17 | 0.957422 | 0.829 | 0.524 | 2.58E-13 | 3 M1 _ | Sod1 |
| Elk3 | 1.18E-17 | 1.009282 | 0.817 | 0.488 | 2.37E-13 | 3 M1 _ | Elk3 |
| Tcf7l1 | 1.16E-17 | 0.82666 | 0.671 | 0.287 | 2.34E-13 | 3 M1 _ | Tcf7l1 |
| 9930111J21Rik21 | 1.05E-17 | 0.817481 | 0.841 | 0.428 | 2.12E-13 | 3 M1 _ | 9930111J21Rik2 |
| Afdn | 8.97E-18 | 1.092136 | 0.744 | 0.398 | 1.81E-13 | 3 M1 _ | Afdn |
| Lpcat1 | 8.30E-18 | 0.701691 | 0.561 | 0.193 | 1.67E-13 | 3 M1 _ | Lpcat1 |
| Bcl2l1 | 8.21E-18 | 0.704193 | 0.537 | 0.183 | 1.65E-13 | 3 M1 _ | Bcl2l1 |
| Cpd1 | 8.13E-18 | 0.939305 | 0.793 | 0.473 | 1.64E-13 | 3 M1 _ | Cpd |
| Sh2d3c | 7.59E-18 | 0.620447 | 0.476 | 0.14 | 1.53E-13 | 3 M1 _ | Sh2d3c |
| Ndufc2 | 7.02E-18 | 0.924088 | 0.951 | 0.703 | 1.41E-13 | 3 M1 _ | Ndufc2 |
| Rai14 | 6.61E-18 | 0.894644 | 0.72 | 0.345 | 1.33E-13 | 3 M1 _ | Rai14 |
| Crip21 | 6.26E-18 | 0.837985 | 0.951 | 0.765 | 1.26E-13 | 3 M1 _ | Crip2 |
| Cyb5r1 | 5.62E-18 | 0.526424 | 0.415 | 0.106 | 1.13E-13 | 3 M1 _ | Cyb5r1 |
| Col4a12 | 4.72E-18 | 0.872841 | 1 | 0.869 | 9.51E-14 | 3 M1 _ | Col4a1 |
| Anp32a | 4.68E-18 | 1.01488 | 0.878 | 0.657 | 9.42E-14 | 3 M1 _ | Anp32a |
| Gbp4 | 4.56E-18 | 0.629366 | 0.549 | 0.176 | 9.18E-14 | 3 M1 _ | Gbp4 |
| Fam117b | 4.32E-18 | 0.801797 | 0.585 | 0.222 | 8.71E-14 | 3 M1 _ | Fam117b |

| | | | | | | | |
|---|---|---|---|---|---|---|---|
| Luzp1 | 4.28E-18 | 0.856862 | 0.878 | 0.59 | 8.61E-14 | 3 M1 _ | Luzp1 |
| Iqgap1 | 3.90E-18 | 0.92114 | 0.951 | 0.751 | 7.85E-14 | 3 M1 _ | Iqgap1 |
| Pdgfd | 3.31E-18 | 0.540773 | 0.415 | 0.106 | 6.66E-14 | 3 M1 _ | Pdgfd |
| Ctnnbip1 | 3.26E-18 | 0.786071 | 0.659 | 0.269 | 6.57E-14 | 3 M1 _ | Ctnnbip1 |
| Tspan18 | 2.20E-18 | 0.512123 | 0.402 | 0.101 | 4.43E-14 | 3 M1 _ | Tspan18 |
| Lats2 | 2.10E-18 | 0.834137 | 0.707 | 0.324 | 4.24E-14 | 3 M1 _ | Lats2 |
| Car7 | 1.96E-18 | 0.496752 | 0.22 | 0.032 | 3.95E-14 | 3 M1 _ | Car7 |
| Cmip | 1.78E-18 | 0.822933 | 0.634 | 0.255 | 3.58E-14 | 3 M1 _ | Cmip |
| Tpm41 | 1.53E-18 | 1.017727 | 0.902 | 0.751 | 3.07E-14 | 3 M1 _ | Tpm4 |
| Ifi44 | 1.50E-18 | 0.680734 | 0.549 | 0.166 | 3.02E-14 | 3 M1 _ | Ifi44 |
| Cebpg | 1.32E-18 | 0.837788 | 0.768 | 0.393 | 2.66E-14 | 3 M1 _ | Cebpg |
| Tmtc2 | 1.18E-18 | 0.336355 | 0.22 | 0.031 | 2.38E-14 | 3 M1 _ | Tmtc2 |
| 2900026A02Rik | 9.55E-19 | 0.789148 | 0.707 | 0.291 | 1.92E-14 | 3 M1 _ | 2900026A02Rik |
| Gcnt21 | 9.45E-19 | 0.558847 | 0.622 | 0.191 | 1.90E-14 | 3 M1 _ | Gcnt2 |
| 3110062M04Rik2 | 8.96E-19 | 0.295227 | 0.22 | 0.031 | 1.80E-14 | 3 M1 _ | 3110062M04Rik |
| Irx3 | 8.09E-19 | 0.436856 | 0.305 | 0.057 | 1.63E-14 | 3 M1 _ | Irx3 |
| Thsd1 | 6.32E-19 | 0.575131 | 0.305 | 0.058 | 1.27E-14 | 3 M1 _ | Thsd1 |
| Wars | 4.52E-19 | 0.781175 | 0.549 | 0.186 | 9.10E-15 | 3 M1 _ | Wars |
| Piezo2 | 4.12E-19 | 0.967147 | 0.744 | 0.346 | 8.29E-15 | 3 M1 _ | Piezo2 |
| Efna1 | 3.00E-19 | 0.543818 | 0.463 | 0.118 | 6.04E-15 | 3 M1 _ | Efna1 |
| Sox7 | 2.25E-19 | 0.40638 | 0.256 | 0.04 | 4.54E-15 | 3 M1 _ | Sox7 |
| Rtl8a | 2.08E-19 | 1.213659 | 0.793 | 0.477 | 4.19E-15 | 3 M1 _ | Rtl8a |
| Csrnp1 | 1.27E-19 | 0.386753 | 0.256 | 0.041 | 2.56E-15 | 3 M1 _ | Csrnp1 |
| Qk | 1.18E-19 | 0.930914 | 0.902 | 0.675 | 2.38E-15 | 3 M1 _ | Qk |
| AY0361182 | 1.01E-19 | 1.685148 | 0.89 | 0.555 | 2.03E-15 | 3 M1 _ | AY036118 |
| Adgrl2 | 9.84E-20 | 0.886933 | 0.683 | 0.27 | 1.98E-15 | 3 M1 _ | Adgrl2 |
| Ier3 | 9.62E-20 | 0.969444 | 0.512 | 0.152 | 1.94E-15 | 3 M1 _ | Ier3 |
| Utrn1 | 9.03E-20 | 1.061699 | 0.854 | 0.478 | 1.82E-15 | 3 M1 _ | Utrn |
| Phactr4 | 8.63E-20 | 0.82145 | 0.622 | 0.23 | 1.74E-15 | 3 M1 _ | Phactr4 |
| Nck1 | 8.53E-20 | 1.06806 | 0.829 | 0.513 | 1.72E-15 | 3 M1 _ | Nck1 |
| Ndufa8 | 7.37E-20 | 0.947702 | 0.89 | 0.594 | 1.48E-15 | 3 M1 _ | Ndufa8 |
| Spag72 | 5.93E-20 | 0.628729 | 0.561 | 0.17 | 1.19E-15 | 3 M1 _ | Spag7 |
| Eepd1 | 5.87E-20 | 0.42185 | 0.256 | 0.039 | 1.18E-15 | 3 M1 _ | Eepd1 |
| Stc1 | 5.32E-20 | 0.660552 | 0.5 | 0.133 | 1.07E-15 | 3 M1 _ | Stc1 |
| Klf7 | 4.64E-20 | 1.081865 | 0.866 | 0.539 | 9.34E-16 | 3 M1 _ | Klf7 |
| Jam31 | 4.35E-20 | 1.068704 | 0.78 | 0.393 | 8.77E-16 | 3 M1 _ | Jam3 |
| Spata13 | 4.21E-20 | 0.548003 | 0.415 | 0.098 | 8.48E-16 | 3 M1 _ | Spata13 |
| Clu | 3.63E-20 | 0.846916 | 0.463 | 0.122 | 7.31E-16 | 3 M1 _ | Clu |
| Igfbp7 | 3.54E-20 | 0.833601 | 0.988 | 0.943 | 7.13E-16 | 3 M1 _ | Igfbp7 |
| Hspa12b | 3.37E-20 | 0.40735 | 0.354 | 0.072 | 6.79E-16 | 3 M1 _ | Hspa12b |
| Jcad | 3.37E-20 | 0.759265 | 0.622 | 0.222 | 6.79E-16 | 3 M1 _ | Jcad |

| | | | | | | | |
|---|---|---|---|---|---|---|---|
| Rgs12 | 2.96E-20 | 0.596263 | 0.476 | 0.132 | 5.96E-16 | 3 M1 _ | Rgs12 |
| App1 | 2.91E-20 | 0.906957 | 0.988 | 0.835 | 5.87E-16 | 3 M1 _ | App |
| Dgkh | 2.71E-20 | 0.744953 | 0.622 | 0.211 | 5.45E-16 | 3 M1 _ | Dgkh |
| Fhod1 | 2.66E-20 | 0.504957 | 0.317 | 0.058 | 5.36E-16 | 3 M1 _ | Fhod1 |
| St8sia4 | 2.31E-20 | 0.774759 | 0.634 | 0.208 | 4.66E-16 | 3 M1 _ | St8sia4 |
| Lama5 | 2.22E-20 | 0.376613 | 0.329 | 0.061 | 4.48E-16 | 3 M1 _ | Lama5 |
| Arhgef12 | 2.01E-20 | 1.06404 | 0.878 | 0.663 | 4.04E-16 | 3 M1 _ | Arhgef12 |
| Mertk | 1.55E-20 | 0.65916 | 0.5 | 0.137 | 3.13E-16 | 3 M1 _ | Mertk |
| Plxnd1 | 1.23E-20 | 1.063538 | 0.78 | 0.369 | 2.47E-16 | 3 M1 _ | Plxnd1 |
| Unc119b | 1.19E-20 | 0.947023 | 0.61 | 0.227 | 2.39E-16 | 3 M1 _ | Unc119b |
| Lars22 | 1.12E-20 | 1.244504 | 0.976 | 0.808 | 2.26E-16 | 3 M1 _ | Lars2 |
| Slc12a7 | 9.47E-21 | 0.416506 | 0.354 | 0.07 | 1.91E-16 | 3 M1 _ | Slc12a7 |
| Smad7 | 9.26E-21 | 1.007337 | 0.829 | 0.398 | 1.87E-16 | 3 M1 _ | Smad7 |
| Abcb1a | 8.15E-21 | 0.677033 | 0.415 | 0.096 | 1.64E-16 | 3 M1 _ | Abcb1a |
| Syn31 | 8.14E-21 | 0.689872 | 0.427 | 0.1 | 1.64E-16 | 3 M1 _ | Syn3 |
| Nav2 | 7.42E-21 | 0.575256 | 0.354 | 0.071 | 1.49E-16 | 3 M1 _ | Nav2 |
| BC028528 | 7.35E-21 | 0.762964 | 0.598 | 0.189 | 1.48E-16 | 3 M1 _ | BC028528 |
| Gnb1 | 7.01E-21 | 0.871239 | 0.927 | 0.686 | 1.41E-16 | 3 M1 _ | Gnb1 |
| Ppm1f | 6.67E-21 | 0.52477 | 0.378 | 0.083 | 1.34E-16 | 3 M1 _ | Ppm1f |
| Epb411 | 4.25E-21 | 0.557511 | 0.573 | 0.165 | 8.55E-17 | 3 M1 _ | Epb41 |
| Plcg1 | 3.53E-21 | 0.693606 | 0.671 | 0.243 | 7.12E-17 | 3 M1 _ | Plcg1 |
| Ripor1 | 2.18E-21 | 0.869106 | 0.671 | 0.259 | 4.39E-17 | 3 M1 _ | Ripor1 |
| Scarf1 | 2.13E-21 | 0.415114 | 0.305 | 0.051 | 4.30E-17 | 3 M1 _ | Scarf1 |
| Fam69a2 | 1.87E-21 | 0.295794 | 0.329 | 0.057 | 3.77E-17 | 3 M1 _ | Fam69a |
| Mapk3 | 1.26E-21 | 0.937121 | 0.902 | 0.537 | 2.53E-17 | 3 M1 _ | Mapk3 |
| Pparg1 | 1.08E-21 | 0.341735 | 0.293 | 0.045 | 2.18E-17 | 3 M1 _ | Pparg |
| Rnf144b | 1.08E-21 | 0.280979 | 0.195 | 0.021 | 2.17E-17 | 3 M1 _ | Rnf144b |
| Fut4 | 9.52E-22 | 0.53091 | 0.28 | 0.044 | 1.92E-17 | 3 M1 _ | Fut4 |
| Tmem55b2 | 8.15E-22 | 0.322928 | 0.28 | 0.043 | 1.64E-17 | 3 M1 _ | Tmem55b |
| Ccnyl1 | 7.30E-22 | 0.577622 | 0.366 | 0.074 | 1.47E-17 | 3 M1 _ | Ccnyl1 |
| Map3k11 | 5.62E-22 | 0.557824 | 0.427 | 0.098 | 1.13E-17 | 3 M1 _ | Map3k11 |
| Myo1c | 5.00E-22 | 0.98708 | 0.805 | 0.403 | 1.01E-17 | 3 M1 _ | Myo1c |
| Gch1 | 4.78E-22 | 0.456377 | 0.354 | 0.067 | 9.62E-18 | 3 M1 _ | Gch1 |
| Smagp1 | 4.66E-22 | 0.641181 | 0.537 | 0.139 | 9.39E-18 | 3 M1 _ | Smagp |
| Hspg21 | 4.26E-22 | 1.1064 | 0.963 | 0.69 | 8.59E-18 | 3 M1 _ | Hspg2 |
| Chst2 | 3.93E-22 | 0.894247 | 0.549 | 0.162 | 7.91E-18 | 3 M1 _ | Chst2 |
| Chtf82 | 3.22E-22 | 0.60298 | 0.39 | 0.082 | 6.48E-18 | 3 M1 _ | Chtf8 |
| Sez6l2 | 2.95E-22 | 0.380694 | 0.183 | 0.018 | 5.94E-18 | 3 M1 _ | Sez6l2 |
| mt-Atp82 | 2.95E-22 | 1.03947 | 0.915 | 0.596 | 5.94E-18 | 3 M1 _ | mt-Atp8 |
| Nfkbia | 2.82E-22 | 1.204716 | 0.866 | 0.531 | 5.68E-18 | 3 M1 _ | Nfkbia |
| Sema3g | 2.82E-22 | 0.696666 | 0.256 | 0.035 | 5.68E-18 | 3 M1 _ | Sema3g |

| | | | | | | | |
|---|---|---|---|---|---|---|---|
| Acacb | 2.23E-22 | 0.384636 | 0.256 | 0.035 | 4.50E-18 | 3 M1 _ | Acacb |
| Tspan9 | 2.16E-22 | 0.948883 | 0.695 | 0.283 | 4.35E-18 | 3 M1 _ | Tspan9 |
| Trpv4 | 1.95E-22 | 0.286709 | 0.22 | 0.026 | 3.92E-18 | 3 M1 _ | Trpv4 |
| Fas | 1.16E-22 | 0.715284 | 0.549 | 0.158 | 2.33E-18 | 3 M1 _ | Fas |
| Dnm3 | 1.03E-22 | 0.859875 | 0.683 | 0.25 | 2.07E-18 | 3 M1 _ | Dnm3 |
| Skiv2l22 | 9.04E-23 | 0.476763 | 0.317 | 0.052 | 1.82E-18 | 3 M1 _ | Skiv2l2 |
| Lmo2 | 6.39E-23 | 0.732069 | 0.537 | 0.148 | 1.29E-18 | 3 M1 _ | Lmo2 |
| Uaca | 5.87E-23 | 1.043719 | 0.707 | 0.262 | 1.18E-18 | 3 M1 _ | Uaca |
| 2810403A07Rik | 5.26E-23 | 0.274478 | 0.22 | 0.025 | 1.06E-18 | 3 M1 _ | 2810403A07Rik |
| Tmcc3 | 3.92E-23 | 0.779584 | 0.549 | 0.154 | 7.90E-19 | 3 M1 _ | Tmcc3 |
| C2cd21 | 3.22E-23 | 0.652463 | 0.622 | 0.187 | 6.48E-19 | 3 M1 _ | C2cd2 |
| Ephb1 | 2.38E-23 | 0.315927 | 0.22 | 0.024 | 4.80E-19 | 3 M1 _ | Ephb1 |
| Ets1 | 1.56E-23 | 1.092875 | 0.951 | 0.597 | 3.14E-19 | 3 M1 _ | Ets1 |
| C1qtnf9 | 1.41E-23 | 0.366529 | 0.146 | 0.01 | 2.84E-19 | 3 M1 _ | C1qtnf9 |
| Tcf15 | 1.21E-23 | 0.771219 | 0.488 | 0.111 | 2.43E-19 | 3 M1 _ | Tcf15 |
| Tal1 | 1.09E-23 | 0.258428 | 0.195 | 0.019 | 2.19E-19 | 3 M1 _ | Tal1 |
| Syne21 | 9.88E-24 | 1.156968 | 0.854 | 0.406 | 1.99E-19 | 3 M1 _ | Syne2 |
| Mall | 9.59E-24 | 0.461323 | 0.305 | 0.046 | 1.93E-19 | 3 M1 _ | Mall |
| Klf13 | 8.74E-24 | 1.046224 | 0.915 | 0.54 | 1.76E-19 | 3 M1 _ | Klf13 |
| Ctnna1 | 8.48E-24 | 1.059239 | 0.951 | 0.662 | 1.71E-19 | 3 M1 _ | Ctnna1 |
| Dennd5b | 6.96E-24 | 0.856211 | 0.646 | 0.213 | 1.40E-19 | 3 M1 _ | Dennd5b |
| Rasal21 | 6.38E-24 | 0.860523 | 0.805 | 0.32 | 1.28E-19 | 3 M1 _ | Rasal2 |
| Dscr3 | 6.12E-24 | 0.28448 | 0.22 | 0.024 | 1.23E-19 | 3 M1 _ | Dscr3 |
| Ltbp41 | 6.08E-24 | 1.350535 | 0.927 | 0.663 | 1.22E-19 | 3 M1 _ | Ltbp4 |
| Plk2 | 5.80E-24 | 0.813711 | 0.512 | 0.126 | 1.17E-19 | 3 M1 _ | Plk2 |
| Efnb1 | 4.59E-24 | 1.079237 | 0.659 | 0.226 | 9.25E-20 | 3 M1 _ | Efnb1 |
| Sema6a | 4.59E-24 | 0.873184 | 0.561 | 0.153 | 9.24E-20 | 3 M1 _ | Sema6a |
| Ly6e | 3.55E-24 | 1.012602 | 1 | 0.874 | 7.15E-20 | 3 M1 _ | Ly6e |
| Gm26669 | 3.39E-24 | 0.281495 | 0.146 | 0.01 | 6.83E-20 | 3 M1 _ | Gm26669 |
| Slc43a31 | 3.21E-24 | 1.301482 | 0.902 | 0.495 | 6.47E-20 | 3 M1 _ | Slc43a3 |
| Tmem47 | 2.76E-24 | 0.98108 | 0.78 | 0.321 | 5.56E-20 | 3 M1 _ | Tmem47 |
| Papd42 | 2.61E-24 | 0.333929 | 0.268 | 0.035 | 5.26E-20 | 3 M1 _ | Papd4 |
| Dopey12 | 2.48E-24 | 0.44969 | 0.305 | 0.046 | 5.00E-20 | 3 M1 _ | Dopey1 |
| Tnfaip11 | 2.43E-24 | 0.990492 | 0.817 | 0.366 | 4.89E-20 | 3 M1 _ | Tnfaip1 |
| Mfhas1 | 2.40E-24 | 0.844209 | 0.61 | 0.189 | 4.83E-20 | 3 M1 _ | Mfhas1 |
| Sox131 | 2.30E-24 | 0.560007 | 0.415 | 0.081 | 4.63E-20 | 3 M1 _ | Sox13 |
| Slfn51 | 2.30E-24 | 1.219896 | 0.939 | 0.709 | 4.62E-20 | 3 M1 _ | Slfn5 |
| Rapgef21 | 2.18E-24 | 1.045395 | 0.646 | 0.214 | 4.39E-20 | 3 M1 _ | Rapgef2 |
| Helz2 | 1.94E-24 | 0.652071 | 0.512 | 0.128 | 3.91E-20 | 3 M1 _ | Helz2 |
| Klhl5 | 1.45E-24 | 0.936907 | 0.659 | 0.227 | 2.92E-20 | 3 M1 _ | Klhl5 |
| Actn41 | 1.43E-24 | 1.125904 | 0.939 | 0.712 | 2.88E-20 | 3 M1 _ | Actn4 |

| | | | | | | | |
|---|---|---|---|---|---|---|---|
| Foxp1 | 1.36E-24 | 1.171668 | 0.902 | 0.544 | 2.73E-20 | 3 M1 _ | Foxp1 |
| Prdm16 | 1.12E-24 | 0.550007 | 0.305 | 0.045 | 2.26E-20 | 3 M1 _ | Prdm16 |
| Ddah1 | 8.55E-25 | 0.434061 | 0.232 | 0.026 | 1.72E-20 | 3 M1 _ | Ddah1 |
| Ackr3 | 5.68E-25 | 0.654032 | 0.5 | 0.113 | 1.14E-20 | 3 M1 _ | Ackr3 |
| Ssu2 | 5.10E-25 | 0.40663 | 0.28 | 0.037 | 1.03E-20 | 3 M1 _ | Ssu2 |
| Dopey21 | 4.13E-25 | 0.320562 | 0.195 | 0.018 | 8.31E-21 | 3 M1 _ | Dopey2 |
| Efhd1 | 3.62E-25 | 0.716073 | 0.268 | 0.035 | 7.29E-21 | 3 M1 _ | Efhd1 |
| Aplp2 | 3.35E-25 | 1.155317 | 0.939 | 0.696 | 6.74E-21 | 3 M1 _ | Aplp2 |
| Cdc42bpb | 2.99E-25 | 0.892887 | 0.756 | 0.296 | 6.01E-21 | 3 M1 _ | Cdc42bpb |
| Filip11 | 2.86E-25 | 0.734914 | 0.634 | 0.179 | 5.76E-21 | 3 M1 _ | Filip1 |
| Alpl | 2.57E-25 | 0.763537 | 0.378 | 0.067 | 5.17E-21 | 3 M1 _ | Alpl |
| Stard81 | 2.24E-25 | 0.871651 | 0.671 | 0.199 | 4.52E-21 | 3 M1 _ | Stard8 |
| Trib1 | 1.93E-25 | 0.696375 | 0.5 | 0.117 | 3.89E-21 | 3 M1 _ | Trib1 |
| Dgke | 1.83E-25 | 0.388554 | 0.329 | 0.05 | 3.68E-21 | 3 M1 _ | Dgke |
| Pcdh11 | 1.81E-25 | 0.732183 | 0.61 | 0.165 | 3.65E-21 | 3 M1 _ | Pcdh1 |
| Als2cl | 1.59E-25 | 0.495383 | 0.329 | 0.051 | 3.20E-21 | 3 M1 _ | Als2cl |
| Id3 | 1.57E-25 | 1.415778 | 0.963 | 0.754 | 3.15E-21 | 3 M1 _ | Id3 |
| Rps6ka2 | 1.47E-25 | 0.466379 | 0.305 | 0.044 | 2.97E-21 | 3 M1 _ | Rps6ka2 |
| Eif4g21 | 1.38E-25 | 1.069507 | 0.988 | 0.772 | 2.78E-21 | 3 M1 _ | Eif4g2 |
| Tox3 | 1.26E-25 | 0.333185 | 0.207 | 0.02 | 2.55E-21 | 3 M1 _ | Tox3 |
| Plscr1 | 1.19E-25 | 0.917643 | 0.61 | 0.185 | 2.40E-21 | 3 M1 _ | Plscr1 |
| Atp2b41 | 6.69E-26 | 0.994861 | 0.793 | 0.296 | 1.35E-21 | 3 M1 _ | Atp2b4 |
| 1500011K16Rik2 | 5.02E-26 | 0.347447 | 0.305 | 0.041 | 1.01E-21 | 3 M1 _ | 1500011K16Rik |
| Pyurf3 | 4.77E-26 | 0.437305 | 0.293 | 0.039 | 9.60E-22 | 3 M1 _ | Pyurf |
| Fat4 | 3.71E-26 | 0.57191 | 0.451 | 0.092 | 7.47E-22 | 3 M1 _ | Fat4 |
| Phlpp1 | 3.46E-26 | 0.739937 | 0.585 | 0.157 | 6.98E-22 | 3 M1 _ | Phlpp1 |
| Fry1 | 2.83E-26 | 0.717433 | 0.659 | 0.175 | 5.70E-22 | 3 M1 _ | Fry |
| Aaed12 | 2.82E-26 | 0.517229 | 0.378 | 0.063 | 5.69E-22 | 3 M1 _ | Aaed1 |
| Smad6 | 2.77E-26 | 0.530184 | 0.305 | 0.043 | 5.59E-22 | 3 M1 _ | Smad6 |
| Hip1 | 2.62E-26 | 1.055013 | 0.805 | 0.34 | 5.28E-22 | 3 M1 _ | Hip1 |
| Hyal2 | 2.19E-26 | 0.790919 | 0.5 | 0.118 | 4.41E-22 | 3 M1 _ | Hyal2 |
| AI3141802 | 2.12E-26 | 0.502087 | 0.378 | 0.063 | 4.28E-22 | 3 M1 _ | AI314180 |
| Ahr | 2.10E-26 | 0.968304 | 0.646 | 0.187 | 4.24E-22 | 3 M1 _ | Ahr |
| Shroom4 | 1.85E-26 | 0.477703 | 0.329 | 0.048 | 3.73E-22 | 3 M1 _ | Shroom4 |
| Fam58b2 | 1.82E-26 | 0.403551 | 0.268 | 0.032 | 3.67E-22 | 3 M1 _ | Fam58b |
| 4931406P16Rik1 | 1.61E-26 | 1.206082 | 0.866 | 0.432 | 3.23E-22 | 3 M1 _ | 4931406P16Rik |
| Fam43a | 1.02E-26 | 0.829237 | 0.61 | 0.163 | 2.05E-22 | 3 M1 _ | Fam43a |
| Clec2d | 7.49E-27 | 1.479963 | 0.927 | 0.641 | 1.51E-22 | 3 M1 _ | Clec2d |
| Fut8 | 5.81E-27 | 0.990712 | 0.793 | 0.314 | 1.17E-22 | 3 M1 _ | Fut8 |
| Ppp1r13b | 2.51E-27 | 0.583017 | 0.524 | 0.118 | 5.05E-23 | 3 M1 _ | Ppp1r13b |
| 6430548M08Rik | 1.49E-27 | 0.659749 | 0.427 | 0.082 | 3.00E-23 | 3 M1 _ | 6430548M08Rik |

| | | | | | | | |
|---|---|---|---|---|---|---|---|
| Spns2 | 1.29E-27 | 0.501131 | 0.329 | 0.047 | 2.60E-23 | 3 M1 _ | Spns2 |
| Cd341 | 1.20E-27 | 1.186386 | 0.927 | 0.432 | 2.41E-23 | 3 M1 _ | Cd34 |
| Cd38 | 1.13E-27 | 0.88004 | 0.695 | 0.217 | 2.27E-23 | 3 M1 _ | Cd38 |
| Sdccag32 | 6.14E-28 | 0.377105 | 0.305 | 0.039 | 1.24E-23 | 3 M1 _ | Sdccag3 |
| Acvrl1 | 5.66E-28 | 1.249699 | 0.841 | 0.391 | 1.14E-23 | 3 M1 _ | Acvrl1 |
| Fez21 | 5.56E-28 | 1.147296 | 0.854 | 0.363 | 1.12E-23 | 3 M1 _ | Fez2 |
| Kif26a | 5.09E-28 | 0.628421 | 0.341 | 0.049 | 1.02E-23 | 3 M1 _ | Kif26a |
| Sptan1 | 3.56E-28 | 1.307695 | 0.951 | 0.658 | 7.18E-24 | 3 M1 _ | Sptan1 |
| Clstn11 | 3.21E-28 | 1.159639 | 0.927 | 0.557 | 6.46E-24 | 3 M1 _ | Clstn1 |
| Arhgap29 | 1.70E-28 | 1.317556 | 0.927 | 0.611 | 3.43E-24 | 3 M1 _ | Arhgap29 |
| Zbtb162 | 1.42E-28 | 0.648595 | 0.463 | 0.087 | 2.86E-24 | 3 M1 _ | Zbtb16 |
| Arhgef3 | 1.41E-28 | 0.650524 | 0.488 | 0.1 | 2.84E-24 | 3 M1 _ | Arhgef3 |
| Endou | 1.04E-28 | 0.437091 | 0.293 | 0.035 | 2.09E-24 | 3 M1 _ | Endou |
| Lnx2 | 9.04E-29 | 0.632127 | 0.402 | 0.069 | 1.82E-24 | 3 M1 _ | Lnx2 |
| Tmcc2 | 7.13E-29 | 0.571648 | 0.402 | 0.068 | 1.44E-24 | 3 M1 _ | Tmcc2 |
| BC0033312 | 5.67E-29 | 0.534893 | 0.378 | 0.057 | 1.14E-24 | 3 M1 _ | BC003331 |
| Ptprm | 5.59E-29 | 0.789057 | 0.598 | 0.153 | 1.12E-24 | 3 M1 _ | Ptprm |
| Gpr4 | 5.40E-29 | 0.443592 | 0.317 | 0.041 | 1.09E-24 | 3 M1 _ | Gpr4 |
| Sox17 | 4.93E-29 | 0.838197 | 0.451 | 0.082 | 9.93E-25 | 3 M1 _ | Sox17 |
| Rhbdl2 | 2.84E-29 | 0.456662 | 0.256 | 0.026 | 5.72E-25 | 3 M1 _ | Rhbdl2 |
| Cited21 | 2.70E-29 | 1.09664 | 0.78 | 0.269 | 5.44E-25 | 3 M1 _ | Cited2 |
| Mgea53 | 2.41E-29 | 0.560757 | 0.341 | 0.048 | 4.85E-25 | 3 M1 _ | Mgea5 |
| Dusp3 | 2.13E-29 | 1.408965 | 0.817 | 0.345 | 4.29E-25 | 3 M1 _ | Dusp3 |
| Ncoa7 | 1.43E-29 | 1.517954 | 0.829 | 0.355 | 2.88E-25 | 3 M1 _ | Ncoa7 |
| Msn1 | 1.42E-29 | 1.286183 | 0.963 | 0.791 | 2.86E-25 | 3 M1 _ | Msn |
| Gm32688 | 1.31E-29 | 0.442179 | 0.28 | 0.031 | 2.63E-25 | 3 M1 _ | Gm32688 |
| Fam103a13 | 1.29E-29 | 0.476835 | 0.378 | 0.056 | 2.60E-25 | 3 M1 _ | Fam103a1 |
| Ptgs1 | 1.26E-29 | 1.261661 | 0.732 | 0.26 | 2.53E-25 | 3 M1 _ | Ptgs1 |
| 2810474O19Rik3 | 8.40E-30 | 0.860498 | 0.415 | 0.068 | 1.69E-25 | 3 M1 _ | 2810474O19Rik |
| Adam15 | 7.10E-30 | 1.235531 | 0.805 | 0.343 | 1.43E-25 | 3 M1 _ | Adam15 |
| Gnai2 | 5.24E-30 | 1.075432 | 0.976 | 0.923 | 1.06E-25 | 3 M1 _ | Gnai2 |
| Ephb4 | 5.01E-30 | 0.863564 | 0.585 | 0.144 | 1.01E-25 | 3 M1 _ | Ephb4 |
| Deup1 | 5.00E-30 | 0.251651 | 0.146 | 0.007 | 1.01E-25 | 3 M1 _ | Deup1 |
| St6galnac3 | 3.57E-30 | 0.513109 | 0.341 | 0.046 | 7.19E-26 | 3 M1 _ | St6galnac3 |
| Slco2a1 | 3.09E-30 | 0.950214 | 0.573 | 0.138 | 6.23E-26 | 3 M1 _ | Slco2a1 |
| Zfp979 | 1.77E-30 | 0.719215 | 0.378 | 0.056 | 3.57E-26 | 3 M1 _ | Zfp979 |
| Jup | 1.38E-30 | 1.538205 | 0.829 | 0.379 | 2.78E-26 | 3 M1 _ | Jup |
| St3gal5 | 1.02E-30 | 0.522831 | 0.341 | 0.046 | 2.05E-26 | 3 M1 _ | St3gal5 |
| Chml | 7.34E-31 | 0.536967 | 0.366 | 0.052 | 1.48E-26 | 3 M1 _ | Chml |
| Ripply3 | 6.08E-31 | 0.782257 | 0.512 | 0.105 | 1.22E-26 | 3 M1 _ | Ripply3 |
| 0610009O20Rik2 | 5.01E-31 | 0.409051 | 0.28 | 0.03 | 1.01E-26 | 3 M1 _ | 0610009O20Rik |

| | | | | | | | |
|---|---|---|---|---|---|---|---|
| Hmbox1 | 4.39E-31 | 1.10648 | 0.768 | 0.256 | 8.84E-27 | 3 M1 _ | Hmbox1 |
| Bmpr2 | 3.61E-31 | 1.440104 | 0.939 | 0.638 | 7.27E-27 | 3 M1 _ | Bmpr2 |
| Ccdc28b | 3.40E-31 | 1.258046 | 0.829 | 0.326 | 6.84E-27 | 3 M1 _ | Ccdc28b |
| Vamp5 | 3.15E-31 | 1.327735 | 0.927 | 0.457 | 6.35E-27 | 3 M1 _ | Vamp5 |
| Itpkb | 2.77E-31 | 1.316526 | 0.878 | 0.387 | 5.59E-27 | 3 M1 _ | Itpkb |
| 0610037L13Rik3 | 2.64E-31 | 0.484376 | 0.341 | 0.044 | 5.32E-27 | 3 M1 _ | 0610037L13Rik |
| Gpr146 | 1.67E-31 | 0.713808 | 0.5 | 0.097 | 3.36E-27 | 3 M1 _ | Gpr146 |
| Apold11 | 1.13E-31 | 0.536738 | 0.439 | 0.069 | 2.28E-27 | 3 M1 _ | Apold1 |
| Ndrg1 | 1.03E-31 | 1.213579 | 0.817 | 0.299 | 2.07E-27 | 3 M1 _ | Ndrg1 |
| Ctla2a | 7.22E-32 | 1.366884 | 0.927 | 0.336 | 1.45E-27 | 3 M1 _ | Ctla2a |
| Gm9917 | 6.25E-32 | 0.338481 | 0.232 | 0.019 | 1.26E-27 | 3 M1 _ | Gm9917 |
| Pitpnm2 | 4.42E-32 | 0.805369 | 0.573 | 0.125 | 8.89E-28 | 3 M1 _ | Pitpnm2 |
| Ramp21 | 4.07E-32 | 1.435999 | 0.939 | 0.482 | 8.19E-28 | 3 M1 _ | Ramp2 |
| Zcchc113 | 4.03E-32 | 0.489535 | 0.415 | 0.062 | 8.12E-28 | 3 M1 _ | Zcchc11 |
| Akr1c14 | 3.20E-32 | 0.671321 | 0.415 | 0.067 | 6.44E-28 | 3 M1 _ | Akr1c14 |
| Nat6 | 2.58E-32 | 0.261776 | 0.232 | 0.019 | 5.20E-28 | 3 M1 _ | Nat6 |
| Meox1 | 1.50E-32 | 1.122058 | 0.744 | 0.224 | 3.02E-28 | 3 M1 _ | Meox1 |
| Myo18a | 7.64E-33 | 1.42364 | 0.744 | 0.251 | 1.54E-28 | 3 M1 _ | Myo18a |
| Ly6a | 2.48E-33 | 1.671361 | 1 | 0.724 | 5.00E-29 | 3 M1 _ | Ly6a |
| Slc16a1 | 2.21E-33 | 0.839395 | 0.341 | 0.043 | 4.46E-29 | 3 M1 _ | Slc16a1 |
| Rapgef5 | 1.57E-33 | 0.931144 | 0.61 | 0.127 | 3.16E-29 | 3 M1 _ | Rapgef5 |
| Dlc11 | 1.38E-33 | 1.551995 | 0.976 | 0.696 | 2.77E-29 | 3 M1 _ | Dlc1 |
| Pla2g162 | 9.15E-34 | 0.44895 | 0.317 | 0.035 | 1.84E-29 | 3 M1 _ | Pla2g16 |
| Cdh13 | 8.89E-34 | 0.801918 | 0.415 | 0.062 | 1.79E-29 | 3 M1 _ | Cdh13 |
| Tmem54 | 7.74E-34 | 0.661799 | 0.463 | 0.075 | 1.56E-29 | 3 M1 _ | Tmem5 |
| AU0198233 | 7.60E-34 | 0.459947 | 0.305 | 0.032 | 1.53E-29 | 3 M1 _ | AU019823 |
| Cracr2b | 6.81E-34 | 0.770579 | 0.585 | 0.121 | 1.37E-29 | 3 M1 _ | Cracr2b |
| Heg1 | 7.07E-35 | 1.585225 | 0.829 | 0.322 | 1.42E-30 | 3 M1 _ | Heg1 |
| Npr1 | 5.86E-35 | 0.846915 | 0.549 | 0.109 | 1.18E-30 | 3 M1 _ | Npr1 |
| Unc45b | 5.17E-35 | 0.926092 | 0.537 | 0.103 | 1.04E-30 | 3 M1 _ | Unc45b |
| Rassf3 | 4.98E-35 | 0.890893 | 0.573 | 0.115 | 1.00E-30 | 3 M1 _ | Rassf3 |
| Scarb1 | 4.18E-35 | 0.840494 | 0.72 | 0.165 | 8.41E-31 | 3 M1 _ | Scarb1 |
| Nxpe4 | 3.63E-35 | 0.723319 | 0.378 | 0.05 | 7.30E-31 | 3 M1 _ | Nxpe4 |
| Snrk1 | 3.17E-35 | 1.582523 | 0.988 | 0.596 | 6.38E-31 | 3 M1 _ | Snrk |
| Timp31 | 2.81E-35 | 1.689011 | 1 | 0.784 | 5.66E-31 | 3 M1 _ | Timp3 |
| Mpped2 | 2.54E-35 | 0.436693 | 0.244 | 0.019 | 5.11E-31 | 3 M1 _ | Mpped2 |
| Slc27a3 | 2.51E-35 | 0.706065 | 0.378 | 0.05 | 5.06E-31 | 3 M1 _ | Slc27a3 |
| Mfsd6 | 1.03E-35 | 0.907668 | 0.537 | 0.101 | 2.08E-31 | 3 M1 _ | Mfsd6 |
| Hoxd8 | 9.32E-36 | 0.615057 | 0.427 | 0.062 | 1.88E-31 | 3 M1 _ | Hoxd8 |
| Igfbp3 | 4.53E-36 | 1.323391 | 0.829 | 0.252 | 9.13E-32 | 3 M1 _ | Igfbp3 |
| Abi3 | 4.40E-36 | 1.056553 | 0.707 | 0.177 | 8.86E-32 | 3 M1 _ | Abi3 |

| | | | | | | | |
|---|---|---|---|---|---|---|---|
| Notch4 | 4.29E-36 | 0.983864 | 0.622 | 0.135 | 8.64E-32 | 3 M1 _ | Notch4 |
| Calcrl | 2.94E-36 | 1.441992 | 0.817 | 0.298 | 5.93E-32 | 3 M1 _ | Calcrl |
| Prkch | 2.74E-36 | 0.799869 | 0.707 | 0.164 | 5.53E-32 | 3 M1 _ | Prkch |
| Pkp4 | 2.30E-36 | 1.026964 | 0.634 | 0.142 | 4.63E-32 | 3 M1 _ | Pkp4 |
| Epas11 | 1.96E-36 | 1.796681 | 0.951 | 0.515 | 3.95E-32 | 3 M1 _ | Epas1 |
| Acss1 | 1.60E-36 | 0.435032 | 0.293 | 0.027 | 3.23E-32 | 3 M1 _ | Acss1 |
| Nos3 | 1.45E-36 | 0.640479 | 0.378 | 0.047 | 2.91E-32 | 3 M1 _ | Nos3 |
| 1110004E09Rik1 | 1.13E-36 | 0.442066 | 0.293 | 0.027 | 2.28E-32 | 3 M1 _ | 1110004E09Rik |
| Prickle1 | 1.09E-36 | 0.82389 | 0.549 | 0.106 | 2.20E-32 | 3 M1 _ | Prickle1 |
| Nrp1 | 1.02E-36 | 1.829335 | 0.976 | 0.794 | 2.06E-32 | 3 M1 _ | Nrp1 |
| 1110008F13Rik3 | 9.92E-37 | 0.760635 | 0.5 | 0.082 | 2.00E-32 | 3 M1 _ | 1110008F13Rik |
| Ttll71 | 9.57E-37 | 0.970434 | 0.5 | 0.083 | 1.93E-32 | 3 M1 _ | Ttll7 |
| Klf2 | 8.34E-37 | 2.083978 | 0.927 | 0.45 | 1.68E-32 | 3 M1 _ | Klf2 |
| Nfib1 | 4.65E-37 | 1.629491 | 1 | 0.753 | 9.37E-33 | 3 M1 _ | Nfib |
| Dennd3 | 3.35E-37 | 0.559623 | 0.415 | 0.055 | 6.74E-33 | 3 M1 _ | Dennd3 |
| Tm4sf11 | 2.40E-37 | 1.906983 | 0.988 | 0.685 | 4.83E-33 | 3 M1 _ | Tm4sf1 |
| Jam2 | 2.38E-37 | 1.430941 | 0.866 | 0.319 | 4.79E-33 | 3 M1 _ | Jam2 |
| Hspb11 | 2.16E-37 | 1.832418 | 0.988 | 0.59 | 4.36E-33 | 3 M1 _ | Hspb1 |
| Tspan121 | 1.80E-37 | 0.991399 | 0.659 | 0.139 | 3.62E-33 | 3 M1 _ | Tspan12 |
| Sorbs21 | 1.78E-37 | 0.974153 | 0.744 | 0.174 | 3.59E-33 | 3 M1 _ | Sorbs2 |
| Entpd1 | 1.15E-37 | 1.271077 | 0.732 | 0.185 | 2.32E-33 | 3 M1 _ | Entpd1 |
| Clic4 | 6.88E-38 | 1.69877 | 0.951 | 0.572 | 1.38E-33 | 3 M1 _ | Clic4 |
| Arap3 | 5.72E-38 | 1.066177 | 0.707 | 0.169 | 1.15E-33 | 3 M1 _ | Arap3 |
| Sybu | 5.24E-38 | 0.598482 | 0.354 | 0.039 | 1.06E-33 | 3 M1 _ | Sybu |
| Ppp1r16b | 4.79E-38 | 0.797036 | 0.549 | 0.098 | 9.65E-34 | 3 M1 _ | Ppp1r16b |
| Calm11 | 3.95E-38 | 1.496157 | 0.988 | 0.95 | 7.95E-34 | 3 M1 _ | Calm1 |
| Fam208b2 | 3.30E-38 | 0.549234 | 0.305 | 0.028 | 6.65E-34 | 3 M1 _ | Fam208b |
| Anxa31 | 2.37E-38 | 1.34183 | 0.878 | 0.296 | 4.77E-34 | 3 M1 _ | Anxa3 |
| Fli1 | 2.32E-38 | 1.078762 | 0.805 | 0.223 | 4.68E-34 | 3 M1 _ | Fli1 |
| Lipe | 2.11E-38 | 0.996133 | 0.549 | 0.1 | 4.25E-34 | 3 M1 _ | Lipe |
| Ccm2l | 1.35E-38 | 0.72556 | 0.451 | 0.064 | 2.72E-34 | 3 M1 _ | Ccm2l |
| Rapgef3 | 4.59E-39 | 1.082126 | 0.634 | 0.135 | 9.23E-35 | 3 M1 _ | Rapgef3 |
| Scube3 | 2.28E-39 | 0.341193 | 0.207 | 0.011 | 4.59E-35 | 3 M1 _ | Scube3 |
| Ankrd29 | 2.15E-39 | 0.684539 | 0.39 | 0.046 | 4.33E-35 | 3 M1 _ | Ankrd29 |
| Tmem255b | 2.11E-39 | 0.644398 | 0.427 | 0.055 | 4.24E-35 | 3 M1 _ | Tmem255b |
| Arhgap27 | 8.52E-40 | 0.849561 | 0.549 | 0.094 | 1.72E-35 | 3 M1 _ | Arhgap27 |
| Plxna2 | 8.03E-40 | 1.192037 | 0.634 | 0.136 | 1.62E-35 | 3 M1 _ | Plxna2 |
| Mef2c1 | 7.22E-40 | 1.992195 | 0.988 | 0.64 | 1.45E-35 | 3 M1 _ | Mef2c |
| Ptprg | 5.26E-40 | 1.539824 | 0.927 | 0.405 | 1.06E-35 | 3 M1 _ | Ptprg |
| Abcg2 | 2.08E-40 | 1.674733 | 0.89 | 0.33 | 4.19E-36 | 3 M1 _ | Abcg2 |
| Sptbn1 | 1.75E-40 | 1.684171 | 1 | 0.898 | 3.52E-36 | 3 M1 _ | Sptbn1 |

| | | | | | | | |
|---|---|---|---|---|---|---|---|
| Plpp31 | 1.47E-40 | 2.411758 | 0.988 | 0.722 | 2.96E-36 | 3 M1 _ | Plpp3 |
| Ece1 | 9.89E-41 | 1.667696 | 0.902 | 0.349 | 1.99E-36 | 3 M1 _ | Ece1 |
| Afap1l1 | 6.48E-41 | 0.83681 | 0.488 | 0.071 | 1.30E-36 | 3 M1 _ | Afap1l1 |
| Cxcl12 | 6.38E-41 | 3.080739 | 0.988 | 0.787 | 1.28E-36 | 3 M1 _ | Cxcl12 |
| Kitl1 | 3.71E-41 | 1.885658 | 0.915 | 0.391 | 7.47E-37 | 3 M1 _ | Kitl |
| Rapgef4 | 2.55E-41 | 0.973223 | 0.537 | 0.087 | 5.13E-37 | 3 M1 _ | Rapgef4 |
| Esam1 | 2.11E-41 | 1.447976 | 0.963 | 0.314 | 4.25E-37 | 3 M1 _ | Esam |
| Tmem572 | 1.58E-41 | 0.538911 | 0.341 | 0.033 | 3.17E-37 | 3 M1 _ | Tmem57 |
| Jag2 | 1.54E-41 | 0.870555 | 0.488 | 0.069 | 3.10E-37 | 3 M1 _ | Jag2 |
| Pik3r3 | 1.45E-41 | 0.709653 | 0.5 | 0.074 | 2.93E-37 | 3 M1 _ | Pik3r3 |
| Sipa1l2 | 1.42E-41 | 1.008771 | 0.524 | 0.085 | 2.87E-37 | 3 M1 _ | Sipa1l2 |
| Cav21 | 1.21E-41 | 1.162373 | 0.768 | 0.188 | 2.44E-37 | 3 M1 _ | Cav2 |
| Syne1 | 8.32E-42 | 1.774331 | 0.89 | 0.322 | 1.68E-37 | 3 M1 _ | Syne1 |
| Eng | 7.65E-42 | 1.778519 | 1 | 0.662 | 1.54E-37 | 3 M1 _ | Eng |
| Sdk2 | 5.34E-42 | 0.591878 | 0.354 | 0.035 | 1.08E-37 | 3 M1 _ | Sdk2 |
| Fam96a1 | 5.22E-42 | 0.456491 | 0.354 | 0.035 | 1.05E-37 | 3 M1 _ | Fam96a |
| Gnas2 | 1.68E-42 | 1.358998 | 1 | 0.96 | 3.38E-38 | 3 M1 _ | Gnas |
| Tmtc1 | 1.52E-42 | 0.725259 | 0.524 | 0.079 | 3.07E-38 | 3 M1 _ | Tmtc1 |
| Gpr150 | 1.36E-42 | 0.307836 | 0.183 | 0.007 | 2.73E-38 | 3 M1 _ | Gpr150 |
| Plpp1 | 6.00E-43 | 2.791234 | 0.951 | 0.588 | 1.21E-38 | 3 M1 _ | Plpp1 |
| Bcam1 | 4.36E-43 | 1.520976 | 0.951 | 0.309 | 8.79E-39 | 3 M1 _ | Bcam |
| Etl4 | 3.38E-43 | 1.212575 | 0.707 | 0.163 | 6.81E-39 | 3 M1 _ | Etl4 |
| Tspan13 | 1.40E-43 | 1.384688 | 0.744 | 0.172 | 2.81E-39 | 3 M1 _ | Tspan13 |
| Gipc3 | 9.64E-44 | 0.465756 | 0.305 | 0.024 | 1.94E-39 | 3 M1 _ | Gipc3 |
| Exoc3l | 9.39E-44 | 0.553001 | 0.317 | 0.026 | 1.89E-39 | 3 M1 _ | Exoc3l |
| Ccnd11 | 8.84E-44 | 1.469225 | 0.732 | 0.157 | 1.78E-39 | 3 M1 _ | Ccnd1 |
| Pcp4l11 | 4.19E-44 | 1.492326 | 0.829 | 0.205 | 8.43E-40 | 3 M1 _ | Pcp4l1 |
| Bvht | 4.18E-44 | 1.214955 | 0.695 | 0.154 | 8.42E-40 | 3 M1 _ | Bvht |
| Efr3b | 1.35E-44 | 0.618827 | 0.415 | 0.046 | 2.72E-40 | 3 M1 _ | Efr3b |
| Minos13 | 1.15E-44 | 0.974521 | 0.646 | 0.11 | 2.33E-40 | 3 M1 _ | Minos1 |
| 9030624J02Rik2 | 8.81E-45 | 0.475578 | 0.366 | 0.035 | 1.78E-40 | 3 M1 _ | 9030624J02Rik |
| Fzd41 | 8.21E-45 | 1.648944 | 0.939 | 0.327 | 1.65E-40 | 3 M1 _ | Fzd4 |
| Efnb2 | 5.10E-45 | 1.044343 | 0.646 | 0.117 | 1.03E-40 | 3 M1 _ | Efnb2 |
| Papd51 | 4.96E-45 | 0.497889 | 0.293 | 0.021 | 9.99E-41 | 3 M1 _ | Papd5 |
| Klf4 | 4.44E-45 | 1.843763 | 0.902 | 0.295 | 8.95E-41 | 3 M1 _ | Klf4 |
| Fabp51 | 4.21E-45 | 2.321958 | 0.805 | 0.233 | 8.49E-41 | 3 M1 _ | Fabp5 |
| Tinagl11 | 3.04E-45 | 1.671647 | 0.963 | 0.286 | 6.11E-41 | 3 M1 _ | Tinagl1 |
| Cyr612 | 1.33E-45 | 0.558793 | 0.329 | 0.027 | 2.69E-41 | 3 M1 _ | Cyr61 |
| Sgpp2 | 1.19E-45 | 0.274905 | 0.183 | 0.006 | 2.39E-41 | 3 M1 _ | Sgpp2 |
| Enpp6 | 1.05E-45 | 0.445181 | 0.22 | 0.01 | 2.12E-41 | 3 M1 _ | Enpp6 |
| Lrrc3b | 7.96E-46 | 0.389059 | 0.268 | 0.016 | 1.60E-41 | 3 M1 _ | Lrrc3b |

| | | | | | | | |
|---|---|---|---|---|---|---|---|
| Sema3c | 4.34E-46 | 1.277553 | 0.61 | 0.104 | 8.75E-42 | 3 M1 _ | Sema3c |
| Tdrp | 4.29E-46 | 0.74852 | 0.439 | 0.051 | 8.64E-42 | 3 M1 _ | Tdrp |
| 1810022K09Rik3 | 3.39E-46 | 0.79729 | 0.585 | 0.087 | 6.82E-42 | 3 M1 _ | 1810022K09Rik |
| Mum13 | 2.92E-46 | 0.706049 | 0.427 | 0.046 | 5.89E-42 | 3 M1 _ | Mum1 |
| Cd361 | 2.28E-46 | 2.066302 | 0.793 | 0.183 | 4.60E-42 | 3 M1 _ | Cd36 |
| Srgn1 | 9.13E-47 | 2.127405 | 0.976 | 0.401 | 1.84E-42 | 3 M1 _ | Srgn |
| Nr5a2 | 7.69E-47 | 0.71058 | 0.256 | 0.015 | 1.55E-42 | 3 M1 _ | Nr5a2 |
| Sema3f | 4.78E-47 | 0.938953 | 0.561 | 0.084 | 9.63E-43 | 3 M1 _ | Sema3f |
| Dok4 | 3.15E-47 | 1.009302 | 0.61 | 0.1 | 6.35E-43 | 3 M1 _ | Dok4 |
| Plekho1 | 2.79E-47 | 1.119263 | 0.744 | 0.15 | 5.63E-43 | 3 M1 _ | Plekho1 |
| Egfl7 | 2.30E-47 | 2.200471 | 0.976 | 0.373 | 4.64E-43 | 3 M1 _ | Egfl7 |
| Mmp15 | 1.61E-47 | 0.890948 | 0.524 | 0.071 | 3.25E-43 | 3 M1 _ | Mmp15 |
| C130074G19Rik | 1.23E-47 | 1.198087 | 0.585 | 0.091 | 2.48E-43 | 3 M1 _ | C130074G19Rik |
| Zfp366 | 1.11E-47 | 0.717631 | 0.463 | 0.054 | 2.24E-43 | 3 M1 _ | Zfp366 |
| Thbd1 | 9.56E-48 | 2.317095 | 0.963 | 0.397 | 1.92E-43 | 3 M1 _ | Thbd |
| Impdh1 | 4.29E-48 | 1.152306 | 0.732 | 0.153 | 8.63E-44 | 3 M1 _ | Impdh1 |
| Fgd5 | 4.01E-48 | 1.310431 | 0.756 | 0.17 | 8.08E-44 | 3 M1 _ | Fgd5 |
| Nostrin | 2.99E-48 | 1.509548 | 0.78 | 0.171 | 6.02E-44 | 3 M1 _ | Nostrin |
| Enah | 2.42E-48 | 0.962342 | 0.549 | 0.078 | 4.87E-44 | 3 M1 _ | Enah |
| Rasgrf2 | 1.30E-48 | 0.652145 | 0.451 | 0.049 | 2.62E-44 | 3 M1 _ | Rasgrf2 |
| Gpihbp1 | 1.15E-48 | 1.958321 | 0.829 | 0.204 | 2.32E-44 | 3 M1 _ | Gpihbp1 |
| Rnf1521 | 1.00E-48 | 0.858908 | 0.524 | 0.068 | 2.02E-44 | 3 M1 _ | Rnf152 |
| She | 9.62E-49 | 1.007315 | 0.646 | 0.108 | 1.94E-44 | 3 M1 _ | She |
| Rdx | 7.59E-49 | 3.013101 | 0.988 | 0.796 | 1.53E-44 | 3 M1 _ | Rdx |
| Hlx | 5.70E-49 | 1.004945 | 0.561 | 0.08 | 1.15E-44 | 3 M1 _ | Hlx |
| Flt4 | 4.51E-49 | 0.693187 | 0.512 | 0.064 | 9.08E-45 | 3 M1 _ | Flt4 |
| Plvap | 3.54E-49 | 2.893919 | 1 | 0.51 | 7.12E-45 | 3 M1 _ | Plvap |
| Lrrc8c1 | 3.42E-49 | 1.438191 | 0.866 | 0.227 | 6.88E-45 | 3 M1 _ | Lrrc8c |
| Shank3 | 3.04E-49 | 0.819392 | 0.439 | 0.047 | 6.13E-45 | 3 M1 _ | Shank3 |
| Rmst | 1.90E-49 | 0.53242 | 0.268 | 0.015 | 3.83E-45 | 3 M1 _ | Rmst |
| Rtp3 | 1.35E-49 | 0.683465 | 0.402 | 0.038 | 2.72E-45 | 3 M1 _ | Rtp3 |
| Tnfsf10 | 1.16E-49 | 1.578094 | 0.841 | 0.208 | 2.33E-45 | 3 M1 _ | Tnfsf10 |
| Slc26a10 | 7.16E-50 | 0.733159 | 0.39 | 0.035 | 1.44E-45 | 3 M1 _ | Slc26a10 |
| Kctd12b | 2.48E-50 | 1.86657 | 0.915 | 0.307 | 4.99E-46 | 3 M1 _ | Kctd12b |
| Gcn1l13 | 2.25E-50 | 0.587202 | 0.39 | 0.035 | 4.53E-46 | 3 M1 _ | Gcn1l1 |
| Cav11 | 1.15E-50 | 1.899306 | 0.939 | 0.289 | 2.31E-46 | 3 M1 _ | Cav1 |
| Ecscr | 9.42E-51 | 1.338428 | 0.878 | 0.211 | 1.90E-46 | 3 M1 _ | Ecscr |
| 2010107E04Rik3 | 4.75E-51 | 0.956862 | 0.695 | 0.11 | 9.57E-47 | 3 M1 _ | 2010107E04Rik |
| Arhgap31 | 3.81E-51 | 2.001532 | 0.951 | 0.341 | 7.67E-47 | 3 M1 _ | Arhgap31 |
| Clec14a | 1.51E-51 | 2.056745 | 0.976 | 0.362 | 3.04E-47 | 3 M1 _ | Clec14a |
| Pim3 | 1.68E-52 | 1.369691 | 0.744 | 0.149 | 3.38E-48 | 3 M1 _ | Pim3 |

| | | | | | | | |
|---|---|---|---|---|---|---|---|
| Gatsl3 | 1.20E-52 | 0.451732 | 0.305 | 0.019 | 2.41E-48 | 3 M1 _ | Gatsl3 |
| Slc9a3r2 | 3.79E-53 | 2.634004 | 0.988 | 0.453 | 7.63E-49 | 3 M1 _ | Slc9a3r2 |
| Grrp1 | 2.26E-53 | 1.221655 | 0.695 | 0.11 | 4.56E-49 | 3 M1 _ | Grrp1 |
| Ly6c1 | 5.14E-54 | 2.721514 | 1 | 0.406 | 1.04E-49 | 3 M1 _ | Ly6c1 |
| Oraov11 | 2.25E-54 | 0.54501 | 0.329 | 0.021 | 4.54E-50 | 3 M1 _ | Oraov1 |
| Pkn3 | 1.53E-54 | 0.602727 | 0.415 | 0.036 | 3.09E-50 | 3 M1 _ | Pkn3 |
| Crim11 | 1.30E-54 | 2.211794 | 0.915 | 0.275 | 2.61E-50 | 3 M1 _ | Crim1 |
| Sema7a | 7.66E-55 | 1.703875 | 0.866 | 0.217 | 1.54E-50 | 3 M1 _ | Sema7a |
| 2010111I01Rik2 | 5.70E-55 | 0.93735 | 0.537 | 0.063 | 1.15E-50 | 3 M1 _ | 2010111I01Rik |
| Adcy4 | 2.15E-55 | 1.081266 | 0.768 | 0.139 | 4.33E-51 | 3 M1 _ | Adcy4 |
| Sox18 | 1.83E-55 | 0.858033 | 0.573 | 0.071 | 3.70E-51 | 3 M1 _ | Sox18 |
| Chrm3 | 1.28E-55 | 0.458057 | 0.244 | 0.01 | 2.58E-51 | 3 M1 _ | Chrm3 |
| Gata2 | 9.74E-56 | 0.906698 | 0.512 | 0.058 | 1.96E-51 | 3 M1 _ | Gata2 |
| Pltp | 8.81E-56 | 2.232135 | 0.951 | 0.304 | 1.77E-51 | 3 M1 _ | Pltp |
| Ehd4 | 6.03E-56 | 2.711368 | 0.951 | 0.331 | 1.21E-51 | 3 M1 _ | Ehd4 |
| Fam60a1 | 4.66E-56 | 0.355969 | 0.22 | 0.007 | 9.38E-52 | 3 M1 _ | Fam60a |
| 2410015M20Rik3 | 1.36E-56 | 0.891132 | 0.683 | 0.098 | 2.74E-52 | 3 M1 _ | 2410015M20Rik |
| B4galt4 | 1.32E-56 | 1.351613 | 0.744 | 0.138 | 2.66E-52 | 3 M1 _ | B4galt4 |
| D17Wsu92e3 | 9.81E-57 | 0.734992 | 0.476 | 0.047 | 1.98E-52 | 3 M1 _ | D17Wsu92e |
| Plscr2 | 6.37E-57 | 2.63546 | 0.976 | 0.362 | 1.28E-52 | 3 M1 _ | Plscr2 |
| Tcim | 4.61E-57 | 2.153882 | 0.927 | 0.261 | 9.29E-53 | 3 M1 _ | Tcim |
| Ushbp1 | 4.61E-57 | 1.437574 | 0.78 | 0.141 | 9.28E-53 | 3 M1 _ | Ushbp1 |
| Gngt2 | 4.59E-57 | 1.757485 | 0.902 | 0.239 | 9.24E-53 | 3 M1 _ | Gngt2 |
| Usmg53 | 1.98E-57 | 1.269469 | 0.732 | 0.112 | 3.98E-53 | 3 M1 _ | Usmg5 |
| Tek | 1.85E-57 | 1.397099 | 0.793 | 0.143 | 3.72E-53 | 3 M1 _ | Tek |
| Cdkn1a1 | 7.18E-58 | 1.687151 | 0.817 | 0.166 | 1.45E-53 | 3 M1 _ | Cdkn1a |
| Caskin2 | 6.08E-58 | 1.292735 | 0.744 | 0.131 | 1.22E-53 | 3 M1 _ | Caskin2 |
| Atp5o.13 | 5.11E-58 | 1.037798 | 0.707 | 0.105 | 1.03E-53 | 3 M1 _ | Atp5o.1 |
| Fabp4 | 3.87E-58 | 3.876567 | 1 | 0.435 | 7.79E-54 | 3 M1 _ | Fabp4 |
| Samd12 | 2.01E-58 | 0.762818 | 0.524 | 0.055 | 4.06E-54 | 3 M1 _ | Samd12 |
| Rflnb | 1.49E-58 | 2.734064 | 0.951 | 0.3 | 3.00E-54 | 3 M1 _ | Rflnb |
| Fam96b3 | 1.22E-58 | 0.84038 | 0.524 | 0.056 | 2.45E-54 | 3 M1 _ | Fam96b |
| Nova2 | 1.04E-58 | 0.795117 | 0.524 | 0.056 | 2.10E-54 | 3 M1 _ | Nova2 |
| Lhx6 | 6.52E-59 | 0.760791 | 0.317 | 0.018 | 1.31E-54 | 3 M1 _ | Lhx6 |
| Smco4 | 2.39E-59 | 2.03169 | 0.915 | 0.242 | 4.82E-55 | 3 M1 _ | Smco4 |
| Slc6a6 | 8.43E-60 | 3.102767 | 0.963 | 0.343 | 1.70E-55 | 3 M1 _ | Slc6a6 |
| Chst15 | 4.85E-60 | 1.276507 | 0.744 | 0.124 | 9.76E-56 | 3 M1 _ | Chst15 |
| F11r | 9.77E-61 | 1.205651 | 0.756 | 0.126 | 1.97E-56 | 3 M1 _ | F11r |
| Ptprr | 6.95E-61 | 0.946184 | 0.415 | 0.032 | 1.40E-56 | 3 M1 _ | Ptprr |
| Mcc | 3.14E-61 | 1.656111 | 0.78 | 0.152 | 6.32E-57 | 3 M1 _ | Mcc |
| Tbxa2r | 2.76E-61 | 0.776277 | 0.512 | 0.051 | 5.57E-57 | 3 M1 _ | Tbxa2r |

| | | | | | | | |
|---|---|---|---|---|---|---|---|
| Hecw2 | 1.01E-61 | 1.582948 | 0.829 | 0.165 | 2.03E-57 | 3 M1 _ | Hecw2 |
| Trove22 | 7.18E-62 | 1.049805 | 0.585 | 0.067 | 1.45E-57 | 3 M1 _ | Trove2 |
| Olfr1396 | 3.15E-62 | 0.808434 | 0.341 | 0.02 | 6.35E-58 | 3 M1 _ | Olfr1396 |
| Cavin2 | 3.78E-63 | 2.294266 | 0.988 | 0.265 | 7.61E-59 | 3 M1 _ | Cavin2 |
| Car8 | 2.88E-63 | 0.893013 | 0.524 | 0.051 | 5.81E-59 | 3 M1 _ | Car8 |
| Rsad2 | 2.42E-63 | 2.632818 | 0.89 | 0.199 | 4.87E-59 | 3 M1 _ | Rsad2 |
| Lama3 | 1.13E-63 | 0.952465 | 0.561 | 0.06 | 2.27E-59 | 3 M1 _ | Lama3 |
| Pecam1 | 8.60E-64 | 2.357342 | 0.988 | 0.257 | 1.73E-59 | 3 M1 _ | Pecam1 |
| Sgk1 | 5.44E-64 | 2.773484 | 0.939 | 0.269 | 1.10E-59 | 3 M1 _ | Sgk1 |
| Fam208a3 | 5.23E-64 | 1.0833 | 0.598 | 0.068 | 1.05E-59 | 3 M1 _ | Fam208a |
| Icam2 | 3.80E-64 | 1.349581 | 0.829 | 0.139 | 7.65E-60 | 3 M1 _ | Icam2 |
| Syt15 | 2.31E-64 | 0.740485 | 0.439 | 0.034 | 4.64E-60 | 3 M1 _ | Syt15 |
| Cd200 | 7.82E-65 | 2.240684 | 0.976 | 0.249 | 1.57E-60 | 3 M1 _ | Cd200 |
| Itga6 | 3.66E-65 | 1.704248 | 0.854 | 0.156 | 7.36E-61 | 3 M1 _ | Itga6 |
| Aes3 | 1.32E-66 | 1.456532 | 0.78 | 0.115 | 2.67E-62 | 3 M1 _ | Aes |
| Emcn | 6.03E-67 | 2.884653 | 0.963 | 0.256 | 1.22E-62 | 3 M1 _ | Emcn |
| Ptprb | 4.15E-67 | 2.688812 | 0.988 | 0.26 | 8.35E-63 | 3 M1 _ | Ptprb |
| Id1 | 1.40E-67 | 2.710077 | 0.963 | 0.263 | 2.83E-63 | 3 M1 _ | Id1 |
| Dysf | 7.18E-68 | 1.66509 | 0.756 | 0.118 | 1.45E-63 | 3 M1 _ | Dysf |
| Grin2a | 4.69E-68 | 0.345451 | 0.183 | 0.002 | 9.44E-64 | 3 M1 _ | Grin2a |
| Kank3 | 1.26E-68 | 1.43511 | 0.829 | 0.137 | 2.54E-64 | 3 M1 _ | Kank3 |
| Clec1a | 9.29E-69 | 1.918281 | 0.829 | 0.15 | 1.87E-64 | 3 M1 _ | Clec1a |
| Adgrf5 | 7.10E-69 | 2.289982 | 0.951 | 0.214 | 1.43E-64 | 3 M1 _ | Adgrf5 |
| Palmd | 7.98E-70 | 1.452152 | 0.72 | 0.096 | 1.61E-65 | 3 M1 _ | Palmd |
| Vwa3a | 7.79E-70 | 0.544149 | 0.305 | 0.012 | 1.57E-65 | 3 M1 _ | Vwa3a |
| Ccdc85a | 3.70E-71 | 0.965042 | 0.561 | 0.053 | 7.45E-67 | 3 M1 _ | Ccdc85a |
| Fam212a1 | 3.37E-71 | 0.716245 | 0.39 | 0.022 | 6.78E-67 | 3 M1 _ | Fam212a |
| Pcdh17 | 1.48E-71 | 1.783992 | 0.805 | 0.12 | 2.97E-67 | 3 M1 _ | Pcdh17 |
| Plcb1 | 8.93E-72 | 1.645986 | 0.829 | 0.133 | 1.80E-67 | 3 M1 _ | Plcb1 |
| Ceacam1 | 5.87E-72 | 1.284626 | 0.707 | 0.086 | 1.18E-67 | 3 M1 _ | Ceacam1 |
| Mmrn2 | 4.94E-72 | 1.602746 | 0.89 | 0.149 | 9.95E-68 | 3 M1 _ | Mmrn2 |
| Rasip1 | 7.50E-73 | 1.393653 | 0.841 | 0.127 | 1.51E-68 | 3 M1 _ | Rasip1 |
| Myct1 | 4.41E-73 | 1.338292 | 0.817 | 0.117 | 8.87E-69 | 3 M1 _ | Myct1 |
| Robo4 | 4.16E-73 | 1.216443 | 0.683 | 0.081 | 8.38E-69 | 3 M1 _ | Robo4 |
| Mgll | 2.07E-73 | 3.167218 | 0.976 | 0.254 | 4.17E-69 | 3 M1 _ | Mgll |
| Flt1 | 9.10E-74 | 3.47485 | 1 | 0.267 | 1.83E-69 | 3 M1 _ | Flt1 |
| Gm12002 | 2.54E-74 | 1.135431 | 0.561 | 0.05 | 5.11E-70 | 3 M1 _ | Gm12002 |
| Alox12 | 3.35E-75 | 1.089989 | 0.561 | 0.048 | 6.75E-71 | 3 M1 _ | Alox12 |
| Acer2 | 5.42E-76 | 0.954235 | 0.5 | 0.037 | 1.09E-71 | 3 M1 _ | Acer2 |
| Parvb | 3.41E-77 | 1.076357 | 0.61 | 0.059 | 6.86E-73 | 3 M1 _ | Parvb |
| Lims2 | 1.86E-77 | 1.452784 | 0.866 | 0.13 | 3.74E-73 | 3 M1 _ | Lims2 |

| | | | | | | | |
|---|---|---|---|---|---|---|---|
| S1pr1 | 1.41E-77 | 2.291023 | 0.915 | 0.17 | 2.84E-73 | 3 M1 _ | S1pr1 |
| Rgcc | 7.35E-78 | 3.662971 | 0.988 | 0.243 | 1.48E-73 | 3 M1 _ | Rgcc |
| Adgrl4 | 2.34E-78 | 2.161645 | 0.976 | 0.18 | 4.71E-74 | 3 M1 _ | Adgrl4 |
| Proser2 | 2.12E-78 | 1.019509 | 0.549 | 0.045 | 4.27E-74 | 3 M1 _ | Proser2 |
| Col13a1 | 5.32E-80 | 2.103377 | 0.902 | 0.154 | 1.07E-75 | 3 M1 _ | Col13a1 |
| 4930578C19Rik | 4.29E-80 | 0.31933 | 0.159 | 0 | 8.64E-76 | 3 M1 _ | 4930578C19Rik |
| Pdgfb | 4.09E-80 | 2.544089 | 0.951 | 0.193 | 8.23E-76 | 3 M1 _ | Pdgfb |
| Cdh5 | 2.85E-80 | 2.237979 | 0.963 | 0.174 | 5.73E-76 | 3 M1 _ | Cdh5 |
| Ssfa21 | 2.83E-80 | 0.706379 | 0.427 | 0.023 | 5.70E-76 | 3 M1 _ | Ssfa2 |
| Fam167b | 1.24E-81 | 1.86797 | 0.805 | 0.107 | 2.50E-77 | 3 M1 _ | Fam167b |
| Akap21 | 7.66E-82 | 0.665999 | 0.354 | 0.014 | 1.54E-77 | 3 M1 _ | Akap2 |
| Cd93 | 4.46E-82 | 2.137187 | 0.939 | 0.158 | 8.98E-78 | 3 M1 _ | Cd93 |
| Magix | 5.15E-83 | 0.76326 | 0.427 | 0.022 | 1.04E-78 | 3 M1 _ | Magix |
| Ramp3 | 1.07E-84 | 1.86512 | 0.878 | 0.127 | 2.16E-80 | 3 M1 _ | Ramp3 |
| Tie1 | 5.51E-85 | 2.133856 | 0.951 | 0.161 | 1.11E-80 | 3 M1 _ | Tie1 |
| Car4 | 4.07E-85 | 1.923353 | 0.683 | 0.07 | 8.20E-81 | 3 M1 _ | Car4 |
| Kdr | 1.13E-85 | 3.589535 | 0.988 | 0.211 | 2.27E-81 | 3 M1 _ | Kdr |
| Rasgrp3 | 1.73E-86 | 1.853898 | 0.854 | 0.116 | 3.48E-82 | 3 M1 _ | Rasgrp3 |
| Cmtm8 | 4.93E-87 | 1.851554 | 0.805 | 0.102 | 9.92E-83 | 3 M1 _ | Cmtm8 |
| Rbp7 | 1.38E-87 | 3.471194 | 1 | 0.193 | 2.78E-83 | 3 M1 _ | Rbp7 |
| Cd300lg | 2.39E-89 | 2.479438 | 0.939 | 0.156 | 4.82E-85 | 3 M1 _ | Cd300lg |
| Adgrg1 | 2.01E-90 | 1.443022 | 0.78 | 0.089 | 4.06E-86 | 3 M1 _ | Adgrg1 |
| Podxl | 1.08E-90 | 3.083257 | 0.976 | 0.18 | 2.17E-86 | 3 M1 _ | Podxl |
| Tmem88 | 1.57E-91 | 1.522528 | 0.817 | 0.094 | 3.16E-87 | 3 M1 _ | Tmem88 |
| Fam198b1 | 4.34E-93 | 1.621172 | 0.768 | 0.074 | 8.75E-89 | 3 M1 _ | Fam198b |
| 1700020I14Rik2 | 8.07E-94 | 1.51086 | 0.793 | 0.084 | 1.63E-89 | 3 M1 _ | 1700020I14Rik |
| Arhgef15 | 4.98E-95 | 1.576205 | 0.854 | 0.101 | 1.00E-90 | 3 M1 _ | Arhgef15 |
| Esm1 | 5.09E-96 | 1.467349 | 0.5 | 0.027 | 1.02E-91 | 3 M1 _ | Esm1 |
| Cyyr1 | 7.46E-97 | 2.588824 | 0.963 | 0.145 | 1.50E-92 | 3 M1 _ | Cyyr1 |
| Clic5 | 2.11E-100 | 2.331695 | 0.951 | 0.131 | 4.25E-96 | 3 M1 _ | Clic5 |
| Thrsp | 2.23E-101 | 2.873073 | 0.951 | 0.141 | 4.49E-97 | 3 M1 _ | Thrsp |
| Btnl9 | 1.15E-102 | 3.064075 | 0.915 | 0.124 | 2.32E-98 | 3 M1 _ | Btnl9 |
| Mcf2l | 5.94E-104 | 1.862431 | 0.878 | 0.103 | 1.20E-99 | 3 M1 _ | Mcf2l |
| Ces2e | 2.72E-104 | 1.622393 | 0.744 | 0.067 | 5.47E-100 | 3 M1 _ | Ces2e |
| Cd24a | 2.61E-107 | 2.470109 | 0.915 | 0.114 | 5.26E-103 | 3 M1 _ | Cd24a |
| Zcchc63 | 5.83E-108 | 1.483826 | 0.805 | 0.075 | 1.17E-103 | 3 M1 _ | Zcchc6 |
| Exoc3l2 | 5.50E-112 | 1.98132 | 0.866 | 0.09 | 1.11E-107 | 3 M1 _ | Exoc3l2 |
| Tmem21 | 2.74E-123 | 0.744372 | 0.451 | 0.013 | 5.52E-119 | 3 M1 _ | Tmem2 |
| Mkl21 | 1.30E-140 | 1.461867 | 0.671 | 0.033 | 2.61E-136 | 3 M1 _ | Mkl2 |
| 8430408G22Rik | 1.37E-245 | 1.410893 | 0.512 | 0.001 | 2.77E-241 | 3 M1 _ | 8430408G22Rik |
| Steap32 | 2.32E-06 | 0.88659 | 0.941 | 0.36 | 0.046685 | 4 L1 _ | Steap3 |

| | | | | | | | |
|---|---|---|---|---|---|---|---|
| Acyp1 | 2.06E-06 | 0.260375 | 0.529 | 0.12 | 0.04148 | 4 L1 _ | Acyp1 |
| Adam17 | 1.87E-06 | 0.588332 | 0.882 | 0.336 | 0.037729 | 4 L1 _ | Adam17 |
| Serinc5 | 1.82E-06 | 0.486668 | 0.471 | 0.108 | 0.036664 | 4 L1 _ | Serinc5 |
| Trim30d1 | 1.82E-06 | 0.615831 | 0.471 | 0.111 | 0.036621 | 4 L1 _ | Trim30d |
| Galnt171 | 1.80E-06 | 0.409765 | 0.412 | 0.085 | 0.036152 | 4 L1 _ | Galnt17 |
| Dock42 | 1.74E-06 | 0.400405 | 0.529 | 0.124 | 0.035141 | 4 L1 _ | Dock4 |
| Amotl21 | 1.52E-06 | 0.640423 | 0.706 | 0.225 | 0.030526 | 4 L1 _ | Amotl2 |
| Rpl37a2 | 1.48E-06 | 0.640381 | 1 | 0.985 | 0.029815 | 4 L1 _ | Rpl37a |
| Dock111 | 1.44E-06 | 0.503889 | 0.588 | 0.15 | 0.02891 | 4 L1 _ | Dock11 |
| Rarres23 | 1.37E-06 | 0.956697 | 1 | 0.689 | 0.027659 | 4 L1 _ | Rarres2 |
| Tmem43 | 1.36E-06 | 0.567359 | 0.647 | 0.191 | 0.027308 | 4 L1 _ | Tmem43 |
| Fpgt | 1.32E-06 | 0.342721 | 0.353 | 0.062 | 0.026533 | 4 L1 _ | Fpgt |
| Rab11fip5 | 1.27E-06 | 0.57325 | 0.529 | 0.132 | 0.025653 | 4 L1 _ | Rab11fip5 |
| Fbln23 | 1.27E-06 | 1.053499 | 0.941 | 0.543 | 0.025585 | 4 L1 _ | Fbln2 |
| Tmem19 | 1.27E-06 | 0.361074 | 0.588 | 0.146 | 0.025482 | 4 L1 _ | Tmem19 |
| Cercam | 1.17E-06 | 0.589471 | 0.471 | 0.108 | 0.023648 | 4 L1 _ | Cercam |
| Sestd1 | 1.01E-06 | 0.529993 | 0.529 | 0.131 | 0.020298 | 4 L1 _ | Sestd1 |
| Prss231 | 9.87E-07 | 1.357291 | 0.941 | 0.72 | 0.01987 | 4 L1 _ | Prss23 |
| Atp10a | 9.55E-07 | 0.331447 | 0.471 | 0.098 | 0.019228 | 4 L1 _ | Atp10a |
| Triobp1 | 8.70E-07 | 0.684939 | 0.765 | 0.273 | 0.017531 | 4 L1 _ | Triobp |
| Rack11 | 7.18E-07 | 0.737294 | 1 | 0.928 | 0.014461 | 4 L1 _ | Rack1 |
| Bicd11 | 7.03E-07 | 0.401253 | 0.294 | 0.044 | 0.014153 | 4 L1 _ | Bicd1 |
| Fgfr12 | 6.94E-07 | 0.969663 | 1 | 0.487 | 0.01398 | 4 L1 _ | Fgfr1 |
| Mcub | 6.66E-07 | 0.308252 | 0.353 | 0.059 | 0.013413 | 4 L1 _ | Mcub |
| Cpz1 | 6.11E-07 | 0.416222 | 0.471 | 0.095 | 0.012307 | 4 L1 _ | Cpz |
| Rps282 | 6.01E-07 | 0.718733 | 1 | 0.962 | 0.012108 | 4 L1 _ | Rps28 |
| Abca61 | 5.99E-07 | 0.473043 | 0.529 | 0.122 | 0.01207 | 4 L1 _ | Abca6 |
| Tmeff1 | 5.49E-07 | 0.394826 | 0.235 | 0.029 | 0.011052 | 4 L1 _ | Tmeff1 |
| Wnt11 | 5.12E-07 | 0.334305 | 0.235 | 0.029 | 0.010311 | 4 L1 _ | Wnt11 |
| Radil | 4.74E-07 | 0.295032 | 0.412 | 0.074 | 0.00954 | 4 L1 _ | Radil |
| Ptgs2 | 4.67E-07 | 0.318467 | 0.176 | 0.016 | 0.009408 | 4 L1 _ | Ptgs2 |
| Thbs21 | 4.45E-07 | 0.75465 | 1 | 0.374 | 0.008965 | 4 L1 _ | Thbs2 |
| Trnp1 | 3.95E-07 | 0.413477 | 0.353 | 0.059 | 0.007949 | 4 L1 _ | Trnp1 |
| Mgst13 | 3.93E-07 | 0.973787 | 1 | 0.498 | 0.007914 | 4 L1 _ | Mgst1 |
| Gm50012 | 3.79E-07 | 0.394996 | 0.235 | 0.028 | 0.007639 | 4 L1 _ | Gm50012 |
| Anks1 | 3.66E-07 | 0.4579 | 0.588 | 0.135 | 0.007374 | 4 L1 _ | Anks1 |
| Ehd11 | 3.60E-07 | 0.629939 | 0.882 | 0.286 | 0.007254 | 4 L1 _ | Ehd1 |
| Palm3 | 3.57E-07 | 0.764554 | 0.765 | 0.248 | 0.007188 | 4 L1 _ | Palm |
| Gsdme | 3.57E-07 | 0.293449 | 0.412 | 0.074 | 0.007181 | 4 L1 _ | Gsdme |
| Flrt22 | 3.46E-07 | 0.862576 | 0.941 | 0.376 | 0.006972 | 4 L1 _ | Flrt2 |
| Add32 | 2.87E-07 | 1.079693 | 1 | 0.505 | 0.005778 | 4 L1 _ | Add3 |

| | | | | | | | |
|---|---|---|---|---|---|---|---|
| Npc21 | 2.82E-07 | 0.957223 | 1 | 0.753 | 0.005677 | 4 L1 _ | Npc2 |
| Plat1 | 2.56E-07 | 0.963217 | 1 | 0.523 | 0.005161 | 4 L1 _ | Plat |
| Sh3d192 | 2.42E-07 | 0.905805 | 0.941 | 0.408 | 0.004865 | 4 L1 _ | Sh3d19 |
| Jpt11 | 2.39E-07 | 0.987368 | 1 | 0.378 | 0.004817 | 4 L1 _ | Jpt1 |
| Plce11 | 2.31E-07 | 0.764958 | 0.706 | 0.204 | 0.004654 | 4 L1 _ | Plce1 |
| Plagl1 | 2.21E-07 | 0.377398 | 0.353 | 0.058 | 0.004449 | 4 L1 _ | Plagl1 |
| Spsb1 | 1.92E-07 | 0.777457 | 0.706 | 0.204 | 0.003862 | 4 L1 _ | Spsb1 |
| Mcc2 | 1.83E-07 | 0.279491 | 0.706 | 0.17 | 0.003694 | 4 L1 _ | Mcc |
| Ddhd1 | 1.79E-07 | 0.522647 | 0.529 | 0.117 | 0.003604 | 4 L1 _ | Ddhd1 |
| Rab7b1 | 1.76E-07 | 0.571085 | 0.706 | 0.194 | 0.003544 | 4 L1 _ | Rab7b |
| C430049B03Rik1 | 1.64E-07 | 0.810824 | 0.294 | 0.041 | 0.003311 | 4 L1 _ | C430049B03Rik |
| Rps261 | 1.24E-07 | 0.83449 | 1 | 0.958 | 0.002506 | 4 L1 _ | Rps26 |
| Lacc1 | 1.11E-07 | 0.403572 | 0.471 | 0.091 | 0.002226 | 4 L1 _ | Lacc1 |
| Col1a12 | 1.00E-07 | 1.802513 | 1 | 0.672 | 0.002023 | 4 L1 _ | Col1a1 |
| Daglb1 | 9.78E-08 | 0.70984 | 0.471 | 0.095 | 0.00197 | 4 L1 _ | Daglb |
| Gpsm2 | 9.40E-08 | 0.357994 | 0.353 | 0.054 | 0.001893 | 4 L1 _ | Gpsm2 |
| Col1a22 | 9.37E-08 | 1.690726 | 1 | 0.749 | 0.001887 | 4 L1 _ | Col1a2 |
| Vgll31 | 9.37E-08 | 1.013289 | 0.824 | 0.31 | 0.001887 | 4 L1 _ | Vgll3 |
| Ogn2 | 9.28E-08 | 1.448634 | 1 | 0.544 | 0.001869 | 4 L1 _ | Ogn |
| Sod32 | 8.61E-08 | 1.062786 | 0.824 | 0.295 | 0.001734 | 4 L1 _ | Sod3 |
| Eef22 | 8.59E-08 | 0.825579 | 1 | 0.898 | 0.001729 | 4 L1 _ | Eef2 |
| Sulf11 | 8.20E-08 | 0.826682 | 0.941 | 0.346 | 0.001651 | 4 L1 _ | Sulf1 |
| Ssc5d1 | 8.15E-08 | 0.606653 | 0.588 | 0.145 | 0.001642 | 4 L1 _ | Ssc5d |
| Glce1 | 8.14E-08 | 0.687527 | 0.529 | 0.116 | 0.00164 | 4 L1 _ | Glce |
| Nmt2 | 7.99E-08 | 0.572633 | 0.765 | 0.229 | 0.00161 | 4 L1 _ | Nmt2 |
| Zfp976 | 7.74E-08 | 0.261417 | 0.235 | 0.025 | 0.001558 | 4 L1 _ | Zfp976 |
| Cacnb4 | 7.24E-08 | 0.267096 | 0.294 | 0.037 | 0.001458 | 4 L1 _ | Cacnb4 |
| Vasn1 | 6.79E-08 | 1.138175 | 0.824 | 0.297 | 0.001367 | 4 L1 _ | Vasn |
| Pdxk1 | 6.63E-08 | 0.451866 | 0.529 | 0.107 | 0.001336 | 4 L1 _ | Pdxk |
| Has2 | 6.10E-08 | 0.293058 | 0.294 | 0.038 | 0.001228 | 4 L1 _ | Has2 |
| Pdpn2 | 6.02E-08 | 1.225702 | 0.882 | 0.377 | 0.001213 | 4 L1 _ | Pdpn |
| Ndst2 | 5.93E-08 | 0.385828 | 0.412 | 0.07 | 0.001195 | 4 L1 _ | Ndst2 |
| Nqo22 | 5.78E-08 | 0.516991 | 0.588 | 0.137 | 0.001164 | 4 L1 _ | Nqo2 |
| Meg32 | 5.40E-08 | 1.584149 | 1 | 0.648 | 0.001088 | 4 L1 _ | Meg3 |
| Gm45187 | 5.23E-08 | 0.316938 | 0.176 | 0.014 | 0.001054 | 4 L1 _ | Gm45187 |
| Prcp | 4.46E-08 | 0.468886 | 0.588 | 0.13 | 0.000899 | 4 L1 _ | Prcp |
| Ttc39c1 | 4.24E-08 | 0.294473 | 0.412 | 0.066 | 0.000854 | 4 L1 _ | Ttc39c |
| Tmem1581 | 4.24E-08 | 0.835027 | 0.824 | 0.263 | 0.000854 | 4 L1 _ | Tmem158 |
| Rps151 | 4.12E-08 | 0.954493 | 1 | 0.951 | 0.000831 | 4 L1 _ | Rps15 |
| Plac82 | 3.96E-08 | 0.854616 | 1 | 0.361 | 0.000797 | 4 L1 _ | Plac8 |
| Hdac73 | 3.83E-08 | 1.147702 | 1 | 0.448 | 0.000771 | 4 L1 _ | Hdac7 |

| | | | | | | | |
|---|---|---|---|---|---|---|---|
| Rbpj1 | 3.32E-08 | 1.024413 | 1 | 0.442 | 0.00067 | 4 L1 _ | Rbpj |
| Plpp33 | 2.89E-08 | 1.383127 | 1 | 0.729 | 0.000581 | 4 L1 _ | Plpp3 |
| Agpat4 | 2.59E-08 | 0.995757 | 0.765 | 0.241 | 0.000522 | 4 L1 _ | Agpat4 |
| Ptprj | 2.55E-08 | 0.38856 | 0.471 | 0.083 | 0.000513 | 4 L1 _ | Ptprj |
| Sh3pxd2b2 | 2.32E-08 | 0.807654 | 0.941 | 0.324 | 0.000467 | 4 L1 _ | Sh3pxd2b |
| B3galnt11 | 2.11E-08 | 0.393667 | 0.471 | 0.082 | 0.000425 | 4 L1 _ | B3galnt1 |
| Arhgap422 | 2.05E-08 | 0.662543 | 0.765 | 0.199 | 0.000413 | 4 L1 _ | Arhgap42 |
| Ahnak21 | 2.05E-08 | 1.342545 | 0.882 | 0.35 | 0.000413 | 4 L1 _ | Ahnak2 |
| Six1 | 1.61E-08 | 0.293356 | 0.235 | 0.023 | 0.000323 | 4 L1 _ | Six1 |
| Gga2 | 1.51E-08 | 0.538568 | 0.647 | 0.149 | 0.000304 | 4 L1 _ | Gga2 |
| Rps121 | 1.49E-08 | 0.896084 | 1 | 0.974 | 0.0003 | 4 L1 _ | Rps12 |
| Ahnak4 | 1.37E-08 | 1.046011 | 1 | 0.894 | 0.000275 | 4 L1 _ | Ahnak |
| Notch21 | 1.28E-08 | 0.658108 | 0.765 | 0.21 | 0.000257 | 4 L1 _ | Notch2 |
| Kank11 | 1.22E-08 | 0.543485 | 0.588 | 0.122 | 0.000246 | 4 L1 _ | Kank1 |
| 9130008F23Rik | 1.17E-08 | 0.381259 | 0.353 | 0.048 | 0.000236 | 4 L1 _ | 9130008F23Rik |
| Alcam | 1.14E-08 | 0.286299 | 0.294 | 0.034 | 0.000231 | 4 L1 _ | Alcam |
| Disp1 | 1.05E-08 | 0.536434 | 0.471 | 0.083 | 0.000212 | 4 L1 _ | Disp1 |
| Fbln52 | 9.15E-09 | 0.997408 | 1 | 0.386 | 0.000184 | 4 L1 _ | Fbln5 |
| Col5a13 | 8.84E-09 | 1.329154 | 0.941 | 0.512 | 0.000178 | 4 L1 _ | Col5a1 |
| Col3a12 | 8.56E-09 | 2.08133 | 1 | 0.756 | 0.000172 | 4 L1 _ | Col3a1 |
| Dkk2 | 8.55E-09 | 0.502965 | 0.118 | 0.006 | 0.000172 | 4 L1 _ | Dkk2 |
| S100a111 | 8.37E-09 | 1.250021 | 1 | 0.863 | 0.000169 | 4 L1 _ | S100a11 |
| Fyn1 | 8.37E-09 | 1.254978 | 0.882 | 0.398 | 0.000169 | 4 L1 _ | Fyn |
| Igfbp52 | 8.29E-09 | 2.292937 | 0.941 | 0.515 | 0.000167 | 4 L1 _ | Igfbp5 |
| Adamts53 | 7.10E-09 | 1.472048 | 1 | 0.523 | 0.000143 | 4 L1 _ | Adamts5 |
| Ly6a2 | 6.82E-09 | 1.628061 | 1 | 0.731 | 0.000137 | 4 L1 _ | Ly6a |
| Adamtsl51 | 5.46E-09 | 0.414452 | 0.647 | 0.132 | 0.00011 | 4 L1 _ | Adamtsl5 |
| Plekho2 | 5.26E-09 | 0.911535 | 0.706 | 0.181 | 0.000106 | 4 L1 _ | Plekho2 |
| Fam102b2 | 5.18E-09 | 1.027984 | 0.882 | 0.313 | 0.000104 | 4 L1 _ | Fam102b |
| Gpnmb1 | 4.30E-09 | 0.415003 | 0.529 | 0.093 | 8.66E-05 | 4 L1 _ | Gpnmb |
| Slco3a12 | 4.28E-09 | 0.872541 | 0.941 | 0.319 | 8.63E-05 | 4 L1 _ | Slco3a1 |
| Loxl22 | 4.14E-09 | 1.036245 | 0.941 | 0.326 | 8.34E-05 | 4 L1 _ | Loxl2 |
| Ablim13 | 4.05E-09 | 1.21337 | 1 | 0.395 | 8.16E-05 | 4 L1 _ | Ablim1 |
| Ly6c12 | 4.05E-09 | 1.34386 | 1 | 0.423 | 8.15E-05 | 4 L1 _ | Ly6c1 |
| Cavin13 | 3.60E-09 | 1.265676 | 1 | 0.48 | 7.25E-05 | 4 L1 _ | Cavin1 |
| Lama41 | 3.48E-09 | 1.501146 | 1 | 0.517 | 7.00E-05 | 4 L1 _ | Lama4 |
| Ifi2052 | 3.02E-09 | 0.950907 | 0.941 | 0.294 | 6.08E-05 | 4 L1 _ | Ifi205 |
| Ugp2 | 2.42E-09 | 1.025285 | 0.882 | 0.314 | 4.87E-05 | 4 L1 _ | Ugp2 |
| Serpinb6a3 | 2.31E-09 | 1.315362 | 1 | 0.712 | 4.66E-05 | 4 L1 _ | Serpinb6a |
| Itsn11 | 2.24E-09 | 0.68963 | 0.824 | 0.214 | 4.51E-05 | 4 L1 _ | Itsn1 |
| Tbx181 | 1.99E-09 | 0.395186 | 0.412 | 0.058 | 4.02E-05 | 4 L1 _ | Tbx18 |

| | | | | | | | |
|---|---|---|---|---|---|---|---|
| Arhgap202 | 1.97E-09 | 0.987245 | 0.824 | 0.241 | 3.96E-05 | 4 L1 _ | Arhgap20 |
| Mfap52 | 1.86E-09 | 2.348267 | 0.941 | 0.535 | 3.75E-05 | 4 L1 _ | Mfap5 |
| Il1rl2 | 1.58E-09 | 1.030387 | 0.588 | 0.129 | 3.19E-05 | 4 L1 _ | Il1rl2 |
| Sntb22 | 1.47E-09 | 1.105992 | 1 | 0.549 | 2.96E-05 | 4 L1 _ | Sntb2 |
| Pde8a3 | 1.31E-09 | 0.874405 | 0.765 | 0.199 | 2.64E-05 | 4 L1 _ | Pde8a |
| Lims22 | 1.16E-09 | 0.735812 | 0.706 | 0.152 | 2.34E-05 | 4 L1 _ | Lims2 |
| Marcks2 | 1.10E-09 | 1.470108 | 1 | 0.823 | 2.21E-05 | 4 L1 _ | Marcks |
| S100a102 | 1.01E-09 | 1.656086 | 1 | 0.69 | 2.03E-05 | 4 L1 _ | S100a10 |
| Gngt22 | 9.43E-10 | 1.06836 | 0.882 | 0.257 | 1.90E-05 | 4 L1 _ | Gngt2 |
| St3gal2 | 8.36E-10 | 0.828949 | 0.882 | 0.266 | 1.68E-05 | 4 L1 _ | St3gal2 |
| Scara53 | 7.81E-10 | 1.593532 | 1 | 0.405 | 1.57E-05 | 4 L1 _ | Scara5 |
| Mmp22 | 7.73E-10 | 1.721493 | 1 | 0.56 | 1.56E-05 | 4 L1 _ | Mmp2 |
| Gm101151 | 6.34E-10 | 0.801989 | 0.588 | 0.115 | 1.28E-05 | 4 L1 _ | Gm10115 |
| Uap12 | 5.25E-10 | 1.003202 | 0.882 | 0.263 | 1.06E-05 | 4 L1 _ | Uap1 |
| Axl3 | 4.91E-10 | 1.411504 | 1 | 0.577 | 9.89E-06 | 4 L1 _ | Axl |
| Pla1a1 | 4.57E-10 | 2.011282 | 0.882 | 0.414 | 9.20E-06 | 4 L1 _ | Pla1a |
| Lurap1l2 | 3.13E-10 | 0.516089 | 0.471 | 0.068 | 6.31E-06 | 4 L1 _ | Lurap1l |
| Ppp1r14b1 | 3.00E-10 | 1.869972 | 1 | 0.483 | 6.05E-06 | 4 L1 _ | Ppp1r14b |
| Mrgprf | 2.74E-10 | 0.395954 | 0.588 | 0.098 | 5.51E-06 | 4 L1 _ | Mrgprf |
| Bmp13 | 2.38E-10 | 1.566358 | 0.941 | 0.439 | 4.78E-06 | 4 L1 _ | Bmp1 |
| Lvrn | 2.19E-10 | 0.596819 | 0.471 | 0.071 | 4.41E-06 | 4 L1 _ | Lvrn |
| Basp13 | 2.19E-10 | 1.42166 | 0.941 | 0.373 | 4.41E-06 | 4 L1 _ | Basp1 |
| Aebp11 | 1.91E-10 | 1.80296 | 1 | 0.629 | 3.84E-06 | 4 L1 _ | Aebp1 |
| Klhl132 | 1.05E-10 | 0.704378 | 0.706 | 0.148 | 2.12E-06 | 4 L1 _ | Klhl13 |
| Dcbld2 | 8.11E-11 | 0.812964 | 0.882 | 0.227 | 1.63E-06 | 4 L1 _ | Dcbld2 |
| Nt5dc3 | 6.61E-11 | 0.258187 | 0.353 | 0.037 | 1.33E-06 | 4 L1 _ | Nt5dc3 |
| Hspb81 | 5.50E-11 | 0.585444 | 0.647 | 0.118 | 1.11E-06 | 4 L1 _ | Hspb8 |
| Lmnb1 | 5.02E-11 | 0.469036 | 0.471 | 0.066 | 1.01E-06 | 4 L1 _ | Lmnb1 |
| Kctd9 | 4.88E-11 | 0.537135 | 0.529 | 0.082 | 9.83E-07 | 4 L1 _ | Kctd9 |
| Tgfbr23 | 4.38E-11 | 1.700508 | 1 | 0.651 | 8.83E-07 | 4 L1 _ | Tgfbr2 |
| Poglut31 | 3.97E-11 | 0.991028 | 0.882 | 0.245 | 8.00E-07 | 4 L1 _ | Poglut3 |
| Dnase2a | 3.85E-11 | 0.618937 | 0.647 | 0.12 | 7.75E-07 | 4 L1 _ | Dnase2a |
| Gan | 3.78E-11 | 0.703689 | 0.647 | 0.122 | 7.62E-07 | 4 L1 _ | Gan |
| Ecm12 | 2.78E-11 | 2.344265 | 0.941 | 0.48 | 5.60E-07 | 4 L1 _ | Ecm1 |
| Sema3d | 2.41E-11 | 0.645867 | 0.529 | 0.082 | 4.86E-07 | 4 L1 _ | Sema3d |
| Gxylt21 | 2.36E-11 | 0.913674 | 0.765 | 0.173 | 4.75E-07 | 4 L1 _ | Gxylt2 |
| Anxa13 | 1.69E-11 | 1.994467 | 1 | 0.571 | 3.40E-07 | 4 L1 _ | Anxa1 |
| Ttyh2 | 1.54E-11 | 0.558197 | 0.588 | 0.094 | 3.09E-07 | 4 L1 _ | Ttyh2 |
| Nid13 | 1.25E-11 | 2.022071 | 1 | 0.733 | 2.51E-07 | 4 L1 _ | Nid1 |
| Zyx2 | 1.11E-11 | 1.421526 | 1 | 0.433 | 2.23E-07 | 4 L1 _ | Zyx |
| C33 | 9.27E-12 | 2.684961 | 1 | 0.395 | 1.87E-07 | 4 L1 _ | C3 |

| | | | | | | | |
|---|---|---|---|---|---|---|---|
| Gnpnat1 | 8.87E-12 | 0.894615 | 0.824 | 0.197 | 1.79E-07 | 4 L1 _ | Gnpnat1 |
| Epdr1 | 8.10E-12 | 0.577854 | 0.471 | 0.062 | 1.63E-07 | 4 L1 _ | Epdr1 |
| Dclk12 | 7.47E-12 | 1.068478 | 0.882 | 0.216 | 1.51E-07 | 4 L1 _ | Dclk1 |
| Eya4 | 6.69E-12 | 0.329875 | 0.235 | 0.016 | 1.35E-07 | 4 L1 _ | Eya4 |
| Fstl12 | 5.44E-12 | 2.087933 | 1 | 0.698 | 1.10E-07 | 4 L1 _ | Fstl1 |
| Grb102 | 5.25E-12 | 1.238077 | 1 | 0.353 | 1.06E-07 | 4 L1 _ | Grb10 |
| Sulf22 | 5.10E-12 | 0.834476 | 0.882 | 0.2 | 1.03E-07 | 4 L1 _ | Sulf2 |
| Rian | 4.77E-12 | 1.533597 | 0.765 | 0.177 | 9.60E-08 | 4 L1 _ | Rian |
| Gfpt23 | 2.90E-12 | 1.471783 | 1 | 0.317 | 5.84E-08 | 4 L1 _ | Gfpt2 |
| Sec22c | 2.74E-12 | 0.465725 | 0.412 | 0.046 | 5.52E-08 | 4 L1 _ | Sec22c |
| Cd344 | 2.72E-12 | 2.141541 | 1 | 0.445 | 5.47E-08 | 4 L1 _ | Cd34 |
| Plxna11 | 1.98E-12 | 0.753371 | 0.882 | 0.199 | 3.98E-08 | 4 L1 _ | Plxna1 |
| Tmem1004 | 1.73E-12 | 1.752872 | 0.941 | 0.282 | 3.48E-08 | 4 L1 _ | Tmem100 |
| Cd276 | 1.22E-12 | 0.389117 | 0.412 | 0.045 | 2.46E-08 | 4 L1 _ | Cd276 |
| Dbn11 | 1.12E-12 | 1.468456 | 0.882 | 0.258 | 2.25E-08 | 4 L1 _ | Dbn1 |
| Itpr32 | 1.10E-12 | 0.601959 | 0.588 | 0.087 | 2.22E-08 | 4 L1 _ | Itpr3 |
| Timp23 | 1.07E-12 | 2.853402 | 1 | 0.704 | 2.15E-08 | 4 L1 _ | Timp2 |
| Adam331 | 8.53E-13 | 0.572745 | 0.412 | 0.044 | 1.72E-08 | 4 L1 _ | Adam33 |
| B830012L14Rik | 7.87E-13 | 0.465476 | 0.412 | 0.044 | 1.58E-08 | 4 L1 _ | B830012L14Rik |
| Igfbp62 | 6.59E-13 | 2.748685 | 1 | 0.486 | 1.33E-08 | 4 L1 _ | Igfbp6 |
| Medag3 | 5.18E-13 | 1.643665 | 1 | 0.329 | 1.04E-08 | 4 L1 _ | Medag |
| Npr13 | 4.85E-13 | 0.826511 | 0.706 | 0.12 | 9.76E-09 | 4 L1 _ | Npr1 |
| Ace5 | 3.50E-13 | 1.719893 | 1 | 0.394 | 7.05E-09 | 4 L1 _ | Ace |
| Fbn13 | 3.34E-13 | 3.033151 | 1 | 0.62 | 6.73E-09 | 4 L1 _ | Fbn1 |
| Sdk11 | 3.19E-13 | 0.330933 | 0.412 | 0.041 | 6.42E-09 | 4 L1 _ | Sdk1 |
| Heg13 | 2.55E-13 | 1.58578 | 1 | 0.335 | 5.13E-09 | 4 L1 _ | Heg1 |
| Atp1b12 | 2.49E-13 | 0.939554 | 0.706 | 0.122 | 5.02E-09 | 4 L1 _ | Atp1b1 |
| Xpnpep21 | 2.07E-13 | 0.489422 | 0.529 | 0.066 | 4.17E-09 | 4 L1 _ | Xpnpep2 |
| Bmp61 | 1.79E-13 | 0.50086 | 0.412 | 0.043 | 3.60E-09 | 4 L1 _ | Bmp6 |
| Ddr22 | 1.46E-13 | 1.861772 | 1 | 0.44 | 2.94E-09 | 4 L1 _ | Ddr2 |
| Tnfaip61 | 1.10E-13 | 0.938984 | 0.765 | 0.147 | 2.22E-09 | 4 L1 _ | Tnfaip6 |
| Col14a13 | 5.19E-14 | 2.73025 | 1 | 0.456 | 1.04E-09 | 4 L1 _ | Col14a1 |
| Adamts23 | 4.45E-14 | 1.80584 | 0.941 | 0.305 | 8.97E-10 | 4 L1 _ | Adamts2 |
| Galnt163 | 3.67E-14 | 0.788785 | 0.765 | 0.126 | 7.39E-10 | 4 L1 _ | Galnt16 |
| Gm48617 | 3.21E-14 | 0.260424 | 0.235 | 0.013 | 6.46E-10 | 4 L1 _ | Gm48617 |
| Has2os | 2.97E-14 | 0.345562 | 0.353 | 0.029 | 5.98E-10 | 4 L1 _ | Has2os |
| Tek3 | 2.29E-14 | 0.981087 | 0.882 | 0.161 | 4.61E-10 | 4 L1 _ | Tek |
| Cul9 | 2.28E-14 | 0.623022 | 0.588 | 0.078 | 4.60E-10 | 4 L1 _ | Cul9 |
| Car83 | 1.34E-14 | 1.023029 | 0.529 | 0.065 | 2.70E-10 | 4 L1 _ | Car8 |
| Irak3 | 1.20E-14 | 0.845708 | 0.882 | 0.168 | 2.42E-10 | 4 L1 _ | Irak3 |
| Gm9780 | 1.17E-14 | 0.280988 | 0.353 | 0.028 | 2.35E-10 | 4 L1 _ | Gm9780 |

| | | | | | | | |
|---|---|---|---|---|---|---|---|
| Sirpa1 | 9.25E-15 | 0.695273 | 0.588 | 0.079 | 1.86E-10 | 4 L1 _ | Sirpa |
| Clec3b3 | 7.53E-15 | 3.315716 | 1 | 0.433 | 1.52E-10 | 4 L1 _ | Clec3b |
| Klf43 | 5.51E-15 | 2.339315 | 1 | 0.311 | 1.11E-10 | 4 L1 _ | Klf4 |
| Cd2482 | 5.25E-15 | 2.939691 | 1 | 0.421 | 1.06E-10 | 4 L1 _ | Cd248 |
| Efemp13 | 4.33E-15 | 2.665256 | 1 | 0.353 | 8.73E-11 | 4 L1 _ | Efemp1 |
| Ebf23 | 4.14E-15 | 1.495473 | 0.882 | 0.174 | 8.33E-11 | 4 L1 _ | Ebf2 |
| Igdcc4 | 4.10E-15 | 0.420777 | 0.471 | 0.048 | 8.25E-11 | 4 L1 _ | Igdcc4 |
| Glipr22 | 4.05E-15 | 0.886252 | 0.824 | 0.154 | 8.15E-11 | 4 L1 _ | Glipr2 |
| Mmp271 | 3.44E-15 | 0.288127 | 0.235 | 0.012 | 6.93E-11 | 4 L1 _ | Mmp27 |
| Gas72 | 3.17E-15 | 2.525652 | 1 | 0.351 | 6.37E-11 | 4 L1 _ | Gas7 |
| Mmp232 | 2.68E-15 | 1.491312 | 0.882 | 0.204 | 5.40E-11 | 4 L1 _ | Mmp23 |
| Rhou2 | 1.07E-15 | 0.825292 | 0.824 | 0.138 | 2.16E-11 | 4 L1 _ | Rhou |
| Procr2 | 7.93E-16 | 1.191487 | 0.765 | 0.123 | 1.60E-11 | 4 L1 _ | Procr |
| Antxr12 | 7.77E-16 | 1.963724 | 1 | 0.302 | 1.57E-11 | 4 L1 _ | Antxr1 |
| Dact11 | 6.79E-16 | 1.141566 | 0.941 | 0.204 | 1.37E-11 | 4 L1 _ | Dact1 |
| Man1a2 | 6.40E-16 | 1.819727 | 1 | 0.333 | 1.29E-11 | 4 L1 _ | Man1a |
| Tubb2b1 | 6.38E-16 | 0.964885 | 0.765 | 0.122 | 1.28E-11 | 4 L1 _ | Tubb2b |
| Csrp21 | 5.08E-16 | 2.131391 | 1 | 0.286 | 1.02E-11 | 4 L1 _ | Csrp2 |
| Cdon2 | 3.92E-16 | 1.234215 | 0.706 | 0.106 | 7.89E-12 | 4 L1 _ | Cdon |
| Tubb4a2 | 2.13E-16 | 0.414017 | 0.353 | 0.025 | 4.30E-12 | 4 L1 _ | Tubb4a |
| Gprc5b1 | 1.62E-16 | 0.765337 | 0.588 | 0.07 | 3.27E-12 | 4 L1 _ | Gprc5b |
| AW551984 | 1.60E-16 | 0.77146 | 0.529 | 0.057 | 3.23E-12 | 4 L1 _ | AW551984 |
| Ugdh2 | 1.46E-16 | 2.602996 | 0.941 | 0.24 | 2.94E-12 | 4 L1 _ | Ugdh |
| Zfpm2 | 1.22E-16 | 0.767278 | 0.647 | 0.086 | 2.45E-12 | 4 L1 _ | Zfpm2 |
| C1qtnf12 | 1.16E-16 | 1.125706 | 0.882 | 0.162 | 2.33E-12 | 4 L1 _ | C1qtnf1 |
| Bdh21 | 9.99E-17 | 0.504387 | 0.529 | 0.053 | 2.01E-12 | 4 L1 _ | Bdh2 |
| Sytl23 | 9.97E-17 | 0.426518 | 0.647 | 0.075 | 2.01E-12 | 4 L1 _ | Sytl2 |
| Rras2 | 6.99E-17 | 0.853575 | 0.706 | 0.097 | 1.41E-12 | 4 L1 _ | Rras2 |
| Fez12 | 6.51E-17 | 0.513539 | 0.588 | 0.065 | 1.31E-12 | 4 L1 _ | Fez1 |
| Anxa34 | 6.03E-17 | 2.871307 | 1 | 0.311 | 1.22E-12 | 4 L1 _ | Anxa3 |
| Scara31 | 3.18E-17 | 1.887871 | 0.941 | 0.232 | 6.40E-13 | 4 L1 _ | Scara3 |
| Hebp2 | 2.43E-17 | 0.451512 | 0.588 | 0.063 | 4.89E-13 | 4 L1 _ | Hebp2 |
| Metrnl | 7.61E-18 | 1.789003 | 1 | 0.26 | 1.53E-13 | 4 L1 _ | Metrnl |
| Zfp385a1 | 5.81E-18 | 1.33047 | 0.765 | 0.121 | 1.17E-13 | 4 L1 _ | Zfp385a |
| Pi162 | 2.95E-18 | 3.569863 | 1 | 0.263 | 5.95E-14 | 4 L1 _ | Pi16 |
| Thbs31 | 1.95E-18 | 1.250653 | 0.882 | 0.153 | 3.92E-14 | 4 L1 _ | Thbs3 |
| Osr12 | 2.31E-19 | 1.96781 | 0.941 | 0.188 | 4.64E-15 | 4 L1 _ | Osr1 |
| Srgap12 | 2.54E-20 | 0.927014 | 0.647 | 0.069 | 5.11E-16 | 4 L1 _ | Srgap1 |
| Olfml2b3 | 2.33E-20 | 1.219085 | 0.882 | 0.126 | 4.69E-16 | 4 L1 _ | Olfml2b |
| Tnxb2 | 1.59E-20 | 3.241293 | 1 | 0.235 | 3.20E-16 | 4 L1 _ | Tnxb |
| Flnc2 | 6.99E-21 | 0.954711 | 0.824 | 0.116 | 1.41E-16 | 4 L1 _ | Flnc |

| | | | | | | | |
|---|---|---|---|---|---|---|---|
| Lgi21 | 6.55E-21 | 1.555193 | 0.824 | 0.12 | 1.32E-16 | 4 L1 _ | Lgi2 |
| C4b2 | 4.01E-21 | 2.122394 | 0.706 | 0.084 | 8.08E-17 | 4 L1 _ | C4b |
| Tppp32 | 3.99E-21 | 1.860873 | 1 | 0.187 | 8.04E-17 | 4 L1 _ | Tppp3 |
| Twist2 | 3.60E-21 | 0.593676 | 0.529 | 0.044 | 7.26E-17 | 4 L1 _ | Twist2 |
| 5730507C01Rik | 3.46E-21 | 0.320508 | 0.235 | 0.008 | 6.97E-17 | 4 L1 _ | 5730507C01Rik |
| Lamc21 | 3.13E-21 | 0.439028 | 0.235 | 0.008 | 6.31E-17 | 4 L1 _ | Lamc2 |
| Nhsl11 | 1.96E-21 | 0.989859 | 0.765 | 0.095 | 3.95E-17 | 4 L1 _ | Nhsl1 |
| Mustn13 | 5.57E-22 | 1.224018 | 0.882 | 0.116 | 1.12E-17 | 4 L1 _ | Mustn1 |
| Nox41 | 5.26E-22 | 0.93541 | 0.706 | 0.075 | 1.06E-17 | 4 L1 _ | Nox4 |
| Kif21a1 | 3.32E-22 | 0.502649 | 0.412 | 0.025 | 6.68E-18 | 4 L1 _ | Kif21a |
| Gm12122 | 1.06E-22 | 0.402596 | 0.471 | 0.031 | 2.14E-18 | 4 L1 _ | Gm12122 |
| Arl4d2 | 4.78E-23 | 0.710691 | 0.588 | 0.049 | 9.62E-19 | 4 L1 _ | Arl4d |
| Megf91 | 3.49E-23 | 0.730176 | 0.706 | 0.07 | 7.04E-19 | 4 L1 _ | Megf9 |
| Pi15 | 3.28E-23 | 1.572167 | 0.765 | 0.088 | 6.61E-19 | 4 L1 _ | Pi15 |
| Il6ra1 | 2.48E-23 | 0.528055 | 0.471 | 0.031 | 4.99E-19 | 4 L1 _ | Il6ra |
| Ildr21 | 1.48E-23 | 1.625999 | 0.765 | 0.091 | 2.98E-19 | 4 L1 _ | Ildr2 |
| Ndrg41 | 8.75E-24 | 0.493173 | 0.471 | 0.03 | 1.76E-19 | 4 L1 _ | Ndrg4 |
| Wnt10b | 8.36E-24 | 0.429525 | 0.412 | 0.023 | 1.68E-19 | 4 L1 _ | Wnt10b |
| Arsi1 | 3.79E-24 | 0.906493 | 0.588 | 0.049 | 7.63E-20 | 4 L1 _ | Arsi |
| Rab32 | 2.16E-24 | 0.672663 | 0.529 | 0.038 | 4.35E-20 | 4 L1 _ | Rab32 |
| Qpct1 | 3.58E-25 | 1.536949 | 0.882 | 0.119 | 7.21E-21 | 4 L1 _ | Qpct |
| Gpc32 | 2.41E-25 | 1.54556 | 0.765 | 0.08 | 4.86E-21 | 4 L1 _ | Gpc3 |
| Sel1l3 | 2.33E-25 | 0.375309 | 0.176 | 0.003 | 4.68E-21 | 4 L1 _ | Sel1l3 |
| Pygl1 | 2.07E-25 | 0.915745 | 0.706 | 0.066 | 4.18E-21 | 4 L1 _ | Pygl |
| Wnt21 | 1.69E-25 | 1.156696 | 0.647 | 0.056 | 3.39E-21 | 4 L1 _ | Wnt2 |
| Prdm81 | 1.42E-25 | 0.622075 | 0.412 | 0.022 | 2.86E-21 | 4 L1 _ | Prdm8 |
| Emilin22 | 2.30E-26 | 2.544684 | 1 | 0.169 | 4.63E-22 | 4 L1 _ | Emilin2 |
| Itga111 | 9.99E-27 | 1.058857 | 0.765 | 0.075 | 2.01E-22 | 4 L1 _ | Itga11 |
| Dpt2 | 2.97E-27 | 4.387907 | 1 | 0.165 | 5.98E-23 | 4 L1 _ | Dpt |
| Fn12 | 7.88E-28 | 4.737653 | 1 | 0.159 | 1.59E-23 | 4 L1 _ | Fn1 |
| Cadm32 | 1.07E-28 | 0.941645 | 0.706 | 0.056 | 2.15E-24 | 4 L1 _ | Cadm3 |
| Stxbp61 | 1.83E-30 | 0.531314 | 0.588 | 0.035 | 3.69E-26 | 4 L1 _ | Stxbp6 |
| Gda3 | 1.71E-30 | 1.514098 | 0.824 | 0.078 | 3.45E-26 | 4 L1 _ | Gda |
| Ackr33 | 7.08E-32 | 2.375167 | 1 | 0.12 | 1.43E-27 | 4 L1 _ | Ackr3 |
| Slc4a41 | 3.48E-32 | 1.050363 | 0.647 | 0.044 | 7.01E-28 | 4 L1 _ | Slc4a4 |
| Limch12 | 3.09E-32 | 1.265872 | 0.882 | 0.08 | 6.22E-28 | 4 L1 _ | Limch1 |
| 5033421B08Rik1 | 1.76E-32 | 0.335146 | 0.235 | 0.005 | 3.55E-28 | 4 L1 _ | 5033421B08Rik |
| Pamr11 | 9.29E-33 | 1.798363 | 0.882 | 0.088 | 1.87E-28 | 4 L1 _ | Pamr1 |
| Tnfrsf11b | 2.69E-34 | 0.355425 | 0.412 | 0.015 | 5.42E-30 | 4 L1 _ | Tnfrsf11b |
| Stmn41 | 1.13E-34 | 0.335737 | 0.235 | 0.004 | 2.28E-30 | 4 L1 _ | Stmn4 |
| Sema3c3 | 4.81E-35 | 3.228118 | 1 | 0.116 | 9.69E-31 | 4 L1 _ | Sema3c |

| | | | | | | | |
|---|---|---|---|---|---|---|---|
| Cd552 | 1.35E-35 | 2.641034 | 1 | 0.114 | 2.72E-31 | 4 L1 _ | Cd55 |
| Fhl13 | 9.78E-36 | 1.094054 | 0.882 | 0.07 | 1.97E-31 | 4 L1 _ | Fhl1 |
| Hoxb31 | 3.13E-37 | 0.280678 | 0.235 | 0.004 | 6.30E-33 | 4 L1 _ | Hoxb3 |
| Itgb3 | 4.04E-38 | 0.880249 | 0.882 | 0.069 | 8.15E-34 | 4 L1 _ | Itgb3 |
| Fndc11 | 3.32E-38 | 3.527999 | 1 | 0.106 | 6.69E-34 | 4 L1 _ | Fndc1 |
| Pcdh20 | 2.24E-38 | 0.598472 | 0.353 | 0.009 | 4.51E-34 | 4 L1 _ | Pcdh20 |
| Dmkn1 | 1.84E-39 | 1.154061 | 0.176 | 0.002 | 3.70E-35 | 4 L1 _ | Dmkn |
| Adra1b1 | 2.47E-40 | 0.27629 | 0.235 | 0.003 | 4.97E-36 | 4 L1 _ | Adra1b |
| Krtdap | 3.27E-41 | 1.569889 | 0.118 | 0 | 6.58E-37 | 4 L1 _ | Krtdap |
| Vldlr2 | 2.04E-41 | 1.492355 | 0.882 | 0.064 | 4.11E-37 | 4 L1 _ | Vldlr |
| Ackr11 | 1.01E-41 | 0.461857 | 0.471 | 0.016 | 2.04E-37 | 4 L1 _ | Ackr1 |
| Sfrp22 | 5.19E-43 | 3.42122 | 1 | 0.089 | 1.05E-38 | 4 L1 _ | Sfrp2 |
| Kcnk51 | 3.49E-43 | 0.634727 | 0.588 | 0.025 | 7.02E-39 | 4 L1 _ | Kcnk5 |
| Chrdl11 | 1.01E-44 | 0.717095 | 0.353 | 0.008 | 2.02E-40 | 4 L1 _ | Chrdl1 |
| Vegfd1 | 3.61E-45 | 0.717319 | 0.588 | 0.024 | 7.28E-41 | 4 L1 _ | Vegfd |
| Tbx5 | 7.86E-46 | 0.308476 | 0.176 | 0.001 | 1.58E-41 | 4 L1 _ | Tbx5 |
| Fzd10 | 7.86E-46 | 0.262312 | 0.176 | 0.001 | 1.58E-41 | 4 L1 _ | Fzd10 |
| Ano3 | 6.59E-46 | 0.319128 | 0.176 | 0.001 | 1.33E-41 | 4 L1 _ | Ano3 |
| Pcsk62 | 2.06E-46 | 2.2421 | 0.882 | 0.059 | 4.15E-42 | 4 L1 _ | Pcsk6 |
| Ackr21 | 5.29E-47 | 0.574672 | 0.353 | 0.007 | 1.07E-42 | 4 L1 _ | Ackr2 |
| Clip41 | 4.61E-48 | 1.241983 | 0.824 | 0.047 | 9.29E-44 | 4 L1 _ | Clip4 |
| Rnf165 | 1.51E-49 | 0.622857 | 0.529 | 0.017 | 3.03E-45 | 4 L1 _ | Rnf165 |
| Cmah2 | 1.42E-52 | 1.581066 | 0.882 | 0.049 | 2.86E-48 | 4 L1 _ | Cmah |
| Gm267711 | 5.65E-53 | 0.766776 | 0.588 | 0.02 | 1.14E-48 | 4 L1 _ | Gm26771 |
| Sema3b | 1.22E-53 | 0.919961 | 0.529 | 0.016 | 2.45E-49 | 4 L1 _ | Sema3b |
| Gpr11 | 6.13E-54 | 0.417401 | 0.294 | 0.004 | 1.23E-49 | 4 L1 _ | Gpr1 |
| Adamts16 | 6.81E-55 | 0.284603 | 0.176 | 0.001 | 1.37E-50 | 4 L1 _ | Adamts16 |
| Ston21 | 3.98E-55 | 0.983861 | 0.588 | 0.019 | 8.02E-51 | 4 L1 _ | Ston2 |
| Lrrc172 | 2.21E-55 | 1.813236 | 1 | 0.063 | 4.44E-51 | 4 L1 _ | Lrrc17 |
| Sbsn1 | 2.83E-56 | 2.853742 | 0.941 | 0.055 | 5.70E-52 | 4 L1 _ | Sbsn |
| Lrrn4cl1 | 1.46E-57 | 1.712321 | 0.824 | 0.039 | 2.95E-53 | 4 L1 _ | Lrrn4cl |
| Dact21 | 1.50E-59 | 1.36252 | 0.706 | 0.026 | 3.02E-55 | 4 L1 _ | Dact2 |
| Smpd31 | 1.17E-59 | 2.334909 | 0.471 | 0.01 | 2.35E-55 | 4 L1 _ | Smpd3 |
| Gm156751 | 7.50E-60 | 0.325077 | 0.353 | 0.005 | 1.51E-55 | 4 L1 _ | Gm15675 |
| Sfrp41 | 1.23E-61 | 1.701221 | 0.588 | 0.016 | 2.49E-57 | 4 L1 _ | Sfrp4 |
| Ccn3 | 5.12E-62 | 1.303772 | 0.588 | 0.016 | 1.03E-57 | 4 L1 _ | Ccn3 |
| Cthrc1 | 1.74E-62 | 1.52729 | 0.588 | 0.016 | 3.50E-58 | 4 L1 _ | Cthrc1 |
| Creb51 | 1.24E-62 | 2.845729 | 1 | 0.056 | 2.50E-58 | 4 L1 _ | Creb5 |
| En11 | 6.50E-63 | 0.573903 | 0.294 | 0.003 | 1.31E-58 | 4 L1 _ | En1 |
| Gap43 | 3.55E-63 | 0.765104 | 0.471 | 0.009 | 7.14E-59 | 4 L1 _ | Gap43 |
| Efhd13 | 1.01E-65 | 1.5986 | 0.882 | 0.037 | 2.03E-61 | 4 L1 _ | Efhd1 |

| | | | | | | | |
|---|---|---|---|---|---|---|---|
| Gm15902 | 3.08E-67 | 0.402922 | 0.353 | 0.004 | 6.21E-63 | 4 L1 _ | Gm15902 |
| Rorb1 | 1.28E-67 | 0.760623 | 0.353 | 0.004 | 2.58E-63 | 4 L1 _ | Rorb |
| 1700019D03Rik1 | 4.00E-74 | 1.289143 | 0.765 | 0.024 | 8.06E-70 | 4 L1 _ | 1700019D03Rik |
| Efna51 | 2.24E-74 | 0.805183 | 0.588 | 0.013 | 4.52E-70 | 4 L1 _ | Efna5 |
| Adgrg21 | 2.15E-75 | 0.413009 | 0.294 | 0.002 | 4.32E-71 | 4 L1 _ | Adgrg2 |
| Dmrt2 | 1.80E-75 | 0.440102 | 0.294 | 0.002 | 3.62E-71 | 4 L1 _ | Dmrt2 |
| Edn12 | 5.20E-76 | 1.267375 | 0.824 | 0.026 | 1.05E-71 | 4 L1 _ | Edn1 |
| Adamtsl41 | 4.51E-83 | 1.377788 | 0.941 | 0.033 | 9.09E-79 | 4 L1 _ | Adamtsl4 |
| Tmeff21 | 4.04E-83 | 1.259145 | 0.824 | 0.024 | 8.14E-79 | 4 L1 _ | Tmeff2 |
| Adgrd11 | 3.95E-83 | 1.868371 | 1 | 0.039 | 7.95E-79 | 4 L1 _ | Adgrd1 |
| Il181 | 1.44E-84 | 0.898147 | 0.765 | 0.02 | 2.91E-80 | 4 L1 _ | Il18 |
| Aldh1a31 | 3.66E-87 | 1.272624 | 0.765 | 0.019 | 7.37E-83 | 4 L1 _ | Aldh1a3 |
| Fzd10os | 3.43E-92 | 0.567243 | 0.412 | 0.004 | 6.91E-88 | 4 L1 _ | Fzd10os |
| Aif1l1 | 3.78E-94 | 0.952993 | 0.765 | 0.017 | 7.60E-90 | 4 L1 _ | Aif1l |
| Sema3e | 3.89E-95 | 0.663828 | 0.471 | 0.005 | 7.83E-91 | 4 L1 _ | Sema3e |
| Dpp41 | 4.95E-104 | 2.708736 | 1 | 0.03 | 9.98E-100 | 4 L1 _ | Dpp4 |
| Nova11 | 9.33E-111 | 1.5914 | 0.824 | 0.017 | 1.88E-106 | 4 L1 _ | Nova1 |
| Opcml1 | 5.46E-114 | 1.975968 | 0.941 | 0.022 | 1.10E-109 | 4 L1 _ | Opcml |
| Gabra2 | 8.32E-121 | 0.343529 | 0.235 | 0 | 1.68E-116 | 4 L1 _ | Gabra2 |
| Krt19 | 1.22E-133 | 1.006143 | 0.412 | 0.002 | 2.45E-129 | 4 L1 _ | Krt19 |
| Duoxa11 | 3.41E-135 | 0.917152 | 0.529 | 0.004 | 6.87E-131 | 4 L1 _ | Duoxa1 |
| Pcolce21 | 6.45E-142 | 3.118156 | 1 | 0.019 | 1.30E-137 | 4 L1 _ | Pcolce2 |
| Mrgprg | 1.41E-150 | 0.713379 | 0.294 | 0 | 2.84E-146 | 4 L1 _ | Mrgprg |
| Dhrs9 | 1.25E-154 | 0.573111 | 0.353 | 0 | 2.53E-150 | 4 L1 _ | Dhrs9 |
| Lgi3 | 6.10E-160 | 0.533451 | 0.471 | 0.002 | 1.23E-155 | 4 L1 _ | Lgi3 |
| Bmp31 | 1.61E-172 | 0.999039 | 0.647 | 0.004 | 3.24E-168 | 4 L1 _ | Bmp3 |
| Gm43050 | 2.29E-180 | 0.431297 | 0.353 | 0 | 4.62E-176 | 4 L1 _ | Gm43050 |
| Krt36 | 2.29E-180 | 0.369443 | 0.353 | 0 | 4.62E-176 | 4 L1 _ | Krt36 |
| Duox11 | 2.10E-186 | 0.746668 | 0.529 | 0.002 | 4.24E-182 | 4 L1 _ | Duox1 |
| Islr21 | 1.65E-187 | 1.146162 | 0.765 | 0.006 | 3.32E-183 | 4 L1 _ | Islr2 |
| Akr1c18 | 0 | 3.021516 | 0.824 | 0.001 | 0 | 4 L1 _ | Akr1c18 |
| Itgb71 | 2.23E-06 | 0.68378 | 0.5 | 0.084 | 0.044978 | 4 M1 _ | Itgb7 |
| Basp11 | 2.15E-06 | 1.010263 | 1 | 0.375 | 0.043358 | 4 M1 _ | Basp1 |
| Col5a11 | 2.15E-06 | 1.500699 | 1 | 0.513 | 0.043209 | 4 M1 _ | Col5a1 |
| Gstm12 | 2.14E-06 | 1.747277 | 1 | 0.665 | 0.043076 | 4 M1 _ | Gstm1 |
| Fam126a1 | 2.07E-06 | 0.789595 | 0.7 | 0.164 | 0.041731 | 4 M1 _ | Fam126a |
| Antxr21 | 2.00E-06 | 1.73798 | 0.9 | 0.394 | 0.040324 | 4 M1 _ | Antxr2 |
| Lrrc8b1 | 2.00E-06 | 0.514652 | 0.4 | 0.053 | 0.040219 | 4 M1 _ | Lrrc8b |
| Egfr1 | 1.98E-06 | 1.012504 | 0.9 | 0.311 | 0.039818 | 4 M1 _ | Egfr |
| Sytl21 | 1.90E-06 | 0.383055 | 0.5 | 0.077 | 0.038242 | 4 M1 _ | Sytl2 |
| Ndrg4 | 1.88E-06 | 0.414759 | 0.3 | 0.032 | 0.037854 | 4 M1 _ | Ndrg4 |

| | | | | | | | |
|---|---|---|---|---|---|---|---|
| Adamts51 | 1.84E-06 | 1.812988 | 1 | 0.524 | 0.037043 | 4 M1 _ | Adamts5 |
| Bdh2 | 1.83E-06 | 0.557715 | 0.4 | 0.055 | 0.036763 | 4 M1 _ | Bdh2 |
| Zyx | 1.81E-06 | 1.364988 | 1 | 0.435 | 0.036421 | 4 M1 _ | Zyx |
| Glce | 1.71E-06 | 0.71429 | 0.6 | 0.117 | 0.034352 | 4 M1 _ | Glce |
| Ccdc174 | 1.62E-06 | 0.911141 | 1 | 0.334 | 0.032645 | 4 M1 _ | Ccdc174 |
| Il17ra | 1.47E-06 | 0.825026 | 0.6 | 0.12 | 0.029562 | 4 M1 _ | Il17ra |
| 2810403A07Rik1 | 1.40E-06 | 0.30716 | 0.3 | 0.031 | 0.028147 | 4 M1 _ | 2810403A07Rik |
| Rnase41 | 1.35E-06 | 1.512482 | 1 | 0.601 | 0.027158 | 4 M1 _ | Rnase4 |
| Cep295nl | 1.29E-06 | 0.282322 | 0.2 | 0.014 | 0.025896 | 4 M1 _ | Cep295nl |
| B3galnt1 | 1.22E-06 | 0.607004 | 0.5 | 0.083 | 0.024534 | 4 M1 _ | B3galnt1 |
| Glipr21 | 1.21E-06 | 0.719146 | 0.7 | 0.156 | 0.024362 | 4 M1 _ | Glipr2 |
| Chst11 | 1.20E-06 | 0.706921 | 0.6 | 0.113 | 0.024121 | 4 M1 _ | Chst1 |
| 1700020I14Rik3 | 1.09E-06 | 0.772789 | 0.6 | 0.107 | 0.022048 | 4 M1 _ | 1700020I14Rik |
| Mmp23 | 1.04E-06 | 0.968367 | 0.8 | 0.207 | 0.020911 | 4 M1 _ | Mmp23 |
| Mgst11 | 1.01E-06 | 1.641805 | 1 | 0.5 | 0.020301 | 4 M1 _ | Mgst1 |
| Serpinb6a1 | 9.45E-07 | 1.724218 | 1 | 0.712 | 0.019033 | 4 M1 _ | Serpinb6a |
| 1810022K09Rik4 | 8.61E-07 | 0.631056 | 0.6 | 0.102 | 0.017346 | 4 M1 _ | 1810022K09Rik |
| Mfap5 | 8.08E-07 | 2.238491 | 1 | 0.536 | 0.016264 | 4 M1 _ | Mfap5 |
| Il1r2 | 7.21E-07 | 0.87028 | 0.4 | 0.055 | 0.014515 | 4 M1 _ | Il1r2 |
| Oxsm | 6.95E-07 | 0.403949 | 0.4 | 0.051 | 0.013996 | 4 M1 _ | Oxsm |
| Lgi41 | 6.65E-07 | 0.858864 | 0.7 | 0.151 | 0.013402 | 4 M1 _ | Lgi4 |
| Sod31 | 6.28E-07 | 0.820639 | 1 | 0.295 | 0.012655 | 4 M1 _ | Sod3 |
| Efs1 | 6.23E-07 | 0.53611 | 0.6 | 0.11 | 0.012543 | 4 M1 _ | Efs |
| Minos14 | 6.09E-07 | 0.609778 | 0.7 | 0.126 | 0.012269 | 4 M1 _ | Minos1 |
| Hdac71 | 5.91E-07 | 1.513884 | 1 | 0.45 | 0.011908 | 4 M1 _ | Hdac7 |
| Nid11 | 5.89E-07 | 1.922222 | 1 | 0.734 | 0.011857 | 4 M1 _ | Nid1 |
| Ppp1r14b | 5.53E-07 | 1.432392 | 1 | 0.485 | 0.011147 | 4 M1 _ | Ppp1r14b |
| Zcchc114 | 5.40E-07 | 0.494541 | 0.5 | 0.072 | 0.010883 | 4 M1 _ | Zcchc11 |
| Coro1b3 | 5.27E-07 | 0.535392 | 0.6 | 0.103 | 0.010612 | 4 M1 _ | Coro1b |
| Tmem55b3 | 5.25E-07 | 0.432078 | 0.4 | 0.049 | 0.010567 | 4 M1 _ | Tmem55b |
| Sulf2 | 5.03E-07 | 1.088807 | 0.8 | 0.202 | 0.010137 | 4 M1 _ | Sulf2 |
| Anxa12 | 5.01E-07 | 2.09083 | 1 | 0.573 | 0.01008 | 4 M1 _ | Anxa1 |
| Adamtsl5 | 4.81E-07 | 0.484078 | 0.7 | 0.134 | 0.009696 | 4 M1 _ | Adamtsl5 |
| Bmp11 | 3.67E-07 | 1.578271 | 1 | 0.441 | 0.007386 | 4 M1 _ | Bmp1 |
| Snhg11 | 3.27E-07 | 0.477017 | 0.2 | 0.013 | 0.006584 | 4 M1 _ | Snhg11 |
| Adgra3 | 2.93E-07 | 0.748793 | 0.7 | 0.143 | 0.005902 | 4 M1 _ | Adgra3 |
| D630033O11Rik | 2.88E-07 | 0.732351 | 0.6 | 0.104 | 0.005802 | 4 M1 _ | D630033O11Rik |
| Atp5o.14 | 2.79E-07 | 0.639437 | 0.7 | 0.123 | 0.005626 | 4 M1 _ | Atp5o.1 |
| Tmod2 | 2.44E-07 | 0.592553 | 0.7 | 0.138 | 0.004905 | 4 M1 _ | Tmod2 |
| Scrn1 | 2.36E-07 | 0.556286 | 0.5 | 0.074 | 0.004747 | 4 M1 _ | Scrn1 |
| Trim30d | 2.28E-07 | 1.026382 | 0.6 | 0.111 | 0.004592 | 4 M1 _ | Trim30d |

| | | | | | | | |
|---|---|---|---|---|---|---|---|
| Col14a11 | 2.03E-07 | 2.063766 | 1 | 0.458 | 0.004095 | 4 M1 _ | Col14a1 |
| Fbn11 | 1.96E-07 | 2.760873 | 1 | 0.621 | 0.003941 | 4 M1 _ | Fbn1 |
| Mt21 | 1.49E-07 | 1.19708 | 1 | 0.342 | 0.003004 | 4 M1 _ | Mt2 |
| Add31 | 1.22E-07 | 1.534489 | 1 | 0.507 | 0.002464 | 4 M1 _ | Add3 |
| Ddr21 | 1.09E-07 | 2.046759 | 1 | 0.442 | 0.002196 | 4 M1 _ | Ddr2 |
| Bmp6 | 9.71E-08 | 0.505647 | 0.4 | 0.044 | 0.001956 | 4 M1 _ | Bmp6 |
| Gng7 | 9.34E-08 | 0.29079 | 0.3 | 0.025 | 0.001882 | 4 M1 _ | Gng7 |
| Heg11 | 8.80E-08 | 1.781271 | 1 | 0.337 | 0.001773 | 4 M1 _ | Heg1 |
| Nqo21 | 8.57E-08 | 0.90846 | 0.7 | 0.138 | 0.001726 | 4 M1 _ | Nqo2 |
| Tgfbr21 | 8.21E-08 | 1.862673 | 1 | 0.652 | 0.001654 | 4 M1 _ | Tgfbr2 |
| Rhou1 | 7.87E-08 | 0.976563 | 0.7 | 0.14 | 0.001584 | 4 M1 _ | Rhou |
| Axl1 | 7.67E-08 | 2.006641 | 1 | 0.578 | 0.001545 | 4 M1 _ | Axl |
| Carhsp1 | 7.44E-08 | 0.967522 | 1 | 0.26 | 0.001499 | 4 M1 _ | Carhsp1 |
| Fam103a14 | 7.25E-08 | 0.623649 | 0.5 | 0.066 | 0.001461 | 4 M1 _ | Fam103a1 |
| Bicd1 | 6.95E-08 | 0.57863 | 0.4 | 0.044 | 0.001399 | 4 M1 _ | Bicd1 |
| Fam208a4 | 6.65E-08 | 0.388482 | 0.6 | 0.085 | 0.00134 | 4 M1 _ | Fam208a |
| C31 | 6.15E-08 | 3.002608 | 1 | 0.397 | 0.001238 | 4 M1 _ | C3 |
| Ebf21 | 5.99E-08 | 1.245827 | 0.8 | 0.177 | 0.001205 | 4 M1 _ | Ebf2 |
| Cd342 | 5.36E-08 | 2.135385 | 1 | 0.447 | 0.00108 | 4 M1 _ | Cd34 |
| Scara51 | 5.33E-08 | 2.00899 | 1 | 0.407 | 0.001074 | 4 M1 _ | Scara5 |
| Timp21 | 4.44E-08 | 2.721025 | 1 | 0.705 | 0.000895 | 4 M1 _ | Timp2 |
| Exd2 | 4.34E-08 | 0.438099 | 0.5 | 0.065 | 0.000873 | 4 M1 _ | Exd2 |
| Ace3 | 2.92E-08 | 2.061176 | 1 | 0.396 | 0.000588 | 4 M1 _ | Ace |
| Prkca | 2.84E-08 | 0.990026 | 0.9 | 0.213 | 0.000573 | 4 M1 _ | Prkca |
| Srd5a1 | 2.18E-08 | 0.372556 | 0.2 | 0.011 | 0.000439 | 4 M1 _ | Srd5a1 |
| Galnt161 | 2.03E-08 | 1.21939 | 0.7 | 0.128 | 0.00041 | 4 M1 _ | Galnt16 |
| Dbn1 | 1.91E-08 | 1.504961 | 0.9 | 0.26 | 0.000386 | 4 M1 _ | Dbn1 |
| Rtn1 | 1.88E-08 | 0.542594 | 0.4 | 0.043 | 0.00038 | 4 M1 _ | Rtn1 |
| Medag1 | 1.86E-08 | 1.397662 | 1 | 0.331 | 0.000375 | 4 M1 _ | Medag |
| Folh1 | 1.80E-08 | 0.277392 | 0.1 | 0.003 | 0.000362 | 4 M1 _ | Folh1 |
| Adamts21 | 1.77E-08 | 1.781133 | 1 | 0.307 | 0.000357 | 4 M1 _ | Adamts2 |
| Igfbp6 | 1.64E-08 | 2.678025 | 1 | 0.488 | 0.00033 | 4 M1 _ | Igfbp6 |
| Uap11 | 1.60E-08 | 1.671771 | 0.9 | 0.265 | 0.000323 | 4 M1 _ | Uap1 |
| Galnt15 | 1.52E-08 | 1.087915 | 0.7 | 0.132 | 0.000307 | 4 M1 _ | Galnt15 |
| Agap1 | 1.52E-08 | 1.106691 | 0.8 | 0.165 | 0.000307 | 4 M1 _ | Agap1 |
| Efemp11 | 1.45E-08 | 2.551018 | 1 | 0.355 | 0.000293 | 4 M1 _ | Efemp1 |
| Dmpk2 | 1.22E-08 | 0.571566 | 0.7 | 0.113 | 0.000246 | 4 M1 _ | Dmpk |
| Wnt2 | 8.87E-09 | 0.412859 | 0.5 | 0.059 | 0.000179 | 4 M1 _ | Wnt2 |
| Vasn | 8.80E-09 | 1.719349 | 1 | 0.298 | 0.000177 | 4 M1 _ | Vasn |
| Gfpt21 | 8.41E-09 | 1.926354 | 1 | 0.319 | 0.000169 | 4 M1 _ | Gfpt2 |
| Taf6l2 | 7.84E-09 | 0.724437 | 0.6 | 0.089 | 0.000158 | 4 M1 _ | Taf6l |

| | | | | | | | |
|---|---|---|---|---|---|---|---|
| Dact1 | 7.52E-09 | 1.433616 | 0.9 | 0.206 | 0.000151 | 4 M1 _ | Dact1 |
| Gas7 | 7.06E-09 | 1.928091 | 1 | 0.353 | 0.000142 | 4 M1 _ | Gas7 |
| D17Wsu92e4 | 6.29E-09 | 0.649565 | 0.5 | 0.06 | 0.000127 | 4 M1 _ | D17Wsu92e |
| Clec3b1 | 6.07E-09 | 2.615731 | 1 | 0.435 | 0.000122 | 4 M1 _ | Clec3b |
| Tbx18 | 2.96E-09 | 0.734627 | 0.5 | 0.059 | 5.97E-05 | 4 M1 _ | Tbx18 |
| Klf41 | 2.56E-09 | 2.246031 | 1 | 0.313 | 5.15E-05 | 4 M1 _ | Klf4 |
| Cd2481 | 2.47E-09 | 2.63618 | 1 | 0.423 | 4.96E-05 | 4 M1 _ | Cd248 |
| Gm10073 | 2.29E-09 | 0.406864 | 0.2 | 0.009 | 4.61E-05 | 4 M1 _ | Gm10073 |
| Galnt17 | 2.25E-09 | 0.692291 | 0.6 | 0.085 | 4.54E-05 | 4 M1 _ | Galnt17 |
| Tek1 | 1.70E-09 | 0.963359 | 0.9 | 0.163 | 3.43E-05 | 4 M1 _ | Tek |
| Angptl41 | 1.57E-09 | 0.96097 | 0.6 | 0.08 | 3.16E-05 | 4 M1 _ | Angptl4 |
| Klhl131 | 1.52E-09 | 0.911853 | 0.8 | 0.15 | 3.07E-05 | 4 M1 _ | Klhl13 |
| Mst1r | 1.42E-09 | 0.257497 | 0.3 | 0.02 | 2.85E-05 | 4 M1 _ | Mst1r |
| Fam208b3 | 1.39E-09 | 0.583556 | 0.4 | 0.036 | 2.79E-05 | 4 M1 _ | Fam208b |
| Hoxc4 | 1.07E-09 | 0.25568 | 0.1 | 0.002 | 2.15E-05 | 4 M1 _ | Hoxc4 |
| Ifi2051 | 6.52E-10 | 2.529903 | 1 | 0.295 | 1.31E-05 | 4 M1 _ | Ifi205 |
| Cdon1 | 5.66E-10 | 1.058046 | 0.7 | 0.107 | 1.14E-05 | 4 M1 _ | Cdon |
| Man1a1 | 5.64E-10 | 1.939153 | 1 | 0.335 | 1.13E-05 | 4 M1 _ | Man1a |
| Coro2b1 | 5.06E-10 | 0.699439 | 0.6 | 0.081 | 1.02E-05 | 4 M1 _ | Coro2b |
| Uchl1 | 3.63E-10 | 0.544116 | 0.4 | 0.035 | 7.31E-06 | 4 M1 _ | Uchl1 |
| Aaed13 | 3.02E-10 | 0.713282 | 0.6 | 0.072 | 6.09E-06 | 4 M1 _ | Aaed1 |
| Osr11 | 2.84E-10 | 1.47383 | 0.9 | 0.19 | 5.72E-06 | 4 M1 _ | Osr1 |
| Anxa32 | 2.78E-10 | 2.62631 | 1 | 0.313 | 5.60E-06 | 4 M1 _ | Anxa3 |
| Sgms2 | 2.27E-10 | 0.392591 | 0.4 | 0.033 | 4.57E-06 | 4 M1 _ | Sgms2 |
| Nox4 | 2.04E-10 | 1.113638 | 0.6 | 0.078 | 4.10E-06 | 4 M1 _ | Nox4 |
| 2410015M20Rik4 | 1.66E-10 | 0.766106 | 0.8 | 0.116 | 3.35E-06 | 4 M1 _ | 2410015M20Rik |
| Srgap11 | 1.37E-10 | 0.947902 | 0.6 | 0.071 | 2.75E-06 | 4 M1 _ | Srgap1 |
| Ildr2 | 1.16E-10 | 0.69903 | 0.7 | 0.093 | 2.33E-06 | 4 M1 _ | Ildr2 |
| Lurap1l1 | 1.04E-10 | 0.760607 | 0.6 | 0.069 | 2.09E-06 | 4 M1 _ | Lurap1l |
| Arl4d1 | 9.96E-11 | 0.795171 | 0.5 | 0.051 | 2.01E-06 | 4 M1 _ | Arl4d |
| Nhsl1 | 9.86E-11 | 0.9139 | 0.7 | 0.098 | 1.99E-06 | 4 M1 _ | Nhsl1 |
| Tmem1002 | 8.52E-11 | 2.408157 | 1 | 0.284 | 1.72E-06 | 4 M1 _ | Tmem100 |
| Emb | 8.23E-11 | 0.718564 | 0.5 | 0.049 | 1.66E-06 | 4 M1 _ | Emb |
| Npr11 | 8.16E-11 | 0.971925 | 0.8 | 0.121 | 1.64E-06 | 4 M1 _ | Npr1 |
| Il6ra | 6.43E-11 | 0.648446 | 0.4 | 0.032 | 1.29E-06 | 4 M1 _ | Il6ra |
| Arhgap201 | 5.33E-11 | 1.544576 | 1 | 0.242 | 1.07E-06 | 4 M1 _ | Arhgap20 |
| Mustn11 | 4.67E-11 | 1.035519 | 0.8 | 0.119 | 9.40E-07 | 4 M1 _ | Mustn1 |
| Gm26870 | 1.92E-11 | 0.276736 | 0.1 | 0.002 | 3.87E-07 | 4 M1 _ | Gm26870 |
| Daglb | 1.57E-11 | 1.132388 | 0.7 | 0.095 | 3.17E-07 | 4 M1 _ | Daglb |
| Fez11 | 1.38E-11 | 0.695542 | 0.6 | 0.067 | 2.77E-07 | 4 M1 _ | Fez1 |
| Smtnl2 | 1.00E-11 | 0.319179 | 0.2 | 0.007 | 2.02E-07 | 4 M1 _ | Smtnl2 |

| | | | | | | | |
|---|---|---|---|---|---|---|---|
| Dscr31 | 9.43E-12 | 0.409747 | 0.4 | 0.029 | 1.90E-07 | 4 M1 _ | Dscr3 |
| Hspb8 | 9.19E-12 | 0.912944 | 0.8 | 0.119 | 1.85E-07 | 4 M1 _ | Hspb8 |
| Ugdh1 | 7.48E-12 | 1.757183 | 1 | 0.242 | 1.51E-07 | 4 M1 _ | Ugdh |
| Atp6v0e2 | 4.68E-12 | 0.470129 | 0.4 | 0.028 | 9.43E-08 | 4 M1 _ | Atp6v0e2 |
| Pde8a1 | 4.29E-12 | 1.21762 | 1 | 0.199 | 8.64E-08 | 4 M1 _ | Pde8a |
| Pla2g163 | 3.41E-12 | 0.640368 | 0.5 | 0.043 | 6.88E-08 | 4 M1 _ | Pla2g16 |
| Pi16 | 3.25E-12 | 4.679027 | 1 | 0.265 | 6.54E-08 | 4 M1 _ | Pi16 |
| Qpct | 3.01E-12 | 1.251781 | 0.8 | 0.121 | 6.05E-08 | 4 M1 _ | Qpct |
| Tnxb1 | 2.82E-12 | 3.020776 | 1 | 0.238 | 5.69E-08 | 4 M1 _ | Tnxb |
| Gria4 | 2.67E-12 | 0.391534 | 0.2 | 0.007 | 5.38E-08 | 4 M1 _ | Gria4 |
| Adh7 | 2.61E-12 | 0.331264 | 0.2 | 0.007 | 5.27E-08 | 4 M1 _ | Adh7 |
| Flnc1 | 2.44E-12 | 0.969709 | 0.8 | 0.118 | 4.91E-08 | 4 M1 _ | Flnc |
| Kcnk5 | 2.24E-12 | 0.486537 | 0.4 | 0.028 | 4.50E-08 | 4 M1 _ | Kcnk5 |
| Fam220a3 | 1.03E-12 | 0.74497 | 0.6 | 0.057 | 2.08E-08 | 4 M1 _ | Fam220a |
| Sirpa | 5.97E-13 | 0.699126 | 0.7 | 0.08 | 1.20E-08 | 4 M1 _ | Sirpa |
| Kif21a | 4.81E-13 | 0.636532 | 0.4 | 0.026 | 9.69E-09 | 4 M1 _ | Kif21a |
| Mkl11 | 4.25E-13 | 0.397218 | 0.4 | 0.025 | 8.56E-09 | 4 M1 _ | Mkl1 |
| Mkl22 | 4.02E-13 | 0.464695 | 0.6 | 0.053 | 8.09E-09 | 4 M1 _ | Mkl2 |
| Gpc31 | 2.81E-13 | 1.53287 | 0.7 | 0.083 | 5.67E-09 | 4 M1 _ | Gpc3 |
| Tppp31 | 2.64E-13 | 1.953479 | 1 | 0.19 | 5.31E-09 | 4 M1 _ | Tppp3 |
| Ints5.1 | 1.85E-13 | 0.365104 | 0.3 | 0.014 | 3.72E-09 | 4 M1 _ | Ints5.1 |
| Gda1 | 7.26E-14 | 1.668077 | 0.7 | 0.08 | 1.46E-09 | 4 M1 _ | Gda |
| 2810474O19Rik4 | 3.46E-14 | 1.569514 | 0.7 | 0.077 | 6.97E-10 | 4 M1 _ | 2810474O19Rik |
| Il18 | 2.55E-14 | 0.627418 | 0.4 | 0.024 | 5.14E-10 | 4 M1 _ | Il18 |
| Nat8f3 | 1.50E-14 | 0.281438 | 0.2 | 0.006 | 3.03E-10 | 4 M1 _ | Nat8f3 |
| 5033421B08Rik | 1.22E-14 | 0.338036 | 0.2 | 0.006 | 2.45E-10 | 4 M1 _ | 5033421B08Rik |
| Aes4 | 8.68E-15 | 1.439021 | 1 | 0.135 | 1.75E-10 | 4 M1 _ | Aes |
| Gm26771 | 8.00E-15 | 0.51266 | 0.4 | 0.022 | 1.61E-10 | 4 M1 _ | Gm26771 |
| Arsi | 6.21E-15 | 0.751605 | 0.6 | 0.051 | 1.25E-10 | 4 M1 _ | Arsi |
| Gprc5b | 4.75E-15 | 0.905622 | 0.7 | 0.071 | 9.57E-11 | 4 M1 _ | Gprc5b |
| Fbxo182 | 4.62E-15 | 0.642641 | 0.6 | 0.049 | 9.30E-11 | 4 M1 _ | Fbxo18 |
| Sfrp21 | 3.66E-15 | 2.10719 | 0.8 | 0.093 | 7.37E-11 | 4 M1 _ | Sfrp2 |
| 2010107E04Rik4 | 1.39E-15 | 1.344128 | 1 | 0.127 | 2.80E-11 | 4 M1 _ | 2010107E04Rik |
| Sdccag33 | 1.07E-15 | 0.554225 | 0.6 | 0.046 | 2.16E-11 | 4 M1 _ | Sdccag3 |
| Adora2b | 1.06E-15 | 0.596945 | 0.5 | 0.033 | 2.14E-11 | 4 M1 _ | Adora2b |
| Lrrc171 | 6.47E-16 | 1.407871 | 0.7 | 0.067 | 1.30E-11 | 4 M1 _ | Lrrc17 |
| Pamr1 | 5.10E-16 | 1.921924 | 0.8 | 0.091 | 1.03E-11 | 4 M1 _ | Pamr1 |
| Dpt1 | 4.79E-16 | 3.549243 | 1 | 0.167 | 9.65E-12 | 4 M1 _ | Dpt |
| Ism1 | 3.84E-16 | 1.102832 | 0.5 | 0.034 | 7.73E-12 | 4 M1 _ | Ism1 |
| Zfp385a | 3.75E-16 | 2.069098 | 0.9 | 0.123 | 7.55E-12 | 4 M1 _ | Zfp385a |
| Emilin21 | 2.94E-16 | 2.628175 | 1 | 0.171 | 5.92E-12 | 4 M1 _ | Emilin2 |

| | | | | | | | |
|---|---|---|---|---|---|---|---|
| C4b1 | 2.69E-16 | 1.705626 | 0.8 | 0.086 | 5.42E-12 | 4 M1 _ | C4b |
| Ackr31 | 1.71E-16 | 2.516615 | 0.9 | 0.123 | 3.45E-12 | 4 M1 _ | Ackr3 |
| Fn11 | 1.26E-16 | 3.909252 | 1 | 0.162 | 2.53E-12 | 4 M1 _ | Fn1 |
| Hoxb3 | 1.24E-16 | 0.272015 | 0.2 | 0.005 | 2.50E-12 | 4 M1 _ | Hoxb3 |
| Gm15564 | 9.70E-17 | 0.317656 | 0.2 | 0.005 | 1.95E-12 | 4 M1 _ | Gm15564 |
| AI464131 | 4.65E-17 | 0.350128 | 0.3 | 0.011 | 9.36E-13 | 4 M1 _ | AI464131 |
| Fam213b1 | 1.78E-17 | 0.630504 | 0.4 | 0.019 | 3.58E-13 | 4 M1 _ | Fam213b |
| Sfrp4 | 1.66E-17 | 1.477214 | 0.4 | 0.019 | 3.35E-13 | 4 M1 _ | Sfrp4 |
| Zbtb163 | 1.52E-17 | 1.252849 | 0.9 | 0.097 | 3.07E-13 | 4 M1 _ | Zbtb16 |
| Edn11 | 1.35E-17 | 2.202648 | 0.5 | 0.03 | 2.71E-13 | 4 M1 _ | Edn1 |
| Adra1b | 6.52E-18 | 0.311806 | 0.2 | 0.004 | 1.31E-13 | 4 M1 _ | Adra1b |
| En1 | 5.90E-18 | 0.272633 | 0.2 | 0.004 | 1.19E-13 | 4 M1 _ | En1 |
| Sdk1 | 4.87E-18 | 0.780388 | 0.6 | 0.042 | 9.82E-14 | 4 M1 _ | Sdk1 |
| Sbsn | 1.50E-18 | 2.080147 | 0.7 | 0.059 | 3.03E-14 | 4 M1 _ | Sbsn |
| Megf9 | 1.46E-18 | 0.832358 | 0.8 | 0.071 | 2.94E-14 | 4 M1 _ | Megf9 |
| Fhl12 | 1.05E-18 | 1.590903 | 0.8 | 0.073 | 2.12E-14 | 4 M1 _ | Fhl1 |
| Fam46a2 | 9.28E-19 | 1.268693 | 0.7 | 0.054 | 1.87E-14 | 4 M1 _ | Fam46a |
| Pygl | 1.05E-19 | 1.003157 | 0.8 | 0.068 | 2.11E-15 | 4 M1 _ | Pygl |
| Gm16894 | 7.56E-20 | 0.38213 | 0.3 | 0.009 | 1.52E-15 | 4 M1 _ | Gm16894 |
| Tubb4a1 | 5.81E-20 | 0.601649 | 0.5 | 0.025 | 1.17E-15 | 4 M1 _ | Tubb4a |
| Car81 | 4.57E-20 | 1.058159 | 0.8 | 0.065 | 9.19E-16 | 4 M1 _ | Car8 |
| Vldlr1 | 2.64E-20 | 1.298497 | 0.8 | 0.067 | 5.32E-16 | 4 M1 _ | Vldlr |
| Lamc2 | 8.28E-21 | 0.593 | 0.3 | 0.009 | 1.67E-16 | 4 M1 _ | Lamc2 |
| Efna5 | 4.49E-21 | 0.623257 | 0.4 | 0.015 | 9.04E-17 | 4 M1 _ | Efna5 |
| Sema3c1 | 3.82E-21 | 3.120442 | 1 | 0.118 | 7.70E-17 | 4 M1 _ | Sema3c |
| Cd551 | 3.57E-21 | 2.560169 | 1 | 0.117 | 7.19E-17 | 4 M1 _ | Cd55 |
| Fndc1 | 4.50E-23 | 3.525262 | 1 | 0.108 | 9.07E-19 | 4 M1 _ | Fndc1 |
| Irf4 | 9.06E-24 | 0.441536 | 0.2 | 0.003 | 1.82E-19 | 4 M1 _ | Irf4 |
| Kdelc23 | 4.39E-24 | 1.369485 | 0.9 | 0.071 | 8.83E-20 | 4 M1 _ | Kdelc2 |
| CT025619.11 | 2.14E-24 | 0.912207 | 0.6 | 0.03 | 4.30E-20 | 4 M1 _ | CT025619.1 |
| 1-Nov | 2.87E-25 | 1.063457 | 0.4 | 0.012 | 5.79E-21 | 4 M1 _ | Nov |
| Dact2 | 4.49E-26 | 1.381105 | 0.6 | 0.029 | 9.03E-22 | 4 M1 _ | Dact2 |
| Gm15675 | 2.87E-26 | 0.368541 | 0.3 | 0.006 | 5.79E-22 | 4 M1 _ | Gm15675 |
| Spsb4 | 1.24E-26 | 0.498472 | 0.2 | 0.003 | 2.49E-22 | 4 M1 _ | Spsb4 |
| Opcml | 2.97E-27 | 1.535823 | 0.6 | 0.027 | 5.98E-23 | 4 M1 _ | Opcml |
| 5830416I19Rik | 3.04E-28 | 0.378803 | 0.4 | 0.011 | 6.13E-24 | 4 M1 _ | 5830416I19Rik |
| Vegfd | 3.09E-29 | 1.568998 | 0.6 | 0.025 | 6.22E-25 | 4 M1 _ | Vegfd |
| Dmkn | 3.64E-30 | 0.285188 | 0.2 | 0.002 | 7.32E-26 | 4 M1 _ | Dmkn |
| Slc4a4 | 3.55E-30 | 1.749685 | 0.8 | 0.045 | 7.15E-26 | 4 M1 _ | Slc4a4 |
| Creb5 | 3.40E-30 | 2.523531 | 0.9 | 0.06 | 6.84E-26 | 4 M1 _ | Creb5 |
| Adam33 | 2.76E-31 | 1.54425 | 0.8 | 0.044 | 5.57E-27 | 4 M1 _ | Adam33 |

| | | | | | | | |
|---|---|---|---|---|---|---|---|
| Ppp2r2b | 8.98E-32 | 0.615706 | 0.4 | 0.009 | 1.81E-27 | 4 M1 _ | Ppp2r2b |
| Efhd11 | 1.60E-32 | 1.431097 | 0.8 | 0.04 | 3.21E-28 | 4 M1 _ | Efhd1 |
| Aldh1a3 | 1.44E-32 | 1.361196 | 0.6 | 0.022 | 2.89E-28 | 4 M1 _ | Aldh1a3 |
| Prkg2 | 2.88E-33 | 0.658658 | 0.4 | 0.009 | 5.79E-29 | 4 M1 _ | Prkg2 |
| Stmn4 | 1.54E-33 | 0.466519 | 0.3 | 0.005 | 3.10E-29 | 4 M1 _ | Stmn4 |
| Ston2 | 1.89E-34 | 0.98446 | 0.6 | 0.021 | 3.80E-30 | 4 M1 _ | Ston2 |
| Fam46b | 4.83E-35 | 0.334542 | 0.2 | 0.002 | 9.73E-31 | 4 M1 _ | Fam46b |
| Clip4 | 4.05E-35 | 1.572447 | 0.9 | 0.049 | 8.16E-31 | 4 M1 _ | Clip4 |
| Stxbp6 | 5.92E-36 | 1.201559 | 0.8 | 0.036 | 1.19E-31 | 4 M1 _ | Stxbp6 |
| Cadm31 | 1.52E-36 | 1.494623 | 1 | 0.057 | 3.07E-32 | 4 M1 _ | Cadm3 |
| Adamtsl4 | 1.50E-36 | 1.551281 | 0.8 | 0.036 | 3.01E-32 | 4 M1 _ | Adamtsl4 |
| Pcsk61 | 5.96E-37 | 3.003431 | 1 | 0.061 | 1.20E-32 | 4 M1 _ | Pcsk6 |
| Adgrd1 | 2.10E-40 | 1.867996 | 0.9 | 0.042 | 4.23E-36 | 4 M1 _ | Adgrd1 |
| Cmah | 8.56E-41 | 1.871298 | 1 | 0.051 | 1.72E-36 | 4 M1 _ | Cmah |
| Ackr1 | 3.62E-41 | 0.83389 | 0.6 | 0.017 | 7.28E-37 | 4 M1 _ | Ackr1 |
| Serpina3c | 6.23E-42 | 0.438436 | 0.2 | 0.001 | 1.25E-37 | 4 M1 _ | Serpina3c |
| Ptx3 | 7.49E-44 | 0.758436 | 0.4 | 0.006 | 1.51E-39 | 4 M1 _ | Ptx3 |
| Prdm8 | 3.71E-44 | 1.165419 | 0.7 | 0.021 | 7.48E-40 | 4 M1 _ | Prdm8 |
| Sntg1 | 5.12E-46 | 0.684606 | 0.4 | 0.006 | 1.03E-41 | 4 M1 _ | Sntg1 |
| Adgrg2 | 8.25E-47 | 0.507858 | 0.3 | 0.003 | 1.66E-42 | 4 M1 _ | Adgrg2 |
| Tmeff2 | 2.03E-47 | 1.504091 | 0.8 | 0.027 | 4.08E-43 | 4 M1 _ | Tmeff2 |
| Aif1l | 1.38E-47 | 1.146094 | 0.7 | 0.02 | 2.77E-43 | 4 M1 _ | Aif1l |
| 1700019D03Rik | 6.74E-48 | 1.201507 | 0.8 | 0.026 | 1.36E-43 | 4 M1 _ | 1700019D03Rik |
| Rorb | 1.12E-51 | 0.798668 | 0.4 | 0.005 | 2.25E-47 | 4 M1 _ | Rorb |
| Lrrn4cl | 7.47E-52 | 2.024279 | 1 | 0.04 | 1.50E-47 | 4 M1 _ | Lrrn4cl |
| Miat | 3.61E-52 | 0.399217 | 0.2 | 0.001 | 7.26E-48 | 4 M1 _ | Miat |
| Gm36888 | 1.79E-52 | 0.260133 | 0.1 | 0 | 3.60E-48 | 4 M1 _ | Gm36888 |
| Chrdl1 | 1.49E-53 | 1.424684 | 0.5 | 0.008 | 3.01E-49 | 4 M1 _ | Chrdl1 |
| Gpr1 | 4.47E-59 | 0.65698 | 0.4 | 0.004 | 9.01E-55 | 4 M1 _ | Gpr1 |
| Bmp3 | 1.19E-60 | 0.653715 | 0.5 | 0.007 | 2.39E-56 | 4 M1 _ | Bmp3 |
| Dpp4 | 5.56E-62 | 2.645448 | 1 | 0.033 | 1.12E-57 | 4 M1 _ | Dpp4 |
| Duox1 | 1.88E-63 | 0.660672 | 0.4 | 0.004 | 3.78E-59 | 4 M1 _ | Duox1 |
| Eppk1 | 3.50E-66 | 0.314584 | 0.3 | 0.002 | 7.05E-62 | 4 M1 _ | Eppk1 |
| Islr2 | 2.69E-68 | 0.79063 | 0.6 | 0.009 | 5.41E-64 | 4 M1 _ | Islr2 |
| Mmp27 | 2.47E-78 | 0.88424 | 0.7 | 0.011 | 4.98E-74 | 4 M1 _ | Mmp27 |
| Ackr2 | 8.78E-81 | 1.24855 | 0.6 | 0.007 | 1.77E-76 | 4 M1 _ | Ackr2 |
| Pcolce2 | 2.81E-84 | 3.036075 | 1 | 0.022 | 5.66E-80 | 4 M1 _ | Pcolce2 |
| Htr2a | 1.48E-92 | 0.451531 | 0.3 | 0.001 | 2.99E-88 | 4 M1 _ | Htr2a |
| Nova1 | 1.60E-98 | 2.103567 | 1 | 0.018 | 3.21E-94 | 4 M1 _ | Nova1 |
| Smpd3 | 3.52E-103 | 3.563491 | 0.8 | 0.01 | 7.09E-99 | 4 M1 _ | Smpd3 |
| Duoxa1 | 1.87E-138 | 0.757994 | 0.7 | 0.005 | 3.77E-134 | 4 M1 _ | Duoxa1 |

**Supplementary Table 4. Differentially expressed matrisome genes in stromal subsetted mock and ligated cells**

Differentially expressed matrisome-associated gene are shown for the stromal cell subset gene lists from Supplementary Table 3. Genes colored red are expressed in both ligated and mock cells within the cluster and those with no colors are unique to the condition.

| Cluster 0 | | Cluster 1 | | Cluster 2 | | Cluster 3 | | Cluster 4 | |
|---|---|---|---|---|---|---|---|---|---|
| **Ligated** | **Mock** | **Ligated** | **Mock** | **Ligated** | **Mock** | **Ligated** | **Mock** | **Ligated** | **Mock** |
| Abi3bp | Bmper | Fgl2 | Srgn | Sparcl1 | Crim1 | Agrn | Crim1 | Aebp1 | Dpt |
| Aebp1 | Crispld2 | Lamb3 | | **Ecm2** | Ctgf | Fbln2 | Ctgf | AW551984 | Efemp1 |
| Cilp | Ctgf | Srgn | | **Emid1** | Cyr61 | Gas6 | Cyr61 | Cthrc1 | Emilin2 |
| Creld2 | Cyr61 | | | **Lgi1** | Ecm2 | Hmcn1 | Ecm2 | Ecm1 | Fbn1 |
| Crispld1 | Dpt | | | **Mfge8** | Edil3 | Igfbp3 | Edil3 | Fbln2 | Fn1 |
| Efemp2 | Ecm1 | | | **Tinagl1** | Emid1 | Igfbp7 | Emid1 | Fbln5 | Fndc1 |
| Egflam | Efemp1 | | | **Vtn** | Lamb2 | Lama3 | Lamb2 | Igfbp5 | Igfbp6 |
| Eln | Emilin2 | | | **Aspn** | Lgi1 | Lama5 | Lgi1 | Lama4 | Lamc2 |
| Emilin1 | Fbln1 | | | | Mfge8 | Lrg1 | Mfge8 | Lgi2 | Lgi4 |
| Fbn2 | Fbln2 | | | | Ntn4 | Ltbp4 | Ntn4 | Lgi3 | Mfap5 |
| Fgl2 | Fbln5 | | | | Postn | Mfap3 | Postn | Thbs2 | Nid1 |
| Gas6 | Fbln7 | | | | Sparc | Mmrn2 | Sparc | Thbs3 | Nov |
| Igfbp6 | Fbn1 | | | | Thsd4 | Vwa1 | Thsd4 | Tnfaip6 | Pcolce2 |
| Igfbp7 | Fn1 | | | | Tinagl1 | Col13a1 | Tinagl1 | Col1a1 | Tnxb |
| Ltbp2 | Igfbp4 | | | | Vtn | Esm1 | Vtn | Col1a2 | Col14a1 |
| Mfap2 | Igfbp5 | | | | Col4a1 | Hspg2 | Col4a1 | Col3a1 | Col5a1 |
| Mfap4 | Igsf10 | | | | Col4a2 | Srgn | Col4a2 | Ogn | |
| Mfap5 | Lama2 | | | | Aspn | **Crim1** | Aspn | **Dpt** | |
| Mgp | Lama4 | | | | | **Tinagl1** | | **Efemp1** | |
| Postn | Lamb1 | | | | | **Col4a1** | | **Emilin2** | |
| Slit2 | Lamc1 | | | | | **Col4a2** | | **Fbn1** | |
| Sparc | Lamc3 | | | | | | | **Fn1** | |
| Spp1 | Lgi2 | | | | | | | **Fndc1** | |
| Svep1 | Lgi4 | | | | | | | **Igfbp6** | |
| Thbs2 | Ltbp1 | | | | | | | **Lamc2** | |
| Thbs4 | Ltbp3 | | | | | | | **Mfap5** | |
| Col11a1 | Ltbp4 | | | | | | | **Nid1** | |
| Col12a1 | Nid1 | | | | | | | **Pcolce2** | |
| Col16a1 | Nid2 | | | | | | | **Tnxb** | |
| Col28a1 | Pcolce | | | | | | | **Col14a1** | |
| Col4a3 | Pxdn | | | | | | | **Col5a1** | |
| Col4a4 | Rspo1 | | | | | | | | |
| Col4a5 | Rspo3 | | | | | | | | |
| Col8a1 | Slit3 | | | | | | | | |
| Col8a2 | Smoc2 | | | | | | | | |
| Bgn | Sned1 | | | | | | | | |
| Fmod | Spon1 | | | | | | | | |
| Omd | Spon2 | | | | | | | | |
| **Crispld2** | Srpx | | | | | | | | |
| **Ecm1** | Srpx2 | | | | | | | | |
| **Efemp1** | Tgfbi | | | | | | | | |
| **Fbln1** | Thbs1 | | | | | | | | |
| **Fbln5** | Thbs3 | | | | | | | | |
| **Fbln7** | Tnfaip6 | | | | | | | | |

| | |
|---|---|
| **Fbn1** | Tnxb |
| **Igfbp4** | Vit |
| **Igfbp5** | Vwa5a |
| **Igsf10** | Wisp1 |
| **Lama2** | Col14a1 |
| **Lamb1** | Col15a1 |
| **Lamc1** | Col18a1 |
| **Lamc3** | Col1a1 |
| **Ltbp1** | Col1a2 |
| **Ltbp3** | Col23a1 |
| **Nid1** | Col27a1 |
| **Nid2** | Col3a1 |
| **Pcolce** | Col4a1 |
| **Rspo3** | Col5a1 |
| **Slit3** | Col5a2 |
| **Smoc2** | Col5a3 |
| **Sned1** | Col6a1 |
| **Spon1** | Col6a2 |
| **Srpx** | Col6a3 |
| **Srpx2** | Col6a5 |
| **Tgfbi** | Col6a6 |
| **Thbs1** | Dcn |
| **Col14a1** | Hspg2 |
| **Col15a1** | Lum |
| **Col1a1** | Ogn |
| **Col1a2** | Podn |
| **Col27a1** | Prelp |
| **Col3a1** | Prg4 |
| **Col5a1** | Spock2 |
| **Col5a2** | Vcan |
| **Col5a3** | |
| **Col6a1** | |
| **Col6a2** | |
| **Col6a3** | |
| **Col6a6** | |
| **Dcn** | |
| **Hspg2** | |
| **Lum** | |
| **Ogn** | |
| **Podn** | |
| **Prelp** | |
| **Vcan** | |
